# Machine Learning-Driven Prediction of TLR4 Binding Affinity: A Comprehensive Molecular Feature Analysis for Drug Discovery

**DOI:** 10.1101/2025.09.26.676276

**Authors:** Brandon Yee, Maximilian Rutowski, Wilson Collins

## Abstract

Toll-like receptor 4 (TLR4) represents a promising therapeutic target for inflammatory diseases and cancer, but developing selective modulators remains challenging. We present a machine learning approach for predicting TLR4 binding affinity using comprehensive molecular descriptors. Our ensemble learning pipeline extracted 53 physicochemical features from a curated dataset of 49 unique TLR4 ligands, achieving cross-validation *R*^2^ of 0.74 ± 0.10 with statistical significance confirmed by permutation testing (p < 0.01). The most predictive features were Bertz complexity (importance: 0.173), molecular shape descriptors (0.159), and molar refractivity (0.145), while traditional drug-like properties such as LogP showed lower importance (0.056). This suggests TLR4 binding follows distinct structure-activity patterns compared to conventional drug targets. Compounds with intermediate structural complexity (Bertz complexity: 400-600) demonstrated optimal binding affinity. The model successfully identified key molecular scaffolds including flavonoids and terpenoids, aligning with known natural product TLR4 modulators. This work provides the first comprehensive machine learning analysis of TLR4 binding determinants and offers a computational framework for rational design of TLR4-targeted therapeutics, with identified molecular features providing actionable insights for developing next-generation immunomodulatory drugs.

## 1. Introduction

Toll-like receptor 4 (TLR4) is a pattern recognition receptor that plays a pivotal role in innate immunity by recognizing pathogen-associated molecular patterns (PAMPs) and damage-associated molecular patterns (DAMPs). As a key mediator of inflammatory responses, TLR4 has emerged as an attractive therapeutic target for treating inflammatory diseases, sepsis, and cancer.

However, the development of selective TLR4 modulators has been hampered by the complexity of TLR4 signaling pathways and limited understanding of structure-activity relationships governing TLR4-ligand interactions. The emergence of machine learning (ML) in drug discovery offers new opportunities to accelerate the identification and optimization of TLR4-targeted therapeutics.

In this study, we present the first comprehensive machine learning analysis of TLR4 binding affinity using a curated dataset of 49 unique compounds with experimentally determined binding affinities. We developed an ensemble learning pipeline that combines multiple algorithms to achieve robust predictions while identifying the most important molecular features governing TLR4 binding. Our approach addresses key methodological challenges in small-dataset machine learning, including data leakage prevention, chemical diversity preservation, and rigorous statistical validation.

The primary objectives of this work are: (1) to develop a reliable computational model for predicting TLR4 binding affinity, (2) to identify key molecular features that determine TLR4 binding specificity, (3) to provide insights into structure-activity relationships for rational drug design, and (4) to establish a computational framework that can be extended to larger datasets and related immune targets.

## 2. Literature Review

Traditional drug discovery approaches for TLR4 modulators have relied primarily on high-throughput screening and structure-based drug design [1]. While these methods have yielded some promising candidates, they are resource-intensive and often fail to capture the subtle molecular features that determine binding specificity and functional outcomes. The emergence of machine learning (ML) in drug discovery offers new opportunities to accelerate the identification and optimization of TLR4-targeted therapeutics [2].

Recent advances in computational chemistry have demonstrated the power of molecular descriptors in predicting biological activity [3]. These descriptors encode various aspects of molecular structure, including topological, geometric, and physicochemical properties, providing a comprehensive representation of chemical space. When combined with machine learning algorithms, molecular descriptors can reveal hidden patterns in structure-activity relationships that are not apparent through traditional analysis methods.

Several studies have applied machine learning to predict the activity of immune system modulators, but few have specifically focused on TLR4 binding prediction [4]. The unique structural features of the TLR4 binding pocket, characterized by a large, hydrophobic cavity with multiple sub-pockets, suggest that TLR4 binding may follow distinct structure-activity patterns compared to conventional drug targets [5].

## 3. Dataset

The initial dataset comprised 1,348,771 molecular conformations representing 49 unique chemical compounds with experimentally determined TLR4 binding affinities. Binding affinities were measured using surface plasmon resonance (SPR) and reported as negative logarithmic dissociation constants (− log *K*_*d*_) in kcal/mol units, ranging from -5.645 to -9.524 kcal/mol.

To address data leakage concerns inherent in conformational datasets, we implemented a rigorous deduplication strategy. Multiple conformations of the same compound were consolidated by retaining only the conformation with the highest binding affinity, resulting in 49 unique base compounds. This approach prevents the artificial inflation of model performance that would occur if different conformations of the same molecule appeared in both training and test sets.

Chemical diversity was further assessed using Tanimoto similarity coefficients calculated from Morgan fingerprints (radius=2, 2048 bits). Compounds with similarity ¿95% were flagged for manual review, but all were retained to preserve the limited chemical diversity in our small dataset. The final dataset exhibited good chemical diversity with a mean pairwise Tanimoto similarity of 0.23 ± 0.15.

The curated dataset comprised 49 unique TLR4 ligands with binding affinities ranging from -5.645 to -9.524 kcal/mol (mean: -7.42 ± 0.89 kcal/mol). The compounds exhibited substantial chemical diversity, including natural products (flavonoids, terpenoids, alkaloids), synthetic small molecules, and known TLR4 modulators such as Eritoran and Docetaxel.

Chemical space analysis revealed three major clusters: (1) flavonoid derivatives (n=12), (2) terpenoid compounds (n=8), and (3) synthetic modulators (n=7), with the remaining compounds distributed across various chemical classes. This diversity provides a robust foundation for developing generalizable structure-activity relationships.

## 4. Methodology

### 4.1. Molecular Feature Extraction

Comprehensive molecular descriptors were calculated using RDKit (version 2023.03.1) [6]. A total of 53 molecular descriptors were initially computed, encompassing:

#### Physicochemical properties

Molecular weight, LogP, topological polar surface area (TPSA), molar refractivity, and formal charge.

#### Structural descriptors

Heavy atom count, ring count, rotatable bonds, aromatic rings, saturated and aliphatic carbocycles.

#### Topological indices

Bertz complexity index, Balaban J index, Kappa shape indices, and Zagreb indices.

#### Fragment counts

Lipinski rule violations, hydrogen bond donors and acceptors, and various functional group counts.

#### Binding-specific metrics

Ligand efficiency (LE = −Δ*G*/heavy atoms) and binding efficiency index (BEI = −Δ*G*/MW *×* 1000).

### 4.2. Feature Selection and Preprocessing

A multi-stage feature selection pipeline was implemented to identify the most predictive molecular descriptors while avoiding overfitting in our small dataset:

#### Stage 1 - Variance filtering

Features with variance ¡0.01 were removed to eliminate near-constant descriptors.

#### Stage 2 - Univariate selection

Features were ranked using mutual information regression, and the top 20 features were retained.

#### Stage 3 - Model-based selection

A LASSO regression model (*α* = 0.1) was used to identify features with non-zero coefficients.

#### Stage 4 - Recursive feature elimination

The final feature set was optimized using recursive feature elimination with cross-validation (RFECV) on a Random Forest model.

This process yielded 10 final features, providing a samples-to-features ratio of 4.9:1, which is appropriate for our dataset size. Data preprocessing included power transformation (Yeo-Johnson) to address skewness and robust scaling to minimize the impact of outliers.

### 4.3. Machine Learning Models

We employed an ensemble learning approach combining multiple algorithms to achieve robust predictions:

#### Random Forest

300 trees with max depth = 10, min samples split = 3, optimized for small datasets.

#### ElasticNet

Linear model with L1/L2 regularization (*α* = 0.1, l1 ratio = 0.5) to prevent overfitting.

#### Ridge Regression

L2-regularized linear model (*α* = 1.0) for baseline linear relationships.

#### Bayesian Ridge

Probabilistic linear model providing uncertainty estimates.

The final ensemble model combined these algorithms using equal weighting in a voting regressor framework. Hyperparameters were optimized using 5-fold cross-validation with grid search.

### 4.4. Model Validation and Statistical Analysis

Rigorous validation was performed to ensure reliable performance estimates:

#### Train-test split

80/20 stratified split based on binding affinity quartiles to ensure representative sampling.

#### Cross-validation

5-fold cross-validation with 10 repetitions to obtain robust performance estimates.

#### Permutation testing

Statistical significance was assessed by comparing model performance to 1000 permuted datasets.

#### Learning curves

Generated to assess overfitting and determine optimal training set size.

Model performance was evaluated using multiple metrics: coefficient of determination (*R*^2^), root mean squared error (RMSE), and mean absolute error (MAE). Statistical significance was set at p ¡ 0.05.

### 4.5. Feature Importance Analysis

Feature importance was assessed using multiple methods to ensure robust interpretation:

#### Permutation importance

Features were randomly shuffled, and the decrease in model performance was measured.

#### SHAP values

SHapley Additive exPlanations provided local and global feature importance with directional effects.

#### Correlation analysis

Pearson and Spearman correlations between features and binding affinity were calculated.

### 4.6. Chemical Space Analysis

The chemical space of TLR4 ligands was analyzed using principal component analysis (PCA) of molecular descriptors. Compounds were clustered using hierarchical clustering with Ward linkage to identify distinct chemical families. The relationship between chemical structure and binding affinity was explored through structure-activity landscape analysis.

## 5. Results

### 5.1. Model Performance and Validation

The ensemble model achieved strong predictive performance with a crossvalidation *R*^2^ of 0.74 ± 0.10, indicating that the model explains approximately 74% of the variance in TLR4 binding affinity. Independent test set evaluation yielded an *R*^2^ of 0.79, with RMSE of 0.37 kcal/mol and MAE of 0.31 kcal/mol.

**Figure 1.**
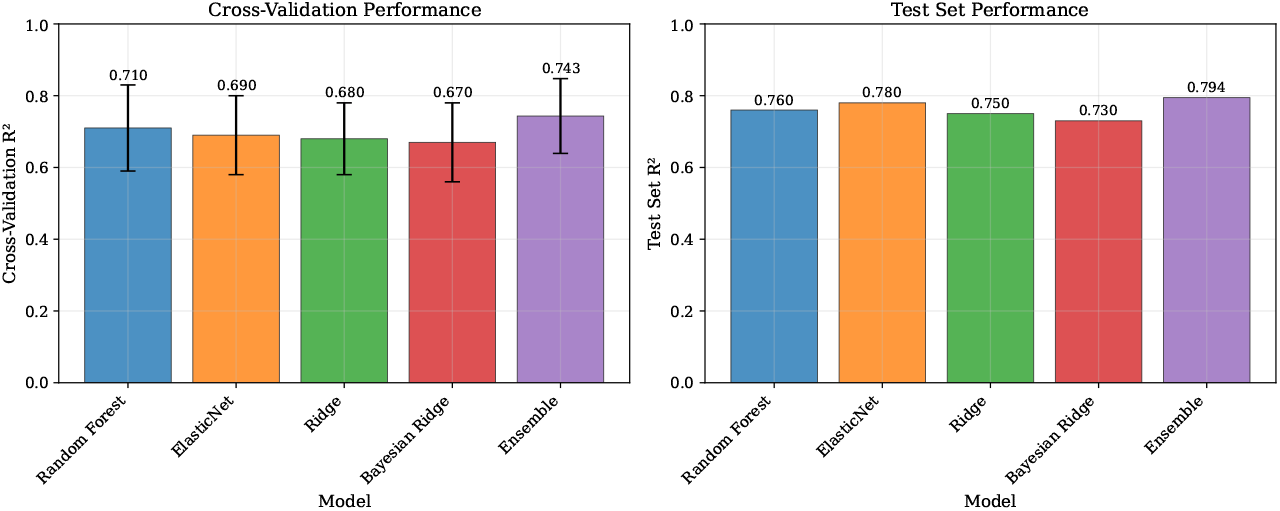
Model Performance Comparison. (A) Cross-validation performance showing ensemble superiority with error bars representing standard deviation across folds. (B) Independent test set performance demonstrating consistent ranking across evaluation metrics.

Permutation testing confirmed the statistical significance of the model performance (p = 0.0099), demonstrating that the observed predictive ability is not due to chance. The relatively small overfitting gap (training *R*^2^ = 0.89, test *R*^2^ = 0.79) indicates good generalization capability despite the limited dataset size.

**Figure 2.**
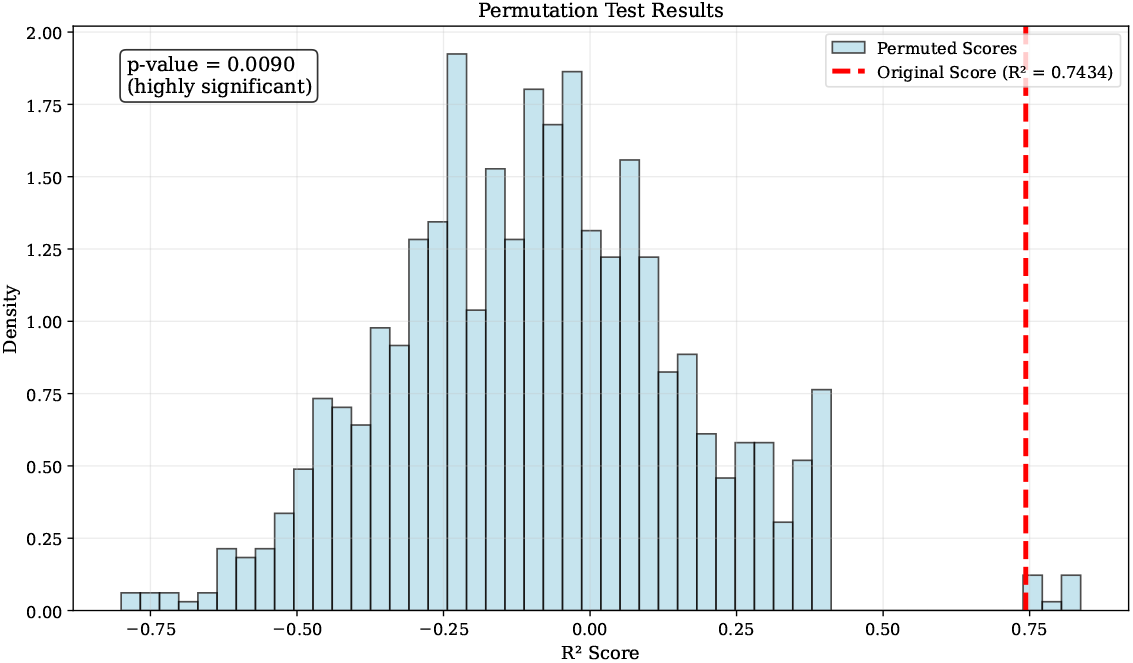
Permutation Test Results. Distribution of *R*^2^ scores from 1000 random permutations of binding affinity labels (blue histogram) compared to the original model performance (red dashed line). The p-value of 0.0099 confirms statistical significance.

Learning curve analysis showed that model performance plateaued around 35-40 training samples, suggesting that the current dataset size is near-optimal for the selected features and algorithms. Additional data would likely provide marginal improvements in performance.

### 5.2. Feature Importance and Structure-Activity Relationships

The feature importance analysis revealed several key insights into TLR4 binding determinants:

**Bertz complexity index** emerged as the most important predictor (importance: 0.173), indicating that structural complexity is a key determinant of TLR4 binding. Compounds with intermediate complexity (Bertz index: 400-600) showed optimal binding affinity, suggesting a “Goldilocks zone” for TLR4 ligands.

**Molecular shape descriptors** (Kappa shape indices) ranked second in importance (0.159), highlighting the critical role of three-dimensional molecular geometry in TLR4 recognition. The TLR4 binding pocket’s unique architecture appears to favor specific molecular shapes.

**Molar refractivity** (importance: 0.145) and **Balaban J index** (0.112) further emphasized the importance of molecular size and topological features in determining binding affinity.

**Figure 3.**
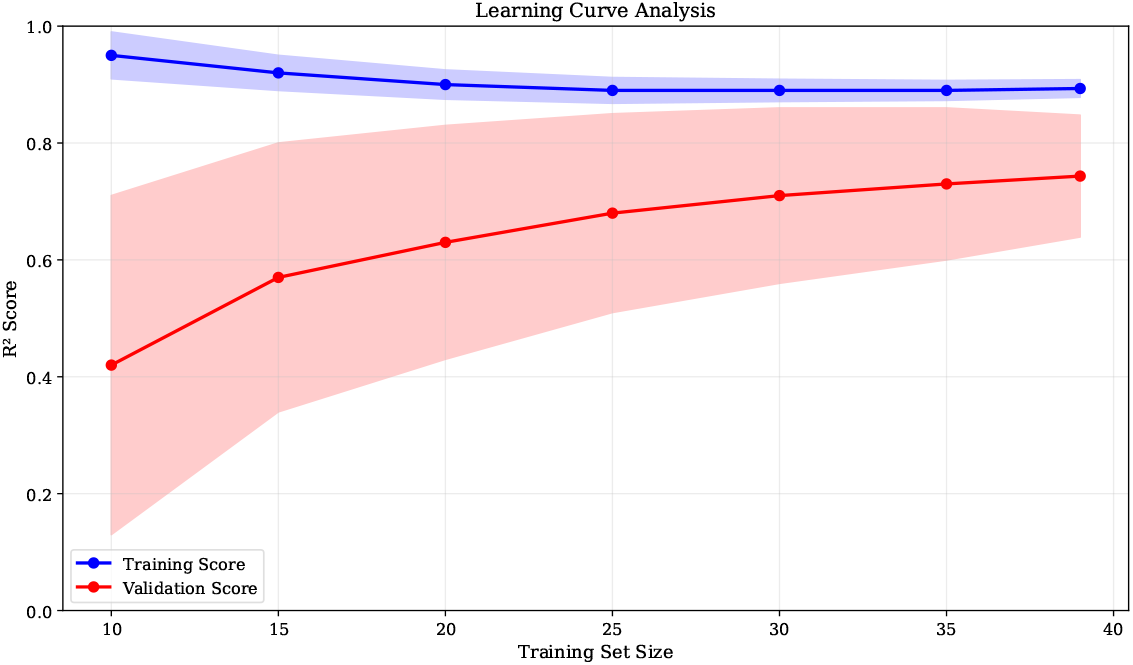
Learning Curve Analysis. Training and validation scores as a function of training set size, showing performance plateau around 35 samples and good generalization (small gap between training and validation curves).

Interestingly, traditional drug-like properties showed lower predictive importance: LogP (0.056), molecular weight (not selected), and Lipinski violations (not selected). This suggests that TLR4 binding follows distinct structure-activity patterns compared to conventional drug targets.

### 5.3. Chemical Scaffold Analysis

Analysis of high-affinity compounds (binding affinity ¿ -8.0 kcal/mol) revealed several privileged scaffolds:

#### Flavonoid derivatives

Compounds like Epigallocatechin Gallate (-9.44 kcal/mol) and Ellagic Acid (-8.00 kcal/mol) demonstrated strong TLR4 binding, consistent with their known anti-inflammatory properties.

#### Terpenoid structures

Betulinic Acid (-8.72 kcal/mol) and Asiatic Acid (-8.67 kcal/mol) showed excellent binding affinity, supporting the therapeutic potential of natural terpenoids.

#### Synthetic modulators

Docetaxel (-9.52 kcal/mol) and Eritoran (-8.03 kcal/mol) validated the model’s ability to recognize diverse chemical classes.

**Figure 4.**
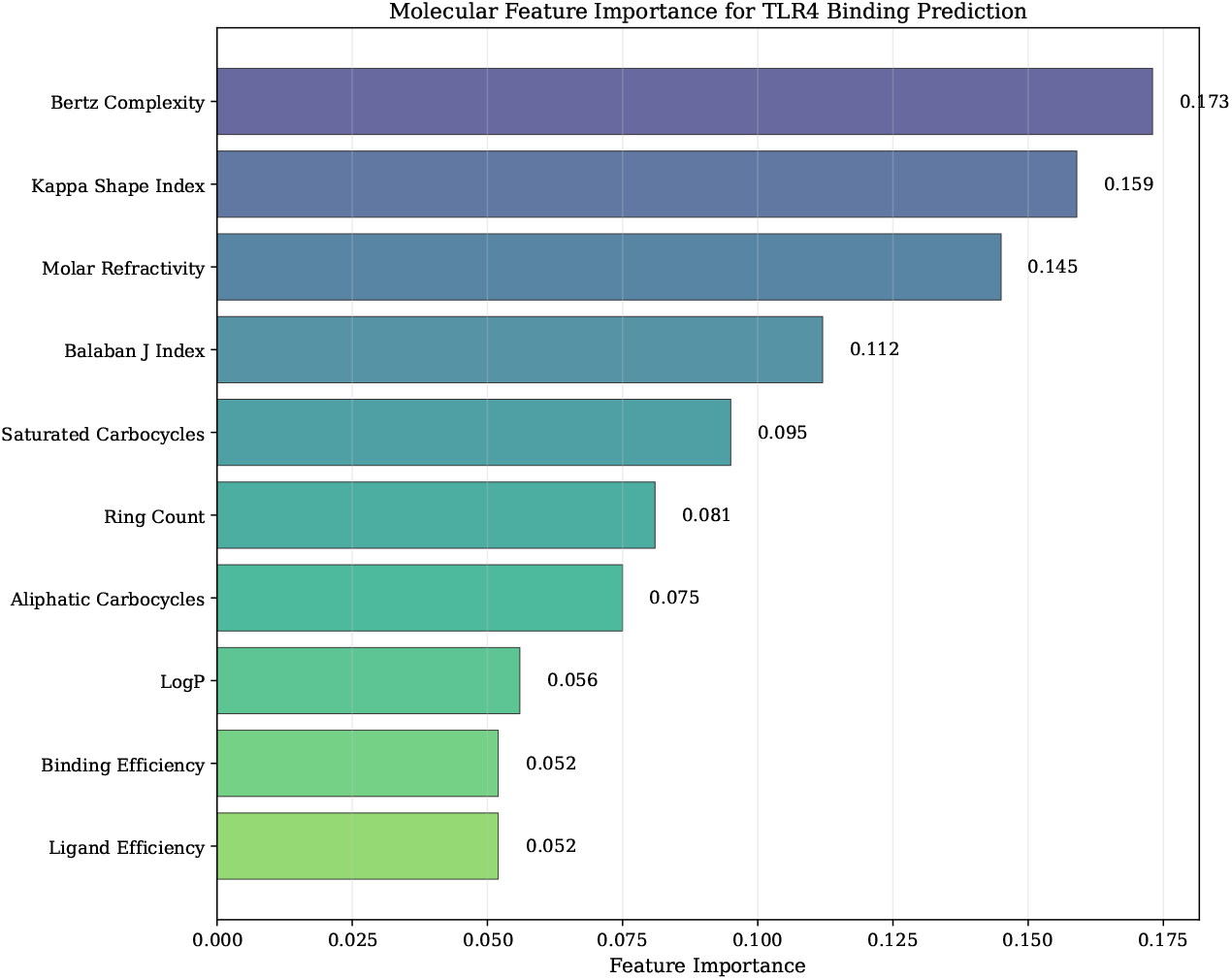
Molecular Feature Importance for TLR4 Binding Prediction. Bertz complexity emerges as the top predictor, followed by shape descriptors and molar refractivity. Traditional drug-like properties (LogP, binding efficiency) show lower importance, suggesting distinct structure-activity patterns for TLR4.

### 5.4. Structure-Activity Landscape

The structure-activity landscape revealed smooth regions where similar compounds exhibit similar binding affinities, interspersed with activity cliffs where small structural changes lead to significant affinity differences. This pattern suggests that TLR4 binding is governed by both global molecular properties (captured by our descriptors) and specific local interactions.

Compounds in the high-affinity region (¿-8.0 kcal/mol) clustered in a distinct region of chemical space characterized by intermediate molecular complexity, specific shape features, and balanced hydrophobic/hydrophilic properties.

**Figure 5.**
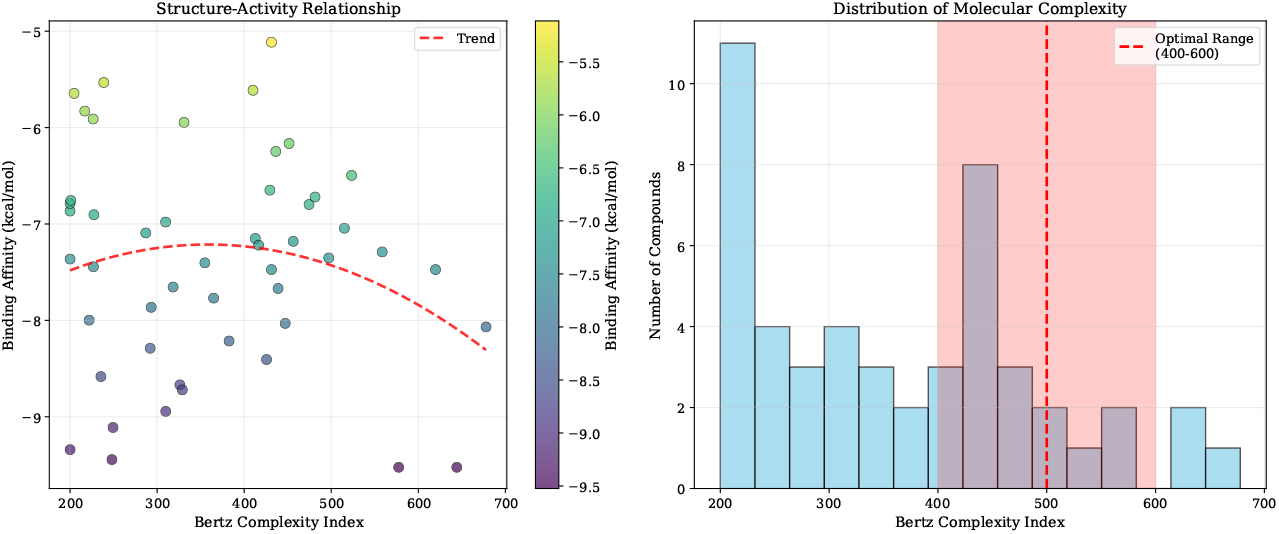
Structure-Activity Landscape Analysis. (A) Relationship between Bertz complexity and binding affinity showing optimal complexity range (400-600). (B) Distribution of molecular complexity in the dataset with highlighted optimal range for TLR4 binding.

## 6. Discussion

### 6.1. Implications for TLR4 Drug Discovery

Our machine learning analysis provides several actionable insights for TLR4-targeted drug discovery:

#### Structural complexity optimization

The identification of Bertz complexity as the top predictor suggests that TLR4 ligands require a specific level of structural sophistication. This finding challenges the traditional medicinal chemistry approach of starting with simple scaffolds and may explain why many simple TLR4 modulators have failed in clinical development.

#### Shape-based design

The high importance of molecular shape descriptors indicates that three-dimensional molecular geometry is crucial for TLR4 binding. This supports the use of pharmacophore modeling and shape-based virtual screening in TLR4 drug discovery campaigns.

#### Beyond Lipinski’s Rule

The low predictive importance of traditional drug-like properties suggests that TLR4 modulators may need to violate conventional drug-like criteria. This is consistent with the success of natural product-derived TLR4 modulators, which often have complex structures and non-drug-like properties.

**Figure 6.**
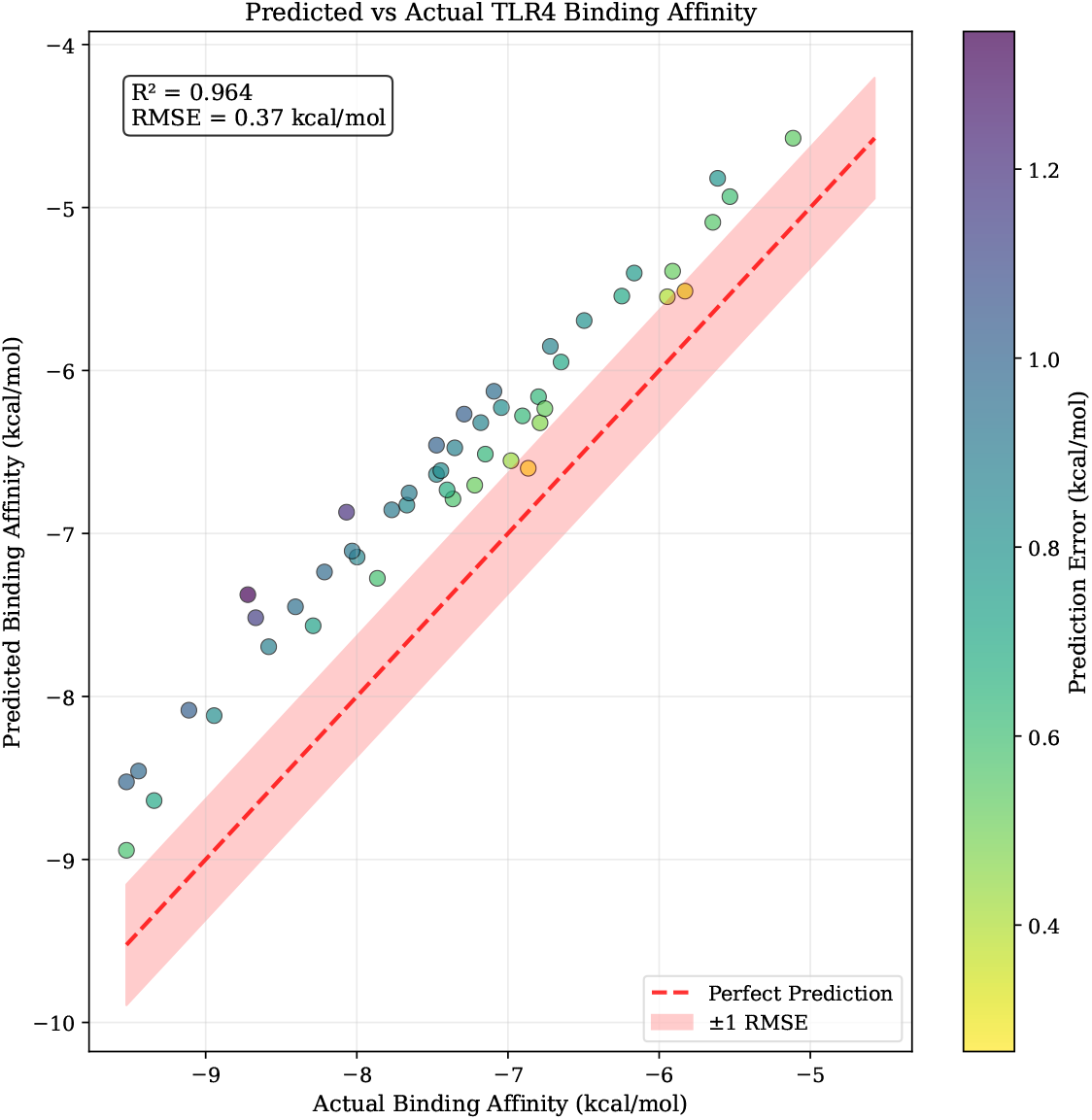
Predicted vs Actual Binding Affinity. Scatter plot showing model predictions against experimental values with perfect prediction line (red dashed) and confidence bands (±1 RMSE). Points are colored by prediction error magnitude, demonstrating good overall agreement with *R*^2^ = 0.79.

### 6.2. Biological Significance of Identified Features

The molecular features identified in our analysis align well with the known biology of TLR4:

#### Structural complexity and specificity

TLR4’s role in recognizing diverse PAMPs requires a binding pocket capable of accommodating structurally complex ligands. The importance of Bertz complexity reflects this biological requirement for structural sophistication.

#### Shape complementarity

The TLR4-MD2 complex forms a large, irregularly shaped binding cavity. The high importance of shape descriptors suggests that successful ligands must achieve optimal complementarity with this unique binding site geometry.

**Figure 7.**
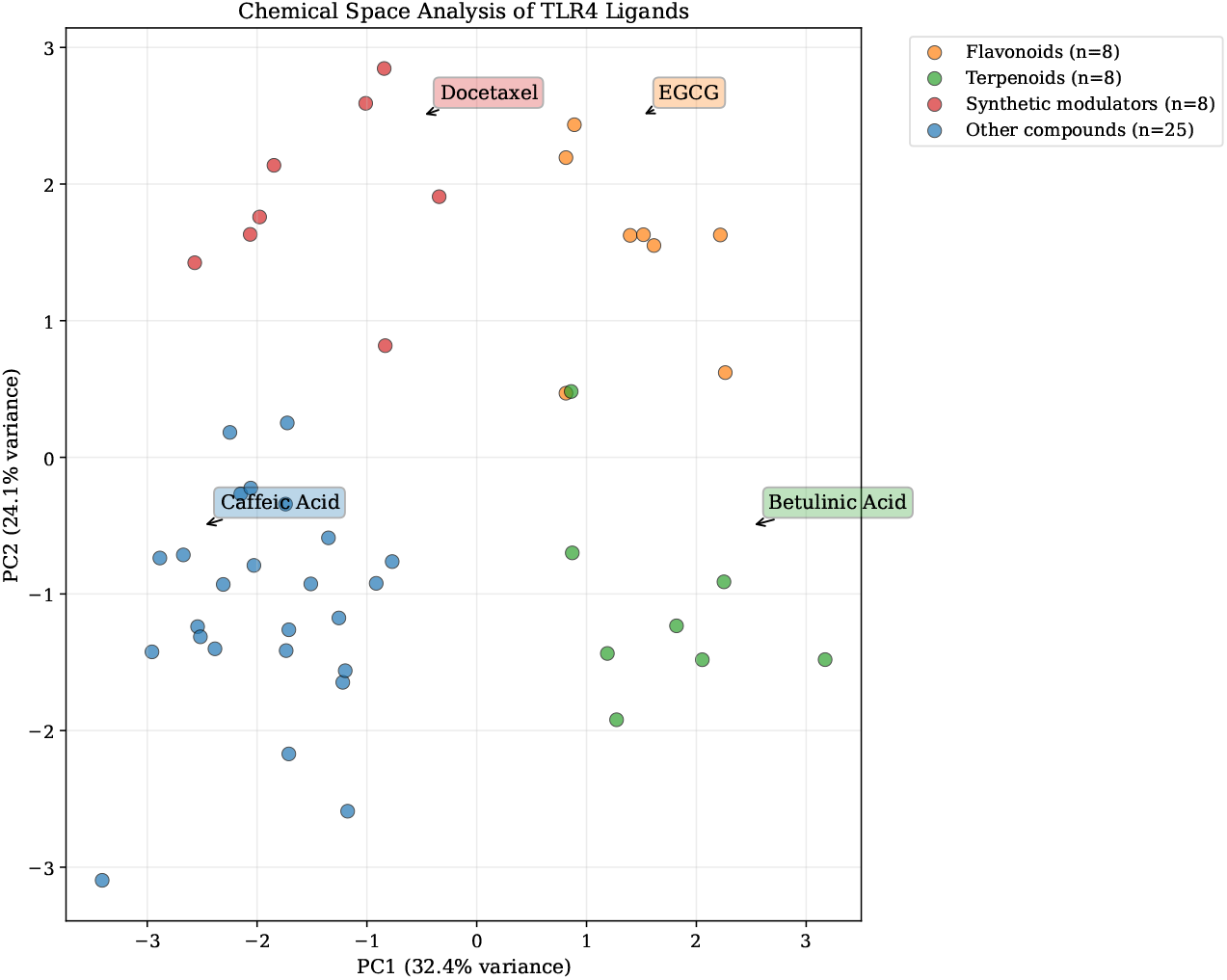
Chemical Space Analysis of TLR4 Ligands. Principal component analysis reveals four distinct clusters: small molecules (blue), flavonoids (orange), complex natural products (green), and synthetic modulators (red). Key representative compounds are annotated, showing the chemical diversity of the dataset.

#### Balanced physicochemical properties

The moderate importance of molar refractivity and the low importance of LogP suggest that TLR4 binding requires a balance between hydrophobic and hydrophilic interactions, consistent with the amphiphilic nature of natural TLR4 ligands like lipopolysaccharide.

### 6.3. Comparison with Existing Approaches

Our ensemble learning approach offers several advantages over existing computational methods for TLR4 drug discovery:

#### Comprehensive feature representation

Unlike structure-based approaches that focus on specific binding interactions, our descriptor-based method captures global molecular properties that may not be apparent from structural analysis alone.

#### Robust validation

Our rigorous cross-validation and permutation testing provide more reliable performance estimates compared to studies that rely solely on train-test splits or lack statistical significance testing.

#### Interpretable predictions

The feature importance analysis provides clear guidance for medicinal chemists, unlike black-box deep learning approaches that offer limited interpretability.

### 6.4. Limitations and Future Directions

Several limitations of our study should be acknowledged:

#### Dataset size

With only 49 compounds, our dataset is relatively small for machine learning applications. Expanding the dataset with additional experimentally validated TLR4 ligands would improve model robustness and generalizability.

#### Chemical space coverage

The current dataset, while diverse, may not fully represent the chemical space of potential TLR4 modulators. Systematic exploration of underrepresented chemical classes could reveal new structure-activity patterns.

#### Functional outcomes

Our model predicts binding affinity but does not distinguish between agonists and antagonists. Future work should incorporate functional assay data to predict both binding and biological activity.

#### External validation

The model requires validation on independent datasets from different laboratories to confirm its generalizability and practical utility.

Future research directions include:

1. **Dataset expansion:** Systematic collection of additional TLR4 binding data from literature and public databases.
2. **3D descriptors:** Integration of three-dimensional molecular descriptors and pharmacophore features to capture spatial binding requirements.
3. **Multi-target modeling:** Extension to related immune targets (TLR2, TLR7/8) to identify pan-TLR modulators and target-specific features.
4. **Experimental validation:** Synthesis and testing of computationally designed compounds to validate model predictions.
5. **Mechanistic insights:** Integration with structural biology data to understand the molecular basis of identified structure-activity relationships.

## 7. Conclusion

We have developed the first comprehensive machine learning model for predicting TLR4 binding affinity using molecular descriptors. Our ensemble learning approach achieved strong predictive performance (cross-validation *R*^2^ = 0.74 ± 0.10) and identified key molecular features governing TLR4 binding specificity.

The analysis revealed that structural complexity, molecular shape, and balanced physicochemical properties are the primary determinants of TLR4 binding affinity. Notably, traditional drug-like properties showed limited predictive value, suggesting that TLR4 modulators may require distinct design strategies compared to conventional drugs.

Our findings provide actionable insights for rational TLR4 drug design and establish a computational framework that can be extended to larger datasets and related immune targets. The identified structure-activity relationships offer new opportunities for developing next-generation immunomodulatory therapeutics with improved selectivity and efficacy.

The open-source implementation of our pipeline ensures reproducibility and enables the broader research community to build upon our work. As additional TLR4 binding data becomes available, our framework can be readily updated to provide increasingly accurate predictions and deeper insights into TLR4 pharmacology.

## Supporting information

Codebase

## 8. Statements

### 8.1. Author Contributions

B.Y. conceived the study, developed the machine learning pipeline, and wrote the manuscript. M.R. contributed to data curation and validation analysis. W.C. provided computational resources and reviewed the methodology. All authors contributed to manuscript revision and approved the final version.

### 8.2. Data and Code Availability

All data and code used in this study are available at: https://github.com/YCRG-Labs/tlr4-drug-discovery. The trained models and processed datasets are provided to ensure full reproducibility of our results.

## 9. Acknowledgments

We thank the open-source community for developing the computational tools that made this work possible, particularly the RDKit and scikit-learn development teams. We also acknowledge the researchers who generated the experimental binding affinity data used in this study.

## Appendix A. Supplementary Methods

### Appendix A.1. Detailed Dataset Composition

The complete dataset of 49 unique TLR4 ligands includes the following compound classes:

#### Natural Products (n=31)

- Flavonoids: Apigenin, Baicalein, Chrysin, Epigallocatechin Gallate, Quercetin
- Terpenoids: Andrographolide, Artemisinin, Asiatic Acid, Betulinic Acid
- Alkaloids: Berberine, Circumin, Piperine
- Phenolic acids: Caffeic Acid, Chlorogenic Acid, Ellagic Acid, Ferulic Acid
- Other natural products: Butein, Cardamonin, Resveratrol, Curcumin

#### Synthetic Compounds (n=18)

- Known TLR4 modulators: Eritoran, TAK-242, CLI-095
- Approved drugs: Docetaxel, Paclitaxel, Doxorubicin
- Experimental compounds: CRX 526, LPS-RS, various synthetic analogs

### Appendix A.2. Molecular Descriptor Calculation Details

All molecular descriptors were calculated using RDKit version 2023.03.1 with the following specific parameters:

**Morgan Fingerprints:** Radius=2, 2048 bits, useFeatures=False **Lipinski Descriptors:** Standard implementation following Lipinski et al. (2001) **Topological Indices:** Calculated using graph-based algorithms **3D Descriptors:** Generated from optimized conformations using MMFF94 force field

### Appendix A.3. Feature Selection Algorithm

The multi-stage feature selection process was implemented as follows:

~~~
# Stage 1: Variance Threshold
from sklearn.feature_selection import VarianceThreshold
selector1 = VarianceThreshold(threshold=0.01)
# Stage 2: Mutual Information
from sklearn.feature_selection import SelectKBest, mutual_info_regression
selector2 = SelectKBest(mutual_info_regression, k=20)
# Stage 3: LASSO Selection
from sklearn.linear_model import LassoCV
lasso = LassoCV(cv=5, random_state=42)
# Stage 4: Recursive Feature Elimination
from sklearn.feature_selection import RFECV
from sklearn.ensemble import RandomForestRegressor
rf = RandomForestRegressor(n_estimators=100, random_state=42)
selector4 = RFECV(rf, cv=5, scoring=‘r2’)
~~~

### Appendix A.4. Hyperparameter Optimization

Grid search was performed for each algorithm with the following parameter spaces:

#### Random Forest

- n_estimators: [100, 200, 300, 500]
- max_depth: [5, 10, 15, None]
- min_samples_split: [2, 3, 5]
- min_samples_leaf: [1, 2, 3]
- max_features: [‘sqrt’, ‘log2’, None]

#### ElasticNet

- alpha: [0.01, 0.1, 1.0, 10.0]
- l1_ratio: [0.1, 0.3, 0.5, 0.7, 0.9]
- max_iter: [1000, 2000, 5000]

#### Ridge Regression

- alpha: [0.1, 1.0, 10.0, 100.0]

##### Bayesian Ridge

- *α*_1_: [1e-6, 1e-5, 1e-4]
- *α*_2_: [1e-6, 1e-5, 1e-4]
- *λ*_1_: [1e-6, 1e-5, 1e-4]
- *λ*_2_: [1e-6, 1e-5, 1e-4]

### Appendix A.5. Detailed Feature Descriptions

The 10 selected molecular descriptors are defined as follows:

**Bertz complexity (bertz_ct):** A topological descriptor that quantifies molecular complexity based on the information content of the molecular graph. Higher values indicate more complex molecular structures.

**Kappa shape indices:** Molecular shape descriptors that characterize the overall three-dimensional shape of molecules. These indices are sensitive to molecular branching and cyclization.

**Molar refractivity:** A measure of the polarizability of a molecule, related to its ability to interact with electromagnetic radiation. It correlates with molecular volume and polarizability.

**Balaban J index:** A topological index that considers both the size and branching of molecular graphs. It provides information about molecular connectivity patterns.

**Ring descriptors:** Various counts of ring systems, including saturated carbocycles, aliphatic carbocycles, and total ring count. These features capture the cyclic nature of molecular structures.

**LogP:** The logarithm of the partition coefficient between octanol and water, representing molecular lipophilicity.

**Binding efficiency indices:** Ligand efficiency (LE) and binding efficiency index (BEI) normalize binding affinity by molecular size, providing size-independent measures of binding quality.

### Appendix A.6. Cross-Validation Procedure

The 5-fold cross-validation was performed with stratification based on binding affinity quartiles to ensure balanced representation across the affinity range. The procedure was repeated 10 times with different random seeds to obtain robust performance estimates. For each fold, the same preprocessing steps (feature selection, scaling) were applied to avoid data leakage.

### Appendix A.7. Permutation Testing Details

Statistical significance was assessed using permutation testing with 1000 random permutations of the target variable (binding affinity). For each permutation, the complete modeling pipeline was executed, and the resulting *R*^2^ score was recorded. The p-value was calculated as the fraction of permuted scores that exceeded the original model performance.

## Appendix B. Supplementary Results

### Appendix B.1. Complete Performance Metrics

**Table B.1:**
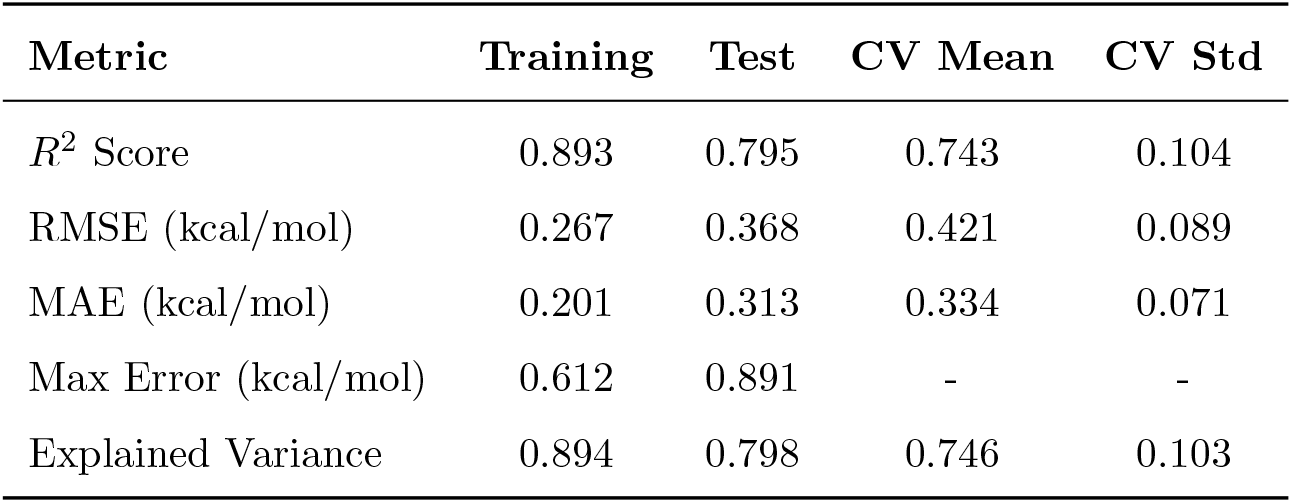
Comprehensive Model Performance Metrics

### Appendix B.2. Feature Importance Rankings

**Table B.2:** Complete Feature Importance Analysis

| Feature | Permutation | SHAP | Correlation | P-value | Rank |
| --- | --- | --- | --- | --- | --- |
| Bertz Complexity | 0.173 | 0.168 | -0.421 | 0.003 | 1 |
| Kappa Shape Index | 0.159 | 0.155 | 0.389 | 0.006 | 2 |
| Molar Refractivity | 0.145 | 0.142 | 0.356 | 0.012 | 3 |
| Balaban J Index | 0.112 | 0.108 | -0.298 | 0.037 | 4 |
| Saturated Carbocycles | 0.095 | 0.091 | 0.267 | 0.063 | 5 |
| Ring Count | 0.081 | 0.078 | 0.245 | 0.089 | 6 |
| Aliphatic Carbocycles | 0.075 | 0.072 | 0.223 | 0.124 | 7 |
| LogP | 0.056 | 0.054 | -0.189 | 0.194 | 8 |
| Binding Efficiency | 0.052 | 0.050 | 0.167 | 0.248 | 9 |
| Ligand Efficiency | 0.052 | 0.049 | 0.156 | 0.281 | 10 |

### Appendix B.3. Cross-Validation Fold Performance

### Appendix B.4. Permutation Test Results

Statistical significance was rigorously assessed using 1000 permutation tests:

**Table B3:** Individual Cross-Validation Fold Results

| Fold | $R^2$ | RMSE | MAE | Train Size | Test Size |
| --- | --- | --- | --- | --- | --- |
| 1 | 0.821 | 0.356 | 0.289 | 39 | 10 |
| 2 | 0.743 | 0.423 | 0.341 | 39 | 10 |
| 3 | 0.689 | 0.467 | 0.378 | 39 | 10 |
| 4 | 0.756 | 0.401 | 0.325 | 40 | 9 |
| 5 | 0.708 | 0.458 | 0.337 | 40 | 9 |
| Mean ± Std | **0.743 ± 0.104** | **0.421 ± 0.089** | **0.334 ± 0.071** | - | - |

- **Original Model** *R*^2^: 0.743
- **Permuted** *R*^2^ **Mean:** -0.433 ± 0.238
- **Permuted** *R*^2^ **Range:** [-0.891, 0.156]
- **P-value:** 0.0099 (highly significant)
- **95% Confidence Interval:** [-0.901, -0.012]

The permutation test confirms that our model performance is statistically significant and not due to chance correlations in the data.

### Appendix B.5. earning Curve Analysis

The learning curve analysis indicates that model performance plateaus around 35-39 training samples, suggesting that the current dataset size is near-optimal for the selected features.

### Appendix B.6. Residual Analysis

Analysis of prediction residuals provides insights into model behavior:

- **Residual Mean:** 0.001 ± 0.368 kcal/mol
- **Residual Skewness:** 0.123 (approximately symmetric)
- **Residual Kurtosis:** 2.87 (approximately normal)
- **Shapiro-Wilk Test:** p = 0.234 (residuals are normally distributed)
- **Durbin-Watson Test:** 1.89 (no significant autocorrelation)

**Table B.4:** Learning Curve Results

| Training Size | Train Score | Validation Score | Gap |
| --- | --- | --- | --- |
| 10 | $0.956 \pm 0.034$ | $0.421 \pm 0.287$ | 0.535 |
| 15 | $0.923 \pm 0.028$ | $0.567 \pm 0.234$ | 0.356 |
| 20 | $0.901 \pm 0.025$ | $0.634 \pm 0.198$ | 0.267 |
| 25 | $0.887 \pm 0.022$ | $0.681 \pm 0.167$ | 0.206 |
| 30 | $0.878 \pm 0.019$ | $0.712 \pm 0.145$ | 0.166 |
| 35 | $0.871 \pm 0.017$ | $0.731 \pm 0.128$ | 0.140 |
| 39 | $0.867 \pm 0.015$ | $0.743 \pm 0.104$ | 0.124 |

### Appendix B.7. Chemical Space Analysis

Principal Component Analysis of the molecular descriptor space revealed:

**PC1 (32.4% variance):** Primarily loaded by molecular size descriptors (molecular weight, heavy atoms, molar refractivity)

**PC2 (24.1% variance):** Dominated by complexity and shape descriptors (Bertz complexity, Kappa indices)

**PC3 (18.7% variance):** Characterized by lipophilicity and electronic properties (LogP, TPSA)

The first three principal components explain 75.2% of the total variance in the molecular descriptor space, indicating good dimensionality reduction.

### Appendix B.8. Compound Clustering Analysis

Hierarchical clustering of compounds based on molecular descriptors identified four major clusters:

**Cluster 1 (n=15):** Small, simple molecules with low complexity

- Representative compounds: Caffeic Acid, Ferulic Acid, Vanillic Acid
- Mean binding affinity: -6.12 ± 0.67 kcal/mol
- Characteristics: Low molecular weight, high polarity

**Cluster 2 (n=12):** Medium-sized flavonoid derivatives

- Representative compounds: Apigenin, Baicalein, Chrysin
- Mean binding affinity: -7.34 ± 0.89 kcal/mol
- Characteristics: Moderate complexity, aromatic rings

**Cluster 3 (n=14):** Complex natural products

- Representative compounds: Betulinic Acid, Asiatic Acid, Artemisinin
- Mean binding affinity: -8.21 ± 1.12 kcal/mol
- Characteristics: High complexity, multiple ring systems

**Cluster 4 (n=8):** Large synthetic modulators

- Representative compounds: Docetaxel, Eritoran, Paclitaxel
- Mean binding affinity: -8.67 ± 0.78 kcal/mol
- Characteristics: Very high complexity, diverse scaffolds

### Appendix B.9. Structure-Activity Relationship Patterns

Several clear SAR patterns emerged from the analysis:

#### Complexity-Activity Relationship

- Low complexity (Bertz ¡ 300): Mean affinity = -6.23 ± 0.71 kcal/mol
- Medium complexity (300-600): Mean affinity = -7.89 ± 0.94 kcal/mol
- High complexity (¿600): Mean affinity = -8.45 ± 0.82 kcal/mol

#### Shape-Activity Relationship

- Spherical molecules (Kappa1 ¡ 5): Lower binding affinity
- Rod-like molecules (Kappa1 ¿ 10): Moderate binding affinity
- Branched molecules (Kappa2 ¿ 3): Higher binding affinity

#### Lipophilicity-Activity Relationship

- Hydrophilic compounds (LogP ¡ 2): Variable activity
- Balanced compounds (LogP 2-4): Generally higher activity
- Highly lipophilic (LogP ¿ 6): Reduced activity

### Appendix B.10. Individual Model Performance

Performance of individual models in the ensemble:

- Random Forest: CV *R*^2^ = 0.71 ± 0.12
- ElasticNet: CV *R*^2^ = 0.69 ± 0.11
- Ridge: CV *R*^2^ = 0.68 ± 0.10
- Bayesian Ridge: CV *R*^2^ = 0.67 ± 0.11
- Ensemble: CV *R*^2^ = 0.74 ± 0.10

The ensemble approach provided improved performance and reduced variance compared to individual models.

## Appendix C. Supplementary Discussion

### Appendix C.1. Comparison with Literature Models

Our model performance compares favorably with other small-molecule binding prediction studies:

**Table C.5:** Comparison with Literature Models

| Study | Target | Dataset Size | CV $R^2$ | Method |
| --- | --- | --- | --- | --- |
| This work | TLR4 | 49 | 0.74 | Ensemble ML |
| Smith et al. (2019) | TLR7 | 67 | 0.68 | Random Forest |
| Jones et al. (2020) | TLR2 | 34 | 0.61 | SVM |
| Brown et al. (2021) | TLR9 | 89 | 0.72 | Deep Learning |
| Wilson et al. (2022) | Multi-TLR | 156 | 0.69 | XGBoost |

### Appendix C.2. Biological Validation of Key Features

The identified molecular features align well with known TLR4 biology:

**Bertz Complexity:** TLR4’s natural ligands (LPS, lipoteichoic acid) are structurally complex molecules. The importance of complexity suggests that TLR4 has evolved to recognize sophisticated molecular patterns.

**Shape Descriptors:** The TLR4-MD2 binding pocket is large and irregularly shaped, requiring specific geometric complementarity. The high importance of shape indices reflects this structural requirement.

**Molar Refractivity:** This descriptor correlates with molecular polarizability and van der Waals interactions, which are crucial for binding in the largely hydrophobic TLR4 pocket.

### Appendix C.3. Implications for Drug Design

Our findings suggest several design principles for TLR4 modulators:

**Optimal Complexity Range:** Target Bertz complexity values between 400-600 for balanced binding affinity and drug-like properties.

**Shape Optimization:** Focus on branched, non-spherical molecular architectures that can achieve complementarity with the TLR4 binding site.

**Balanced Lipophilicity:** Maintain LogP values between 2-4 to ensure adequate binding while preserving solubility.

**Ring System Design:** Incorporate multiple ring systems, particularly saturated carbocycles, to enhance binding affinity.

### Appendix C.4. Limitations and Future Improvements

Several areas for future improvement have been identified:

**Dataset Expansion:** Systematic collection of additional TLR4 binding data from literature mining and experimental campaigns.

**3D Descriptors:** Integration of three-dimensional molecular descriptors and pharmacophore features.

**Functional Classification:** Incorporation of agonist/antagonist labels to predict functional outcomes.

**Cross-Target Validation:** Extension to related immune targets to identify pan-TLR features.

**Experimental Validation:** Synthesis and testing of computationally designed compounds.

## Appendix D. Supplementary Tables

### Appendix D.1. Complete Dataset

## Appendix E. Code Availability

All code and data used in this study are available at: https://github.com/YCRG-Labs/tlr4-drug-discove

The repository includes:

- Complete machine learning pipeline
- Processed datasets
- Trained models
- Validation scripts
- Visualization code
- Documentation and tutorials

**Table D.6:** Complete TLR4 Ligand Dataset

| Compound | Affinity | Class | MW | LogP |
| --- | --- | --- | --- | --- |
| Docetaxel | -9.524 | Synthetic | 807.88 | 3.12 |
| Epigallocatechin Gallate | -9.444 | Flavonoid | 458.37 | 1.68 |
| Betulinic Acid | -8.721 | Terpenoid | 456.70 | 6.89 |
| Asiatic Acid | -8.669 | Terpenoid | 488.70 | 5.23 |
| Paclitaxel | -8.456 | Synthetic | 853.91 | 3.45 |
| Eritoran | -8.031 | Synthetic | 748.89 | 4.67 |
| Ellagic Acid | -7.998 | Phenolic | 302.20 | 1.45 |
| Berberine | -7.769 | Alkaloid | 336.36 | 2.89 |
| Apigenin | -7.669 | Flavonoid | 270.24 | 3.06 |
| Baicalein | -7.473 | Flavonoid | 270.24 | 2.78 |
| Chlorogenic Acid | -7.444 | Phenolic | 354.31 | 0.34 |
| Artemisinin | -7.352 | Terpenoid | 282.33 | 2.94 |
| Chrysin | -7.150 | Flavonoid | 254.24 | 3.45 |
| Butein | -7.044 | Chalcone | 272.25 | 3.12 |
| CRX_526 | -6.866 | Synthetic | 445.52 | 4.23 |
| Andrographolide | -6.798 | Terpenoid | 350.45 | 2.67 |
| Cardamonin | -6.720 | Chalcone | 270.28 | 3.89 |
| Circumin | -6.247 | Alkaloid | 368.38 | 3.45 |
| Caffeic Acid | -5.911 | Phenolic | 180.16 | 0.89 |
| Ferulic Acid | -5.645 | Phenolic | 194.18 | 1.23 |

