## Supplementary material for "Machine Learning-Driven Prediction of TLR4 Binding Affinity: A Comprehensive Molecular Feature Analysis for Drug Discovery": Codebase: academic_validation.py

```
#!/usr/bin/env python3
"""
Academic Validation of TLR4 Binding Prediction Pipeline.

This script performs rigorous academic validation to ensure:
1. No data leakage or overfitting
2. Proper statistical validation
3. Realistic performance expectations
4. Methodological soundness
5. Reproducibility

Author: Kiro AI Assistant
"""

import pandas as pd
import numpy as np
import logging
from pathlib import Path
import joblib
import json
from sklearn.model_selection import (
    cross_val_score, LeaveOneOut, StratifiedKFold, 
    permutation_test_score, learning_curve
)
from sklearn.metrics import r2_score, mean_squared_error, mean_absolute_error
from scipy import stats
from scipy.stats import pearsonr, spearmanr
import matplotlib.pyplot as plt
import seaborn as sns
import warnings

warnings.filterwarnings('ignore')

logging.basicConfig(level=logging.INFO)
logger = logging.getLogger(__name__)


def load_publication_results():
    """Load the publication model and results."""
    try:
        model = joblib.load('results/publication_model.pkl')
        scaler = joblib.load('results/publication_scaler.pkl')
        
        with open('results/publication_features.json', 'r') as f:
            selected_features = json.load(f)
        
        with open('results/publication_metrics.json', 'r') as f:
            metrics = json.load(f)
        
        return model, scaler, selected_features, metrics
    except Exception as e:
        logger.error(f"Failed to load publication results: {e}")
        return None, None, None, None


def validate_data_integrity():
    """Validate data integrity and check for potential issues."""
    logger.info("=== DATA INTEGRITY VALIDATION ===")
    
    # Load processed data
    binding_df = pd.read_csv('data/processed/processed_logs.csv')
    
    # Check for data leakage
    logger.info("1. Checking for data leakage...")
    
    # Count conformations per base compound
    binding_df['base_compound'] = binding_df['ligand'].apply(
        lambda x: x.split('_conf_')[0] if '_conf_' in x else x
    )
    
    conformation_counts = binding_df['base_compound'].value_counts()
    total_conformations = len(binding_df)
    unique_compounds = len(conformation_counts)
    
    logger.info(f"  Total records: {total_conformations}")
    logger.info(f"  Unique base compounds: {unique_compounds}")
    logger.info(f"  Average conformations per compound: {total_conformations/unique_compounds:.2f}")
    
    if total_conformations > unique_compounds * 2:
        logger.warning(f"  ⚠️  High conformation ratio detected - ensure proper deduplication")
    else:
        logger.info(f"  ✅ Reasonable conformation ratio")
    
    # Check affinity distribution
    logger.info("2. Checking affinity distribution...")
    affinities = binding_df['affinity']
    
    logger.info(f"  Affinity range: {affinities.min():.3f} to {affinities.max():.3f} kcal/mol")
    logger.info(f"  Mean ± SD: {affinities.mean():.3f} ± {affinities.std():.3f} kcal/mol")
    logger.info(f"  Median: {affinities.median():.3f} kcal/mol")
    
    # Check for outliers
    q1, q3 = affinities.quantile([0.25, 0.75])
    iqr = q3 - q1
    outlier_threshold_low = q1 - 1.5 * iqr
    outlier_threshold_high = q3 + 1.5 * iqr
    
    outliers = affinities[(affinities < outlier_threshold_low) | (affinities > outlier_threshold_high)]
    logger.info(f"  Outliers (IQR method): {len(outliers)} ({len(outliers)/len(affinities)*100:.1f}%)")
    
    # Check for suspicious patterns
    if affinities.std() < 0.5:
        logger.warning("  ⚠️  Very low affinity variance - check data quality")
    elif affinities.std() > 3.0:
        logger.warning("  ⚠️  Very high affinity variance - check for outliers")
    else:
        logger.info("  ✅ Reasonable affinity variance")
    
    return {
        'total_records': total_conformations,
        'unique_compounds': unique_compounds,
        'affinity_stats': {
            'mean': affinities.mean(),
            'std': affinities.std(),
            'min': affinities.min(),
            'max': affinities.max(),
            'outliers': len(outliers)
        }
    }


def validate_model_performance(model, scaler, selected_features, metrics):
    """Validate model performance with rigorous statistical tests."""
    logger.info("=== MODEL PERFORMANCE VALIDATION ===")
    
    # Reload the integrated data
    from publication_ready_pipeline import (
        ComprehensiveFeatureExtractor, load_binding_data, integrate_features_and_binding
    )
    
    # Extract features
    extractor = ComprehensiveFeatureExtractor()
    features_df = extractor.extract_features_from_files("data/raw/pdbqt")
    binding_df = load_binding_data("data/processed/processed_logs.csv")
    integrated_df = integrate_features_and_binding(features_df, binding_df)
    
    # Prepare data
    X = integrated_df[selected_features]
    y = integrated_df['affinity']
    
    logger.info(f"Dataset: {len(X)} samples, {len(selected_features)} features")
    
    # 1. Leave-One-Out Cross-Validation (most rigorous for small datasets)
    logger.info("1. Leave-One-Out Cross-Validation...")
    loo = LeaveOneOut()
    loo_scores = cross_val_score(model, scaler.transform(X), y, cv=loo, scoring='r2')
    
    logger.info(f"  LOO CV R²: {loo_scores.mean():.4f} ± {loo_scores.std():.4f}")
    logger.info(f"  LOO CV range: {loo_scores.min():.4f} to {loo_scores.max():.4f}")
    
    # 2. Permutation Test (test for chance correlation)
    logger.info("2. Permutation Test...")
    perm_score, perm_scores, perm_pvalue = permutation_test_score(
        model, scaler.transform(X), y, scoring='r2', cv=5, n_permutations=100, random_state=42
    )
    
    logger.info(f"  Original R²: {perm_score:.4f}")
    logger.info(f"  Permuted R² mean: {perm_scores.mean():.4f} ± {perm_scores.std():.4f}")
    logger.info(f"  P-value: {perm_pvalue:.6f}")
    
    if perm_pvalue < 0.001:
        logger.info("  ✅ Highly significant (p < 0.001)")
    elif perm_pvalue < 0.01:
        logger.info("  ✅ Very significant (p < 0.01)")
    elif perm_pvalue < 0.05:
        logger.info("  ✅ Significant (p < 0.05)")
    else:
        logger.warning("  ⚠️  Not significant (p ≥ 0.05)")
    
    # 3. Learning Curve Analysis
    logger.info("3. Learning Curve Analysis...")
    train_sizes = np.linspace(0.3, 1.0, 8)
    train_sizes_abs, train_scores, val_scores = learning_curve(
        model, scaler.transform(X), y, train_sizes=train_sizes, cv=5, scoring='r2'
    )
    
    train_mean = train_scores.mean(axis=1)
    train_std = train_scores.std(axis=1)
    val_mean = val_scores.mean(axis=1)
    val_std = val_scores.std(axis=1)
    
    logger.info(f"  Final training R²: {train_mean[-1]:.4f} ± {train_std[-1]:.4f}")
    logger.info(f"  Final validation R²: {val_mean[-1]:.4f} ± {val_std[-1]:.4f}")
    
    # Check for overfitting
    overfitting_gap = train_mean[-1] - val_mean[-1]
    if overfitting_gap > 0.3:
        logger.warning(f"  ⚠️  High overfitting detected (gap = {overfitting_gap:.3f})")
    elif overfitting_gap > 0.15:
        logger.warning(f"  ⚠️  Moderate overfitting detected (gap = {overfitting_gap:.3f})")
    else:
        logger.info(f"  ✅ Acceptable overfitting (gap = {overfitting_gap:.3f})")
    
    # 4. Feature Stability Analysis
    logger.info("4. Feature Stability Analysis...")
    from sklearn.utils import resample
    
    feature_importance_bootstrap = []
    n_bootstrap = 50
    
    for i in range(n_bootstrap):
        # Bootstrap sample
        X_boot, y_boot = resample(X, y, random_state=i)
        X_boot_scaled = scaler.transform(X_boot)
        
        # Fit model and get feature importance
        model.fit(X_boot_scaled, y_boot)
        if hasattr(model.named_estimators_['rf'], 'feature_importances_'):
            importance = model.named_estimators_['rf'].feature_importances_
            feature_importance_bootstrap.append(importance)
    
    if feature_importance_bootstrap:
        importance_array = np.array(feature_importance_bootstrap)
        importance_mean = importance_array.mean(axis=0)
        importance_std = importance_array.std(axis=0)
        
        logger.info("  Top 5 most stable features:")
        for i in np.argsort(importance_mean)[-5:][::-1]:
            feature_name = selected_features[i]
            mean_imp = importance_mean[i]
            std_imp = importance_std[i]
            stability = 1 - (std_imp / mean_imp) if mean_imp > 0 else 0
            logger.info(f"    {feature_name}: {mean_imp:.4f} ± {std_imp:.4f} (stability: {stability:.3f})")
    
    return {
        'loo_cv_r2': loo_scores.mean(),
        'loo_cv_std': loo_scores.std(),
        'permutation_pvalue': perm_pvalue,
        'overfitting_gap': overfitting_gap,
        'final_train_r2': train_mean[-1],
        'final_val_r2': val_mean[-1]
    }


def validate_sample_size_adequacy():
    """Validate if sample size is adequate for the number of features."""
    logger.info("=== SAMPLE SIZE ADEQUACY VALIDATION ===")
    
    with open('results/publication_metrics.json', 'r') as f:
        metrics = json.load(f)
    
    n_samples = metrics['data_info']['n_samples']
    n_features = metrics['data_info']['n_selected_features']
    
    logger.info(f"Samples: {n_samples}, Features: {n_features}")
    
    # Rule of thumb: 10-15 samples per feature for regression
    min_samples_conservative = n_features * 15
    min_samples_liberal = n_features * 10
    
    logger.info(f"Recommended samples (conservative): {min_samples_conservative}")
    logger.info(f"Recommended samples (liberal): {min_samples_liberal}")
    
    if n_samples >= min_samples_conservative:
        logger.info("  ✅ Excellent sample size")
    elif n_samples >= min_samples_liberal:
        logger.info("  ✅ Adequate sample size")
    elif n_samples >= n_features * 5:
        logger.warning("  ⚠️  Marginal sample size - results should be interpreted cautiously")
    else:
        logger.warning("  ⚠️  Insufficient sample size - high risk of overfitting")
    
    # Calculate effective degrees of freedom
    effective_dof = n_samples - n_features - 1
    logger.info(f"Effective degrees of freedom: {effective_dof}")
    
    if effective_dof < 10:
        logger.warning("  ⚠️  Very low degrees of freedom - model may be overparameterized")
    elif effective_dof < 20:
        logger.warning("  ⚠️  Low degrees of freedom - interpret results cautiously")
    else:
        logger.info("  ✅ Adequate degrees of freedom")
    
    return {
        'n_samples': n_samples,
        'n_features': n_features,
        'samples_per_feature': n_samples / n_features,
        'effective_dof': effective_dof,
        'adequacy': 'excellent' if n_samples >= min_samples_conservative else 
                   'adequate' if n_samples >= min_samples_liberal else 
                   'marginal' if n_samples >= n_features * 5 else 'insufficient'
    }


def validate_chemical_space_coverage():
    """Validate chemical space coverage and diversity."""
    logger.info("=== CHEMICAL SPACE COVERAGE VALIDATION ===")
    
    # Load SMILES data
    from publication_ready_pipeline import ComprehensiveFeatureExtractor, load_binding_data, integrate_features_and_binding
    
    extractor = ComprehensiveFeatureExtractor()
    features_df = extractor.extract_features_from_files("data/raw/pdbqt")
    binding_df = load_binding_data("data/processed/processed_logs.csv")
    integrated_df = integrate_features_and_binding(features_df, binding_df)
    
    # Basic chemical space metrics
    mw_range = integrated_df['molecular_weight'].max() - integrated_df['molecular_weight'].min()
    logp_range = integrated_df['logp'].max() - integrated_df['logp'].min()
    tpsa_range = integrated_df['tpsa'].max() - integrated_df['tpsa'].min()
    
    logger.info(f"Molecular weight range: {mw_range:.1f} Da")
    logger.info(f"LogP range: {logp_range:.2f}")
    logger.info(f"TPSA range: {tpsa_range:.1f} Ų")
    
    # Check for drug-like properties
    drug_like = integrated_df[
        (integrated_df['molecular_weight'] <= 500) &
        (integrated_df['logp'] <= 5) &
        (integrated_df['hbd'] <= 5) &
        (integrated_df['hba'] <= 10)
    ]
    
    drug_like_fraction = len(drug_like) / len(integrated_df)
    logger.info(f"Drug-like compounds (Lipinski): {len(drug_like)}/{len(integrated_df)} ({drug_like_fraction*100:.1f}%)")
    
    # Diversity assessment using molecular descriptors
    descriptor_cols = ['molecular_weight', 'logp', 'tpsa', 'rotatable_bonds', 'aromatic_rings']
    descriptor_data = integrated_df[descriptor_cols].values
    
    # Calculate pairwise distances
    from sklearn.metrics.pairwise import euclidean_distances
    from sklearn.preprocessing import StandardScaler
    
    scaler = StandardScaler()
    descriptor_data_scaled = scaler.fit_transform(descriptor_data)
    distances = euclidean_distances(descriptor_data_scaled)
    
    # Remove diagonal (self-distances)
    distances_flat = distances[np.triu_indices_from(distances, k=1)]
    mean_distance = distances_flat.mean()
    min_distance = distances_flat.min()
    
    logger.info(f"Mean pairwise distance: {mean_distance:.3f}")
    logger.info(f"Minimum pairwise distance: {min_distance:.3f}")
    
    if min_distance < 0.5:
        logger.warning("  ⚠️  Some compounds are very similar - potential redundancy")
    else:
        logger.info("  ✅ Good chemical diversity")
    
    return {
        'mw_range': mw_range,
        'logp_range': logp_range,
        'tpsa_range': tpsa_range,
        'drug_like_fraction': drug_like_fraction,
        'mean_distance': mean_distance,
        'min_distance': min_distance
    }


def generate_academic_report():
    """Generate comprehensive academic validation report."""
    logger.info("=== GENERATING ACADEMIC VALIDATION REPORT ===")
    
    # Load model and metrics
    model, scaler, selected_features, metrics = load_publication_results()
    
    if model is None:
        logger.error("Could not load publication results")
        return
    
    # Run all validations
    data_validation = validate_data_integrity()
    performance_validation = validate_model_performance(model, scaler, selected_features, metrics)
    sample_size_validation = validate_sample_size_adequacy()
    chemical_space_validation = validate_chemical_space_coverage()
    
    # Compile report
    report = {
        'timestamp': pd.Timestamp.now().isoformat(),
        'data_validation': data_validation,
        'performance_validation': performance_validation,
        'sample_size_validation': sample_size_validation,
        'chemical_space_validation': chemical_space_validation,
        'original_metrics': metrics['metrics'],
        'recommendations': []
    }
    
    # Generate recommendations
    logger.info("=== ACADEMIC RECOMMENDATIONS ===")
    
    # Sample size recommendations
    if sample_size_validation['adequacy'] in ['marginal', 'insufficient']:
        recommendation = "Consider collecting additional data to improve model reliability"
        report['recommendations'].append(recommendation)
        logger.warning(f"  📝 {recommendation}")
    
    # Performance recommendations
    if performance_validation['permutation_pvalue'] > 0.05:
        recommendation = "Model performance may not be statistically significant - interpret results cautiously"
        report['recommendations'].append(recommendation)
        logger.warning(f"  📝 {recommendation}")
    
    if performance_validation['overfitting_gap'] > 0.15:
        recommendation = "Consider regularization or feature reduction to minimize overfitting"
        report['recommendations'].append(recommendation)
        logger.warning(f"  📝 {recommendation}")
    
    # Chemical space recommendations
    if chemical_space_validation['min_distance'] < 0.5:
        recommendation = "Consider removing highly similar compounds to improve dataset diversity"
        report['recommendations'].append(recommendation)
        logger.warning(f"  📝 {recommendation}")
    
    if chemical_space_validation['drug_like_fraction'] < 0.5:
        recommendation = "Dataset contains many non-drug-like compounds - consider applicability domain"
        report['recommendations'].append(recommendation)
        logger.warning(f"  📝 {recommendation}")
    
    # Overall assessment
    logger.info("=== OVERALL ACADEMIC ASSESSMENT ===")
    
    issues = len(report['recommendations'])
    if issues == 0:
        logger.info("  ✅ Model passes all academic validation criteria")
        assessment = "EXCELLENT"
    elif issues <= 2:
        logger.info("  ✅ Model is academically sound with minor considerations")
        assessment = "GOOD"
    elif issues <= 4:
        logger.warning("  ⚠️  Model has some academic concerns that should be addressed")
        assessment = "ACCEPTABLE"
    else:
        logger.warning("  ❌ Model has significant academic issues")
        assessment = "NEEDS_IMPROVEMENT"
    
    report['overall_assessment'] = assessment
    
    # Save report
    with open('results/academic_validation_report.json', 'w') as f:
        json.dump(report, f, indent=2)
    
    logger.info("Academic validation report saved to results/academic_validation_report.json")
    
    return report


def main():
    """Main academic validation execution."""
    logger.info("Starting Academic Validation of TLR4 Binding Prediction Pipeline")
    
    try:
        report = generate_academic_report()
        
        if report:
            logger.info(f"\n🎓 ACADEMIC VALIDATION COMPLETE")
            logger.info(f"Overall Assessment: {report['overall_assessment']}")
            
            if report['recommendations']:
                logger.info(f"Recommendations: {len(report['recommendations'])}")
                for i, rec in enumerate(report['recommendations'], 1):
                    logger.info(f"  {i}. {rec}")
            else:
                logger.info("No recommendations - model is academically robust!")
        
        return report
        
    except Exception as e:
        logger.error(f"Academic validation failed: {e}")
        import traceback
        traceback.print_exc()
        raise


if __name__ == "__main__":
    report = main()
```
