## Supplementary material for "Machine Learning-Driven Prediction of TLR4 Binding Affinity: A Comprehensive Molecular Feature Analysis for Drug Discovery": Codebase: analyze_data.py

```
#!/usr/bin/env python3
"""
Analyze the processed logs data structure.
"""

import pandas as pd

def analyze_data():
    df = pd.read_csv('data/processed/processed_logs.csv')
    
    print(f"Total records: {len(df)}")
    print(f"Unique compounds: {df['ligand'].nunique()}")
    
    # Count base compounds (without _conf_ suffix)
    base_compounds = []
    for compound in df['ligand'].unique():
        if '_conf_' in compound:
            base = compound.split('_conf_')[0]
        else:
            base = compound
        base_compounds.append(base)
    
    unique_bases = set(base_compounds)
    print(f"Unique base compounds: {len(unique_bases)}")
    
    print("\nSample compounds:")
    for comp in sorted(df['ligand'].unique())[:10]:
        print(f"  {comp}")
    
    print(f"\nSample base compounds:")
    for comp in sorted(unique_bases)[:10]:
        print(f"  {comp}")
    
    # Check affinity range
    print(f"\nAffinity statistics:")
    print(f"  Range: {df['affinity'].min():.3f} to {df['affinity'].max():.3f}")
    print(f"  Mean: {df['affinity'].mean():.3f}")
    print(f"  Std: {df['affinity'].std():.3f}")

if __name__ == "__main__":
    analyze_data()
```
