## Supplementary material for "Machine Learning-Driven Prediction of TLR4 Binding Affinity: A Comprehensive Molecular Feature Analysis for Drug Discovery": Codebase: demo_ablation_study.py

```
#!/usr/bin/env python3
"""
Demo script for TLR4 Binding Prediction Ablation Study Framework

This script demonstrates the comprehensive ablation study framework including:
- Feature ablation studies
- Data size ablation studies  
- Hyperparameter sensitivity analysis
- Statistical significance testing
- Comprehensive reporting and visualization

Usage:
    python demo_ablation_study.py
"""

import sys
import os
import numpy as np
import pandas as pd
import matplotlib.pyplot as plt
from pathlib import Path
import warnings
warnings.filterwarnings('ignore')

# Add src to path
sys.path.append(str(Path(__file__).parent.parent / "src"))

from tlr4_binding.ml_components.ablation_study import (
    AblationStudyFramework, AblationConfig, AblationResult
)
from tlr4_binding.ml_components.trainer import MLModelTrainer
from tlr4_binding.ml_components.feature_engineering import FeatureEngineeringPipeline
from tlr4_binding.data_processing.data_loader import BindingDataLoader
from tlr4_binding.molecular_analysis.feature_extractor import MolecularFeatureExtractor

def create_synthetic_data(n_samples=500, n_features=50):
    """Create synthetic dataset for ablation study demonstration"""
    print("Creating synthetic molecular binding dataset...")
    
    # Generate synthetic features (molecular descriptors)
    np.random.seed(42)
    
    # Create feature groups for more realistic ablation study
    feature_groups = {
        "molecular_weight": ["mol_weight", "mol_weight_log"],
        "lipophilicity": ["logp", "logp_squared", "logp_cube"],
        "polar_surface_area": ["tpsa", "tpsa_log", "tpsa_sqrt"],
        "hydrogen_bonds": ["hbd", "hba", "hbd_hba_ratio"],
        "rotatable_bonds": ["rot_bonds", "rot_bonds_norm"],
        "charge_properties": ["formal_charge", "partial_charge_mean", "partial_charge_std"],
        "structural_features": ["aromatic_atoms", "ring_count", "stereo_centers"],
        "3d_properties": ["radius_gyration", "molecular_volume", "surface_area", "asphericity"],
        "conformational": ["energy_min", "energy_max", "energy_std"],
        "additional_descriptors": [f"descriptor_{i}" for i in range(30)]
    }
    
    # Generate features with realistic correlations
    X_data = {}
    for group_name, features in feature_groups.items():
        # Create correlated features within groups
        base_feature = np.random.normal(0, 1, n_samples)
        for i, feature in enumerate(features):
            # Add some correlation within groups
            noise = np.random.normal(0, 0.3, n_samples)
            X_data[feature] = base_feature + i * 0.1 + noise
    
    X = pd.DataFrame(X_data)
    
    # Create target with realistic relationships to some features
    # Strongest predictors (molecular weight, lipophilicity, 3D properties)
    strong_predictors = X["mol_weight"] * 0.3 + X["logp"] * 0.25 + X["radius_gyration"] * 0.2
    
    # Medium predictors (polar surface area, hydrogen bonds)
    medium_predictors = X["tpsa"] * 0.15 + X["hbd"] * 0.1
    
    # Weak predictors (remaining features)
    weak_predictors = X.iloc[:, 10:].sum(axis=1) * 0.01
    
    # Add noise
    noise = np.random.normal(0, 0.5, n_samples)
    
    # Create binding affinity (lower = stronger binding)
    y = -(strong_predictors + medium_predictors + weak_predictors + noise)
    
    print(f"Generated dataset: {X.shape[0]} samples, {X.shape[1]} features")
    print(f"Target range: {y.min():.2f} to {y.max():.2f}")
    
    return X, y, feature_groups

def run_feature_ablation_demo():
    """Demonstrate feature ablation study"""
    print("\n" + "="*60)
    print("FEATURE ABLATION STUDY DEMONSTRATION")
    print("="*60)
    
    # Create synthetic data
    X, y, feature_groups = create_synthetic_data(n_samples=300, n_features=50)
    
    # Initialize ablation framework
    config = AblationConfig(
        cv_folds=3,  # Reduced for demo
        n_iterations=5,
        random_state=42
    )
    
    ablation_framework = AblationStudyFramework(config)
    
    # Initialize model trainer (mock for demo)
    model_trainer = MLModelTrainer()
    
    print("\nRunning feature ablation study...")
    
    # Run feature ablation by groups
    feature_results = ablation_framework.feature_study.run_feature_ablation(
        X, y, model_trainer, feature_groups
    )
    
    print(f"\nFeature ablation results ({len(feature_results)} experiments):")
    print("-" * 80)
    print(f"{'Feature Group':<25} {'Impact':<10} {'P-value':<10} {'Significant':<12}")
    print("-" * 80)
    
    for result in sorted(feature_results, key=lambda x: abs(x.score_difference), reverse=True):
        significant = "Yes" if result.p_value and result.p_value < 0.05 else "No"
        print(f"{result.component_name:<25} {result.score_difference:<10.4f} "
              f"{result.p_value:<10.4f} {significant:<12}")
    
    # Run sequential feature selection
    print("\nRunning sequential feature selection...")
    sequential_results = ablation_framework.feature_study.run_sequential_feature_ablation(
        X, y, model_trainer, method="forward"
    )
    
    print(f"\nForward selection results (top 5 features):")
    for i, result in enumerate(sequential_results[:5]):
        features = result.metadata["selected_features"]
        print(f"Step {i+1}: {result.baseline_score:.4f} (features: {len(features)})")
    
    return {"feature_groups": feature_results, "sequential": sequential_results}

def run_data_size_ablation_demo():
    """Demonstrate data size ablation study"""
    print("\n" + "="*60)
    print("DATA SIZE ABLATION STUDY DEMONSTRATION")
    print("="*60)
    
    # Create larger dataset for data size study
    X, y, _ = create_synthetic_data(n_samples=400, n_features=30)
    
    config = AblationConfig(cv_folds=3, random_state=42)
    ablation_framework = AblationStudyFramework(config)
    model_trainer = MLModelTrainer()
    
    print("\nRunning data size ablation study...")
    
    data_size_results = ablation_framework.data_size_study.run_data_size_ablation(
        X, y, model_trainer
    )
    
    print(f"\nData size ablation results:")
    print("-" * 60)
    print(f"{'Sample Size':<12} {'R² Score':<10} {'Performance':<15}")
    print("-" * 60)
    
    for result in data_size_results:
        sample_size = result.metadata["sample_size"]
        score = result.ablated_score
        performance = "Poor" if score < 0.5 else "Good" if score > 0.7 else "Fair"
        print(f"{sample_size:<12} {score:<10.4f} {performance:<15}")
    
    return data_size_results

def run_hyperparameter_ablation_demo():
    """Demonstrate hyperparameter ablation study"""
    print("\n" + "="*60)
    print("HYPERPARAMETER ABLATION STUDY DEMONSTRATION")
    print("="*60)
    
    # Create dataset for hyperparameter study
    X, y, _ = create_synthetic_data(n_samples=200, n_features=25)
    
    config = AblationConfig(cv_folds=3, random_state=42)
    ablation_framework = AblationStudyFramework(config)
    model_trainer = MLModelTrainer()
    
    # Define hyperparameter grids for demo
    param_grids = {
        "n_estimators": [50, 100, 200],
        "max_depth": [3, 5, 10],
        "min_samples_split": [2, 5, 10],
        "learning_rate": [0.01, 0.1, 0.3]
    }
    
    print("\nRunning hyperparameter ablation study...")
    
    hyperparameter_results = ablation_framework.hyperparameter_study.run_hyperparameter_ablation(
        X, y, model_trainer, param_grids
    )
    
    # Group results by parameter
    param_groups = {}
    for result in hyperparameter_results:
        param_name = result.metadata["parameter"]
        if param_name not in param_groups:
            param_groups[param_name] = []
        param_groups[param_name].append(result)
    
    print(f"\nHyperparameter sensitivity analysis:")
    print("-" * 70)
    print(f"{'Parameter':<20} {'Best Value':<12} {'Score Range':<15} {'Sensitivity':<12}")
    print("-" * 70)
    
    for param_name, param_results in param_groups.items():
        scores = [r.ablated_score for r in param_results]
        best_result = max(param_results, key=lambda x: x.ablated_score)
        best_value = best_result.metadata["value"]
        score_range = max(scores) - min(scores)
        sensitivity = "High" if score_range > 0.1 else "Medium" if score_range > 0.05 else "Low"
        
        print(f"{param_name:<20} {best_value:<12} {score_range:<15.4f} {sensitivity:<12}")
    
    return hyperparameter_results

def run_comprehensive_ablation_demo():
    """Run comprehensive ablation study with all components"""
    print("\n" + "="*60)
    print("COMPREHENSIVE ABLATION STUDY DEMONSTRATION")
    print("="*60)
    
    # Create dataset
    X, y, feature_groups = create_synthetic_data(n_samples=300, n_features=40)
    
    # Configure ablation study
    config = AblationConfig(
        cv_folds=3,
        n_iterations=5,
        random_state=42,
        min_effect_size=0.01
    )
    
    ablation_framework = AblationStudyFramework(config)
    model_trainer = MLModelTrainer()
    
    print("\nRunning comprehensive ablation study...")
    
    # Run all ablation studies
    all_results = ablation_framework.run_comprehensive_ablation(
        X, y, model_trainer,
        study_types=["feature", "data_size", "hyperparameter"]
    )
    
    print(f"\nComprehensive ablation study completed!")
    print(f"Total experiments run: {sum(len(r) for r in all_results.values())}")
    
    # Generate comprehensive report
    print("\nGenerating ablation study report...")
    report = ablation_framework.generate_ablation_report(
        all_results, 
        save_path="results/ablation_study"
    )
    
    # Print summary
    print("\n" + "="*60)
    print("ABLATION STUDY SUMMARY")
    print("="*60)
    
    summary = report["summary"]
    print(f"Total studies: {summary['total_studies']}")
    print(f"Total experiments: {summary['total_experiments']}")
    print(f"Significant effects: {summary['significant_effects']}")
    
    if summary["largest_effect"]:
        largest = summary["largest_effect"]
        print(f"Largest effect: {largest.component_name} (ΔR² = {largest.score_difference:.4f})")
    
    if summary["most_sensitive_component"]:
        sensitive = summary["most_sensitive_component"]
        print(f"Most sensitive: {sensitive.component_name} ({sensitive.relative_change:.2%} change)")
    
    # Print recommendations
    print(f"\nRecommendations:")
    for i, rec in enumerate(report["recommendations"], 1):
        print(f"{i}. {rec}")
    
    # Create visualizations
    print(f"\nCreating ablation study visualizations...")
    ablation_framework.plot_ablation_results(
        all_results,
        save_path="results/ablation_study/ablation_plots.png"
    )
    
    return all_results, report

def demonstrate_statistical_analysis():
    """Demonstrate statistical analysis capabilities"""
    print("\n" + "="*60)
    print("STATISTICAL ANALYSIS DEMONSTRATION")
    print("="*60)
    
    # Create some mock ablation results for statistical demonstration
    from tlr4_binding.ml_components.ablation_study import AblationResult
    
    # Simulate feature ablation results with different effect sizes
    results = [
        AblationResult("mol_weight", 0.75, 0.60, 0.15, 0.20, 0.001),
        AblationResult("logp", 0.75, 0.68, 0.07, 0.09, 0.02),
        AblationResult("tpsa", 0.75, 0.72, 0.03, 0.04, 0.15),
        AblationResult("hbd", 0.75, 0.73, 0.02, 0.03, 0.25),
        AblationResult("rot_bonds", 0.75, 0.74, 0.01, 0.01, 0.45),
    ]
    
    print("\nStatistical significance analysis:")
    print("-" * 50)
    print(f"{'Feature':<15} {'Effect Size':<12} {'P-value':<10} {'Significant':<12}")
    print("-" * 50)
    
    significant_count = 0
    for result in results:
        significant = result.p_value < 0.05 if result.p_value else False
        if significant:
            significant_count += 1
        
        sig_text = "Yes" if significant else "No"
        print(f"{result.component_name:<15} {result.score_difference:<12.4f} "
              f"{result.p_value:<10.4f} {sig_text:<12}")
    
    print(f"\nSummary: {significant_count}/{len(results)} features show statistically significant effects")
    
    # Effect size interpretation
    print(f"\nEffect size interpretation:")
    for result in results:
        effect_size = abs(result.score_difference)
        if effect_size > 0.1:
            interpretation = "Large"
        elif effect_size > 0.05:
            interpretation = "Medium"
        else:
            interpretation = "Small"
        print(f"{result.component_name}: {interpretation} effect (ΔR² = {effect_size:.4f})")

def main():
    """Main demonstration function"""
    print("TLR4 Binding Prediction - Ablation Study Framework Demo")
    print("=" * 70)
    
    # Create results directory
    Path("results/ablation_study").mkdir(parents=True, exist_ok=True)
    
    try:
        # Run individual ablation studies
        feature_results = run_feature_ablation_demo()
        data_size_results = run_data_size_ablation_demo()
        hyperparameter_results = run_hyperparameter_ablation_demo()
        
        # Run comprehensive study
        comprehensive_results, report = run_comprehensive_ablation_demo()
        
        # Demonstrate statistical analysis
        demonstrate_statistical_analysis()
        
        print("\n" + "="*70)
        print("ABLATION STUDY DEMONSTRATION COMPLETED SUCCESSFULLY!")
        print("="*70)
        
        print(f"\nResults saved to: results/ablation_study/")
        print(f"- ablation_report.json: Comprehensive analysis report")
        print(f"- ablation_results.csv: Detailed results table")
        print(f"- ablation_plots.png: Visualization plots")
        
        print(f"\nKey findings:")
        print(f"- Feature ablation identified critical molecular descriptors")
        print(f"- Data size analysis shows learning curve characteristics")
        print(f"- Hyperparameter sensitivity varies across parameters")
        print(f"- Statistical significance testing validates findings")
        
        return True
        
    except Exception as e:
        print(f"\nError during ablation study demonstration: {e}")
        import traceback
        traceback.print_exc()
        return False

if __name__ == "__main__":
    success = main()
    sys.exit(0 if success else 1)
```
