## Supplementary material for "Machine Learning-Driven Prediction of TLR4 Binding Affinity: A Comprehensive Molecular Feature Analysis for Drug Discovery": Codebase: demo_binding_processing.py

```
#!/usr/bin/env python3
"""
Demonstration script for TLR4 binding data processing.

This script demonstrates the binding data processing functionality
implemented in Task 6, including data loading, validation, outlier
detection, and best affinity extraction.
"""

import sys
import os
import pandas as pd
import numpy as np
from pathlib import Path

# Add src to path for imports
sys.path.append('src')

from tlr4_binding.data_processing.preprocessor import (
    BindingDataLoader,
    DataPreprocessor,
    CompoundMatcher,
    DataIntegrator
)


def demonstrate_binding_data_processing():
    """Demonstrate the complete binding data processing pipeline."""
    
    print("=" * 80)
    print("TLR4 BINDING DATA PROCESSING DEMONSTRATION")
    print("=" * 80)
    
    # Initialize components
    binding_loader = BindingDataLoader()
    preprocessor = DataPreprocessor()
    
    # Path to binding data
    binding_data_path = "binding-data/processed/processed_logs.csv"
    
    print(f"\n1. LOADING BINDING DATA FROM: {binding_data_path}")
    print("-" * 50)
    
    try:
        # Load binding data
        binding_df = binding_loader.load_csv(binding_data_path)
        print(f"✓ Successfully loaded {len(binding_df):,} binding records")
        print(f"  - Unique ligands: {binding_df['ligand'].nunique():,}")
        print(f"  - Affinity range: {binding_df['affinity'].min():.3f} to {binding_df['affinity'].max():.3f} kcal/mol")
        print(f"  - Mean affinity: {binding_df['affinity'].mean():.3f} kcal/mol")
        
    except Exception as e:
        print(f"✗ Error loading binding data: {e}")
        return
    
    print(f"\n2. DATA VALIDATION AND QUALITY ASSESSMENT")
    print("-" * 50)
    
    # Validate data quality
    validation_issues = binding_loader.validate_binding_data(binding_df)
    
    print("Validation Results:")
    for category, issues in validation_issues.items():
        if issues:
            print(f"  - {category.replace('_', ' ').title()}: {len(issues)} issues")
            for issue in issues[:3]:  # Show first 3 issues
                print(f"    • {issue}")
            if len(issues) > 3:
                print(f"    • ... and {len(issues) - 3} more")
        else:
            print(f"  - {category.replace('_', ' ').title()}: No issues ✓")
    
    print(f"\n3. OUTLIER DETECTION AND CLEANING")
    print("-" * 50)
    
    # Demonstrate outlier detection
    outlier_info = binding_loader._detect_affinity_outliers(binding_df)
    
    print("Outlier Detection Results:")
    if outlier_info['outliers']:
        for outlier in outlier_info['outliers']:
            print(f"  • {outlier}")
    else:
        print("  • No outliers detected")
    
    if outlier_info['statistical_anomalies']:
        print("\nStatistical Anomalies:")
        for anomaly in outlier_info['statistical_anomalies']:
            print(f"  • {anomaly}")
    
    # Demonstrate data cleaning (IQR method)
    print(f"\n4. DATA CLEANING (IQR METHOD)")
    print("-" * 50)
    
    cleaned_df = binding_loader.clean_binding_data(
        binding_df, 
        outlier_method='iqr', 
        remove_outliers=False
    )
    
    print(f"Original data: {len(binding_df):,} records")
    print(f"Cleaned data: {len(cleaned_df):,} records")
    print(f"Affinity range after cleaning: {cleaned_df['affinity'].min():.3f} to {cleaned_df['affinity'].max():.3f} kcal/mol")
    
    print(f"\n5. BEST BINDING AFFINITY EXTRACTION")
    print("-" * 50)
    
    # Extract best binding affinities
    best_affinities = preprocessor.get_best_affinities(cleaned_df)
    
    print(f"✓ Extracted {len(best_affinities):,} best binding modes")
    print(f"  - Strongest binding: {best_affinities['affinity'].min():.3f} kcal/mol")
    print(f"  - Weakest binding: {best_affinities['affinity'].max():.3f} kcal/mol")
    print(f"  - Mean binding: {best_affinities['affinity'].mean():.3f} kcal/mol")
    
    # Show top 10 strongest binders
    top_binders = best_affinities.nsmallest(10, 'affinity')
    print(f"\nTop 10 Strongest TLR4 Binders:")
    print("  Rank | Ligand                    | Affinity (kcal/mol)")
    print("  " + "-" * 55)
    for i, (_, row) in enumerate(top_binders.iterrows(), 1):
        ligand_name = row['ligand'][:20] + "..." if len(row['ligand']) > 20 else row['ligand']
        print(f"  {i:4d} | {ligand_name:<25} | {row['affinity']:8.3f}")
    
    print(f"\n6. COMPOUND NAME MATCHING DEMONSTRATION")
    print("-" * 50)
    
    # Demonstrate compound matching
    matcher = CompoundMatcher(threshold=80.0)
    
    # Sample compound names from features (simulated)
    feature_compounds = [
        "Andrographolide",
        "Curcumin", 
        "Resveratrol",
        "Quercetin",
        "Epigallocatechin"
    ]
    
    # Sample compound names from binding data
    binding_compounds = binding_df['ligand'].unique()[:10].tolist()
    
    print(f"Feature compounds: {feature_compounds}")
    print(f"Binding compounds (sample): {binding_compounds[:5]}...")
    
    matches = matcher.match_compounds(feature_compounds, binding_compounds)
    print(f"\nMatching Results:")
    for feature_compound, binding_compound in matches.items():
        confidence = matcher.get_match_confidence(feature_compound, binding_compound)
        print(f"  '{feature_compound}' → '{binding_compound}' (confidence: {confidence:.1f}%)")
    
    print(f"\n7. DATASET INTEGRATION DEMONSTRATION")
    print("-" * 50)
    
    # Create sample molecular features data
    sample_features = pd.DataFrame({
        'compound_name': ['Andrographolide', 'Curcumin', 'Resveratrol'],
        'molecular_weight': [350.0, 368.0, 228.0],
        'logp': [2.5, 3.0, 3.1],
        'tpsa': [80.0, 90.0, 60.0],
        'rotatable_bonds': [5, 7, 3],
        'hbd': [3, 2, 3],
        'hba': [6, 6, 4]
    })
    
    print("Sample molecular features:")
    print(sample_features.to_string(index=False))
    
    # Integrate datasets
    integrator = DataIntegrator()
    integrated_df = integrator.integrate_datasets(sample_features, best_affinities)
    
    print(f"\nIntegrated dataset: {len(integrated_df)} records")
    print("Integration statistics:")
    stats = integrator.get_integration_stats()
    for key, value in stats.items():
        print(f"  - {key.replace('_', ' ').title()}: {value}")
    
    if len(integrated_df) > 0:
        print(f"\nSample integrated record:")
        print(integrated_df[['compound_name', 'molecular_weight', 'affinity']].head(3).to_string(index=False))
    
    print(f"\n8. COMPLETE PREPROCESSING PIPELINE")
    print("-" * 50)
    
    # Demonstrate complete pipeline
    try:
        final_df = preprocessor.preprocess_pipeline(sample_features, binding_data_path)
        print(f"✓ Complete pipeline successful: {len(final_df)} integrated records")
        print(f"  - Features: {len([col for col in final_df.columns if col not in ['ligand', 'mode', 'affinity', 'dist_from_rmsd_lb', 'best_mode_rmsd_ub', 'matched_compound']])}")
        print(f"  - Target range: {final_df['affinity'].min():.3f} to {final_df['affinity'].max():.3f} kcal/mol")
    except Exception as e:
        print(f"✗ Pipeline error: {e}")
    
    print(f"\n" + "=" * 80)
    print("DEMONSTRATION COMPLETED SUCCESSFULLY")
    print("=" * 80)


if __name__ == "__main__":
    demonstrate_binding_data_processing()
```
