## Supplementary material for "Machine Learning-Driven Prediction of TLR4 Binding Affinity: A Comprehensive Molecular Feature Analysis for Drug Discovery": Codebase: demo_comprehensive_evaluation.py

```
"""
Demo script for comprehensive model evaluation framework.

This script demonstrates the comprehensive evaluation capabilities including:
- Learning curves and validation curves
- Cross-validation evaluation
- Statistical significance testing
- Performance comparison plots
- Comprehensive evaluation reports
"""

import sys
import os
import numpy as np
import pandas as pd
import matplotlib.pyplot as plt
from pathlib import Path

# Add src to path
sys.path.append(str(Path(__file__).parent.parent / "src"))

from tlr4_binding.ml_components.comprehensive_evaluator import ComprehensiveEvaluator
from tlr4_binding.ml_components.evaluator import ModelEvaluator, PerformanceMetrics
from tlr4_binding.ml_components.trainer import MLModelTrainer
from tlr4_binding.ml_components.deep_learning_trainer import DeepLearningTrainer
from tlr4_binding.ml_components.ensemble_models import EnsembleTrainer
from sklearn.model_selection import train_test_split
from sklearn.datasets import make_regression
from sklearn.ensemble import RandomForestRegressor
from sklearn.linear_model import LinearRegression
from sklearn.svm import SVR
from sklearn.metrics import mean_squared_error, r2_score
import warnings
warnings.filterwarnings('ignore')

def create_synthetic_data(n_samples=1000, n_features=50, noise=0.1):
    """Create synthetic regression data for demonstration."""
    print("Creating synthetic molecular binding data...")
    
    # Create synthetic molecular features and binding affinities
    X, y = make_regression(
        n_samples=n_samples,
        n_features=n_features,
        noise=noise,
        random_state=42
    )
    
    # Make binding affinities more realistic (negative values for stronger binding)
    y = y * -0.5 - 5  # Scale to typical binding affinity range
    
    # Create feature names
    feature_names = [f"mol_feature_{i:02d}" for i in range(n_features)]
    
    return X, y, feature_names

def demonstrate_individual_evaluation():
    """Demonstrate individual model evaluation capabilities."""
    print("\n" + "="*60)
    print("DEMONSTRATION 1: Individual Model Evaluation")
    print("="*60)
    
    # Create synthetic data
    X, y, feature_names = create_synthetic_data(n_samples=500, n_features=20)
    X_train, X_test, y_train, y_test = train_test_split(
        X, y, test_size=0.2, random_state=42
    )
    
    # Initialize evaluator
    evaluator = ModelEvaluator()
    
    # Train a simple model for demonstration
    model = RandomForestRegressor(n_estimators=50, random_state=42)
    model.fit(X_train, y_train)
    
    # Make predictions
    y_pred = model.predict(X_test)
    
    # Evaluate model
    print("Evaluating Random Forest model...")
    metrics = evaluator.evaluate(y_test, y_pred, "Random Forest Demo")
    
    # Print key metrics
    print(f"\nModel Performance Summary:")
    print(f"R² Score: {metrics.metrics['r2']:.4f}")
    print(f"RMSE: {metrics.metrics['rmse']:.4f}")
    print(f"MAE: {metrics.metrics['mae']:.4f}")
    print(f"Pearson Correlation: {metrics.metrics['pearson_r']:.4f}")
    print(f"Spearman Correlation: {metrics.metrics['spearman_r']:.4f}")
    
    # Generate residual plots
    print("\nGenerating residual analysis plots...")
    evaluator.plot_residuals(metrics, save_path="results/residual_analysis_demo.png")
    
    return metrics

def demonstrate_cross_validation():
    """Demonstrate cross-validation evaluation."""
    print("\n" + "="*60)
    print("DEMONSTRATION 2: Cross-Validation Evaluation")
    print("="*60)
    
    # Create synthetic data
    X, y, feature_names = create_synthetic_data(n_samples=800, n_features=25)
    
    # Initialize evaluator
    evaluator = ModelEvaluator()
    
    # Create multiple models for comparison
    models = {
        'Random Forest': RandomForestRegressor(n_estimators=100, random_state=42),
        'Linear Regression': LinearRegression(),
        'SVR': SVR(kernel='rbf', C=1.0, gamma='scale')
    }
    
    # Perform cross-validation for each model
    cv_results = {}
    for name, model in models.items():
        print(f"Performing 5-fold cross-validation for {name}...")
        cv_result = evaluator.cross_validate_model(model, X, y, name, cv=5)
        cv_results[name] = cv_result
        
        print(f"  Mean R²: {cv_result['r2']['mean']:.4f} ± {cv_result['r2']['std']:.4f}")
        print(f"  Mean RMSE: {cv_result['rmse']['mean']:.4f} ± {cv_result['rmse']['std']:.4f}")
    
    return cv_results

def demonstrate_learning_curves():
    """Demonstrate learning curves generation."""
    print("\n" + "="*60)
    print("DEMONSTRATION 3: Learning Curves")
    print("="*60)
    
    # Create synthetic data
    X, y, feature_names = create_synthetic_data(n_samples=1000, n_features=30)
    
    # Initialize evaluator
    evaluator = ModelEvaluator()
    
    # Create a model for learning curves
    model = RandomForestRegressor(n_estimators=100, random_state=42)
    
    print("Generating learning curves for Random Forest...")
    learning_curve_data = evaluator.generate_learning_curves(
        model, X, y, "Random Forest Demo", 
        save_path="results/learning_curves_demo.png"
    )
    
    print(f"Learning curve data generated with {len(learning_curve_data['train_sizes'])} training sizes")
    print(f"Final training RMSE: {learning_curve_data['train_mean'][-1]:.4f}")
    print(f"Final validation RMSE: {learning_curve_data['val_mean'][-1]:.4f}")
    
    return learning_curve_data

def demonstrate_validation_curves():
    """Demonstrate validation curves generation."""
    print("\n" + "="*60)
    print("DEMONSTRATION 4: Validation Curves")
    print("="*60)
    
    # Create synthetic data
    X, y, feature_names = create_synthetic_data(n_samples=800, n_features=25)
    
    # Initialize evaluator
    evaluator = ModelEvaluator()
    
    # Create a Random Forest model for validation curves
    model = RandomForestRegressor(random_state=42)
    
    # Test different values of n_estimators
    param_range = [10, 20, 50, 100, 200]
    
    print("Generating validation curves for Random Forest (n_estimators)...")
    validation_curve_data = evaluator.generate_validation_curves(
        model, X, y, 'n_estimators', param_range, "Random Forest Demo",
        save_path="results/validation_curves_demo.png"
    )
    
    print(f"Validation curve data generated for {len(param_range)} parameter values")
    print(f"Best parameter value (lowest validation RMSE): {param_range[np.argmin(validation_curve_data['val_mean'])]}")
    
    return validation_curve_data

def demonstrate_statistical_testing():
    """Demonstrate statistical significance testing."""
    print("\n" + "="*60)
    print("DEMONSTRATION 5: Statistical Significance Testing")
    print("="*60)
    
    # Create synthetic data
    X, y, feature_names = create_synthetic_data(n_samples=600, n_features=20)
    X_train, X_test, y_train, y_test = train_test_split(
        X, y, test_size=0.3, random_state=42
    )
    
    # Initialize evaluator
    evaluator = ModelEvaluator()
    
    # Train two different models
    model1 = RandomForestRegressor(n_estimators=50, random_state=42)
    model2 = LinearRegression()
    
    model1.fit(X_train, y_train)
    model2.fit(X_train, y_train)
    
    # Make predictions
    y_pred1 = model1.predict(X_test)
    y_pred2 = model2.predict(X_test)
    
    # Evaluate both models
    metrics1 = evaluator.evaluate(y_test, y_pred1, "Random Forest")
    metrics2 = evaluator.evaluate(y_test, y_pred2, "Linear Regression")
    
    # Perform statistical significance test
    print("Performing statistical significance test...")
    test_result = evaluator.statistical_significance_test(metrics1, metrics2)
    
    print(f"\nStatistical Test Results:")
    print(f"Paired t-test p-value: {test_result['paired_t_test']['p_value']:.4f}")
    print(f"Significant difference: {test_result['paired_t_test']['significant']}")
    print(f"Model 1 R²: {test_result['model1_r2']:.4f}")
    print(f"Model 2 R²: {test_result['model2_r2']:.4f}")
    print(f"R² difference: {test_result['r2_difference']:.4f}")
    
    return test_result

def demonstrate_comprehensive_evaluation():
    """Demonstrate comprehensive evaluation framework."""
    print("\n" + "="*60)
    print("DEMONSTRATION 6: Comprehensive Evaluation Framework")
    print("="*60)
    
    # Create synthetic data
    X, y, feature_names = create_synthetic_data(n_samples=1200, n_features=40)
    
    # Split data
    X_train, X_temp, y_train, y_temp = train_test_split(
        X, y, test_size=0.4, random_state=42
    )
    X_val, X_test, y_val, y_test = train_test_split(
        X_temp, y_temp, test_size=0.5, random_state=42
    )
    
    # Initialize comprehensive evaluator
    evaluator = ComprehensiveEvaluator(
        output_dir="results/comprehensive_evaluation_demo"
    )
    
    print("Starting comprehensive evaluation of all model types...")
    
    # Define model configurations
    model_configs = {
        'traditional_ml': {
            'random_forest': {'n_estimators': 50, 'max_depth': 10},
            'linear_regression': {},
            'svr': {'C': 1.0, 'gamma': 'scale'}
        }
    }
    
    # Perform comprehensive evaluation
    results = evaluator.evaluate_all_models(
        X_train, y_train, X_val, y_val, X_test, y_test,
        feature_names=feature_names,
        model_configs=model_configs
    )
    
    # Generate evaluation report
    print("\nGenerating comprehensive evaluation report...")
    report = evaluator.generate_evaluation_report(results)
    
    # Save report to file
    with open("results/comprehensive_evaluation_report.txt", "w") as f:
        f.write(report)
    
    print("\nComprehensive evaluation completed!")
    print("Results saved to results/comprehensive_evaluation_demo/")
    print("Report saved to results/comprehensive_evaluation_report.txt")
    
    # Print summary
    summary = results['summary']
    print(f"\nEvaluation Summary:")
    print(f"Total models evaluated: {summary['total_models_evaluated']}")
    print(f"Best model: {summary['best_overall_model']}")
    print(f"Best R² score: {summary['best_r2_score']:.4f}")
    
    return results

def main():
    """Run all demonstrations."""
    print("TLR4 Binding Prediction - Comprehensive Evaluation Framework Demo")
    print("=" * 80)
    
    # Create results directory
    os.makedirs("results", exist_ok=True)
    
    try:
        # Run individual demonstrations
        demonstrate_individual_evaluation()
        demonstrate_cross_validation()
        demonstrate_learning_curves()
        demonstrate_validation_curves()
        demonstrate_statistical_testing()
        
        # Run comprehensive evaluation
        demonstrate_comprehensive_evaluation()
        
        print("\n" + "="*80)
        print("ALL DEMONSTRATIONS COMPLETED SUCCESSFULLY!")
        print("="*80)
        print("\nGenerated files:")
        print("- results/residual_analysis_demo.png")
        print("- results/learning_curves_demo.png")
        print("- results/validation_curves_demo.png")
        print("- results/comprehensive_evaluation_demo/ (directory)")
        print("- results/comprehensive_evaluation_report.txt")
        
    except Exception as e:
        print(f"\nError during demonstration: {e}")
        import traceback
        traceback.print_exc()

if __name__ == "__main__":
    main()
```
