## Supplementary material for "Machine Learning-Driven Prediction of TLR4 Binding Affinity: A Comprehensive Molecular Feature Analysis for Drug Discovery": Codebase: demo_data_splitting.py

```
#!/usr/bin/env python3
"""
Demo script for Data Splitting and Validation Framework

This script demonstrates the comprehensive data splitting, cross-validation,
and quality reporting capabilities of the TLR4 binding prediction system.
"""

import numpy as np
import pandas as pd
import sys
import os

# Add src to path
sys.path.insert(0, os.path.join(os.path.dirname(__file__), 'src'))

from tlr4_binding.ml_components.data_splitting import (
    DataValidationFramework,
    DataSplitConfig,
    CrossValidationConfig
)


def create_sample_dataset(n_samples=1000):
    """Create a realistic sample dataset for TLR4 binding prediction."""
    print(f"Creating sample dataset with {n_samples} compounds...")
    
    np.random.seed(42)
    
    # Create features similar to molecular descriptors
    X = pd.DataFrame({
        'molecular_weight': np.random.normal(300, 100, n_samples),
        'logp': np.random.normal(2, 1, n_samples),
        'tpsa': np.random.uniform(0, 150, n_samples),
        'rotatable_bonds': np.random.poisson(5, n_samples),
        'hbd': np.random.poisson(3, n_samples),
        'hba': np.random.poisson(6, n_samples),
        'radius_of_gyration': np.random.uniform(3, 8, n_samples),
        'molecular_volume': np.random.uniform(200, 800, n_samples),
        'surface_area': np.random.uniform(100, 400, n_samples),
        'asphericity': np.random.uniform(0.1, 0.8, n_samples)
    })
    
    # Create binding affinities (lower values = stronger binding)
    # Add some correlation with molecular properties
    y = pd.Series(
        -6.0 + 
        0.01 * X['molecular_weight'] + 
        0.5 * X['logp'] + 
        -0.02 * X['tpsa'] + 
        np.random.normal(0, 1.5, n_samples),
        name='binding_affinity'
    )
    
    # Create compound names
    compound_names = pd.Series([f'TLR4_compound_{i:04d}' for i in range(n_samples)])
    
    print(f"✓ Created dataset: {X.shape[0]} samples, {X.shape[1]} features")
    print(f"✓ Binding affinity range: [{y.min():.2f}, {y.max():.2f}] kcal/mol")
    
    return X, y, compound_names


def demo_basic_data_splitting():
    """Demonstrate basic data splitting functionality."""
    print("\n" + "="*60)
    print("DEMO 1: Basic Data Splitting")
    print("="*60)
    
    # Create sample data
    X, y, compound_names = create_sample_dataset(500)
    
    # Create data splitter with default configuration
    framework = DataValidationFramework()
    
    # Process dataset
    results = framework.process_dataset(X, y, compound_names, generate_quality_report=True)
    
    # Display results
    splits = results['data_splits']
    split_info = results['split_info']
    
    print(f"\nData Split Results:")
    print(f"  Train: {len(splits['X_train'])} samples ({len(splits['X_train'])/len(X)*100:.1f}%)")
    print(f"  Validation: {len(splits['X_val'])} samples ({len(splits['X_val'])/len(X)*100:.1f}%)")
    print(f"  Test: {len(splits['X_test'])} samples ({len(splits['X_test'])/len(X)*100:.1f}%)")
    
    print(f"\nTarget Statistics:")
    print(f"  Train mean: {splits['y_train'].mean():.2f} ± {splits['y_train'].std():.2f}")
    print(f"  Validation mean: {splits['y_val'].mean():.2f} ± {splits['y_val'].std():.2f}")
    print(f"  Test mean: {splits['y_test'].mean():.2f} ± {splits['y_test'].std():.2f}")
    
    # Validation status
    validation = results['split_validation']
    print(f"\nSplit Validation:")
    print(f"  Overall valid: {'✓' if validation['overall_valid'] else '✗'}")
    print(f"  Size validation: {'✓' if validation['size_validation']['status'] else '✗'}")
    print(f"  Distribution validation: {'✓' if validation['distribution_validation']['status'] else '✗'}")
    print(f"  No overlap: {'✓' if validation['overlap_validation']['status'] else '✗'}")


def demo_stratified_splitting():
    """Demonstrate stratified data splitting."""
    print("\n" + "="*60)
    print("DEMO 2: Stratified Data Splitting")
    print("="*60)
    
    # Create sample data with more extreme values for better stratification
    X, y, compound_names = create_sample_dataset(1000)
    
    # Add some extreme binding affinities
    y.iloc[0:50] = np.random.normal(-10, 1, 50)  # Very strong binders
    y.iloc[950:1000] = np.random.normal(-2, 1, 50)  # Weak binders
    
    # Create splitter with stratification enabled
    split_config = DataSplitConfig(
        test_size=0.15,
        validation_size=0.15,
        train_size=0.70,
        stratify=True,
        n_bins=5
    )
    
    framework = DataValidationFramework(split_config=split_config)
    results = framework.process_dataset(X, y, compound_names, generate_quality_report=False)
    
    splits = results['data_splits']
    
    print(f"\nStratified Split Results:")
    print(f"  Original target range: [{y.min():.2f}, {y.max():.2f}]")
    print(f"  Train range: [{splits['y_train'].min():.2f}, {splits['y_train'].max():.2f}]")
    print(f"  Validation range: [{splits['y_val'].min():.2f}, {splits['y_val'].max():.2f}]")
    print(f"  Test range: [{splits['y_test'].min():.2f}, {splits['y_test'].max():.2f}]")
    
    # Check distribution similarity
    train_mean = splits['y_train'].mean()
    val_mean = splits['y_val'].mean()
    test_mean = splits['y_test'].mean()
    overall_std = y.std()
    
    print(f"\nDistribution Similarity (should be close for stratified splitting):")
    print(f"  Train mean: {train_mean:.2f}")
    print(f"  Validation mean: {val_mean:.2f}")
    print(f"  Test mean: {test_mean:.2f}")
    print(f"  Mean difference tolerance: {0.2 * overall_std:.2f}")


def demo_cross_validation():
    """Demonstrate cross-validation setup."""
    print("\n" + "="*60)
    print("DEMO 3: Cross-Validation Setup")
    print("="*60)
    
    # Create sample data
    X, y, compound_names = create_sample_dataset(800)
    
    # Create framework with custom CV configuration
    cv_config = CrossValidationConfig(
        n_folds=5,
        cv_type='stratified_kfold',
        stratify=True
    )
    
    framework = DataValidationFramework(cv_config=cv_config)
    results = framework.process_dataset(X, y, compound_names, generate_quality_report=False)
    
    # Get CV splits
    splits = results['data_splits']
    cv = results['cv_setup']
    cv_splits = framework.cv_setup.get_cv_splits(splits['X_train'], splits['y_train'])
    
    print(f"\nCross-Validation Setup:")
    print(f"  CV type: {cv_config.cv_type}")
    print(f"  Number of folds: {cv_config.n_folds}")
    print(f"  Training samples for CV: {len(splits['X_train'])}")
    
    print(f"\nCV Fold Sizes:")
    for i, (train_idx, val_idx) in enumerate(cv_splits):
        print(f"  Fold {i+1}: Train={len(train_idx)}, Val={len(val_idx)}")
    
    # CV validation
    cv_validation = results['cv_validation']
    print(f"\nCV Validation:")
    print(f"  Overall valid: {'✓' if cv_validation['overall_valid'] else '✗'}")
    print(f"  Dataset size adequate: {'✓' if cv_validation['dataset_size_check']['status'] else '✗'}")
    print(f"  CV type appropriate: {'✓' if cv_validation['cv_type_appropriateness']['status'] else '✗'}")


def demo_quality_reporting():
    """Demonstrate data quality reporting."""
    print("\n" + "="*60)
    print("DEMO 4: Data Quality Reporting")
    print("="*60)
    
    # Create sample data with some issues
    X, y, compound_names = create_sample_dataset(600)
    
    # Introduce some data quality issues
    X.iloc[0:10, 0] = np.nan  # Missing values
    X.iloc[20:25, 1] = 1000   # Outliers
    X.iloc[30:35, 2] = -50    # More outliers
    
    framework = DataValidationFramework()
    results = framework.process_dataset(X, y, compound_names, generate_quality_report=True)
    
    quality_report = results['quality_report']
    
    print(f"\nDataset Overview:")
    overview = quality_report['dataset_overview']
    print(f"  Samples: {overview['n_samples']}")
    print(f"  Features: {overview['n_features']}")
    print(f"  Memory usage: {overview['memory_usage_mb']:.2f} MB")
    print(f"  Target range: [{overview['target_range'][0]:.2f}, {overview['target_range'][1]:.2f}]")
    
    print(f"\nMissing Data Analysis:")
    missing = quality_report['missing_data']
    print(f"  Features with missing values: {missing['features_with_missing']}")
    print(f"  Samples with missing values: {missing['samples_with_missing']}")
    print(f"  Missing percentage: {missing['missing_percentage']['features']:.1f}%")
    
    print(f"\nOutlier Analysis:")
    outliers = quality_report['outlier_analysis']
    print(f"  Target outliers: {outliers['target_outliers']['count']} ({outliers['target_outliers']['percentage']:.1f}%)")
    
    # Print formatted summary
    framework.quality_reporter.print_summary(quality_report)


def demo_custom_configurations():
    """Demonstrate custom configurations."""
    print("\n" + "="*60)
    print("DEMO 5: Custom Configurations")
    print("="*60)
    
    # Create sample data
    X, y, compound_names = create_sample_dataset(400)
    
    # Custom split configuration
    split_config = DataSplitConfig(
        test_size=0.2,
        validation_size=0.1,
        train_size=0.7,
        random_state=123,
        stratify=False  # Disable stratification
    )
    
    # Custom CV configuration
    cv_config = CrossValidationConfig(
        n_folds=3,
        cv_type='kfold',
        stratify=False
    )
    
    framework = DataValidationFramework(split_config, cv_config)
    results = framework.process_dataset(X, y, compound_names, generate_quality_report=False)
    
    splits = results['data_splits']
    split_info = results['split_info']
    
    print(f"\nCustom Split Configuration:")
    print(f"  Test size: {split_config.test_size}")
    print(f"  Validation size: {split_config.validation_size}")
    print(f"  Train size: {split_config.train_size}")
    print(f"  Stratify: {split_config.stratify}")
    
    print(f"\nActual Split Results:")
    print(f"  Train: {len(splits['X_train'])} ({len(splits['X_train'])/len(X)*100:.1f}%)")
    print(f"  Validation: {len(splits['X_val'])} ({len(splits['X_val'])/len(X)*100:.1f}%)")
    print(f"  Test: {len(splits['X_test'])} ({len(splits['X_test'])/len(X)*100:.1f}%)")
    
    print(f"\nCustom CV Configuration:")
    print(f"  CV type: {cv_config.cv_type}")
    print(f"  Number of folds: {cv_config.n_folds}")
    print(f"  Stratify: {cv_config.stratify}")


def demo_framework_summary():
    """Demonstrate framework summary functionality."""
    print("\n" + "="*60)
    print("DEMO 6: Framework Summary")
    print("="*60)
    
    # Create sample data
    X, y, compound_names = create_sample_dataset(750)
    
    framework = DataValidationFramework()
    results = framework.process_dataset(X, y, compound_names, generate_quality_report=True)
    
    # Get and display framework summary
    summary = framework.get_framework_summary()
    
    print(f"\nFramework Summary:")
    print(f"  Dataset: {summary['dataset_overview']['n_samples']} samples, {summary['dataset_overview']['n_features']} features")
    
    split_summary = summary['split_summary']
    if 'split_sizes' in split_summary:
        sizes = split_summary['split_sizes']
        print(f"  Splits: {sizes['train']} train, {sizes['validation']} val, {sizes['test']} test")
    
    validation_status = summary['validation_status']
    print(f"  Split validation: {'✓' if validation_status['split_valid'] else '✗'}")
    print(f"  CV validation: {'✓' if validation_status['cv_valid'] else '✗'}")
    
    # Print formatted summary
    print(f"\nFormatted Summary:")
    framework.print_framework_summary()


def main():
    """Run all demos."""
    print("TLR4 Binding Prediction - Data Splitting and Validation Framework Demo")
    print("="*80)
    
    try:
        demo_basic_data_splitting()
        demo_stratified_splitting()
        demo_cross_validation()
        demo_quality_reporting()
        demo_custom_configurations()
        demo_framework_summary()
        
        print("\n" + "="*80)
        print("✓ All demos completed successfully!")
        print("The data splitting and validation framework is ready for use.")
        print("="*80)
        
    except Exception as e:
        print(f"\n❌ Demo failed with error: {e}")
        import traceback
        traceback.print_exc()


if __name__ == "__main__":
    main()
```
