## Supplementary material for "Machine Learning-Driven Prediction of TLR4 Binding Affinity: A Comprehensive Molecular Feature Analysis for Drug Discovery": Codebase: demo_deep_learning_models.py

```
#!/usr/bin/env python3
"""
Demo script for deep learning models in TLR4 binding prediction.

This script demonstrates the usage of CNN, Transformer, and Multi-task
neural network models for molecular binding prediction.
"""

import sys
import os
import logging
import numpy as np
import pandas as pd
from pathlib import Path

# Add src to path
sys.path.append(str(Path(__file__).parent.parent / "src"))

from tlr4_binding.ml_components.deep_learning_models import (
    MolecularVoxelizer, VoxelDataset, SMILESDataset, SMILESData,
    MolecularCNN3D, MolecularTransformer, MultiTaskNeuralNetwork,
    DeepLearningTrainer
)
from tlr4_binding.ml_components.deep_learning_trainer import DeepLearningModelTrainer

# Set up logging
logging.basicConfig(level=logging.INFO)
logger = logging.getLogger(__name__)


def create_sample_pdbqt_file(filepath: str, compound_name: str = "sample") -> None:
    """Create a sample PDBQT file for testing."""
    content = f"""REMARK  VINA RESULT:    -6.5      0.000      0.000
REMARK  VINA RESULT:    -6.2      0.000      0.000
REMARK  VINA RESULT:    -5.8      0.000      0.000
ATOM      1  C   {compound_name}     1      12.345   23.456   34.567  1.00  0.00     0.000 C
ATOM      2  N   {compound_name}     1      13.345   24.456   35.567  1.00  0.00     0.000 N
ATOM      3  O   {compound_name}     1      14.345   25.456   36.567  1.00  0.00     0.000 O
ATOM      4  S   {compound_name}     1      15.345   26.456   37.567  1.00  0.00     0.000 S
ATOM      5  H   {compound_name}     1      11.345   22.456   33.567  1.00  0.00     0.000 H
ATOM      6  H   {compound_name}     1      12.345   24.456   35.567  1.00  0.00     0.000 H
"""
    
    with open(filepath, 'w') as f:
        f.write(content)


def demo_voxelizer():
    """Demonstrate molecular voxelizer functionality."""
    logger.info("=== Molecular Voxelizer Demo ===")
    
    # Create sample PDBQT files
    pdbqt_files = []
    binding_affinities = []
    
    for i in range(5):
        filepath = f"temp_compound_{i}.pdbqt"
        create_sample_pdbqt_file(filepath, f"compound_{i}")
        pdbqt_files.append(filepath)
        binding_affinities.append(-5.0 - i * 0.5)
    
    try:
        # Initialize voxelizer
        voxelizer = MolecularVoxelizer(grid_size=32, resolution=1.0)
        
        # Voxelize each file
        voxel_data = []
        for pdbqt_file, affinity in zip(pdbqt_files, binding_affinities):
            voxel_item = voxelizer.voxelize_pdbqt(pdbqt_file)
            if voxel_item is not None:
                voxel_item = voxel_item._replace(binding_affinity=affinity)
                voxel_data.append(voxel_item)
                logger.info(f"Voxelized {pdbqt_file}: shape {voxel_item.voxel_grid.shape}")
        
        # Create dataset
        dataset = VoxelDataset(voxel_data)
        logger.info(f"Created voxel dataset with {len(dataset)} samples")
        
        # Test dataset access
        sample_voxel, sample_affinity = dataset[0]
        logger.info(f"Sample voxel shape: {sample_voxel.shape}, affinity: {sample_affinity}")
        
    finally:
        # Clean up temporary files
        for filepath in pdbqt_files:
            if os.path.exists(filepath):
                os.unlink(filepath)


def demo_smiles_dataset():
    """Demonstrate SMILES dataset functionality."""
    logger.info("=== SMILES Dataset Demo ===")
    
    # Sample SMILES data
    smiles_data = [
        SMILESData(smiles="CCO", compound_name="ethanol", binding_affinity=-5.5),
        SMILESData(smiles="CC(=O)O", compound_name="acetic_acid", binding_affinity=-6.2),
        SMILESData(smiles="c1ccccc1", compound_name="benzene", binding_affinity=-4.8),
        SMILESData(smiles="CCN(CC)CC", compound_name="triethylamine", binding_affinity=-7.1),
        SMILESData(smiles="CC(C)CO", compound_name="isobutanol", binding_affinity=-5.9)
    ]
    
    # Create dataset
    dataset = SMILESDataset(smiles_data, vocab_size=100, max_length=64)
    logger.info(f"Created SMILES dataset with {len(dataset)} samples")
    logger.info(f"Vocabulary size: {dataset.vocab_size}, Max length: {dataset.max_length}")
    
    # Test tokenization
    tokens = dataset._tokenize_smiles("CCO")
    logger.info(f"Tokenized 'CCO': {tokens}")
    
    # Test dataset access
    sample_tokens, sample_affinity = dataset[0]
    logger.info(f"Sample tokens shape: {sample_tokens.shape}, affinity: {sample_affinity}")


def demo_cnn3d_model():
    """Demonstrate 3D CNN model."""
    logger.info("=== 3D CNN Model Demo ===")
    
    try:
        # Create model
        model = MolecularCNN3D(input_channels=1, num_classes=1, dropout=0.1)
        logger.info(f"Created CNN3D model with {sum(p.numel() for p in model.parameters())} parameters")
        
        # Test forward pass
        import torch
        x = torch.randn(2, 1, 32, 32, 32)  # batch_size=2, channels=1, 32x32x32 voxel grid
        output = model(x)
        logger.info(f"Input shape: {x.shape}, Output shape: {output.shape}")
        
        # Test without batch normalization
        model_no_bn = MolecularCNN3D(input_channels=1, num_classes=1, use_batch_norm=False)
        output_no_bn = model_no_bn(x)
        logger.info(f"Model without batch norm - Output shape: {output_no_bn.shape}")
        
    except ImportError:
        logger.warning("PyTorch not available. Skipping CNN3D model demo.")


def demo_transformer_model():
    """Demonstrate molecular transformer model."""
    logger.info("=== Molecular Transformer Demo ===")
    
    try:
        # Create model
        model = MolecularTransformer(vocab_size=100, d_model=128, nhead=4, num_layers=2, max_length=64)
        logger.info(f"Created Transformer model with {sum(p.numel() for p in model.parameters())} parameters")
        
        # Test forward pass
        import torch
        x = torch.randint(0, 100, (2, 10))  # batch_size=2, sequence_length=10
        output = model(x)
        logger.info(f"Input shape: {x.shape}, Output shape: {output.shape}")
        
        # Test with different sequence lengths
        for seq_len in [5, 15, 30]:
            x = torch.randint(0, 100, (2, seq_len))
            output = model(x)
            logger.info(f"Sequence length {seq_len} - Output shape: {output.shape}")
        
    except ImportError:
        logger.warning("PyTorch not available. Skipping Transformer model demo.")


def demo_multitask_model():
    """Demonstrate multi-task neural network."""
    logger.info("=== Multi-task Neural Network Demo ===")
    
    try:
        # Create model
        model = MultiTaskNeuralNetwork(input_dim=100, hidden_dims=[64, 32], num_tasks=3)
        logger.info(f"Created Multi-task model with {sum(p.numel() for p in model.parameters())} parameters")
        logger.info(f"Task names: {model.task_names}")
        
        # Test forward pass
        import torch
        x = torch.randn(4, 100)  # batch_size=4, input_dim=100
        outputs = model(x)
        logger.info(f"Input shape: {x.shape}, Number of outputs: {len(outputs)}")
        
        for i, output in enumerate(outputs):
            logger.info(f"Task {i} ({model.task_names[i]}): {output.shape}")
        
        # Test single task prediction
        for task_idx in range(3):
            single_output = model.predict_single_task(x, task_idx)
            logger.info(f"Single task {task_idx} prediction shape: {single_output.shape}")
        
    except ImportError:
        logger.warning("PyTorch not available. Skipping Multi-task model demo.")


def demo_deep_learning_trainer():
    """Demonstrate deep learning trainer."""
    logger.info("=== Deep Learning Trainer Demo ===")
    
    try:
        # Create trainer
        trainer = DeepLearningModelTrainer(models_dir="temp_models")
        logger.info("Created deep learning model trainer")
        
        # Create sample data
        pdbqt_files = []
        binding_affinities = []
        smiles_list = []
        compound_names = []
        
        for i in range(5):
            # PDBQT files
            filepath = f"temp_compound_{i}.pdbqt"
            create_sample_pdbqt_file(filepath, f"compound_{i}")
            pdbqt_files.append(filepath)
            binding_affinities.append(-5.0 - i * 0.5)
            
            # SMILES data
            smiles_list.append(["CCO", "CC(=O)O", "c1ccccc1", "CCN(CC)CC", "CC(C)CO"][i])
            compound_names.append(f"compound_{i}")
        
        # Create features DataFrame
        features_df = pd.DataFrame({
            'molecular_weight': [46.07, 60.05, 78.11, 101.19, 74.12],
            'logp': [-0.31, -0.17, 2.13, 0.15, 0.76],
            'tpsa': [20.23, 37.30, 0.00, 3.24, 20.23],
            'rotatable_bonds': [0, 0, 0, 2, 1],
            'hbd': [1, 1, 0, 0, 1],
            'hba': [1, 2, 0, 1, 1]
        })
        
        # Prepare data
        train_indices = [0, 1, 2]
        val_indices = [3, 4]
        
        logger.info("Preparing voxel data...")
        train_voxel_dataset = trainer.prepare_voxel_data(
            [pdbqt_files[i] for i in train_indices],
            [binding_affinities[i] for i in train_indices]
        )
        
        logger.info("Preparing SMILES data...")
        train_smiles_dataset = trainer.prepare_smiles_data(
            [smiles_list[i] for i in train_indices],
            [compound_names[i] for i in train_indices],
            [binding_affinities[i] for i in train_indices]
        )
        
        logger.info("Preparing multi-task data...")
        features_tensor, targets_tensors = trainer.prepare_multitask_data(
            features_df.iloc[train_indices],
            pd.Series([binding_affinities[i] for i in train_indices])
        )
        
        logger.info(f"Voxel dataset size: {len(train_voxel_dataset)}")
        logger.info(f"SMILES dataset size: {len(train_smiles_dataset)}")
        logger.info(f"Features tensor shape: {features_tensor.shape}")
        logger.info(f"Number of target tasks: {len(targets_tensors)}")
        
    except Exception as e:
        logger.error(f"Error in deep learning trainer demo: {str(e)}")
    
    finally:
        # Clean up temporary files
        for filepath in pdbqt_files:
            if os.path.exists(filepath):
                os.unlink(filepath)
        
        # Clean up models directory
        import shutil
        if os.path.exists("temp_models"):
            shutil.rmtree("temp_models")


def demo_model_training():
    """Demonstrate model training (if PyTorch is available)."""
    logger.info("=== Model Training Demo ===")
    
    try:
        import torch
        from torch.utils.data import DataLoader
        
        # Create simple training data
        voxelizer = MolecularVoxelizer(grid_size=16, resolution=2.0)
        
        # Create sample PDBQT files
        pdbqt_files = []
        binding_affinities = []
        
        for i in range(10):
            filepath = f"temp_training_{i}.pdbqt"
            create_sample_pdbqt_file(filepath, f"training_{i}")
            pdbqt_files.append(filepath)
            binding_affinities.append(-5.0 - i * 0.3)
        
        try:
            # Prepare voxel data
            voxel_data = []
            for pdbqt_file, affinity in zip(pdbqt_files, binding_affinities):
                voxel_item = voxelizer.voxelize_pdbqt(pdbqt_file)
                if voxel_item is not None:
                    voxel_item = voxel_item._replace(binding_affinity=affinity)
                    voxel_data.append(voxel_item)
            
            dataset = VoxelDataset(voxel_data)
            data_loader = DataLoader(dataset, batch_size=2, shuffle=True, num_workers=0)
            
            # Create and train CNN3D model
            trainer = DeepLearningTrainer(model_type="cnn3d", device="cpu")
            trainer.model = trainer.create_model(input_channels=1, num_classes=1)
            
            logger.info("Training CNN3D model for 3 epochs...")
            history = trainer.train(data_loader, epochs=3, lr=0.001)
            
            logger.info(f"Training completed. Final train loss: {history['train_loss'][-1]:.4f}")
            logger.info(f"Final validation loss: {history['val_loss'][-1]:.4f}")
            
        finally:
            # Clean up
            for filepath in pdbqt_files:
                if os.path.exists(filepath):
                    os.unlink(filepath)
    
    except ImportError:
        logger.warning("PyTorch not available. Skipping model training demo.")
    except Exception as e:
        logger.error(f"Error in model training demo: {str(e)}")


def main():
    """Run all demos."""
    logger.info("Starting Deep Learning Models Demo")
    logger.info("=" * 50)
    
    # Run individual demos
    demo_voxelizer()
    print()
    
    demo_smiles_dataset()
    print()
    
    demo_cnn3d_model()
    print()
    
    demo_transformer_model()
    print()
    
    demo_multitask_model()
    print()
    
    demo_deep_learning_trainer()
    print()
    
    demo_model_training()
    print()
    
    logger.info("Deep Learning Models Demo completed!")


if __name__ == "__main__":
    main()
```
