## Supplementary material for "Machine Learning-Driven Prediction of TLR4 Binding Affinity: A Comprehensive Molecular Feature Analysis for Drug Discovery": Codebase: demo_ensemble_models.py

```
"""
Demo script for ensemble and hybrid models.

This script demonstrates the usage of various ensemble methods
including stacked ensembles, weighted ensembles, and physics-informed models.
"""

import pandas as pd
import numpy as np
import logging
from pathlib import Path
import matplotlib.pyplot as plt
import seaborn as sns
from typing import Dict, List, Optional

# Set up logging
logging.basicConfig(level=logging.INFO)
logger = logging.getLogger(__name__)

# Import ensemble models
from src.tlr4_binding.ml_components.ensemble_models import (
    StackedEnsemble, WeightedEnsemble, PhysicsInformedNeuralNetwork,
    PhysicsInformedEnsemble, EnsembleModelTrainer
)

# Import base model trainers
from src.tlr4_binding.ml_components.trainer import (
    RandomForestTrainer, SVRTrainer, XGBoostTrainer, LightGBMTrainer
)

# Import data processing components
from src.tlr4_binding.data_processing.data_loader import BindingDataLoader
from src.tlr4_binding.molecular_analysis.feature_extractor import MolecularFeatureExtractor
from src.tlr4_binding.ml_components.feature_engineering import FeatureEngineeringPipeline
from src.tlr4_binding.ml_components.data_splitting import DataSplitter


def create_synthetic_data(n_samples: int = 1000, random_state: int = 42) -> tuple:
    """
    Create synthetic molecular binding data for demonstration.
    
    Args:
        n_samples: Number of samples to generate
        random_state: Random state for reproducibility
        
    Returns:
        Tuple of (features_df, binding_affinities)
    """
    np.random.seed(random_state)
    
    # Generate molecular features
    features_data = {
        'molecular_weight': np.random.uniform(100, 800, n_samples),
        'logp': np.random.uniform(-2, 6, n_samples),
        'tpsa': np.random.uniform(20, 200, n_samples),
        'rotatable_bonds': np.random.randint(0, 15, n_samples),
        'hbd': np.random.randint(0, 8, n_samples),
        'hba': np.random.randint(0, 12, n_samples),
        'radius_of_gyration': np.random.uniform(2, 8, n_samples),
        'molecular_volume': np.random.uniform(100, 1000, n_samples),
        'surface_area': np.random.uniform(200, 800, n_samples),
        'asphericity': np.random.uniform(0, 1, n_samples),
        'ring_count': np.random.randint(0, 6, n_samples),
        'aromatic_rings': np.random.randint(0, 4, n_samples),
        'formal_charge': np.random.randint(-2, 3, n_samples)
    }
    
    features_df = pd.DataFrame(features_data)
    
    # Generate binding affinities with some realistic relationships
    binding_affinities = (
        -2.0 * features_df['molecular_weight'] / 1000 +  # Heavier molecules bind better
        0.5 * features_df['logp'] +  # Lipophilicity effect
        -0.01 * features_df['tpsa'] +  # TPSA penalty
        0.1 * features_df['rotatable_bonds'] +  # Flexibility penalty
        -0.2 * features_df['hbd'] +  # H-bond donors help
        -0.1 * features_df['hba'] +  # H-bond acceptors help
        np.random.normal(0, 0.5, n_samples)  # Random noise
    )
    
    return features_df, binding_affinities


def create_base_models() -> Dict[str, any]:
    """
    Create base models for ensemble training.
    
    Returns:
        Dictionary of base model trainers
    """
    base_models = {}
    
    try:
        base_models['random_forest'] = RandomForestTrainer(n_jobs=1, random_state=42)
        logger.info("Added Random Forest trainer")
    except Exception as e:
        logger.warning(f"Could not create Random Forest trainer: {e}")
    
    try:
        base_models['svr'] = SVRTrainer(random_state=42)
        logger.info("Added SVR trainer")
    except Exception as e:
        logger.warning(f"Could not create SVR trainer: {e}")
    
    try:
        base_models['xgboost'] = XGBoostTrainer(random_state=42)
        logger.info("Added XGBoost trainer")
    except Exception as e:
        logger.warning(f"Could not create XGBoost trainer: {e}")
    
    try:
        base_models['lightgbm'] = LightGBMTrainer(random_state=42)
        logger.info("Added LightGBM trainer")
    except Exception as e:
        logger.warning(f"Could not create LightGBM trainer: {e}")
    
    return base_models


def train_individual_models(X_train: pd.DataFrame, y_train: pd.Series,
                          X_val: pd.DataFrame, y_val: pd.Series,
                          base_models: Dict[str, any]) -> Dict[str, any]:
    """
    Train individual base models for comparison.
    
    Args:
        X_train: Training features
        y_train: Training targets
        X_val: Validation features
        y_val: Validation targets
        base_models: Dictionary of base model trainers
        
    Returns:
        Dictionary of trained models and their performance
    """
    logger.info("Training individual base models")
    
    trained_models = {}
    model_performance = {}
    
    for model_name, trainer in base_models.items():
        try:
            logger.info(f"Training {model_name}")
            
            # Train model
            model = trainer.train(X_train, y_train, X_val, y_val)
            trained_models[model_name] = model
            
            # Evaluate on validation set
            val_predictions = trainer.predict(model, X_val)
            
            # Calculate metrics
            mse = np.mean((y_val - val_predictions) ** 2)
            rmse = np.sqrt(mse)
            mae = np.mean(np.abs(y_val - val_predictions))
            
            # R² score
            ss_res = np.sum((y_val - val_predictions) ** 2)
            ss_tot = np.sum((y_val - np.mean(y_val)) ** 2)
            r2 = 1 - (ss_res / ss_tot) if ss_tot != 0 else 0
            
            model_performance[model_name] = {
                'mse': mse,
                'rmse': rmse,
                'mae': mae,
                'r2': r2
            }
            
            logger.info(f"{model_name} - R²: {r2:.4f}, RMSE: {rmse:.4f}")
            
        except Exception as e:
            logger.error(f"Error training {model_name}: {e}")
            continue
    
    return trained_models, model_performance


def train_ensemble_models(X_train: pd.DataFrame, y_train: pd.Series,
                         X_val: pd.DataFrame, y_val: pd.Series,
                         base_models: Dict[str, any]) -> Dict[str, any]:
    """
    Train ensemble models.
    
    Args:
        X_train: Training features
        y_train: Training targets
        X_val: Validation features
        y_val: Validation targets
        base_models: Dictionary of base model trainers
        
    Returns:
        Dictionary of trained ensemble models and their performance
    """
    logger.info("Training ensemble models")
    
    # Create ensemble trainer
    ensemble_trainer = EnsembleModelTrainer(
        models_dir="models/ensemble_demo",
        include_physics_informed=True
    )
    
    # Train ensembles
    ensemble_models = ensemble_trainer.train_ensembles(
        X_train, y_train, X_val, y_val, base_models
    )
    
    # Evaluate ensembles
    ensemble_performance = ensemble_trainer.evaluate_ensembles(X_val, y_val)
    
    return ensemble_models, ensemble_performance


def compare_model_performance(individual_performance: Dict[str, Dict],
                            ensemble_performance: Dict[str, Dict]) -> pd.DataFrame:
    """
    Compare performance of individual and ensemble models.
    
    Args:
        individual_performance: Performance of individual models
        ensemble_performance: Performance of ensemble models
        
    Returns:
        DataFrame with performance comparison
    """
    comparison_data = []
    
    # Add individual model performance
    for model_name, metrics in individual_performance.items():
        comparison_data.append({
            'model_name': model_name,
            'model_type': 'Individual',
            **metrics
        })
    
    # Add ensemble model performance
    for ensemble_name, results in ensemble_performance.items():
        if 'metrics' in results:
            comparison_data.append({
                'model_name': ensemble_name,
                'model_type': 'Ensemble',
                **results['metrics']
            })
    
    return pd.DataFrame(comparison_data)


def plot_performance_comparison(comparison_df: pd.DataFrame, save_path: str = None):
    """
    Plot performance comparison between models.
    
    Args:
        comparison_df: DataFrame with performance metrics
        save_path: Path to save the plot
    """
    fig, axes = plt.subplots(2, 2, figsize=(15, 10))
    fig.suptitle('Model Performance Comparison', fontsize=16)
    
    # R² Score comparison
    axes[0, 0].bar(range(len(comparison_df)), comparison_df['r2'])
    axes[0, 0].set_title('R² Score')
    axes[0, 0].set_xticks(range(len(comparison_df)))
    axes[0, 0].set_xticklabels(comparison_df['model_name'], rotation=45, ha='right')
    axes[0, 0].set_ylabel('R² Score')
    
    # RMSE comparison
    axes[0, 1].bar(range(len(comparison_df)), comparison_df['rmse'])
    axes[0, 1].set_title('RMSE')
    axes[0, 1].set_xticks(range(len(comparison_df)))
    axes[0, 1].set_xticklabels(comparison_df['model_name'], rotation=45, ha='right')
    axes[0, 1].set_ylabel('RMSE')
    
    # MAE comparison
    axes[1, 0].bar(range(len(comparison_df)), comparison_df['mae'])
    axes[1, 0].set_title('MAE')
    axes[1, 0].set_xticks(range(len(comparison_df)))
    axes[1, 0].set_xticklabels(comparison_df['model_name'], rotation=45, ha='right')
    axes[1, 0].set_ylabel('MAE')
    
    # Model type distribution
    model_type_counts = comparison_df['model_type'].value_counts()
    axes[1, 1].pie(model_type_counts.values, labels=model_type_counts.index, autopct='%1.1f%%')
    axes[1, 1].set_title('Model Type Distribution')
    
    plt.tight_layout()
    
    if save_path:
        plt.savefig(save_path, dpi=300, bbox_inches='tight')
        logger.info(f"Performance comparison plot saved to {save_path}")
    
    plt.show()


def demonstrate_uncertainty_quantification(ensemble_models: Dict[str, any],
                                         X_test: pd.DataFrame,
                                         y_test: pd.Series,
                                         n_samples: int = 10):
    """
    Demonstrate uncertainty quantification for ensemble models.
    
    Args:
        ensemble_models: Dictionary of trained ensemble models
        X_test: Test features
        y_test: Test targets
        n_samples: Number of samples to show
    """
    logger.info("Demonstrating uncertainty quantification")
    
    # Select a subset of test samples
    test_subset = X_test.iloc[:n_samples]
    true_values = y_test.iloc[:n_samples]
    
    for ensemble_name, ensemble in ensemble_models.items():
        try:
            logger.info(f"Analyzing uncertainty for {ensemble_name}")
            
            # Get predictions with uncertainty
            uncertainty_predictions = ensemble.predict_with_uncertainty(test_subset)
            
            # Print results
            print(f"\n{ensemble_name} Uncertainty Analysis:")
            print("-" * 50)
            print(f"{'Sample':<8} {'True':<8} {'Pred':<8} {'Uncert':<8} {'CI_Lower':<10} {'CI_Upper':<10}")
            print("-" * 50)
            
            for i, pred in enumerate(uncertainty_predictions):
                print(f"{i+1:<8} {true_values.iloc[i]:<8.3f} {pred.prediction:<8.3f} "
                      f"{pred.uncertainty:<8.3f} {pred.confidence_interval[0]:<10.3f} "
                      f"{pred.confidence_interval[1]:<10.3f}")
            
            # Calculate coverage (percentage of true values within confidence interval)
            coverage = 0
            for i, pred in enumerate(uncertainty_predictions):
                if pred.confidence_interval[0] <= true_values.iloc[i] <= pred.confidence_interval[1]:
                    coverage += 1
            
            coverage_pct = (coverage / len(uncertainty_predictions)) * 100
            print(f"\nCoverage: {coverage_pct:.1f}% (expected: ~95%)")
            
        except Exception as e:
            logger.error(f"Error analyzing uncertainty for {ensemble_name}: {e}")


def main():
    """Main demonstration function."""
    logger.info("Starting ensemble models demonstration")
    
    # Create synthetic data
    logger.info("Creating synthetic molecular binding data")
    features_df, binding_affinities = create_synthetic_data(n_samples=1000)
    
    # Split data
    from sklearn.model_selection import train_test_split
    X_train, X_test, y_train, y_test = train_test_split(
        features_df, binding_affinities, test_size=0.2, random_state=42
    )
    X_train, X_val, y_train, y_val = train_test_split(
        X_train, y_train, test_size=0.25, random_state=42
    )
    
    logger.info(f"Data split - Train: {len(X_train)}, Val: {len(X_val)}, Test: {len(X_test)}")
    
    # Create base models
    base_models = create_base_models()
    if not base_models:
        logger.error("No base models available. Please check dependencies.")
        return
    
    # Train individual models
    individual_models, individual_performance = train_individual_models(
        X_train, y_train, X_val, y_val, base_models
    )
    
    # Train ensemble models
    ensemble_models, ensemble_performance = train_ensemble_models(
        X_train, y_train, X_val, y_val, base_models
    )
    
    # Compare performance
    comparison_df = compare_model_performance(individual_performance, ensemble_performance)
    print("\nModel Performance Comparison:")
    print("=" * 60)
    print(comparison_df.to_string(index=False, float_format='%.4f'))
    
    # Plot performance comparison
    plot_performance_comparison(comparison_df, "ensemble_performance_comparison.png")
    
    # Demonstrate uncertainty quantification
    demonstrate_uncertainty_quantification(ensemble_models, X_test, y_test)
    
    # Find best model
    best_r2_idx = comparison_df['r2'].idxmax()
    best_model = comparison_df.iloc[best_r2_idx]
    
    print(f"\nBest Model: {best_model['model_name']} ({best_model['model_type']})")
    print(f"R² Score: {best_model['r2']:.4f}")
    print(f"RMSE: {best_model['rmse']:.4f}")
    print(f"MAE: {best_model['mae']:.4f}")
    
    logger.info("Ensemble models demonstration completed successfully")


if __name__ == "__main__":
    main()
```
