## Supplementary material for "Machine Learning-Driven Prediction of TLR4 Binding Affinity: A Comprehensive Molecular Feature Analysis for Drug Discovery": Codebase: demo_gnn_models.py

```
#!/usr/bin/env python3
"""
Demo script for Graph Neural Network models in TLR4 binding prediction.

This script demonstrates the GNN functionality including:
- Molecular graph construction from PDBQT files
- Training of various GNN architectures
- Model evaluation and visualization
- Integration with traditional ML models
"""

import sys
import logging
from pathlib import Path
import numpy as np
import pandas as pd
import matplotlib.pyplot as plt

# Add src to path
sys.path.append(str(Path(__file__).parent.parent / "src"))

from tlr4_binding.ml_components.trainer import MLModelTrainer
from tlr4_binding.ml_components.gnn_models import (
    MolecularGraphBuilder, GNNModelTrainer, GraphDataset
)
from tlr4_binding.ml_components.gnn_visualization import (
    GraphVisualizer, GNNModelAnalyzer
)
from tlr4_binding.ml_components.data_splitting import DataSplitter
from tlr4_binding.data_processing.binding_data_loader import BindingDataLoader
from tlr4_binding.molecular_analysis.molecular_feature_extractor import MolecularFeatureExtractor

# Set up logging
logging.basicConfig(level=logging.INFO, format='%(asctime)s - %(levelname)s - %(message)s')
logger = logging.getLogger(__name__)


def create_sample_pdbqt_files(num_files: int = 10) -> list:
    """Create sample PDBQT files for demonstration."""
    sample_dir = Path("data/sample_pdbqt")
    sample_dir.mkdir(parents=True, exist_ok=True)
    
    pdbqt_files = []
    
    for i in range(num_files):
        pdbqt_file = sample_dir / f"compound_{i:03d}.pdbqt"
        
        # Create a simple PDBQT file with random coordinates
        with open(pdbqt_file, 'w') as f:
            f.write("REMARK  Generated sample PDBQT file\n")
            f.write("REMARK  Compound: sample_compound\n")
            
            # Add some random atoms
            for j in range(5 + i % 10):  # 5-14 atoms per molecule
                x = np.random.uniform(-5, 5)
                y = np.random.uniform(-5, 5)
                z = np.random.uniform(-5, 5)
                atom_type = np.random.choice(['C', 'N', 'O', 'H'])
                
                f.write(f"ATOM  {j+1:5d}  {atom_type}   LIG A   1    "
                       f"{x:8.3f}{y:8.3f}{z:8.3f}  1.00  0.00          {atom_type}  \n")
        
        pdbqt_files.append(str(pdbqt_file))
    
    return pdbqt_files


def create_sample_binding_data(num_compounds: int = 10) -> list:
    """Create sample binding affinity data."""
    # Generate realistic binding affinities (kcal/mol)
    # Lower values = stronger binding
    binding_affinities = np.random.uniform(-12.0, -2.0, num_compounds)
    return binding_affinities.tolist()


def demo_molecular_graph_construction():
    """Demonstrate molecular graph construction from PDBQT files."""
    logger.info("=== Molecular Graph Construction Demo ===")
    
    # Create sample PDBQT files
    pdbqt_files = create_sample_pdbqt_files(5)
    logger.info(f"Created {len(pdbqt_files)} sample PDBQT files")
    
    # Initialize graph builder
    graph_builder = MolecularGraphBuilder()
    
    # Build graphs from PDBQT files
    graphs = []
    for pdbqt_file in pdbqt_files:
        graph = graph_builder.build_graph_from_pdbqt(pdbqt_file)
        if graph is not None:
            graphs.append(graph)
            logger.info(f"Built graph from {Path(pdbqt_file).name}: "
                      f"{graph.num_nodes} nodes, {graph.num_edges} edges")
    
    logger.info(f"Successfully built {len(graphs)} molecular graphs")
    return graphs, pdbqt_files


def demo_gnn_model_training():
    """Demonstrate GNN model training."""
    logger.info("=== GNN Model Training Demo ===")
    
    # Create sample data
    pdbqt_files = create_sample_pdbqt_files(20)
    binding_affinities = create_sample_binding_data(20)
    
    # Initialize GNN trainer
    gnn_trainer = GNNModelTrainer(models_dir="models/gnn_demo")
    
    # Prepare graph dataset
    logger.info("Preparing graph dataset...")
    dataset = gnn_trainer.prepare_graph_data(pdbqt_files, binding_affinities)
    logger.info(f"Created dataset with {len(dataset)} graphs")
    
    # Split data
    train_size = int(0.7 * len(dataset))
    val_size = int(0.15 * len(dataset))
    
    train_dataset = GraphDataset(dataset.graphs[:train_size], 
                               dataset.targets[:train_size])
    val_dataset = GraphDataset(dataset.graphs[train_size:train_size+val_size], 
                             dataset.targets[train_size:train_size+val_size])
    test_dataset = GraphDataset(dataset.graphs[train_size+val_size:], 
                              dataset.targets[train_size+val_size:])
    
    logger.info(f"Data split: {len(train_dataset)} train, {len(val_dataset)} val, {len(test_dataset)} test")
    
    # Train GNN models
    logger.info("Training GNN models...")
    model_types = ["graphconv", "mpnn", "attentivefp"]
    
    try:
        gnn_models = gnn_trainer.train_models(
            train_dataset, val_dataset, model_types,
            epochs=50, lr=0.001, batch_size=8
        )
        logger.info(f"Successfully trained {len(gnn_models)} GNN models")
        
        # Evaluate models
        logger.info("Evaluating GNN models...")
        evaluation_results = gnn_trainer.evaluate_models(test_dataset)
        
        # Print results
        for model_name, results in evaluation_results.items():
            metrics = results['metrics']
            logger.info(f"{model_name}: R² = {metrics['r2']:.3f}, "
                       f"RMSE = {metrics['rmse']:.3f}, MAE = {metrics['mae']:.3f}")
        
        return gnn_trainer, evaluation_results
        
    except Exception as e:
        logger.error(f"Error training GNN models: {str(e)}")
        logger.info("This is expected if PyTorch/PyTorch Geometric are not installed")
        return None, {}


def demo_gnn_visualization():
    """Demonstrate GNN visualization capabilities."""
    logger.info("=== GNN Visualization Demo ===")
    
    # Create sample graphs
    graphs, pdbqt_files = demo_molecular_graph_construction()
    
    if not graphs:
        logger.warning("No graphs available for visualization")
        return
    
    # Initialize visualizer
    visualizer = GraphVisualizer(output_dir="results/gnn_visualizations")
    
    # Visualize first few graphs
    for i, graph in enumerate(graphs[:3]):
        compound_name = f"compound_{i:03d}"
        save_path = f"results/gnn_visualizations/{compound_name}_graph.png"
        
        try:
            fig = visualizer.visualize_molecular_graph(graph, compound_name, save_path)
            if fig is not None:
                logger.info(f"Visualized graph for {compound_name}")
                plt.close(fig)
        except Exception as e:
            logger.warning(f"Could not visualize graph {i}: {str(e)}")
    
    # Test feature extraction
    feature_extractor = GraphFeatureExtractor()
    for i, graph in enumerate(graphs[:2]):
        features = feature_extractor.extract_graph_features(graph)
        logger.info(f"Graph {i} features: {list(features.keys())}")


def demo_integrated_training():
    """Demonstrate integrated training with both traditional ML and GNN models."""
    logger.info("=== Integrated Training Demo ===")
    
    # Create sample data
    pdbqt_files = create_sample_pdbqt_files(30)
    binding_affinities = create_sample_binding_data(30)
    
    # Create some mock molecular features for traditional ML
    num_features = 20
    X = pd.DataFrame(np.random.randn(30, num_features), 
                    columns=[f"feature_{i}" for i in range(num_features)])
    y = pd.Series(binding_affinities)
    
    # Initialize integrated trainer
    trainer = MLModelTrainer(models_dir="models/integrated_demo", include_gnn=True)
    
    # Split data
    splitter = DataSplitter()
    train_indices, val_indices, test_indices = splitter.split_data(
        X, y, train_size=0.7, val_size=0.15, test_size=0.15, random_state=42
    )
    
    # Train traditional ML models
    logger.info("Training traditional ML models...")
    try:
        X_train = X.iloc[train_indices]
        y_train = y.iloc[train_indices]
        X_val = X.iloc[val_indices]
        y_val = y.iloc[val_indices]
        
        traditional_models = trainer.train_models(X_train, y_train, X_val, y_val)
        logger.info(f"Trained {len(traditional_models)} traditional ML models")
    except Exception as e:
        logger.error(f"Error training traditional ML models: {str(e)}")
        traditional_models = {}
    
    # Train GNN models
    logger.info("Training GNN models...")
    try:
        gnn_models = trainer.train_gnn_models(
            pdbqt_files, binding_affinities,
            train_indices, val_indices, test_indices,
            model_types=["graphconv", "mpnn"]
        )
        logger.info(f"Trained {len(gnn_models)} GNN models")
    except Exception as e:
        logger.error(f"Error training GNN models: {str(e)}")
        gnn_models = {}
    
    # Get comprehensive model summary
    if traditional_models or gnn_models:
        summary_df = trainer.get_model_summary(include_gnn=True)
        logger.info("Model Performance Summary:")
        logger.info(f"\n{summary_df.to_string(index=False)}")
        
        # Get best model
        try:
            best_model_name, best_model = trainer.get_best_model(metric='r2', include_gnn=True)
            logger.info(f"Best model: {best_model_name}")
        except Exception as e:
            logger.warning(f"Could not determine best model: {str(e)}")
    
    return trainer


def main():
    """Main demo function."""
    logger.info("Starting GNN Models Demo for TLR4 Binding Prediction")
    logger.info("=" * 60)
    
    # Create output directories
    Path("results/gnn_visualizations").mkdir(parents=True, exist_ok=True)
    Path("models/gnn_demo").mkdir(parents=True, exist_ok=True)
    Path("models/integrated_demo").mkdir(parents=True, exist_ok=True)
    
    try:
        # Demo 1: Molecular graph construction
        demo_molecular_graph_construction()
        
        # Demo 2: GNN model training
        demo_gnn_model_training()
        
        # Demo 3: GNN visualization
        demo_gnn_visualization()
        
        # Demo 4: Integrated training
        demo_integrated_training()
        
        logger.info("=" * 60)
        logger.info("GNN Models Demo completed successfully!")
        logger.info("Check the 'results/' and 'models/' directories for outputs.")
        
    except Exception as e:
        logger.error(f"Demo failed with error: {str(e)}")
        logger.info("Note: Some features require PyTorch and PyTorch Geometric to be installed")
        logger.info("Install with: pip install torch torch-geometric")


if __name__ == "__main__":
    main()
```
