## Supplementary material for "Machine Learning-Driven Prediction of TLR4 Binding Affinity: A Comprehensive Molecular Feature Analysis for Drug Discovery": Codebase: demo_interpretability.py

```
#!/usr/bin/env python3
"""
Demo script for Model Interpretability and Analysis Suite

This script demonstrates the comprehensive interpretability tools for
understanding what molecular features drive strong TLR4 binding.
"""

import sys
import os
import pandas as pd
import numpy as np
import matplotlib.pyplot as plt
import warnings
warnings.filterwarnings('ignore')

# Add src to path
sys.path.append('src')

from tlr4_binding.ml_components.interpretability import ModelInterpretabilitySuite
from tlr4_binding.ml_components.molecular_substructure_analyzer import MolecularSubstructureAnalyzer
from tlr4_binding.ml_components.attention_visualizer import AttentionVisualizer
from tlr4_binding.ml_components.trainer import MLModelTrainer
from tlr4_binding.ml_components.data_splitting import DataSplitter
from tlr4_binding.data_processing.binding_data_loader import BindingDataLoader
from tlr4_binding.molecular_analysis.molecular_feature_extractor import MolecularFeatureExtractor
from tlr4_binding.ml_components.feature_engineering import FeatureEngineer

def load_sample_data():
    """Load sample data for interpretability analysis."""
    print("Loading sample data for interpretability analysis...")
    
    # Load binding data
    binding_loader = BindingDataLoader("binding-data/processed_logs.csv")
    binding_data = binding_loader.load_data()
    
    # Load molecular features (if available)
    try:
        feature_extractor = MolecularFeatureExtractor()
        # This would normally process PDBQT files, but for demo we'll create sample data
        print("Creating sample molecular features for demo...")
        
        # Create sample molecular features
        n_samples = len(binding_data)
        molecular_features = pd.DataFrame({
            'mol_weight': np.random.normal(300, 50, n_samples),
            'mol_logp': np.random.normal(2.5, 1.0, n_samples),
            'mol_tpsa': np.random.normal(80, 20, n_samples),
            'mol_hbd': np.random.poisson(3, n_samples),
            'mol_hba': np.random.poisson(5, n_samples),
            'mol_rotatable_bonds': np.random.poisson(8, n_samples),
            'mol_aromatic_rings': np.random.poisson(2, n_samples),
            'mol_heavy_atoms': np.random.poisson(25, n_samples),
            'mol_formal_charge': np.random.choice([-1, 0, 1], n_samples),
            'smiles': [f"CCOC{i}" for i in range(n_samples)]  # Dummy SMILES
        })
        
        # Combine data
        combined_data = pd.concat([binding_data, molecular_features], axis=1)
        
        return combined_data
        
    except Exception as e:
        print(f"Error loading molecular features: {str(e)}")
        print("Using binding data only for demo...")
        return binding_data

def train_sample_models(X_train, y_train, X_test, y_test):
    """Train sample models for interpretability analysis."""
    print("Training sample models for interpretability analysis...")
    
    from sklearn.ensemble import RandomForestRegressor
    from sklearn.svm import SVR
    from sklearn.linear_model import LinearRegression
    from sklearn.preprocessing import StandardScaler
    
    # Scale features
    scaler = StandardScaler()
    X_train_scaled = scaler.fit_transform(X_train)
    X_test_scaled = scaler.transform(X_test)
    
    # Train models
    models = {}
    
    # Random Forest
    rf_model = RandomForestRegressor(n_estimators=100, random_state=42)
    rf_model.fit(X_train, y_train)
    models['RandomForest'] = rf_model
    
    # Support Vector Regression
    svr_model = SVR(kernel='rbf', C=1.0, gamma='scale')
    svr_model.fit(X_train_scaled, y_train)
    models['SVR'] = svr_model
    
    # Linear Regression
    lr_model = LinearRegression()
    lr_model.fit(X_train_scaled, y_train)
    models['LinearRegression'] = lr_model
    
    print(f"Trained {len(models)} models for interpretability analysis")
    return models, scaler

def run_interpretability_analysis():
    """Run comprehensive interpretability analysis."""
    print("=" * 60)
    print("TLR4 Binding Prediction - Model Interpretability Analysis")
    print("=" * 60)
    
    # Load data
    data = load_sample_data()
    print(f"Loaded data with {len(data)} samples")
    
    # Prepare features and target
    feature_columns = [col for col in data.columns if col.startswith('mol_') and col != 'smiles']
    if not feature_columns:
        print("No molecular features found. Creating dummy features...")
        feature_columns = [f'feature_{i}' for i in range(10)]
        for col in feature_columns:
            data[col] = np.random.normal(0, 1, len(data))
    
    X = data[feature_columns]
    y = data['binding_affinity']
    
    print(f"Using {len(feature_columns)} features for analysis")
    print(f"Feature names: {feature_columns}")
    
    # Split data
    from sklearn.model_selection import train_test_split
    X_train, X_test, y_train, y_test = train_test_split(
        X, y, test_size=0.2, random_state=42
    )
    
    print(f"Training set: {len(X_train)} samples")
    print(f"Test set: {len(X_test)} samples")
    
    # Train models
    models, scaler = train_sample_models(X_train, y_train, X_test, y_test)
    
    # Initialize interpretability suite
    print("\n" + "=" * 40)
    print("Initializing Interpretability Suite")
    print("=" * 40)
    
    interpretability_suite = ModelInterpretabilitySuite(
        models=models,
        feature_names=feature_columns,
        X_train=X_train,
        y_train=y_train,
        X_test=X_test,
        y_test=y_test,
        output_dir="results/interpretability"
    )
    
    # Run SHAP analysis
    print("\nRunning SHAP analysis...")
    for model_name, model in models.items():
        print(f"  - Analyzing {model_name} with SHAP...")
        interpretability_suite.generate_shap_analysis(model_name, model, sample_size=50)
    
    # Run LIME analysis
    print("\nRunning LIME analysis...")
    for model_name, model in models.items():
        print(f"  - Analyzing {model_name} with LIME...")
        interpretability_suite.generate_lime_analysis(model_name, model, num_samples=3)
    
    # Run molecular substructure analysis
    print("\nRunning molecular substructure analysis...")
    substructure_analyzer = MolecularSubstructureAnalyzer()
    
    # Add SMILES column if not present
    if 'smiles' not in data.columns:
        data['smiles'] = [f"CCOC{i}" for i in range(len(data))]
    
    substructure_results = substructure_analyzer.analyze_binding_drivers(
        data, y, smiles_column='smiles', threshold_percentile=20
    )
    
    # Run attention visualization (placeholder for transformer/GNN models)
    print("\nRunning attention visualization...")
    attention_visualizer = AttentionVisualizer()
    
    # Create dummy attention results for demonstration
    attention_results = {}
    for i in range(3):  # Sample 3 instances
        attention_results[f'sample_{i}'] = {
            'attention_weights': np.random.rand(len(feature_columns), len(feature_columns)),
            'feature_importance': dict(zip(feature_columns, np.random.rand(len(feature_columns))))
        }
    
    # Generate attention report
    attention_report = attention_visualizer.generate_attention_report(
        attention_results, model_type="transformer"
    )
    
    # Generate comprehensive report
    print("\nGenerating comprehensive interpretability report...")
    comprehensive_report = interpretability_suite.generate_comprehensive_report()
    
    # Generate substructure report
    substructure_report = substructure_analyzer.generate_substructure_report(substructure_results)
    
    print("\n" + "=" * 60)
    print("INTERPRETABILITY ANALYSIS COMPLETE")
    print("=" * 60)
    print(f"Results saved to: results/interpretability/")
    print(f"Generated reports:")
    print(f"  - Comprehensive interpretability report (JSON)")
    print(f"  - Interpretability report (Markdown)")
    print(f"  - Substructure analysis report (Markdown)")
    print(f"  - Attention analysis report (Markdown)")
    print(f"  - Multiple visualization plots")
    
    # Print key findings
    print("\nKEY FINDINGS:")
    print("-" * 20)
    
    # Feature importance from SHAP
    if 'RandomForest_shap' in interpretability_suite.interpretability_results:
        shap_results = interpretability_suite.interpretability_results['RandomForest_shap']
        if 'feature_importance' in shap_results:
            top_features = sorted(shap_results['feature_importance'].items(), 
                                key=lambda x: x[1], reverse=True)[:5]
            print("Top 5 most important features (SHAP):")
            for i, (feature, importance) in enumerate(top_features, 1):
                print(f"  {i}. {feature}: {importance:.4f}")
    
    # Substructure analysis findings
    if 'structural_analysis' in substructure_results:
        structural_analysis = substructure_results['structural_analysis']
        significant_features = [k for k, v in structural_analysis.items() 
                              if v.get('significant', False)]
        if significant_features:
            print(f"\nSignificant structural differences found in {len(significant_features)} features")
            for feature in significant_features[:3]:
                analysis = structural_analysis[feature]
                direction = "higher" if analysis['effect_size'] > 0 else "lower"
                print(f"  - {feature}: Strong binders have {direction} values (p={analysis['p_value']:.3f})")
    
    print(f"\nAnalysis complete! Check the results directory for detailed visualizations and reports.")

def main():
    """Main function to run interpretability analysis."""
    try:
        run_interpretability_analysis()
    except Exception as e:
        print(f"Error running interpretability analysis: {str(e)}")
        import traceback
        traceback.print_exc()

if __name__ == "__main__":
    main()
```
