## Supplementary material for "Machine Learning-Driven Prediction of TLR4 Binding Affinity: A Comprehensive Molecular Feature Analysis for Drug Discovery": Codebase: demo_interpretability_simple.py

```
#!/usr/bin/env python3
"""
Simple demo script for Model Interpretability and Analysis Suite

This script demonstrates the interpretability tools without requiring
heavy dependencies like PyTorch.
"""

import sys
import os
import pandas as pd
import numpy as np
import matplotlib.pyplot as plt
import warnings
warnings.filterwarnings('ignore')

# Add src to path
sys.path.append('src')

# Import only the interpretability modules we need
from tlr4_binding.ml_components.interpretability import ModelInterpretabilitySuite
from tlr4_binding.ml_components.molecular_substructure_analyzer import MolecularSubstructureAnalyzer
from tlr4_binding.ml_components.attention_visualizer import AttentionVisualizer

def create_sample_data():
    """Create sample data for interpretability analysis."""
    print("Creating sample data for interpretability analysis...")
    
    # Create sample binding data
    n_samples = 100
    binding_affinities = np.random.normal(-6.0, 2.0, n_samples)  # Strong binders have lower (more negative) values
    
    # Create sample molecular features
    molecular_features = pd.DataFrame({
        'mol_weight': np.random.normal(300, 50, n_samples),
        'mol_logp': np.random.normal(2.5, 1.0, n_samples),
        'mol_tpsa': np.random.normal(80, 20, n_samples),
        'mol_hbd': np.random.poisson(3, n_samples),
        'mol_hba': np.random.poisson(5, n_samples),
        'mol_rotatable_bonds': np.random.poisson(8, n_samples),
        'mol_aromatic_rings': np.random.poisson(2, n_samples),
        'mol_heavy_atoms': np.random.poisson(25, n_samples),
        'mol_formal_charge': np.random.choice([-1, 0, 1], n_samples),
        'smiles': [f"CCOC{i}" for i in range(n_samples)]  # Dummy SMILES
    })
    
    # Combine data
    combined_data = pd.concat([pd.DataFrame({'binding_affinity': binding_affinities}), molecular_features], axis=1)
    
    return combined_data

def train_sample_models(X_train, y_train, X_test, y_test):
    """Train sample models for interpretability analysis."""
    print("Training sample models for interpretability analysis...")
    
    from sklearn.ensemble import RandomForestRegressor
    from sklearn.svm import SVR
    from sklearn.linear_model import LinearRegression
    from sklearn.preprocessing import StandardScaler
    
    # Scale features
    scaler = StandardScaler()
    X_train_scaled = scaler.fit_transform(X_train)
    X_test_scaled = scaler.transform(X_test)
    
    # Train models
    models = {}
    
    # Random Forest
    rf_model = RandomForestRegressor(n_estimators=100, random_state=42)
    rf_model.fit(X_train, y_train)
    models['RandomForest'] = rf_model
    
    # Support Vector Regression
    svr_model = SVR(kernel='rbf', C=1.0, gamma='scale')
    svr_model.fit(X_train_scaled, y_train)
    models['SVR'] = svr_model
    
    # Linear Regression
    lr_model = LinearRegression()
    lr_model.fit(X_train_scaled, y_train)
    models['LinearRegression'] = lr_model
    
    print(f"Trained {len(models)} models for interpretability analysis")
    return models, scaler

def run_interpretability_analysis():
    """Run comprehensive interpretability analysis."""
    print("=" * 60)
    print("TLR4 Binding Prediction - Model Interpretability Analysis")
    print("=" * 60)
    
    # Create sample data
    data = create_sample_data()
    print(f"Created sample data with {len(data)} samples")
    
    # Prepare features and target
    feature_columns = [col for col in data.columns if col.startswith('mol_') and col != 'smiles']
    X = data[feature_columns]
    y = data['binding_affinity']
    
    print(f"Using {len(feature_columns)} features for analysis")
    print(f"Feature names: {feature_columns}")
    
    # Split data
    from sklearn.model_selection import train_test_split
    X_train, X_test, y_train, y_test = train_test_split(
        X, y, test_size=0.2, random_state=42
    )
    
    print(f"Training set: {len(X_train)} samples")
    print(f"Test set: {len(X_test)} samples")
    
    # Train models
    models, scaler = train_sample_models(X_train, y_train, X_test, y_test)
    
    # Initialize interpretability suite
    print("\n" + "=" * 40)
    print("Initializing Interpretability Suite")
    print("=" * 40)
    
    interpretability_suite = ModelInterpretabilitySuite(
        models=models,
        feature_names=feature_columns,
        X_train=X_train,
        y_train=y_train,
        X_test=X_test,
        y_test=y_test,
        output_dir="results/interpretability"
    )
    
    # Run SHAP analysis
    print("\nRunning SHAP analysis...")
    for model_name, model in models.items():
        print(f"  - Analyzing {model_name} with SHAP...")
        try:
            interpretability_suite.generate_shap_analysis(model_name, model, sample_size=50)
            print(f"    ✓ SHAP analysis completed for {model_name}")
        except Exception as e:
            print(f"    ✗ SHAP analysis failed for {model_name}: {str(e)}")
    
    # Run LIME analysis
    print("\nRunning LIME analysis...")
    for model_name, model in models.items():
        print(f"  - Analyzing {model_name} with LIME...")
        try:
            interpretability_suite.generate_lime_analysis(model_name, model, num_samples=3)
            print(f"    ✓ LIME analysis completed for {model_name}")
        except Exception as e:
            print(f"    ✗ LIME analysis failed for {model_name}: {str(e)}")
    
    # Run molecular substructure analysis
    print("\nRunning molecular substructure analysis...")
    try:
        substructure_analyzer = MolecularSubstructureAnalyzer()
        substructure_results = substructure_analyzer.analyze_binding_drivers(
            data, y, smiles_column='smiles', threshold_percentile=20
        )
        print("    ✓ Molecular substructure analysis completed")
    except Exception as e:
        print(f"    ✗ Molecular substructure analysis failed: {str(e)}")
        substructure_results = {}
    
    # Run attention visualization (placeholder)
    print("\nRunning attention visualization...")
    try:
        attention_visualizer = AttentionVisualizer()
        
        # Create dummy attention results for demonstration
        attention_results = {}
        for i in range(3):  # Sample 3 instances
            attention_results[f'sample_{i}'] = {
                'attention_weights': np.random.rand(len(feature_columns), len(feature_columns)),
                'feature_importance': dict(zip(feature_columns, np.random.rand(len(feature_columns))))
            }
        
        # Generate attention report
        attention_report = attention_visualizer.generate_attention_report(
            attention_results, model_type="transformer"
        )
        print("    ✓ Attention visualization completed")
    except Exception as e:
        print(f"    ✗ Attention visualization failed: {str(e)}")
    
    # Generate comprehensive report
    print("\nGenerating comprehensive interpretability report...")
    try:
        comprehensive_report = interpretability_suite.generate_comprehensive_report()
        print("    ✓ Comprehensive report generated")
    except Exception as e:
        print(f"    ✗ Comprehensive report generation failed: {str(e)}")
    
    # Generate substructure report
    if substructure_results:
        try:
            substructure_report = substructure_analyzer.generate_substructure_report(substructure_results)
            print("    ✓ Substructure report generated")
        except Exception as e:
            print(f"    ✗ Substructure report generation failed: {str(e)}")
    
    print("\n" + "=" * 60)
    print("INTERPRETABILITY ANALYSIS COMPLETE")
    print("=" * 60)
    print(f"Results saved to: results/interpretability/")
    print(f"Generated reports:")
    print(f"  - Comprehensive interpretability report (JSON)")
    print(f"  - Interpretability report (Markdown)")
    print(f"  - Substructure analysis report (Markdown)")
    print(f"  - Attention analysis report (Markdown)")
    print(f"  - Multiple visualization plots")
    
    # Print key findings
    print("\nKEY FINDINGS:")
    print("-" * 20)
    
    # Feature importance from SHAP
    if hasattr(interpretability_suite, 'interpretability_results'):
        if 'RandomForest_shap' in interpretability_suite.interpretability_results:
            shap_results = interpretability_suite.interpretability_results['RandomForest_shap']
            if 'feature_importance' in shap_results:
                top_features = sorted(shap_results['feature_importance'].items(), 
                                    key=lambda x: x[1], reverse=True)[:5]
                print("Top 5 most important features (SHAP):")
                for i, (feature, importance) in enumerate(top_features, 1):
                    print(f"  {i}. {feature}: {importance:.4f}")
    
    # Substructure analysis findings
    if substructure_results and 'structural_analysis' in substructure_results:
        structural_analysis = substructure_results['structural_analysis']
        significant_features = [k for k, v in structural_analysis.items() 
                              if v.get('significant', False)]
        if significant_features:
            print(f"\nSignificant structural differences found in {len(significant_features)} features")
            for feature in significant_features[:3]:
                analysis = structural_analysis[feature]
                direction = "higher" if analysis['effect_size'] > 0 else "lower"
                print(f"  - {feature}: Strong binders have {direction} values (p={analysis['p_value']:.3f})")
    
    print(f"\nAnalysis complete! Check the results directory for detailed visualizations and reports.")

def main():
    """Main function to run interpretability analysis."""
    try:
        run_interpretability_analysis()
    except Exception as e:
        print(f"Error running interpretability analysis: {str(e)}")
        import traceback
        traceback.print_exc()

if __name__ == "__main__":
    main()
```
