## Supplementary material for "Machine Learning-Driven Prediction of TLR4 Binding Affinity: A Comprehensive Molecular Feature Analysis for Drug Discovery": Codebase: demo_ml_baseline.py

```
#!/usr/bin/env python3
"""
Demo script for traditional ML baseline models.

This script demonstrates the training and evaluation of traditional ML models
for TLR4 binding affinity prediction, including Random Forest, SVR, XGBoost,
and LightGBM with hyperparameter optimization and feature importance analysis.
"""

import sys
import logging
from pathlib import Path
import pandas as pd
import numpy as np
import matplotlib.pyplot as plt
import seaborn as sns
from sklearn.model_selection import train_test_split
from sklearn.preprocessing import StandardScaler
import warnings

# Add src to path
sys.path.insert(0, str(Path(__file__).parent.parent / "src"))

from tlr4_binding.ml_components.trainer import MLModelTrainer
from tlr4_binding.ml_components.feature_engineering import FeatureEngineeringPipeline
from tlr4_binding.data_processing.preprocessor import DataPreprocessor

# Suppress warnings for cleaner output
warnings.filterwarnings('ignore')

# Set up logging
logging.basicConfig(
    level=logging.INFO,
    format='%(asctime)s - %(name)s - %(levelname)s - %(message)s'
)
logger = logging.getLogger(__name__)


def create_synthetic_dataset(n_samples=500, n_features=20, noise_level=0.1):
    """
    Create synthetic dataset for demonstration.
    
    Args:
        n_samples: Number of samples
        n_features: Number of features
        noise_level: Amount of noise to add
        
    Returns:
        Tuple of (X, y) where X is features and y is targets
    """
    np.random.seed(42)
    
    # Create features with some structure
    X = np.random.randn(n_samples, n_features)
    
    # Create target with non-linear relationships
    y = (
        X[:, 0] * 2.0 +                    # Linear term
        X[:, 1] * 1.5 +                    # Linear term
        X[:, 2] ** 2 * 0.5 +               # Quadratic term
        X[:, 3] * X[:, 4] * 0.8 +          # Interaction term
        np.sin(X[:, 5]) * 1.2 +            # Non-linear term
        np.random.randn(n_samples) * noise_level  # Noise
    )
    
    # Create feature names
    feature_names = [f'mol_feature_{i}' for i in range(n_features)]
    X_df = pd.DataFrame(X, columns=feature_names)
    y_series = pd.Series(y, name='binding_affinity')
    
    return X_df, y_series


def plot_model_performance(results, save_path=None):
    """
    Create performance comparison plots.
    
    Args:
        results: Dictionary of model evaluation results
        save_path: Path to save plot (optional)
    """
    # Extract metrics for plotting
    model_names = list(results.keys())
    metrics = ['mse', 'rmse', 'mae', 'r2']
    
    # Create subplots
    fig, axes = plt.subplots(2, 2, figsize=(15, 10))
    fig.suptitle('Traditional ML Models Performance Comparison', fontsize=16)
    
    for i, metric in enumerate(metrics):
        row, col = i // 2, i % 2
        ax = axes[row, col]
        
        values = [results[model]['metrics'][metric] for model in model_names]
        
        bars = ax.bar(model_names, values, alpha=0.7)
        ax.set_title(f'{metric.upper()} Comparison')
        ax.set_ylabel(metric.upper())
        ax.tick_params(axis='x', rotation=45)
        
        # Add value labels on bars
        for bar, value in zip(bars, values):
            height = bar.get_height()
            ax.text(bar.get_x() + bar.get_width()/2., height,
                   f'{value:.3f}', ha='center', va='bottom')
    
    plt.tight_layout()
    
    if save_path:
        plt.savefig(save_path, dpi=300, bbox_inches='tight')
        logger.info(f"Performance plot saved to {save_path}")
    
    plt.show()


def plot_feature_importance(results, feature_names, top_n=10, save_path=None):
    """
    Plot feature importance for tree-based models.
    
    Args:
        results: Dictionary of model evaluation results
        feature_names: List of feature names
        top_n: Number of top features to show
        save_path: Path to save plot (optional)
    """
    # Get tree-based models
    tree_models = ['random_forest', 'xgboost', 'lightgbm']
    
    fig, axes = plt.subplots(1, len(tree_models), figsize=(15, 5))
    if len(tree_models) == 1:
        axes = [axes]
    
    fig.suptitle('Feature Importance Comparison (Tree-based Models)', fontsize=16)
    
    for i, model_name in enumerate(tree_models):
        if model_name in results and results[model_name]['feature_importance']:
            ax = axes[i]
            
            # Get feature importance
            importance = results[model_name]['feature_importance']
            
            # Sort by importance and get top N
            sorted_features = sorted(importance.items(), key=lambda x: x[1], reverse=True)[:top_n]
            features, importances = zip(*sorted_features)
            
            # Create horizontal bar plot
            y_pos = np.arange(len(features))
            bars = ax.barh(y_pos, importances, alpha=0.7)
            ax.set_yticks(y_pos)
            ax.set_yticklabels(features)
            ax.set_xlabel('Importance')
            ax.set_title(f'{model_name.replace("_", " ").title()}')
            ax.invert_yaxis()
            
            # Add value labels
            for bar, value in zip(bars, importances):
                width = bar.get_width()
                ax.text(width, bar.get_y() + bar.get_height()/2.,
                       f'{value:.3f}', ha='left', va='center')
    
    plt.tight_layout()
    
    if save_path:
        plt.savefig(save_path, dpi=300, bbox_inches='tight')
        logger.info(f"Feature importance plot saved to {save_path}")
    
    plt.show()


def plot_predictions_vs_actual(results, y_test, save_path=None):
    """
    Plot predictions vs actual values for all models.
    
    Args:
        results: Dictionary of model evaluation results
        y_test: True test values
        save_path: Path to save plot (optional)
    """
    n_models = len(results)
    n_cols = min(3, n_models)
    n_rows = (n_models + n_cols - 1) // n_cols
    
    fig, axes = plt.subplots(n_rows, n_cols, figsize=(5*n_cols, 5*n_rows))
    if n_models == 1:
        axes = [axes]
    elif n_rows == 1:
        axes = axes.reshape(1, -1)
    
    fig.suptitle('Predictions vs Actual Values', fontsize=16)
    
    for i, (model_name, result) in enumerate(results.items()):
        row, col = i // n_cols, i % n_cols
        ax = axes[row, col] if n_rows > 1 else axes[col]
        
        predictions = result['predictions']
        
        # Create scatter plot
        ax.scatter(y_test, predictions, alpha=0.6)
        
        # Add perfect prediction line
        min_val = min(y_test.min(), predictions.min())
        max_val = max(y_test.max(), predictions.max())
        ax.plot([min_val, max_val], [min_val, max_val], 'r--', alpha=0.8)
        
        # Add R² score
        r2 = result['metrics']['r2']
        ax.text(0.05, 0.95, f'R² = {r2:.3f}', transform=ax.transAxes,
               bbox=dict(boxstyle='round', facecolor='white', alpha=0.8))
        
        ax.set_xlabel('Actual')
        ax.set_ylabel('Predicted')
        ax.set_title(f'{model_name.replace("_", " ").title()}')
    
    # Hide empty subplots
    for i in range(n_models, n_rows * n_cols):
        row, col = i // n_cols, i % n_cols
        ax = axes[row, col] if n_rows > 1 else axes[col]
        ax.set_visible(False)
    
    plt.tight_layout()
    
    if save_path:
        plt.savefig(save_path, dpi=300, bbox_inches='tight')
        logger.info(f"Predictions plot saved to {save_path}")
    
    plt.show()


def main():
    """Main demonstration function."""
    logger.info("Starting Traditional ML Baseline Demo")
    
    # Create synthetic dataset
    logger.info("Creating synthetic dataset...")
    X, y = create_synthetic_dataset(n_samples=500, n_features=20)
    
    # Split data
    X_train, X_test, y_train, y_test = train_test_split(
        X, y, test_size=0.2, random_state=42
    )
    
    logger.info(f"Dataset created: {X.shape[0]} samples, {X.shape[1]} features")
    logger.info(f"Train set: {X_train.shape[0]} samples")
    logger.info(f"Test set: {X_test.shape[0]} samples")
    
    # Apply feature scaling
    logger.info("Applying feature scaling...")
    scaler = StandardScaler()
    X_train_scaled = pd.DataFrame(
        scaler.fit_transform(X_train),
        columns=X_train.columns,
        index=X_train.index
    )
    X_test_scaled = pd.DataFrame(
        scaler.transform(X_test),
        columns=X_test.columns,
        index=X_test.index
    )
    
    # Train models
    logger.info("Training traditional ML models...")
    trainer = MLModelTrainer(models_dir="models/demo")
    
    # Train all available models
    trained_models = trainer.train_models(X_train_scaled, y_train)
    logger.info(f"Successfully trained {len(trained_models)} models")
    
    # Evaluate models
    logger.info("Evaluating models on test set...")
    results = trainer.evaluate_models(X_test_scaled, y_test)
    
    # Print performance summary
    logger.info("\n" + "="*60)
    logger.info("MODEL PERFORMANCE SUMMARY")
    logger.info("="*60)
    
    summary_df = trainer.get_model_summary()
    print(summary_df.round(4))
    
    # Get best model
    best_name, best_model = trainer.get_best_model('r2')
    logger.info(f"\nBest model: {best_name} (R² = {summary_df[summary_df['model_name'] == best_name]['r2'].iloc[0]:.4f})")
    
    # Create visualizations
    logger.info("Creating performance visualizations...")
    
    # Set style
    plt.style.use('seaborn-v0_8')
    
    # Performance comparison
    plot_model_performance(results, save_path="results/ml_performance_comparison.png")
    
    # Feature importance (for tree-based models)
    plot_feature_importance(results, X.columns, save_path="results/feature_importance.png")
    
    # Predictions vs actual
    plot_predictions_vs_actual(results, y_test, save_path="results/predictions_vs_actual.png")
    
    # Detailed analysis for best model
    logger.info(f"\nDetailed analysis for best model: {best_name}")
    best_result = results[best_name]
    
    logger.info(f"R² Score: {best_result['metrics']['r2']:.4f}")
    logger.info(f"RMSE: {best_result['metrics']['rmse']:.4f}")
    logger.info(f"MAE: {best_result['metrics']['mae']:.4f}")
    
    if best_result['feature_importance']:
        logger.info("\nTop 5 Most Important Features:")
        sorted_features = sorted(
            best_result['feature_importance'].items(),
            key=lambda x: x[1], reverse=True
        )[:5]
        
        for feature, importance in sorted_features:
            logger.info(f"  {feature}: {importance:.4f}")
    
    # Save results
    results_dir = Path("results")
    results_dir.mkdir(exist_ok=True)
    
    # Save model summary
    summary_df.to_csv(results_dir / "model_performance_summary.csv", index=False)
    logger.info(f"Model summary saved to {results_dir / 'model_performance_summary.csv'}")
    
    # Save detailed results
    detailed_results = {}
    for model_name, result in results.items():
        detailed_results[model_name] = {
            'metrics': result['metrics'],
            'feature_importance': result['feature_importance']
        }
    
    import json
    with open(results_dir / "detailed_results.json", 'w') as f:
        json.dump(detailed_results, f, indent=2, default=str)
    logger.info(f"Detailed results saved to {results_dir / 'detailed_results.json'}")
    
    logger.info("\n" + "="*60)
    logger.info("DEMO COMPLETED SUCCESSFULLY")
    logger.info("="*60)
    logger.info("Check the 'results/' directory for saved plots and data")
    logger.info("All models have been trained and evaluated with hyperparameter optimization")


if __name__ == "__main__":
    main()
```
