## Supplementary material for "Machine Learning-Driven Prediction of TLR4 Binding Affinity: A Comprehensive Molecular Feature Analysis for Drug Discovery": Codebase: demo_pipeline_orchestrator.py

```
#!/usr/bin/env python3
"""
Demo script for Research Pipeline Orchestrator

This script demonstrates the complete research pipeline orchestrator
with MLflow integration, hyperparameter optimization, and automated reporting.
"""

import sys
import os
import numpy as np
import pandas as pd
from pathlib import Path

# Add src to path
sys.path.append(str(Path(__file__).parent.parent / "src"))

from tlr4_binding.ml_components.pipeline_orchestrator import (
    ResearchPipelineOrchestrator, 
    create_experiment_config
)
from tlr4_binding.data_processing.data_loader import DataLoader
from tlr4_binding.molecular_analysis.feature_extractor import MolecularFeatureExtractor

def create_sample_data(n_samples: int = 100, n_features: int = 20) -> tuple:
    """Create sample data for testing the pipeline orchestrator"""
    print(f"Creating sample dataset with {n_samples} samples and {n_features} features")
    
    # Generate synthetic molecular features
    np.random.seed(42)
    feature_names = [
        'molecular_weight', 'logp', 'tpsa', 'rotatable_bonds', 'hbd', 'hba',
        'radius_of_gyration', 'molecular_volume', 'surface_area', 'asphericity',
        'ring_count', 'aromatic_rings', 'formal_charge', 'polar_surface_area',
        'molecular_flexibility', 'hydrogen_bond_donors', 'hydrogen_bond_acceptors',
        'topological_polar_surface_area', 'molecular_connectivity', 'electrotopological_state'
    ]
    
    # Ensure we have enough feature names
    if n_features > len(feature_names):
        feature_names.extend([f'feature_{i}' for i in range(len(feature_names), n_features)])
    
    feature_names = feature_names[:n_features]
    
    # Generate realistic molecular features
    X = pd.DataFrame(
        np.random.randn(n_samples, n_features),
        columns=feature_names
    )
    
    # Add some realistic constraints
    X['molecular_weight'] = np.random.uniform(100, 800, n_samples)
    X['logp'] = np.random.uniform(-2, 6, n_samples)
    X['tpsa'] = np.random.uniform(0, 200, n_samples)
    X['rotatable_bonds'] = np.random.randint(0, 15, n_samples)
    X['hbd'] = np.random.randint(0, 8, n_samples)
    X['hba'] = np.random.randint(0, 12, n_samples)
    
    # Generate binding affinities with some correlation to features
    # Lower (more negative) values indicate stronger binding
    y = (
        -0.5 * X['molecular_weight'] / 100 +
        -0.3 * X['logp'] +
        -0.2 * X['tpsa'] / 100 +
        -0.1 * X['rotatable_bonds'] +
        np.random.normal(0, 1, n_samples)
    )
    
    # Ensure reasonable binding affinity range (kcal/mol)
    y = np.clip(y, -15, 0)
    
    print(f"Generated dataset: {X.shape[0]} samples, {X.shape[1]} features")
    print(f"Binding affinity range: {y.min():.2f} to {y.max():.2f} kcal/mol")
    
    return X, y

def run_basic_pipeline_demo():
    """Run a basic pipeline demonstration"""
    print("=" * 60)
    print("TLR4 Binding Prediction - Pipeline Orchestrator Demo")
    print("=" * 60)
    
    # Create sample data
    X, y = create_sample_data(n_samples=200, n_features=15)
    
    # Create experiment configuration
    config = create_experiment_config(
        experiment_name="tlr4_pipeline_demo",
        description="Demonstration of TLR4 binding prediction pipeline orchestrator",
        data_path="./data",
        output_path="./results/pipeline_demo",
        n_trials=20,  # Reduced for demo
        cv_folds=3,   # Reduced for demo
        models_to_test=['random_forest', 'xgboost', 'neural_network']
    )
    
    print(f"\nExperiment Configuration:")
    print(f"- Name: {config.experiment_name}")
    print(f"- Description: {config.description}")
    print(f"- Models: {config.models_to_test}")
    print(f"- CV Folds: {config.cv_folds}")
    print(f"- Optimization Trials: {config.n_trials}")
    
    # Create orchestrator
    print(f"\nInitializing pipeline orchestrator...")
    orchestrator = ResearchPipelineOrchestrator(config)
    
    # Save configuration
    config_path = "./results/pipeline_demo/experiment_config.yaml"
    os.makedirs(os.path.dirname(config_path), exist_ok=True)
    orchestrator.save_config(config_path)
    print(f"Configuration saved to: {config_path}")
    
    # Run the complete pipeline
    print(f"\nRunning complete research pipeline...")
    try:
        results = orchestrator.run_complete_pipeline(X, y)
        
        print(f"\nPipeline completed successfully!")
        print(f"Results keys: {list(results.keys())}")
        
        # Display model performance
        if 'model_evaluation' in results:
            print(f"\nModel Performance Summary:")
            print("-" * 40)
            for model_name, result in results['model_evaluation'].items():
                if 'error' not in result:
                    metrics = result['metrics']
                    print(f"{model_name:15} | R²: {metrics['r2_score']:6.3f} | RMSE: {metrics['rmse']:6.3f}")
                else:
                    print(f"{model_name:15} | ERROR: {result['error']}")
        
        # Display nested CV results
        if 'nested_cv' in results:
            print(f"\nNested Cross-Validation Results:")
            print("-" * 40)
            for model_name, scores in results['nested_cv'].items():
                if 'error' not in scores:
                    print(f"{model_name:15} | {scores['mean_score']:6.3f} ± {scores['std_score']:6.3f}")
                else:
                    print(f"{model_name:15} | ERROR: {scores['error']}")
        
        print(f"\nResearch report generated at: ./results/pipeline_demo/research_report.md")
        
    except Exception as e:
        print(f"Pipeline failed: {e}")
        import traceback
        traceback.print_exc()

def run_mlflow_demo():
    """Demonstrate MLflow integration"""
    print("\n" + "=" * 60)
    print("MLflow Integration Demo")
    print("=" * 60)
    
    import mlflow
    
    # Check MLflow setup
    print(f"MLflow tracking URI: {mlflow.get_tracking_uri()}")
    
    # List experiments
    try:
        experiments = mlflow.search_experiments()
        print(f"\nFound {len(experiments)} experiments:")
        for exp in experiments:
            print(f"- {exp.name} (ID: {exp.experiment_id})")
    except Exception as e:
        print(f"Could not list experiments: {e}")

def run_optuna_demo():
    """Demonstrate Optuna integration"""
    print("\n" + "=" * 60)
    print("Optuna Integration Demo")
    print("=" * 60)
    
    try:
        import optuna
        print(f"Optuna version: {optuna.__version__}")
        
        # Create a simple study
        def objective(trial):
            x = trial.suggest_float('x', -10, 10)
            return (x - 2) ** 2
        
        study = optuna.create_study(direction='minimize')
        study.optimize(objective, n_trials=10)
        
        print(f"Best value: {study.best_value}")
        print(f"Best params: {study.best_params}")
        
    except Exception as e:
        print(f"Optuna demo failed: {e}")

def main():
    """Main demo function"""
    print("Starting TLR4 Binding Prediction Pipeline Orchestrator Demo")
    
    # Create output directory
    os.makedirs("./results/pipeline_demo", exist_ok=True)
    
    try:
        # Run basic pipeline demo
        run_basic_pipeline_demo()
        
        # Run MLflow demo
        run_mlflow_demo()
        
        # Run Optuna demo
        run_optuna_demo()
        
        print("\n" + "=" * 60)
        print("Demo completed successfully!")
        print("=" * 60)
        print("\nNext steps:")
        print("1. Check MLflow UI: mlflow ui --backend-store-uri file:./mlruns")
        print("2. View research report: ./results/pipeline_demo/research_report.md")
        print("3. Check experiment config: ./results/pipeline_demo/experiment_config.yaml")
        
    except Exception as e:
        print(f"Demo failed: {e}")
        import traceback
        traceback.print_exc()

if __name__ == "__main__":
    main()
```
