## Supplementary material for "Machine Learning-Driven Prediction of TLR4 Binding Affinity: A Comprehensive Molecular Feature Analysis for Drug Discovery": Codebase: demo_uncertainty_quantification.py

```
#!/usr/bin/env python3
"""
Demo script for uncertainty quantification methods in TLR4 binding prediction.

This script demonstrates various uncertainty quantification techniques including:
- Monte Carlo Dropout for neural networks
- Bootstrap aggregating for traditional ML models
- Conformal prediction intervals
- Ensemble-based uncertainty quantification
- Calibration analysis and reliability diagrams
"""

import sys
import os
import numpy as np
import pandas as pd
import torch
import torch.nn as nn
from sklearn.ensemble import RandomForestRegressor, GradientBoostingRegressor
from sklearn.model_selection import train_test_split
from sklearn.preprocessing import StandardScaler
import matplotlib.pyplot as plt
import seaborn as sns
from pathlib import Path

# Add src to path
sys.path.append(os.path.join(os.path.dirname(__file__), 'src'))

from tlr4_binding.ml_components.uncertainty_quantification import (
    MonteCarloDropout,
    BootstrapUncertainty,
    ConformalPrediction,
    EnsembleUncertainty,
    UncertaintyQuantifier,
    UncertaintyCalibration
)
from tlr4_binding.data_processing.data_loader import BindingDataLoader
from tlr4_binding.molecular_analysis.feature_extractor import MolecularFeatureExtractor
from tlr4_binding.ml_components.feature_engineering import FeatureScaler
from tlr4_binding.ml_components.trainer import MLTrainer
from tlr4_binding.ml_components.deep_learning_models import DeepLearningModel
from tlr4_binding.ml_components.ensemble_models import EnsembleModel

# Set up logging
import logging
logging.basicConfig(level=logging.INFO)
logger = logging.getLogger(__name__)

# Set random seeds for reproducibility
np.random.seed(42)
torch.manual_seed(42)


class SimpleNeuralNetwork(nn.Module):
    """Simple neural network for demonstration purposes."""
    
    def __init__(self, input_size, hidden_sizes=[64, 32], dropout_rate=0.2):
        super().__init__()
        
        layers = []
        prev_size = input_size
        
        for hidden_size in hidden_sizes:
            layers.extend([
                nn.Linear(prev_size, hidden_size),
                nn.ReLU(),
                nn.Dropout(dropout_rate)
            ])
            prev_size = hidden_size
            
        layers.append(nn.Linear(prev_size, 1))
        
        self.network = nn.Sequential(*layers)
        
    def forward(self, x):
        return self.network(x).squeeze()


def load_sample_data():
    """Load sample data for demonstration."""
    logger.info("Loading sample data...")
    
    # Try to load real data if available
    try:
        data_loader = BindingDataLoader()
        binding_data = data_loader.load_binding_data()
        
        # Use a subset for demonstration
        if len(binding_data) > 1000:
            binding_data = binding_data.sample(n=1000, random_state=42)
            
        logger.info(f"Loaded {len(binding_data)} samples from real data")
        return binding_data
        
    except Exception as e:
        logger.warning(f"Could not load real data: {e}")
        logger.info("Generating synthetic data for demonstration...")
        
        # Generate synthetic data
        np.random.seed(42)
        n_samples = 500
        n_features = 20
        
        # Generate features
        X = np.random.randn(n_samples, n_features)
        
        # Generate target with some noise
        true_coeffs = np.random.randn(n_features)
        y = X @ true_coeffs + 0.1 * np.random.randn(n_samples)
        
        # Create DataFrame
        feature_names = [f'feature_{i}' for i in range(n_features)]
        data = pd.DataFrame(X, columns=feature_names)
        data['binding_affinity'] = y
        
        return data


def prepare_data(data):
    """Prepare data for training and testing."""
    logger.info("Preparing data...")
    
    # Separate features and target
    feature_cols = [col for col in data.columns if col != 'binding_affinity']
    X = data[feature_cols].values
    y = data['binding_affinity'].values
    
    # Split data
    X_train, X_test, y_train, y_test = train_test_split(
        X, y, test_size=0.2, random_state=42
    )
    
    # Scale features
    scaler = StandardScaler()
    X_train_scaled = scaler.fit_transform(X_train)
    X_test_scaled = scaler.transform(X_test)
    
    return X_train_scaled, X_test_scaled, y_train, y_test, scaler


def demo_monte_carlo_dropout(X_train, X_test, y_train, y_test):
    """Demonstrate Monte Carlo Dropout uncertainty quantification."""
    logger.info("Demonstrating Monte Carlo Dropout...")
    
    # Create and train neural network
    input_size = X_train.shape[1]
    model = SimpleNeuralNetwork(input_size, hidden_sizes=[64, 32], dropout_rate=0.2)
    
    # Convert to tensors
    X_train_tensor = torch.FloatTensor(X_train)
    y_train_tensor = torch.FloatTensor(y_train)
    X_test_tensor = torch.FloatTensor(X_test)
    
    # Train model
    criterion = nn.MSELoss()
    optimizer = torch.optim.Adam(model.parameters(), lr=0.001)
    
    model.train()
    for epoch in range(100):
        optimizer.zero_grad()
        outputs = model(X_train_tensor)
        loss = criterion(outputs, y_train_tensor)
        loss.backward()
        optimizer.step()
        
        if (epoch + 1) % 20 == 0:
            logger.info(f"Epoch {epoch + 1}, Loss: {loss.item():.4f}")
    
    # Monte Carlo Dropout uncertainty quantification
    mc_dropout = MonteCarloDropout(model, n_samples=100, dropout_rate=0.2)
    uncertainty_result = mc_dropout.predict_with_uncertainty(X_test_tensor)
    
    # Calculate metrics
    mse = np.mean((y_test - uncertainty_result.predictions) ** 2)
    mae = np.mean(np.abs(y_test - uncertainty_result.predictions))
    
    logger.info(f"MC Dropout - MSE: {mse:.4f}, MAE: {mae:.4f}")
    logger.info(f"MC Dropout - Mean Uncertainty: {np.mean(uncertainty_result.uncertainties):.4f}")
    
    return uncertainty_result, model


def demo_bootstrap_uncertainty(X_train, X_test, y_train, y_test):
    """Demonstrate Bootstrap uncertainty quantification."""
    logger.info("Demonstrating Bootstrap Uncertainty...")
    
    # Bootstrap uncertainty with Random Forest
    bootstrap_rf = BootstrapUncertainty(
        RandomForestRegressor,
        n_bootstrap=50,
        random_state=42,
        n_estimators=100
    )
    
    bootstrap_rf.fit(X_train, y_train)
    uncertainty_result = bootstrap_rf.predict_with_uncertainty(X_test)
    
    # Calculate metrics
    mse = np.mean((y_test - uncertainty_result.predictions) ** 2)
    mae = np.mean(np.abs(y_test - uncertainty_result.predictions))
    
    logger.info(f"Bootstrap RF - MSE: {mse:.4f}, MAE: {mae:.4f}")
    logger.info(f"Bootstrap RF - Mean Uncertainty: {np.mean(uncertainty_result.uncertainties):.4f}")
    
    return uncertainty_result


def demo_conformal_prediction(X_train, X_test, y_train, y_test):
    """Demonstrate Conformal Prediction uncertainty quantification."""
    logger.info("Demonstrating Conformal Prediction...")
    
    # Train base model
    base_model = GradientBoostingRegressor(n_estimators=100, random_state=42)
    base_model.fit(X_train, y_train)
    
    # Conformal prediction
    conformal_pred = ConformalPrediction(base_model, alpha=0.05, method='quantile')
    conformal_pred.fit(X_train, y_train)
    uncertainty_result = conformal_pred.predict_with_uncertainty(X_test)
    
    # Calculate metrics
    mse = np.mean((y_test - uncertainty_result.predictions) ** 2)
    mae = np.mean(np.abs(y_test - uncertainty_result.predictions))
    
    logger.info(f"Conformal Prediction - MSE: {mse:.4f}, MAE: {mae:.4f}")
    logger.info(f"Conformal Prediction - Mean Uncertainty: {np.mean(uncertainty_result.uncertainties):.4f}")
    
    return uncertainty_result


def demo_ensemble_uncertainty(X_train, X_test, y_train, y_test):
    """Demonstrate Ensemble uncertainty quantification."""
    logger.info("Demonstrating Ensemble Uncertainty...")
    
    # Train multiple models
    models = []
    
    # Random Forest
    rf = RandomForestRegressor(n_estimators=100, random_state=42)
    rf.fit(X_train, y_train)
    models.append(rf)
    
    # Gradient Boosting
    gb = GradientBoostingRegressor(n_estimators=100, random_state=43)
    gb.fit(X_train, y_train)
    models.append(gb)
    
    # Another Random Forest with different parameters
    rf2 = RandomForestRegressor(n_estimators=50, max_depth=10, random_state=44)
    rf2.fit(X_train, y_train)
    models.append(rf2)
    
    # Ensemble uncertainty
    ensemble_uncertainty = EnsembleUncertainty(models)
    uncertainty_result = ensemble_uncertainty.predict_with_uncertainty(X_test)
    
    # Calculate metrics
    mse = np.mean((y_test - uncertainty_result.predictions) ** 2)
    mae = np.mean(np.abs(y_test - uncertainty_result.predictions))
    
    logger.info(f"Ensemble - MSE: {mse:.4f}, MAE: {mae:.4f}")
    logger.info(f"Ensemble - Mean Uncertainty: {np.mean(uncertainty_result.uncertainties):.4f}")
    
    return uncertainty_result


def plot_uncertainty_comparison(results_dict, y_test, save_dir="results/uncertainty"):
    """Plot comparison of different uncertainty quantification methods."""
    logger.info("Creating uncertainty comparison plots...")
    
    # Create save directory
    Path(save_dir).mkdir(parents=True, exist_ok=True)
    
    # Set up the plot
    fig, axes = plt.subplots(2, 2, figsize=(15, 12))
    axes = axes.flatten()
    
    colors = ['blue', 'red', 'green', 'orange']
    
    for i, (method, result) in enumerate(results_dict.items()):
        ax = axes[i]
        
        # Plot predictions vs actual
        ax.scatter(y_test, result.predictions, alpha=0.6, color=colors[i])
        
        # Plot uncertainty bars
        ax.errorbar(y_test, result.predictions, 
                   yerr=1.96 * np.sqrt(result.uncertainties),
                   fmt='none', alpha=0.3, color=colors[i])
        
        # Plot perfect prediction line
        min_val = min(y_test.min(), result.predictions.min())
        max_val = max(y_test.max(), result.predictions.max())
        ax.plot([min_val, max_val], [min_val, max_val], 'k--', alpha=0.5)
        
        ax.set_xlabel('True Values')
        ax.set_ylabel('Predicted Values')
        ax.set_title(f'{method} - Predictions vs Actual')
        ax.grid(True, alpha=0.3)
        
        # Calculate R²
        r2 = 1 - np.sum((y_test - result.predictions) ** 2) / np.sum((y_test - np.mean(y_test)) ** 2)
        ax.text(0.05, 0.95, f'R² = {r2:.3f}', transform=ax.transAxes, 
                bbox=dict(boxstyle='round', facecolor='white', alpha=0.8))
    
    plt.tight_layout()
    plt.savefig(f"{save_dir}/uncertainty_comparison.png", dpi=300, bbox_inches='tight')
    plt.show()
    
    # Plot uncertainty distributions
    fig, ax = plt.subplots(figsize=(10, 6))
    
    for i, (method, result) in enumerate(results_dict.items()):
        ax.hist(result.uncertainties, bins=30, alpha=0.6, 
               label=method, color=colors[i])
    
    ax.set_xlabel('Uncertainty')
    ax.set_ylabel('Frequency')
    ax.set_title('Uncertainty Distribution Comparison')
    ax.legend()
    ax.grid(True, alpha=0.3)
    
    plt.tight_layout()
    plt.savefig(f"{save_dir}/uncertainty_distributions.png", dpi=300, bbox_inches='tight')
    plt.show()


def demo_calibration_analysis(results_dict, y_test, save_dir="results/uncertainty"):
    """Demonstrate uncertainty calibration analysis."""
    logger.info("Demonstrating calibration analysis...")
    
    calibrator = UncertaintyCalibration()
    
    for method, result in results_dict.items():
        logger.info(f"\nCalibration analysis for {method}:")
        
        # Calculate calibration metrics
        metrics = calibrator.calculate_calibration_metrics(
            y_test, result.predictions, result.uncertainties
        )
        
        for metric, value in metrics.items():
            logger.info(f"  {metric}: {value:.4f}")
        
        # Plot reliability diagram
        fig = calibrator.plot_reliability_diagram(
            y_test, result.predictions, result.uncertainties
        )
        fig.suptitle(f'{method} - Reliability Diagram')
        plt.savefig(f"{save_dir}/{method.lower().replace(' ', '_')}_reliability.png", 
                   dpi=300, bbox_inches='tight')
        plt.show()


def main():
    """Main demonstration function."""
    logger.info("Starting Uncertainty Quantification Demo")
    
    # Load data
    data = load_sample_data()
    
    # Prepare data
    X_train, X_test, y_train, y_test, scaler = prepare_data(data)
    
    logger.info(f"Training set size: {X_train.shape[0]}")
    logger.info(f"Test set size: {X_test.shape[0]}")
    logger.info(f"Number of features: {X_train.shape[1]}")
    
    # Run different uncertainty quantification methods
    results = {}
    
    # Monte Carlo Dropout
    try:
        mc_result, nn_model = demo_monte_carlo_dropout(X_train, X_test, y_train, y_test)
        results['Monte Carlo Dropout'] = mc_result
    except Exception as e:
        logger.error(f"Monte Carlo Dropout failed: {e}")
    
    # Bootstrap Uncertainty
    try:
        bootstrap_result = demo_bootstrap_uncertainty(X_train, X_test, y_train, y_test)
        results['Bootstrap RF'] = bootstrap_result
    except Exception as e:
        logger.error(f"Bootstrap Uncertainty failed: {e}")
    
    # Conformal Prediction
    try:
        conformal_result = demo_conformal_prediction(X_train, X_test, y_train, y_test)
        results['Conformal Prediction'] = conformal_result
    except Exception as e:
        logger.error(f"Conformal Prediction failed: {e}")
    
    # Ensemble Uncertainty
    try:
        ensemble_result = demo_ensemble_uncertainty(X_train, X_test, y_train, y_test)
        results['Ensemble'] = ensemble_result
    except Exception as e:
        logger.error(f"Ensemble Uncertainty failed: {e}")
    
    # Create comparison plots
    if results:
        plot_uncertainty_comparison(results, y_test)
        demo_calibration_analysis(results, y_test)
        
        # Summary statistics
        logger.info("\n" + "="*50)
        logger.info("UNCERTAINTY QUANTIFICATION SUMMARY")
        logger.info("="*50)
        
        for method, result in results.items():
            mse = np.mean((y_test - result.predictions) ** 2)
            mae = np.mean(np.abs(y_test - result.predictions))
            mean_uncertainty = np.mean(result.uncertainties)
            
            logger.info(f"\n{method}:")
            logger.info(f"  MSE: {mse:.4f}")
            logger.info(f"  MAE: {mae:.4f}")
            logger.info(f"  Mean Uncertainty: {mean_uncertainty:.4f}")
            logger.info(f"  Uncertainty Std: {np.std(result.uncertainties):.4f}")
    
    logger.info("\nUncertainty quantification demo completed!")


if __name__ == "__main__":
    main()
```
