## Supplementary material for "Machine Learning-Driven Prediction of TLR4 Binding Affinity: A Comprehensive Molecular Feature Analysis for Drug Discovery": Codebase: feature_engineering_example.py

```
"""
Example usage of the feature engineering pipeline for TLR4 binding prediction.

This script demonstrates how to use the FeatureEngineeringPipeline to preprocess
molecular descriptor data for machine learning model training.
"""

import sys
import os
import numpy as np
import pandas as pd

# Add src to path for imports
sys.path.insert(0, os.path.join(os.path.dirname(__file__), '..', 'src'))

from tlr4_binding.ml_components.feature_engineering import (
    FeatureEngineeringPipeline,
    FeatureEngineeringConfig
)


def create_sample_data(n_samples=200):
    """Create sample molecular descriptor data for demonstration."""
    np.random.seed(42)
    
    # Create base features with different importance levels
    important_feature = np.random.normal(0, 1, n_samples)
    noise_feature = np.random.normal(0, 1, n_samples)
    
    # Create target (binding affinity) that depends on important features
    y = (2 * important_feature + 
         0.1 * noise_feature + 
         np.random.normal(0, 0.1, n_samples))
    
    # Create features with correlations and different importance
    X = pd.DataFrame({
        # Important features
        'molecular_weight': important_feature * 100 + 200,
        'logp': important_feature * 0.5 + 2,
        'tpsa': important_feature * 20 + 50,
        
        # Correlated features (should be removed)
        'mw_correlated': important_feature * 100 + 200 + np.random.normal(0, 0.1, n_samples),
        'logp_correlated': important_feature * 0.5 + 2 + np.random.normal(0, 0.01, n_samples),
        
        # Noise features (less important)
        'noise1': noise_feature,
        'noise2': np.random.normal(0, 1, n_samples),
        'noise3': np.random.normal(0, 1, n_samples),
        'noise4': np.random.normal(0, 1, n_samples),
        
        # Additional molecular descriptors
        'rotatable_bonds': np.random.poisson(5, n_samples),
        'hbd': np.random.poisson(2, n_samples),
        'hba': np.random.poisson(4, n_samples),
        'ring_count': np.random.poisson(2, n_samples),
    })
    
    return X, pd.Series(y, name='binding_affinity')


def main():
    """Demonstrate feature engineering pipeline usage."""
    print("=== Feature Engineering Pipeline Example ===\n")
    
    # Create sample data
    print("1. Creating sample molecular descriptor data...")
    X, y = create_sample_data()
    print(f"   Original data shape: {X.shape}")
    print(f"   Features: {list(X.columns)}")
    print(f"   Target range: [{y.min():.2f}, {y.max():.2f}]\n")
    
    # Configure pipeline
    print("2. Configuring feature engineering pipeline...")
    config = FeatureEngineeringConfig(
        correlation_threshold=0.95,
        mutual_info_k=8,
        use_robust_scaling=False,
        apply_pca=False
    )
    print(f"   Correlation threshold: {config.correlation_threshold}")
    print(f"   Features to select: {config.mutual_info_k}")
    print(f"   Robust scaling: {config.use_robust_scaling}")
    print(f"   Apply PCA: {config.apply_pca}\n")
    
    # Create and fit pipeline
    print("3. Fitting feature engineering pipeline...")
    pipeline = FeatureEngineeringPipeline(config)
    X_transformed = pipeline.fit_transform(X, y)
    print(f"   Transformed data shape: {X_transformed.shape}")
    print(f"   Selected features: {list(X_transformed.columns)}\n")
    
    # Show correlation analysis results
    print("4. Correlation analysis results...")
    corr_analysis = pipeline.get_correlation_analysis()
    print(f"   Highly correlated pairs found: {len(corr_analysis['correlated_pairs'])}")
    for pair in corr_analysis['correlated_pairs']:
        print(f"     {pair['feature1']} <-> {pair['feature2']}: {pair['correlation']:.4f}")
    print(f"   Features removed: {corr_analysis['features_removed']}\n")
    
    # Show feature importance
    print("5. Feature importance scores...")
    importance_df = pipeline.get_feature_importance()
    print("   Top features by mutual information:")
    for _, row in importance_df.head().iterrows():
        print(f"     {row['feature']}: {row['score']:.4f}")
    print()
    
    # Show pipeline summary
    print("6. Pipeline summary...")
    summary = pipeline.get_pipeline_summary()
    print(f"   Total features removed: {summary['total_features_removed']}")
    print(f"   Final features: {len(summary['final_features'])}")
    print(f"   Scaling method: {'Robust' if config.use_robust_scaling else 'Standard'}")
    print()
    
    # Demonstrate transform on new data
    print("7. Transforming new data...")
    X_new, _ = create_sample_data(50)  # Smaller dataset
    X_new_transformed = pipeline.transform(X_new)
    print(f"   New data shape: {X_new.shape} -> {X_new_transformed.shape}")
    print(f"   Features preserved: {list(X_new_transformed.columns)}\n")
    
    print("=== Feature Engineering Complete ===")


if __name__ == "__main__":
    main()
```
