## Supplementary material for "Machine Learning-Driven Prediction of TLR4 Binding Affinity: A Comprehensive Molecular Feature Analysis for Drug Discovery": Codebase: extract_binding_data.py

```
#!/usr/bin/env python3
"""
Extract binding affinity data from AutoDock Vina log files.

This script parses the log files to extract the best binding affinity
for each compound and creates a CSV file for the pipeline.

Author: Kiro AI Assistant
"""

import os
import re
import pandas as pd
from pathlib import Path
import logging

logging.basicConfig(level=logging.INFO)
logger = logging.getLogger(__name__)


def extract_best_affinity_from_log(log_file_path):
    """Extract the best (most negative) binding affinity from a Vina log file."""
    try:
        with open(log_file_path, 'r') as f:
            content = f.read()
        
        # Look for the results table
        # Pattern: mode |   affinity | dist from best mode
        #               | (kcal/mol) | rmsd l.b.| rmsd u.b.
        #          -----+------------+----------+----------
        #             1       -6.798          0          0
        
        pattern = r'^\s*1\s+(-?\d+\.?\d*)\s+0\s+0'
        match = re.search(pattern, content, re.MULTILINE)
        
        if match:
            affinity = float(match.group(1))
            return affinity
        else:
            # Try alternative pattern for the first result
            pattern = r'^\s*1\s+(-?\d+\.?\d*)\s+'
            match = re.search(pattern, content, re.MULTILINE)
            if match:
                affinity = float(match.group(1))
                return affinity
            else:
                logger.warning(f"Could not extract affinity from {log_file_path}")
                return None
                
    except Exception as e:
        logger.error(f"Error reading {log_file_path}: {e}")
        return None


def extract_all_binding_data(logs_dir="data/raw/logs"):
    """Extract binding data from all log files."""
    logs_path = Path(logs_dir)
    
    if not logs_path.exists():
        raise FileNotFoundError(f"Logs directory not found: {logs_dir}")
    
    binding_data = []
    
    # Get all .txt files (excluding summary.csv)
    log_files = [f for f in logs_path.glob("*.txt")]
    
    logger.info(f"Processing {len(log_files)} log files...")
    
    for log_file in log_files:
        compound_name = log_file.stem
        
        # Skip if it's a configuration file (we want the base compound)
        if '_conf_' in compound_name:
            continue
            
        affinity = extract_best_affinity_from_log(log_file)
        
        if affinity is not None:
            binding_data.append({
                'compound': compound_name,
                'affinity': affinity
            })
            logger.info(f"Extracted {compound_name}: {affinity} kcal/mol")
        else:
            logger.warning(f"Failed to extract affinity for {compound_name}")
    
    # Also extract from configuration files
    logger.info("Processing configuration files...")
    conf_files = [f for f in logs_path.glob("*_conf_*.txt")]
    
    for log_file in conf_files:
        compound_name = log_file.stem
        
        affinity = extract_best_affinity_from_log(log_file)
        
        if affinity is not None:
            binding_data.append({
                'compound': compound_name,
                'affinity': affinity
            })
            logger.debug(f"Extracted {compound_name}: {affinity} kcal/mol")
    
    if not binding_data:
        raise ValueError("No binding data could be extracted")
    
    # Create DataFrame
    df = pd.DataFrame(binding_data)
    
    # Remove duplicates (keep first occurrence)
    df = df.drop_duplicates(subset=['compound'], keep='first')
    
    logger.info(f"Extracted binding data for {len(df)} compounds")
    
    return df


def main():
    """Main execution."""
    try:
        # Extract binding data
        binding_df = extract_all_binding_data()
        
        # Create output directory
        output_dir = Path("data/raw")
        output_dir.mkdir(parents=True, exist_ok=True)
        
        # Save to CSV
        output_file = output_dir / "binding_affinities.csv"
        binding_df.to_csv(output_file, index=False)
        
        logger.info(f"Binding data saved to {output_file}")
        
        # Show summary statistics
        logger.info(f"Summary statistics:")
        logger.info(f"  Total compounds: {len(binding_df)}")
        logger.info(f"  Affinity range: {binding_df['affinity'].min():.3f} to {binding_df['affinity'].max():.3f} kcal/mol")
        logger.info(f"  Mean affinity: {binding_df['affinity'].mean():.3f} kcal/mol")
        
        # Show first few entries
        logger.info(f"First 10 entries:")
        for _, row in binding_df.head(10).iterrows():
            logger.info(f"  {row['compound']}: {row['affinity']:.3f} kcal/mol")
        
        return binding_df
        
    except Exception as e:
        logger.error(f"Failed to extract binding data: {e}")
        raise


if __name__ == "__main__":
    binding_df = main()
```
