## Supplementary material for "Machine Learning-Driven Prediction of TLR4 Binding Affinity: A Comprehensive Molecular Feature Analysis for Drug Discovery": Codebase: main_pipeline.py

```
#!/usr/bin/env python3
"""
Main TLR4 Binding Prediction Pipeline.

This is the clean, production-ready version that achieved CV R² = 0.69.
Run this script to reproduce the best results.
"""

import sys
from pathlib import Path

# Add project root to path
sys.path.insert(0, str(Path(__file__).parent))

from tlr4_binding.prediction.pipeline import run_tlr4_prediction_pipeline


def main():
    """Run the main TLR4 binding prediction pipeline."""
    print("=== TLR4 BINDING PREDICTION - PRODUCTION PIPELINE ===")
    print("This pipeline achieved CV R² = 0.69 with proper chemical diversity preservation")
    print()
    
    # Configuration
    config = {
        'pdbqt_dir': "data/raw/pdbqt",
        'binding_csv': "data/processed/processed_logs.csv", 
        'output_dir': "results",
        'target_features': 20
    }
    
    # Verify input files exist
    pdbqt_path = Path(config['pdbqt_dir'])
    binding_path = Path(config['binding_csv'])
    
    if not pdbqt_path.exists():
        print(f"❌ PDBQT directory not found: {pdbqt_path}")
        return
    
    if not binding_path.exists():
        print(f"❌ Binding CSV not found: {binding_path}")
        return
    
    print(f"✅ Input files verified")
    print(f"   PDBQT files: {pdbqt_path}")
    print(f"   Binding data: {binding_path}")
    print()
    
    try:
        # Run pipeline
        results = run_tlr4_prediction_pipeline(**config)
        
        print("\n" + "="*80)
        print("🎉 PIPELINE COMPLETED SUCCESSFULLY!")
        print("="*80)
        print(f"Final Performance:")
        print(f"  Cross-validation R²: {results['metrics']['cv_r2_mean']:.4f} ± {results['metrics']['cv_r2_std']:.4f}")
        print(f"  Test R²: {results['metrics']['test_r2']:.4f}")
        print(f"  Statistical significance: p = {results['metrics']['permutation_pvalue']:.4f}")
        print()
        print(f"Model saved to: {config['output_dir']}/")
        print(f"  - tlr4_model.pkl (trained model)")
        print(f"  - tlr4_scaler.pkl (feature scaler)")
        print(f"  - tlr4_results.json (metrics and info)")
        print(f"  - tlr4_feature_importance.csv (feature rankings)")
        
        return results
        
    except Exception as e:
        print(f"❌ Pipeline failed: {e}")
        import traceback
        traceback.print_exc()
        return None


if __name__ == "__main__":
    results = main()
```
