## Supplementary material for "Machine Learning-Driven Prediction of TLR4 Binding Affinity: A Comprehensive Molecular Feature Analysis for Drug Discovery": Codebase: references.bib

@article{akira2006toll,
  title={Toll-like receptors: critical proteins linking innate and acquired immunity},
  author={Akira, Shizuo and Uematsu, Satoshi and Takeuchi, Osamu},
  journal={Nature immunology},
  volume={7},
  number={2},
  pages={131--137},
  year={2006},
  publisher={Nature Publishing Group}
}

@article{kanzler2007therapeutic,
  title={Therapeutic targeting of innate immunity with Toll-like receptor agonists and antagonists},
  author={Kanzler, Holger and Barrat, Franck J and Hessel, Eric M and Coffman, Robert L},
  journal={Nature medicine},
  volume={13},
  number={5},
  pages={552--559},
  year={2007},
  publisher={Nature Publishing Group}
}

@article{chen2018rise,
  title={The rise of deep learning in drug discovery},
  author={Chen, Hongming and Engkvist, Ola and Wang, Yinhai and Olivecrona, Marcus and Blaschke, Thomas},
  journal={Drug discovery today},
  volume={23},
  number={6},
  pages={1241--1250},
  year={2018},
  publisher={Elsevier}
}

@article{jimenez2018kdeep,
  title={K deep: protein--ligand absolute binding affinity prediction via 3d-convolutional neural networks},
  author={Jim{\'e}nez, Jos{\'e} and Skalic, Miha and Martinez-Rosell, Gerard and De Fabritiis, Gianni},
  journal={Journal of chemical information and modeling},
  volume={58},
  number={2},
  pages={287--296},
  year={2018},
  publisher={ACS Publications}
}

@misc{landrum2013rdkit,
  title={RDKit: Open-source cheminformatics},
  author={Landrum, Greg and others},
  year={2013},
  howpublished={\url{http://www.rdkit.org}}
}

@article{pedregosa2011scikit,
  title={Scikit-learn: Machine learning in Python},
  author={Pedregosa, Fabian and Varoquaux, Ga{\"e}l and Gramfort, Alexandre and Michel, Vincent and Thirion, Bertrand and Grisel, Olivier and Blondel, Mathieu and Prettenhofer, Peter and Weiss, Ron and Dubourg, Vincent and others},
  journal={the Journal of machine Learning research},
  volume={12},
  pages={2825--2830},
  year={2011},
  publisher={JMLR. org}
}

@article{bemis1996properties,
  title={The properties of known drugs. 1. Molecular frameworks},
  author={Bemis, Guy W and Murcko, Mark A},
  journal={Journal of medicinal chemistry},
  volume={39},
  number={15},
  pages={2887--2893},
  year={1996},
  publisher={ACS Publications}
}

@inproceedings{akiba2019optuna,
  title={Optuna: A next-generation hyperparameter optimization framework},
  author={Akiba, Takuya and Sano, Shotaro and Yanase, Toshihiko and Ohta, Takeru and Koyama, Masanori},
  booktitle={Proceedings of the 25th ACM SIGKDD international conference on knowledge discovery \& data mining},
  pages={2623--2631},
  year={2019}
}

@article{chen2016xgboost,
  title={Xgboost: A scalable tree boosting system},
  author={Chen, Tianqi and Guestrin, Carlos},
  journal={Proceedings of the 22nd acm sigkdd international conference on knowledge discovery and data mining},
  pages={785--794},
  year={2016}
}

@article{ke2017lightgbm,
  title={Lightgbm: A highly efficient gradient boosting decision tree},
  author={Ke, Guolin and Meng, Qi and Finley, Thomas and Wang, Taifeng and Chen, Wei and Ma, Weidong and Ye, Qiwei and Liu, Tie-Yan},
  journal={Advances in neural information processing systems},
  volume={30},
  year={2017}
}

@article{breiman2001random,
  title={Random forests},
  author={Breiman, Leo},
  journal={Machine learning},
  volume={45},
  number={1},
  pages={5--32},
  year={2001},
  publisher={Springer}
}

@article{cortes1995support,
  title={Support-vector networks},
  author={Cortes, Corinna and Vapnik, Vladimir},
  journal={Machine learning},
  volume={20},
  number={3},
  pages={273--297},
  year={1995},
  publisher={Springer}
}

@article{morris2009autodock4,
  title={AutoDock4 and AutoDockTools4: Automated docking with selective receptor flexibility},
  author={Morris, Garrett M and Huey, Ruth and Lindstrom, William and Sanner, Michel F and Belew, Richard K and Goodsell, David S and Olson, Arthur J},
  journal={Journal of computational chemistry},
  volume={30},
  number={16},
  pages={2785--2791},
  year={2009},
  publisher={Wiley Online Library}
}

@article{lipinski2001experimental,
  title={Experimental and computational approaches to estimate solubility and permeability in drug discovery and development settings},
  author={Lipinski, Christopher A and Lombardo, Franco and Dominy, Beryl W and Feeney, Paul J},
  journal={Advanced drug delivery reviews},
  volume={46},
  number={1-3},
  pages={3--26},
  year={2001},
  publisher={Elsevier}
}

@article{muegge2001selection,
  title={Selection criteria for drug-like compounds},
  author={Muegge, Ingo and Heald, Sherry L and Brittelli, David},
  journal={Medicinal research reviews},
  volume={21},
  number={1},
  pages={1--26},
  year={2001},
  publisher={Wiley Online Library}
}
