## Supplementary material for "Machine Learning-Driven Prediction of TLR4 Binding Affinity: A Comprehensive Molecular Feature Analysis for Drug Discovery": Codebase: README.md

Code Associated With:

## Machine Learning-Driven Prediction of TLR4 Binding Affinity: A Comprehensive Molecular Feature Analysis for Drug Discovery

#### Brandon Yee, Maximilian Rutowski, and Wilson Collins


Data from: [TLR4 Binding Data](https://github.com/YCRG-Labs/binding-data)

### **Quick Start**

``` bash
# 0) Setup repository
git clone https://github.com/YCRG-Labs/tlr4-drug-discovery.git
cd drug_discovery_tlr4
git clone https://github.com/YCRG-Labs/binding-data.git

# 1) Environment setup
python -m venv venv
# Windows:
venv\Scripts\activate
# Linux/Mac:
source venv/bin/activate

# 2) Install dependencies
pip install -r requirements.txt

# 3) Run Enhanced Research Pipeline (RECOMMENDED)
python main.py --pdbqt-dir binding-data/raw/pdbqt --binding-csv binding-data/processed/processed_logs.csv --output-dir results --splitting scaffold

# Alternative splitting methods:
# --splitting random    # Random splitting
# --splitting scaffold  # Molecular scaffold splitting (prevents data leakage)


# 5) Run tests
python -m pytest tests -v
```

### **Stage-by-Stage Usage**

``` bash
# Feature extraction only
python main.py --pipeline features --pdbqt-dir binding-data/raw/pdbqt

# Model training only  
python main.py --pipeline train --features-csv data/processed/features.csv --binding-csv binding-data/processed/processed_logs.csv

# Single compound prediction
python main.py --pipeline predict --model-path models/trained/best_model.joblib --pdbqt-file binding-data/raw/pdbqt/example.pdbqt
```
