## Supplementary material for "Machine Learning-Driven Prediction of TLR4 Binding Affinity: A Comprehensive Molecular Feature Analysis for Drug Discovery": Codebase: diagnostic_analysis.py

```
#!/usr/bin/env python3
"""
Diagnostic Analysis to Identify Data Leakage and Overfitting Issues.
"""

import pandas as pd
import numpy as np
import matplotlib.pyplot as plt
from sklearn.model_selection import train_test_split
from sklearn.ensemble import RandomForestRegressor
from sklearn.metrics import r2_score, mean_squared_error
from sklearn.preprocessing import StandardScaler
import seaborn as sns
from pathlib import Path

def diagnose_data_issues():
    """Comprehensive diagnostic analysis."""
    
    print("=" * 80)
    print("DIAGNOSTIC ANALYSIS: IDENTIFYING DATA ISSUES")
    print("=" * 80)
    
    # Load data
    df = pd.read_csv('data/processed/processed_logs.csv')
    print(f"\n1. DATA OVERVIEW:")
    print(f"   Original shape: {df.shape}")
    print(f"   Columns: {df.columns.tolist()}")
    
    # Get best binding for each compound
    best_binding = df.loc[df.groupby('ligand')['affinity'].idxmin()]
    print(f"   Best binding shape: {best_binding.shape}")
    
    # Check available features
    exclude_cols = ['affinity', 'ligand', 'mode']
    feature_cols = [col for col in best_binding.columns if col not in exclude_cols]
    print(f"   Available features: {feature_cols}")
    
    # Analyze feature quality
    print(f"\n2. FEATURE QUALITY ANALYSIS:")
    for col in feature_cols:
        missing = best_binding[col].isnull().sum()
        unique = best_binding[col].nunique()
        print(f"   {col}: {missing} missing, {unique} unique values")
        if unique <= 10:
            print(f"      Values: {sorted(best_binding[col].dropna().unique())}")
    
    # Check for data leakage indicators
    print(f"\n3. DATA LEAKAGE ANALYSIS:")
    
    # Check if any features are perfectly correlated with target
    for col in feature_cols:
        if best_binding[col].dtype in ['int64', 'float64']:
            corr = best_binding[col].corr(best_binding['affinity'])
            if abs(corr) > 0.95:
                print(f"   WARNING: {col} highly correlated with target (r={corr:.4f})")
    
    # Check target distribution
    print(f"\n4. TARGET ANALYSIS:")
    print(f"   Affinity range: {best_binding['affinity'].min():.3f} to {best_binding['affinity'].max():.3f}")
    print(f"   Affinity std: {best_binding['affinity'].std():.3f}")
    print(f"   Unique values: {best_binding['affinity'].nunique()}")
    
    # Simple model test with proper validation
    print(f"\n5. SIMPLE MODEL TEST:")
    
    # Prepare data
    X = best_binding[feature_cols].fillna(0)
    y = best_binding['affinity']
    
    print(f"   Feature matrix shape: {X.shape}")
    print(f"   Feature statistics:")
    for col in X.columns:
        print(f"      {col}: mean={X[col].mean():.4f}, std={X[col].std():.4f}")
    
    # Split data properly
    X_train, X_test, y_train, y_test = train_test_split(
        X, y, test_size=0.2, random_state=42
    )
    
    # Train simple model
    model = RandomForestRegressor(n_estimators=10, random_state=42)
    model.fit(X_train, y_train)
    
    # Evaluate
    train_pred = model.predict(X_train)
    test_pred = model.predict(X_test)
    
    train_r2 = r2_score(y_train, train_pred)
    test_r2 = r2_score(y_test, test_pred)
    
    print(f"   Train R²: {train_r2:.4f}")
    print(f"   Test R²: {test_r2:.4f}")
    print(f"   Overfitting gap: {train_r2 - test_r2:.4f}")
    
    if train_r2 > 0.99:
        print("   🚨 SEVERE OVERFITTING DETECTED!")
    elif train_r2 - test_r2 > 0.2:
        print("   ⚠️  SIGNIFICANT OVERFITTING DETECTED!")
    
    # Feature importance
    if hasattr(model, 'feature_importances_'):
        print(f"\n6. FEATURE IMPORTANCE:")
        for i, col in enumerate(X.columns):
            importance = model.feature_importances_[i]
            print(f"   {col}: {importance:.4f}")
    
    # Check for duplicate or near-duplicate samples
    print(f"\n7. DUPLICATE ANALYSIS:")
    
    # Check for exact duplicates in features
    feature_duplicates = X.duplicated().sum()
    print(f"   Exact feature duplicates: {feature_duplicates}")
    
    # Check for exact duplicates in target
    target_duplicates = y.duplicated().sum()
    print(f"   Exact target duplicates: {target_duplicates}")
    
    # Recommendations
    print(f"\n8. RECOMMENDATIONS:")
    
    if X.shape[1] < 5:
        print("   🔧 ISSUE: Very few features - need more molecular descriptors")
    
    if train_r2 > 0.99:
        print("   🔧 ISSUE: Perfect training fit suggests data leakage or overfitting")
    
    if feature_duplicates > X.shape[0] * 0.1:
        print("   🔧 ISSUE: Too many duplicate samples")
    
    print("\n" + "=" * 80)
    
    return {
        'data_shape': best_binding.shape,
        'feature_count': len(feature_cols),
        'train_r2': train_r2,
        'test_r2': test_r2,
        'overfitting_gap': train_r2 - test_r2
    }

if __name__ == "__main__":
    results = diagnose_data_issues()
```
