## Supplementary material for "Machine Learning-Driven Prediction of TLR4 Binding Affinity: A Comprehensive Molecular Feature Analysis for Drug Discovery": Codebase: README.md

# Scripts Directory

This directory contains utility and analysis scripts for the TLR4 binding prediction project.

## Analysis Scripts

### `diagnostic_analysis.py`
- **Purpose**: Identifies data leakage and overfitting issues
- **Usage**: `python scripts/diagnostic_analysis.py`
- **Output**: Diagnostic reports on data quality and model performance

### `verification_analysis.py`
- **Purpose**: Verifies that no artificial data generation is occurring
- **Usage**: `python scripts/verification_analysis.py`
- **Output**: Verification report ensuring realistic performance

### `publication_analysis.py`
- **Purpose**: Generates publication-ready analysis and figures
- **Features**: Feature importance, literature comparison, SAR analysis
- **Usage**: `python scripts/publication_analysis.py`

### `create_publication_figures.py`
- **Purpose**: Creates high-quality figures for publication
- **Usage**: `python scripts/create_publication_figures.py`
- **Output**: Publication-ready plots and visualizations

## Legacy Pipeline Scripts

The following scripts are legacy implementations kept for reference:

- `corrected_realistic_pipeline.py`: Early realistic pipeline implementation
- `realistic_research_pipeline.py`: Research-grade pipeline attempt
- `improved_research_pipeline.py`: Improved pipeline with calibration
- `final_improved_pipeline.py`: Final improved pipeline version
- `fixed_smiles_pipeline.py`: SMILES-based molecular descriptor pipeline

**Note**: These legacy scripts are superseded by the main pipeline in `main.py`. They are kept for historical reference and comparison purposes.

## Usage

All scripts should be run from the project root directory:

```bash
# Run diagnostic analysis
python scripts/diagnostic_analysis.py

# Generate publication figures
python scripts/create_publication_figures.py

# Verify results authenticity
python scripts/verification_analysis.py
```

## Dependencies

Scripts may require additional dependencies beyond the main project requirements. Install with:

```bash
pip install -r requirements.txt
```
