## Supplementary material for "Machine Learning-Driven Prediction of TLR4 Binding Affinity: A Comprehensive Molecular Feature Analysis for Drug Discovery": Codebase: test_compound_matching_integration.py

```
"""
Integration tests for compound name matching and data integration.

Tests the complete pipeline from molecular feature extraction to
binding data integration with real-world scenarios.
"""

import pytest
import pandas as pd
import numpy as np
import sys
import os
from pathlib import Path

# Add src to path for imports
sys.path.insert(0, os.path.join(os.path.dirname(__file__), '..', '..', 'src'))

from tlr4_binding.data_processing.preprocessor import (
    DataPreprocessor, 
    BindingDataLoader, 
    CompoundMatcher, 
    DataIntegrator
)
from tlr4_binding.molecular_analysis.features import MolecularFeatures


class TestCompoundMatchingIntegration:
    """Integration tests for compound name matching and data integration."""
    
    def setup_method(self):
        """Set up test fixtures."""
        self.preprocessor = DataPreprocessor()
        self.binding_loader = BindingDataLoader()
        self.compound_matcher = CompoundMatcher(threshold=75.0)
        self.data_integrator = DataIntegrator(self.compound_matcher)
        
        # Path to test data
        self.test_data_dir = Path(__file__).parent.parent.parent / "binding-data" / "processed"
        self.binding_csv_path = self.test_data_dir / "processed_logs.csv"
        
        # Sample molecular features for testing
        self.sample_features = pd.DataFrame({
            'compound_name': [
                'Andrographolide',
                'Curcumin',
                'Resveratrol', 
                'Quercetin',
                'Epigallocatechin',
                'Capsaicin',
                'Gingerol',
                'Caffeic acid',
                'Chlorogenic acid',
                'Ferulic acid',
                'Unknown compound 1',
                'Unknown compound 2'
            ],
            'molecular_weight': [350.4, 368.4, 228.2, 302.2, 458.4, 305.4, 276.4, 180.2, 354.3, 194.2, 100.0, 200.0],
            'logp': [2.3, 2.5, 3.1, 1.8, 2.0, 3.8, 2.9, 1.2, 0.8, 1.4, 1.0, 2.0],
            'tpsa': [74.6, 93.1, 60.7, 131.4, 197.4, 58.6, 50.5, 77.8, 164.7, 66.8, 50.0, 60.0],
            'hbd': [3, 2, 3, 5, 8, 1, 1, 2, 4, 2, 1, 2],
            'hba': [4, 6, 3, 7, 11, 2, 2, 4, 7, 3, 2, 3]
        })
    
    def test_real_binding_data_loading(self):
        """Test loading real binding data from CSV."""
        if not self.binding_csv_path.exists():
            pytest.skip(f"Binding data file not found: {self.binding_csv_path}")
        
        # Load binding data
        binding_df = self.binding_loader.load_csv(str(self.binding_csv_path))
        
        # Validate data structure
        assert len(binding_df) > 0
        assert 'ligand' in binding_df.columns
        assert 'affinity' in binding_df.columns
        assert 'mode' in binding_df.columns
        
        # Validate data quality
        assert binding_df['affinity'].notna().all()
        assert binding_df['ligand'].notna().all()
        assert binding_df['mode'].notna().all()
        
        # Check affinity value ranges (should be negative for binding)
        assert binding_df['affinity'].max() < 10  # Not too positive
        assert binding_df['affinity'].min() > -20  # Not too negative
    
    def test_binding_data_validation(self):
        """Test binding data validation with real data."""
        if not self.binding_csv_path.exists():
            pytest.skip(f"Binding data file not found: {self.binding_csv_path}")
        
        binding_df = self.binding_loader.load_csv(str(self.binding_csv_path))
        
        # Validate data quality
        validation_results = self.binding_loader.validate_binding_data(binding_df)
        
        # Should have some validation results
        assert isinstance(validation_results, dict)
        assert 'missing_values' in validation_results
        assert 'invalid_affinities' in validation_results
        assert 'outliers' in validation_results
    
    def test_best_affinities_extraction(self):
        """Test extraction of best binding affinities from real data."""
        if not self.binding_csv_path.exists():
            pytest.skip(f"Binding data file not found: {self.binding_csv_path}")
        
        binding_df = self.binding_loader.load_csv(str(self.binding_csv_path))
        
        # Extract best affinities
        best_affinities = self.preprocessor.get_best_affinities(binding_df)
        
        # Validate results
        assert len(best_affinities) > 0
        assert len(best_affinities) <= len(binding_df)
        
        # Each compound should have only one best affinity
        assert best_affinities['ligand'].nunique() == len(best_affinities)
        
        # Best affinities should be the minimum (most negative) for each compound
        for _, row in best_affinities.iterrows():
            ligand = row['ligand']
            best_affinity = row['affinity']
            
            # Get all affinities for this ligand
            ligand_affinities = binding_df[binding_df['ligand'] == ligand]['affinity']
            
            # Best affinity should be the minimum
            assert best_affinity == ligand_affinities.min()
    
    def test_compound_matching_with_real_data(self):
        """Test compound matching with real binding data."""
        if not self.binding_csv_path.exists():
            pytest.skip(f"Binding data file not found: {self.binding_csv_path}")
        
        binding_df = self.binding_loader.load_csv(str(self.binding_csv_path))
        
        # Get unique compound names from binding data
        binding_compounds = binding_df['ligand'].unique().tolist()
        
        # Test matching with sample features
        feature_compounds = self.sample_features['compound_name'].tolist()
        
        matches = self.compound_matcher.match_compounds(feature_compounds, binding_compounds)
        
        # Should find some matches
        assert len(matches) > 0
        
        # Check match quality
        for feature_name, binding_name in matches.items():
            confidence = self.compound_matcher.get_match_confidence(feature_name, binding_name)
            assert confidence >= self.compound_matcher.threshold
    
    def test_data_integration_with_real_data(self):
        """Test complete data integration with real binding data."""
        if not self.binding_csv_path.exists():
            pytest.skip(f"Binding data file not found: {self.binding_csv_path}")
        
        # Load and process binding data
        binding_df = self.binding_loader.load_csv(str(self.binding_csv_path))
        best_affinities = self.preprocessor.get_best_affinities(binding_df)
        
        # Integrate with sample features
        integrated_df = self.data_integrator.integrate_datasets(
            self.sample_features,
            best_affinities
        )
        
        # Validate integration results
        assert len(integrated_df) > 0
        
        # Should contain both feature and binding columns
        feature_columns = set(self.sample_features.columns)
        binding_columns = set(best_affinities.columns)
        result_columns = set(integrated_df.columns)
        
        assert feature_columns.issubset(result_columns)
        assert binding_columns.issubset(result_columns)
        
        # Check integration statistics
        stats = self.data_integrator.get_integration_stats()
        assert stats['integrated_records'] == len(integrated_df)
        assert stats['match_rate'] > 0
    
    def test_complete_preprocessing_pipeline(self):
        """Test complete preprocessing pipeline with real data."""
        if not self.binding_csv_path.exists():
            pytest.skip(f"Binding data file not found: {self.binding_csv_path}")
        
        # Run complete preprocessing pipeline
        integrated_df = self.preprocessor.preprocess_pipeline(
            self.sample_features,
            str(self.binding_csv_path)
        )
        
        # Validate pipeline results
        assert len(integrated_df) > 0
        
        # Should have both molecular features and binding data
        assert 'molecular_weight' in integrated_df.columns
        assert 'affinity' in integrated_df.columns
        assert 'compound_name' in integrated_df.columns
        assert 'ligand' in integrated_df.columns
        
        # Should have valid data
        assert integrated_df['affinity'].notna().all()
        assert integrated_df['molecular_weight'].notna().all()
    
    def test_integration_with_different_thresholds(self):
        """Test integration with different matching thresholds."""
        if not self.binding_csv_path.exists():
            pytest.skip(f"Binding data file not found: {self.binding_csv_path}")
        
        binding_df = self.binding_loader.load_csv(str(self.binding_csv_path))
        best_affinities = self.preprocessor.get_best_affinities(binding_df)
        
        # Test with different thresholds
        thresholds = [60.0, 75.0, 85.0, 95.0]
        results = {}
        
        for threshold in thresholds:
            matcher = CompoundMatcher(threshold=threshold)
            integrator = DataIntegrator(matcher)
            
            integrated_df = integrator.integrate_datasets(
                self.sample_features,
                best_affinities
            )
            
            stats = integrator.get_integration_stats()
            results[threshold] = {
                'records': len(integrated_df),
                'match_rate': stats['match_rate']
            }
        
        # Higher thresholds should generally result in fewer matches
        assert results[60.0]['records'] >= results[95.0]['records']
        assert results[60.0]['match_rate'] >= results[95.0]['match_rate']
    
    def test_integration_with_partial_ratio_vs_ratio(self):
        """Test integration with different ratio calculation methods."""
        if not self.binding_csv_path.exists():
            pytest.skip(f"Binding data file not found: {self.binding_csv_path}")
        
        binding_df = self.binding_loader.load_csv(str(self.binding_csv_path))
        best_affinities = self.preprocessor.get_best_affinities(binding_df)
        
        # Test with partial ratio
        matcher_partial = CompoundMatcher(threshold=75.0, use_partial_ratio=True)
        integrator_partial = DataIntegrator(matcher_partial)
        result_partial = integrator_partial.integrate_datasets(
            self.sample_features,
            best_affinities
        )
        
        # Test with regular ratio
        matcher_ratio = CompoundMatcher(threshold=75.0, use_partial_ratio=False)
        integrator_ratio = DataIntegrator(matcher_ratio)
        result_ratio = integrator_ratio.integrate_datasets(
            self.sample_features,
            best_affinities
        )
        
        # Both should work, but may have different results
        assert len(result_partial) >= 0
        assert len(result_ratio) >= 0
    
    def test_integration_error_handling(self):
        """Test error handling in integration pipeline."""
        # Test with invalid CSV path
        with pytest.raises(FileNotFoundError):
            self.preprocessor.preprocess_pipeline(
                self.sample_features,
                "nonexistent_file.csv"
            )
        
        # Test with empty features DataFrame
        empty_features = pd.DataFrame(columns=['compound_name', 'molecular_weight'])
        if self.binding_csv_path.exists():
            with pytest.raises(KeyError):
                self.data_integrator.integrate_datasets(
                    empty_features,
                    pd.DataFrame(columns=['ligand', 'affinity'])
                )
    
    def test_integration_data_quality_validation(self):
        """Test data quality validation in integrated dataset."""
        if not self.binding_csv_path.exists():
            pytest.skip(f"Binding data file not found: {self.binding_csv_path}")
        
        # Run integration
        integrated_df = self.preprocessor.preprocess_pipeline(
            self.sample_features,
            str(self.binding_csv_path)
        )
        
        if len(integrated_df) > 0:
            # Check for missing values
            missing_values = integrated_df.isnull().sum()
            assert missing_values['affinity'] == 0  # No missing affinities
            assert missing_values['compound_name'] == 0  # No missing compound names
            
            # Check for duplicate compound names
            duplicates = integrated_df['compound_name'].duplicated().sum()
            assert duplicates == 0  # No duplicate compounds
            
            # Check affinity value ranges
            assert integrated_df['affinity'].min() < 0  # Should have negative affinities
            assert integrated_df['affinity'].max() < 10  # Should not be too positive
    
    def test_integration_performance_with_large_data(self):
        """Test integration performance with larger datasets."""
        if not self.binding_csv_path.exists():
            pytest.skip(f"Binding data file not found: {self.binding_csv_path}")
        
        # Create larger feature dataset
        large_features = pd.concat([self.sample_features] * 5).reset_index(drop=True)
        large_features['compound_name'] = [
            f"{name}_{i//len(self.sample_features)}" 
            for i, name in enumerate(large_features['compound_name'])
        ]
        
        # Run integration
        start_time = pd.Timestamp.now()
        integrated_df = self.preprocessor.preprocess_pipeline(
            large_features,
            str(self.binding_csv_path)
        )
        end_time = pd.Timestamp.now()
        
        # Should complete in reasonable time (less than 30 seconds)
        processing_time = (end_time - start_time).total_seconds()
        assert processing_time < 30
        
        # Should have some results
        assert len(integrated_df) >= 0
    
    def test_integration_with_special_characters(self):
        """Test integration with special characters in compound names."""
        # Create features with special characters
        special_features = pd.DataFrame({
            'compound_name': [
                'Compound-123',
                'Ligand@Name',
                'Molecule#Test',
                'Drug$Special',
                'Test%Chars'
            ],
            'molecular_weight': [100.0, 200.0, 300.0, 400.0, 500.0],
            'logp': [1.0, 2.0, 3.0, 4.0, 5.0]
        })
        
        # Create corresponding binding data
        special_binding = pd.DataFrame({
            'ligand': [
                'compound 123',
                'ligand name',
                'molecule test',
                'drug special',
                'test chars'
            ],
            'mode': [1, 1, 1, 1, 1],
            'affinity': [-5.0, -6.0, -7.0, -8.0, -9.0]
        })
        
        # Test integration
        integrated_df = self.data_integrator.integrate_datasets(
            special_features,
            special_binding
        )
        
        # Should handle special characters
        assert len(integrated_df) > 0
    
    def test_integration_statistics_accuracy(self):
        """Test accuracy of integration statistics."""
        if not self.binding_csv_path.exists():
            pytest.skip(f"Binding data file not found: {self.binding_csv_path}")
        
        binding_df = self.binding_loader.load_csv(str(self.binding_csv_path))
        best_affinities = self.preprocessor.get_best_affinities(binding_df)
        
        # Run integration
        integrated_df = self.data_integrator.integrate_datasets(
            self.sample_features,
            best_affinities
        )
        
        # Get statistics
        stats = self.data_integrator.get_integration_stats()
        
        # Validate statistics accuracy
        assert stats['total_features'] == len(self.sample_features)
        assert stats['total_binding'] == len(best_affinities)
        assert stats['integrated_records'] == len(integrated_df)
        
        # Match rate should be between 0 and 1
        assert 0 <= stats['match_rate'] <= 1
        
        # If we have matches, match rate should be positive
        if len(integrated_df) > 0:
            assert stats['match_rate'] > 0


if __name__ == "__main__":
    pytest.main([__file__, "-v"])
```
