## Supplementary material for "Machine Learning-Driven Prediction of TLR4 Binding Affinity: A Comprehensive Molecular Feature Analysis for Drug Discovery": Codebase: test_feature_extractor_integration.py

```
#!/usr/bin/env python3
"""
Simple integration test script for the comprehensive molecular feature extractor.

This script tests the MolecularFeatureExtractor with actual PDBQT files
from the binding-data directory to verify end-to-end functionality.
"""

import sys
import logging
from pathlib import Path
import pandas as pd

# Add src to path
sys.path.insert(0, str(Path(__file__).parent.parent.parent / "src"))

from tlr4_binding.molecular_analysis.extractor import MolecularFeatureExtractor

# Set up logging
logging.basicConfig(
    level=logging.INFO,
    format='%(asctime)s - %(name)s - %(levelname)s - %(message)s'
)
logger = logging.getLogger(__name__)


def test_single_file_extraction():
    """Test extraction from a single PDBQT file."""
    logger.info("=== Testing Single File Extraction ===")
    
    # Initialize extractor
    extractor = MolecularFeatureExtractor(
        include_2d_features=True,
        include_3d_features=True,
        include_advanced_features=True
    )
    
    # Find a sample PDBQT file
    pdbqt_dir = Path("binding-data/raw/pdbqt")
    if not pdbqt_dir.exists():
        logger.error("PDBQT directory not found: binding-data/raw/pdbqt")
        return False
    
    pdbqt_files = list(pdbqt_dir.glob("*.pdbqt"))
    if not pdbqt_files:
        logger.error("No PDBQT files found")
        return False
    
    # Test with first file
    test_file = str(pdbqt_files[0])
    logger.info(f"Testing with file: {test_file}")
    
    try:
        # Extract features
        features = extractor.extract_features(test_file)
        
        # Display results
        logger.info(f"Successfully extracted features for: {features.compound_name}")
        logger.info(f"Molecular weight: {features.molecular_weight}")
        logger.info(f"LogP: {features.logp}")
        logger.info(f"TPSA: {features.tpsa}")
        logger.info(f"Radius of gyration: {features.radius_of_gyration}")
        
        # Check statistics
        stats = extractor.get_extraction_stats()
        logger.info(f"Processing time: {stats['processing_times'][0]:.2f}s")
        logger.info(f"Memory used: {stats['memory_usage'][0]:.1f}MB")
        
        return True
        
    except Exception as e:
        logger.error(f"Failed to extract features: {str(e)}")
        return False


def test_batch_extraction():
    """Test batch extraction from multiple PDBQT files."""
    logger.info("=== Testing Batch Extraction ===")
    
    # Initialize extractor
    extractor = MolecularFeatureExtractor()
    
    # Find PDBQT files
    pdbqt_dir = Path("binding-data/raw/pdbqt")
    if not pdbqt_dir.exists():
        logger.error("PDBQT directory not found")
        return False
    
    # Test with first 3 files
    pdbqt_files = list(pdbqt_dir.glob("*.pdbqt"))[:3]
    if not pdbqt_files:
        logger.error("No PDBQT files found")
        return False
    
    logger.info(f"Testing batch extraction with {len(pdbqt_files)} files")
    
    try:
        # Extract features from list
        features_df = extractor.extract_features_from_list([str(f) for f in pdbqt_files])
        
        # Display results
        logger.info(f"Successfully extracted features for {len(features_df)} compounds")
        logger.info(f"DataFrame shape: {features_df.shape}")
        logger.info(f"Columns: {list(features_df.columns)}")
        
        # Display feature summary
        summary = extractor.get_feature_summary(features_df)
        logger.info(f"Feature summary generated for {len(summary)} features")
        
        # Display performance report
        performance_report = extractor.get_performance_report()
        batch_summary = performance_report['batch_summary']
        logger.info(f"Batch processing completed:")
        logger.info(f"  Success rate: {batch_summary['success_rate']:.1f}%")
        logger.info(f"  Total time: {batch_summary['total_time']:.2f}s")
        
        # Display efficiency metrics
        efficiency = extractor.get_feature_extraction_efficiency()
        if efficiency:
            logger.info(f"Efficiency metrics:")
            for key, value in efficiency.items():
                logger.info(f"  {key}: {value:.2f}")
        
        return True
        
    except Exception as e:
        logger.error(f"Batch extraction failed: {str(e)}")
        return False


def test_feature_validation():
    """Test feature validation functionality."""
    logger.info("=== Testing Feature Validation ===")
    
    # Initialize extractor
    extractor = MolecularFeatureExtractor()
    
    # Find PDBQT files
    pdbqt_dir = Path("binding-data/raw/pdbqt")
    if not pdbqt_dir.exists():
        logger.error("PDBQT directory not found")
        return False
    
    # Test with first 5 files
    pdbqt_files = list(pdbqt_dir.glob("*.pdbqt"))[:5]
    if not pdbqt_files:
        logger.error("No PDBQT files found")
        return False
    
    try:
        # Extract features
        features_df = extractor.extract_features_from_list([str(f) for f in pdbqt_files])
        
        # Validate features
        validation_results = extractor.validate_features(features_df)
        
        # Display validation results
        total_issues = sum(len(issues) for issues in validation_results.values())
        logger.info(f"Feature validation completed. Total issues found: {total_issues}")
        
        for issue_type, issues in validation_results.items():
            if issues:
                logger.info(f"{issue_type}: {len(issues)} issues")
                for issue in issues[:3]:  # Show first 3 issues
                    logger.info(f"  - {issue}")
        
        return True
        
    except Exception as e:
        logger.error(f"Feature validation failed: {str(e)}")
        return False


def main():
    """Run all integration tests."""
    logger.info("Starting MolecularFeatureExtractor Integration Tests")
    
    tests = [
        test_single_file_extraction,
        test_batch_extraction,
        test_feature_validation
    ]
    
    passed = 0
    failed = 0
    
    for test_func in tests:
        try:
            if test_func():
                passed += 1
                logger.info(f"✓ {test_func.__name__} PASSED")
            else:
                failed += 1
                logger.error(f"✗ {test_func.__name__} FAILED")
        except Exception as e:
            failed += 1
            logger.error(f"✗ {test_func.__name__} FAILED with exception: {str(e)}")
        
        logger.info("-" * 60)
    
    # Summary
    logger.info(f"Integration Tests Summary:")
    logger.info(f"  Passed: {passed}")
    logger.info(f"  Failed: {failed}")
    logger.info(f"  Total: {passed + failed}")
    
    if failed == 0:
        logger.info("🎉 All integration tests passed!")
        return True
    else:
        logger.error(f"❌ {failed} integration tests failed!")
        return False


if __name__ == "__main__":
    success = main()
    sys.exit(0 if success else 1)
```
