## Supplementary material for "Machine Learning-Driven Prediction of TLR4 Binding Affinity: A Comprehensive Molecular Feature Analysis for Drug Discovery": Codebase: test_ml_pipeline.py

```
"""
Integration tests for ML model training pipeline.

This module tests the complete ML pipeline including data preprocessing,
feature engineering, model training, and evaluation.
"""

import unittest
import sys
import tempfile
import shutil
from pathlib import Path
import pandas as pd
import numpy as np
from unittest.mock import patch, MagicMock

# Add src to path
sys.path.insert(0, str(Path(__file__).parent.parent.parent.parent / "src"))

from tlr4_binding.ml_components.trainer import MLModelTrainer
from tlr4_binding.ml_components.feature_engineering import FeatureEngineeringPipeline
from tlr4_binding.data_processing.preprocessor import DataPreprocessor
from tlr4_binding.molecular_analysis.features import MolecularFeatures, BindingData


class TestMLPipelineIntegration(unittest.TestCase):
    """Test complete ML pipeline integration."""
    
    def setUp(self):
        """Set up test data and temporary directory."""
        np.random.seed(42)
        self.n_samples = 200
        self.n_features = 15
        
        # Create realistic synthetic molecular data
        self.molecular_data = self._create_molecular_data()
        self.binding_data = self._create_binding_data()
        
        # Create temporary directory
        self.temp_dir = tempfile.mkdtemp()
    
    def tearDown(self):
        """Clean up temporary directory."""
        shutil.rmtree(self.temp_dir)
    
    def _create_molecular_data(self):
        """Create synthetic molecular features data."""
        data = []
        for i in range(self.n_samples):
            features = MolecularFeatures(
                compound_name=f"compound_{i:03d}",
                molecular_weight=100 + np.random.randn() * 50,
                logp=-2 + np.random.randn() * 3,
                tpsa=20 + np.random.randn() * 30,
                rotatable_bonds=int(np.random.poisson(5)),
                hbd=int(np.random.poisson(3)),
                hba=int(np.random.poisson(4)),
                formal_charge=int(np.random.choice([-2, -1, 0, 1, 2])),
                radius_of_gyration=3 + np.random.randn() * 2,
                molecular_volume=100 + np.random.randn() * 50,
                surface_area=150 + np.random.randn() * 40,
                asphericity=0.1 + np.random.rand() * 0.8,
                ring_count=int(np.random.poisson(2)),
                aromatic_rings=int(np.random.poisson(1)),
                branching_index=0.1 + np.random.rand() * 0.9,
                dipole_moment=np.random.randn() * 2,
                polarizability=20 + np.random.randn() * 10
            )
            data.append(features.to_dict())
        
        return pd.DataFrame(data)
    
    def _create_binding_data(self):
        """Create synthetic binding affinity data."""
        data = []
        for i in range(self.n_samples):
            # Create multiple binding modes per compound
            n_modes = np.random.randint(1, 4)
            for mode in range(n_modes):
                binding = BindingData(
                    ligand=f"compound_{i:03d}",
                    mode=mode + 1,
                    affinity=-8 + np.random.randn() * 2,  # Most compounds have strong binding
                    rmsd_lb=0.0,
                    rmsd_ub=2.0 + np.random.rand() * 2
                )
                data.append(binding.to_dict())
        
        return pd.DataFrame(data)
    
    def test_complete_pipeline(self):
        """Test complete ML pipeline from data to predictions."""
        # Step 1: Data preprocessing
        preprocessor = DataPreprocessor()
        
        # Get best binding affinities
        best_affinities = preprocessor.get_best_affinities(self.binding_data)
        self.assertGreater(len(best_affinities), 0)
        
        # Integrate datasets
        integrated_data = preprocessor.integrate_datasets(
            self.molecular_data, best_affinities
        )
        self.assertGreater(len(integrated_data), 0)
        self.assertIn('affinity', integrated_data.columns)
        
        # Step 2: Feature engineering
        feature_cols = [col for col in integrated_data.columns 
                       if col not in ['compound_name', 'affinity']]
        X = integrated_data[feature_cols]
        y = integrated_data['affinity']
        
        # Apply feature engineering
        feature_pipeline = FeatureEngineeringPipeline()
        X_processed = feature_pipeline.fit_transform(X, y)
        
        self.assertEqual(X_processed.shape[0], X.shape[0])
        self.assertGreater(X_processed.shape[1], 0)
        
        # Step 3: Train-test split
        from sklearn.model_selection import train_test_split
        X_train, X_test, y_train, y_test = train_test_split(
            X_processed, y, test_size=0.2, random_state=42
        )
        
        # Step 4: Model training
        trainer = MLModelTrainer(models_dir=self.temp_dir)
        trained_models = trainer.train_models(X_train, y_train)
        
        self.assertGreater(len(trained_models), 0)
        
        # Step 5: Model evaluation
        results = trainer.evaluate_models(X_test, y_test)
        
        self.assertEqual(len(results), len(trained_models))
        
        # Check that all models have reasonable performance
        for model_name, result in results.items():
            r2 = result['metrics']['r2']
            self.assertGreater(r2, -2.0)  # Allow for some negative R²
            self.assertLess(r2, 1.1)  # Should not exceed 1.0
    
    def test_pipeline_with_missing_data(self):
        """Test pipeline robustness with missing data."""
        # Introduce missing values
        molecular_data_missing = self.molecular_data.copy()
        molecular_data_missing.loc[0:10, 'molecular_weight'] = np.nan
        molecular_data_missing.loc[5:15, 'logp'] = np.nan
        
        # Test that pipeline handles missing data
        preprocessor = DataPreprocessor()
        best_affinities = preprocessor.get_best_affinities(self.binding_data)
        integrated_data = preprocessor.integrate_datasets(
            molecular_data_missing, best_affinities
        )
        
        # Should still work with missing data
        self.assertGreater(len(integrated_data), 0)
        
        # Feature engineering should handle missing values
        feature_cols = [col for col in integrated_data.columns 
                       if col not in ['compound_name', 'affinity']]
        X = integrated_data[feature_cols]
        y = integrated_data['affinity']
        
        feature_pipeline = FeatureEngineeringPipeline()
        X_processed = feature_pipeline.fit_transform(X, y)
        
        # Should not have NaN values after processing
        self.assertFalse(X_processed.isnull().any().any())
    
    def test_pipeline_with_outliers(self):
        """Test pipeline robustness with outliers."""
        # Introduce outliers
        binding_data_outliers = self.binding_data.copy()
        binding_data_outliers.loc[0, 'affinity'] = -20  # Extreme outlier
        binding_data_outliers.loc[1, 'affinity'] = 10   # Positive outlier (unrealistic)
        
        # Test that pipeline handles outliers
        preprocessor = DataPreprocessor()
        best_affinities = preprocessor.get_best_affinities(binding_data_outliers)
        
        # Should filter out unrealistic values
        self.assertLess(best_affinities['affinity'].max(), 0)  # All affinities should be negative
        self.assertGreater(best_affinities['affinity'].min(), -15)  # Not too extreme
    
    def test_model_persistence(self):
        """Test model saving and loading."""
        # Prepare data
        preprocessor = DataPreprocessor()
        best_affinities = preprocessor.get_best_affinities(self.binding_data)
        integrated_data = preprocessor.integrate_datasets(
            self.molecular_data, best_affinities
        )
        
        feature_cols = [col for col in integrated_data.columns 
                       if col not in ['compound_name', 'affinity']]
        X = integrated_data[feature_cols]
        y = integrated_data['affinity']
        
        feature_pipeline = FeatureEngineeringPipeline()
        X_processed = feature_pipeline.fit_transform(X, y)
        
        # Split data
        from sklearn.model_selection import train_test_split
        X_train, X_test, y_train, y_test = train_test_split(
            X_processed, y, test_size=0.2, random_state=42
        )
        
        # Train models
        trainer = MLModelTrainer(models_dir=self.temp_dir)
        trained_models = trainer.train_models(X_train, y_train)
        
        # Test model loading
        for model_name in trained_models.keys():
            loaded_model = trainer.load_model(model_name)
            
            # Test that loaded model works
            original_pred = trained_models[model_name].predict(X_test.iloc[:5])
            loaded_pred = loaded_model.predict(X_test.iloc[:5])
            
            np.testing.assert_array_almost_equal(original_pred, loaded_pred)
    
    def test_cross_validation_consistency(self):
        """Test that cross-validation results are consistent."""
        # Prepare data
        preprocessor = DataPreprocessor()
        best_affinities = preprocessor.get_best_affinities(self.binding_data)
        integrated_data = preprocessor.integrate_datasets(
            self.molecular_data, best_affinities
        )
        
        feature_cols = [col for col in integrated_data.columns 
                       if col not in ['compound_name', 'affinity']]
        X = integrated_data[feature_cols]
        y = integrated_data['affinity']
        
        feature_pipeline = FeatureEngineeringPipeline()
        X_processed = feature_pipeline.fit_transform(X, y)
        
        # Test multiple runs with same random state
        trainer1 = MLModelTrainer(models_dir=self.temp_dir)
        trainer2 = MLModelTrainer(models_dir=self.temp_dir)
        
        # Split data consistently
        from sklearn.model_selection import train_test_split
        X_train, X_test, y_train, y_test = train_test_split(
            X_processed, y, test_size=0.2, random_state=42
        )
        
        # Train models
        models1 = trainer1.train_models(X_train, y_train)
        models2 = trainer2.train_models(X_train, y_train)
        
        # Results should be similar
        results1 = trainer1.evaluate_models(X_test, y_test)
        results2 = trainer2.evaluate_models(X_test, y_test)
        
        for model_name in models1.keys():
            if model_name in results1 and model_name in results2:
                r2_1 = results1[model_name]['metrics']['r2']
                r2_2 = results2[model_name]['metrics']['r2']
                self.assertAlmostEqual(r2_1, r2_2, places=2)


class TestMLPipelineRobustness(unittest.TestCase):
    """Test ML pipeline robustness and error handling."""
    
    def setUp(self):
        """Set up test data."""
        np.random.seed(42)
        self.n_samples = 50
        self.n_features = 10
        
        # Create minimal dataset
        X = np.random.randn(self.n_samples, self.n_features)
        y = np.random.randn(self.n_samples)
        
        self.X = pd.DataFrame(X, columns=[f'feature_{i}' for i in range(self.n_features)])
        self.y = pd.Series(y)
    
    def test_pipeline_with_empty_data(self):
        """Test pipeline behavior with empty data."""
        trainer = MLModelTrainer()
        
        # Test with empty training data
        empty_X = pd.DataFrame()
        empty_y = pd.Series(dtype=float)
        
        with self.assertRaises((ValueError, IndexError)):
            trainer.train_models(empty_X, empty_y)
    
    def test_pipeline_with_single_sample(self):
        """Test pipeline behavior with single sample."""
        trainer = MLModelTrainer()
        
        # Test with single sample
        single_X = self.X.iloc[:1]
        single_y = self.y.iloc[:1]
        
        # Should handle single sample gracefully or raise informative error
        try:
            models = trainer.train_models(single_X, single_y)
            # If it succeeds, check that models are created
            self.assertGreaterEqual(len(models), 0)
        except Exception as e:
            # Should raise informative error
            self.assertIn("sample", str(e).lower())
    
    def test_pipeline_with_constant_features(self):
        """Test pipeline behavior with constant features."""
        # Create data with constant features
        X_constant = self.X.copy()
        X_constant['constant_feature'] = 1.0  # All values are 1.0
        X_constant['another_constant'] = 0.0  # All values are 0.0
        
        trainer = MLModelTrainer()
        
        # Should handle constant features
        try:
            models = trainer.train_models(X_constant, self.y)
            self.assertGreaterEqual(len(models), 0)
        except Exception as e:
            # Should handle constant features gracefully
            self.assertNotIn("constant", str(e).lower())
    
    def test_pipeline_with_high_dimensional_data(self):
        """Test pipeline behavior with high-dimensional data."""
        # Create high-dimensional data
        n_features = 1000
        X_high_dim = pd.DataFrame(
            np.random.randn(self.n_samples, n_features),
            columns=[f'feature_{i}' for i in range(n_features)]
        )
        
        trainer = MLModelTrainer()
        
        # Should handle high-dimensional data
        try:
            models = trainer.train_models(X_high_dim, self.y)
            self.assertGreaterEqual(len(models), 0)
        except Exception as e:
            # Should handle high dimensions or raise informative error
            self.assertIsInstance(e, (ValueError, MemoryError, RuntimeError))


if __name__ == '__main__':
    # Run tests with verbose output
    unittest.main(verbosity=2)
```
