## Supplementary material for "Machine Learning-Driven Prediction of TLR4 Binding Affinity: A Comprehensive Molecular Feature Analysis for Drug Discovery": Codebase: test_molecular_feature_extractor_integration.py

```
"""
Integration tests for the comprehensive molecular feature extractor.

Tests the MolecularFeatureExtractor with actual PDBQT files from the binding-data directory
to ensure end-to-end functionality and performance.
"""

import pytest
import pandas as pd
import numpy as np
from pathlib import Path
import tempfile
import json
import logging

from src.tlr4_binding.molecular_analysis.extractor import MolecularFeatureExtractor
from src.tlr4_binding.molecular_analysis.features import MolecularFeatures

# Set up logging for tests
logging.basicConfig(level=logging.INFO)
logger = logging.getLogger(__name__)


class TestMolecularFeatureExtractorIntegration:
    """Integration tests for MolecularFeatureExtractor."""
    
    @pytest.fixture
    def extractor(self):
        """Create MolecularFeatureExtractor instance for testing."""
        return MolecularFeatureExtractor(
            include_2d_features=True,
            include_3d_features=True,
            include_advanced_features=True
        )
    
    @pytest.fixture
    def sample_pdbqt_files(self):
        """Get sample PDBQT files from binding-data directory."""
        pdbqt_dir = Path("binding-data/raw/pdbqt")
        if not pdbqt_dir.exists():
            pytest.skip("PDBQT directory not found")
        
        # Get a few sample files for testing
        pdbqt_files = list(pdbqt_dir.glob("*.pdbqt"))[:5]
        if not pdbqt_files:
            pytest.skip("No PDBQT files found for testing")
        
        return [str(f) for f in pdbqt_files]
    
    def test_single_file_extraction(self, extractor, sample_pdbqt_files):
        """Test extraction from a single PDBQT file."""
        pdbqt_file = sample_pdbqt_files[0]
        logger.info(f"Testing single file extraction: {pdbqt_file}")
        
        # Extract features
        features = extractor.extract_features(pdbqt_file)
        
        # Validate MolecularFeatures object
        assert isinstance(features, MolecularFeatures)
        assert features.compound_name is not None
        assert features.pdbqt_file == pdbqt_file
        
        # Check that basic features are present
        assert not np.isnan(features.molecular_weight) or features.molecular_weight is not None
        assert not np.isnan(features.logp) or features.logp is not None
        assert not np.isnan(features.tpsa) or features.tpsa is not None
        
        # Check statistics were updated
        stats = extractor.get_extraction_stats()
        assert stats['successful_extractions'] == 1
        assert stats['total_files_processed'] == 1
        assert len(stats['processing_times']) == 1
        
        logger.info(f"Successfully extracted features for {features.compound_name}")
    
    def test_batch_extraction(self, extractor, sample_pdbqt_files):
        """Test batch extraction from multiple PDBQT files."""
        logger.info(f"Testing batch extraction with {len(sample_pdbqt_files)} files")
        
        # Create temporary directory with sample files
        with tempfile.TemporaryDirectory() as temp_dir:
            temp_path = Path(temp_dir)
            
            # Copy sample files to temp directory
            for pdbqt_file in sample_pdbqt_files:
                src = Path(pdbqt_file)
                dst = temp_path / src.name
                dst.write_text(src.read_text())
            
            # Reset statistics
            extractor.reset_stats()
            
            # Perform batch extraction
            features_df = extractor.batch_extract(str(temp_path))
            
            # Validate results
            assert isinstance(features_df, pd.DataFrame)
            assert len(features_df) == len(sample_pdbqt_files)
            assert 'compound_name' in features_df.columns
            assert 'molecular_weight' in features_df.columns
            
            # Check statistics
            stats = extractor.get_extraction_stats()
            assert stats['total_files_processed'] == len(sample_pdbqt_files)
            assert stats['successful_extractions'] == len(sample_pdbqt_files)
            assert len(stats['processing_times']) == len(sample_pdbqt_files)
            
            logger.info(f"Batch extraction completed successfully for {len(features_df)} files")
    
    def test_extract_from_list(self, extractor, sample_pdbqt_files):
        """Test extraction from a list of PDBQT files."""
        logger.info(f"Testing extraction from file list with {len(sample_pdbqt_files)} files")
        
        # Reset statistics
        extractor.reset_stats()
        
        # Extract features from list
        features_df = extractor.extract_features_from_list(sample_pdbqt_files)
        
        # Validate results
        assert isinstance(features_df, pd.DataFrame)
        assert len(features_df) == len(sample_pdbqt_files)
        
        # Check that all expected columns are present
        expected_columns = [
            'compound_name', 'molecular_weight', 'logp', 'tpsa',
            'rotatable_bonds', 'hbd', 'hba', 'formal_charge',
            'radius_of_gyration', 'molecular_volume', 'surface_area', 'asphericity'
        ]
        
        for col in expected_columns:
            assert col in features_df.columns, f"Missing expected column: {col}"
        
        # Check statistics
        stats = extractor.get_extraction_stats()
        assert stats['total_files_processed'] == len(sample_pdbqt_files)
        
        logger.info(f"List extraction completed successfully for {len(features_df)} files")
    
    def test_feature_validation(self, extractor, sample_pdbqt_files):
        """Test feature validation functionality."""
        logger.info("Testing feature validation")
        
        # Extract features from a few files
        features_df = extractor.extract_features_from_list(sample_pdbqt_files[:3])
        
        # Validate features
        validation_results = extractor.validate_features(features_df)
        
        # Check validation results structure
        assert isinstance(validation_results, dict)
        expected_keys = ['missing_values', 'infinite_values', 'outliers', 'invalid_ranges']
        for key in expected_keys:
            assert key in validation_results
            assert isinstance(validation_results[key], list)
        
        logger.info(f"Feature validation completed. Issues found: {len(sum(validation_results.values(), []))}")
    
    def test_feature_summary(self, extractor, sample_pdbqt_files):
        """Test feature summary generation."""
        logger.info("Testing feature summary generation")
        
        # Extract features
        features_df = extractor.extract_features_from_list(sample_pdbqt_files[:3])
        
        # Get feature summary
        summary_df = extractor.get_feature_summary(features_df)
        
        # Validate summary
        assert isinstance(summary_df, pd.DataFrame)
        assert 'feature' in summary_df.columns
        assert 'count' in summary_df.columns
        assert 'mean' in summary_df.columns
        assert 'std' in summary_df.columns
        assert 'min' in summary_df.columns
        assert 'max' in summary_df.columns
        
        # Check that summary includes numerical features
        assert len(summary_df) > 0
        
        logger.info(f"Feature summary generated for {len(summary_df)} features")
    
    def test_performance_reporting(self, extractor, sample_pdbqt_files):
        """Test performance reporting functionality."""
        logger.info("Testing performance reporting")
        
        # Reset statistics
        extractor.reset_stats()
        
        # Extract features from multiple files
        features_df = extractor.extract_features_from_list(sample_pdbqt_files)
        
        # Get performance report
        performance_report = extractor.get_performance_report()
        
        # Validate report structure
        assert isinstance(performance_report, dict)
        assert 'batch_summary' in performance_report
        
        batch_summary = performance_report['batch_summary']
        assert 'total_files' in batch_summary
        assert 'successful' in batch_summary
        assert 'failed' in batch_summary
        assert 'success_rate' in batch_summary
        
        # Check performance metrics
        assert batch_summary['total_files'] == len(sample_pdbqt_files)
        assert batch_summary['successful'] == len(features_df)
        
        # Test efficiency metrics
        efficiency = extractor.get_feature_extraction_efficiency()
        assert isinstance(efficiency, dict)
        
        logger.info(f"Performance report generated. Success rate: {batch_summary['success_rate']:.1f}%")
    
    def test_performance_report_saving(self, extractor, sample_pdbqt_files):
        """Test saving performance report to file."""
        logger.info("Testing performance report saving")
        
        # Extract features
        extractor.extract_features_from_list(sample_pdbqt_files[:2])
        
        # Save performance report
        with tempfile.NamedTemporaryFile(mode='w', suffix='.json', delete=False) as temp_file:
            temp_path = temp_file.name
        
        try:
            extractor.save_performance_report(temp_path)
            
            # Verify file was created and contains valid JSON
            assert Path(temp_path).exists()
            
            with open(temp_path, 'r') as f:
                saved_report = json.load(f)
            
            assert isinstance(saved_report, dict)
            assert 'batch_summary' in saved_report
            
            logger.info("Performance report saved successfully")
            
        finally:
            # Clean up temporary file
            Path(temp_path).unlink(missing_ok=True)
    
    def test_error_handling(self, extractor):
        """Test error handling with invalid files."""
        logger.info("Testing error handling")
        
        # Test with non-existent file
        with pytest.raises(ValueError):
            extractor.extract_features("non_existent_file.pdbqt")
        
        # Test with invalid directory
        with pytest.raises(ValueError):
            extractor.batch_extract("non_existent_directory")
        
        # Create empty directory
        with tempfile.TemporaryDirectory() as temp_dir:
            with pytest.raises(ValueError):
                extractor.batch_extract(temp_dir)
        
        logger.info("Error handling tests passed")
    
    def test_feature_extraction_modes(self):
        """Test different feature extraction modes."""
        logger.info("Testing different feature extraction modes")
        
        # Test with only 2D features
        extractor_2d = MolecularFeatureExtractor(
            include_2d_features=True,
            include_3d_features=False,
            include_advanced_features=True
        )
        
        # Test with only 3D features
        extractor_3d = MolecularFeatureExtractor(
            include_2d_features=False,
            include_3d_features=True,
            include_advanced_features=False
        )
        
        # Test with minimal features
        extractor_minimal = MolecularFeatureExtractor(
            include_2d_features=False,
            include_3d_features=False,
            include_advanced_features=False
        )
        
        assert extractor_2d.include_2d_features is True
        assert extractor_2d.include_3d_features is False
        
        assert extractor_3d.include_2d_features is False
        assert extractor_3d.include_3d_features is True
        
        assert extractor_minimal.include_2d_features is False
        assert extractor_minimal.include_3d_features is False
        
        logger.info("Feature extraction modes tested successfully")
    
    def test_memory_and_performance_tracking(self, extractor, sample_pdbqt_files):
        """Test memory usage and performance tracking."""
        logger.info("Testing memory and performance tracking")
        
        # Reset statistics
        extractor.reset_stats()
        
        # Extract features from multiple files
        features_df = extractor.extract_features_from_list(sample_pdbqt_files[:3])
        
        # Check performance tracking
        stats = extractor.get_extraction_stats()
        
        # Verify timing data
        assert len(stats['processing_times']) == len(sample_pdbqt_files[:3])
        assert len(stats['memory_usage']) == len(sample_pdbqt_files[:3])
        
        # Verify feature extraction timing breakdown
        feature_times = stats['feature_extraction_times']
        assert '2d_features' in feature_times
        assert '3d_features' in feature_times
        assert 'parsing' in feature_times
        
        # All timing lists should have the same length
        for timing_list in feature_times.values():
            assert len(timing_list) == len(sample_pdbqt_files[:3])
        
        # Check that timing values are positive
        for timing_list in stats['processing_times']:
            assert timing_list > 0
        
        logger.info("Memory and performance tracking verified")


if __name__ == "__main__":
    # Run integration tests
    pytest.main([__file__, "-v", "--tb=short"])
```
