## Supplementary material for "Machine Learning-Driven Prediction of TLR4 Binding Affinity: A Comprehensive Molecular Feature Analysis for Drug Discovery": Codebase: README.md

# Test Organization

This directory contains all test files for the TLR4 Binding Prediction project, organized into unit tests and integration tests.

## Directory Structure

```
tests/
├── __init__.py
├── test_basic.py                    # Basic functionality tests
├── unit/                           # Unit tests for individual components
│   ├── test_3d_features.py         # 3D feature extraction unit tests
│   ├── test_3d_simple.py           # Simple 3D feature tests
│   ├── test_3d_standalone.py       # Standalone 3D feature tests
│   ├── test_extractor_simple.py    # Simple extractor functionality tests
│   ├── test_extractor_standalone.py # Standalone extractor tests
│   ├── test_molecular_descriptors.py # Molecular descriptor unit tests
│   ├── test_molecular_descriptors_minimal.py # Minimal descriptor tests
│   ├── test_pdbqt_parser.py        # PDBQT parser unit tests
│   ├── test_pdbqt_parser_simple.py # Simple parser tests
│   ├── test_pdbqt_parser_standalone.py # Standalone parser tests
│   ├── test_structural_features.py # Structural feature unit tests
│   ├── test_structural_standalone.py # Standalone structural tests
│   └── standalone_pdbqt_parser.py  # Standalone parser utility
└── integration/                    # Integration tests for full workflows
    ├── test_feature_extractor_integration.py # Feature extractor integration tests
    └── test_molecular_feature_extractor_integration.py # Comprehensive integration tests
```

## Test Categories

### Unit Tests (`tests/unit/`)
Unit tests focus on testing individual components in isolation:
- **PDBQT Parser Tests**: Test file parsing functionality
- **Molecular Descriptor Tests**: Test 2D molecular property calculations
- **3D Feature Tests**: Test 3D structural feature extraction
- **Structural Feature Tests**: Test PyMOL-based structural analysis
- **Extractor Tests**: Test individual extractor methods

### Integration Tests (`tests/integration/`)
Integration tests verify that components work together correctly:
- **Feature Extractor Integration**: End-to-end feature extraction workflows
- **Molecular Feature Extractor Integration**: Comprehensive pipeline testing

## Running Tests

### Run All Tests
```bash
python -m pytest tests/ -v
```

### Run Unit Tests Only
```bash
python -m pytest tests/unit/ -v
```

### Run Integration Tests Only
```bash
python -m pytest tests/integration/ -v
```

### Run Specific Test File
```bash
python -m pytest tests/unit/test_pdbqt_parser.py -v
```

## Test Naming Conventions

- **`test_*.py`**: Standard pytest test files
- **`*_simple.py`**: Simplified tests that don't require full dependencies
- **`*_standalone.py`**: Tests that can run independently without package imports
- **`*_integration.py`**: Tests that verify component integration

## Validation Scripts

The `validate_extractor.py` script in the project root is a utility for validating implementation completeness and is not a test file.

## Notes

- All test files follow pytest conventions
- Tests are designed to be runnable even with missing optional dependencies (RDKit, PyMOL)
- Integration tests may require actual PDBQT files from the `binding-data/raw/pdbqt` directory
- Some tests include fallback mechanisms for environments with limited dependencies
