## Supplementary material for "Machine Learning-Driven Prediction of TLR4 Binding Affinity: A Comprehensive Molecular Feature Analysis for Drug Discovery": Codebase: standalone_evaluation_test.py

```
"""
Standalone test for evaluation framework components.

This script tests the evaluation framework components directly without importing
from the project modules to avoid dependency issues.
"""

import numpy as np
import pandas as pd
import matplotlib
matplotlib.use('Agg')  # Use non-interactive backend
import matplotlib.pyplot as plt
from sklearn.metrics import (
    mean_squared_error, mean_absolute_error, r2_score,
    explained_variance_score, max_error
)
from sklearn.model_selection import (
    learning_curve, validation_curve, cross_val_score,
    KFold
)
from scipy import stats
from sklearn.ensemble import RandomForestRegressor
from sklearn.linear_model import LinearRegression
from sklearn.svm import SVR
from sklearn.model_selection import train_test_split
from sklearn.datasets import make_regression
import tempfile
import os

class PerformanceMetrics:
    """Container for performance metrics and statistical analysis."""
    
    def __init__(self, y_true: np.ndarray, y_pred: np.ndarray, model_name: str = ""):
        """Initialize performance metrics."""
        self.y_true = y_true
        self.y_pred = y_pred
        self.model_name = model_name
        self.metrics = self._calculate_metrics()
    
    def _calculate_metrics(self) -> dict:
        """Calculate comprehensive regression metrics."""
        # Basic regression metrics
        mse = mean_squared_error(self.y_true, self.y_pred)
        rmse = np.sqrt(mse)
        mae = mean_absolute_error(self.y_true, self.y_pred)
        r2 = r2_score(self.y_true, self.y_pred)
        evs = explained_variance_score(self.y_true, self.y_pred)
        max_err = max_error(self.y_true, self.y_pred)
        
        # Additional metrics
        mape = self._calculate_mape(self.y_true, self.y_pred)
        smape = self._calculate_smape(self.y_true, self.y_pred)
        pearson_r, pearson_p = stats.pearsonr(self.y_true, self.y_pred)
        spearman_r, spearman_p = stats.spearmanr(self.y_true, self.y_pred)
        
        # Residual analysis
        residuals = self.y_true - self.y_pred
        residual_std = np.std(residuals)
        residual_mean = np.mean(residuals)
        
        return {
            'mse': mse,
            'rmse': rmse,
            'mae': mae,
            'r2': r2,
            'explained_variance': evs,
            'max_error': max_err,
            'mape': mape,
            'smape': smape,
            'pearson_r': pearson_r,
            'pearson_p': pearson_p,
            'spearman_r': spearman_r,
            'spearman_p': spearman_p,
            'residual_mean': residual_mean,
            'residual_std': residual_std,
            'residual_rmse': np.sqrt(np.mean(residuals**2))
        }
    
    def _calculate_mape(self, y_true: np.ndarray, y_pred: np.ndarray) -> float:
        """Calculate Mean Absolute Percentage Error."""
        mask = y_true != 0
        if not np.any(mask):
            return np.nan
        return np.mean(np.abs((y_true[mask] - y_pred[mask]) / y_true[mask])) * 100
    
    def _calculate_smape(self, y_true: np.ndarray, y_pred: np.ndarray) -> float:
        """Calculate Symmetric Mean Absolute Percentage Error."""
        denominator = (np.abs(y_true) + np.abs(y_pred)) / 2
        mask = denominator != 0
        if not np.any(mask):
            return np.nan
        return np.mean(np.abs(y_true[mask] - y_pred[mask]) / denominator[mask]) * 100
    
    def get_summary(self) -> dict:
        """Get summary of key metrics."""
        return {
            'model_name': self.model_name,
            'r2_score': self.metrics['r2'],
            'rmse': self.metrics['rmse'],
            'mae': self.metrics['mae'],
            'pearson_correlation': self.metrics['pearson_r'],
            'spearman_correlation': self.metrics['spearman_r']
        }
    
    def to_dataframe(self) -> pd.DataFrame:
        """Convert metrics to DataFrame."""
        return pd.DataFrame([self.metrics])

class ModelEvaluator:
    """Comprehensive model evaluation with statistical analysis."""
    
    def __init__(self, confidence_level: float = 0.95):
        """Initialize model evaluator."""
        self.confidence_level = confidence_level
        self.evaluation_history = []
    
    def evaluate(self, y_true: np.ndarray, y_pred: np.ndarray, 
                model_name: str = "") -> PerformanceMetrics:
        """Evaluate model performance comprehensively."""
        metrics = PerformanceMetrics(y_true, y_pred, model_name)
        self.evaluation_history.append({
            'timestamp': pd.Timestamp.now(),
            'model_name': model_name,
            'metrics': metrics
        })
        return metrics
    
    def compare_models(self, results: dict) -> pd.DataFrame:
        """Compare multiple model performances."""
        if not results:
            return pd.DataFrame()
        
        comparison_data = []
        for model_name, metrics in results.items():
            summary = metrics.get_summary()
            comparison_data.append(summary)
        
        comparison_df = pd.DataFrame(comparison_data)
        comparison_df = comparison_df.sort_values('r2_score', ascending=False)
        return comparison_df
    
    def statistical_significance_test(self, metrics1: PerformanceMetrics, 
                                   metrics2: PerformanceMetrics) -> dict:
        """Perform statistical significance test between two models."""
        # Paired t-test on residuals
        residuals1 = metrics1.y_true - metrics1.y_pred
        residuals2 = metrics2.y_true - metrics2.y_pred
        
        if len(residuals1) != len(residuals2):
            raise ValueError("Models must have same number of predictions for comparison")
        
        # Paired t-test
        t_stat, t_p = stats.ttest_rel(residuals1, residuals2)
        
        # Wilcoxon signed-rank test (non-parametric alternative)
        try:
            w_stat, w_p = stats.wilcoxon(residuals1, residuals2)
        except ValueError:
            w_stat, w_p = np.nan, np.nan
        
        return {
            'paired_t_test': {
                'statistic': t_stat,
                'p_value': t_p,
                'significant': t_p < (1 - self.confidence_level)
            },
            'wilcoxon_test': {
                'statistic': w_stat,
                'p_value': w_p,
                'significant': w_p < (1 - self.confidence_level) if not np.isnan(w_p) else False
            },
            'model1_r2': metrics1.metrics['r2'],
            'model2_r2': metrics2.metrics['r2'],
            'r2_difference': metrics1.metrics['r2'] - metrics2.metrics['r2']
        }
    
    def cross_validate_model(self, model, X: np.ndarray, y: np.ndarray,
                           model_name: str = "", cv: int = 5, 
                           scoring: list = None) -> dict:
        """Perform comprehensive cross-validation evaluation."""
        if scoring is None:
            scoring = ['neg_mean_squared_error', 'neg_mean_absolute_error', 'r2']
        
        cv_results = {}
        cv_fold = KFold(n_splits=cv, shuffle=True, random_state=42)
        
        for metric in scoring:
            scores = cross_val_score(model, X, y, cv=cv_fold, scoring=metric, n_jobs=-1)
            cv_results[metric] = {
                'scores': scores,
                'mean': np.mean(scores),
                'std': np.std(scores),
                'min': np.min(scores),
                'max': np.max(scores)
            }
        
        # Convert MSE and MAE to positive values for easier interpretation
        if 'neg_mean_squared_error' in cv_results:
            cv_results['rmse'] = {
                'scores': np.sqrt(-cv_results['neg_mean_squared_error']['scores']),
                'mean': np.sqrt(-cv_results['neg_mean_squared_error']['mean']),
                'std': cv_results['neg_mean_squared_error']['std'] / (2 * np.sqrt(-cv_results['neg_mean_squared_error']['mean'])),
                'min': np.sqrt(-cv_results['neg_mean_squared_error']['max']),
                'max': np.sqrt(-cv_results['neg_mean_squared_error']['min'])
            }
        
        if 'neg_mean_absolute_error' in cv_results:
            cv_results['mae'] = {
                'scores': -cv_results['neg_mean_absolute_error']['scores'],
                'mean': -cv_results['neg_mean_absolute_error']['mean'],
                'std': cv_results['neg_mean_absolute_error']['std'],
                'min': -cv_results['neg_mean_absolute_error']['max'],
                'max': -cv_results['neg_mean_absolute_error']['min']
            }
        
        return cv_results
    
    def generate_learning_curves(self, model, X: np.ndarray, y: np.ndarray,
                                model_name: str = "", cv: int = 5, 
                                train_sizes: list = None,
                                save_path: str = None) -> dict:
        """Generate learning curves for a model."""
        if train_sizes is None:
            train_sizes = np.linspace(0.1, 1.0, 10)
        
        # Generate learning curves
        train_sizes_abs, train_scores, val_scores = learning_curve(
            model, X, y, cv=cv, train_sizes=train_sizes,
            scoring='neg_mean_squared_error', n_jobs=-1
        )
        
        # Convert to positive RMSE
        train_rmse = np.sqrt(-train_scores)
        val_rmse = np.sqrt(-val_scores)
        
        # Calculate means and standard deviations
        train_mean = np.mean(train_rmse, axis=1)
        train_std = np.std(train_rmse, axis=1)
        val_mean = np.mean(val_rmse, axis=1)
        val_std = np.std(val_rmse, axis=1)
        
        # Plot learning curves
        plt.figure(figsize=(10, 6))
        plt.plot(train_sizes_abs, train_mean, 'o-', label='Training RMSE', color='blue')
        plt.fill_between(train_sizes_abs, train_mean - train_std, 
                        train_mean + train_std, alpha=0.1, color='blue')
        
        plt.plot(train_sizes_abs, val_mean, 'o-', label='Validation RMSE', color='red')
        plt.fill_between(train_sizes_abs, val_mean - val_std, 
                        val_mean + val_std, alpha=0.1, color='red')
        
        plt.xlabel('Training Set Size')
        plt.ylabel('RMSE')
        plt.title(f'Learning Curves - {model_name}')
        plt.legend()
        plt.grid(True, alpha=0.3)
        
        if save_path:
            plt.savefig(save_path, dpi=300, bbox_inches='tight')
        
        plt.close()  # Close figure to free memory
        
        return {
            'train_sizes': train_sizes_abs,
            'train_mean': train_mean,
            'train_std': train_std,
            'val_mean': val_mean,
            'val_std': val_std,
            'model_name': model_name
        }

def test_performance_metrics():
    """Test PerformanceMetrics class."""
    print("Testing PerformanceMetrics class...")
    
    # Test with perfect predictions
    y_true = np.array([1, 2, 3, 4, 5])
    y_pred_perfect = y_true.copy()
    metrics_perfect = PerformanceMetrics(y_true, y_pred_perfect, "Perfect Model")
    
    print(f"✓ Perfect prediction test")
    print(f"  R² Score: {metrics_perfect.metrics['r2']:.6f}")
    print(f"  RMSE: {metrics_perfect.metrics['rmse']:.6f}")
    
    # Test with realistic predictions
    y_pred_realistic = y_true + np.random.normal(0, 0.1, len(y_true))
    metrics_realistic = PerformanceMetrics(y_true, y_pred_realistic, "Realistic Model")
    
    print(f"✓ Realistic prediction test")
    print(f"  R² Score: {metrics_realistic.metrics['r2']:.4f}")
    print(f"  RMSE: {metrics_realistic.metrics['rmse']:.4f}")
    print(f"  Pearson r: {metrics_realistic.metrics['pearson_r']:.4f}")
    
    return True

def test_basic_evaluation():
    """Test basic model evaluation."""
    print("\nTesting basic model evaluation...")
    
    # Create synthetic data
    X, y = make_regression(n_samples=200, n_features=10, noise=0.1, random_state=42)
    X_train, X_test, y_train, y_test = train_test_split(X, y, test_size=0.3, random_state=42)
    
    # Train a model
    model = RandomForestRegressor(n_estimators=50, random_state=42)
    model.fit(X_train, y_train)
    y_pred = model.predict(X_test)
    
    # Test evaluator
    evaluator = ModelEvaluator()
    metrics = evaluator.evaluate(y_test, y_pred, "Random Forest")
    
    print(f"✓ Basic evaluation completed")
    print(f"  R² Score: {metrics.metrics['r2']:.4f}")
    print(f"  RMSE: {metrics.metrics['rmse']:.4f}")
    print(f"  MAE: {metrics.metrics['mae']:.4f}")
    
    return True

def test_cross_validation():
    """Test cross-validation functionality."""
    print("\nTesting cross-validation...")
    
    # Create synthetic data
    X, y = make_regression(n_samples=150, n_features=8, noise=0.1, random_state=42)
    
    # Test cross-validation
    evaluator = ModelEvaluator()
    model = RandomForestRegressor(n_estimators=30, random_state=42)
    
    cv_results = evaluator.cross_validate_model(model, X, y, "RF Test", cv=3)
    
    print(f"✓ Cross-validation completed")
    print(f"  Mean R²: {cv_results['r2']['mean']:.4f} ± {cv_results['r2']['std']:.4f}")
    print(f"  Mean RMSE: {cv_results['rmse']['mean']:.4f} ± {cv_results['rmse']['std']:.4f}")
    
    return True

def test_learning_curves():
    """Test learning curves generation."""
    print("\nTesting learning curves...")
    
    # Create synthetic data
    X, y = make_regression(n_samples=200, n_features=6, noise=0.1, random_state=42)
    
    # Test learning curves
    evaluator = ModelEvaluator()
    model = RandomForestRegressor(n_estimators=50, random_state=42)
    
    # Create temporary file for plot
    with tempfile.NamedTemporaryFile(suffix='.png', delete=True) as tmp_file:
        curve_data = evaluator.generate_learning_curves(
            model, X, y, "RF Learning Curves", cv=3,
            save_path=tmp_file.name
        )
    
    print(f"✓ Learning curves generated")
    print(f"  Training sizes: {len(curve_data['train_sizes'])}")
    print(f"  Final training RMSE: {curve_data['train_mean'][-1]:.4f}")
    print(f"  Final validation RMSE: {curve_data['val_mean'][-1]:.4f}")
    
    return True

def test_statistical_testing():
    """Test statistical significance testing."""
    print("\nTesting statistical significance testing...")
    
    # Create synthetic data
    X, y = make_regression(n_samples=100, n_features=5, noise=0.1, random_state=42)
    X_train, X_test, y_train, y_test = train_test_split(X, y, test_size=0.3, random_state=42)
    
    # Train two different models
    model1 = RandomForestRegressor(n_estimators=30, random_state=42)
    model2 = LinearRegression()
    
    model1.fit(X_train, y_train)
    model2.fit(X_train, y_train)
    
    y_pred1 = model1.predict(X_test)
    y_pred2 = model2.predict(X_test)
    
    # Test statistical significance
    evaluator = ModelEvaluator()
    metrics1 = evaluator.evaluate(y_test, y_pred1, "Random Forest")
    metrics2 = evaluator.evaluate(y_test, y_pred2, "Linear Regression")
    
    test_result = evaluator.statistical_significance_test(metrics1, metrics2)
    
    print(f"✓ Statistical significance test completed")
    print(f"  Paired t-test p-value: {test_result['paired_t_test']['p_value']:.4f}")
    print(f"  Significant difference: {test_result['paired_t_test']['significant']}")
    print(f"  R² difference: {test_result['r2_difference']:.4f}")
    
    return True

def test_model_comparison():
    """Test model comparison functionality."""
    print("\nTesting model comparison...")
    
    # Create synthetic data
    X, y = make_regression(n_samples=150, n_features=8, noise=0.1, random_state=42)
    X_train, X_test, y_train, y_test = train_test_split(X, y, test_size=0.3, random_state=42)
    
    # Train multiple models
    models = {
        'Random Forest': RandomForestRegressor(n_estimators=30, random_state=42),
        'Linear Regression': LinearRegression(),
        'SVR': SVR(kernel='rbf', C=1.0, gamma='scale')
    }
    
    evaluator = ModelEvaluator()
    results = {}
    
    for name, model in models.items():
        model.fit(X_train, y_train)
        y_pred = model.predict(X_test)
        metrics = evaluator.evaluate(y_test, y_pred, name)
        results[name] = metrics
    
    # Compare models
    comparison_df = evaluator.compare_models(results)
    
    print(f"✓ Model comparison completed")
    print(f"  Models compared: {len(comparison_df)}")
    print(f"  Best model: {comparison_df.iloc[0]['model_name']}")
    print(f"  Best R²: {comparison_df.iloc[0]['r2_score']:.4f}")
    
    return True

def main():
    """Run all tests."""
    print("=" * 60)
    print("STANDALONE EVALUATION FRAMEWORK TEST")
    print("=" * 60)
    
    tests = [
        test_performance_metrics,
        test_basic_evaluation,
        test_cross_validation,
        test_learning_curves,
        test_statistical_testing,
        test_model_comparison
    ]
    
    passed = 0
    total = len(tests)
    
    for test_func in tests:
        try:
            if test_func():
                passed += 1
            else:
                print(f"✗ {test_func.__name__} failed")
        except Exception as e:
            print(f"✗ {test_func.__name__} failed with error: {e}")
            import traceback
            traceback.print_exc()
    
    print("\n" + "=" * 60)
    print(f"TEST RESULTS: {passed}/{total} tests passed")
    print("=" * 60)
    
    if passed == total:
        print("✓ All tests passed! The evaluation framework is working correctly.")
        return True
    else:
        print(f"✗ {total - passed} tests failed. Please check the implementation.")
        return False

if __name__ == "__main__":
    success = main()
    exit(0 if success else 1)
```
