## Supplementary material for "Machine Learning-Driven Prediction of TLR4 Binding Affinity: A Comprehensive Molecular Feature Analysis for Drug Discovery": Codebase: test_ablation_study.py

```
"""
Unit tests for TLR4 Binding Prediction Ablation Study Framework

Tests the comprehensive ablation study functionality including:
- Feature ablation studies
- Data size ablation studies
- Hyperparameter sensitivity analysis
- Statistical significance testing
- Report generation
"""

import pytest
import numpy as np
import pandas as pd
from unittest.mock import Mock, patch, MagicMock
import tempfile
import shutil
from pathlib import Path

from src.tlr4_binding.ml_components.ablation_study import (
    AblationStudyFramework,
    FeatureAblationStudy,
    DataSizeAblationStudy,
    HyperparameterAblationStudy,
    AblationConfig,
    AblationResult
)

class TestAblationConfig:
    """Test AblationConfig dataclass"""
    
    def test_default_config(self):
        """Test default configuration values"""
        config = AblationConfig()
        assert config.cv_folds == 5
        assert config.n_iterations == 10
        assert config.random_state == 42
        assert config.confidence_level == 0.95
        assert config.min_effect_size == 0.01
        assert config.parallel_jobs == -1
        assert config.save_models is False
        assert config.verbose is True
    
    def test_custom_config(self):
        """Test custom configuration values"""
        config = AblationConfig(
            cv_folds=3,
            n_iterations=5,
            random_state=123,
            confidence_level=0.99
        )
        assert config.cv_folds == 3
        assert config.n_iterations == 5
        assert config.random_state == 123
        assert config.confidence_level == 0.99

class TestAblationResult:
    """Test AblationResult dataclass"""
    
    def test_ablation_result_creation(self):
        """Test creating AblationResult instance"""
        result = AblationResult(
            component_name="test_component",
            baseline_score=0.8,
            ablated_score=0.7,
            score_difference=0.1,
            relative_change=0.125,
            p_value=0.05,
            confidence_interval=(0.05, 0.15),
            metadata={"test": "data"}
        )
        
        assert result.component_name == "test_component"
        assert result.baseline_score == 0.8
        assert result.ablated_score == 0.7
        assert result.score_difference == 0.1
        assert result.relative_change == 0.125
        assert result.p_value == 0.05
        assert result.confidence_interval == (0.05, 0.15)
        assert result.metadata == {"test": "data"}

class TestFeatureAblationStudy:
    """Test FeatureAblationStudy class"""
    
    def setup_method(self):
        """Set up test fixtures"""
        self.config = AblationConfig(cv_folds=3, random_state=42)
        self.study = FeatureAblationStudy(self.config)
        
        # Create mock data
        np.random.seed(42)
        self.X = pd.DataFrame({
            'feature_1': np.random.normal(0, 1, 100),
            'feature_2': np.random.normal(0, 1, 100),
            'feature_3': np.random.normal(0, 1, 100),
            'feature_4': np.random.normal(0, 1, 100)
        })
        self.y = pd.Series(np.random.normal(0, 1, 100))
        
        # Mock model trainer
        self.mock_trainer = Mock()
        self.mock_model = Mock()
        self.mock_trainer.train_single_model.return_value = self.mock_model
        self.mock_model.predict.return_value = np.random.normal(0, 1, 10)
    
    def test_feature_ablation_initialization(self):
        """Test FeatureAblationStudy initialization"""
        assert self.study.config == self.config
        assert isinstance(self.study.evaluator, Mock)  # Mocked in setup
        assert self.study.results == []
    
    @patch('src.tlr4_binding.ml_components.ablation_study.r2_score')
    def test_get_baseline_performance(self, mock_r2_score):
        """Test baseline performance calculation"""
        mock_r2_score.return_value = 0.8
        
        scores = self.study._get_baseline_performance(self.X, self.y, self.mock_trainer)
        
        assert len(scores) == self.config.cv_folds
        assert all(score == 0.8 for score in scores)
        assert self.mock_trainer.train_single_model.call_count == self.config.cv_folds
    
    def test_calculate_ablation_stats(self):
        """Test statistical calculation for ablation results"""
        baseline_scores = [0.8, 0.75, 0.82, 0.78, 0.79]
        ablated_scores = [0.7, 0.68, 0.72, 0.71, 0.69]
        metadata = {"test": "data"}
        
        result = self.study._calculate_ablation_stats(
            "test_component", baseline_scores, ablated_scores, metadata
        )
        
        assert result.component_name == "test_component"
        assert abs(result.baseline_score - 0.788) < 0.01  # Mean of baseline_scores
        assert abs(result.ablated_score - 0.7) < 0.01     # Mean of ablated_scores
        assert abs(result.score_difference - 0.088) < 0.01
        assert result.p_value is not None
        assert result.confidence_interval is not None
        assert result.metadata == metadata
    
    @patch('src.tlr4_binding.ml_components.ablation_study.r2_score')
    def test_feature_ablation_by_groups(self, mock_r2_score):
        """Test feature ablation by groups"""
        mock_r2_score.return_value = 0.8
        
        feature_groups = {
            "group_1": ["feature_1", "feature_2"],
            "group_2": ["feature_3"]
        }
        
        results = self.study.run_feature_ablation(
            self.X, self.y, self.mock_trainer, feature_groups
        )
        
        assert len(results) == 2
        assert results[0].component_name == "group_1"
        assert results[1].component_name == "group_2"
        
        # Check that features were actually removed
        assert self.mock_trainer.train_single_model.call_count > 0
    
    @patch('src.tlr4_binding.ml_components.ablation_study.r2_score')
    def test_forward_feature_selection(self, mock_r2_score):
        """Test forward feature selection"""
        mock_r2_score.return_value = 0.8
        
        results = self.study.run_sequential_feature_ablation(
            self.X, self.y, self.mock_trainer, method="forward"
        )
        
        assert len(results) == len(self.X.columns)
        # Check that features are added incrementally
        for i, result in enumerate(results):
            assert len(result.metadata["selected_features"]) == i + 1

class TestDataSizeAblationStudy:
    """Test DataSizeAblationStudy class"""
    
    def setup_method(self):
        """Set up test fixtures"""
        self.config = AblationConfig(cv_folds=3, random_state=42)
        self.study = DataSizeAblationStudy(self.config)
        
        # Create mock data
        np.random.seed(42)
        self.X = pd.DataFrame(np.random.normal(0, 1, (100, 10)))
        self.y = pd.Series(np.random.normal(0, 1, 100))
        
        # Mock model trainer
        self.mock_trainer = Mock()
        self.mock_model = Mock()
        self.mock_trainer.train_single_model.return_value = self.mock_model
        self.mock_model.predict.return_value = np.random.normal(0, 1, 10)
    
    def test_data_size_ablation_initialization(self):
        """Test DataSizeAblationStudy initialization"""
        assert self.study.config == self.config
        assert self.study.results == []
    
    @patch('src.tlr4_binding.ml_components.ablation_study.r2_score')
    def test_data_size_ablation(self, mock_r2_score):
        """Test data size ablation study"""
        mock_r2_score.return_value = 0.8
        
        sample_sizes = [20, 50, 80]
        results = self.study.run_data_size_ablation(
            self.X, self.y, self.mock_trainer, sample_sizes
        )
        
        assert len(results) == len(sample_sizes)
        
        for i, result in enumerate(results):
            assert result.component_name == f"data_size_{sample_sizes[i]}"
            assert result.metadata["sample_size"] == sample_sizes[i]
    
    def test_default_sample_sizes(self):
        """Test default sample size generation"""
        results = self.study.run_data_size_ablation(
            self.X, self.y, self.mock_trainer
        )
        
        # Should generate multiple sample sizes based on data size
        assert len(results) > 3
        assert all("data_size_" in result.component_name for result in results)

class TestHyperparameterAblationStudy:
    """Test HyperparameterAblationStudy class"""
    
    def setup_method(self):
        """Set up test fixtures"""
        self.config = AblationConfig(cv_folds=3, random_state=42)
        self.study = HyperparameterAblationStudy(self.config)
        
        # Create mock data
        np.random.seed(42)
        self.X = pd.DataFrame(np.random.normal(0, 1, (100, 10)))
        self.y = pd.Series(np.random.normal(0, 1, 100))
        
        # Mock model trainer
        self.mock_trainer = Mock()
        self.mock_model = Mock()
        self.mock_trainer.train_single_model.return_value = self.mock_model
        self.mock_model.predict.return_value = np.random.normal(0, 1, 10)
    
    def test_hyperparameter_ablation_initialization(self):
        """Test HyperparameterAblationStudy initialization"""
        assert self.study.config == self.config
        assert self.study.results == []
    
    @patch('src.tlr4_binding.ml_components.ablation_study.r2_score')
    def test_hyperparameter_ablation(self, mock_r2_score):
        """Test hyperparameter ablation study"""
        mock_r2_score.return_value = 0.8
        
        param_grids = {
            "n_estimators": [50, 100],
            "max_depth": [3, 5]
        }
        
        results = self.study.run_hyperparameter_ablation(
            self.X, self.y, self.mock_trainer, param_grids
        )
        
        # Should test 2 parameters with 2 values each = 4 experiments
        assert len(results) == 4
        
        # Check that parameters are tested
        param_names = set(result.metadata["parameter"] for result in results)
        assert param_names == {"n_estimators", "max_depth"}

class TestAblationStudyFramework:
    """Test main AblationStudyFramework class"""
    
    def setup_method(self):
        """Set up test fixtures"""
        self.config = AblationConfig(cv_folds=3, random_state=42)
        self.framework = AblationStudyFramework(self.config)
        
        # Create mock data
        np.random.seed(42)
        self.X = pd.DataFrame(np.random.normal(0, 1, (100, 10)))
        self.y = pd.Series(np.random.normal(0, 1, 100))
        
        # Mock model trainer
        self.mock_trainer = Mock()
        self.mock_model = Mock()
        self.mock_trainer.train_single_model.return_value = self.mock_model
        self.mock_model.predict.return_value = np.random.normal(0, 1, 10)
    
    def test_framework_initialization(self):
        """Test framework initialization"""
        assert self.framework.config == self.config
        assert isinstance(self.framework.feature_study, FeatureAblationStudy)
        assert isinstance(self.framework.architecture_study, ArchitectureAblationStudy)
        assert isinstance(self.framework.data_size_study, DataSizeAblationStudy)
        assert isinstance(self.framework.hyperparameter_study, HyperparameterAblationStudy)
        assert self.framework.all_results == []
    
    @patch('src.tlr4_binding.ml_components.ablation_study.r2_score')
    def test_comprehensive_ablation(self, mock_r2_score):
        """Test comprehensive ablation study"""
        mock_r2_score.return_value = 0.8
        
        results = self.framework.run_comprehensive_ablation(
            self.X, self.y, self.mock_trainer,
            study_types=["feature", "data_size"]
        )
        
        assert "feature" in results
        assert "data_size" in results
        assert len(results["feature"]) > 0
        assert len(results["data_size"]) > 0
        assert len(self.framework.all_results) > 0
    
    def test_generate_summary(self):
        """Test summary generation"""
        # Create mock results
        mock_results = {
            "feature": [
                AblationResult("feat1", 0.8, 0.7, 0.1, 0.125, 0.05),
                AblationResult("feat2", 0.8, 0.75, 0.05, 0.0625, 0.15)
            ],
            "data_size": [
                AblationResult("size1", 0.8, 0.7, 0.1, 0.125, 0.05)
            ]
        }
        
        summary = self.framework._generate_summary(mock_results)
        
        assert summary["total_studies"] == 2
        assert summary["total_experiments"] == 3
        assert summary["significant_effects"] == 2  # Two results with p < 0.05
    
    def test_analyze_feature_importance(self):
        """Test feature importance analysis"""
        feature_results = [
            AblationResult("feat1", 0.8, 0.6, 0.2, 0.25, 0.01),
            AblationResult("feat2", 0.8, 0.75, 0.05, 0.0625, 0.1),
            AblationResult("feat3", 0.8, 0.78, 0.02, 0.025, 0.3)
        ]
        
        analysis = self.framework._analyze_feature_importance(feature_results)
        
        assert len(analysis["most_important_features"]) == 3
        assert len(analysis["least_important_features"]) == 3
        assert "mean_impact" in analysis["feature_impact_distribution"]
        assert "std_impact" in analysis["feature_impact_distribution"]
        
        # Most important should be sorted by impact
        most_important = analysis["most_important_features"]
        assert most_important[0].component_name == "feat1"  # Highest impact
    
    def test_analyze_statistical_significance(self):
        """Test statistical significance analysis"""
        # Add some results to framework
        self.framework.all_results = [
            AblationResult("comp1", 0.8, 0.7, 0.1, 0.125, 0.01),  # Significant
            AblationResult("comp2", 0.8, 0.75, 0.05, 0.0625, 0.15),  # Not significant
            AblationResult("comp3", 0.8, 0.72, 0.08, 0.1, 0.03)   # Significant
        ]
        
        analysis = self.framework._analyze_statistical_significance()
        
        assert analysis["significant_results_count"] == 2
        assert analysis["total_results_count"] == 3
        assert abs(analysis["significance_rate"] - 2/3) < 0.01
    
    def test_generate_recommendations(self):
        """Test recommendation generation"""
        mock_results = {
            "feature": [
                AblationResult("feat1", 0.8, 0.6, 0.2, 0.25, 0.01)
            ],
            "data_size": [
                AblationResult("size1", 0.8, 0.7, 0.1, 0.125, 0.05)
            ],
            "hyperparameter": [
                AblationResult("param1", 0.8, 0.75, 0.05, 0.0625, 0.1)
            ]
        }
        
        recommendations = self.framework._generate_recommendations(mock_results)
        
        assert len(recommendations) > 0
        assert all(isinstance(rec, str) for rec in recommendations)
    
    def test_save_and_load_report(self):
        """Test saving and loading ablation reports"""
        # Create temporary directory
        with tempfile.TemporaryDirectory() as temp_dir:
            save_path = Path(temp_dir) / "test_report"
            
            # Create mock results
            mock_results = {
                "feature": [
                    AblationResult("feat1", 0.8, 0.7, 0.1, 0.125, 0.05)
                ]
            }
            
            # Add results to framework
            self.framework.all_results = mock_results["feature"]
            
            # Generate and save report
            report = self.framework.generate_ablation_report(mock_results, save_path)
            
            # Check that files were created
            assert (save_path / "ablation_report.json").exists()
            assert (save_path / "ablation_results.csv").exists()
            
            # Check report structure
            assert "summary" in report
            assert "feature_analysis" in report
            assert "statistical_significance" in report
            assert "recommendations" in report

class TestAblationStudyIntegration:
    """Integration tests for ablation study framework"""
    
    def setup_method(self):
        """Set up integration test fixtures"""
        self.config = AblationConfig(cv_folds=2, random_state=42)
        
        # Create realistic test data
        np.random.seed(42)
        n_samples = 50
        n_features = 20
        
        self.X = pd.DataFrame(
            np.random.normal(0, 1, (n_samples, n_features)),
            columns=[f"feature_{i}" for i in range(n_features)]
        )
        
        # Create target with some feature relationships
        self.y = pd.Series(
            self.X.iloc[:, :5].sum(axis=1) * 0.5 + np.random.normal(0, 0.1, n_samples)
        )
        
        # Mock model trainer that returns a simple model
        self.mock_trainer = Mock()
        self.mock_model = Mock()
        self.mock_trainer.train_single_model.return_value = self.mock_model
        
        # Mock model predictions with some correlation to target
        def mock_predict(X):
            return X.iloc[:, 0] * 0.5 + np.random.normal(0, 0.1, len(X))
        
        self.mock_model.predict.side_effect = mock_predict
    
    @patch('src.tlr4_binding.ml_components.ablation_study.r2_score')
    def test_end_to_end_ablation_study(self, mock_r2_score):
        """Test complete end-to-end ablation study"""
        # Mock R² score to return reasonable values
        mock_r2_score.return_value = 0.7
        
        framework = AblationStudyFramework(self.config)
        
        # Run comprehensive ablation study
        results = framework.run_comprehensive_ablation(
            self.X, self.y, self.mock_trainer,
            study_types=["feature", "data_size"]
        )
        
        # Verify results structure
        assert "feature" in results
        assert "data_size" in results
        assert len(results["feature"]) > 0
        assert len(results["data_size"]) > 0
        
        # Generate report
        report = framework.generate_ablation_report(results)
        
        # Verify report structure
        assert "summary" in report
        assert "feature_analysis" in report
        assert "data_size_analysis" in report
        assert "statistical_significance" in report
        assert "recommendations" in report
        
        # Verify summary statistics
        summary = report["summary"]
        assert summary["total_studies"] == 2
        assert summary["total_experiments"] > 0

if __name__ == "__main__":
    pytest.main([__file__, "-v"])
```
