## Supplementary material for "Machine Learning-Driven Prediction of TLR4 Binding Affinity: A Comprehensive Molecular Feature Analysis for Drug Discovery": Codebase: test_basic.py

```
"""
Basic tests for TLR4 Binding Affinity Prediction System.

This module contains basic functionality tests to ensure
the core components are working correctly.
"""

import unittest
import sys
from pathlib import Path
import pandas as pd
import numpy as np

# Add src to path
sys.path.insert(0, str(Path(__file__).parent.parent.parent / "src"))

from tlr4_binding.molecular_analysis.features import MolecularFeatures, BindingData, PredictionResult
from tlr4_binding.molecular_analysis.parser import PDBQTParser
from tlr4_binding.molecular_analysis.descriptors import MolecularDescriptorCalculator
from tlr4_binding.data_processing.preprocessor import DataPreprocessor, BindingDataLoader
from tlr4_binding.config import Config


class TestMolecularFeatures(unittest.TestCase):
    """Test MolecularFeatures data class."""
    
    def test_molecular_features_creation(self):
        """Test creating MolecularFeatures object."""
        features = MolecularFeatures(
            compound_name="test_compound",
            molecular_weight=200.0,
            logp=2.5,
            tpsa=50.0,
            rotatable_bonds=5,
            hbd=2,
            hba=4,
            formal_charge=0,
            radius_of_gyration=5.0,
            molecular_volume=150.0,
            surface_area=200.0,
            asphericity=0.3,
            ring_count=2,
            aromatic_rings=1,
            branching_index=0.5,
            dipole_moment=2.0,
            polarizability=25.0
        )
        
        self.assertEqual(features.compound_name, "test_compound")
        self.assertEqual(features.molecular_weight, 200.0)
        self.assertEqual(features.logp, 2.5)
    
    def test_molecular_features_to_dict(self):
        """Test converting MolecularFeatures to dictionary."""
        features = MolecularFeatures(
            compound_name="test_compound",
            molecular_weight=200.0,
            logp=2.5,
            tpsa=50.0,
            rotatable_bonds=5,
            hbd=2,
            hba=4,
            formal_charge=0,
            radius_of_gyration=5.0,
            molecular_volume=150.0,
            surface_area=200.0,
            asphericity=0.3,
            ring_count=2,
            aromatic_rings=1,
            branching_index=0.5,
            dipole_moment=2.0,
            polarizability=25.0
        )
        
        features_dict = features.to_dict()
        self.assertIsInstance(features_dict, dict)
        self.assertEqual(features_dict['compound_name'], "test_compound")
        self.assertEqual(features_dict['molecular_weight'], 200.0)


class TestBindingData(unittest.TestCase):
    """Test BindingData data class."""
    
    def test_binding_data_creation(self):
        """Test creating BindingData object."""
        binding = BindingData(
            ligand="test_ligand",
            mode=1,
            affinity=-7.5,
            rmsd_lb=0.0,
            rmsd_ub=2.0
        )
        
        self.assertEqual(binding.ligand, "test_ligand")
        self.assertEqual(binding.affinity, -7.5)
        self.assertTrue(binding.is_strong_binding())
    
    def test_strong_binding_detection(self):
        """Test strong binding detection."""
        strong_binding = BindingData(
            ligand="strong",
            mode=1,
            affinity=-8.0,
            rmsd_lb=0.0,
            rmsd_ub=2.0
        )
        
        weak_binding = BindingData(
            ligand="weak",
            mode=1,
            affinity=-5.0,
            rmsd_lb=0.0,
            rmsd_ub=2.0
        )
        
        self.assertTrue(strong_binding.is_strong_binding())
        self.assertFalse(weak_binding.is_strong_binding())


class TestPredictionResult(unittest.TestCase):
    """Test PredictionResult data class."""
    
    def test_prediction_result_creation(self):
        """Test creating PredictionResult object."""
        result = PredictionResult(
            compound_name="test_compound",
            predicted_affinity=-6.5,
            confidence_interval_lower=-7.0,
            confidence_interval_upper=-6.0,
            model_used="random_forest"
        )
        
        self.assertEqual(result.compound_name, "test_compound")
        self.assertEqual(result.predicted_affinity, -6.5)
        self.assertEqual(result.get_confidence_interval_width(), 1.0)


class TestPDBQTParser(unittest.TestCase):
    """Test PDBQTParser functionality."""
    
    def test_parser_initialization(self):
        """Test PDBQTParser initialization."""
        parser = PDBQTParser()
        self.assertIsNotNone(parser)
        self.assertTrue(parser.strict_validation)
    
    def test_validate_file_nonexistent(self):
        """Test validation of non-existent file."""
        parser = PDBQTParser()
        is_valid, errors = parser.validate_file("nonexistent.pdbqt")
        self.assertFalse(is_valid)
        self.assertIn("File not found", errors[0])


class TestBindingDataLoader(unittest.TestCase):
    """Test BindingDataLoader functionality."""
    
    def test_loader_initialization(self):
        """Test BindingDataLoader initialization."""
        loader = BindingDataLoader()
        self.assertEqual(loader.affinity_column, "affinity")
        self.assertEqual(loader.ligand_column, "ligand")
        self.assertEqual(loader.mode_column, "mode")
    
    def test_validate_binding_data_empty(self):
        """Test validation of empty DataFrame."""
        loader = BindingDataLoader()
        empty_df = pd.DataFrame()
        issues = loader.validate_binding_data(empty_df)
        self.assertIn('missing_values', issues)


class TestConfig(unittest.TestCase):
    """Test configuration management."""
    
    def test_config_initialization(self):
        """Test Config initialization."""
        config = Config()
        self.assertIsNotNone(config.paths)
        self.assertIsNotNone(config.data)
        self.assertIsNotNone(config.model)
    
    def test_path_config(self):
        """Test PathConfig functionality."""
        from tlr4_binding.config.settings import PathConfig
        path_config = PathConfig()
        self.assertIsInstance(path_config.project_root, Path)
        self.assertIsInstance(path_config.data_dir, Path)


class TestDataPreprocessor(unittest.TestCase):
    """Test DataPreprocessor functionality."""
    
    def test_preprocessor_initialization(self):
        """Test DataPreprocessor initialization."""
        preprocessor = DataPreprocessor()
        self.assertIsNotNone(preprocessor.binding_loader)
        self.assertIsNotNone(preprocessor.compound_matcher)
        self.assertIsNotNone(preprocessor.data_integrator)


if __name__ == '__main__':
    unittest.main()
```
