## Supplementary material for "Machine Learning-Driven Prediction of TLR4 Binding Affinity: A Comprehensive Molecular Feature Analysis for Drug Discovery": Codebase: test_binding_data_processing.py

```
"""
Unit tests for binding data processing functionality.

Tests the BindingDataLoader, DataPreprocessor, and related classes
for TLR4 binding affinity prediction.
"""

import pytest
import pandas as pd
import numpy as np
import tempfile
import os
from pathlib import Path
from unittest.mock import patch, MagicMock

from src.tlr4_binding.data_processing.preprocessor import (
    BindingDataLoader,
    DataPreprocessor,
    CompoundMatcher,
    DataIntegrator
)


class TestBindingDataLoader:
    """Test cases for BindingDataLoader class."""
    
    def setup_method(self):
        """Set up test fixtures."""
        self.loader = BindingDataLoader()
        self.sample_data = pd.DataFrame({
            'ligand': ['Compound_A', 'Compound_A', 'Compound_B', 'Compound_B'],
            'mode': [1, 2, 1, 2],
            'affinity': [-7.2, -6.8, -8.1, -7.9],
            'dist_from_rmsd_lb': [0.0, 2.5, 0.0, 1.8],
            'best_mode_rmsd_ub': [0.0, 5.2, 0.0, 4.1]
        })
    
    def test_initialization(self):
        """Test BindingDataLoader initialization."""
        loader = BindingDataLoader(
            affinity_column='binding_energy',
            ligand_column='molecule',
            mode_column='conformation'
        )
        
        assert loader.affinity_column == 'binding_energy'
        assert loader.ligand_column == 'molecule'
        assert loader.mode_column == 'conformation'
        assert loader.required_columns == ['binding_energy', 'molecule', 'conformation']
    
    def test_load_csv_success(self):
        """Test successful CSV loading."""
        with tempfile.NamedTemporaryFile(mode='w', suffix='.csv', delete=False) as f:
            self.sample_data.to_csv(f.name, index=False)
            temp_path = f.name
        
        try:
            result = self.loader.load_csv(temp_path)
            
            assert len(result) == 4
            assert 'ligand' in result.columns
            assert 'mode' in result.columns
            assert 'affinity' in result.columns
            assert result['affinity'].dtype in [np.float64, np.int64]
        finally:
            os.unlink(temp_path)
    
    def test_load_csv_file_not_found(self):
        """Test handling of missing file."""
        with pytest.raises(FileNotFoundError):
            self.loader.load_csv('nonexistent_file.csv')
    
    def test_load_csv_missing_columns(self):
        """Test handling of missing required columns."""
        incomplete_data = pd.DataFrame({
            'ligand': ['Compound_A', 'Compound_B'],
            'affinity': [-7.2, -8.1]
            # Missing 'mode' column
        })
        
        with tempfile.NamedTemporaryFile(mode='w', suffix='.csv', delete=False) as f:
            incomplete_data.to_csv(f.name, index=False)
            temp_path = f.name
        
        try:
            with pytest.raises(ValueError, match="Missing required columns"):
                self.loader.load_csv(temp_path)
        finally:
            os.unlink(temp_path)
    
    def test_load_csv_invalid_affinity_values(self):
        """Test handling of invalid affinity values."""
        invalid_data = pd.DataFrame({
            'ligand': ['Compound_A', 'Compound_B', 'Compound_C'],
            'mode': [1, 1, 1],
            'affinity': [-7.2, 'invalid', np.nan]
        })
        
        with tempfile.NamedTemporaryFile(mode='w', suffix='.csv', delete=False) as f:
            invalid_data.to_csv(f.name, index=False)
            temp_path = f.name
        
        try:
            result = self.loader.load_csv(temp_path)
            # Should remove rows with invalid affinity values
            assert len(result) == 1
            assert result['affinity'].iloc[0] == -7.2
        finally:
            os.unlink(temp_path)
    
    def test_validate_binding_data_clean(self):
        """Test validation of clean binding data."""
        issues = self.loader.validate_binding_data(self.sample_data)
        
        assert len(issues['missing_values']) == 0
        assert len(issues['invalid_affinities']) == 0
        assert len(issues['duplicate_entries']) == 0
    
    def test_validate_binding_data_missing_values(self):
        """Test validation with missing values."""
        data_with_missing = self.sample_data.copy()
        data_with_missing.loc[0, 'affinity'] = np.nan
        
        issues = self.loader.validate_binding_data(data_with_missing)
        
        assert len(issues['missing_values']) > 0
        assert 'affinity' in str(issues['missing_values'])
    
    def test_validate_binding_data_outliers(self):
        """Test outlier detection."""
        data_with_outliers = pd.DataFrame({
            'ligand': ['Compound_A'] * 10,
            'mode': range(1, 11),
            'affinity': [-7.0, -6.8, -6.9, -7.1, -6.7, -6.5, -6.6, -6.4, -6.3, -2.0]  # -2.0 is outlier
        })
        
        issues = self.loader.validate_binding_data(data_with_outliers)
        
        assert len(issues['outliers']) > 0
        assert any('outlier' in issue.lower() for issue in issues['outliers'])
    
    def test_detect_affinity_outliers_iqr(self):
        """Test IQR-based outlier detection."""
        data = pd.DataFrame({
            'ligand': ['A'] * 10,
            'mode': range(1, 11),
            'affinity': [-7.0, -6.8, -6.9, -7.1, -6.7, -6.5, -6.6, -6.4, -6.3, -2.0]
        })
        
        outlier_info = self.loader._detect_affinity_outliers(data)
        
        assert len(outlier_info['outliers']) > 0
        assert any('IQR method' in outlier for outlier in outlier_info['outliers'])
    
    def test_detect_affinity_outliers_zscore(self):
        """Test Z-score based outlier detection."""
        data = pd.DataFrame({
            'ligand': ['A'] * 10,
            'mode': range(1, 11),
            'affinity': [-7.0, -6.8, -6.9, -7.1, -6.7, -6.5, -6.6, -6.4, -6.3, -2.0]
        })
        
        outlier_info = self.loader._detect_affinity_outliers(data)
        
        assert len(outlier_info['outliers']) > 0
        # Check that at least one outlier detection method found outliers
        assert any('outlier' in outlier.lower() for outlier in outlier_info['outliers'])
    
    def test_clean_binding_data_iqr_remove(self):
        """Test IQR-based data cleaning with outlier removal."""
        data = pd.DataFrame({
            'ligand': ['A'] * 10,
            'mode': range(1, 11),
            'affinity': [-7.0, -6.8, -6.9, -7.1, -6.7, -6.5, -6.6, -6.4, -6.3, -2.0]
        })
        
        cleaned = self.loader.clean_binding_data(data, outlier_method='iqr', remove_outliers=True)
        
        assert len(cleaned) < len(data)
        assert -2.0 not in cleaned['affinity'].values
    
    def test_clean_binding_data_iqr_cap(self):
        """Test IQR-based data cleaning with outlier capping."""
        data = pd.DataFrame({
            'ligand': ['A'] * 10,
            'mode': range(1, 11),
            'affinity': [-7.0, -6.8, -6.9, -7.1, -6.7, -6.5, -6.6, -6.4, -6.3, -2.0]
        })
        
        cleaned = self.loader.clean_binding_data(data, outlier_method='iqr', remove_outliers=False)
        
        assert len(cleaned) == len(data)
        # Check that the outlier (-2.0) was capped to a different value
        assert -2.0 not in cleaned['affinity'].values  # Original outlier should be capped


class TestDataPreprocessor:
    """Test cases for DataPreprocessor class."""
    
    def setup_method(self):
        """Set up test fixtures."""
        self.preprocessor = DataPreprocessor()
        self.sample_binding_data = pd.DataFrame({
            'ligand': ['Compound_A', 'Compound_A', 'Compound_B', 'Compound_B', 'Compound_C'],
            'mode': [1, 2, 1, 2, 1],
            'affinity': [-7.2, -6.8, -8.1, -7.9, -6.5],
            'dist_from_rmsd_lb': [0.0, 2.5, 0.0, 1.8, 0.0],
            'best_mode_rmsd_ub': [0.0, 5.2, 0.0, 4.1, 0.0]
        })
        
        self.sample_features_data = pd.DataFrame({
            'compound_name': ['Compound_A', 'Compound_B', 'Compound_C'],
            'molecular_weight': [300.0, 250.0, 400.0],
            'logp': [2.5, 1.8, 3.2],
            'tpsa': [80.0, 60.0, 100.0]
        })
    
    def test_get_best_affinities(self):
        """Test extraction of best binding affinities."""
        best_affinities = self.preprocessor.get_best_affinities(self.sample_binding_data)
        
        assert len(best_affinities) == 3  # One per unique ligand
        assert best_affinities['affinity'].min() == -8.1  # Strongest binding
        assert best_affinities['affinity'].max() == -6.5  # Weakest binding
        
        # Check that we get the minimum affinity for each compound
        for ligand in best_affinities['ligand']:
            ligand_data = self.sample_binding_data[self.sample_binding_data['ligand'] == ligand]
            best_affinity = best_affinities[best_affinities['ligand'] == ligand]['affinity'].iloc[0]
            assert best_affinity == ligand_data['affinity'].min()
    
    def test_get_best_affinities_validation(self):
        """Test validation in get_best_affinities method."""
        # Test with invalid affinity data
        invalid_data = pd.DataFrame({
            'ligand': ['Compound_A'],
            'mode': [1],
            'affinity': [np.nan]  # Invalid affinity
        })
        
        with pytest.raises(ValueError):
            self.preprocessor.get_best_affinities(invalid_data)
    
    def test_validate_affinity_data(self):
        """Test affinity data validation."""
        # Test with valid data
        self.preprocessor._validate_affinity_data(self.sample_binding_data)
        
        # Test with non-numeric affinity
        invalid_data = self.sample_binding_data.copy()
        invalid_data['affinity'] = ['invalid'] * len(invalid_data)
        
        with pytest.raises(ValueError, match="must contain numeric values"):
            self.preprocessor._validate_affinity_data(invalid_data)
    
    def test_validate_best_affinities(self):
        """Test validation of best affinities extraction."""
        best_affinities = self.preprocessor.get_best_affinities(self.sample_binding_data)
        
        # This should not raise any exceptions
        self.preprocessor._validate_best_affinities(best_affinities, self.sample_binding_data)
    
    def test_integrate_datasets(self):
        """Test dataset integration."""
        integrated = self.preprocessor.integrate_datasets(
            self.sample_features_data, 
            self.sample_binding_data
        )
        
        # Should have matches for all compounds (may have multiple modes per compound)
        assert len(integrated) >= 3  # At least one match per compound
        assert 'affinity' in integrated.columns
        assert 'molecular_weight' in integrated.columns
        assert 'compound_name' in integrated.columns
    
    def test_preprocess_pipeline(self):
        """Test complete preprocessing pipeline."""
        with tempfile.NamedTemporaryFile(mode='w', suffix='.csv', delete=False) as f:
            self.sample_binding_data.to_csv(f.name, index=False)
            temp_path = f.name
        
        try:
            result = self.preprocessor.preprocess_pipeline(self.sample_features_data, temp_path)
            
            assert len(result) == 3
            assert 'affinity' in result.columns
            assert 'molecular_weight' in result.columns
            assert result['affinity'].min() == -8.1  # Strongest binding
        finally:
            os.unlink(temp_path)


class TestCompoundMatcher:
    """Test cases for CompoundMatcher class."""
    
    def setup_method(self):
        """Set up test fixtures."""
        self.matcher = CompoundMatcher(threshold=80.0)
    
    def test_match_compounds_exact(self):
        """Test exact compound name matching."""
        names1 = ['Compound_A', 'Compound_B']
        names2 = ['Compound_A', 'Compound_B', 'Compound_C']
        
        matches = self.matcher.match_compounds(names1, names2)
        
        assert matches['Compound_A'] == 'Compound_A'
        assert matches['Compound_B'] == 'Compound_B'
    
    def test_match_compounds_fuzzy(self):
        """Test fuzzy compound name matching."""
        names1 = ['Compound A', 'Compound B']
        names2 = ['Compound_A', 'Compound_B', 'Compound_C']
        
        matches = self.matcher.match_compounds(names1, names2)
        
        assert 'Compound A' in matches
        assert 'Compound B' in matches
    
    def test_match_compounds_no_match(self):
        """Test handling when no matches are found."""
        names1 = ['Unknown_Compound']
        names2 = ['Compound_A', 'Compound_B']
        
        matches = self.matcher.match_compounds(names1, names2)
        
        assert 'Unknown_Compound' not in matches
    
    def test_clean_name(self):
        """Test compound name cleaning."""
        # Test basic cleaning
        assert self.matcher._clean_name('Compound_A') == 'a'  # 'compound' prefix removed
        assert self.matcher._clean_name('  Compound B  ') == 'b'  # 'compound' prefix removed
        
        # Test prefix/suffix removal
        assert self.matcher._clean_name('compound_ligand_A') == 'liganda'  # 'compound' prefix removed, 'ligand' becomes 'liganda'
        assert self.matcher._clean_name('molecule_B_docked') == 'b'  # 'molecule' prefix removed, 'docked' suffix removed
        
        # Test special character removal
        assert self.matcher._clean_name('Compound-A_123') == 'a123'  # 'compound' prefix removed, special chars removed
    
    def test_get_match_confidence(self):
        """Test match confidence calculation."""
        confidence = self.matcher.get_match_confidence('Compound A', 'Compound_A')
        assert confidence > 80  # Should be high confidence
        
        confidence = self.matcher.get_match_confidence('Compound A', 'Different_Compound')
        assert confidence < 80  # Should be low confidence


class TestDataIntegrator:
    """Test cases for DataIntegrator class."""
    
    def setup_method(self):
        """Set up test fixtures."""
        self.integrator = DataIntegrator()
        self.features_data = pd.DataFrame({
            'compound_name': ['Compound_A', 'Compound_B', 'Compound_C'],
            'molecular_weight': [300.0, 250.0, 400.0],
            'logp': [2.5, 1.8, 3.2]
        })
        self.binding_data = pd.DataFrame({
            'ligand': ['Compound_A', 'Compound_B', 'Compound_D'],
            'affinity': [-7.2, -8.1, -6.5],
            'mode': [1, 1, 1]
        })
    
    def test_integrate_datasets(self):
        """Test dataset integration."""
        integrated = self.integrator.integrate_datasets(
            self.features_data, 
            self.binding_data
        )
        
        assert len(integrated) == 2  # Only Compound_A and Compound_B should match
        assert 'molecular_weight' in integrated.columns
        assert 'affinity' in integrated.columns
    
    def test_integration_stats(self):
        """Test integration statistics."""
        self.integrator.integrate_datasets(self.features_data, self.binding_data)
        stats = self.integrator.get_integration_stats()
        
        assert 'total_features' in stats
        assert 'total_binding' in stats
        assert 'successful_matches' in stats
        assert 'integrated_records' in stats
        assert 'match_rate' in stats
        
        assert stats['total_features'] == 3
        assert stats['total_binding'] == 3
        assert stats['integrated_records'] == 2


class TestBindingDataProcessingIntegration:
    """Integration tests for binding data processing."""
    
    def test_end_to_end_processing(self):
        """Test complete end-to-end binding data processing."""
        # Create sample data
        binding_data = pd.DataFrame({
            'ligand': ['Andrographolide', 'Andrographolide', 'Curcumin', 'Curcumin'],
            'mode': [1, 2, 1, 2],
            'affinity': [-7.203, -7.197, -8.5, -8.2],
            'dist_from_rmsd_lb': [0.0, 2.783, 0.0, 1.5],
            'best_mode_rmsd_ub': [0.0, 6.674, 0.0, 4.2]
        })
        
        features_data = pd.DataFrame({
            'compound_name': ['Andrographolide', 'Curcumin'],
            'molecular_weight': [350.0, 368.0],
            'logp': [2.5, 3.0],
            'tpsa': [80.0, 90.0]
        })
        
        # Test preprocessing
        preprocessor = DataPreprocessor()
        
        # Test best affinities extraction
        best_affinities = preprocessor.get_best_affinities(binding_data)
        assert len(best_affinities) == 2
        assert best_affinities['affinity'].min() == -8.5  # Strongest binding
        
        # Test dataset integration
        integrated = preprocessor.integrate_datasets(features_data, best_affinities)
        assert len(integrated) == 2
        assert 'affinity' in integrated.columns
        assert 'molecular_weight' in integrated.columns
        
        # Verify strongest binding is preserved
        strongest_binding = integrated.loc[integrated['affinity'].idxmin()]
        assert strongest_binding['compound_name'] == 'Curcumin'
        assert strongest_binding['affinity'] == -8.5


if __name__ == '__main__':
    pytest.main([__file__, '-v'])
```
