## Supplementary material for "Machine Learning-Driven Prediction of TLR4 Binding Affinity: A Comprehensive Molecular Feature Analysis for Drug Discovery": Codebase: test_compound_analysis.py

```
import numpy as np
import pandas as pd

from src.tlr4_binding.ml_components import CompoundAnalysis


def test_tanimoto_similarity_fallback():
    ca = CompoundAnalysis()
    sim = ca.compute_tanimoto_similarity(["CCO", "CCN", None])
    assert sim.shape == (3, 3)
    assert np.allclose(np.diag(sim), 1.0)


def test_ranking_with_ci():
    ca = CompoundAnalysis()
    df = pd.DataFrame({
        "compound_name": ["a", "b", "c"],
        "predicted_affinity": [-7.1, -8.2, -8.2],
        "confidence_interval_lower": [-7.3, -8.5, -8.1],
        "confidence_interval_upper": [-6.9, -7.9, -7.7],
    })
    ranked = ca.rank_compounds(df)
    assert list(ranked["compound_name"])[:2] == ["b", "c"]
    assert ranked.iloc[0]["rank"] == 1


def test_chemical_space_pca_fallback():
    ca = CompoundAnalysis()
    X = np.random.RandomState(0).randn(10, 5)
    emb = ca.chemical_space(X)
    assert emb.shape == (10, 2)
```
