## Supplementary material for "Machine Learning-Driven Prediction of TLR4 Binding Affinity: A Comprehensive Molecular Feature Analysis for Drug Discovery": Codebase: test_compound_matching_real_data.py

```
#!/usr/bin/env python3
"""
Test script for compound name matching and data integration with real data.

This script validates the compound name matching and data integration functionality
using actual data from the binding-data directory.
"""

import sys
import os
import pandas as pd
import numpy as np
from pathlib import Path
import logging

# Add src to path for imports
sys.path.insert(0, os.path.join(os.path.dirname(__file__), 'src'))

from tlr4_binding.data_processing.preprocessor import (
    DataPreprocessor, 
    BindingDataLoader, 
    CompoundMatcher, 
    DataIntegrator
)

# Set up logging
logging.basicConfig(level=logging.INFO, format='%(asctime)s - %(levelname)s - %(message)s')
logger = logging.getLogger(__name__)


def create_sample_molecular_features():
    """Create sample molecular features for testing."""
    return pd.DataFrame({
        'compound_name': [
            'Andrographolide',
            'Curcumin',
            'Resveratrol',
            'Quercetin',
            'Epigallocatechin',
            'Capsaicin',
            'Gingerol',
            'Caffeic acid',
            'Chlorogenic acid',
            'Ferulic acid',
            'Rutin',
            'Kaempferol',
            'Apigenin',
            'Luteolin',
            'Naringenin',
            'Hesperidin',
            'Catechin',
            'Epicatechin',
            'Gallic acid',
            'Cinnamic acid'
        ],
        'molecular_weight': [
            350.4, 368.4, 228.2, 302.2, 458.4, 305.4, 276.4, 180.2, 354.3, 194.2,
            610.5, 286.2, 270.2, 286.2, 272.3, 610.6, 290.3, 290.3, 170.1, 148.2
        ],
        'logp': [
            2.3, 2.5, 3.1, 1.8, 2.0, 3.8, 2.9, 1.2, 0.8, 1.4,
            1.1, 2.0, 2.0, 2.0, 2.0, 1.0, 1.0, 1.0, 0.7, 1.9
        ],
        'tpsa': [
            74.6, 93.1, 60.7, 131.4, 197.4, 58.6, 50.5, 77.8, 164.7, 66.8,
            210.5, 111.1, 90.9, 111.1, 90.9, 210.5, 110.4, 110.4, 97.1, 37.3
        ],
        'hbd': [3, 2, 3, 5, 8, 1, 1, 2, 4, 2, 10, 4, 3, 4, 3, 10, 5, 5, 3, 1],
        'hba': [4, 6, 3, 7, 11, 2, 2, 4, 7, 3, 16, 6, 5, 6, 5, 16, 6, 6, 4, 2]
    })


def test_compound_matching():
    """Test compound name matching functionality."""
    logger.info("Testing compound name matching...")
    
    # Create sample features
    features_df = create_sample_molecular_features()
    
    # Load real binding data
    binding_csv_path = Path("binding-data/processed/processed_logs.csv")
    if not binding_csv_path.exists():
        logger.error(f"Binding data file not found: {binding_csv_path}")
        return False
    
    binding_loader = BindingDataLoader()
    binding_df = binding_loader.load_csv(str(binding_csv_path))
    
    # Get unique compound names from binding data
    binding_compounds = binding_df['ligand'].unique().tolist()
    feature_compounds = features_df['compound_name'].tolist()
    
    logger.info(f"Feature compounds: {len(feature_compounds)}")
    logger.info(f"Binding compounds: {len(binding_compounds)}")
    
    # Test compound matching with different thresholds
    thresholds = [60.0, 75.0, 85.0, 95.0]
    
    for threshold in thresholds:
        logger.info(f"\nTesting with threshold: {threshold}")
        
        matcher = CompoundMatcher(threshold=threshold)
        matches = matcher.match_compounds(feature_compounds, binding_compounds)
        
        logger.info(f"Matches found: {len(matches)}")
        logger.info(f"Match rate: {len(matches)/len(feature_compounds):.2%}")
        
        # Show some example matches
        if matches:
            logger.info("Example matches:")
            for i, (feature_name, binding_name) in enumerate(list(matches.items())[:5]):
                confidence = matcher.get_match_confidence(feature_name, binding_name)
                logger.info(f"  {feature_name} -> {binding_name} (confidence: {confidence:.1f})")
    
    return True


def test_data_integration():
    """Test data integration functionality."""
    logger.info("\nTesting data integration...")
    
    # Create sample features
    features_df = create_sample_molecular_features()
    
    # Load and process binding data
    binding_csv_path = Path("binding-data/processed/processed_logs.csv")
    if not binding_csv_path.exists():
        logger.error(f"Binding data file not found: {binding_csv_path}")
        return False
    
    preprocessor = DataPreprocessor()
    
    # Load binding data
    binding_df = preprocessor.load_binding_data(str(binding_csv_path))
    logger.info(f"Loaded {len(binding_df)} binding records")
    
    # Get best affinities
    best_affinities = preprocessor.get_best_affinities(binding_df)
    logger.info(f"Extracted {len(best_affinities)} best binding modes")
    
    # Test integration with different matching strategies
    strategies = [
        ("Partial Ratio", CompoundMatcher(threshold=75.0, use_partial_ratio=True)),
        ("Regular Ratio", CompoundMatcher(threshold=75.0, use_partial_ratio=False)),
        ("High Threshold", CompoundMatcher(threshold=90.0, use_partial_ratio=True)),
        ("Low Threshold", CompoundMatcher(threshold=60.0, use_partial_ratio=True))
    ]
    
    for strategy_name, matcher in strategies:
        logger.info(f"\nTesting integration with {strategy_name}:")
        
        integrator = DataIntegrator(matcher)
        integrated_df = integrator.integrate_datasets(features_df, best_affinities)
        
        stats = integrator.get_integration_stats()
        logger.info(f"  Integrated records: {stats['integrated_records']}")
        logger.info(f"  Match rate: {stats['match_rate']:.2%}")
        
        if len(integrated_df) > 0:
            logger.info(f"  Affinity range: {integrated_df['affinity'].min():.3f} to {integrated_df['affinity'].max():.3f} kcal/mol")
            logger.info(f"  Average affinity: {integrated_df['affinity'].mean():.3f} kcal/mol")
            
            # Show some integrated records
            logger.info("  Sample integrated records:")
            sample_cols = ['compound_name', 'molecular_weight', 'logp', 'affinity']
            available_cols = [col for col in sample_cols if col in integrated_df.columns]
            logger.info(f"  {integrated_df[available_cols].head(3).to_string(index=False)}")
    
    return True


def test_complete_pipeline():
    """Test complete preprocessing pipeline."""
    logger.info("\nTesting complete preprocessing pipeline...")
    
    # Create sample features
    features_df = create_sample_molecular_features()
    
    # Run complete pipeline
    binding_csv_path = Path("binding-data/processed/processed_logs.csv")
    if not binding_csv_path.exists():
        logger.error(f"Binding data file not found: {binding_csv_path}")
        return False
    
    preprocessor = DataPreprocessor()
    
    try:
        integrated_df = preprocessor.preprocess_pipeline(features_df, str(binding_csv_path))
        
        logger.info(f"Pipeline completed successfully!")
        logger.info(f"Integrated records: {len(integrated_df)}")
        
        if len(integrated_df) > 0:
            logger.info(f"Columns: {list(integrated_df.columns)}")
            logger.info(f"Data types: {integrated_df.dtypes.to_dict()}")
            
            # Check data quality
            missing_values = integrated_df.isnull().sum()
            if missing_values.any():
                logger.warning(f"Missing values found: {missing_values[missing_values > 0].to_dict()}")
            else:
                logger.info("No missing values found")
            
            # Check for duplicates
            duplicates = integrated_df['compound_name'].duplicated().sum()
            if duplicates > 0:
                logger.warning(f"Found {duplicates} duplicate compound names")
            else:
                logger.info("No duplicate compound names found")
            
            # Show summary statistics
            logger.info("\nSummary statistics:")
            numeric_cols = integrated_df.select_dtypes(include=[np.number]).columns
            if len(numeric_cols) > 0:
                logger.info(f"{integrated_df[numeric_cols].describe().to_string()}")
        
        return True
        
    except Exception as e:
        logger.error(f"Pipeline failed: {str(e)}")
        return False


def test_data_validation():
    """Test data validation functionality."""
    logger.info("\nTesting data validation...")
    
    binding_csv_path = Path("binding-data/processed/processed_logs.csv")
    if not binding_csv_path.exists():
        logger.error(f"Binding data file not found: {binding_csv_path}")
        return False
    
    binding_loader = BindingDataLoader()
    binding_df = binding_loader.load_csv(str(binding_csv_path))
    
    # Validate binding data
    validation_results = binding_loader.validate_binding_data(binding_df)
    
    logger.info("Data validation results:")
    for category, issues in validation_results.items():
        if issues:
            logger.info(f"  {category}: {len(issues)} issues")
            for issue in issues[:3]:  # Show first 3 issues
                logger.info(f"    - {issue}")
        else:
            logger.info(f"  {category}: No issues found")
    
    return True


def main():
    """Run all tests."""
    logger.info("Starting compound name matching and data integration tests...")
    
    tests = [
        ("Compound Matching", test_compound_matching),
        ("Data Integration", test_data_integration),
        ("Complete Pipeline", test_complete_pipeline),
        ("Data Validation", test_data_validation)
    ]
    
    results = {}
    
    for test_name, test_func in tests:
        logger.info(f"\n{'='*50}")
        logger.info(f"Running {test_name} test...")
        logger.info(f"{'='*50}")
        
        try:
            success = test_func()
            results[test_name] = success
            if success:
                logger.info(f"✅ {test_name} test passed")
            else:
                logger.error(f"❌ {test_name} test failed")
        except Exception as e:
            logger.error(f"❌ {test_name} test failed with exception: {str(e)}")
            results[test_name] = False
    
    # Summary
    logger.info(f"\n{'='*50}")
    logger.info("Test Summary:")
    logger.info(f"{'='*50}")
    
    passed = sum(results.values())
    total = len(results)
    
    for test_name, success in results.items():
        status = "✅ PASSED" if success else "❌ FAILED"
        logger.info(f"{test_name}: {status}")
    
    logger.info(f"\nOverall: {passed}/{total} tests passed")
    
    if passed == total:
        logger.info("🎉 All tests passed!")
        return 0
    else:
        logger.error("💥 Some tests failed!")
        return 1


if __name__ == "__main__":
    exit(main())
```
