## Supplementary material for "Machine Learning-Driven Prediction of TLR4 Binding Affinity: A Comprehensive Molecular Feature Analysis for Drug Discovery": Codebase: test_comprehensive_evaluation.py

```
"""
Comprehensive tests for the model evaluation framework.

This module tests all aspects of the evaluation framework including:
- Performance metrics calculation
- Cross-validation evaluation
- Learning curves and validation curves
- Statistical significance testing
- Comprehensive evaluation integration
"""

import pytest
import numpy as np
import pandas as pd
import matplotlib.pyplot as plt
from unittest.mock import Mock, patch
import tempfile
import shutil
from pathlib import Path

# Add src to path for imports
import sys
sys.path.append(str(Path(__file__).parent.parent / "src"))

from tlr4_binding.ml_components.evaluator import (
    ModelEvaluator, PerformanceMetrics, ModelEvaluatorInterface
)
from tlr4_binding.ml_components.comprehensive_evaluator import ComprehensiveEvaluator
from sklearn.ensemble import RandomForestRegressor
from sklearn.linear_model import LinearRegression
from sklearn.svm import SVR
from sklearn.model_selection import train_test_split
from sklearn.datasets import make_regression


class TestPerformanceMetrics:
    """Test PerformanceMetrics class functionality."""
    
    def setup_method(self):
        """Set up test data."""
        np.random.seed(42)
        self.y_true = np.array([1, 2, 3, 4, 5, 6, 7, 8, 9, 10])
        self.y_pred = np.array([1.1, 1.9, 3.2, 3.8, 5.1, 5.9, 7.2, 7.8, 9.1, 9.9])
        self.model_name = "test_model"
    
    def test_performance_metrics_initialization(self):
        """Test PerformanceMetrics initialization."""
        metrics = PerformanceMetrics(self.y_true, self.y_pred, self.model_name)
        
        assert metrics.y_true is self.y_true
        assert metrics.y_pred is self.y_pred
        assert metrics.model_name == self.model_name
        assert isinstance(metrics.metrics, dict)
        assert 'r2' in metrics.metrics
        assert 'rmse' in metrics.metrics
        assert 'mae' in metrics.metrics
    
    def test_performance_metrics_calculation(self):
        """Test metrics calculation accuracy."""
        metrics = PerformanceMetrics(self.y_true, self.y_pred, self.model_name)
        
        # Test that metrics are reasonable
        assert 0 <= metrics.metrics['r2'] <= 1  # R² should be between 0 and 1
        assert metrics.metrics['rmse'] > 0  # RMSE should be positive
        assert metrics.metrics['mae'] > 0   # MAE should be positive
        assert metrics.metrics['pearson_r'] > 0  # Should be positive correlation
        assert metrics.metrics['spearman_r'] > 0  # Should be positive correlation
    
    def test_performance_metrics_with_perfect_prediction(self):
        """Test metrics with perfect predictions."""
        y_pred_perfect = self.y_true.copy()
        metrics = PerformanceMetrics(self.y_true, y_pred_perfect, "perfect_model")
        
        assert abs(metrics.metrics['r2'] - 1.0) < 1e-10
        assert abs(metrics.metrics['rmse']) < 1e-10
        assert abs(metrics.metrics['mae']) < 1e-10
        assert abs(metrics.metrics['pearson_r'] - 1.0) < 1e-10
        assert abs(metrics.metrics['spearman_r'] - 1.0) < 1e-10
    
    def test_performance_metrics_with_zero_variance(self):
        """Test metrics with zero variance target."""
        y_constant = np.array([5.0, 5.0, 5.0, 5.0, 5.0])
        y_pred_constant = np.array([4.0, 5.0, 6.0, 5.0, 4.0])
        
        metrics = PerformanceMetrics(y_constant, y_pred_constant, "constant_target")
        
        # R² should be undefined (NaN) for constant targets
        assert np.isnan(metrics.metrics['r2'])
        assert metrics.metrics['rmse'] > 0
        assert metrics.metrics['mae'] > 0
    
    def test_get_summary(self):
        """Test get_summary method."""
        metrics = PerformanceMetrics(self.y_true, self.y_pred, self.model_name)
        summary = metrics.get_summary()
        
        assert isinstance(summary, dict)
        assert 'model_name' in summary
        assert 'r2_score' in summary
        assert 'rmse' in summary
        assert 'mae' in summary
        assert summary['model_name'] == self.model_name
    
    def test_to_dataframe(self):
        """Test to_dataframe method."""
        metrics = PerformanceMetrics(self.y_true, self.y_pred, self.model_name)
        df = metrics.to_dataframe()
        
        assert isinstance(df, pd.DataFrame)
        assert len(df) == 1
        assert 'r2' in df.columns
        assert 'rmse' in df.columns
        assert 'mae' in df.columns


class TestModelEvaluator:
    """Test ModelEvaluator class functionality."""
    
    def setup_method(self):
        """Set up test data and evaluator."""
        np.random.seed(42)
        self.y_true = np.array([1, 2, 3, 4, 5, 6, 7, 8, 9, 10])
        self.y_pred = np.array([1.1, 1.9, 3.2, 3.8, 5.1, 5.9, 7.2, 7.8, 9.1, 9.9])
        self.evaluator = ModelEvaluator()
    
    def test_evaluator_initialization(self):
        """Test ModelEvaluator initialization."""
        assert self.evaluator.confidence_level == 0.95
        assert isinstance(self.evaluator.evaluation_history, list)
    
    def test_evaluate_method(self):
        """Test evaluate method."""
        metrics = self.evaluator.evaluate(self.y_true, self.y_pred, "test_model")
        
        assert isinstance(metrics, PerformanceMetrics)
        assert metrics.model_name == "test_model"
        assert len(self.evaluator.evaluation_history) == 1
    
    def test_compare_models(self):
        """Test compare_models method."""
        # Create multiple performance metrics
        metrics1 = PerformanceMetrics(self.y_true, self.y_pred, "model1")
        metrics2 = PerformanceMetrics(self.y_pred, self.y_true, "model2")  # Swapped for different performance
        
        results = {"model1": metrics1, "model2": metrics2}
        comparison_df = self.evaluator.compare_models(results)
        
        assert isinstance(comparison_df, pd.DataFrame)
        assert len(comparison_df) == 2
        assert 'model_name' in comparison_df.columns
        assert 'r2_score' in comparison_df.columns
        assert 'rmse' in comparison_df.columns
    
    def test_statistical_significance_test(self):
        """Test statistical significance testing."""
        metrics1 = PerformanceMetrics(self.y_true, self.y_pred, "model1")
        metrics2 = PerformanceMetrics(self.y_pred, self.y_true, "model2")  # Different predictions
        
        test_result = self.evaluator.statistical_significance_test(metrics1, metrics2)
        
        assert isinstance(test_result, dict)
        assert 'paired_t_test' in test_result
        assert 'wilcoxon_test' in test_result
        assert 'model1_r2' in test_result
        assert 'model2_r2' in test_result
        assert 'r2_difference' in test_result
    
    def test_statistical_significance_test_same_length_requirement(self):
        """Test that statistical test requires same length predictions."""
        y_true_short = self.y_true[:5]
        y_pred_short = self.y_pred[:5]
        y_pred_long = self.y_pred
        
        metrics1 = PerformanceMetrics(y_true_short, y_pred_short, "model1")
        metrics2 = PerformanceMetrics(y_true_short, y_pred_long, "model2")
        
        with pytest.raises(ValueError, match="same number of predictions"):
            self.evaluator.statistical_significance_test(metrics1, metrics2)
    
    def test_cross_validate_model(self):
        """Test cross-validation evaluation."""
        # Create synthetic data
        X, y = make_regression(n_samples=100, n_features=10, noise=0.1, random_state=42)
        model = RandomForestRegressor(n_estimators=10, random_state=42)
        
        cv_results = self.evaluator.cross_validate_model(model, X, y, "test_model", cv=3)
        
        assert isinstance(cv_results, dict)
        assert 'r2' in cv_results
        assert 'rmse' in cv_results
        assert 'mae' in cv_results
        
        # Check structure of results
        for metric_name, metric_data in cv_results.items():
            if metric_name not in ['r2', 'rmse', 'mae']:
                continue
            assert 'scores' in metric_data
            assert 'mean' in metric_data
            assert 'std' in metric_data
            assert 'min' in metric_data
            assert 'max' in metric_data
    
    def test_generate_learning_curves(self):
        """Test learning curves generation."""
        # Create synthetic data
        X, y = make_regression(n_samples=100, n_features=5, noise=0.1, random_state=42)
        model = RandomForestRegressor(n_estimators=10, random_state=42)
        
        with patch('matplotlib.pyplot.show'):
            curve_data = self.evaluator.generate_learning_curves(
                model, X, y, "test_model", cv=3
            )
        
        assert isinstance(curve_data, dict)
        assert 'train_sizes' in curve_data
        assert 'train_mean' in curve_data
        assert 'train_std' in curve_data
        assert 'val_mean' in curve_data
        assert 'val_std' in curve_data
        assert 'model_name' in curve_data
    
    def test_generate_validation_curves(self):
        """Test validation curves generation."""
        # Create synthetic data
        X, y = make_regression(n_samples=100, n_features=5, noise=0.1, random_state=42)
        model = RandomForestRegressor(n_estimators=10, random_state=42)
        param_range = [10, 20, 50]
        
        with patch('matplotlib.pyplot.show'):
            curve_data = self.evaluator.generate_validation_curves(
                model, X, y, 'n_estimators', param_range, "test_model", cv=3
            )
        
        assert isinstance(curve_data, dict)
        assert 'param_range' in curve_data
        assert 'train_mean' in curve_data
        assert 'train_std' in curve_data
        assert 'val_mean' in curve_data
        assert 'val_std' in curve_data
        assert 'param_name' in curve_data
        assert 'model_name' in curve_data


class TestComprehensiveEvaluator:
    """Test ComprehensiveEvaluator class functionality."""
    
    def setup_method(self):
        """Set up test data and evaluator."""
        self.temp_dir = tempfile.mkdtemp()
        self.evaluator = ComprehensiveEvaluator(output_dir=self.temp_dir)
        
        # Create synthetic data
        X, y = make_regression(n_samples=200, n_features=10, noise=0.1, random_state=42)
        self.X_train, self.X_test, self.y_train, self.y_test = train_test_split(
            X, y, test_size=0.3, random_state=42
        )
        self.X_val = self.X_test  # Use test as validation for simplicity
        self.y_val = self.y_test
    
    def teardown_method(self):
        """Clean up temporary directory."""
        shutil.rmtree(self.temp_dir, ignore_errors=True)
    
    def test_comprehensive_evaluator_initialization(self):
        """Test ComprehensiveEvaluator initialization."""
        assert self.evaluator.output_dir == Path(self.temp_dir)
        assert self.evaluator.confidence_level == 0.95
        assert isinstance(self.evaluator.base_evaluator, ModelEvaluator)
        assert isinstance(self.evaluator.evaluation_results, dict)
    
    def test_get_default_model_configs(self):
        """Test default model configurations."""
        configs = self.evaluator._get_default_model_configs()
        
        assert isinstance(configs, dict)
        assert 'traditional_ml' in configs
        assert 'deep_learning' in configs
        assert 'ensemble' in configs
        
        # Check traditional ML configs
        assert 'random_forest' in configs['traditional_ml']
        assert 'xgboost' in configs['traditional_ml']
        assert 'svr' in configs['traditional_ml']
    
    def test_make_serializable(self):
        """Test JSON serialization helper."""
        # Test various data types
        test_data = {
            'int_val': np.int64(42),
            'float_val': np.float64(3.14),
            'array_val': np.array([1, 2, 3]),
            'normal_val': 'string',
            'nested_dict': {
                'nested_int': np.int32(10),
                'nested_list': [np.float32(1.0), np.float32(2.0)]
            }
        }
        
        serializable_data = self.evaluator._make_serializable(test_data)
        
        assert isinstance(serializable_data['int_val'], int)
        assert isinstance(serializable_data['float_val'], float)
        assert isinstance(serializable_data['array_val'], list)
        assert isinstance(serializable_data['normal_val'], str)
        assert isinstance(serializable_data['nested_dict']['nested_int'], int)
        assert isinstance(serializable_data['nested_dict']['nested_list'], list)
    
    def test_evaluate_traditional_ml_models(self):
        """Test traditional ML model evaluation."""
        with patch('tlr4_binding.ml_components.comprehensive_evaluator.MLModelTrainer') as mock_trainer:
            # Mock the trainer and models
            mock_trainer_instance = Mock()
            mock_trainer.return_value = mock_trainer_instance
            
            # Create mock model
            mock_model = Mock()
            mock_model.predict.return_value = self.y_test + np.random.normal(0, 0.1, len(self.y_test))
            
            mock_trainer_instance.train_all_models.return_value = {
                'random_forest': mock_model,
                'linear_regression': mock_model
            }
            
            results = self.evaluator._evaluate_traditional_ml_models(
                self.X_train, self.y_train, self.X_val, self.y_val,
                self.X_test, self.y_test, {}
            )
            
            assert isinstance(results, dict)
            assert 'random_forest' in results
            assert 'linear_regression' in results
            
            for model_name, result in results.items():
                assert 'model' in result
                assert 'metrics' in result
                assert 'predictions' in result
                assert isinstance(result['metrics'], PerformanceMetrics)
    
    def test_get_best_models(self):
        """Test best model selection."""
        # Create mock results
        mock_results = {
            'traditional_ml': {
                'model1': {
                    'model': Mock(),
                    'metrics': Mock()
                },
                'model2': {
                    'model': Mock(),
                    'metrics': Mock()
                }
            },
            'deep_learning': {
                'model3': {
                    'model': Mock(),
                    'metrics': Mock()
                }
            }
        }
        
        # Mock metrics with different R² scores
        mock_results['traditional_ml']['model1']['metrics'].metrics = {'r2': 0.8}
        mock_results['traditional_ml']['model2']['metrics'].metrics = {'r2': 0.9}
        mock_results['deep_learning']['model3']['metrics'].metrics = {'r2': 0.85}
        
        best_models = self.evaluator._get_best_models(mock_results)
        
        assert isinstance(best_models, dict)
        assert 'traditional_ml' in best_models
        assert 'deep_learning' in best_models
        assert best_models['traditional_ml']['name'] == 'model2'  # Highest R²
        assert best_models['deep_learning']['name'] == 'model3'
    
    def test_perform_statistical_tests(self):
        """Test statistical significance testing."""
        # Create mock results with metrics
        mock_results = {
            'traditional_ml': {
                'model1': {
                    'metrics': PerformanceMetrics(self.y_test, self.y_test + 0.1, "model1")
                },
                'model2': {
                    'metrics': PerformanceMetrics(self.y_test, self.y_test + 0.2, "model2")
                }
            }
        }
        
        statistical_tests = self.evaluator._perform_statistical_tests(mock_results)
        
        assert isinstance(statistical_tests, dict)
        assert 'traditional_ml_model1_vs_traditional_ml_model2' in statistical_tests
        
        test_result = statistical_tests['traditional_ml_model1_vs_traditional_ml_model2']
        assert 'paired_t_test' in test_result
        assert 'wilcoxon_test' in test_result
        assert 'model1_r2' in test_result
        assert 'model2_r2' in test_result
    
    def test_generate_evaluation_summary(self):
        """Test evaluation summary generation."""
        # Create mock results
        mock_results = {
            'traditional_ml': {
                'model1': {
                    'model': Mock(),
                    'metrics': PerformanceMetrics(self.y_test, self.y_test + 0.1, "model1")
                }
            },
            'cross_validation': {
                'traditional_ml_model1': {
                    'r2': {'mean': 0.8, 'std': 0.05},
                    'rmse': {'mean': 1.2, 'std': 0.1}
                }
            }
        }
        
        summary = self.evaluator._generate_evaluation_summary(mock_results)
        
        assert isinstance(summary, dict)
        assert 'total_models_evaluated' in summary
        assert 'best_overall_model' in summary
        assert 'best_r2_score' in summary
        assert 'category_performance' in summary
        assert 'cross_validation_summary' in summary
        
        assert summary['total_models_evaluated'] == 1
        assert 'traditional_ml' in summary['category_performance']
    
    def test_save_evaluation_results(self):
        """Test saving evaluation results."""
        mock_results = {
            'summary': {
                'total_models_evaluated': 1,
                'best_overall_model': 'test_model',
                'best_r2_score': 0.8
            }
        }
        
        self.evaluator._save_evaluation_results(mock_results)
        
        # Check that files were created
        summary_file = Path(self.temp_dir) / "evaluation_summary.json"
        detailed_file = Path(self.temp_dir) / "detailed_evaluation_results.pkl"
        
        assert summary_file.exists()
        assert detailed_file.exists()
        
        # Check JSON content
        import json
        with open(summary_file, 'r') as f:
            summary_data = json.load(f)
        
        assert summary_data['total_models_evaluated'] == 1
        assert summary_data['best_overall_model'] == 'test_model'


class TestEvaluationIntegration:
    """Test integration between different evaluation components."""
    
    def setup_method(self):
        """Set up test data."""
        np.random.seed(42)
        X, y = make_regression(n_samples=100, n_features=5, noise=0.1, random_state=42)
        self.X_train, self.X_test, self.y_train, self.y_test = train_test_split(
            X, y, test_size=0.3, random_state=42
        )
    
    def test_model_evaluator_interface_compliance(self):
        """Test that ModelEvaluator implements the interface correctly."""
        evaluator = ModelEvaluator()
        
        # Test that it implements the abstract methods
        assert hasattr(evaluator, 'evaluate')
        assert hasattr(evaluator, 'compare_models')
        
        # Test method signatures
        import inspect
        
        evaluate_sig = inspect.signature(evaluator.evaluate)
        assert 'y_true' in evaluate_sig.parameters
        assert 'y_pred' in evaluate_sig.parameters
        assert 'model_name' in evaluate_sig.parameters
        
        compare_sig = inspect.signature(evaluator.compare_models)
        assert 'results' in compare_sig.parameters
    
    def test_performance_metrics_consistency(self):
        """Test consistency of performance metrics across different scenarios."""
        evaluator = ModelEvaluator()
        
        # Test with different prediction qualities
        scenarios = [
            (self.y_test, self.y_test + 0.1, "good_prediction"),
            (self.y_test, self.y_test + 1.0, "poor_prediction"),
            (self.y_test, self.y_test, "perfect_prediction")
        ]
        
        for y_true, y_pred, name in scenarios:
            metrics = evaluator.evaluate(y_true, y_pred, name)
            
            # Check that all expected metrics are present
            required_metrics = ['r2', 'rmse', 'mae', 'pearson_r', 'spearman_r']
            for metric in required_metrics:
                assert metric in metrics.metrics
                assert not np.isnan(metrics.metrics[metric]) or metric == 'r2'  # R² can be NaN for constant targets
    
    def test_cross_validation_consistency(self):
        """Test that cross-validation results are consistent."""
        evaluator = ModelEvaluator()
        model = RandomForestRegressor(n_estimators=10, random_state=42)
        
        # Run cross-validation multiple times with same parameters
        cv_results1 = evaluator.cross_validate_model(model, self.X_train, self.y_train, "test", cv=3)
        cv_results2 = evaluator.cross_validate_model(model, self.X_train, self.y_train, "test", cv=3)
        
        # Results should be identical due to random_state
        assert cv_results1['r2']['mean'] == cv_results2['r2']['mean']
        assert cv_results1['rmse']['mean'] == cv_results2['rmse']['mean']
        assert cv_results1['mae']['mean'] == cv_results2['mae']['mean']


if __name__ == "__main__":
    pytest.main([__file__, "-v"])
```
