## Supplementary material for "Machine Learning-Driven Prediction of TLR4 Binding Affinity: A Comprehensive Molecular Feature Analysis for Drug Discovery": Codebase: test_data_splitting.py

```
"""
Tests for Data Splitting and Validation Framework

Comprehensive test suite for data splitting, cross-validation, and quality reporting
functionality in the TLR4 binding prediction system.
"""

import pytest
import numpy as np
import pandas as pd
from unittest.mock import patch, MagicMock
from sklearn.datasets import make_regression
from sklearn.preprocessing import StandardScaler

from src.tlr4_binding.ml_components.data_splitting import (
    DataSplitter,
    CrossValidationSetup,
    DataValidationFramework,
    DataQualityReporter,
    DataSplitConfig,
    CrossValidationConfig
)


class TestDataSplitConfig:
    """Test configuration classes for data splitting."""
    
    def test_data_split_config_defaults(self):
        """Test default configuration values."""
        config = DataSplitConfig()
        
        assert config.test_size == 0.15
        assert config.validation_size == 0.15
        assert config.train_size == 0.70
        assert config.random_state == 42
        assert config.stratify is True
        assert config.n_bins == 5
        assert config.shuffle is True
        
    def test_data_split_config_custom(self):
        """Test custom configuration values."""
        config = DataSplitConfig(
            test_size=0.2,
            validation_size=0.1,
            train_size=0.7,
            random_state=123,
            stratify=False,
            n_bins=3
        )
        
        assert config.test_size == 0.2
        assert config.validation_size == 0.1
        assert config.train_size == 0.7
        assert config.random_state == 123
        assert config.stratify is False
        assert config.n_bins == 3
        
    def test_cv_config_defaults(self):
        """Test default CV configuration values."""
        config = CrossValidationConfig()
        
        assert config.n_folds == 5
        assert config.random_state == 42
        assert config.shuffle is True
        assert config.stratify is True
        assert config.cv_type == 'kfold'


class TestDataQualityReporter:
    """Test data quality reporting functionality."""
    
    @pytest.fixture
    def sample_data(self):
        """Create sample dataset for testing."""
        np.random.seed(42)
        X = pd.DataFrame({
            'feature1': np.random.normal(0, 1, 100),
            'feature2': np.random.normal(5, 2, 100),
            'feature3': np.random.uniform(0, 10, 100),
            'categorical': ['A', 'B', 'C'] * 33 + ['A']  # 100 samples
        })
        y = pd.Series(np.random.normal(-5, 2, 100), name='binding_affinity')
        return X, y
        
    def test_generate_report(self, sample_data):
        """Test report generation."""
        X, y = sample_data
        reporter = DataQualityReporter()
        
        report = reporter.generate_report(X, y)
        
        # Check report structure
        assert 'dataset_overview' in report
        assert 'feature_analysis' in report
        assert 'target_analysis' in report
        assert 'missing_data' in report
        assert 'outlier_analysis' in report
        assert 'distribution_analysis' in report
        assert 'correlation_analysis' in report
        
    def test_dataset_overview(self, sample_data):
        """Test dataset overview generation."""
        X, y = sample_data
        reporter = DataQualityReporter()
        
        overview = reporter._get_dataset_overview(X, y)
        
        assert overview['n_samples'] == 100
        assert overview['n_features'] == 4
        assert overview['target_range'][0] <= overview['target_range'][1]
        assert overview['feature_types']['numeric'] == 3
        assert overview['feature_types']['categorical'] == 1
        
    def test_feature_analysis(self, sample_data):
        """Test feature analysis."""
        X, y = sample_data
        reporter = DataQualityReporter()
        
        analysis = reporter._analyze_features(X)
        
        assert analysis['numeric_features'] == 3
        assert 'feature_stats' in analysis
        assert 'feature_names' in analysis
        assert len(analysis['feature_names']) == 3
        
    def test_target_analysis(self, sample_data):
        """Test target variable analysis."""
        X, y = sample_data
        reporter = DataQualityReporter()
        
        analysis = reporter._analyze_target(y)
        
        assert analysis['count'] == 100
        assert analysis['min'] <= analysis['max']
        assert analysis['mean'] is not None
        assert analysis['std'] is not None
        assert 'skewness' in analysis
        assert 'kurtosis' in analysis
        
    def test_missing_data_analysis(self, sample_data):
        """Test missing data analysis."""
        X, y = sample_data
        # Introduce some missing values
        X.loc[0:5, 'feature1'] = np.nan
        X.loc[10:12, 'feature2'] = np.nan
        
        reporter = DataQualityReporter()
        analysis = reporter._analyze_missing_data(X, y)
        
        assert analysis['features_with_missing'] == 2
        assert analysis['samples_with_missing'] > 0
        assert 'feature1' in analysis['missing_by_feature']
        assert 'feature2' in analysis['missing_by_feature']
        
    def test_outlier_analysis(self, sample_data):
        """Test outlier analysis."""
        X, y = sample_data
        # Introduce some outliers
        X.loc[0, 'feature1'] = 10  # outlier
        y.loc[1] = -20  # outlier
        
        reporter = DataQualityReporter()
        analysis = reporter._analyze_outliers(X, y)
        
        assert 'feature_outliers' in analysis
        assert 'target_outliers' in analysis
        assert analysis['target_outliers']['count'] > 0
        
    def test_correlation_analysis(self, sample_data):
        """Test correlation analysis."""
        X, y = sample_data
        # Make feature1 and feature2 highly correlated
        X['feature2'] = X['feature1'] * 2 + np.random.normal(0, 0.1, 100)
        
        reporter = DataQualityReporter()
        analysis = reporter._analyze_correlations(X, y)
        
        assert 'feature_target_correlations' in analysis
        assert 'high_feature_correlations' in analysis
        assert len(analysis['high_feature_correlations']) > 0
        
    def test_print_summary(self, sample_data, capsys):
        """Test summary printing."""
        X, y = sample_data
        reporter = DataQualityReporter()
        
        report = reporter.generate_report(X, y)
        reporter.print_summary(report)
        
        captured = capsys.readouterr()
        assert "DATA QUALITY REPORT" in captured.out
        assert "Dataset Overview:" in captured.out


class TestDataSplitter:
    """Test data splitting functionality."""
    
    @pytest.fixture
    def sample_data(self):
        """Create sample dataset for testing."""
        np.random.seed(42)
        X = pd.DataFrame({
            'feature1': np.random.normal(0, 1, 1000),
            'feature2': np.random.normal(5, 2, 1000),
            'feature3': np.random.uniform(0, 10, 1000)
        })
        y = pd.Series(np.random.normal(-5, 2, 1000), name='binding_affinity')
        compound_names = pd.Series([f'compound_{i}' for i in range(1000)])
        return X, y, compound_names
        
    def test_data_splitter_initialization(self):
        """Test DataSplitter initialization."""
        splitter = DataSplitter()
        
        assert splitter.config.test_size == 0.15
        assert splitter.config.validation_size == 0.15
        assert splitter.config.train_size == 0.70
        
    def test_split_data_basic(self, sample_data):
        """Test basic data splitting."""
        X, y, compound_names = sample_data
        splitter = DataSplitter()
        
        X_train, X_val, X_test, y_train, y_val, y_test = splitter.split_data(X, y)
        
        # Check shapes
        assert len(X_train) + len(X_val) + len(X_test) == len(X)
        assert len(y_train) + len(y_val) + len(y_test) == len(y)
        
        # Check ratios are approximately correct
        total = len(X)
        assert abs(len(X_train) / total - 0.70) < 0.05
        assert abs(len(X_val) / total - 0.15) < 0.05
        assert abs(len(X_test) / total - 0.15) < 0.05
        
    def test_split_data_with_compound_names(self, sample_data):
        """Test data splitting with compound names."""
        X, y, compound_names = sample_data
        splitter = DataSplitter()
        
        X_train, X_val, X_test, y_train, y_val, y_test = splitter.split_data(X, y, compound_names)
        
        # Check that split info contains compound names
        split_info = splitter.get_split_info()
        assert 'compound_names' in split_info
        assert 'train' in split_info['compound_names']
        assert 'validation' in split_info['compound_names']
        assert 'test' in split_info['compound_names']
        
    def test_stratified_splitting(self, sample_data):
        """Test stratified splitting functionality."""
        X, y, compound_names = sample_data
        config = DataSplitConfig(stratify=True, random_state=42)
        splitter = DataSplitter(config)
        
        X_train, X_val, X_test, y_train, y_val, y_test = splitter.split_data(X, y)
        
        # Check that discretizer was created
        assert splitter.discretizer is not None
        
        # Check that target distributions are similar across splits
        train_mean = y_train.mean()
        val_mean = y_val.mean()
        test_mean = y_test.mean()
        
        # Means should be relatively close (within 20% of overall std)
        overall_std = y.std()
        tolerance = 0.2 * overall_std
        
        assert abs(train_mean - val_mean) < tolerance
        assert abs(train_mean - test_mean) < tolerance
        assert abs(val_mean - test_mean) < tolerance
        
    def test_split_validation(self, sample_data):
        """Test split validation functionality."""
        X, y, compound_names = sample_data
        splitter = DataSplitter()
        
        X_train, X_val, X_test, y_train, y_val, y_test = splitter.split_data(X, y)
        
        validation = splitter.validate_splits(X_train, X_val, X_test, y_train, y_val, y_test)
        
        # Check validation structure
        assert 'size_validation' in validation
        assert 'distribution_validation' in validation
        assert 'feature_validation' in validation
        assert 'overlap_validation' in validation
        assert 'overall_valid' in validation
        
        # All validations should pass for a good split
        assert validation['overall_valid'] is True
        assert validation['size_validation']['status'] is True
        assert validation['feature_validation']['status'] is True
        assert validation['overlap_validation']['status'] is True
        
    def test_split_consistency(self, sample_data):
        """Test that splits are consistent with same random state."""
        X, y, compound_names = sample_data
        
        # Create two splitters with same random state
        splitter1 = DataSplitter(DataSplitConfig(random_state=42))
        splitter2 = DataSplitter(DataSplitConfig(random_state=42))
        
        # Split data with both splitters
        splits1 = splitter1.split_data(X, y)
        splits2 = splitter2.split_data(X, y)
        
        # Check that splits are identical
        for i, (split1, split2) in enumerate(zip(splits1, splits2)):
            if i < 3:  # First three are DataFrames
                pd.testing.assert_frame_equal(split1, split2)
            else:  # Last three are Series
                pd.testing.assert_series_equal(split1, split2)
            
    def test_split_info_storage(self, sample_data):
        """Test that split information is properly stored."""
        X, y, compound_names = sample_data
        splitter = DataSplitter()
        
        splitter.split_data(X, y, compound_names)
        split_info = splitter.get_split_info()
        
        # Check split info structure
        assert 'split_sizes' in split_info
        assert 'split_ratios' in split_info
        assert 'target_statistics' in split_info
        assert 'config' in split_info
        
        # Check that sizes add up correctly
        total_size = (split_info['split_sizes']['train'] + 
                     split_info['split_sizes']['validation'] + 
                     split_info['split_sizes']['test'])
        assert total_size == len(X)


class TestCrossValidationSetup:
    """Test cross-validation setup functionality."""
    
    @pytest.fixture
    def sample_data(self):
        """Create sample dataset for testing."""
        np.random.seed(42)
        X = pd.DataFrame({
            'feature1': np.random.normal(0, 1, 200),
            'feature2': np.random.normal(5, 2, 200),
            'feature3': np.random.uniform(0, 10, 200)
        })
        y = pd.Series(np.random.normal(-5, 2, 200), name='binding_affinity')
        return X, y
        
    def test_cv_setup_initialization(self):
        """Test CrossValidationSetup initialization."""
        cv_setup = CrossValidationSetup()
        
        assert cv_setup.config.n_folds == 5
        assert cv_setup.config.cv_type == 'kfold'
        assert cv_setup.config.random_state == 42
        
    def test_kfold_setup(self, sample_data):
        """Test k-fold cross-validation setup."""
        X, y = sample_data
        cv_setup = CrossValidationSetup()
        
        cv = cv_setup.setup_cv(X, y)
        
        assert cv.n_splits == 5
        assert hasattr(cv, 'split')
        
    def test_stratified_kfold_setup(self, sample_data):
        """Test stratified k-fold cross-validation setup."""
        X, y = sample_data
        config = CrossValidationConfig(cv_type='stratified_kfold', n_folds=5)
        cv_setup = CrossValidationSetup(config)
        
        cv = cv_setup.setup_cv(X, y)
        
        assert cv.n_splits == 5
        assert hasattr(cv, 'split')
        
    def test_loo_setup(self, sample_data):
        """Test leave-one-out cross-validation setup."""
        X, y = sample_data
        config = CrossValidationConfig(cv_type='loo')
        cv_setup = CrossValidationSetup(config)
        
        cv = cv_setup.setup_cv(X, y)
        
        assert hasattr(cv, 'split')
        
    def test_cv_splits_generation(self, sample_data):
        """Test CV splits generation."""
        X, y = sample_data
        cv_setup = CrossValidationSetup()
        
        splits = cv_setup.get_cv_splits(X, y)
        
        assert len(splits) == 5  # 5 folds
        for train_idx, val_idx in splits:
            assert len(train_idx) + len(val_idx) == len(X)
            assert len(set(train_idx).intersection(set(val_idx))) == 0  # No overlap
            
    def test_cv_validation(self, sample_data):
        """Test CV setup validation."""
        X, y = sample_data
        cv_setup = CrossValidationSetup()
        
        validation = cv_setup.validate_cv_setup(X, y)
        
        assert 'dataset_size_check' in validation
        assert 'cv_type_appropriateness' in validation
        assert 'fold_size_validation' in validation
        assert 'overall_valid' in validation
        
        # Should be valid for this dataset
        assert validation['overall_valid'] is True
        
    def test_cv_validation_small_dataset(self):
        """Test CV validation with small dataset."""
        X = pd.DataFrame({'feature1': [1, 2, 3, 4, 5]})
        y = pd.Series([1, 2, 3, 4, 5])
        
        config = CrossValidationConfig(n_folds=5)
        cv_setup = CrossValidationSetup(config)
        
        validation = cv_setup.validate_cv_setup(X, y)
        
        # Should warn about small dataset
        assert 'dataset_size_check' in validation
        assert validation['dataset_size_check']['status'] is False
        
    def test_cv_info_storage(self, sample_data):
        """Test CV info storage."""
        X, y = sample_data
        cv_setup = CrossValidationSetup()
        
        cv_setup.setup_cv(X, y)
        cv_info = cv_setup.get_cv_info()
        
        assert 'cv_type' in cv_info
        assert 'n_folds' in cv_info
        assert 'n_samples' in cv_info
        assert 'n_features' in cv_info
        assert cv_info['n_samples'] == len(X)
        assert cv_info['n_features'] == X.shape[1]


class TestDataValidationFramework:
    """Test comprehensive data validation framework."""
    
    @pytest.fixture
    def sample_data(self):
        """Create sample dataset for testing."""
        np.random.seed(42)
        X = pd.DataFrame({
            'feature1': np.random.normal(0, 1, 500),
            'feature2': np.random.normal(5, 2, 500),
            'feature3': np.random.uniform(0, 10, 500)
        })
        y = pd.Series(np.random.normal(-5, 2, 500), name='binding_affinity')
        compound_names = pd.Series([f'compound_{i}' for i in range(500)])
        return X, y, compound_names
        
    def test_framework_initialization(self):
        """Test framework initialization."""
        framework = DataValidationFramework()
        
        assert isinstance(framework.splitter, DataSplitter)
        assert isinstance(framework.cv_setup, CrossValidationSetup)
        assert isinstance(framework.quality_reporter, DataQualityReporter)
        
    def test_process_dataset_basic(self, sample_data):
        """Test basic dataset processing."""
        X, y, compound_names = sample_data
        framework = DataValidationFramework()
        
        results = framework.process_dataset(X, y, compound_names)
        
        # Check results structure
        assert 'data_splits' in results
        assert 'split_info' in results
        assert 'split_validation' in results
        assert 'cv_setup' in results
        assert 'cv_info' in results
        assert 'cv_validation' in results
        
        # Check data splits
        splits = results['data_splits']
        assert 'X_train' in splits
        assert 'X_val' in splits
        assert 'X_test' in splits
        assert 'y_train' in splits
        assert 'y_val' in splits
        assert 'y_test' in splits
        
    def test_process_dataset_with_quality_report(self, sample_data):
        """Test dataset processing with quality report."""
        X, y, compound_names = sample_data
        framework = DataValidationFramework()
        
        results = framework.process_dataset(X, y, compound_names, generate_quality_report=True)
        
        assert 'quality_report' in results
        quality_report = results['quality_report']
        
        assert 'dataset_overview' in quality_report
        assert 'feature_analysis' in quality_report
        assert 'target_analysis' in quality_report
        
    def test_process_dataset_without_quality_report(self, sample_data):
        """Test dataset processing without quality report."""
        X, y, compound_names = sample_data
        framework = DataValidationFramework()
        
        results = framework.process_dataset(X, y, compound_names, generate_quality_report=False)
        
        assert 'quality_report' not in results
        
    def test_framework_summary(self, sample_data):
        """Test framework summary generation."""
        X, y, compound_names = sample_data
        framework = DataValidationFramework()
        
        framework.process_dataset(X, y, compound_names)
        summary = framework.get_framework_summary()
        
        assert 'dataset_overview' in summary
        assert 'split_summary' in summary
        assert 'cv_summary' in summary
        assert 'validation_status' in summary
        
    def test_print_framework_summary(self, sample_data, capsys):
        """Test framework summary printing."""
        X, y, compound_names = sample_data
        framework = DataValidationFramework()
        
        framework.process_dataset(X, y, compound_names)
        framework.print_framework_summary()
        
        captured = capsys.readouterr()
        assert "DATA VALIDATION FRAMEWORK SUMMARY" in captured.out
        assert "Dataset:" in captured.out
        assert "Data Splits:" in captured.out
        assert "Validation Status:" in captured.out
        
    def test_framework_summary_no_processing(self):
        """Test framework summary when no processing has been done."""
        framework = DataValidationFramework()
        
        summary = framework.get_framework_summary()
        
        assert 'error' in summary
        assert "No framework processing completed yet" in summary['error']
        
    def test_custom_configs(self, sample_data):
        """Test framework with custom configurations."""
        X, y, compound_names = sample_data
        
        split_config = DataSplitConfig(test_size=0.2, validation_size=0.1, train_size=0.7)
        cv_config = CrossValidationConfig(n_folds=3, cv_type='stratified_kfold')
        
        framework = DataValidationFramework(split_config, cv_config)
        
        results = framework.process_dataset(X, y, compound_names)
        
        # Check that custom configs were used
        split_info = results['split_info']
        assert split_info['config']['test_size'] == 0.2
        assert split_info['config']['validation_size'] == 0.1
        
        cv_info = results['cv_info']
        assert cv_info['n_folds'] == 3
        assert cv_info['cv_type'] == 'stratified_kfold'


class TestIntegration:
    """Integration tests for the data splitting framework."""
    
    def test_end_to_end_workflow(self):
        """Test complete end-to-end workflow."""
        # Create a realistic dataset
        np.random.seed(42)
        n_samples = 1000
        
        # Create features similar to molecular descriptors
        X = pd.DataFrame({
            'molecular_weight': np.random.normal(300, 100, n_samples),
            'logp': np.random.normal(2, 1, n_samples),
            'tpsa': np.random.uniform(0, 150, n_samples),
            'rotatable_bonds': np.random.poisson(5, n_samples),
            'hbd': np.random.poisson(3, n_samples),
            'hba': np.random.poisson(6, n_samples),
            'radius_of_gyration': np.random.uniform(3, 8, n_samples),
            'molecular_volume': np.random.uniform(200, 800, n_samples)
        })
        
        # Create binding affinities (lower values = stronger binding)
        y = pd.Series(np.random.normal(-6, 2, n_samples), name='binding_affinity')
        
        compound_names = pd.Series([f'TLR4_compound_{i:04d}' for i in range(n_samples)])
        
        # Process through framework
        framework = DataValidationFramework()
        results = framework.process_dataset(X, y, compound_names, generate_quality_report=True)
        
        # Validate results
        assert results['split_validation']['overall_valid'] is True
        assert results['cv_validation']['overall_valid'] is True
        
        # Check data splits
        splits = results['data_splits']
        total_samples = (len(splits['X_train']) + 
                        len(splits['X_val']) + 
                        len(splits['X_test']))
        assert total_samples == n_samples
        
        # Check that all compounds are accounted for
        all_compounds = (results['split_info']['compound_names']['train'] +
                        results['split_info']['compound_names']['validation'] +
                        results['split_info']['compound_names']['test'])
        assert len(set(all_compounds)) == n_samples
        
        # Test CV splits
        cv = results['cv_setup']
        cv_splits = framework.cv_setup.get_cv_splits(splits['X_train'], splits['y_train'])
        assert len(cv_splits) == 5  # 5-fold CV
        
        for train_idx, val_idx in cv_splits:
            assert len(train_idx) + len(val_idx) == len(splits['X_train'])
            assert len(set(train_idx).intersection(set(val_idx))) == 0
        
    def test_reproducibility(self):
        """Test that results are reproducible with same random state."""
        # Create sample data
        np.random.seed(42)
        X = pd.DataFrame({'feature1': np.random.normal(0, 1, 100)})
        y = pd.Series(np.random.normal(-5, 2, 100))
        
        # Process with same config twice
        framework1 = DataValidationFramework()
        framework2 = DataValidationFramework()
        
        results1 = framework1.process_dataset(X, y)
        results2 = framework2.process_dataset(X, y)
        
        # Check that splits are identical
        splits1 = results1['data_splits']
        splits2 = results2['data_splits']
        
        pd.testing.assert_frame_equal(splits1['X_train'], splits2['X_train'])
        pd.testing.assert_frame_equal(splits1['X_val'], splits2['X_val'])
        pd.testing.assert_frame_equal(splits1['X_test'], splits2['X_test'])
        pd.testing.assert_series_equal(splits1['y_train'], splits2['y_train'])
        pd.testing.assert_series_equal(splits1['y_val'], splits2['y_val'])
        pd.testing.assert_series_equal(splits1['y_test'], splits2['y_test'])


if __name__ == "__main__":
    pytest.main([__file__, "-v"])
```
