## Supplementary material for "Machine Learning-Driven Prediction of TLR4 Binding Affinity: A Comprehensive Molecular Feature Analysis for Drug Discovery": Codebase: test_deep_learning_direct.py

```
#!/usr/bin/env python3
"""
Direct test for deep learning models without package imports.

This script tests the deep learning models by importing them directly.
"""

import sys
import os
import logging
import numpy as np
import pandas as pd
from pathlib import Path

# Set up logging
logging.basicConfig(level=logging.INFO)
logger = logging.getLogger(__name__)

def test_deep_learning_models_direct():
    """Test deep learning models by importing them directly."""
    logger.info("Testing Deep Learning Models (Direct Import)")
    logger.info("=" * 50)
    
    try:
        # Import deep learning models directly
        sys.path.append(str(Path(__file__).parent.parent / "src" / "tlr4_binding" / "ml_components"))
        
        from deep_learning_models import (
            MolecularVoxelizer, VoxelDataset, SMILESDataset, SMILESData,
            MolecularCNN3D, MolecularTransformer, MultiTaskNeuralNetwork,
            DeepLearningTrainer
        )
        logger.info("✓ Successfully imported deep learning models directly")
        
        # Test voxelizer
        voxelizer = MolecularVoxelizer(grid_size=16, resolution=2.0)
        logger.info(f"✓ Created voxelizer: grid_size={voxelizer.grid_size}, resolution={voxelizer.resolution}")
        
        # Test atom type conversion
        assert voxelizer._atom_type_to_number('C') == 6
        assert voxelizer._atom_type_to_number('N') == 7
        assert voxelizer._atom_type_to_number('O') == 8
        logger.info("✓ Atom type conversion working correctly")
        
        # Test PDBQT parsing
        sample_pdbqt_content = """REMARK  VINA RESULT:    -6.5      0.000      0.000
ATOM      1  C   UNL     1      12.345   23.456   34.567  1.00  0.00     0.000 C
ATOM      2  N   UNL     1      13.345   24.456   35.567  1.00  0.00     0.000 N
ATOM      3  O   UNL     1      14.345   25.456   36.567  1.00  0.00     0.000 O
"""
        
        with open("temp_test.pdbqt", 'w') as f:
            f.write(sample_pdbqt_content)
        
        try:
            coords, atom_types = voxelizer._parse_pdbqt("temp_test.pdbqt")
            assert coords is not None
            assert len(coords) == 3
            assert len(atom_types) == 3
            assert coords.shape == (3, 3)
            assert atom_types == [6, 7, 8]  # C, N, O
            logger.info("✓ PDBQT parsing working correctly")
            
            # Test voxel grid creation
            voxel_grid = voxelizer._create_voxel_grid(coords, atom_types)
            assert voxel_grid.shape == (16, 16, 16)
            assert voxel_grid.dtype == np.float32
            logger.info("✓ Voxel grid creation working correctly")
            
        finally:
            if os.path.exists("temp_test.pdbqt"):
                os.unlink("temp_test.pdbqt")
        
        # Test SMILES dataset
        smiles_data = [
            SMILESData(smiles="CCO", compound_name="ethanol", binding_affinity=-5.5),
            SMILESData(smiles="CC(=O)O", compound_name="acetic_acid", binding_affinity=-6.2),
        ]
        dataset = SMILESDataset(smiles_data, vocab_size=100, max_length=64)
        logger.info(f"✓ Created SMILES dataset with {len(dataset)} samples")
        
        # Test tokenization
        tokens = dataset._tokenize_smiles("CCO")
        assert isinstance(tokens, np.ndarray)
        assert len(tokens) > 0
        logger.info(f"✓ SMILES tokenization working: {tokens}")
        
        # Test PyTorch models if available
        try:
            import torch
            
            # Test CNN3D model
            cnn_model = MolecularCNN3D(input_channels=1, num_classes=1)
            x = torch.randn(2, 1, 16, 16, 16)
            output = cnn_model(x)
            assert output.shape == (2, 1)
            logger.info(f"✓ CNN3D model working: input {x.shape} -> output {output.shape}")
            
            # Test Transformer model
            transformer_model = MolecularTransformer(vocab_size=100, d_model=128, nhead=4, num_layers=2)
            x = torch.randint(0, 100, (2, 10))
            output = transformer_model(x)
            assert output.shape == (2, 1)
            logger.info(f"✓ Transformer model working: input {x.shape} -> output {output.shape}")
            
            # Test Multi-task model
            multitask_model = MultiTaskNeuralNetwork(input_dim=100, hidden_dims=[64, 32], num_tasks=3)
            x = torch.randn(4, 100)
            outputs = multitask_model(x)
            assert len(outputs) == 3
            for i, output in enumerate(outputs):
                assert output.shape == (4, 1)
            logger.info(f"✓ Multi-task model working: input {x.shape} -> {len(outputs)} outputs")
            
            # Test trainers
            cnn_trainer = DeepLearningTrainer(model_type="cnn3d", device="cpu")
            transformer_trainer = DeepLearningTrainer(model_type="transformer", device="cpu")
            multitask_trainer = DeepLearningTrainer(model_type="multitask", device="cpu")
            
            logger.info("✓ All deep learning trainers created successfully")
            
            # Test model creation
            cnn_model_created = cnn_trainer.create_model(input_channels=1, num_classes=1)
            assert isinstance(cnn_model_created, MolecularCNN3D)
            logger.info("✓ CNN3D model creation through trainer working")
            
            transformer_model_created = transformer_trainer.create_model(vocab_size=100, d_model=128)
            assert isinstance(transformer_model_created, MolecularTransformer)
            logger.info("✓ Transformer model creation through trainer working")
            
            multitask_model_created = multitask_trainer.create_model(input_dim=100, num_tasks=3)
            assert isinstance(multitask_model_created, MultiTaskNeuralNetwork)
            logger.info("✓ Multi-task model creation through trainer working")
            
        except ImportError:
            logger.warning("PyTorch not available - skipping PyTorch model tests")
        
        # Test deep learning trainer coordinator
        from deep_learning_trainer import DeepLearningModelTrainer
        
        dl_trainer = DeepLearningModelTrainer(models_dir="test_models")
        logger.info("✓ Deep learning model trainer coordinator created")
        
        # Test data preparation methods
        pdbqt_files = ["test1.pdbqt", "test2.pdbqt"]
        binding_affinities = [-5.5, -6.2]
        
        # Create sample PDBQT content
        sample_content = """REMARK  VINA RESULT:    -6.5      0.000      0.000
ATOM      1  C   UNL     1      12.345   23.456   34.567  1.00  0.00     0.000 C
ATOM      2  N   UNL     1      13.345   24.456   35.567  1.00  0.00     0.000 N
ATOM      3  O   UNL     1      14.345   25.456   36.567  1.00  0.00     0.000 O
"""
        
        # Create temporary files
        for filepath in pdbqt_files:
            with open(filepath, 'w') as f:
                f.write(sample_content)
        
        try:
            # Test voxel data preparation
            voxel_dataset = dl_trainer.prepare_voxel_data(pdbqt_files, binding_affinities)
            logger.info(f"✓ Voxel data preparation working: {len(voxel_dataset)} samples")
            
            # Test SMILES data preparation
            smiles_list = ["CCO", "CC(=O)O"]
            compound_names = ["ethanol", "acetic_acid"]
            smiles_dataset = dl_trainer.prepare_smiles_data(smiles_list, compound_names, binding_affinities)
            logger.info(f"✓ SMILES data preparation working: {len(smiles_dataset)} samples")
            
            # Test multi-task data preparation
            features_df = pd.DataFrame({
                'molecular_weight': [46.07, 60.05],
                'logp': [-0.31, -0.17],
                'tpsa': [20.23, 37.30]
            })
            features_tensor, targets_tensors = dl_trainer.prepare_multitask_data(features_df, pd.Series(binding_affinities))
            logger.info(f"✓ Multi-task data preparation working: features {features_tensor.shape}, targets {len(targets_tensors)}")
            
        finally:
            # Clean up temporary files
            for filepath in pdbqt_files:
                if os.path.exists(filepath):
                    os.unlink(filepath)
        
        logger.info("✓ All deep learning model tests passed!")
        return True
        
    except Exception as e:
        logger.error(f"✗ Test failed with error: {str(e)}")
        import traceback
        traceback.print_exc()
        return False


def test_training_workflow():
    """Test a simple training workflow."""
    logger.info("\nTesting Training Workflow")
    logger.info("=" * 30)
    
    try:
        import torch
        from torch.utils.data import DataLoader
        
        # Import models directly
        sys.path.append(str(Path(__file__).parent.parent / "src" / "tlr4_binding" / "ml_components"))
        from deep_learning_models import MolecularVoxelizer, VoxelDataset, DeepLearningTrainer
        
        # Create sample data
        voxelizer = MolecularVoxelizer(grid_size=16, resolution=2.0)
        
        # Create sample PDBQT files
        pdbqt_files = []
        binding_affinities = []
        
        sample_content = """REMARK  VINA RESULT:    -6.5      0.000      0.000
ATOM      1  C   UNL     1      12.345   23.456   34.567  1.00  0.00     0.000 C
ATOM      2  N   UNL     1      13.345   24.456   35.567  1.00  0.00     0.000 N
ATOM      3  O   UNL     1      14.345   25.456   36.567  1.00  0.00     0.000 O
"""
        
        for i in range(5):
            filepath = f"temp_training_{i}.pdbqt"
            with open(filepath, 'w') as f:
                f.write(sample_content)
            pdbqt_files.append(filepath)
            binding_affinities.append(-5.0 - i * 0.3)
        
        try:
            # Prepare voxel data
            voxel_data = []
            for pdbqt_file, affinity in zip(pdbqt_files, binding_affinities):
                voxel_item = voxelizer.voxelize_pdbqt(pdbqt_file)
                if voxel_item is not None:
                    voxel_item = voxel_item._replace(binding_affinity=affinity)
                    voxel_data.append(voxel_item)
            
            dataset = VoxelDataset(voxel_data)
            data_loader = DataLoader(dataset, batch_size=2, shuffle=True, num_workers=0)
            
            # Create and train CNN3D model
            trainer = DeepLearningTrainer(model_type="cnn3d", device="cpu")
            trainer.model = trainer.create_model(input_channels=1, num_classes=1)
            
            logger.info("Training CNN3D model for 2 epochs...")
            history = trainer.train(data_loader, epochs=2, lr=0.001)
            
            logger.info(f"✓ Training completed. Final train loss: {history['train_loss'][-1]:.4f}")
            logger.info(f"✓ Final validation loss: {history['val_loss'][-1]:.4f}")
            
            # Test prediction
            predictions = trainer.predict(data_loader)
            logger.info(f"✓ Predictions shape: {predictions.shape}")
            
        finally:
            # Clean up
            for filepath in pdbqt_files:
                if os.path.exists(filepath):
                    os.unlink(filepath)
        
        logger.info("✓ Training workflow test passed!")
        return True
        
    except ImportError:
        logger.warning("PyTorch not available - skipping training workflow test")
        return True
    except Exception as e:
        logger.error(f"✗ Training workflow test failed with error: {str(e)}")
        import traceback
        traceback.print_exc()
        return False


def main():
    """Run all tests."""
    logger.info("Starting Deep Learning Models Direct Test Suite")
    logger.info("=" * 60)
    
    # Test deep learning models
    success1 = test_deep_learning_models_direct()
    
    # Test training workflow
    success2 = test_training_workflow()
    
    # Clean up
    import shutil
    if os.path.exists("test_models"):
        shutil.rmtree("test_models")
    
    if success1 and success2:
        logger.info("\n🎉 All tests passed successfully!")
        return 0
    else:
        logger.error("\n❌ Some tests failed!")
        return 1


if __name__ == "__main__":
    sys.exit(main())
```
