## Supplementary material for "Machine Learning-Driven Prediction of TLR4 Binding Affinity: A Comprehensive Molecular Feature Analysis for Drug Discovery": Codebase: test_deep_learning_models.py

```
"""
Unit tests for deep learning models.

This module tests CNN, Transformer, and Multi-task neural network
architectures for molecular binding prediction.
"""

import pytest
import numpy as np
import pandas as pd
from pathlib import Path
import tempfile
import os

# Test imports
try:
    import torch
    import torch.nn as nn
    from torch.utils.data import DataLoader
    TORCH_AVAILABLE = True
except ImportError:
    TORCH_AVAILABLE = False

try:
    from rdkit import Chem
    RDKIT_AVAILABLE = True
except ImportError:
    RDKIT_AVAILABLE = False

# Import modules to test
from src.tlr4_binding.ml_components.deep_learning_models import (
    MolecularVoxelizer, VoxelDataset, SMILESDataset, SMILESData,
    MolecularCNN3D, MolecularTransformer, MultiTaskNeuralNetwork,
    DeepLearningTrainer
)
from src.tlr4_binding.ml_components.deep_learning_trainer import DeepLearningModelTrainer


@pytest.fixture
def sample_pdbqt_content():
    """Sample PDBQT content for testing."""
    return """REMARK  VINA RESULT:    -6.5      0.000      0.000
REMARK  VINA RESULT:    -6.2      0.000      0.000
REMARK  VINA RESULT:    -5.8      0.000      0.000
ATOM      1  C   UNL     1      12.345   23.456   34.567  1.00  0.00     0.000 C
ATOM      2  N   UNL     1      13.345   24.456   35.567  1.00  0.00     0.000 N
ATOM      3  O   UNL     1      14.345   25.456   36.567  1.00  0.00     0.000 O
ATOM      4  S   UNL     1      15.345   26.456   37.567  1.00  0.00     0.000 S
"""


@pytest.fixture
def sample_smiles_data():
    """Sample SMILES data for testing."""
    return [
        "CCO",  # Ethanol
        "CC(=O)O",  # Acetic acid
        "c1ccccc1",  # Benzene
        "CCN(CC)CC",  # Triethylamine
        "CC(C)CO"  # Isobutanol
    ]


@pytest.fixture
def sample_features_df():
    """Sample molecular features DataFrame for testing."""
    return pd.DataFrame({
        'molecular_weight': [46.07, 60.05, 78.11, 101.19, 74.12],
        'logp': [-0.31, -0.17, 2.13, 0.15, 0.76],
        'tpsa': [20.23, 37.30, 0.00, 3.24, 20.23],
        'rotatable_bonds': [0, 0, 0, 2, 1],
        'hbd': [1, 1, 0, 0, 1],
        'hba': [1, 2, 0, 1, 1]
    })


class TestMolecularVoxelizer:
    """Test molecular voxelizer functionality."""
    
    def test_voxelizer_initialization(self):
        """Test voxelizer initialization."""
        voxelizer = MolecularVoxelizer(grid_size=32, resolution=1.0)
        assert voxelizer.grid_size == 32
        assert voxelizer.resolution == 1.0
    
    def test_atom_type_to_number(self):
        """Test atom type to number conversion."""
        voxelizer = MolecularVoxelizer()
        
        assert voxelizer._atom_type_to_number('C') == 6
        assert voxelizer._atom_type_to_number('N') == 7
        assert voxelizer._atom_type_to_number('O') == 8
        assert voxelizer._atom_type_to_number('S') == 16
        assert voxelizer._atom_type_to_number('H') == 1
        assert voxelizer._atom_type_to_number('Unknown') == 6  # Default
    
    def test_parse_pdbqt(self, sample_pdbqt_content):
        """Test PDBQT parsing."""
        voxelizer = MolecularVoxelizer()
        
        with tempfile.NamedTemporaryFile(mode='w', suffix='.pdbqt', delete=False) as f:
            f.write(sample_pdbqt_content)
            f.flush()
            
            coords, atom_types = voxelizer._parse_pdbqt(f.name)
            
            assert coords is not None
            assert len(coords) == 4
            assert len(atom_types) == 4
            assert coords.shape == (4, 3)
            assert atom_types == [6, 7, 8, 16]  # C, N, O, S
            
            os.unlink(f.name)
    
    def test_create_voxel_grid(self):
        """Test voxel grid creation."""
        voxelizer = MolecularVoxelizer(grid_size=16, resolution=2.0)
        
        coords = np.array([[0.0, 0.0, 0.0], [2.0, 0.0, 0.0], [0.0, 2.0, 0.0]])
        atom_types = [6, 7, 8]  # C, N, O
        
        voxel_grid = voxelizer._create_voxel_grid(coords, atom_types)
        
        assert voxel_grid.shape == (16, 16, 16)
        assert voxel_grid.dtype == np.float32
        assert np.sum(voxel_grid > 0) > 0  # Should have some non-zero voxels
    
    def test_voxelize_pdbqt(self, sample_pdbqt_content):
        """Test complete voxelization process."""
        voxelizer = MolecularVoxelizer(grid_size=16, resolution=2.0)
        
        with tempfile.NamedTemporaryFile(mode='w', suffix='.pdbqt', delete=False) as f:
            f.write(sample_pdbqt_content)
            f.flush()
            
            voxel_data = voxelizer.voxelize_pdbqt(f.name)
            
            assert voxel_data is not None
            assert voxel_data.voxel_grid.shape == (16, 16, 16)
            assert voxel_data.compound_name == Path(f.name).stem
            assert voxel_data.binding_affinity == 0.0
            
            os.unlink(f.name)


class TestVoxelDataset:
    """Test voxel dataset functionality."""
    
    def test_voxel_dataset_creation(self):
        """Test voxel dataset creation."""
        # Create sample voxel data
        voxel_data = [
            type('VoxelData', (), {
                'voxel_grid': np.random.rand(16, 16, 16).astype(np.float32),
                'compound_name': 'test1',
                'binding_affinity': -5.5
            })(),
            type('VoxelData', (), {
                'voxel_grid': np.random.rand(16, 16, 16).astype(np.float32),
                'compound_name': 'test2',
                'binding_affinity': -6.2
            })()
        ]
        
        dataset = VoxelDataset(voxel_data)
        
        assert len(dataset) == 2
        assert dataset[0][1] == -5.5  # binding affinity
        assert dataset[1][1] == -6.2  # binding affinity


class TestSMILESDataset:
    """Test SMILES dataset functionality."""
    
    def test_smiles_dataset_creation(self, sample_smiles_data):
        """Test SMILES dataset creation."""
        smiles_data = [
            SMILESData(smiles=smiles, compound_name=f"compound_{i}", binding_affinity=-5.0 + i)
            for i, smiles in enumerate(sample_smiles_data)
        ]
        
        dataset = SMILESDataset(smiles_data, vocab_size=100, max_length=64)
        
        assert len(dataset) == 5
        assert dataset.vocab_size == 100
        assert dataset.max_length == 64
    
    def test_smiles_tokenization(self, sample_smiles_data):
        """Test SMILES tokenization."""
        smiles_data = [
            SMILESData(smiles=smiles, compound_name=f"compound_{i}", binding_affinity=-5.0 + i)
            for i, smiles in enumerate(sample_smiles_data)
        ]
        
        dataset = SMILESDataset(smiles_data, vocab_size=100, max_length=64)
        
        # Test tokenization
        tokens = dataset._tokenize_smiles("CCO")
        assert isinstance(tokens, np.ndarray)
        assert len(tokens) > 0
    
    def test_smiles_dataset_getitem(self, sample_smiles_data):
        """Test SMILES dataset __getitem__ method."""
        smiles_data = [
            SMILESData(smiles=smiles, compound_name=f"compound_{i}", binding_affinity=-5.0 + i)
            for i, smiles in enumerate(sample_smiles_data)
        ]
        
        dataset = SMILESDataset(smiles_data, vocab_size=100, max_length=64)
        
        # Test getting item
        token_ids, affinity = dataset[0]
        assert isinstance(token_ids, torch.Tensor) if TORCH_AVAILABLE else True
        assert affinity == -5.0


@pytest.mark.skipif(not TORCH_AVAILABLE, reason="PyTorch not available")
class TestMolecularCNN3D:
    """Test 3D CNN model."""
    
    def test_cnn3d_initialization(self):
        """Test CNN3D model initialization."""
        model = MolecularCNN3D(input_channels=1, num_classes=1, dropout=0.1)
        
        assert model.input_channels == 1
        assert model.num_classes == 1
        assert model.dropout == 0.1
        assert model.use_batch_norm == True
    
    def test_cnn3d_forward(self):
        """Test CNN3D forward pass."""
        model = MolecularCNN3D(input_channels=1, num_classes=1)
        
        # Create sample input (batch_size=2, channels=1, depth=16, height=16, width=16)
        x = torch.randn(2, 1, 16, 16, 16)
        
        output = model(x)
        
        assert output.shape == (2, 1)  # batch_size, num_classes
        assert isinstance(output, torch.Tensor)
    
    def test_cnn3d_without_batch_norm(self):
        """Test CNN3D without batch normalization."""
        model = MolecularCNN3D(input_channels=1, num_classes=1, use_batch_norm=False)
        
        x = torch.randn(2, 1, 16, 16, 16)
        output = model(x)
        
        assert output.shape == (2, 1)


@pytest.mark.skipif(not TORCH_AVAILABLE, reason="PyTorch not available")
class TestMolecularTransformer:
    """Test molecular transformer model."""
    
    def test_transformer_initialization(self):
        """Test transformer model initialization."""
        model = MolecularTransformer(vocab_size=100, d_model=128, nhead=4, num_layers=2)
        
        assert model.vocab_size == 100
        assert model.d_model == 128
        assert model.nhead == 4
        assert model.num_layers == 2
    
    def test_transformer_forward(self):
        """Test transformer forward pass."""
        model = MolecularTransformer(vocab_size=100, d_model=128, nhead=4, num_layers=2, max_length=64)
        
        # Create sample input (batch_size=2, sequence_length=10)
        x = torch.randint(0, 100, (2, 10))
        
        output = model(x)
        
        assert output.shape == (2, 1)  # batch_size, num_classes
        assert isinstance(output, torch.Tensor)
    
    def test_transformer_different_sequence_lengths(self):
        """Test transformer with different sequence lengths."""
        model = MolecularTransformer(vocab_size=100, d_model=128, nhead=4, num_layers=2, max_length=64)
        
        # Test with different sequence lengths
        for seq_len in [5, 10, 20, 30]:
            x = torch.randint(0, 100, (2, seq_len))
            output = model(x)
            assert output.shape == (2, 1)


@pytest.mark.skipif(not TORCH_AVAILABLE, reason="PyTorch not available")
class TestMultiTaskNeuralNetwork:
    """Test multi-task neural network."""
    
    def test_multitask_initialization(self):
        """Test multi-task model initialization."""
        model = MultiTaskNeuralNetwork(input_dim=100, hidden_dims=[64, 32], num_tasks=3)
        
        assert model.input_dim == 100
        assert model.hidden_dims == [64, 32]
        assert model.num_tasks == 3
        assert len(model.task_heads) == 3
    
    def test_multitask_forward(self):
        """Test multi-task model forward pass."""
        model = MultiTaskNeuralNetwork(input_dim=100, hidden_dims=[64, 32], num_tasks=3)
        
        # Create sample input (batch_size=4, input_dim=100)
        x = torch.randn(4, 100)
        
        outputs = model(x)
        
        assert len(outputs) == 3  # Number of tasks
        for output in outputs:
            assert output.shape == (4, 1)  # batch_size, 1
            assert isinstance(output, torch.Tensor)
    
    def test_multitask_single_task_prediction(self):
        """Test single task prediction."""
        model = MultiTaskNeuralNetwork(input_dim=100, hidden_dims=[64, 32], num_tasks=3)
        
        x = torch.randn(4, 100)
        
        # Test prediction for each task
        for task_idx in range(3):
            output = model.predict_single_task(x, task_idx)
            assert output.shape == (4, 1)
            assert isinstance(output, torch.Tensor)


@pytest.mark.skipif(not TORCH_AVAILABLE, reason="PyTorch not available")
class TestDeepLearningTrainer:
    """Test deep learning trainer."""
    
    def test_trainer_initialization(self):
        """Test trainer initialization."""
        trainer = DeepLearningTrainer(model_type="cnn3d", device="cpu")
        
        assert trainer.model_type == "cnn3d"
        assert trainer.device.type == "cpu"
        assert trainer.model is None
        assert trainer.optimizer is None
    
    def test_create_model(self):
        """Test model creation."""
        trainer = DeepLearningTrainer(model_type="cnn3d")
        
        model = trainer.create_model(input_channels=1, num_classes=1)
        
        assert isinstance(model, MolecularCNN3D)
        assert model.input_channels == 1
        assert model.num_classes == 1
    
    def test_create_transformer_model(self):
        """Test transformer model creation."""
        trainer = DeepLearningTrainer(model_type="transformer")
        
        model = trainer.create_model(vocab_size=100, d_model=128)
        
        assert isinstance(model, MolecularTransformer)
        assert model.vocab_size == 100
        assert model.d_model == 128
    
    def test_create_multitask_model(self):
        """Test multi-task model creation."""
        trainer = DeepLearningTrainer(model_type="multitask")
        
        model = trainer.create_model(input_dim=100, num_tasks=3)
        
        assert isinstance(model, MultiTaskNeuralNetwork)
        assert model.input_dim == 100
        assert model.num_tasks == 3


class TestDeepLearningModelTrainer:
    """Test deep learning model trainer coordinator."""
    
    def test_trainer_initialization(self):
        """Test trainer coordinator initialization."""
        trainer = DeepLearningModelTrainer(models_dir="test_models")
        
        assert trainer.models_dir == Path("test_models")
        assert isinstance(trainer.voxelizer, MolecularVoxelizer)
        assert len(trainer.trained_models) == 0
        assert len(trainer.training_results) == 0
    
    def test_prepare_voxel_data(self, sample_pdbqt_content):
        """Test voxel data preparation."""
        trainer = DeepLearningModelTrainer()
        
        with tempfile.NamedTemporaryFile(mode='w', suffix='.pdbqt', delete=False) as f:
            f.write(sample_pdbqt_content)
            f.flush()
            
            pdbqt_files = [f.name]
            binding_affinities = [-5.5]
            
            dataset = trainer.prepare_voxel_data(pdbqt_files, binding_affinities)
            
            assert isinstance(dataset, VoxelDataset)
            assert len(dataset) == 1
            
            os.unlink(f.name)
    
    def test_prepare_smiles_data(self, sample_smiles_data):
        """Test SMILES data preparation."""
        trainer = DeepLearningModelTrainer()
        
        compound_names = [f"compound_{i}" for i in range(len(sample_smiles_data))]
        binding_affinities = [-5.0 + i for i in range(len(sample_smiles_data))]
        
        dataset = trainer.prepare_smiles_data(
            sample_smiles_data, compound_names, binding_affinities
        )
        
        assert isinstance(dataset, SMILESDataset)
        assert len(dataset) == len(sample_smiles_data)
    
    def test_prepare_multitask_data(self, sample_features_df):
        """Test multi-task data preparation."""
        trainer = DeepLearningModelTrainer()
        
        binding_affinities = pd.Series([-5.5, -6.2, -4.8, -7.1, -5.9])
        
        features_tensor, targets_tensors = trainer.prepare_multitask_data(
            sample_features_df, binding_affinities
        )
        
        if TORCH_AVAILABLE:
            assert isinstance(features_tensor, torch.Tensor)
            assert features_tensor.shape[0] == len(sample_features_df)
            assert len(targets_tensors) == 1  # Only binding affinity
            assert isinstance(targets_tensors[0], torch.Tensor)
    
    def test_calculate_metrics(self):
        """Test metrics calculation."""
        trainer = DeepLearningModelTrainer()
        
        y_true = [1.0, 2.0, 3.0, 4.0, 5.0]
        y_pred = np.array([1.1, 1.9, 3.1, 3.9, 5.1])
        
        metrics = trainer._calculate_metrics(y_true, y_pred)
        
        assert 'mse' in metrics
        assert 'rmse' in metrics
        assert 'mae' in metrics
        assert 'r2' in metrics
        
        assert metrics['mse'] > 0
        assert metrics['rmse'] > 0
        assert metrics['mae'] > 0
        assert metrics['r2'] > 0  # Should be positive for good predictions


class TestIntegration:
    """Integration tests for deep learning models."""
    
    @pytest.mark.skipif(not TORCH_AVAILABLE, reason="PyTorch not available")
    def test_cnn3d_training_workflow(self, sample_pdbqt_content):
        """Test complete CNN3D training workflow."""
        trainer = DeepLearningModelTrainer()
        
        # Create temporary PDBQT files
        pdbqt_files = []
        binding_affinities = []
        
        for i in range(3):
            with tempfile.NamedTemporaryFile(mode='w', suffix='.pdbqt', delete=False) as f:
                f.write(sample_pdbqt_content)
                f.flush()
                pdbqt_files.append(f.name)
                binding_affinities.append(-5.0 - i)
        
        try:
            # Prepare data
            dataset = trainer.prepare_voxel_data(pdbqt_files, binding_affinities)
            
            # Create data loader
            data_loader = DataLoader(dataset, batch_size=2, shuffle=True, num_workers=0)
            
            # Create and train model
            dl_trainer = DeepLearningTrainer(model_type="cnn3d", device="cpu")
            dl_trainer.model = dl_trainer.create_model()
            
            # Train for a few epochs
            history = dl_trainer.train(data_loader, epochs=2, lr=0.001)
            
            assert 'train_loss' in history
            assert 'val_loss' in history
            assert len(history['train_loss']) == 2
            
        finally:
            # Clean up temporary files
            for f in pdbqt_files:
                if os.path.exists(f):
                    os.unlink(f)
    
    @pytest.mark.skipif(not TORCH_AVAILABLE, reason="PyTorch not available")
    def test_transformer_training_workflow(self, sample_smiles_data):
        """Test complete Transformer training workflow."""
        trainer = DeepLearningModelTrainer()
        
        compound_names = [f"compound_{i}" for i in range(len(sample_smiles_data))]
        binding_affinities = [-5.0 + i for i in range(len(sample_smiles_data))]
        
        # Prepare data
        dataset = trainer.prepare_smiles_data(
            sample_smiles_data, compound_names, binding_affinities
        )
        
        # Create data loader
        data_loader = DataLoader(dataset, batch_size=2, shuffle=True, num_workers=0)
        
        # Create and train model
        dl_trainer = DeepLearningTrainer(model_type="transformer", device="cpu")
        dl_trainer.model = dl_trainer.create_model(vocab_size=100, max_length=64)
        
        # Train for a few epochs
        history = dl_trainer.train(data_loader, epochs=2, lr=0.001)
        
        assert 'train_loss' in history
        assert 'val_loss' in history
        assert len(history['train_loss']) == 2
    
    @pytest.mark.skipif(not TORCH_AVAILABLE, reason="PyTorch not available")
    def test_multitask_training_workflow(self, sample_features_df):
        """Test complete Multi-task training workflow."""
        trainer = DeepLearningModelTrainer()
        
        binding_affinities = pd.Series([-5.5, -6.2, -4.8, -7.1, -5.9])
        
        # Prepare data
        features_tensor, targets_tensors = trainer.prepare_multitask_data(
            sample_features_df, binding_affinities
        )
        
        # Create custom dataset
        class TestDataset:
            def __init__(self, features, targets):
                self.features = features
                self.targets = targets
            
            def __len__(self):
                return len(self.features)
            
            def __getitem__(self, idx):
                return self.features[idx], self.targets[0][idx]
        
        dataset = TestDataset(features_tensor, targets_tensors)
        data_loader = DataLoader(dataset, batch_size=2, shuffle=True, num_workers=0)
        
        # Create and train model
        dl_trainer = DeepLearningTrainer(model_type="multitask", device="cpu")
        dl_trainer.model = dl_trainer.create_model(
            input_dim=features_tensor.shape[1], num_tasks=1
        )
        
        # Train for a few epochs
        history = dl_trainer.train(data_loader, epochs=2, lr=0.001)
        
        assert 'train_loss' in history
        assert 'val_loss' in history
        assert len(history['train_loss']) == 2


if __name__ == "__main__":
    pytest.main([__file__])
```
