## Supplementary material for "Machine Learning-Driven Prediction of TLR4 Binding Affinity: A Comprehensive Molecular Feature Analysis for Drug Discovery": Codebase: test_deep_learning_standalone.py

```
#!/usr/bin/env python3
"""
Standalone test for deep learning models.

This script tests the deep learning models without importing the full package.
"""

import sys
import os
import logging
import numpy as np
import pandas as pd
from pathlib import Path

# Add src to path
sys.path.append(str(Path(__file__).parent.parent / "src"))

# Set up logging
logging.basicConfig(level=logging.INFO)
logger = logging.getLogger(__name__)

def test_deep_learning_models():
    """Test deep learning models directly."""
    logger.info("Testing Deep Learning Models")
    logger.info("=" * 40)
    
    try:
        # Test imports
        from tlr4_binding.ml_components.deep_learning_models import (
            MolecularVoxelizer, VoxelDataset, SMILESDataset, SMILESData,
            MolecularCNN3D, MolecularTransformer, MultiTaskNeuralNetwork,
            DeepLearningTrainer
        )
        logger.info("✓ Successfully imported deep learning models")
        
        # Test voxelizer
        voxelizer = MolecularVoxelizer(grid_size=16, resolution=2.0)
        logger.info(f"✓ Created voxelizer: grid_size={voxelizer.grid_size}, resolution={voxelizer.resolution}")
        
        # Test atom type conversion
        assert voxelizer._atom_type_to_number('C') == 6
        assert voxelizer._atom_type_to_number('N') == 7
        assert voxelizer._atom_type_to_number('O') == 8
        logger.info("✓ Atom type conversion working correctly")
        
        # Test SMILES dataset
        smiles_data = [
            SMILESData(smiles="CCO", compound_name="ethanol", binding_affinity=-5.5),
            SMILESData(smiles="CC(=O)O", compound_name="acetic_acid", binding_affinity=-6.2),
        ]
        dataset = SMILESDataset(smiles_data, vocab_size=100, max_length=64)
        logger.info(f"✓ Created SMILES dataset with {len(dataset)} samples")
        
        # Test tokenization
        tokens = dataset._tokenize_smiles("CCO")
        assert isinstance(tokens, np.ndarray)
        assert len(tokens) > 0
        logger.info(f"✓ SMILES tokenization working: {tokens}")
        
        # Test PyTorch models if available
        try:
            import torch
            
            # Test CNN3D model
            cnn_model = MolecularCNN3D(input_channels=1, num_classes=1)
            x = torch.randn(2, 1, 16, 16, 16)
            output = cnn_model(x)
            assert output.shape == (2, 1)
            logger.info(f"✓ CNN3D model working: input {x.shape} -> output {output.shape}")
            
            # Test Transformer model
            transformer_model = MolecularTransformer(vocab_size=100, d_model=128, nhead=4, num_layers=2)
            x = torch.randint(0, 100, (2, 10))
            output = transformer_model(x)
            assert output.shape == (2, 1)
            logger.info(f"✓ Transformer model working: input {x.shape} -> output {output.shape}")
            
            # Test Multi-task model
            multitask_model = MultiTaskNeuralNetwork(input_dim=100, hidden_dims=[64, 32], num_tasks=3)
            x = torch.randn(4, 100)
            outputs = multitask_model(x)
            assert len(outputs) == 3
            for i, output in enumerate(outputs):
                assert output.shape == (4, 1)
            logger.info(f"✓ Multi-task model working: input {x.shape} -> {len(outputs)} outputs")
            
            # Test trainers
            cnn_trainer = DeepLearningTrainer(model_type="cnn3d", device="cpu")
            transformer_trainer = DeepLearningTrainer(model_type="transformer", device="cpu")
            multitask_trainer = DeepLearningTrainer(model_type="multitask", device="cpu")
            
            logger.info("✓ All deep learning trainers created successfully")
            
        except ImportError:
            logger.warning("PyTorch not available - skipping PyTorch model tests")
        
        # Test deep learning trainer coordinator
        from tlr4_binding.ml_components.deep_learning_trainer import DeepLearningModelTrainer
        
        dl_trainer = DeepLearningModelTrainer(models_dir="test_models")
        logger.info("✓ Deep learning model trainer coordinator created")
        
        # Test data preparation methods
        pdbqt_files = ["test1.pdbqt", "test2.pdbqt"]
        binding_affinities = [-5.5, -6.2]
        
        # Create sample PDBQT content
        sample_content = """REMARK  VINA RESULT:    -6.5      0.000      0.000
ATOM      1  C   UNL     1      12.345   23.456   34.567  1.00  0.00     0.000 C
ATOM      2  N   UNL     1      13.345   24.456   35.567  1.00  0.00     0.000 N
ATOM      3  O   UNL     1      14.345   25.456   36.567  1.00  0.00     0.000 O
"""
        
        # Create temporary files
        for filepath in pdbqt_files:
            with open(filepath, 'w') as f:
                f.write(sample_content)
        
        try:
            # Test voxel data preparation
            voxel_dataset = dl_trainer.prepare_voxel_data(pdbqt_files, binding_affinities)
            logger.info(f"✓ Voxel data preparation working: {len(voxel_dataset)} samples")
            
            # Test SMILES data preparation
            smiles_list = ["CCO", "CC(=O)O"]
            compound_names = ["ethanol", "acetic_acid"]
            smiles_dataset = dl_trainer.prepare_smiles_data(smiles_list, compound_names, binding_affinities)
            logger.info(f"✓ SMILES data preparation working: {len(smiles_dataset)} samples")
            
            # Test multi-task data preparation
            features_df = pd.DataFrame({
                'molecular_weight': [46.07, 60.05],
                'logp': [-0.31, -0.17],
                'tpsa': [20.23, 37.30]
            })
            features_tensor, targets_tensors = dl_trainer.prepare_multitask_data(features_df, pd.Series(binding_affinities))
            logger.info(f"✓ Multi-task data preparation working: features {features_tensor.shape}, targets {len(targets_tensors)}")
            
        finally:
            # Clean up temporary files
            for filepath in pdbqt_files:
                if os.path.exists(filepath):
                    os.unlink(filepath)
        
        logger.info("✓ All deep learning model tests passed!")
        return True
        
    except Exception as e:
        logger.error(f"✗ Test failed with error: {str(e)}")
        import traceback
        traceback.print_exc()
        return False


def test_integration_with_main_trainer():
    """Test integration with main trainer."""
    logger.info("\nTesting Integration with Main Trainer")
    logger.info("=" * 40)
    
    try:
        from tlr4_binding.ml_components.trainer import MLModelTrainer
        
        # Create trainer with deep learning enabled
        trainer = MLModelTrainer(models_dir="test_models", include_deep_learning=True)
        logger.info("✓ Main trainer created with deep learning support")
        
        # Check if deep learning trainer is available
        if trainer.deep_learning_trainer is not None:
            logger.info("✓ Deep learning trainer is available in main trainer")
        else:
            logger.warning("⚠ Deep learning trainer not available in main trainer")
        
        # Test model summary (should include deep learning models)
        summary = trainer.get_model_summary(include_deep_learning=True)
        logger.info(f"✓ Model summary method working: {len(summary)} columns")
        
        logger.info("✓ Integration tests passed!")
        return True
        
    except Exception as e:
        logger.error(f"✗ Integration test failed with error: {str(e)}")
        import traceback
        traceback.print_exc()
        return False


def main():
    """Run all tests."""
    logger.info("Starting Deep Learning Models Test Suite")
    logger.info("=" * 50)
    
    # Test deep learning models
    success1 = test_deep_learning_models()
    
    # Test integration
    success2 = test_integration_with_main_trainer()
    
    # Clean up
    import shutil
    if os.path.exists("test_models"):
        shutil.rmtree("test_models")
    
    if success1 and success2:
        logger.info("\n🎉 All tests passed successfully!")
        return 0
    else:
        logger.error("\n❌ Some tests failed!")
        return 1


if __name__ == "__main__":
    sys.exit(main())
```
