## Supplementary material for "Machine Learning-Driven Prediction of TLR4 Binding Affinity: A Comprehensive Molecular Feature Analysis for Drug Discovery": Codebase: test_ensemble_models.py

```
"""
Unit tests for ensemble and hybrid models.

This module tests the ensemble model implementations including
stacked ensembles, weighted ensembles, and physics-informed models.
"""

import pytest
import pandas as pd
import numpy as np
from unittest.mock import Mock, patch, MagicMock
import tempfile
import shutil
from pathlib import Path

# Import ensemble models
from src.tlr4_binding.ml_components.ensemble_models import (
    StackedEnsemble, WeightedEnsemble, PhysicsInformedNeuralNetwork,
    PhysicsInformedEnsemble, EnsembleModelTrainer, EnsemblePrediction
)

# Import base model trainers for testing
from src.tlr4_binding.ml_components.trainer import (
    RandomForestTrainer, SVRTrainer, XGBoostTrainer, LightGBMTrainer
)


class TestEnsemblePrediction:
    """Test EnsemblePrediction data class."""
    
    def test_ensemble_prediction_creation(self):
        """Test creating EnsemblePrediction object."""
        prediction = EnsemblePrediction(
            prediction=1.5,
            uncertainty=0.2,
            individual_predictions={'model1': 1.4, 'model2': 1.6},
            weights={'model1': 0.6, 'model2': 0.4},
            confidence_interval=(1.1, 1.9)
        )
        
        assert prediction.prediction == 1.5
        assert prediction.uncertainty == 0.2
        assert prediction.individual_predictions == {'model1': 1.4, 'model2': 1.6}
        assert prediction.weights == {'model1': 0.6, 'model2': 0.4}
        assert prediction.confidence_interval == (1.1, 1.9)


class TestStackedEnsemble:
    """Test StackedEnsemble implementation."""
    
    @pytest.fixture
    def sample_data(self):
        """Create sample data for testing."""
        np.random.seed(42)
        X = pd.DataFrame({
            'feature1': np.random.randn(100),
            'feature2': np.random.randn(100),
            'feature3': np.random.randn(100)
        })
        y = pd.Series(2 * X['feature1'] + 3 * X['feature2'] + np.random.randn(100) * 0.1)
        return X, y
    
    @pytest.fixture
    def mock_base_models(self):
        """Create mock base models for testing."""
        models = {}
        
        # Mock Random Forest
        rf_trainer = Mock(spec=RandomForestTrainer)
        rf_model = Mock()
        rf_trainer.train.return_value = rf_model
        rf_trainer.predict.return_value = np.array([1.0, 2.0, 3.0])
        models['random_forest'] = rf_trainer
        
        # Mock SVR
        svr_trainer = Mock(spec=SVRTrainer)
        svr_model = Mock()
        svr_trainer.train.return_value = svr_model
        svr_trainer.predict.return_value = np.array([1.1, 2.1, 3.1])
        models['svr'] = svr_trainer
        
        return models
    
    def test_stacked_ensemble_initialization(self, mock_base_models):
        """Test StackedEnsemble initialization."""
        ensemble = StackedEnsemble(mock_base_models, cv_folds=3)
        
        assert ensemble.base_models == mock_base_models
        assert ensemble.cv_folds == 3
        assert ensemble.trained_base_models == {}
        assert ensemble.meta_learner_trained is None
    
    @patch('src.tlr4_binding.ml_components.ensemble_models.KFold')
    @patch('src.tlr4_binding.ml_components.ensemble_models.Ridge')
    def test_stacked_ensemble_fit(self, mock_ridge, mock_kfold, sample_data, mock_base_models):
        """Test StackedEnsemble fitting."""
        X, y = sample_data
        
        # Mock KFold
        mock_kf = Mock()
        mock_kf.split.return_value = [(range(0, 80), range(80, 100))]
        mock_kfold.return_value = mock_kf
        
        # Mock Ridge
        mock_meta_learner = Mock()
        mock_ridge.return_value = mock_meta_learner
        
        ensemble = StackedEnsemble(mock_base_models, cv_folds=1)
        ensemble.fit(X, y)
        
        # Verify base models were trained
        for model_name, trainer in mock_base_models.items():
            assert trainer.train.call_count >= 1
        
        # Verify meta-learner was fitted
        mock_meta_learner.fit.assert_called_once()
        assert ensemble.meta_learner_trained == mock_meta_learner
    
    def test_stacked_ensemble_predict(self, sample_data, mock_base_models):
        """Test StackedEnsemble prediction."""
        X, y = sample_data
        
        # Create ensemble and fit
        ensemble = StackedEnsemble(mock_base_models, cv_folds=1)
        
        # Mock the trained state
        ensemble.trained_base_models = mock_base_models
        ensemble.meta_learner_trained = Mock()
        ensemble.meta_learner_trained.predict.return_value = np.array([1.5, 2.5, 3.5])
        
        predictions = ensemble.predict(X.iloc[:3])
        
        assert len(predictions) == 3
        assert np.allclose(predictions, [1.5, 2.5, 3.5])
    
    def test_stacked_ensemble_predict_with_uncertainty(self, sample_data, mock_base_models):
        """Test StackedEnsemble prediction with uncertainty."""
        X, y = sample_data
        
        # Create ensemble and fit
        ensemble = StackedEnsemble(mock_base_models, cv_folds=1)
        
        # Mock the trained state
        ensemble.trained_base_models = mock_base_models
        ensemble.meta_learner_trained = Mock()
        ensemble.meta_learner_trained.predict.return_value = np.array([1.5, 2.5])
        
        uncertainty_predictions = ensemble.predict_with_uncertainty(X.iloc[:2])
        
        assert len(uncertainty_predictions) == 2
        assert all(isinstance(pred, EnsemblePrediction) for pred in uncertainty_predictions)
        assert all('prediction' in pred.__dict__ for pred in uncertainty_predictions)
        assert all('uncertainty' in pred.__dict__ for pred in uncertainty_predictions)


class TestWeightedEnsemble:
    """Test WeightedEnsemble implementation."""
    
    @pytest.fixture
    def sample_data(self):
        """Create sample data for testing."""
        np.random.seed(42)
        X = pd.DataFrame({
            'feature1': np.random.randn(100),
            'feature2': np.random.randn(100),
            'feature3': np.random.randn(100)
        })
        y = pd.Series(2 * X['feature1'] + 3 * X['feature2'] + np.random.randn(100) * 0.1)
        return X, y
    
    @pytest.fixture
    def mock_base_models(self):
        """Create mock base models for testing."""
        models = {}
        
        # Mock Random Forest
        rf_trainer = Mock(spec=RandomForestTrainer)
        rf_model = Mock()
        rf_trainer.train.return_value = rf_model
        rf_trainer.predict.return_value = np.array([1.0, 2.0, 3.0])
        models['random_forest'] = rf_trainer
        
        # Mock SVR
        svr_trainer = Mock(spec=SVRTrainer)
        svr_model = Mock()
        svr_trainer.train.return_value = svr_model
        svr_trainer.predict.return_value = np.array([1.1, 2.1, 3.1])
        models['svr'] = svr_trainer
        
        return models
    
    def test_weighted_ensemble_initialization(self, mock_base_models):
        """Test WeightedEnsemble initialization."""
        ensemble = WeightedEnsemble(mock_base_models, weight_method='performance')
        
        assert ensemble.base_models == mock_base_models
        assert ensemble.weight_method == 'performance'
        assert ensemble.trained_base_models == {}
        assert ensemble.weights == {}
    
    def test_weighted_ensemble_fit(self, sample_data, mock_base_models):
        """Test WeightedEnsemble fitting."""
        X, y = sample_data
        X_train, X_val = X.iloc[:80], X.iloc[80:]
        y_train, y_val = y.iloc[:80], y.iloc[80:]
        
        ensemble = WeightedEnsemble(mock_base_models, weight_method='performance')
        ensemble.fit(X_train, y_train, X_val, y_val)
        
        # Verify base models were trained
        for model_name, trainer in mock_base_models.items():
            assert trainer.train.call_count >= 1
        
        # Verify weights were calculated
        assert len(ensemble.weights) > 0
        assert abs(sum(ensemble.weights.values()) - 1.0) < 1e-6  # Weights should sum to 1
    
    def test_weighted_ensemble_predict(self, sample_data, mock_base_models):
        """Test WeightedEnsemble prediction."""
        X, y = sample_data
        
        # Create ensemble and fit
        ensemble = WeightedEnsemble(mock_base_models, weight_method='performance')
        
        # Mock the trained state
        ensemble.trained_base_models = mock_base_models
        ensemble.weights = {'random_forest': 0.6, 'svr': 0.4}
        
        predictions = ensemble.predict(X.iloc[:3])
        
        assert len(predictions) == 3
        # Predictions should be weighted combination
        expected = 0.6 * np.array([1.0, 2.0, 3.0]) + 0.4 * np.array([1.1, 2.1, 3.1])
        assert np.allclose(predictions, expected)
    
    def test_weighted_ensemble_predict_with_uncertainty(self, sample_data, mock_base_models):
        """Test WeightedEnsemble prediction with uncertainty."""
        X, y = sample_data
        
        # Create ensemble and fit
        ensemble = WeightedEnsemble(mock_base_models, weight_method='performance')
        
        # Mock the trained state
        ensemble.trained_base_models = mock_base_models
        ensemble.weights = {'random_forest': 0.6, 'svr': 0.4}
        
        uncertainty_predictions = ensemble.predict_with_uncertainty(X.iloc[:2])
        
        assert len(uncertainty_predictions) == 2
        assert all(isinstance(pred, EnsemblePrediction) for pred in uncertainty_predictions)
        assert all('prediction' in pred.__dict__ for pred in uncertainty_predictions)
        assert all('uncertainty' in pred.__dict__ for pred in uncertainty_predictions)


class TestPhysicsInformedNeuralNetwork:
    """Test PhysicsInformedNeuralNetwork implementation."""
    
    @pytest.fixture
    def sample_data(self):
        """Create sample data for testing."""
        np.random.seed(42)
        X = pd.DataFrame({
            'molecular_weight': np.random.uniform(100, 500, 100),
            'logp': np.random.uniform(-2, 5, 100),
            'tpsa': np.random.uniform(20, 150, 100),
            'feature4': np.random.randn(100)
        })
        y = pd.Series(-2 * X['molecular_weight'] / 1000 + np.random.randn(100) * 0.1)
        return X, y
    
    @pytest.mark.skipif(not hasattr(__import__('torch'), 'torch'), reason="PyTorch not available")
    def test_physics_informed_nn_initialization(self):
        """Test PhysicsInformedNeuralNetwork initialization."""
        model = PhysicsInformedNeuralNetwork(
            input_dim=4,
            hidden_dims=[128, 64],
            dropout=0.1,
            use_thermodynamic_constraints=True
        )
        
        assert model.input_dim == 4
        assert model.hidden_dims == [128, 64]
        assert model.dropout == 0.1
        assert model.use_thermodynamic_constraints is True
    
    @pytest.mark.skipif(not hasattr(__import__('torch'), 'torch'), reason="PyTorch not available")
    def test_physics_informed_nn_forward(self, sample_data):
        """Test PhysicsInformedNeuralNetwork forward pass."""
        X, y = sample_data
        
        model = PhysicsInformedNeuralNetwork(
            input_dim=4,
            hidden_dims=[128, 64],
            dropout=0.1,
            use_thermodynamic_constraints=True
        )
        
        import torch
        X_tensor = torch.tensor(X.values, dtype=torch.float32)
        output = model(X_tensor)
        
        assert output.shape == (len(X), 1)
        # Check that output is within thermodynamic constraints
        assert torch.all(output >= model.min_binding_affinity)
        assert torch.all(output <= model.max_binding_affinity)
    
    @pytest.mark.skipif(not hasattr(__import__('torch'), 'torch'), reason="PyTorch not available")
    def test_physics_informed_nn_physics_loss(self, sample_data):
        """Test PhysicsInformedNeuralNetwork physics loss calculation."""
        X, y = sample_data
        
        model = PhysicsInformedNeuralNetwork(
            input_dim=4,
            hidden_dims=[128, 64],
            dropout=0.1,
            use_thermodynamic_constraints=True
        )
        
        import torch
        X_tensor = torch.tensor(X.values, dtype=torch.float32)
        predictions = model(X_tensor)
        
        physics_loss = model.physics_loss(predictions, X_tensor)
        
        assert isinstance(physics_loss, torch.Tensor)
        assert physics_loss.item() >= 0  # Physics loss should be non-negative


class TestPhysicsInformedEnsemble:
    """Test PhysicsInformedEnsemble implementation."""
    
    @pytest.fixture
    def sample_data(self):
        """Create sample data for testing."""
        np.random.seed(42)
        X = pd.DataFrame({
            'molecular_weight': np.random.uniform(100, 500, 100),
            'logp': np.random.uniform(-2, 5, 100),
            'tpsa': np.random.uniform(20, 150, 100),
            'feature4': np.random.randn(100)
        })
        y = pd.Series(-2 * X['molecular_weight'] / 1000 + np.random.randn(100) * 0.1)
        return X, y
    
    @pytest.fixture
    def mock_base_models(self):
        """Create mock base models for testing."""
        models = {}
        
        # Mock Random Forest
        rf_trainer = Mock(spec=RandomForestTrainer)
        rf_model = Mock()
        rf_trainer.train.return_value = rf_model
        rf_trainer.predict.return_value = np.array([1.0, 2.0, 3.0])
        models['random_forest'] = rf_trainer
        
        return models
    
    @pytest.mark.skipif(not hasattr(__import__('torch'), 'torch'), reason="PyTorch not available")
    def test_physics_informed_ensemble_initialization(self, mock_base_models):
        """Test PhysicsInformedEnsemble initialization."""
        physics_model = PhysicsInformedNeuralNetwork(input_dim=4)
        
        ensemble = PhysicsInformedEnsemble(
            base_models=mock_base_models,
            physics_model=physics_model,
            physics_weight=0.3
        )
        
        assert ensemble.base_models == mock_base_models
        assert ensemble.physics_model == physics_model
        assert ensemble.physics_weight == 0.3
    
    @pytest.mark.skipif(not hasattr(__import__('torch'), 'torch'), reason="PyTorch not available")
    def test_physics_informed_ensemble_fit(self, sample_data, mock_base_models):
        """Test PhysicsInformedEnsemble fitting."""
        X, y = sample_data
        X_train, X_val = X.iloc[:80], X.iloc[80:]
        y_train, y_val = y.iloc[:80], y.iloc[80:]
        
        physics_model = PhysicsInformedNeuralNetwork(input_dim=4)
        ensemble = PhysicsInformedEnsemble(
            base_models=mock_base_models,
            physics_model=physics_model,
            physics_weight=0.3
        )
        
        ensemble.fit(X_train, y_train, X_val, y_val)
        
        # Verify base models were trained
        for model_name, trainer in mock_base_models.items():
            assert trainer.train.call_count >= 1
        
        # Verify physics model was trained
        assert ensemble.trained_physics_model is not None
    
    @pytest.mark.skipif(not hasattr(__import__('torch'), 'torch'), reason="PyTorch not available")
    def test_physics_informed_ensemble_predict(self, sample_data, mock_base_models):
        """Test PhysicsInformedEnsemble prediction."""
        X, y = sample_data
        
        physics_model = PhysicsInformedNeuralNetwork(input_dim=4)
        ensemble = PhysicsInformedEnsemble(
            base_models=mock_base_models,
            physics_model=physics_model,
            physics_weight=0.3
        )
        
        # Mock the trained state
        ensemble.trained_base_models = mock_base_models
        ensemble.trained_physics_model = physics_model.state_dict()
        
        predictions = ensemble.predict(X.iloc[:3])
        
        assert len(predictions) == 3
        # Predictions should be weighted combination of base and physics models
        assert all(isinstance(pred, (int, float)) for pred in predictions)


class TestEnsembleModelTrainer:
    """Test EnsembleModelTrainer implementation."""
    
    @pytest.fixture
    def sample_data(self):
        """Create sample data for testing."""
        np.random.seed(42)
        X = pd.DataFrame({
            'feature1': np.random.randn(100),
            'feature2': np.random.randn(100),
            'feature3': np.random.randn(100)
        })
        y = pd.Series(2 * X['feature1'] + 3 * X['feature2'] + np.random.randn(100) * 0.1)
        return X, y
    
    @pytest.fixture
    def temp_models_dir(self):
        """Create temporary directory for models."""
        temp_dir = tempfile.mkdtemp()
        yield temp_dir
        shutil.rmtree(temp_dir)
    
    def test_ensemble_model_trainer_initialization(self, temp_models_dir):
        """Test EnsembleModelTrainer initialization."""
        trainer = EnsembleModelTrainer(models_dir=temp_models_dir)
        
        assert trainer.models_dir == Path(temp_models_dir)
        assert trainer.include_physics_informed is True
        assert trainer.trained_ensembles == {}
        assert trainer.training_results == {}
    
    @patch('src.tlr4_binding.ml_components.ensemble_models.SKLEARN_AVAILABLE', True)
    def test_ensemble_model_trainer_train_ensembles(self, sample_data, temp_models_dir):
        """Test EnsembleModelTrainer ensemble training."""
        X, y = sample_data
        X_train, X_val = X.iloc[:80], X.iloc[80:]
        y_train, y_val = y.iloc[:80], y.iloc[80:]
        
        trainer = EnsembleModelTrainer(models_dir=temp_models_dir, include_physics_informed=False)
        
        # Mock base models
        base_models = {}
        rf_trainer = Mock(spec=RandomForestTrainer)
        rf_model = Mock()
        rf_trainer.train.return_value = rf_model
        rf_trainer.predict.return_value = np.array([1.0] * 80)
        base_models['random_forest'] = rf_trainer
        
        svr_trainer = Mock(spec=SVRTrainer)
        svr_model = Mock()
        svr_trainer.train.return_value = svr_model
        svr_trainer.predict.return_value = np.array([1.1] * 80)
        base_models['svr'] = svr_trainer
        
        # Train ensembles
        results = trainer.train_ensembles(X_train, y_train, X_val, y_val, base_models)
        
        # Verify ensembles were trained
        assert len(results) > 0
        assert 'stacked' in results or 'weighted' in results
    
    def test_ensemble_model_trainer_evaluate_ensembles(self, sample_data, temp_models_dir):
        """Test EnsembleModelTrainer ensemble evaluation."""
        X, y = sample_data
        X_test = X.iloc[80:]
        y_test = y.iloc[80:]
        
        trainer = EnsembleModelTrainer(models_dir=temp_models_dir)
        
        # Mock trained ensembles
        mock_ensemble = Mock()
        mock_ensemble.predict.return_value = np.array([1.5, 2.5, 3.5])
        mock_ensemble.predict_with_uncertainty.return_value = [
            EnsemblePrediction(1.5, 0.1, {}, {}, (1.3, 1.7)),
            EnsemblePrediction(2.5, 0.2, {}, {}, (2.1, 2.9)),
            EnsemblePrediction(3.5, 0.15, {}, {}, (3.2, 3.8))
        ]
        trainer.trained_ensembles['test_ensemble'] = mock_ensemble
        
        # Evaluate ensembles
        results = trainer.evaluate_ensembles(X_test, y_test)
        
        assert 'test_ensemble' in results
        assert 'predictions' in results['test_ensemble']
        assert 'metrics' in results['test_ensemble']
        assert 'uncertainty_predictions' in results['test_ensemble']
    
    def test_ensemble_model_trainer_get_best_ensemble(self, temp_models_dir):
        """Test EnsembleModelTrainer best ensemble selection."""
        trainer = EnsembleModelTrainer(models_dir=temp_models_dir)
        
        # Mock training results
        trainer.training_results = {
            'ensemble1': {
                'train_metrics': {'r2': 0.8, 'rmse': 1.2}
            },
            'ensemble2': {
                'train_metrics': {'r2': 0.9, 'rmse': 1.0}
            }
        }
        
        trainer.trained_ensembles = {
            'ensemble1': Mock(),
            'ensemble2': Mock()
        }
        
        best_name, best_model = trainer.get_best_ensemble(metric='r2')
        
        assert best_name == 'ensemble2'
        assert best_model == trainer.trained_ensembles['ensemble2']
    
    def test_ensemble_model_trainer_get_ensemble_summary(self, temp_models_dir):
        """Test EnsembleModelTrainer ensemble summary."""
        trainer = EnsembleModelTrainer(models_dir=temp_models_dir)
        
        # Mock training results
        trainer.training_results = {
            'ensemble1': {
                'model_type': 'stacked_ensemble',
                'train_metrics': {'r2': 0.8, 'rmse': 1.2}
            },
            'ensemble2': {
                'model_type': 'weighted_ensemble',
                'train_metrics': {'r2': 0.9, 'rmse': 1.0}
            }
        }
        
        summary = trainer.get_ensemble_summary()
        
        assert isinstance(summary, pd.DataFrame)
        assert len(summary) == 2
        assert 'ensemble_name' in summary.columns
        assert 'model_type' in summary.columns
        assert 'r2' in summary.columns
        assert 'rmse' in summary.columns


class TestEnsembleIntegration:
    """Integration tests for ensemble models."""
    
    @pytest.fixture
    def sample_data(self):
        """Create sample data for testing."""
        np.random.seed(42)
        X = pd.DataFrame({
            'molecular_weight': np.random.uniform(100, 500, 200),
            'logp': np.random.uniform(-2, 5, 200),
            'tpsa': np.random.uniform(20, 150, 200),
            'rotatable_bonds': np.random.randint(0, 10, 200),
            'hbd': np.random.randint(0, 5, 200),
            'hba': np.random.randint(0, 10, 200)
        })
        y = pd.Series(-2 * X['molecular_weight'] / 1000 + 
                     0.5 * X['logp'] + 
                     np.random.randn(200) * 0.1)
        return X, y
    
    @pytest.mark.skipif(not hasattr(__import__('sklearn'), 'sklearn'), reason="scikit-learn not available")
    def test_end_to_end_ensemble_training(self, sample_data, tempfile):
        """Test end-to-end ensemble training and prediction."""
        X, y = sample_data
        X_train, X_test = X.iloc[:150], X.iloc[150:]
        y_train, y_test = y.iloc[:150], y.iloc[150:]
        
        # Create base models
        base_models = {}
        if hasattr(__import__('sklearn'), 'sklearn'):
            from sklearn.ensemble import RandomForestRegressor
            from sklearn.svm import SVR
            
            # Mock trainers that actually work
            class MockTrainer:
                def __init__(self, model_class):
                    self.model_class = model_class
                    self.model = None
                
                def train(self, X, y, X_val=None, y_val=None):
                    self.model = self.model_class()
                    self.model.fit(X, y)
                    return self.model
                
                def predict(self, model, X):
                    return model.predict(X)
            
            base_models['random_forest'] = MockTrainer(RandomForestRegressor)
            base_models['svr'] = MockTrainer(SVR)
        
        # Create ensemble trainer
        with tempfile.TemporaryDirectory() as temp_dir:
            trainer = EnsembleModelTrainer(models_dir=temp_dir, include_physics_informed=False)
            
            # Train ensembles
            results = trainer.train_ensembles(X_train, y_train, base_models=base_models)
            
            # Verify training completed
            assert len(results) > 0
            
            # Evaluate ensembles
            eval_results = trainer.evaluate_ensembles(X_test, y_test)
            
            # Verify evaluation completed
            assert len(eval_results) > 0
            
            # Test prediction with uncertainty
            for ensemble_name, ensemble in trainer.trained_ensembles.items():
                predictions = ensemble.predict(X_test.iloc[:5])
                uncertainty_predictions = ensemble.predict_with_uncertainty(X_test.iloc[:5])
                
                assert len(predictions) == 5
                assert len(uncertainty_predictions) == 5
                assert all(isinstance(pred, EnsemblePrediction) for pred in uncertainty_predictions)


if __name__ == "__main__":
    pytest.main([__file__, "-v"])
```
