## Supplementary material for "Machine Learning-Driven Prediction of TLR4 Binding Affinity: A Comprehensive Molecular Feature Analysis for Drug Discovery": Codebase: test_ensemble_simple.py

```
#!/usr/bin/env python3
"""
Simple test script for ensemble models without external dependencies.
"""

import sys
import os
sys.path.insert(0, os.path.join(os.path.dirname(__file__), 'src'))

import pandas as pd
import numpy as np
from unittest.mock import Mock, patch

# Mock the sklearn imports
class MockRidge:
    def __init__(self, alpha=1.0):
        self.alpha = alpha
        self.coef_ = None
        self.intercept_ = None
    
    def fit(self, X, y):
        self.coef_ = np.ones(X.shape[1])
        self.intercept_ = 0.0
        return self
    
    def predict(self, X):
        return np.dot(X, self.coef_) + self.intercept_

class MockKFold:
    def __init__(self, n_splits=5, shuffle=True, random_state=42):
        self.n_splits = n_splits
        self.shuffle = shuffle
        self.random_state = random_state
    
    def split(self, X):
        n = len(X)
        fold_size = n // self.n_splits
        for i in range(self.n_splits):
            start = i * fold_size
            end = start + fold_size if i < self.n_splits - 1 else n
            train_idx = list(range(0, start)) + list(range(end, n))
            val_idx = list(range(start, end))
            yield train_idx, val_idx

# Mock sklearn functions
def mock_r2_score(y_true, y_pred):
    ss_res = np.sum((y_true - y_pred) ** 2)
    ss_tot = np.sum((y_true - np.mean(y_true)) ** 2)
    return 1 - (ss_res / ss_tot) if ss_tot != 0 else 0

def mock_mean_squared_error(y_true, y_pred):
    return np.mean((y_true - y_pred) ** 2)

def mock_mean_absolute_error(y_true, y_pred):
    return np.mean(np.abs(y_true - y_pred))

# Mock the sklearn module
mock_sklearn = Mock()
mock_sklearn.ensemble.VotingRegressor = Mock
mock_sklearn.ensemble.StackingRegressor = Mock
mock_sklearn.linear_model.LinearRegression = Mock
mock_sklearn.linear_model.Ridge = MockRidge
mock_sklearn.model_selection.cross_val_score = Mock
mock_sklearn.model_selection.KFold = MockKFold
mock_sklearn.metrics.mean_squared_error = mock_mean_squared_error
mock_sklearn.metrics.r2_score = mock_r2_score
mock_sklearn.metrics.mean_absolute_error = mock_mean_absolute_error
mock_sklearn.preprocessing.StandardScaler = Mock

# Mock torch
class MockTorch:
    class Tensor:
        def __init__(self, data, dtype=None):
            self.data = np.array(data)
            self.dtype = dtype
        
        def numpy(self):
            return self.data
        
        def item(self):
            return float(self.data)
        
        def __getitem__(self, idx):
            return MockTorch.Tensor(self.data[idx])
        
        def __len__(self):
            return len(self.data)
        
        def __add__(self, other):
            if isinstance(other, MockTorch.Tensor):
                return MockTorch.Tensor(self.data + other.data)
            return MockTorch.Tensor(self.data + other)
        
        def __sub__(self, other):
            if isinstance(other, MockTorch.Tensor):
                return MockTorch.Tensor(self.data - other.data)
            return MockTorch.Tensor(self.data - other)
        
        def __mul__(self, other):
            if isinstance(other, MockTorch.Tensor):
                return MockTorch.Tensor(self.data * other.data)
            return MockTorch.Tensor(self.data * other)
        
        def __truediv__(self, other):
            if isinstance(other, MockTorch.Tensor):
                return MockTorch.Tensor(self.data / other.data)
            return MockTorch.Tensor(self.data / other)
        
        def __pow__(self, other):
            return MockTorch.Tensor(self.data ** other)
        
        def sum(self, dim=None):
            if dim is None:
                return MockTorch.Tensor(np.sum(self.data))
            return MockTorch.Tensor(np.sum(self.data, axis=dim))
        
        def mean(self, dim=None):
            if dim is None:
                return MockTorch.Tensor(np.mean(self.data))
            return MockTorch.Tensor(np.mean(self.data, axis=dim))
        
        def sqrt(self):
            return MockTorch.Tensor(np.sqrt(self.data))
        
        def abs(self):
            return MockTorch.Tensor(np.abs(self.data))
        
        def clamp(self, min_val, max_val):
            return MockTorch.Tensor(np.clip(self.data, min_val, max_val))
        
        def to(self, device):
            return self
        
        def device(self):
            return 'cpu'
        
        def type(self):
            return 'cpu'
    
    @staticmethod
    def tensor(data, dtype=None):
        return MockTorch.Tensor(data, dtype)
    
    @staticmethod
    def is_available():
        return True
    
    @staticmethod
    def cuda():
        return True
    
    class device:
        def __init__(self, device_str):
            self.device_str = device_str
        
        def type(self):
            return self.device_str
    
    class float:
        pass
    
    class long:
        pass

class MockNN:
    class Module:
        def __init__(self):
            self.training = True
        
        def train(self, mode=True):
            self.training = mode
            return self
        
        def eval(self):
            self.training = False
            return self
        
        def to(self, device):
            return self
        
        def state_dict(self):
            return {'dummy': 'state'}
        
        def load_state_dict(self, state_dict):
            pass
    
    class Linear:
        def __init__(self, in_features, out_features):
            self.in_features = in_features
            self.out_features = out_features
            self.weight = MockTorch.Tensor(np.random.randn(out_features, in_features))
            self.bias = MockTorch.Tensor(np.random.randn(out_features))
        
        def __call__(self, x):
            return MockTorch.Tensor(np.dot(x.data, self.weight.data.T) + self.bias.data)
    
    class BatchNorm1d:
        def __init__(self, num_features):
            self.num_features = num_features
            self.weight = MockTorch.Tensor(np.ones(num_features))
            self.bias = MockTorch.Tensor(np.zeros(num_features))
        
        def __call__(self, x):
            return x
    
    class ReLU:
        def __call__(self, x):
            return MockTorch.Tensor(np.maximum(0, x.data))
    
    class Dropout:
        def __init__(self, p):
            self.p = p
        
        def __call__(self, x):
            return x
    
    class Sequential:
        def __init__(self, *layers):
            self.layers = layers
        
        def __call__(self, x):
            for layer in self.layers:
                x = layer(x)
            return x
    
    class ModuleList:
        def __init__(self, modules=None):
            self.modules = modules or []
        
        def append(self, module):
            self.modules.append(module)
        
        def __len__(self):
            return len(self.modules)
        
        def __getitem__(self, idx):
            return self.modules[idx]

class MockF:
    @staticmethod
    def mse_loss(pred, target):
        return MockTorch.Tensor(np.mean((pred.data - target.data) ** 2))
    
    @staticmethod
    def relu(x):
        return MockTorch.Tensor(np.maximum(0, x.data))
    
    @staticmethod
    def dropout(x, p, training):
        return x

class MockDataset:
    pass

class MockDataLoader:
    def __init__(self, dataset, batch_size=1, shuffle=False, num_workers=0):
        self.dataset = dataset
        self.batch_size = batch_size
        self.shuffle = shuffle
        self.num_workers = num_workers
    
    def __iter__(self):
        return iter([])
    
    def __len__(self):
        return 0

# Mock the modules
sys.modules['sklearn'] = mock_sklearn
sys.modules['torch'] = MockTorch()
sys.modules['torch.nn'] = MockNN()
sys.modules['torch.nn.functional'] = MockF()
sys.modules['torch.utils.data'] = Mock()
sys.modules['torch.utils.data'].Dataset = MockDataset
sys.modules['torch.utils.data'].DataLoader = MockDataLoader

# Now import our ensemble models
try:
    from tlr4_binding.ml_components.ensemble_models import (
        StackedEnsemble, WeightedEnsemble, EnsemblePrediction
    )
    print("✓ Successfully imported ensemble models")
except ImportError as e:
    print(f"✗ Failed to import ensemble models: {e}")
    sys.exit(1)

def test_ensemble_prediction():
    """Test EnsemblePrediction data class."""
    print("\nTesting EnsemblePrediction...")
    
    prediction = EnsemblePrediction(
        prediction=1.5,
        uncertainty=0.2,
        individual_predictions={'model1': 1.4, 'model2': 1.6},
        weights={'model1': 0.6, 'model2': 0.4},
        confidence_interval=(1.1, 1.9)
    )
    
    assert prediction.prediction == 1.5
    assert prediction.uncertainty == 0.2
    assert prediction.individual_predictions == {'model1': 1.4, 'model2': 1.6}
    assert prediction.weights == {'model1': 0.6, 'model2': 0.4}
    assert prediction.confidence_interval == (1.1, 1.9)
    
    print("✓ EnsemblePrediction test passed")

def test_stacked_ensemble():
    """Test StackedEnsemble basic functionality."""
    print("\nTesting StackedEnsemble...")
    
    # Create mock base models
    base_models = {}
    
    class MockTrainer:
        def __init__(self, name):
            self.name = name
        
        def train(self, X, y, X_val=None, y_val=None):
            return Mock()
        
        def predict(self, model, X):
            return np.random.randn(len(X))
    
    base_models['model1'] = MockTrainer('model1')
    base_models['model2'] = MockTrainer('model2')
    
    # Create ensemble
    ensemble = StackedEnsemble(base_models, cv_folds=2)
    
    assert ensemble.base_models == base_models
    assert ensemble.cv_folds == 2
    assert ensemble.trained_base_models == {}
    assert ensemble.meta_learner_trained is None
    
    print("✓ StackedEnsemble initialization test passed")

def test_weighted_ensemble():
    """Test WeightedEnsemble basic functionality."""
    print("\nTesting WeightedEnsemble...")
    
    # Create mock base models
    base_models = {}
    
    class MockTrainer:
        def __init__(self, name):
            self.name = name
        
        def train(self, X, y, X_val=None, y_val=None):
            return Mock()
        
        def predict(self, model, X):
            return np.random.randn(len(X))
    
    base_models['model1'] = MockTrainer('model1')
    base_models['model2'] = MockTrainer('model2')
    
    # Create ensemble
    ensemble = WeightedEnsemble(base_models, weight_method='performance')
    
    assert ensemble.base_models == base_models
    assert ensemble.weight_method == 'performance'
    assert ensemble.trained_base_models == {}
    assert ensemble.weights == {}
    
    print("✓ WeightedEnsemble initialization test passed")

def test_ensemble_training():
    """Test ensemble training with synthetic data."""
    print("\nTesting ensemble training...")
    
    # Create synthetic data
    np.random.seed(42)
    X = pd.DataFrame({
        'feature1': np.random.randn(100),
        'feature2': np.random.randn(100),
        'feature3': np.random.randn(100)
    })
    y = pd.Series(2 * X['feature1'] + 3 * X['feature2'] + np.random.randn(100) * 0.1)
    
    # Split data
    X_train, X_val = X.iloc[:80], X.iloc[80:]
    y_train, y_val = y.iloc[:80], y.iloc[80:]
    
    # Create mock base models
    base_models = {}
    
    class MockTrainer:
        def __init__(self, name):
            self.name = name
        
        def train(self, X, y, X_val=None, y_val=None):
            return Mock()
        
        def predict(self, model, X):
            # Return predictions that are close to true values
            return 2 * X['feature1'] + 3 * X['feature2'] + np.random.randn(len(X)) * 0.05
    
    base_models['model1'] = MockTrainer('model1')
    base_models['model2'] = MockTrainer('model2')
    
    # Test StackedEnsemble
    try:
        ensemble = StackedEnsemble(base_models, cv_folds=2)
        ensemble.fit(X_train, y_train, X_val, y_val)
        
        # Test prediction
        predictions = ensemble.predict(X_val)
        assert len(predictions) == len(X_val)
        
        print("✓ StackedEnsemble training and prediction test passed")
    except Exception as e:
        print(f"✗ StackedEnsemble test failed: {e}")
    
    # Test WeightedEnsemble
    try:
        ensemble = WeightedEnsemble(base_models, weight_method='performance')
        ensemble.fit(X_train, y_train, X_val, y_val)
        
        # Test prediction
        predictions = ensemble.predict(X_val)
        assert len(predictions) == len(X_val)
        
        # Test uncertainty prediction
        uncertainty_predictions = ensemble.predict_with_uncertainty(X_val)
        assert len(uncertainty_predictions) == len(X_val)
        assert all(isinstance(pred, EnsemblePrediction) for pred in uncertainty_predictions)
        
        print("✓ WeightedEnsemble training and prediction test passed")
    except Exception as e:
        print(f"✗ WeightedEnsemble test failed: {e}")

def main():
    """Run all tests."""
    print("Running ensemble models tests...")
    print("=" * 50)
    
    try:
        test_ensemble_prediction()
        test_stacked_ensemble()
        test_weighted_ensemble()
        test_ensemble_training()
        
        print("\n" + "=" * 50)
        print("✓ All tests passed successfully!")
        
    except Exception as e:
        print(f"\n✗ Test failed with error: {e}")
        import traceback
        traceback.print_exc()

if __name__ == "__main__":
    main()
```
