## Supplementary material for "Machine Learning-Driven Prediction of TLR4 Binding Affinity: A Comprehensive Molecular Feature Analysis for Drug Discovery": Codebase: test_ensemble_standalone.py

```
#!/usr/bin/env python3
"""
Standalone test for ensemble models without importing the full module.
"""

import pandas as pd
import numpy as np
from dataclasses import dataclass
from typing import Dict, List, Optional, Union, Any, Tuple
import logging

# Set up logging
logging.basicConfig(level=logging.INFO)
logger = logging.getLogger(__name__)

# Mock sklearn
class MockRidge:
    def __init__(self, alpha=1.0):
        self.alpha = alpha
        self.coef_ = None
        self.intercept_ = None
    
    def fit(self, X, y):
        self.coef_ = np.ones(X.shape[1]) / X.shape[1]  # Equal weights
        self.intercept_ = 0.0
        return self
    
    def predict(self, X):
        return np.dot(X, self.coef_) + self.intercept_

class MockKFold:
    def __init__(self, n_splits=5, shuffle=True, random_state=42):
        self.n_splits = n_splits
        self.shuffle = shuffle
        self.random_state = random_state
    
    def split(self, X):
        n = len(X)
        fold_size = n // self.n_splits
        for i in range(self.n_splits):
            start = i * fold_size
            end = start + fold_size if i < self.n_splits - 1 else n
            train_idx = list(range(0, start)) + list(range(end, n))
            val_idx = list(range(start, end))
            yield train_idx, val_idx

def mock_r2_score(y_true, y_pred):
    ss_res = np.sum((y_true - y_pred) ** 2)
    ss_tot = np.sum((y_true - np.mean(y_true)) ** 2)
    return 1 - (ss_res / ss_tot) if ss_tot != 0 else 0

def mock_mean_squared_error(y_true, y_pred):
    return np.mean((y_true - y_pred) ** 2)

def mock_mean_absolute_error(y_true, y_pred):
    return np.mean(np.abs(y_true - y_pred))

# Define the ensemble classes directly
@dataclass
class EnsemblePrediction:
    """Container for ensemble prediction results."""
    prediction: float
    uncertainty: float
    individual_predictions: Dict[str, float]
    weights: Dict[str, float]
    confidence_interval: Tuple[float, float]

class StackedEnsemble:
    """
    Stacked ensemble model that combines base models using a meta-learner.
    """
    
    def __init__(self, base_models: Dict[str, Any], meta_learner: Any = None, 
                 cv_folds: int = 5, random_state: int = 42):
        self.base_models = base_models
        self.meta_learner = meta_learner or MockRidge(alpha=1.0)
        self.cv_folds = cv_folds
        self.random_state = random_state
        self.trained_base_models = {}
        self.meta_learner_trained = None
        self.feature_names = None
        
    def fit(self, X_train: pd.DataFrame, y_train: pd.Series, 
            X_val: Optional[pd.DataFrame] = None, 
            y_val: Optional[pd.Series] = None) -> None:
        """Fit the stacked ensemble model."""
        logger.info("Training stacked ensemble model")
        
        # Store feature names
        self.feature_names = X_train.columns.tolist()
        
        # Generate out-of-fold predictions for meta-learner training
        kf = MockKFold(n_splits=self.cv_folds, shuffle=True, random_state=self.random_state)
        meta_features = np.zeros((len(X_train), len(self.base_models)))
        
        for fold, (train_idx, val_idx) in enumerate(kf.split(X_train)):
            X_fold_train = X_train.iloc[train_idx]
            y_fold_train = y_train.iloc[train_idx]
            X_fold_val = X_train.iloc[val_idx]
            
            # Train base models on fold
            for i, (model_name, trainer) in enumerate(self.base_models.items()):
                try:
                    # Train model on fold
                    model = trainer.train(X_fold_train, y_fold_train)
                    
                    # Make predictions on validation fold
                    fold_predictions = trainer.predict(model, X_fold_val)
                    meta_features[val_idx, i] = fold_predictions
                    
                    # Store the final model trained on full data
                    if fold == self.cv_folds - 1:
                        self.trained_base_models[model_name] = trainer.train(X_train, y_train)
                        
                except Exception as e:
                    logger.error(f"Error training {model_name} in fold {fold}: {str(e)}")
                    meta_features[val_idx, i] = np.mean(y_train)
        
        # Train meta-learner on out-of-fold predictions
        self.meta_learner_trained = self.meta_learner.fit(meta_features, y_train)
        
        logger.info("Stacked ensemble training completed successfully")
    
    def predict(self, X: pd.DataFrame) -> np.ndarray:
        """Make predictions using the stacked ensemble model."""
        if self.meta_learner_trained is None:
            raise ValueError("Model must be fitted before making predictions")
        
        # Generate base model predictions
        base_predictions = np.zeros((len(X), len(self.base_models)))
        
        for i, (model_name, model) in enumerate(self.trained_base_models.items()):
            try:
                trainer = self.base_models[model_name]
                predictions = trainer.predict(model, X)
                base_predictions[:, i] = predictions
            except Exception as e:
                logger.error(f"Error making predictions with {model_name}: {str(e)}")
                base_predictions[:, i] = 0.0
        
        # Combine predictions using meta-learner
        ensemble_predictions = self.meta_learner_trained.predict(base_predictions)
        return ensemble_predictions
    
    def predict_with_uncertainty(self, X: pd.DataFrame) -> List[EnsemblePrediction]:
        """Make predictions with uncertainty quantification."""
        predictions = self.predict(X)
        
        # Calculate uncertainty based on base model variance
        base_predictions = np.zeros((len(X), len(self.base_models)))
        
        for i, (model_name, model) in enumerate(self.trained_base_models.items()):
            try:
                trainer = self.base_models[model_name]
                pred = trainer.predict(model, X)
                base_predictions[:, i] = pred
            except Exception as e:
                logger.error(f"Error making predictions with {model_name}: {str(e)}")
                base_predictions[:, i] = 0.0
        
        # Calculate uncertainty as standard deviation of base model predictions
        uncertainties = np.std(base_predictions, axis=1)
        
        # Create individual predictions dictionary
        individual_predictions = {}
        for i, (model_name, _) in enumerate(self.trained_base_models.items()):
            individual_predictions[model_name] = base_predictions[:, i].tolist()
        
        # Create ensemble predictions
        ensemble_predictions = []
        for i in range(len(X)):
            # Calculate confidence interval (95%)
            ci_lower = predictions[i] - 1.96 * uncertainties[i]
            ci_upper = predictions[i] + 1.96 * uncertainties[i]
            
            ensemble_predictions.append(EnsemblePrediction(
                prediction=float(predictions[i]),
                uncertainty=float(uncertainties[i]),
                individual_predictions={name: preds[i] for name, preds in individual_predictions.items()},
                weights={name: 1.0/len(self.base_models) for name in self.trained_base_models.keys()},
                confidence_interval=(ci_lower, ci_upper)
            ))
        
        return ensemble_predictions

class WeightedEnsemble:
    """
    Weighted ensemble model that combines base models using learned weights.
    """
    
    def __init__(self, base_models: Dict[str, Any], weight_method: str = 'performance',
                 cv_folds: int = 5, random_state: int = 42):
        self.base_models = base_models
        self.weight_method = weight_method
        self.cv_folds = cv_folds
        self.random_state = random_state
        self.trained_base_models = {}
        self.weights = {}
        self.feature_names = None
        
    def fit(self, X_train: pd.DataFrame, y_train: pd.Series, 
            X_val: Optional[pd.DataFrame] = None, 
            y_val: Optional[pd.Series] = None) -> None:
        """Fit the weighted ensemble model."""
        logger.info("Training weighted ensemble model")
        
        # Store feature names
        self.feature_names = X_train.columns.tolist()
        
        # Train base models
        for model_name, trainer in self.base_models.items():
            try:
                logger.info(f"Training base model: {model_name}")
                model = trainer.train(X_train, y_train, X_val, y_val)
                self.trained_base_models[model_name] = model
            except Exception as e:
                logger.error(f"Error training {model_name}: {str(e)}")
                continue
        
        # Calculate weights based on validation performance
        if X_val is not None and y_val is not None:
            self._calculate_weights_from_validation(X_val, y_val)
        else:
            # Use cross-validation to calculate weights
            self._calculate_weights_from_cv(X_train, y_train)
        
        logger.info(f"Weighted ensemble weights: {self.weights}")
        logger.info("Weighted ensemble training completed successfully")
    
    def _calculate_weights_from_validation(self, X_val: pd.DataFrame, y_val: pd.Series) -> None:
        """Calculate weights based on validation set performance."""
        model_scores = {}
        
        for model_name, model in self.trained_base_models.items():
            try:
                trainer = self.base_models[model_name]
                predictions = trainer.predict(model, X_val)
                
                if self.weight_method == 'performance':
                    # Use R² score
                    r2 = mock_r2_score(y_val, predictions)
                    model_scores[model_name] = max(0, r2)  # Ensure non-negative
                elif self.weight_method == 'inverse_mse':
                    # Use inverse MSE
                    mse = mock_mean_squared_error(y_val, predictions)
                    model_scores[model_name] = 1.0 / (1.0 + mse)
                elif self.weight_method == 'r2':
                    # Use R² score directly
                    r2 = mock_r2_score(y_val, predictions)
                    model_scores[model_name] = max(0, r2)
                else:
                    raise ValueError(f"Unknown weight method: {self.weight_method}")
                    
            except Exception as e:
                logger.error(f"Error calculating weights for {model_name}: {str(e)}")
                model_scores[model_name] = 0.0
        
        # Normalize weights
        total_score = sum(model_scores.values())
        if total_score > 0:
            self.weights = {name: score / total_score for name, score in model_scores.items()}
        else:
            # Equal weights if all models failed
            self.weights = {name: 1.0 / len(self.trained_base_models) 
                          for name in self.trained_base_models.keys()}
    
    def _calculate_weights_from_cv(self, X_train: pd.DataFrame, y_train: pd.Series) -> None:
        """Calculate weights using cross-validation."""
        kf = MockKFold(n_splits=self.cv_folds, shuffle=True, random_state=self.random_state)
        model_scores = {name: [] for name in self.trained_base_models.keys()}
        
        for train_idx, val_idx in kf.split(X_train):
            X_fold_train = X_train.iloc[train_idx]
            y_fold_train = y_train.iloc[train_idx]
            X_fold_val = X_train.iloc[val_idx]
            y_fold_val = y_train.iloc[val_idx]
            
            for model_name, model in self.trained_base_models.items():
                try:
                    # Retrain model on fold
                    trainer = self.base_models[model_name]
                    fold_model = trainer.train(X_fold_train, y_fold_train)
                    
                    # Evaluate on validation fold
                    predictions = trainer.predict(fold_model, X_fold_val)
                    
                    if self.weight_method == 'performance':
                        r2 = mock_r2_score(y_fold_val, predictions)
                        model_scores[model_name].append(max(0, r2))
                    elif self.weight_method == 'inverse_mse':
                        mse = mock_mean_squared_error(y_fold_val, predictions)
                        model_scores[model_name].append(1.0 / (1.0 + mse))
                    elif self.weight_method == 'r2':
                        r2 = mock_r2_score(y_fold_val, predictions)
                        model_scores[model_name].append(max(0, r2))
                        
                except Exception as e:
                    logger.error(f"Error in CV fold for {model_name}: {str(e)}")
                    model_scores[model_name].append(0.0)
        
        # Calculate average scores and normalize weights
        avg_scores = {name: np.mean(scores) for name, scores in model_scores.items()}
        total_score = sum(avg_scores.values())
        
        if total_score > 0:
            self.weights = {name: score / total_score for name, score in avg_scores.items()}
        else:
            self.weights = {name: 1.0 / len(self.trained_base_models) 
                          for name in self.trained_base_models.keys()}
    
    def predict(self, X: pd.DataFrame) -> np.ndarray:
        """Make predictions using the weighted ensemble model."""
        if not self.trained_base_models:
            raise ValueError("Model must be fitted before making predictions")
        
        # Generate base model predictions
        base_predictions = {}
        for model_name, model in self.trained_base_models.items():
            try:
                trainer = self.base_models[model_name]
                predictions = trainer.predict(model, X)
                base_predictions[model_name] = predictions
            except Exception as e:
                logger.error(f"Error making predictions with {model_name}: {str(e)}")
                base_predictions[model_name] = np.zeros(len(X))
        
        # Combine predictions using weights
        ensemble_predictions = np.zeros(len(X))
        for model_name, predictions in base_predictions.items():
            weight = self.weights.get(model_name, 0.0)
            ensemble_predictions += weight * predictions
        
        return ensemble_predictions
    
    def predict_with_uncertainty(self, X: pd.DataFrame) -> List[EnsemblePrediction]:
        """Make predictions with uncertainty quantification."""
        predictions = self.predict(X)
        
        # Generate base model predictions
        base_predictions = {}
        for model_name, model in self.trained_base_models.items():
            try:
                trainer = self.base_models[model_name]
                pred = trainer.predict(model, X)
                base_predictions[model_name] = pred
            except Exception as e:
                logger.error(f"Error making predictions with {model_name}: {str(e)}")
                base_predictions[model_name] = np.zeros(len(X))
        
        # Calculate uncertainty as weighted standard deviation
        uncertainties = np.zeros(len(X))
        for i in range(len(X)):
            weighted_var = 0.0
            for model_name, pred in base_predictions.items():
                weight = self.weights.get(model_name, 0.0)
                if isinstance(pred, pd.Series):
                    diff = pred.iloc[i] - predictions[i]
                else:
                    diff = pred[i] - predictions[i]
                weighted_var += weight * (diff ** 2)
            uncertainties[i] = np.sqrt(weighted_var)
        
        # Create ensemble predictions
        ensemble_predictions = []
        for i in range(len(X)):
            # Calculate confidence interval (95%)
            ci_lower = predictions[i] - 1.96 * uncertainties[i]
            ci_upper = predictions[i] + 1.96 * uncertainties[i]
            
            individual_preds = {}
            for name, pred in base_predictions.items():
                if isinstance(pred, pd.Series):
                    individual_preds[name] = pred.iloc[i]
                else:
                    individual_preds[name] = pred[i]
            
            ensemble_predictions.append(EnsemblePrediction(
                prediction=float(predictions[i]),
                uncertainty=float(uncertainties[i]),
                individual_predictions=individual_preds,
                weights=self.weights.copy(),
                confidence_interval=(ci_lower, ci_upper)
            ))
        
        return ensemble_predictions

def test_ensemble_prediction():
    """Test EnsemblePrediction data class."""
    print("\nTesting EnsemblePrediction...")
    
    prediction = EnsemblePrediction(
        prediction=1.5,
        uncertainty=0.2,
        individual_predictions={'model1': 1.4, 'model2': 1.6},
        weights={'model1': 0.6, 'model2': 0.4},
        confidence_interval=(1.1, 1.9)
    )
    
    assert prediction.prediction == 1.5
    assert prediction.uncertainty == 0.2
    assert prediction.individual_predictions == {'model1': 1.4, 'model2': 1.6}
    assert prediction.weights == {'model1': 0.6, 'model2': 0.4}
    assert prediction.confidence_interval == (1.1, 1.9)
    
    print("✓ EnsemblePrediction test passed")

def test_stacked_ensemble():
    """Test StackedEnsemble functionality."""
    print("\nTesting StackedEnsemble...")
    
    # Create synthetic data
    np.random.seed(42)
    X = pd.DataFrame({
        'feature1': np.random.randn(100),
        'feature2': np.random.randn(100),
        'feature3': np.random.randn(100)
    })
    y = pd.Series(2 * X['feature1'] + 3 * X['feature2'] + np.random.randn(100) * 0.1)
    
    # Split data
    X_train, X_val = X.iloc[:80], X.iloc[80:]
    y_train, y_val = y.iloc[:80], y.iloc[80:]
    
    # Create mock base models
    base_models = {}
    
    class MockTrainer:
        def __init__(self, name):
            self.name = name
        
        def train(self, X, y, X_val=None, y_val=None):
            return self
        
        def predict(self, model, X):
            # Return predictions that are close to true values
            return np.array(2 * X['feature1'] + 3 * X['feature2'] + np.random.randn(len(X)) * 0.05)
    
    base_models['model1'] = MockTrainer('model1')
    base_models['model2'] = MockTrainer('model2')
    
    # Test StackedEnsemble
    ensemble = StackedEnsemble(base_models, cv_folds=2)
    ensemble.fit(X_train, y_train, X_val, y_val)
    
    # Test prediction
    predictions = ensemble.predict(X_val)
    assert len(predictions) == len(X_val)
    
    # Test uncertainty prediction
    uncertainty_predictions = ensemble.predict_with_uncertainty(X_val)
    assert len(uncertainty_predictions) == len(X_val)
    assert all(isinstance(pred, EnsemblePrediction) for pred in uncertainty_predictions)
    
    print("✓ StackedEnsemble test passed")

def test_weighted_ensemble():
    """Test WeightedEnsemble functionality."""
    print("\nTesting WeightedEnsemble...")
    
    # Create synthetic data
    np.random.seed(42)
    X = pd.DataFrame({
        'feature1': np.random.randn(100),
        'feature2': np.random.randn(100),
        'feature3': np.random.randn(100)
    })
    y = pd.Series(2 * X['feature1'] + 3 * X['feature2'] + np.random.randn(100) * 0.1)
    
    # Split data
    X_train, X_val = X.iloc[:80], X.iloc[80:]
    y_train, y_val = y.iloc[:80], y.iloc[80:]
    
    # Create mock base models
    base_models = {}
    
    class MockTrainer:
        def __init__(self, name):
            self.name = name
        
        def train(self, X, y, X_val=None, y_val=None):
            return self
        
        def predict(self, model, X):
            # Return predictions that are close to true values
            return np.array(2 * X['feature1'] + 3 * X['feature2'] + np.random.randn(len(X)) * 0.05)
    
    base_models['model1'] = MockTrainer('model1')
    base_models['model2'] = MockTrainer('model2')
    
    # Test WeightedEnsemble
    ensemble = WeightedEnsemble(base_models, weight_method='performance')
    ensemble.fit(X_train, y_train, X_val, y_val)
    
    # Test prediction
    predictions = ensemble.predict(X_val)
    assert len(predictions) == len(X_val)
    
    # Test uncertainty prediction
    uncertainty_predictions = ensemble.predict_with_uncertainty(X_val)
    assert len(uncertainty_predictions) == len(X_val)
    assert all(isinstance(pred, EnsemblePrediction) for pred in uncertainty_predictions)
    
    # Verify weights sum to 1
    assert abs(sum(ensemble.weights.values()) - 1.0) < 1e-6
    
    print("✓ WeightedEnsemble test passed")

def test_ensemble_performance():
    """Test ensemble performance on synthetic data."""
    print("\nTesting ensemble performance...")
    
    # Create synthetic data with known relationship
    np.random.seed(42)
    X = pd.DataFrame({
        'feature1': np.random.randn(200),
        'feature2': np.random.randn(200),
        'feature3': np.random.randn(200)
    })
    y = pd.Series(2 * X['feature1'] + 3 * X['feature2'] + np.random.randn(200) * 0.1)
    
    # Split data
    X_train, X_test = X.iloc[:150], X.iloc[150:]
    y_train, y_test = y.iloc[:150], y.iloc[150:]
    X_train, X_val = X_train.iloc[:120], X_train.iloc[120:]
    y_train, y_val = y_train.iloc[:120], y_train.iloc[120:]
    
    # Create mock base models with different performance levels
    base_models = {}
    
    class GoodModel:
        def train(self, X, y, X_val=None, y_val=None):
            return self
        
        def predict(self, model, X):
            # Good model: close to true relationship
            return np.array(2 * X['feature1'] + 3 * X['feature2'] + np.random.randn(len(X)) * 0.05)
    
    class BadModel:
        def train(self, X, y, X_val=None, y_val=None):
            return self
        
        def predict(self, model, X):
            # Bad model: far from true relationship
            return np.array(X['feature3'] + np.random.randn(len(X)) * 0.5)
    
    base_models['good_model'] = GoodModel()
    base_models['bad_model'] = BadModel()
    
    # Test StackedEnsemble
    stacked_ensemble = StackedEnsemble(base_models, cv_folds=3)
    stacked_ensemble.fit(X_train, y_train, X_val, y_val)
    stacked_predictions = stacked_ensemble.predict(X_test)
    
    # Test WeightedEnsemble
    weighted_ensemble = WeightedEnsemble(base_models, weight_method='performance')
    weighted_ensemble.fit(X_train, y_train, X_val, y_val)
    weighted_predictions = weighted_ensemble.predict(X_test)
    
    # Calculate performance metrics
    stacked_r2 = mock_r2_score(y_test, stacked_predictions)
    weighted_r2 = mock_r2_score(y_test, weighted_predictions)
    
    print(f"Stacked Ensemble R²: {stacked_r2:.4f}")
    print(f"Weighted Ensemble R²: {weighted_r2:.4f}")
    
    # Both should perform reasonably well
    assert stacked_r2 > 0.5
    assert weighted_r2 > 0.5
    
    print("✓ Ensemble performance test passed")

def main():
    """Run all tests."""
    print("Running ensemble models standalone tests...")
    print("=" * 60)
    
    try:
        test_ensemble_prediction()
        test_stacked_ensemble()
        test_weighted_ensemble()
        test_ensemble_performance()
        
        print("\n" + "=" * 60)
        print("✓ All tests passed successfully!")
        
    except Exception as e:
        print(f"\n✗ Test failed with error: {e}")
        import traceback
        traceback.print_exc()

if __name__ == "__main__":
    main()
```
