## Supplementary material for "Machine Learning-Driven Prediction of TLR4 Binding Affinity: A Comprehensive Molecular Feature Analysis for Drug Discovery": Codebase: test_error_handling_robustness.py

```
"""
Comprehensive tests for error handling and robustness features.

This module tests all error handling, recovery mechanisms, and robustness
features implemented in the TLR4 binding prediction pipeline.
"""

import unittest
import tempfile
import shutil
import pandas as pd
import numpy as np
from pathlib import Path
from unittest.mock import patch, MagicMock
import logging

# Import the modules to test
from src.tlr4_binding.utils.error_handling import (
    RobustnessManager, CheckpointManager, CircuitBreaker,
    robust_execution, safe_execution, graceful_degradation,
    PipelineError, DataQualityError, ModelTrainingError, FeatureExtractionError
)
from src.tlr4_binding.utils.data_quality import (
    DataQualityValidator, DataAnomalyDetector, validate_molecular_data
)
from src.tlr4_binding.molecular_analysis.extractor import MolecularFeatureExtractor

logger = logging.getLogger(__name__)


class TestErrorHandlingUtilities(unittest.TestCase):
    """Test error handling utilities and decorators."""
    
    def setUp(self):
        """Set up test fixtures."""
        self.temp_dir = Path(tempfile.mkdtemp())
        self.robustness_manager = RobustnessManager()
        self.checkpoint_manager = CheckpointManager(str(self.temp_dir / "checkpoints"))
        
    def tearDown(self):
        """Clean up test fixtures."""
        shutil.rmtree(self.temp_dir, ignore_errors=True)
    
    def test_robust_execution_decorator_success(self):
        """Test robust execution decorator with successful operation."""
        @robust_execution(max_retries=3, delay=0.1)
        def successful_function():
            return "success"
        
        result = successful_function()
        self.assertEqual(result, "success")
    
    def test_robust_execution_decorator_retry(self):
        """Test robust execution decorator with retry logic."""
        call_count = 0
        
        @robust_execution(max_retries=3, delay=0.01)
        def failing_then_success():
            nonlocal call_count
            call_count += 1
            if call_count < 3:
                raise ValueError("Temporary failure")
            return "success"
        
        result = failing_then_success()
        self.assertEqual(result, "success")
        self.assertEqual(call_count, 3)
    
    def test_robust_execution_decorator_max_retries(self):
        """Test robust execution decorator with max retries exceeded."""
        @robust_execution(max_retries=2, delay=0.01)
        def always_failing():
            raise ValueError("Always fails")
        
        with self.assertRaises(ValueError):
            always_failing()
    
    def test_safe_execution_context_manager(self):
        """Test safe execution context manager."""
        with safe_execution("test operation", default_return="fallback"):
            raise ValueError("Test error")
        
        # Should not raise exception due to safe execution
        self.assertTrue(True)  # If we get here, safe execution worked
    
    def test_graceful_degradation_decorator(self):
        """Test graceful degradation decorator."""
        @graceful_degradation(fallback_value="degraded")
        def failing_function():
            raise ValueError("Function fails")
        
        result = failing_function()
        self.assertEqual(result, "degraded")
    
    def test_graceful_degradation_with_fallback_func(self):
        """Test graceful degradation with fallback function."""
        def fallback_func():
            return "fallback_result"
        
        @graceful_degradation(fallback_func=fallback_func)
        def failing_function():
            raise ValueError("Function fails")
        
        result = failing_function()
        self.assertEqual(result, "fallback_result")
    
    def test_circuit_breaker_pattern(self):
        """Test circuit breaker pattern."""
        cb = CircuitBreaker(failure_threshold=2, timeout=1)
        
        def failing_function():
            raise ValueError("Always fails")
        
        # First two failures should pass through
        with self.assertRaises(ValueError):
            cb.call(failing_function)
        
        with self.assertRaises(ValueError):
            cb.call(failing_function)
        
        # Third call should be blocked (circuit open)
        with self.assertRaises(Exception) as context:
            cb.call(failing_function)
        
        self.assertIn("Circuit breaker is OPEN", str(context.exception))


class TestCheckpointManager(unittest.TestCase):
    """Test checkpoint manager functionality."""
    
    def setUp(self):
        """Set up test fixtures."""
        self.temp_dir = Path(tempfile.mkdtemp())
        self.checkpoint_manager = CheckpointManager(str(self.temp_dir / "checkpoints"))
        
    def tearDown(self):
        """Clean up test fixtures."""
        shutil.rmtree(self.temp_dir, ignore_errors=True)
    
    def test_save_and_load_checkpoint(self):
        """Test saving and loading checkpoints."""
        checkpoint_id = "test_checkpoint"
        test_data = {
            'model_data': {'weights': [1, 2, 3]},
            'metadata': {'epoch': 10, 'loss': 0.5}
        }
        
        # Save checkpoint
        success = self.checkpoint_manager.save_checkpoint(checkpoint_id, test_data)
        self.assertTrue(success)
        
        # Load checkpoint
        loaded_data = self.checkpoint_manager.load_checkpoint(checkpoint_id)
        self.assertIsNotNone(loaded_data)
        self.assertEqual(loaded_data['model_data'], test_data['model_data'])
        self.assertEqual(loaded_data['metadata'], test_data['metadata'])
    
    def test_load_nonexistent_checkpoint(self):
        """Test loading non-existent checkpoint."""
        loaded_data = self.checkpoint_manager.load_checkpoint("nonexistent")
        self.assertIsNone(loaded_data)
    
    def test_list_checkpoints(self):
        """Test listing checkpoints."""
        # Save multiple checkpoints
        for i in range(3):
            self.checkpoint_manager.save_checkpoint(f"checkpoint_{i}", {'data': i})
        
        checkpoints = self.checkpoint_manager.list_checkpoints()
        self.assertEqual(len(checkpoints), 3)
        self.assertIn("checkpoint_0", checkpoints)
        self.assertIn("checkpoint_1", checkpoints)
        self.assertIn("checkpoint_2", checkpoints)
    
    def test_delete_checkpoint(self):
        """Test deleting checkpoints."""
        checkpoint_id = "test_delete"
        test_data = {'test': 'data'}
        
        # Save and verify
        self.checkpoint_manager.save_checkpoint(checkpoint_id, test_data)
        self.assertIsNotNone(self.checkpoint_manager.load_checkpoint(checkpoint_id))
        
        # Delete and verify
        success = self.checkpoint_manager.delete_checkpoint(checkpoint_id)
        self.assertTrue(success)
        self.assertIsNone(self.checkpoint_manager.load_checkpoint(checkpoint_id))


class TestDataQualityValidator(unittest.TestCase):
    """Test data quality validation functionality."""
    
    def setUp(self):
        """Set up test fixtures."""
        self.validator = DataQualityValidator()
        
        # Create test data
        self.good_data = pd.DataFrame({
            'molecular_weight': [100, 150, 200, 180, 120],
            'logp': [1.0, 2.0, 1.5, 0.5, 2.5],
            'tpsa': [50, 60, 55, 45, 65],
            'compound_name': ['A', 'B', 'C', 'D', 'E']
        })
        
        self.bad_data = pd.DataFrame({
            'molecular_weight': [100, 150, -50, 180, 120],  # Negative value
            'logp': [1.0, 2.0, 1.5, None, 2.5],  # Missing value
            'tpsa': [50, 60, 55, 45, 65],
            'compound_name': ['A', 'B', None, 'D', 'E']  # Missing name
        })
    
    def test_validate_good_data(self):
        """Test validation of good quality data."""
        results = self.validator.validate_dataset(self.good_data, "good_data")
        
        self.assertTrue(results['validation_passed'])
        self.assertGreater(results['quality_score'], 0.8)
        self.assertEqual(len(results['issues']), 0)
    
    def test_validate_bad_data(self):
        """Test validation of poor quality data."""
        results = self.validator.validate_dataset(self.bad_data, "bad_data")
        
        self.assertFalse(results['validation_passed'])
        self.assertLess(results['quality_score'], 0.8)
        self.assertGreater(len(results['issues']), 0)
    
    def test_missing_value_validation(self):
        """Test missing value validation."""
        results = self.validator._validate_missing_values(self.bad_data)
        
        self.assertIn('missing_value_issues', results)
        self.assertIn('missing_value_warnings', results)
        self.assertIn('missing_value_stats', results)
        
        # Check that missing values are detected
        missing_stats = results['missing_value_stats']
        self.assertGreater(missing_stats['total_missing'], 0)
    
    def test_outlier_detection(self):
        """Test outlier detection."""
        # Create data with outliers
        outlier_data = pd.DataFrame({
            'normal_feature': [1, 2, 3, 4, 5, 6, 7, 8, 9, 10],
            'outlier_feature': [1, 2, 3, 4, 5, 6, 7, 8, 9, 100]  # 100 is an outlier
        })
        
        results = self.validator._detect_outliers(outlier_data)
        
        self.assertIn('iqr_outliers', results)
        self.assertIn('zscore_outliers', results)
        self.assertIn('isolation_forest_outliers', results)
        
        # Check that outliers are detected
        iqr_outliers = results['iqr_outliers']
        self.assertGreater(iqr_outliers['outlier_feature'], 0)
    
    def test_establish_and_compare_baseline(self):
        """Test baseline establishment and comparison."""
        # Establish baseline
        baseline = self.validator.establish_baseline(self.good_data)
        self.assertIsNotNone(baseline)
        self.assertIn('data_shape', baseline)
        self.assertIn('missing_rates', baseline)
        
        # Compare similar data
        similar_data = self.good_data.copy()
        similar_data['molecular_weight'] *= 1.1  # Slight change
        
        comparison = self.validator.compare_to_baseline(similar_data, "similar_data")
        self.assertIn('baseline_comparison_passed', comparison)
        self.assertIn('comparison_issues', comparison)
        self.assertIn('comparison_warnings', comparison)


class TestDataAnomalyDetector(unittest.TestCase):
    """Test data anomaly detection functionality."""
    
    def setUp(self):
        """Set up test fixtures."""
        self.detector = DataAnomalyDetector()
        
        # Create baseline data
        np.random.seed(42)
        self.baseline_data = pd.DataFrame({
            'feature_1': np.random.normal(0, 1, 100),
            'feature_2': np.random.normal(5, 2, 100),
            'feature_3': np.random.normal(10, 3, 100)
        })
        
        # Create test data with anomalies
        self.test_data = self.baseline_data.copy()
        self.test_data.loc[0, 'feature_1'] = 10  # Anomaly
        self.test_data.loc[1, 'feature_2'] = -10  # Anomaly
    
    def test_fit_baseline(self):
        """Test fitting baseline data."""
        self.detector.fit_baseline(self.baseline_data)
        
        # Check that detectors are fitted
        self.assertIn('zscore', self.detector.detectors)
        self.assertIn('iqr', self.detector.detectors)
        
        # Check detector parameters
        zscore_detector = self.detector.detectors['zscore']
        self.assertIn('mean', zscore_detector)
        self.assertIn('std', zscore_detector)
    
    def test_detect_anomalies(self):
        """Test anomaly detection."""
        self.detector.fit_baseline(self.baseline_data)
        results = self.detector.detect_anomalies(self.test_data)
        
        self.assertIn('anomaly_detection_methods', results)
        self.assertIn('total_anomalies', results)
        self.assertIn('anomaly_indices', results)
        self.assertIn('column_anomaly_counts', results)
        
        # Should detect some anomalies
        self.assertGreater(results['total_anomalies'], 0)
    
    def test_detect_anomalies_without_baseline(self):
        """Test anomaly detection without baseline."""
        with self.assertRaises(ValueError):
            self.detector.detect_anomalies(self.test_data)


class TestMolecularFeatureExtractorRobustness(unittest.TestCase):
    """Test molecular feature extractor robustness features."""
    
    def setUp(self):
        """Set up test fixtures."""
        self.temp_dir = Path(tempfile.mkdtemp())
        
        # Create a mock PDBQT file
        self.mock_pdbqt = self.temp_dir / "test_compound.pdbqt"
        self.mock_pdbqt.write_text("""REMARK  Generated by test
ATOM      1  C   LIG A   1      20.154  16.967  25.662  0.00  0.00      A    C
ATOM      2  N   LIG A   1      19.032  16.123  26.123  0.00  0.00      A    N
ENDMDL
""")
    
    def tearDown(self):
        """Clean up test fixtures."""
        shutil.rmtree(self.temp_dir, ignore_errors=True)
    
    def test_extractor_initialization_with_robustness(self):
        """Test extractor initialization with robustness features."""
        try:
            extractor = MolecularFeatureExtractor(
                enable_checkpointing=True,
                checkpoint_interval=50,
                robustness_config={'max_retry_attempts': 3}
            )
            
            self.assertTrue(extractor.enable_checkpointing)
            self.assertEqual(extractor.checkpoint_interval, 50)
            self.assertIsNotNone(extractor.robustness_manager)
            self.assertIsNotNone(extractor.checkpoint_manager)
            
        except Exception as e:
            # If initialization fails due to missing dependencies, that's okay for this test
            self.skipTest(f"Extractor initialization failed due to missing dependencies: {e}")
    
    def test_batch_extract_with_checkpointing(self):
        """Test batch extraction with checkpointing."""
        try:
            extractor = MolecularFeatureExtractor(
                enable_checkpointing=True,
                checkpoint_interval=1  # Checkpoint after every file
            )
            
            # Create multiple mock files
            for i in range(3):
                mock_file = self.temp_dir / f"compound_{i}.pdbqt"
                mock_file.write_text("""REMARK  Generated by test
ATOM      1  C   LIG A   1      20.154  16.967  25.662  0.00  0.00      A    C
ENDMDL
""")
            
            # Test batch extraction with checkpointing
            with patch.object(extractor, 'extract_features') as mock_extract:
                mock_extract.return_value = MagicMock()
                mock_extract.return_value.to_dict.return_value = {'compound_name': 'test'}
                
                # This should not raise an exception even if individual extractions fail
                try:
                    results = extractor.batch_extract(str(self.temp_dir))
                    self.assertIsNotNone(results)
                except Exception as e:
                    # If batch extraction fails due to missing dependencies, that's okay
                    self.skipTest(f"Batch extraction failed due to missing dependencies: {e}")
            
        except Exception as e:
            self.skipTest(f"Test skipped due to missing dependencies: {e}")


class TestErrorHandlingIntegration(unittest.TestCase):
    """Integration tests for error handling across components."""
    
    def setUp(self):
        """Set up test fixtures."""
        self.temp_dir = Path(tempfile.mkdtemp())
        
    def tearDown(self):
        """Clean up test fixtures."""
        shutil.rmtree(self.temp_dir, ignore_errors=True)
    
    def test_end_to_end_error_handling(self):
        """Test end-to-end error handling workflow."""
        # Test robustness manager
        robustness_manager = RobustnessManager()
        
        # Test error logging
        test_error = ValueError("Test error")
        robustness_manager.log_error(test_error, {'context': 'test'})
        
        self.assertEqual(len(robustness_manager.error_log), 1)
        self.assertEqual(robustness_manager.error_log[0]['error_type'], 'ValueError')
        
        # Test checkpoint manager
        checkpoint_manager = CheckpointManager(str(self.temp_dir / "checkpoints"))
        
        test_data = {'test': 'data', 'numbers': [1, 2, 3]}
        success = checkpoint_manager.save_checkpoint("test_checkpoint", test_data)
        self.assertTrue(success)
        
        loaded_data = checkpoint_manager.load_checkpoint("test_checkpoint")
        self.assertEqual(loaded_data, test_data)
        
        # Test data quality validator
        validator = DataQualityValidator()
        
        good_data = pd.DataFrame({
            'feature_1': [1, 2, 3, 4, 5],
            'feature_2': [10, 20, 30, 40, 50]
        })
        
        results = validator.validate_dataset(good_data, "test_data")
        self.assertTrue(results['validation_passed'])
        self.assertGreater(results['quality_score'], 0.8)
    
    def test_custom_exception_handling(self):
        """Test custom exception classes."""
        # Test PipelineError
        with self.assertRaises(PipelineError) as context:
            raise PipelineError("Test pipeline error", "TEST_ERROR", {'context': 'test'})
        
        self.assertEqual(context.exception.error_code, "TEST_ERROR")
        self.assertIn('context', context.exception.context)
        
        # Test DataQualityError
        with self.assertRaises(DataQualityError) as context:
            raise DataQualityError("Test quality error", ['issue1', 'issue2'])
        
        self.assertEqual(context.exception.quality_issues, ['issue1', 'issue2'])
        
        # Test ModelTrainingError
        with self.assertRaises(ModelTrainingError) as context:
            raise ModelTrainingError("Test training error", "test_model")
        
        self.assertEqual(context.exception.model_name, "test_model")
        
        # Test FeatureExtractionError
        with self.assertRaises(FeatureExtractionError) as context:
            raise FeatureExtractionError("Test extraction error", "test_compound")
        
        self.assertEqual(context.exception.compound_name, "test_compound")


def run_error_handling_tests():
    """Run all error handling and robustness tests."""
    # Configure logging
    logging.basicConfig(level=logging.WARNING)  # Reduce log noise during tests
    
    # Create test suite
    test_suite = unittest.TestSuite()
    
    # Add test cases
    test_classes = [
        TestErrorHandlingUtilities,
        TestCheckpointManager,
        TestDataQualityValidator,
        TestDataAnomalyDetector,
        TestMolecularFeatureExtractorRobustness,
        TestErrorHandlingIntegration
    ]
    
    for test_class in test_classes:
        tests = unittest.TestLoader().loadTestsFromTestCase(test_class)
        test_suite.addTests(tests)
    
    # Run tests
    runner = unittest.TextTestRunner(verbosity=2)
    result = runner.run(test_suite)
    
    return result.wasSuccessful()


if __name__ == "__main__":
    success = run_error_handling_tests()
    if success:
        print("\n✓ All error handling and robustness tests passed!")
    else:
        print("\n✗ Some error handling and robustness tests failed!")
        exit(1)
```
