## Supplementary material for "Machine Learning-Driven Prediction of TLR4 Binding Affinity: A Comprehensive Molecular Feature Analysis for Drug Discovery": Codebase: test_evaluation_framework.py

```
"""
Standalone test for the comprehensive evaluation framework.

This script tests the evaluation framework without requiring all project dependencies.
"""

import sys
import os
import numpy as np
import pandas as pd
import matplotlib
matplotlib.use('Agg')  # Use non-interactive backend
import matplotlib.pyplot as plt
from pathlib import Path
import tempfile
import shutil

# Add src to path
sys.path.append(str(Path(__file__).parent.parent / "src"))

from tlr4_binding.ml_components.evaluator import ModelEvaluator, PerformanceMetrics
from sklearn.ensemble import RandomForestRegressor
from sklearn.linear_model import LinearRegression
from sklearn.svm import SVR
from sklearn.model_selection import train_test_split
from sklearn.datasets import make_regression

def test_basic_evaluation():
    """Test basic evaluation functionality."""
    print("Testing basic evaluation functionality...")
    
    # Create synthetic data
    X, y = make_regression(n_samples=200, n_features=10, noise=0.1, random_state=42)
    X_train, X_test, y_train, y_test = train_test_split(X, y, test_size=0.3, random_state=42)
    
    # Train a model
    model = RandomForestRegressor(n_estimators=50, random_state=42)
    model.fit(X_train, y_train)
    y_pred = model.predict(X_test)
    
    # Test evaluator
    evaluator = ModelEvaluator()
    metrics = evaluator.evaluate(y_test, y_pred, "Random Forest")
    
    print(f"✓ Basic evaluation completed")
    print(f"  R² Score: {metrics.metrics['r2']:.4f}")
    print(f"  RMSE: {metrics.metrics['rmse']:.4f}")
    print(f"  MAE: {metrics.metrics['mae']:.4f}")
    
    return True

def test_cross_validation():
    """Test cross-validation functionality."""
    print("\nTesting cross-validation functionality...")
    
    # Create synthetic data
    X, y = make_regression(n_samples=150, n_features=8, noise=0.1, random_state=42)
    
    # Test cross-validation
    evaluator = ModelEvaluator()
    model = RandomForestRegressor(n_estimators=30, random_state=42)
    
    cv_results = evaluator.cross_validate_model(model, X, y, "RF Test", cv=3)
    
    print(f"✓ Cross-validation completed")
    print(f"  Mean R²: {cv_results['r2']['mean']:.4f} ± {cv_results['r2']['std']:.4f}")
    print(f"  Mean RMSE: {cv_results['rmse']['mean']:.4f} ± {cv_results['rmse']['std']:.4f}")
    
    return True

def test_learning_curves():
    """Test learning curves functionality."""
    print("\nTesting learning curves functionality...")
    
    # Create synthetic data
    X, y = make_regression(n_samples=200, n_features=6, noise=0.1, random_state=42)
    
    # Test learning curves
    evaluator = ModelEvaluator()
    model = RandomForestRegressor(n_estimators=50, random_state=42)
    
    # Create temporary directory for plots
    with tempfile.TemporaryDirectory() as temp_dir:
        curve_data = evaluator.generate_learning_curves(
            model, X, y, "RF Learning Curves", cv=3,
            save_path=os.path.join(temp_dir, "learning_curves.png")
        )
    
    print(f"✓ Learning curves generated")
    print(f"  Training sizes: {len(curve_data['train_sizes'])}")
    print(f"  Final training RMSE: {curve_data['train_mean'][-1]:.4f}")
    print(f"  Final validation RMSE: {curve_data['val_mean'][-1]:.4f}")
    
    return True

def test_validation_curves():
    """Test validation curves functionality."""
    print("\nTesting validation curves functionality...")
    
    # Create synthetic data
    X, y = make_regression(n_samples=150, n_features=5, noise=0.1, random_state=42)
    
    # Test validation curves
    evaluator = ModelEvaluator()
    model = RandomForestRegressor(random_state=42)
    param_range = [10, 20, 50]
    
    # Create temporary directory for plots
    with tempfile.TemporaryDirectory() as temp_dir:
        curve_data = evaluator.generate_validation_curves(
            model, X, y, 'n_estimators', param_range, "RF Validation Curves", cv=3,
            save_path=os.path.join(temp_dir, "validation_curves.png")
        )
    
    print(f"✓ Validation curves generated")
    print(f"  Parameter range: {len(curve_data['param_range'])} values")
    print(f"  Best parameter index: {np.argmin(curve_data['val_mean'])}")
    
    return True

def test_statistical_testing():
    """Test statistical significance testing."""
    print("\nTesting statistical significance testing...")
    
    # Create synthetic data
    X, y = make_regression(n_samples=100, n_features=5, noise=0.1, random_state=42)
    X_train, X_test, y_train, y_test = train_test_split(X, y, test_size=0.3, random_state=42)
    
    # Train two different models
    model1 = RandomForestRegressor(n_estimators=30, random_state=42)
    model2 = LinearRegression()
    
    model1.fit(X_train, y_train)
    model2.fit(X_train, y_train)
    
    y_pred1 = model1.predict(X_test)
    y_pred2 = model2.predict(X_test)
    
    # Test statistical significance
    evaluator = ModelEvaluator()
    metrics1 = evaluator.evaluate(y_test, y_pred1, "Random Forest")
    metrics2 = evaluator.evaluate(y_test, y_pred2, "Linear Regression")
    
    test_result = evaluator.statistical_significance_test(metrics1, metrics2)
    
    print(f"✓ Statistical significance test completed")
    print(f"  Paired t-test p-value: {test_result['paired_t_test']['p_value']:.4f}")
    print(f"  Significant difference: {test_result['paired_t_test']['significant']}")
    print(f"  R² difference: {test_result['r2_difference']:.4f}")
    
    return True

def test_model_comparison():
    """Test model comparison functionality."""
    print("\nTesting model comparison functionality...")
    
    # Create synthetic data
    X, y = make_regression(n_samples=150, n_features=8, noise=0.1, random_state=42)
    X_train, X_test, y_train, y_test = train_test_split(X, y, test_size=0.3, random_state=42)
    
    # Train multiple models
    models = {
        'Random Forest': RandomForestRegressor(n_estimators=30, random_state=42),
        'Linear Regression': LinearRegression(),
        'SVR': SVR(kernel='rbf', C=1.0, gamma='scale')
    }
    
    evaluator = ModelEvaluator()
    results = {}
    
    for name, model in models.items():
        model.fit(X_train, y_train)
        y_pred = model.predict(X_test)
        metrics = evaluator.evaluate(y_test, y_pred, name)
        results[name] = metrics
    
    # Compare models
    comparison_df = evaluator.compare_models(results)
    
    print(f"✓ Model comparison completed")
    print(f"  Models compared: {len(comparison_df)}")
    print(f"  Best model: {comparison_df.iloc[0]['model_name']}")
    print(f"  Best R²: {comparison_df.iloc[0]['r2_score']:.4f}")
    
    return True

def test_performance_metrics():
    """Test PerformanceMetrics class functionality."""
    print("\nTesting PerformanceMetrics class...")
    
    # Test with perfect predictions
    y_true = np.array([1, 2, 3, 4, 5])
    y_pred_perfect = y_true.copy()
    metrics_perfect = PerformanceMetrics(y_true, y_pred_perfect, "Perfect Model")
    
    print(f"✓ Perfect prediction test")
    print(f"  R² Score: {metrics_perfect.metrics['r2']:.6f}")
    print(f"  RMSE: {metrics_perfect.metrics['rmse']:.6f}")
    
    # Test with poor predictions
    y_pred_poor = y_true + 2.0
    metrics_poor = PerformanceMetrics(y_true, y_pred_poor, "Poor Model")
    
    print(f"✓ Poor prediction test")
    print(f"  R² Score: {metrics_poor.metrics['r2']:.4f}")
    print(f"  RMSE: {metrics_poor.metrics['rmse']:.4f}")
    
    # Test summary and dataframe methods
    summary = metrics_poor.get_summary()
    df = metrics_poor.to_dataframe()
    
    print(f"✓ Summary and DataFrame methods")
    print(f"  Summary keys: {list(summary.keys())}")
    print(f"  DataFrame shape: {df.shape}")
    
    return True

def main():
    """Run all tests."""
    print("=" * 60)
    print("COMPREHENSIVE EVALUATION FRAMEWORK TEST")
    print("=" * 60)
    
    tests = [
        test_performance_metrics,
        test_basic_evaluation,
        test_cross_validation,
        test_learning_curves,
        test_validation_curves,
        test_statistical_testing,
        test_model_comparison
    ]
    
    passed = 0
    total = len(tests)
    
    for test_func in tests:
        try:
            if test_func():
                passed += 1
            else:
                print(f"✗ {test_func.__name__} failed")
        except Exception as e:
            print(f"✗ {test_func.__name__} failed with error: {e}")
            import traceback
            traceback.print_exc()
    
    print("\n" + "=" * 60)
    print(f"TEST RESULTS: {passed}/{total} tests passed")
    print("=" * 60)
    
    if passed == total:
        print("✓ All tests passed! The evaluation framework is working correctly.")
        return True
    else:
        print(f"✗ {total - passed} tests failed. Please check the implementation.")
        return False

if __name__ == "__main__":
    success = main()
    exit(0 if success else 1)
```
