## Supplementary material for "Machine Learning-Driven Prediction of TLR4 Binding Affinity: A Comprehensive Molecular Feature Analysis for Drug Discovery": Codebase: test_feature_analysis.py

```
"""
Tests for feature importance extraction and partial dependence analysis.

This module tests advanced feature analysis capabilities including
feature importance extraction, partial dependence plots, and
model interpretability features for traditional ML models.
"""

import unittest
import sys
import numpy as np
import pandas as pd
from unittest.mock import patch, MagicMock

# Add src to path
sys.path.insert(0, str(Path(__file__).parent.parent / "src"))

from tlr4_binding.ml_components.trainer import (
    RandomForestTrainer,
    XGBoostTrainer,
    LightGBMTrainer,
    MLModelTrainer
)


class TestFeatureImportanceExtraction(unittest.TestCase):
    """Test feature importance extraction for different model types."""
    
    def setUp(self):
        """Set up test data."""
        np.random.seed(42)
        self.n_samples = 100
        self.n_features = 10
        
        # Create synthetic data with known feature importance
        X = np.random.randn(self.n_samples, self.n_features)
        # Make first few features more important
        y = (X[:, 0] * 3.0 + X[:, 1] * 2.0 + X[:, 2] * 1.0 + 
             np.random.randn(self.n_samples) * 0.1)
        
        self.X_train = pd.DataFrame(X, columns=[f'feature_{i}' for i in range(self.n_features)])
        self.y_train = pd.Series(y)
    
    def test_random_forest_feature_importance(self):
        """Test Random Forest feature importance extraction."""
        trainer = RandomForestTrainer(n_jobs=1, random_state=42)
        model = trainer.train(self.X_train, self.y_train)
        
        importance = trainer.get_feature_importance(model)
        
        # Check importance structure
        self.assertIsInstance(importance, dict)
        self.assertEqual(len(importance), self.n_features)
        
        # Check that importance values are non-negative
        for feature, imp in importance.items():
            self.assertGreaterEqual(imp, 0)
            self.assertIn(feature, self.X_train.columns)
        
        # Check that first few features have higher importance
        sorted_features = sorted(importance.items(), key=lambda x: x[1], reverse=True)
        top_features = [f[0] for f in sorted_features[:3]]
        self.assertIn('feature_0', top_features)
        self.assertIn('feature_1', top_features)
    
    def test_xgboost_feature_importance(self):
        """Test XGBoost feature importance extraction."""
        trainer = XGBoostTrainer(random_state=42)
        model = trainer.train(self.X_train, self.y_train)
        
        importance = trainer.get_feature_importance(model)
        
        # Check importance structure
        self.assertIsInstance(importance, dict)
        self.assertEqual(len(importance), self.n_features)
        
        # Check that importance values are non-negative
        for feature, imp in importance.items():
            self.assertGreaterEqual(imp, 0)
            self.assertIn(feature, self.X_train.columns)
    
    def test_lightgbm_feature_importance(self):
        """Test LightGBM feature importance extraction."""
        trainer = LightGBMTrainer(random_state=42)
        model = trainer.train(self.X_train, self.y_train)
        
        importance = trainer.get_feature_importance(model)
        
        # Check importance structure
        self.assertIsInstance(importance, dict)
        self.assertEqual(len(importance), self.n_features)
        
        # Check that importance values are non-negative
        for feature, imp in importance.items():
            self.assertGreaterEqual(imp, 0)
            self.assertIn(feature, self.X_train.columns)
    
    def test_feature_importance_consistency(self):
        """Test that feature importance is consistent across multiple runs."""
        trainer = RandomForestTrainer(n_jobs=1, random_state=42)
        
        # Train multiple models with same random state
        importances = []
        for _ in range(3):
            model = trainer.train(self.X_train, self.y_train)
            importance = trainer.get_feature_importance(model)
            importances.append(importance)
        
        # Check that importances are similar (within tolerance)
        for i in range(1, len(importances)):
            for feature in self.X_train.columns:
                diff = abs(importances[0][feature] - importances[i][feature])
                self.assertLess(diff, 0.1)  # Should be very similar with same random state


class TestPartialDependenceAnalysis(unittest.TestCase):
    """Test partial dependence analysis functionality."""
    
    def setUp(self):
        """Set up test data."""
        np.random.seed(42)
        self.n_samples = 200
        self.n_features = 5
        
        # Create data with known interactions
        X = np.random.randn(self.n_samples, self.n_features)
        y = (X[:, 0] * 2.0 + X[:, 1] * 1.5 + X[:, 0] * X[:, 1] * 0.5 + 
             np.random.randn(self.n_samples) * 0.1)
        
        self.X_train = pd.DataFrame(X, columns=[f'feature_{i}' for i in range(self.n_features)])
        self.y_train = pd.Series(y)
    
    def test_partial_dependence_calculation(self):
        """Test partial dependence calculation for a feature."""
        trainer = RandomForestTrainer(n_jobs=1, random_state=42)
        model = trainer.train(self.X_train, self.y_train)
        
        # Calculate partial dependence for feature_0
        feature_name = 'feature_0'
        feature_idx = self.X_train.columns.get_loc(feature_name)
        
        # Create grid of values for the feature
        feature_values = np.linspace(
            self.X_train[feature_name].min(),
            self.X_train[feature_name].max(),
            20
        )
        
        partial_dependence = []
        
        for value in feature_values:
            # Create modified dataset with this feature set to value
            X_modified = self.X_train.copy()
            X_modified[feature_name] = value
            
            # Get predictions
            predictions = model.predict(X_modified)
            partial_dependence.append(np.mean(predictions))
        
        # Check partial dependence structure
        self.assertEqual(len(partial_dependence), len(feature_values))
        self.assertIsInstance(partial_dependence, list)
        
        # Check that partial dependence values are reasonable
        for pd_value in partial_dependence:
            self.assertIsInstance(pd_value, (int, float, np.number))
            self.assertFalse(np.isnan(pd_value))
    
    def test_partial_dependence_monotonicity(self):
        """Test that partial dependence shows expected monotonicity for linear features."""
        trainer = RandomForestTrainer(n_jobs=1, random_state=42)
        model = trainer.train(self.X_train, self.y_train)
        
        # Test feature_0 which has positive coefficient
        feature_name = 'feature_0'
        feature_values = np.linspace(-2, 2, 10)
        
        partial_dependence = []
        for value in feature_values:
            X_modified = self.X_train.copy()
            X_modified[feature_name] = value
            predictions = model.predict(X_modified)
            partial_dependence.append(np.mean(predictions))
        
        # For a feature with positive coefficient, partial dependence should generally increase
        # (allowing for some noise due to interactions)
        correlation = np.corrcoef(feature_values, partial_dependence)[0, 1]
        self.assertGreater(correlation, 0.3)  # Should be positive correlation


class TestModelInterpretability(unittest.TestCase):
    """Test model interpretability features."""
    
    def setUp(self):
        """Set up test data."""
        np.random.seed(42)
        self.n_samples = 100
        self.n_features = 8
        
        X = np.random.randn(self.n_samples, self.n_features)
        y = (X[:, 0] * 2.0 + X[:, 1] * 1.5 + X[:, 2] * 1.0 + 
             np.random.randn(self.n_samples) * 0.1)
        
        self.X_train = pd.DataFrame(X, columns=[f'feature_{i}' for i in range(self.n_features)])
        self.y_train = pd.Series(y)
        
        self.X_test = self.X_train.iloc[:20]
        self.y_test = self.y_train.iloc[:20]
    
    def test_feature_importance_ranking(self):
        """Test that feature importance ranking makes sense."""
        trainer = RandomForestTrainer(n_jobs=1, random_state=42)
        model = trainer.train(self.X_train, self.y_train)
        
        importance = trainer.get_feature_importance(model)
        
        # Sort features by importance
        sorted_features = sorted(importance.items(), key=lambda x: x[1], reverse=True)
        
        # Check that features are properly ranked
        for i in range(len(sorted_features) - 1):
            self.assertGreaterEqual(
                sorted_features[i][1], 
                sorted_features[i + 1][1]
            )
    
    def test_feature_importance_normalization(self):
        """Test that feature importance values are properly normalized."""
        trainer = RandomForestTrainer(n_jobs=1, random_state=42)
        model = trainer.train(self.X_train, self.y_train)
        
        importance = trainer.get_feature_importance(model)
        importance_values = list(importance.values())
        
        # Check that importance values sum to approximately 1
        total_importance = sum(importance_values)
        self.assertAlmostEqual(total_importance, 1.0, places=2)
        
        # Check that all values are between 0 and 1
        for value in importance_values:
            self.assertGreaterEqual(value, 0)
            self.assertLessEqual(value, 1)
    
    def test_model_explanation_consistency(self):
        """Test that model explanations are consistent across similar inputs."""
        trainer = RandomForestTrainer(n_jobs=1, random_state=42)
        model = trainer.train(self.X_train, self.y_train)
        
        # Test with similar inputs
        X_similar = self.X_test.copy()
        X_similar.iloc[0, 0] += 0.01  # Small change to first feature
        
        pred1 = model.predict(self.X_test.iloc[:1])
        pred2 = model.predict(X_similar.iloc[:1])
        
        # Predictions should be similar for similar inputs
        self.assertLess(abs(pred1[0] - pred2[0]), 0.5)


class TestAdvancedFeatureAnalysis(unittest.TestCase):
    """Test advanced feature analysis capabilities."""
    
    def setUp(self):
        """Set up test data with complex relationships."""
        np.random.seed(42)
        self.n_samples = 300
        self.n_features = 12
        
        # Create complex synthetic data
        X = np.random.randn(self.n_samples, self.n_features)
        y = (
            X[:, 0] * 2.0 +                    # Linear
            X[:, 1] ** 2 * 0.5 +               # Quadratic
            X[:, 2] * X[:, 3] * 0.8 +          # Interaction
            np.sin(X[:, 4]) * 1.2 +            # Non-linear
            np.exp(-X[:, 5] ** 2) * 0.5 +      # Gaussian
            np.random.randn(self.n_samples) * 0.1
        )
        
        self.X_train = pd.DataFrame(X, columns=[f'feature_{i}' for i in range(self.n_features)])
        self.y_train = pd.Series(y)
    
    def test_multi_model_feature_importance_comparison(self):
        """Test comparing feature importance across different models."""
        # Train multiple models
        rf_trainer = RandomForestTrainer(n_jobs=1, random_state=42)
        xgb_trainer = XGBoostTrainer(random_state=42)
        
        rf_model = rf_trainer.train(self.X_train, self.y_train)
        xgb_model = xgb_trainer.train(self.X_train, self.y_train)
        
        rf_importance = rf_trainer.get_feature_importance(rf_model)
        xgb_importance = xgb_trainer.get_feature_importance(xgb_model)
        
        # Both should identify important features
        rf_top_features = sorted(rf_importance.items(), key=lambda x: x[1], reverse=True)[:3]
        xgb_top_features = sorted(xgb_importance.items(), key=lambda x: x[1], reverse=True)[:3]
        
        # Check that both models identify feature_0 as important
        rf_top_feature_names = [f[0] for f in rf_top_features]
        xgb_top_feature_names = [f[0] for f in xgb_top_features]
        
        self.assertIn('feature_0', rf_top_feature_names)
        self.assertIn('feature_0', xgb_top_feature_names)
    
    def test_feature_importance_stability(self):
        """Test stability of feature importance across different random seeds."""
        importances = []
        
        for seed in [42, 123, 456]:
            trainer = RandomForestTrainer(n_jobs=1, random_state=seed)
            model = trainer.train(self.X_train, self.y_train)
            importance = trainer.get_feature_importance(model)
            importances.append(importance)
        
        # Check that top features are consistent across seeds
        top_features_per_seed = []
        for importance in importances:
            top_features = sorted(importance.items(), key=lambda x: x[1], reverse=True)[:5]
            top_features_per_seed.append([f[0] for f in top_features])
        
        # Check overlap in top features
        common_top_features = set(top_features_per_seed[0]) & set(top_features_per_seed[1]) & set(top_features_per_seed[2])
        self.assertGreaterEqual(len(common_top_features), 2)  # At least 2 common top features


if __name__ == '__main__':
    # Run tests with verbose output
    unittest.main(verbosity=2)
```
