## Supplementary material for "Machine Learning-Driven Prediction of TLR4 Binding Affinity: A Comprehensive Molecular Feature Analysis for Drug Discovery": Codebase: test_feature_engineering.py

```
"""
Unit tests for feature engineering pipeline components.

Tests all feature engineering transformations including scaling,
correlation analysis, and feature selection.
"""

import pytest
import numpy as np
import pandas as pd
from unittest.mock import patch
import sys
import os

# Add src to path for imports
sys.path.insert(0, os.path.join(os.path.dirname(__file__), '..', 'src'))

from tlr4_binding.ml_components.feature_engineering import (
    FeatureScaler,
    CorrelationAnalyzer,
    FeatureSelector,
    FeatureEngineeringPipeline,
    FeatureEngineeringConfig
)


class TestFeatureScaler:
    """Test cases for FeatureScaler class."""
    
    def setup_method(self):
        """Set up test data."""
        np.random.seed(42)
        self.X = pd.DataFrame({
            'feature1': np.random.normal(10, 2, 100),
            'feature2': np.random.normal(5, 1, 100),
            'feature3': np.random.normal(0, 0.5, 100)
        })
        
    def test_standard_scaler_fit_transform(self):
        """Test standard scaler fit and transform."""
        scaler = FeatureScaler(use_robust=False)
        X_scaled = scaler.fit_transform(self.X)
        
        # Check that means are approximately 0
        assert np.allclose(X_scaled.mean(), 0, atol=1e-10)
        
        # Check that standard deviations are approximately 1
        assert np.allclose(X_scaled.std(), 1, atol=1e-2)
        
        # Check that column names are preserved
        assert list(X_scaled.columns) == list(self.X.columns)
        
    def test_robust_scaler_fit_transform(self):
        """Test robust scaler fit and transform."""
        scaler = FeatureScaler(use_robust=True)
        X_scaled = scaler.fit_transform(self.X)
        
        # Check that medians are approximately 0
        assert np.allclose(X_scaled.median(), 0, atol=1e-10)
        
        # Check that column names are preserved
        assert list(X_scaled.columns) == list(self.X.columns)
        
    def test_scaler_consistency(self):
        """Test that transform produces same result as fit_transform."""
        scaler1 = FeatureScaler()
        scaler2 = FeatureScaler()
        
        # Fit and transform separately
        scaler1.fit(self.X)
        X_scaled1 = scaler1.transform(self.X)
        
        # Fit and transform together
        X_scaled2 = scaler2.fit_transform(self.X)
        
        # Results should be identical
        pd.testing.assert_frame_equal(X_scaled1, X_scaled2)
        
    def test_scaler_with_different_data(self):
        """Test scaler with different data after fitting."""
        scaler = FeatureScaler()
        scaler.fit(self.X)
        
        # Create new data with different scale
        X_new = self.X * 2 + 5
        X_scaled = scaler.transform(X_new)
        
        # Should be properly scaled
        assert X_scaled.shape == X_new.shape
        assert list(X_scaled.columns) == list(X_new.columns)
        
    def test_scaler_empty_dataframe(self):
        """Test scaler with empty DataFrame."""
        scaler = FeatureScaler()
        empty_df = pd.DataFrame()
        
        with pytest.raises(ValueError, match="Cannot fit scaler on empty DataFrame"):
            scaler.fit(empty_df)
            
    def test_scaler_missing_features(self):
        """Test scaler with missing features in transform."""
        scaler = FeatureScaler()
        scaler.fit(self.X)
        
        # Create data with missing features
        X_missing = self.X[['feature1', 'feature2']]  # Missing feature3
        
        with pytest.raises(ValueError, match="Missing features"):
            scaler.transform(X_missing)
            
    def test_scaler_not_fitted(self):
        """Test scaler transform without fitting."""
        scaler = FeatureScaler()
        
        with pytest.raises(ValueError, match="Scaler must be fitted"):
            scaler.transform(self.X)


class TestCorrelationAnalyzer:
    """Test cases for CorrelationAnalyzer class."""
    
    def setup_method(self):
        """Set up test data with known correlations."""
        np.random.seed(42)
        n_samples = 100
        
        # Create features with known correlations
        base_feature = np.random.normal(0, 1, n_samples)
        
        self.X = pd.DataFrame({
            'feature1': base_feature,
            'feature2': base_feature + np.random.normal(0, 0.01, n_samples),  # Very high correlation
            'feature3': base_feature * 1.1 + np.random.normal(0, 0.01, n_samples),  # Very high correlation
            'feature4': np.random.normal(0, 1, n_samples),  # Low correlation
            'feature5': np.random.normal(0, 1, n_samples)   # Low correlation
        })
        
    def test_find_correlated_features(self):
        """Test finding highly correlated features."""
        analyzer = CorrelationAnalyzer(threshold=0.95)
        result = analyzer.find_correlated_features(self.X)
        
        # Should find highly correlated pairs
        assert len(result['correlated_pairs']) > 0
        assert len(result['features_to_remove']) > 0
        
        # Check that correlations are above threshold
        for pair in result['correlated_pairs']:
            assert pair['correlation'] > 0.95
            
    def test_remove_correlated_features(self):
        """Test removing highly correlated features."""
        analyzer = CorrelationAnalyzer(threshold=0.95)
        X_reduced = analyzer.remove_correlated_features(self.X)
        
        # Should have fewer features
        assert X_reduced.shape[1] < self.X.shape[1]
        
        # Should not have any highly correlated pairs remaining
        corr_matrix = X_reduced.corr().abs()
        upper_tri = corr_matrix.where(
            np.triu(np.ones(corr_matrix.shape), k=1).astype(bool)
        )
        max_corr = upper_tri.max().max()
        assert max_corr <= 0.95 or pd.isna(max_corr)
        
    def test_no_correlated_features(self):
        """Test with data that has no highly correlated features."""
        # Create uncorrelated data
        X_uncorr = pd.DataFrame({
            'feature1': np.random.normal(0, 1, 100),
            'feature2': np.random.normal(0, 1, 100),
            'feature3': np.random.normal(0, 1, 100)
        })
        
        analyzer = CorrelationAnalyzer(threshold=0.9)
        X_reduced = analyzer.remove_correlated_features(X_uncorr)
        
        # Should keep all features
        assert X_reduced.shape[1] == X_uncorr.shape[1]
        assert len(analyzer.features_to_remove) == 0
        
    def test_different_thresholds(self):
        """Test with different correlation thresholds."""
        # Test with very high threshold
        analyzer_high = CorrelationAnalyzer(threshold=0.99)
        X_high = analyzer_high.remove_correlated_features(self.X)
        
        # Test with lower threshold
        analyzer_low = CorrelationAnalyzer(threshold=0.5)
        X_low = analyzer_low.remove_correlated_features(self.X)
        
        # Lower threshold should remove more features
        assert X_low.shape[1] <= X_high.shape[1]


class TestFeatureSelector:
    """Test cases for FeatureSelector class."""
    
    def setup_method(self):
        """Set up test data with known feature importance."""
        np.random.seed(42)
        n_samples = 200
        
        # Create features with different importance levels
        important_feature = np.random.normal(0, 1, n_samples)
        noise_feature = np.random.normal(0, 1, n_samples)
        
        # Create target that depends on important features
        y = (2 * important_feature + 
             0.1 * noise_feature + 
             np.random.normal(0, 0.1, n_samples))
        
        self.X = pd.DataFrame({
            'important1': important_feature,
            'important2': important_feature * 0.8 + np.random.normal(0, 0.1, n_samples),
            'noise1': noise_feature,
            'noise2': np.random.normal(0, 1, n_samples),
            'noise3': np.random.normal(0, 1, n_samples)
        })
        
        self.y = pd.Series(y)
        
    def test_mutual_info_selection(self):
        """Test feature selection using mutual information."""
        selector = FeatureSelector(k=2, method='mutual_info')
        X_selected = selector.fit_transform(self.X, self.y)
        
        # Should select k features
        assert X_selected.shape[1] == 2
        
        # Should select the most important features
        assert 'important1' in X_selected.columns
        assert 'important2' in X_selected.columns
        
    def test_f_regression_selection(self):
        """Test feature selection using F-regression."""
        selector = FeatureSelector(k=2, method='f_regression')
        X_selected = selector.fit_transform(self.X, self.y)
        
        # Should select k features
        assert X_selected.shape[1] == 2
        
    def test_selection_with_insufficient_features(self):
        """Test selection when k > number of features."""
        selector = FeatureSelector(k=10, method='mutual_info')
        X_selected = selector.fit_transform(self.X, self.y)
        
        # Should select all available features
        assert X_selected.shape[1] == self.X.shape[1]
        
    def test_get_feature_importance(self):
        """Test getting feature importance scores."""
        selector = FeatureSelector(k=3, method='mutual_info')
        selector.fit(self.X, self.y)
        
        importance_df = selector.get_feature_importance()
        
        # Should have importance scores
        assert not importance_df.empty
        assert 'feature' in importance_df.columns
        assert 'score' in importance_df.columns
        
        # Should be sorted by score
        assert importance_df['score'].is_monotonic_decreasing
        
    def test_selection_consistency(self):
        """Test that selection is consistent across calls."""
        selector1 = FeatureSelector(k=2, method='mutual_info')
        selector2 = FeatureSelector(k=2, method='mutual_info')
        
        X1 = selector1.fit_transform(self.X, self.y)
        X2 = selector2.fit_transform(self.X, self.y)
        
        # Should select same features
        assert set(X1.columns) == set(X2.columns)
        
    def test_selection_with_missing_features(self):
        """Test selection with missing features in transform."""
        selector = FeatureSelector(k=2, method='mutual_info')
        selector.fit(self.X, self.y)
        
        # Create data with missing features
        X_missing = self.X[['important1', 'noise1']]  # Missing some features
        X_selected = selector.transform(X_missing)
        
        # Should handle missing features gracefully
        assert X_selected.shape[1] <= X_missing.shape[1]


class TestFeatureEngineeringPipeline:
    """Test cases for FeatureEngineeringPipeline class."""
    
    def setup_method(self):
        """Set up test data."""
        np.random.seed(42)
        n_samples = 200
        
        # Create features with correlations and different importance
        base_feature = np.random.normal(0, 1, n_samples)
        
        self.X = pd.DataFrame({
            'important1': base_feature,
            'important2': base_feature * 0.8 + np.random.normal(0, 0.1, n_samples),
            'correlated1': base_feature + np.random.normal(0, 0.05, n_samples),  # High correlation
            'correlated2': base_feature * 1.1 + np.random.normal(0, 0.05, n_samples),  # High correlation
            'noise1': np.random.normal(0, 1, n_samples),
            'noise2': np.random.normal(0, 1, n_samples),
            'noise3': np.random.normal(0, 1, n_samples)
        })
        
        # Create target
        self.y = pd.Series(2 * base_feature + np.random.normal(0, 0.1, n_samples))
        
    def test_pipeline_fit_transform(self):
        """Test complete pipeline fit and transform."""
        config = FeatureEngineeringConfig(
            correlation_threshold=0.9,
            mutual_info_k=3,
            use_robust_scaling=False
        )
        
        pipeline = FeatureEngineeringPipeline(config)
        X_transformed = pipeline.fit_transform(self.X, self.y)
        
        # Should have fewer features than original
        assert X_transformed.shape[1] <= self.X.shape[1]
        
        # Should have same number of samples
        assert X_transformed.shape[0] == self.X.shape[0]
        
    def test_pipeline_with_pca(self):
        """Test pipeline with PCA enabled."""
        config = FeatureEngineeringConfig(
            correlation_threshold=0.9,
            mutual_info_k=5,
            apply_pca=True,
            pca_components=3
        )
        
        pipeline = FeatureEngineeringPipeline(config)
        X_transformed = pipeline.fit_transform(self.X, self.y)
        
        # Should have PCA components
        assert X_transformed.shape[1] == 3
        assert all(col.startswith('PC_') for col in X_transformed.columns)
        
    def test_pipeline_consistency(self):
        """Test that pipeline is consistent across calls."""
        config = FeatureEngineeringConfig(mutual_info_k=3)
        
        pipeline1 = FeatureEngineeringPipeline(config)
        pipeline2 = FeatureEngineeringPipeline(config)
        
        X1 = pipeline1.fit_transform(self.X, self.y)
        X2 = pipeline2.fit_transform(self.X, self.y)
        
        # Should produce same results
        pd.testing.assert_frame_equal(X1, X2)
        
    def test_pipeline_transform_only(self):
        """Test pipeline transform after fitting."""
        pipeline = FeatureEngineeringPipeline()
        pipeline.fit(self.X, self.y)
        
        # Transform same data
        X_transformed = pipeline.transform(self.X)
        
        # Should work without errors
        assert X_transformed.shape[0] == self.X.shape[0]
        
    def test_pipeline_get_summary(self):
        """Test getting pipeline summary."""
        pipeline = FeatureEngineeringPipeline()
        pipeline.fit(self.X, self.y)
        
        summary = pipeline.get_pipeline_summary()
        
        # Should have all expected keys
        expected_keys = ['config', 'correlation_analysis', 'feature_importance', 'final_features']
        for key in expected_keys:
            assert key in summary
            
    def test_pipeline_get_feature_importance(self):
        """Test getting feature importance from pipeline."""
        pipeline = FeatureEngineeringPipeline()
        pipeline.fit(self.X, self.y)
        
        importance_df = pipeline.get_feature_importance()
        
        # Should have importance data
        assert not importance_df.empty
        assert 'feature' in importance_df.columns
        assert 'score' in importance_df.columns
        
    def test_pipeline_not_fitted(self):
        """Test pipeline methods without fitting."""
        pipeline = FeatureEngineeringPipeline()
        
        with pytest.raises(ValueError, match="Pipeline must be fitted"):
            pipeline.transform(self.X)
            
        with pytest.raises(ValueError, match="Pipeline must be fitted"):
            pipeline.get_feature_importance()
            
    def test_pipeline_with_robust_scaling(self):
        """Test pipeline with robust scaling."""
        config = FeatureEngineeringConfig(use_robust_scaling=True)
        pipeline = FeatureEngineeringPipeline(config)
        
        X_transformed = pipeline.fit_transform(self.X, self.y)
        
        # Should work with robust scaling
        assert X_transformed.shape[0] == self.X.shape[0]
        assert X_transformed.shape[1] <= self.X.shape[1]


class TestFeatureEngineeringConfig:
    """Test cases for FeatureEngineeringConfig class."""
    
    def test_default_config(self):
        """Test default configuration values."""
        config = FeatureEngineeringConfig()
        
        assert config.correlation_threshold == 0.95
        assert config.mutual_info_k == 20
        assert config.use_robust_scaling == False
        assert config.apply_pca == False
        assert config.pca_components is None
        assert config.min_variance == 1e-6
        
    def test_custom_config(self):
        """Test custom configuration values."""
        config = FeatureEngineeringConfig(
            correlation_threshold=0.8,
            mutual_info_k=10,
            use_robust_scaling=True,
            apply_pca=True,
            pca_components=5,
            min_variance=1e-4
        )
        
        assert config.correlation_threshold == 0.8
        assert config.mutual_info_k == 10
        assert config.use_robust_scaling == True
        assert config.apply_pca == True
        assert config.pca_components == 5
        assert config.min_variance == 1e-4


if __name__ == "__main__":
    pytest.main([__file__, "-v"])
```
