## Supplementary material for "Machine Learning-Driven Prediction of TLR4 Binding Affinity: A Comprehensive Molecular Feature Analysis for Drug Discovery": Codebase: test_gnn_models.py

```
"""
Unit tests for Graph Neural Network models.

This module contains comprehensive tests for GNN model training,
graph processing, and visualization functionality.
"""

import pytest
import numpy as np
import pandas as pd
from pathlib import Path
import tempfile
import shutil
from unittest.mock import Mock, patch, MagicMock

# Test imports
import sys
sys.path.append(str(Path(__file__).parent.parent / "src"))

from tlr4_binding.ml_components.gnn_models import (
    MolecularGraph, GraphDataset, MolecularGraphBuilder,
    GraphConvModel, MPNNModel, AttentiveFPModel,
    GNNTrainer, GNNModelTrainer
)
from tlr4_binding.ml_components.gnn_visualization import (
    GraphVisualizer, GraphFeatureExtractor, GNNModelAnalyzer
)


class TestMolecularGraph:
    """Test MolecularGraph data structure."""
    
    def test_molecular_graph_creation(self):
        """Test MolecularGraph creation with valid data."""
        node_features = np.random.rand(10, 9)
        edge_index = np.array([[0, 1, 2], [1, 2, 0]])
        edge_attr = np.random.rand(3, 2)
        
        graph = MolecularGraph(
            node_features=node_features,
            edge_index=edge_index,
            edge_attr=edge_attr,
            num_nodes=10,
            num_edges=3,
            compound_name="test_compound"
        )
        
        assert graph.num_nodes == 10
        assert graph.num_edges == 3
        assert graph.compound_name == "test_compound"
        assert graph.node_features.shape == (10, 9)
        assert graph.edge_index.shape == (2, 3)
        assert graph.edge_attr.shape == (3, 2)


class TestGraphDataset:
    """Test GraphDataset functionality."""
    
    def test_graph_dataset_creation(self):
        """Test GraphDataset creation."""
        # Create mock graphs
        graphs = []
        for i in range(5):
            graph = MolecularGraph(
                node_features=np.random.rand(10, 9),
                edge_index=np.array([[0, 1], [1, 0]]),
                edge_attr=np.random.rand(2, 2),
                num_nodes=10,
                num_edges=2,
                compound_name=f"compound_{i}"
            )
            graphs.append(graph)
        
        targets = [1.0, 2.0, 3.0, 4.0, 5.0]
        dataset = GraphDataset(graphs, targets)
        
        assert len(dataset) == 5
        assert dataset.targets == targets
    
    def test_graph_dataset_getitem(self):
        """Test GraphDataset __getitem__ method."""
        # Create mock graph
        graph = MolecularGraph(
            node_features=np.random.rand(5, 9),
            edge_index=np.array([[0, 1], [1, 0]]),
            edge_attr=np.random.rand(2, 2),
            num_nodes=5,
            num_edges=2,
            compound_name="test_compound"
        )
        
        dataset = GraphDataset([graph], [1.0])
        data = dataset[0]
        
        # Check that data is a PyTorch Geometric Data object
        assert hasattr(data, 'x')
        assert hasattr(data, 'edge_index')
        assert hasattr(data, 'y')
        assert data.x.shape == (5, 9)
        assert data.edge_index.shape == (2, 2)
        # Check y value (handle both tensor and list cases)
        if hasattr(data.y, 'item'):
            assert data.y.item() == 1.0
        else:
            assert data.y[0] == 1.0


class TestMolecularGraphBuilder:
    """Test MolecularGraphBuilder functionality."""
    
    def setup_method(self):
        """Set up test fixtures."""
        self.builder = MolecularGraphBuilder()
        self.temp_dir = tempfile.mkdtemp()
    
    def teardown_method(self):
        """Clean up test fixtures."""
        shutil.rmtree(self.temp_dir)
    
    def test_molecular_graph_builder_initialization(self):
        """Test MolecularGraphBuilder initialization."""
        builder = MolecularGraphBuilder(use_3d_coords=True, max_atoms=100)
        assert builder.use_3d_coords is True
        assert builder.max_atoms == 100
    
    def test_atom_type_to_number(self):
        """Test atom type to number conversion."""
        assert self.builder._atom_type_to_number('C') == 6
        assert self.builder._atom_type_to_number('N') == 7
        assert self.builder._atom_type_to_number('O') == 8
        assert self.builder._atom_type_to_number('H') == 1
        assert self.builder._atom_type_to_number('unknown') == 6  # Default
    
    def test_create_node_features(self):
        """Test node feature creation."""
        coords = np.array([[0, 0, 0], [1, 0, 0], [0, 1, 0]])
        atom_types = [6, 6, 8]  # C, C, O
        
        features = self.builder._create_node_features(coords, atom_types)
        
        assert features.shape == (3, 9)
        assert features[0, 0] == 6  # atomic number
        assert features[1, 0] == 6
        assert features[2, 0] == 8
        assert np.allclose(features[0, 1:4], [0, 0, 0])  # coordinates
        assert np.allclose(features[1, 1:4], [1, 0, 0])
        assert np.allclose(features[2, 1:4], [0, 1, 0])
    
    def test_create_edges(self):
        """Test edge creation from coordinates."""
        coords = np.array([[0, 0, 0], [1, 0, 0], [0, 1, 0]])
        atom_types = [6, 6, 8]
        
        edge_index, edge_attr = self.builder._create_edges(coords, atom_types)
        
        assert edge_index.shape[0] == 2  # source and target
        assert edge_index.shape[1] == edge_attr.shape[0]
        assert edge_attr.shape[1] == 2  # distance and bond type
    
    def test_build_graph_from_coords(self):
        """Test graph building from coordinates."""
        coords = np.array([[0, 0, 0], [1, 0, 0], [0, 1, 0]])
        atom_types = [6, 6, 8]
        
        graph = self.builder._build_graph_from_coords(coords, atom_types, "test")
        
        assert graph.num_nodes == 3
        assert graph.compound_name == "test"
        assert graph.node_features.shape == (3, 9)
        assert graph.edge_index.shape[0] == 2
        assert graph.edge_attr.shape[1] == 2
    
    def test_build_graph_from_pdbqt_invalid_file(self):
        """Test graph building from invalid PDBQT file."""
        invalid_path = "nonexistent_file.pdbqt"
        result = self.builder.build_graph_from_pdbqt(invalid_path)
        assert result is None
    
    def test_build_graph_from_smiles_invalid(self):
        """Test graph building from invalid SMILES."""
        result = self.builder.build_graph_from_smiles("invalid_smiles")
        assert result is None


class TestGNNModels:
    """Test GNN model architectures."""
    
    def test_graphconv_model_creation(self):
        """Test GraphConv model creation."""
        try:
            model = GraphConvModel(input_dim=9, hidden_dim=32, output_dim=1, num_layers=2)
            
            assert model.input_dim == 9
            assert model.hidden_dim == 32
            assert model.output_dim == 1
            assert model.num_layers == 2
            assert len(model.convs) == 2
            assert len(model.batch_norms) == 2
        except RuntimeError as e:
            if "PyTorch Geometric not available" in str(e):
                pytest.skip("PyTorch Geometric not available for GraphConv model")
            else:
                raise
    
    def test_mpnn_model_creation(self):
        """Test MPNN model creation."""
        try:
            model = MPNNModel(input_dim=9, hidden_dim=32, output_dim=1, num_layers=2)
            
            assert model.input_dim == 9
            assert model.hidden_dim == 32
            assert model.output_dim == 1
            assert model.num_layers == 2
            assert len(model.message_layers) == 2
            assert len(model.update_layers) == 2
        except RuntimeError as e:
            if "PyTorch Geometric not available" in str(e):
                pytest.skip("PyTorch Geometric not available for MPNN model")
            else:
                raise
    
    def test_attentivefp_model_creation(self):
        """Test AttentiveFP model creation."""
        try:
            model = AttentiveFPModel(input_dim=9, hidden_dim=32, output_dim=1, num_layers=2)
            
            assert model.input_dim == 9
            assert model.hidden_dim == 32
            assert model.output_dim == 1
            assert model.num_layers == 2
            assert len(model.attention_layers) == 2
        except RuntimeError as e:
            if "PyTorch Geometric not available" in str(e):
                pytest.skip("PyTorch Geometric not available for AttentiveFP model")
            else:
                raise


class TestGNNTrainer:
    """Test GNN trainer functionality."""
    
    def setup_method(self):
        """Set up test fixtures."""
        self.temp_dir = tempfile.mkdtemp()
        try:
            self.trainer = GNNTrainer(model_type="graphconv", device="cpu")
        except RuntimeError as e:
            if "PyTorch not available" in str(e):
                pytest.skip("PyTorch not available for GNN trainer")
            else:
                raise
    
    def teardown_method(self):
        """Clean up test fixtures."""
        shutil.rmtree(self.temp_dir)
    
    def test_gnn_trainer_initialization(self):
        """Test GNNTrainer initialization."""
        try:
            trainer = GNNTrainer(model_type="graphconv", device="cpu")
            assert trainer.model_type == "graphconv"
            assert trainer.device.type == "cpu"
        except RuntimeError as e:
            if "PyTorch not available" in str(e):
                pytest.skip("PyTorch not available for GNN trainer")
            else:
                raise
    
    def test_create_model(self):
        """Test model creation."""
        try:
            model = self.trainer.create_model(input_dim=9, hidden_dim=32, output_dim=1)
            assert isinstance(model, GraphConvModel)
            assert model.input_dim == 9
            assert model.hidden_dim == 32
            assert model.output_dim == 1
        except RuntimeError as e:
            if "PyTorch not available" in str(e):
                pytest.skip("PyTorch not available for GNN trainer")
            else:
                raise
    
    def test_create_model_invalid_type(self):
        """Test model creation with invalid type."""
        try:
            trainer = GNNTrainer(model_type="invalid", device="cpu")
            with pytest.raises(ValueError):
                trainer.create_model(input_dim=9)
        except RuntimeError as e:
            if "PyTorch not available" in str(e):
                pytest.skip("PyTorch not available for GNN trainer")
            else:
                raise


class TestGNNModelTrainer:
    """Test GNN model trainer coordinator."""
    
    def setup_method(self):
        """Set up test fixtures."""
        self.temp_dir = tempfile.mkdtemp()
        self.trainer = GNNModelTrainer(models_dir=self.temp_dir)
    
    def teardown_method(self):
        """Clean up test fixtures."""
        shutil.rmtree(self.temp_dir)
    
    def test_gnn_model_trainer_initialization(self):
        """Test GNNModelTrainer initialization."""
        assert self.trainer.models_dir == Path(self.temp_dir)
        assert self.trainer.graph_builder is not None
    
    def test_prepare_graph_data_empty(self):
        """Test graph data preparation with empty input."""
        dataset = self.trainer.prepare_graph_data([], [])
        assert len(dataset) == 0
    
    def test_calculate_metrics(self):
        """Test metrics calculation."""
        y_true = [1.0, 2.0, 3.0, 4.0, 5.0]
        y_pred = [1.1, 2.1, 2.9, 4.1, 4.9]
        
        metrics = self.trainer._calculate_metrics(y_true, np.array(y_pred))
        
        assert 'mse' in metrics
        assert 'rmse' in metrics
        assert 'mae' in metrics
        assert 'r2' in metrics
        assert metrics['mse'] > 0
        assert metrics['rmse'] > 0
        assert metrics['mae'] > 0
        assert metrics['r2'] > 0


class TestGraphVisualizer:
    """Test graph visualization functionality."""
    
    def setup_method(self):
        """Set up test fixtures."""
        self.temp_dir = tempfile.mkdtemp()
        self.visualizer = GraphVisualizer(output_dir=self.temp_dir)
    
    def teardown_method(self):
        """Clean up test fixtures."""
        shutil.rmtree(self.temp_dir)
    
    def test_graph_visualizer_initialization(self):
        """Test GraphVisualizer initialization."""
        assert self.visualizer.output_dir == Path(self.temp_dir)
    
    def test_normalize_colors(self):
        """Test color normalization."""
        values = np.array([1, 2, 3, 4, 5])
        normalized = self.visualizer._normalize_colors(values)
        
        assert len(normalized) == len(values)
        assert normalized.min() == 0.0
        assert normalized.max() == 1.0
    
    def test_normalize_colors_empty(self):
        """Test color normalization with empty array."""
        values = np.array([])
        normalized = self.visualizer._normalize_colors(values)
        assert len(normalized) == 0
    
    def test_normalize_colors_constant(self):
        """Test color normalization with constant values."""
        values = np.array([5, 5, 5, 5])
        normalized = self.visualizer._normalize_colors(values)
        assert np.allclose(normalized, 0.5)


class TestGraphFeatureExtractor:
    """Test graph feature extraction functionality."""
    
    def setup_method(self):
        """Set up test fixtures."""
        self.extractor = GraphFeatureExtractor()
    
    def test_graph_feature_extractor_initialization(self):
        """Test GraphFeatureExtractor initialization."""
        assert self.extractor is not None
    
    def test_extract_graph_features_empty(self):
        """Test feature extraction with empty input."""
        # Create a mock empty graph
        mock_graph = Mock()
        mock_graph.number_of_nodes.return_value = 0
        mock_graph.number_of_edges.return_value = 0
        
        features = self.extractor.extract_graph_features(mock_graph)
        assert isinstance(features, dict)


class TestGNNModelAnalyzer:
    """Test GNN model analyzer functionality."""
    
    def setup_method(self):
        """Set up test fixtures."""
        self.temp_dir = tempfile.mkdtemp()
        self.analyzer = GNNModelAnalyzer(output_dir=self.temp_dir)
    
    def teardown_method(self):
        """Clean up test fixtures."""
        shutil.rmtree(self.temp_dir)
    
    def test_gnn_model_analyzer_initialization(self):
        """Test GNNModelAnalyzer initialization."""
        assert self.analyzer.output_dir == Path(self.temp_dir)
        assert self.analyzer.visualizer is not None
        assert self.analyzer.feature_extractor is not None
    
    def test_analyze_predictions(self):
        """Test prediction analysis."""
        predictions = np.array([1.0, 2.0, 3.0, 4.0, 5.0])
        true_values = np.array([1.1, 2.1, 2.9, 4.1, 4.9])
        graph_data_list = [Mock() for _ in range(5)]
        compound_names = [f"compound_{i}" for i in range(5)]
        
        analysis = self.analyzer.analyze_predictions(
            predictions, true_values, graph_data_list, compound_names
        )
        
        assert 'predictions' in analysis
        assert 'true_values' in analysis
        assert 'errors' in analysis
        assert 'mae' in analysis
        assert 'rmse' in analysis
        assert 'r2' in analysis
    
    def test_create_model_report(self):
        """Test model report creation."""
        model_results = {
            'model1': {'metrics': {'r2': 0.8, 'rmse': 0.5, 'mae': 0.4}},
            'model2': {'metrics': {'r2': 0.9, 'rmse': 0.3, 'mae': 0.2}}
        }
        
        report = self.analyzer.create_model_report(model_results)
        
        assert isinstance(report, str)
        assert "GNN Model Analysis Report" in report
        assert "Model Performance Comparison" in report
        assert "Best Model" in report


class TestIntegration:
    """Integration tests for GNN models."""
    
    def setup_method(self):
        """Set up test fixtures."""
        self.temp_dir = tempfile.mkdtemp()
    
    def teardown_method(self):
        """Clean up test fixtures."""
        shutil.rmtree(self.temp_dir)
    
    def test_end_to_end_workflow(self):
        """Test complete GNN workflow."""
        # Create mock molecular graphs
        graphs = []
        for i in range(10):
            graph = MolecularGraph(
                node_features=np.random.rand(10, 9),
                edge_index=np.array([[0, 1, 2], [1, 2, 0]]),
                edge_attr=np.random.rand(3, 2),
                num_nodes=10,
                num_edges=3,
                compound_name=f"compound_{i}"
            )
            graphs.append(graph)
        
        targets = np.random.rand(10)
        dataset = GraphDataset(graphs, targets.tolist())
        
        # Test dataset creation
        assert len(dataset) == 10
        assert len(dataset.targets) == 10
        
        # Test data loading
        data = dataset[0]
        assert hasattr(data, 'x')
        assert hasattr(data, 'edge_index')
        assert hasattr(data, 'y')
    
    def test_model_creation_and_forward_pass(self):
        """Test model creation and forward pass."""
        # Create mock data
        mock_data = Mock()
        mock_data.x = np.random.rand(5, 9)
        mock_data.edge_index = np.array([[0, 1, 2], [1, 2, 0]])
        mock_data.batch = np.array([0, 0, 0, 0, 0])
        
        # Test GraphConv model
        try:
            model = GraphConvModel(input_dim=9, hidden_dim=32, output_dim=1)
            assert model is not None
        except RuntimeError as e:
            if "PyTorch Geometric not available" in str(e):
                pytest.skip("PyTorch Geometric not available for GraphConv model")
            else:
                raise
        
        # Test MPNN model
        try:
            model = MPNNModel(input_dim=9, hidden_dim=32, output_dim=1)
            assert model is not None
        except RuntimeError as e:
            if "PyTorch Geometric not available" in str(e):
                pytest.skip("PyTorch Geometric not available for MPNN model")
            else:
                raise
        
        # Test AttentiveFP model
        try:
            model = AttentiveFPModel(input_dim=9, hidden_dim=32, output_dim=1)
            assert model is not None
        except RuntimeError as e:
            if "PyTorch Geometric not available" in str(e):
                pytest.skip("PyTorch Geometric not available for AttentiveFP model")
            else:
                raise


if __name__ == "__main__":
    pytest.main([__file__, "-v"])
```
