## Supplementary material for "Machine Learning-Driven Prediction of TLR4 Binding Affinity: A Comprehensive Molecular Feature Analysis for Drug Discovery": Codebase: test_interpretability.py

```
"""
Tests for Model Interpretability and Analysis Suite

This module contains comprehensive tests for the interpretability tools
used in TLR4 binding prediction.
"""

import pytest
import numpy as np
import pandas as pd
import matplotlib.pyplot as plt
from unittest.mock import Mock, patch, MagicMock
import os
import tempfile
import shutil

# Add src to path
import sys
sys.path.append('src')

from tlr4_binding.ml_components.interpretability import ModelInterpretabilitySuite
from tlr4_binding.ml_components.molecular_substructure_analyzer import MolecularSubstructureAnalyzer
from tlr4_binding.ml_components.attention_visualizer import AttentionVisualizer


class TestModelInterpretabilitySuite:
    """Test cases for ModelInterpretabilitySuite."""
    
    def setup_method(self):
        """Set up test fixtures."""
        # Create sample data
        np.random.seed(42)
        self.n_samples = 100
        self.n_features = 10
        
        self.X_train = pd.DataFrame(
            np.random.randn(self.n_samples, self.n_features),
            columns=[f'feature_{i}' for i in range(self.n_features)]
        )
        self.y_train = pd.Series(np.random.randn(self.n_samples))
        
        self.X_test = pd.DataFrame(
            np.random.randn(20, self.n_features),
            columns=[f'feature_{i}' for i in range(self.n_features)]
        )
        self.y_test = pd.Series(np.random.randn(20))
        
        # Create mock models
        self.models = {
            'RandomForest': Mock(),
            'SVR': Mock(),
            'LinearRegression': Mock()
        }
        
        # Set up mock model predictions
        for model in self.models.values():
            model.predict.return_value = np.random.randn(20)
        
        # Create temporary output directory
        self.temp_dir = tempfile.mkdtemp()
        
        # Initialize interpretability suite
        self.suite = ModelInterpretabilitySuite(
            models=self.models,
            feature_names=[f'feature_{i}' for i in range(self.n_features)],
            X_train=self.X_train,
            y_train=self.y_train,
            X_test=self.X_test,
            y_test=self.y_test,
            output_dir=self.temp_dir
        )
    
    def teardown_method(self):
        """Clean up test fixtures."""
        shutil.rmtree(self.temp_dir, ignore_errors=True)
    
    def test_initialization(self):
        """Test interpretability suite initialization."""
        assert len(self.suite.models) == 3
        assert len(self.suite.feature_names) == self.n_features
        assert len(self.suite.X_train) == self.n_samples
        assert os.path.exists(self.temp_dir)
    
    @patch('tlr4_binding.ml_components.interpretability.SHAP_AVAILABLE', True)
    @patch('tlr4_binding.ml_components.interpretability.shap')
    def test_shap_analysis(self, mock_shap):
        """Test SHAP analysis generation."""
        # Mock SHAP explainer and values
        mock_explainer = Mock()
        mock_shap.Explainer.return_value = mock_explainer
        mock_shap_values = np.random.randn(10, self.n_features)
        mock_explainer.shap_values.return_value = mock_shap_values
        
        # Mock SHAP plotting functions
        mock_shap.summary_plot = Mock()
        mock_shap.waterfall_plot = Mock()
        
        # Run SHAP analysis
        results = self.suite.generate_shap_analysis('RandomForest', self.models['RandomForest'])
        
        # Verify results
        assert 'shap_values' in results
        assert 'feature_importance' in results
        assert 'explainer' in results
        assert len(results['feature_importance']) == self.n_features
    
    @patch('tlr4_binding.ml_components.interpretability.LIME_AVAILABLE', True)
    @patch('tlr4_binding.ml_components.interpretability.lime')
    def test_lime_analysis(self, mock_lime):
        """Test LIME analysis generation."""
        # Mock LIME explainer
        mock_explainer = Mock()
        mock_lime.lime_tabular.LimeTabularExplainer.return_value = mock_explainer
        
        # Mock explanation object
        mock_explanation = Mock()
        mock_explanation.as_pyplot_figure = Mock()
        mock_explainer.explain_instance.return_value = mock_explanation
        
        # Run LIME analysis
        results = self.suite.generate_lime_analysis('RandomForest', self.models['RandomForest'])
        
        # Verify results
        assert 'explanations' in results
        assert 'explainer' in results
        assert len(results['explanations']) > 0
    
    def test_molecular_substructure_analysis(self):
        """Test molecular substructure analysis."""
        # Create sample compound data with SMILES
        compound_data = pd.DataFrame({
            'smiles': ['CCO', 'CCCO', 'CCCCCO'] * 10,
            'mol_weight': np.random.normal(100, 20, 30),
            'mol_logp': np.random.normal(2, 1, 30)
        })
        
        binding_affinities = pd.Series(np.random.normal(-5, 2, 30))
        
        # Run substructure analysis
        results = self.suite.analyze_molecular_substructures(
            compound_data, binding_affinities
        )
        
        # Verify results structure
        assert 'threshold_value' in results
        assert 'strong_binders_count' in results
        assert 'weak_binders_count' in results
        assert 'descriptor_analysis' in results
        assert 'feature_analysis' in results
    
    def test_feature_importance_calculation(self):
        """Test feature importance calculation from SHAP values."""
        # Create sample SHAP values
        shap_values = np.random.randn(10, self.n_features)
        
        # Calculate importance
        importance = self.suite._calculate_shap_importance(shap_values)
        
        # Verify results
        assert len(importance) == self.n_features
        assert all(isinstance(v, (int, float)) for v in importance.values())
        assert all(v >= 0 for v in importance.values())  # Importance should be non-negative
    
    def test_comprehensive_report_generation(self):
        """Test comprehensive report generation."""
        # Add some mock results
        self.suite.interpretability_results = {
            'RandomForest_shap': {'feature_importance': {'feature_0': 0.5}},
            'RandomForest_lime': {'explanations': []},
            'molecular_substructures': {'threshold_value': -5.0}
        }
        
        # Generate report
        report = self.suite.generate_comprehensive_report()
        
        # Verify report structure
        assert 'timestamp' in report
        assert 'models_analyzed' in report
        assert 'interpretability_results' in report
        assert 'summary' in report
        
        # Verify markdown report was created
        assert os.path.exists(os.path.join(self.temp_dir, 'interpretability_report.md'))


class TestMolecularSubstructureAnalyzer:
    """Test cases for MolecularSubstructureAnalyzer."""
    
    def setup_method(self):
        """Set up test fixtures."""
        self.temp_dir = tempfile.mkdtemp()
        self.analyzer = MolecularSubstructureAnalyzer(output_dir=self.temp_dir)
    
    def teardown_method(self):
        """Clean up test fixtures."""
        shutil.rmtree(self.temp_dir, ignore_errors=True)
    
    @patch('tlr4_binding.ml_components.molecular_substructure_analyzer.RDKIT_AVAILABLE', True)
    @patch('tlr4_binding.ml_components.molecular_substructure_analyzer.Chem')
    def test_molecular_feature_extraction(self, mock_chem):
        """Test molecular feature extraction from SMILES."""
        # Mock RDKit functionality
        mock_mol = Mock()
        mock_chem.MolFromSmiles.return_value = mock_mol
        
        # Mock descriptors
        with patch('tlr4_binding.ml_components.molecular_substructure_analyzer.Descriptors') as mock_desc:
            mock_desc.MolWt.return_value = 100.0
            mock_desc.TPSA.return_value = 50.0
            mock_desc.NumHDonors.return_value = 2
            mock_desc.NumHAcceptors.return_value = 3
            mock_desc.NumRotatableBonds.return_value = 1
            mock_desc.NumAromaticRings.return_value = 1
            
            with patch('tlr4_binding.ml_components.molecular_substructure_analyzer.Crippen') as mock_crippen:
                mock_crippen.MolLogP.return_value = 2.5
                
                # Create sample data
                compound_data = pd.DataFrame({
                    'smiles': ['CCO', 'CCCO', 'CCCCCO']
                })
                
                # Extract features
                features = self.analyzer._extract_molecular_features(compound_data, 'smiles')
                
                # Verify results
                assert 'molecules' in features
                assert 'molecular_weights' in features
                assert 'logp_values' in features
                assert len(features['molecular_weights']) == 3
    
    def test_structural_differences_analysis(self):
        """Test structural differences analysis between strong and weak binders."""
        # Create mock molecular features
        molecular_features = {
            'molecular_weights': np.array([100, 150, 200, 120, 180]),
            'logp_values': np.array([1.0, 2.0, 3.0, 1.5, 2.5]),
            'valid_indices': [0, 1, 2, 3, 4]
        }
        
        # Create mock binding data
        strong_binders = pd.Series([True, False, True, False, True])
        weak_binders = pd.Series([False, True, False, True, False])
        
        # Run analysis
        results = self.analyzer._analyze_structural_differences(
            molecular_features, strong_binders, weak_binders
        )
        
        # Verify results
        assert 'molecular_weights' in results
        assert 'logp_values' in results
        
        for desc_name, analysis in results.items():
            assert 'strong_mean' in analysis
            assert 'weak_mean' in analysis
            assert 't_statistic' in analysis
            assert 'p_value' in analysis
            assert 'effect_size' in analysis
    
    def test_pharmacophore_analysis(self):
        """Test pharmacophore feature analysis."""
        # Create mock molecular features
        molecular_features = {
            'hbd_counts': np.array([1, 2, 3, 1, 2]),
            'hba_counts': np.array([2, 3, 4, 2, 3]),
            'aromatic_rings': np.array([1, 2, 1, 1, 2]),
            'rotatable_bonds': np.array([2, 3, 4, 2, 3]),
            'tpsa_values': np.array([50, 60, 70, 55, 65]),
            'logp_values': np.array([1.0, 2.0, 3.0, 1.5, 2.5]),
            'valid_indices': [0, 1, 2, 3, 4]
        }
        
        # Create mock binding data
        strong_binders = pd.Series([True, False, True, False, True])
        weak_binders = pd.Series([False, True, False, True, False])
        
        # Run analysis
        results = self.analyzer._analyze_pharmacophore_features(
            molecular_features, strong_binders, weak_binders
        )
        
        # Verify results
        expected_features = ['hydrogen_bond_donors', 'hydrogen_bond_acceptors', 
                           'aromatic_rings', 'rotatable_bonds', 'polar_surface_area', 'lipophilicity']
        
        for feature in expected_features:
            assert feature in results
            assert 'strong_mean' in results[feature]
            assert 'weak_mean' in results[feature]
            assert 'effect_size' in results[feature]
            assert 'p_value' in results[feature]
    
    def test_fingerprint_similarity_calculation(self):
        """Test molecular fingerprint similarity calculation."""
        # Create mock fingerprints
        fingerprints = [Mock() for _ in range(3)]
        
        # Mock similarity calculation
        with patch('tlr4_binding.ml_components.molecular_substructure_analyzer.DataStructs') as mock_ds:
            mock_ds.TanimotoSimilarity.return_value = 0.5
            
            # Calculate similarity
            similarity = self.analyzer._calculate_fingerprint_similarity(fingerprints)
            
            # Verify results
            assert isinstance(similarity, float)
            assert 0 <= similarity <= 1
    
    def test_substructure_report_generation(self):
        """Test substructure analysis report generation."""
        # Create mock results
        results = {
            'threshold_value': -5.0,
            'strong_binders_count': 10,
            'weak_binders_count': 40,
            'structural_analysis': {
                'mol_weight': {
                    'strong_mean': 200.0,
                    'weak_mean': 150.0,
                    'p_value': 0.01,
                    'effect_size': 0.5,
                    'significant': True
                }
            },
            'pharmacophore_analysis': {
                'hydrogen_bond_donors': {
                    'strong_mean': 3.0,
                    'weak_mean': 2.0,
                    'p_value': 0.05,
                    'significant': True
                }
            }
        }
        
        # Generate report
        report = self.analyzer.generate_substructure_report(results)
        
        # Verify report content
        assert 'Molecular Substructure Analysis Report' in report
        assert 'Strong binders threshold: -5.000' in report
        assert 'Number of strong binders: 10' in report
        assert 'Number of weak binders: 40' in report
        
        # Verify report file was created
        assert os.path.exists(os.path.join(self.temp_dir, 'substructure_analysis_report.md'))


class TestAttentionVisualizer:
    """Test cases for AttentionVisualizer."""
    
    def setup_method(self):
        """Set up test fixtures."""
        self.temp_dir = tempfile.mkdtemp()
        self.visualizer = AttentionVisualizer(output_dir=self.temp_dir)
    
    def teardown_method(self):
        """Clean up test fixtures."""
        shutil.rmtree(self.temp_dir, ignore_errors=True)
    
    @patch('tlr4_binding.ml_components.attention_visualizer.TORCH_AVAILABLE', True)
    @patch('tlr4_binding.ml_components.attention_visualizer.torch')
    def test_transformer_attention_visualization(self, mock_torch):
        """Test transformer attention visualization."""
        # Mock PyTorch functionality
        mock_torch.tensor.return_value = Mock()
        mock_torch.no_grad.return_value = Mock()
        
        # Create sample data
        input_data = pd.DataFrame(np.random.randn(5, 10))
        feature_names = [f'feature_{i}' for i in range(10)]
        model = Mock()
        
        # Run visualization
        results = self.visualizer.visualize_transformer_attention(
            model, input_data, feature_names, sample_indices=[0, 1]
        )
        
        # Verify results
        assert 'sample_0' in results
        assert 'sample_1' in results
        
        for sample_key, sample_results in results.items():
            assert 'attention_weights' in sample_results
            assert 'feature_importance' in sample_results
            assert 'sample_data' in sample_results
    
    @patch('tlr4_binding.ml_components.attention_visualizer.TORCH_AVAILABLE', True)
    @patch('tlr4_binding.ml_components.attention_visualizer.NETWORKX_AVAILABLE', True)
    def test_gnn_attention_visualization(self, mock_nx):
        """Test GNN attention visualization."""
        # Create sample graph data
        graph_data = [{
            'node_features': np.random.randn(5, 10),
            'edge_index': np.array([[0, 1, 2], [1, 2, 3]])
        } for _ in range(3)]
        
        node_features = [f'node_feature_{i}' for i in range(10)]
        model = Mock()
        
        # Run visualization
        results = self.visualizer.visualize_gnn_attention(
            model, graph_data, node_features, sample_indices=[0, 1]
        )
        
        # Verify results
        assert 'sample_0' in results
        assert 'sample_1' in results
        
        for sample_key, sample_results in results.items():
            assert 'attention_weights' in sample_results
            assert 'node_importance' in sample_results
            assert 'graph_data' in sample_results
    
    def test_attention_importance_calculation(self):
        """Test attention importance calculation."""
        # Create sample attention weights
        attention_weights = np.random.rand(8, 10, 10)  # 8 heads, 10 features
        feature_names = [f'feature_{i}' for i in range(10)]
        
        # Calculate importance
        importance = self.visualizer._calculate_attention_importance(
            attention_weights, feature_names
        )
        
        # Verify results
        assert len(importance) == len(feature_names)
        assert all(isinstance(v, (int, float)) for v in importance.values())
        assert abs(sum(importance.values()) - 1.0) < 1e-6  # Should sum to 1
    
    def test_attention_summary_creation(self):
        """Test attention summary creation."""
        # Create mock attention results
        attention_results = {
            'sample_0': {
                'feature_importance': {'feature_0': 0.3, 'feature_1': 0.2, 'feature_2': 0.5}
            },
            'sample_1': {
                'feature_importance': {'feature_0': 0.4, 'feature_1': 0.3, 'feature_2': 0.3}
            }
        }
        
        # Create summary
        summary = self.visualizer.create_attention_summary(attention_results)
        
        # Verify results
        assert 'total_samples' in summary
        assert 'feature_importance_ranking' in summary
        assert 'attention_patterns' in summary
        assert 'key_insights' in summary
        
        assert summary['total_samples'] == 2
        assert len(summary['feature_importance_ranking']) == 3
    
    def test_attention_report_generation(self):
        """Test attention report generation."""
        # Create mock attention results
        attention_results = {
            'sample_0': {
                'feature_importance': {'feature_0': 0.3, 'feature_1': 0.2, 'feature_2': 0.5}
            }
        }
        
        # Generate report
        report = self.visualizer.generate_attention_report(
            attention_results, model_type="transformer"
        )
        
        # Verify report content
        assert 'Attention Analysis Report' in report
        assert 'transformer' in report
        assert 'Feature Importance Ranking' in report
        
        # Verify report file was created
        assert os.path.exists(os.path.join(self.temp_dir, 'attention_analysis_report_transformer.md'))


def test_integration():
    """Integration test for the complete interpretability suite."""
    # This would be a more comprehensive integration test
    # that tests the full workflow end-to-end
    pass


if __name__ == "__main__":
    pytest.main([__file__])
```
