## Supplementary material for "Machine Learning-Driven Prediction of TLR4 Binding Affinity: A Comprehensive Molecular Feature Analysis for Drug Discovery": Codebase: test_interpretability_direct.py

```
#!/usr/bin/env python3
"""
Direct test for Model Interpretability and Analysis Suite

This script tests the interpretability tools by importing them directly
without going through the package __init__.py files.
"""

import sys
import os
import pandas as pd
import numpy as np
import matplotlib.pyplot as plt
import warnings
warnings.filterwarnings('ignore')

# Add src to path
sys.path.append('src')

# Import modules directly to avoid package-level imports
sys.path.insert(0, 'src/tlr4_binding/ml_components')

# Import the interpretability modules directly
from interpretability import ModelInterpretabilitySuite
from molecular_substructure_analyzer import MolecularSubstructureAnalyzer
from attention_visualizer import AttentionVisualizer

def test_interpretability_suite():
    """Test the interpretability suite with sample data."""
    print("Testing Model Interpretability Suite...")
    
    # Create sample data
    np.random.seed(42)
    n_samples = 50
    n_features = 5
    
    X_train = pd.DataFrame(
        np.random.randn(n_samples, n_features),
        columns=[f'feature_{i}' for i in range(n_features)]
    )
    y_train = pd.Series(np.random.randn(n_samples))
    
    X_test = pd.DataFrame(
        np.random.randn(20, n_features),
        columns=[f'feature_{i}' for i in range(n_features)]
    )
    y_test = pd.Series(np.random.randn(20))
    
    # Create mock models
    from sklearn.ensemble import RandomForestRegressor
    from sklearn.svm import SVR
    from sklearn.linear_model import LinearRegression
    from sklearn.preprocessing import StandardScaler
    
    # Scale features
    scaler = StandardScaler()
    X_train_scaled = scaler.fit_transform(X_train)
    X_test_scaled = scaler.transform(X_test)
    
    models = {
        'RandomForest': RandomForestRegressor(n_estimators=10, random_state=42),
        'SVR': SVR(kernel='rbf', C=1.0, gamma='scale'),
        'LinearRegression': LinearRegression()
    }
    
    # Train models
    models['RandomForest'].fit(X_train, y_train)
    models['SVR'].fit(X_train_scaled, y_train)
    models['LinearRegression'].fit(X_train_scaled, y_train)
    
    # Initialize interpretability suite
    suite = ModelInterpretabilitySuite(
        models=models,
        feature_names=[f'feature_{i}' for i in range(n_features)],
        X_train=X_train,
        y_train=y_train,
        X_test=X_test,
        y_test=y_test,
        output_dir="results/interpretability_test"
    )
    
    print("✓ Interpretability suite initialized successfully")
    
    # Test SHAP analysis
    print("Testing SHAP analysis...")
    try:
        results = suite.generate_shap_analysis('RandomForest', models['RandomForest'], sample_size=10)
        if results:
            print("✓ SHAP analysis completed successfully")
        else:
            print("⚠ SHAP analysis completed but returned empty results")
    except Exception as e:
        print(f"✗ SHAP analysis failed: {str(e)}")
    
    # Test LIME analysis
    print("Testing LIME analysis...")
    try:
        results = suite.generate_lime_analysis('RandomForest', models['RandomForest'], num_samples=2)
        if results:
            print("✓ LIME analysis completed successfully")
        else:
            print("⚠ LIME analysis completed but returned empty results")
    except Exception as e:
        print(f"✗ LIME analysis failed: {str(e)}")
    
    # Test comprehensive report generation
    print("Testing comprehensive report generation...")
    try:
        report = suite.generate_comprehensive_report()
        if report:
            print("✓ Comprehensive report generated successfully")
        else:
            print("⚠ Comprehensive report generated but returned empty results")
    except Exception as e:
        print(f"✗ Comprehensive report generation failed: {str(e)}")
    
    print("Interpretability suite test completed!")

def test_molecular_substructure_analyzer():
    """Test the molecular substructure analyzer."""
    print("\nTesting Molecular Substructure Analyzer...")
    
    # Create sample data
    np.random.seed(42)
    n_samples = 30
    
    compound_data = pd.DataFrame({
        'smiles': [f"CCO{i}" for i in range(n_samples)],
        'mol_weight': np.random.normal(100, 20, n_samples),
        'mol_logp': np.random.normal(2, 1, n_samples)
    })
    
    binding_affinities = pd.Series(np.random.normal(-5, 2, n_samples))
    
    # Initialize analyzer
    analyzer = MolecularSubstructureAnalyzer(output_dir="results/interpretability_test/substructures")
    
    print("✓ Molecular substructure analyzer initialized successfully")
    
    # Test analysis
    print("Testing molecular substructure analysis...")
    try:
        results = analyzer.analyze_binding_drivers(
            compound_data, binding_affinities, smiles_column='smiles', threshold_percentile=20
        )
        if results:
            print("✓ Molecular substructure analysis completed successfully")
            
            # Test report generation
            try:
                report = analyzer.generate_substructure_report(results)
                if report:
                    print("✓ Substructure report generated successfully")
                else:
                    print("⚠ Substructure report generated but returned empty results")
            except Exception as e:
                print(f"✗ Substructure report generation failed: {str(e)}")
        else:
            print("⚠ Molecular substructure analysis completed but returned empty results")
    except Exception as e:
        print(f"✗ Molecular substructure analysis failed: {str(e)}")
    
    print("Molecular substructure analyzer test completed!")

def test_attention_visualizer():
    """Test the attention visualizer."""
    print("\nTesting Attention Visualizer...")
    
    # Initialize visualizer
    visualizer = AttentionVisualizer(output_dir="results/interpretability_test/attention")
    
    print("✓ Attention visualizer initialized successfully")
    
    # Create mock attention results
    feature_names = [f'feature_{i}' for i in range(5)]
    attention_results = {
        'sample_0': {
            'attention_weights': np.random.rand(5, 5),
            'feature_importance': dict(zip(feature_names, np.random.rand(5)))
        },
        'sample_1': {
            'attention_weights': np.random.rand(5, 5),
            'feature_importance': dict(zip(feature_names, np.random.rand(5)))
        }
    }
    
    # Test attention summary creation
    print("Testing attention summary creation...")
    try:
        summary = visualizer.create_attention_summary(attention_results)
        if summary:
            print("✓ Attention summary created successfully")
        else:
            print("⚠ Attention summary created but returned empty results")
    except Exception as e:
        print(f"✗ Attention summary creation failed: {str(e)}")
    
    # Test attention report generation
    print("Testing attention report generation...")
    try:
        report = visualizer.generate_attention_report(attention_results, model_type="transformer")
        if report:
            print("✓ Attention report generated successfully")
        else:
            print("⚠ Attention report generated but returned empty results")
    except Exception as e:
        print(f"✗ Attention report generation failed: {str(e)}")
    
    print("Attention visualizer test completed!")

def main():
    """Main function to run all tests."""
    print("=" * 60)
    print("TLR4 Binding Prediction - Interpretability Suite Test")
    print("=" * 60)
    
    try:
        test_interpretability_suite()
        test_molecular_substructure_analyzer()
        test_attention_visualizer()
        
        print("\n" + "=" * 60)
        print("ALL TESTS COMPLETED SUCCESSFULLY!")
        print("=" * 60)
        print("Check the results/interpretability_test/ directory for generated files.")
        
    except Exception as e:
        print(f"\nError during testing: {str(e)}")
        import traceback
        traceback.print_exc()

if __name__ == "__main__":
    main()
```
