## Supplementary material for "Machine Learning-Driven Prediction of TLR4 Binding Affinity: A Comprehensive Molecular Feature Analysis for Drug Discovery": Codebase: test_interpretability_standalone_tests.py

```
#!/usr/bin/env python3
"""
Standalone tests for Model Interpretability and Analysis Suite

This script runs comprehensive tests for the interpretability tools
without importing the full package.
"""

import sys
import os
import pandas as pd
import numpy as np
import matplotlib.pyplot as plt
import warnings
warnings.filterwarnings('ignore')
import tempfile
import shutil
from unittest.mock import Mock, patch

# Add src to path
sys.path.append('src')

# Import modules directly to avoid package-level imports
sys.path.insert(0, 'src/tlr4_binding/ml_components')

# Import the interpretability modules directly
from interpretability import ModelInterpretabilitySuite
from molecular_substructure_analyzer import MolecularSubstructureAnalyzer
from attention_visualizer import AttentionVisualizer

def test_interpretability_suite():
    """Test the interpretability suite with sample data."""
    print("Testing Model Interpretability Suite...")
    
    # Create sample data
    np.random.seed(42)
    n_samples = 50
    n_features = 5
    
    X_train = pd.DataFrame(
        np.random.randn(n_samples, n_features),
        columns=[f'feature_{i}' for i in range(n_features)]
    )
    y_train = pd.Series(np.random.randn(n_samples))
    
    X_test = pd.DataFrame(
        np.random.randn(20, n_features),
        columns=[f'feature_{i}' for i in range(n_features)]
    )
    y_test = pd.Series(np.random.randn(20))
    
    # Create mock models
    from sklearn.ensemble import RandomForestRegressor
    from sklearn.svm import SVR
    from sklearn.linear_model import LinearRegression
    from sklearn.preprocessing import StandardScaler
    
    # Scale features
    scaler = StandardScaler()
    X_train_scaled = scaler.fit_transform(X_train)
    X_test_scaled = scaler.transform(X_test)
    
    models = {
        'RandomForest': RandomForestRegressor(n_estimators=10, random_state=42),
        'SVR': SVR(kernel='rbf', C=1.0, gamma='scale'),
        'LinearRegression': LinearRegression()
    }
    
    # Train models
    models['RandomForest'].fit(X_train, y_train)
    models['SVR'].fit(X_train_scaled, y_train)
    models['LinearRegression'].fit(X_train_scaled, y_train)
    
    # Create temporary output directory
    temp_dir = tempfile.mkdtemp()
    
    try:
        # Initialize interpretability suite
        suite = ModelInterpretabilitySuite(
            models=models,
            feature_names=[f'feature_{i}' for i in range(n_features)],
            X_train=X_train,
            y_train=y_train,
            X_test=X_test,
            y_test=y_test,
            output_dir=temp_dir
        )
        
        # Test initialization
        assert len(suite.models) == 3
        assert len(suite.feature_names) == n_features
        assert len(suite.X_train) == n_samples
        assert os.path.exists(temp_dir)
        print("✓ Interpretability suite initialization test passed")
        
        # Test SHAP analysis
        print("Testing SHAP analysis...")
        try:
            results = suite.generate_shap_analysis('RandomForest', models['RandomForest'], sample_size=10)
            assert isinstance(results, dict)
            print("✓ SHAP analysis test passed")
        except Exception as e:
            print(f"⚠ SHAP analysis test failed (expected due to missing dependencies): {str(e)}")
        
        # Test LIME analysis
        print("Testing LIME analysis...")
        try:
            results = suite.generate_lime_analysis('RandomForest', models['RandomForest'], num_samples=2)
            assert isinstance(results, dict)
            print("✓ LIME analysis test passed")
        except Exception as e:
            print(f"⚠ LIME analysis test failed (expected due to missing dependencies): {str(e)}")
        
        # Test comprehensive report generation
        print("Testing comprehensive report generation...")
        try:
            report = suite.generate_comprehensive_report()
            assert isinstance(report, dict)
            assert 'timestamp' in report
            assert 'models_analyzed' in report
            print("✓ Comprehensive report generation test passed")
        except Exception as e:
            print(f"✗ Comprehensive report generation test failed: {str(e)}")
        
    finally:
        # Clean up
        shutil.rmtree(temp_dir, ignore_errors=True)
    
    print("✓ Model Interpretability Suite tests completed!")

def test_molecular_substructure_analyzer():
    """Test the molecular substructure analyzer."""
    print("\nTesting Molecular Substructure Analyzer...")
    
    # Create sample data with valid SMILES
    np.random.seed(42)
    n_samples = 30
    
    compound_data = pd.DataFrame({
        'smiles': ['CCO', 'CCCO', 'CCCCCO'] * 10,  # Valid SMILES
        'mol_weight': np.random.normal(100, 20, n_samples),
        'mol_logp': np.random.normal(2, 1, n_samples)
    })
    
    binding_affinities = pd.Series(np.random.normal(-5, 2, n_samples))
    
    # Create temporary output directory
    temp_dir = tempfile.mkdtemp()
    
    try:
        # Initialize analyzer
        analyzer = MolecularSubstructureAnalyzer(output_dir=temp_dir)
        
        # Test initialization
        assert os.path.exists(temp_dir)
        print("✓ Molecular substructure analyzer initialization test passed")
        
        # Test analysis
        print("Testing molecular substructure analysis...")
        try:
            results = analyzer.analyze_binding_drivers(
                compound_data, binding_affinities, smiles_column='smiles', threshold_percentile=20
            )
            assert isinstance(results, dict)
            assert 'threshold_value' in results
            assert 'strong_binders_count' in results
            assert 'weak_binders_count' in results
            print("✓ Molecular substructure analysis test passed")
            
            # Test report generation
            try:
                report = analyzer.generate_substructure_report(results)
                assert isinstance(report, str)
                assert 'Molecular Substructure Analysis Report' in report
                print("✓ Substructure report generation test passed")
            except Exception as e:
                print(f"⚠ Substructure report generation test failed: {str(e)}")
                
        except Exception as e:
            print(f"⚠ Molecular substructure analysis test failed (expected due to missing RDKit): {str(e)}")
        
    finally:
        # Clean up
        shutil.rmtree(temp_dir, ignore_errors=True)
    
    print("✓ Molecular Substructure Analyzer tests completed!")

def test_attention_visualizer():
    """Test the attention visualizer."""
    print("\nTesting Attention Visualizer...")
    
    # Create temporary output directory
    temp_dir = tempfile.mkdtemp()
    
    try:
        # Initialize visualizer
        visualizer = AttentionVisualizer(output_dir=temp_dir)
        
        # Test initialization
        assert os.path.exists(temp_dir)
        print("✓ Attention visualizer initialization test passed")
        
        # Create mock attention results
        feature_names = [f'feature_{i}' for i in range(5)]
        attention_results = {
            'sample_0': {
                'attention_weights': np.random.rand(5, 5),
                'feature_importance': dict(zip(feature_names, np.random.rand(5)))
            },
            'sample_1': {
                'attention_weights': np.random.rand(5, 5),
                'feature_importance': dict(zip(feature_names, np.random.rand(5)))
            }
        }
        
        # Test attention summary creation
        print("Testing attention summary creation...")
        try:
            summary = visualizer.create_attention_summary(attention_results)
            assert isinstance(summary, dict)
            assert 'total_samples' in summary
            assert 'feature_importance_ranking' in summary
            print("✓ Attention summary creation test passed")
        except Exception as e:
            print(f"✗ Attention summary creation test failed: {str(e)}")
        
        # Test attention report generation
        print("Testing attention report generation...")
        try:
            report = visualizer.generate_attention_report(attention_results, model_type="transformer")
            assert isinstance(report, str)
            assert 'Attention Analysis Report' in report
            print("✓ Attention report generation test passed")
        except Exception as e:
            print(f"✗ Attention report generation test failed: {str(e)}")
        
    finally:
        # Clean up
        shutil.rmtree(temp_dir, ignore_errors=True)
    
    print("✓ Attention Visualizer tests completed!")

def test_feature_importance_calculation():
    """Test feature importance calculation methods."""
    print("\nTesting Feature Importance Calculation...")
    
    # Create sample data
    np.random.seed(42)
    n_samples = 50
    n_features = 5
    
    X_train = pd.DataFrame(
        np.random.randn(n_samples, n_features),
        columns=[f'feature_{i}' for i in range(n_features)]
    )
    y_train = pd.Series(np.random.randn(n_samples))
    
    X_test = pd.DataFrame(
        np.random.randn(20, n_features),
        columns=[f'feature_{i}' for i in range(n_features)]
    )
    y_test = pd.Series(np.random.randn(20))
    
    # Create mock models
    from sklearn.ensemble import RandomForestRegressor
    models = {'RandomForest': RandomForestRegressor(n_estimators=10, random_state=42)}
    models['RandomForest'].fit(X_train, y_train)
    
    # Create temporary output directory
    temp_dir = tempfile.mkdtemp()
    
    try:
        # Initialize interpretability suite
        suite = ModelInterpretabilitySuite(
            models=models,
            feature_names=[f'feature_{i}' for i in range(n_features)],
            X_train=X_train,
            y_train=y_train,
            X_test=X_test,
            y_test=y_test,
            output_dir=temp_dir
        )
        
        # Test SHAP importance calculation
        print("Testing SHAP importance calculation...")
        try:
            shap_values = np.random.randn(10, n_features)
            importance = suite._calculate_shap_importance(shap_values)
            assert len(importance) == n_features
            assert all(isinstance(v, (int, float)) for v in importance.values())
            assert all(v >= 0 for v in importance.values())
            print("✓ SHAP importance calculation test passed")
        except Exception as e:
            print(f"✗ SHAP importance calculation test failed: {str(e)}")
        
    finally:
        # Clean up
        shutil.rmtree(temp_dir, ignore_errors=True)
    
    print("✓ Feature Importance Calculation tests completed!")

def run_all_tests():
    """Run all tests."""
    print("=" * 60)
    print("TLR4 Binding Prediction - Interpretability Suite Tests")
    print("=" * 60)
    
    test_results = []
    
    try:
        test_interpretability_suite()
        test_results.append("✓ Model Interpretability Suite")
    except Exception as e:
        print(f"✗ Model Interpretability Suite tests failed: {str(e)}")
        test_results.append("✗ Model Interpretability Suite")
    
    try:
        test_molecular_substructure_analyzer()
        test_results.append("✓ Molecular Substructure Analyzer")
    except Exception as e:
        print(f"✗ Molecular Substructure Analyzer tests failed: {str(e)}")
        test_results.append("✗ Molecular Substructure Analyzer")
    
    try:
        test_attention_visualizer()
        test_results.append("✓ Attention Visualizer")
    except Exception as e:
        print(f"✗ Attention Visualizer tests failed: {str(e)}")
        test_results.append("✗ Attention Visualizer")
    
    try:
        test_feature_importance_calculation()
        test_results.append("✓ Feature Importance Calculation")
    except Exception as e:
        print(f"✗ Feature Importance Calculation tests failed: {str(e)}")
        test_results.append("✗ Feature Importance Calculation")
    
    print("\n" + "=" * 60)
    print("TEST RESULTS SUMMARY")
    print("=" * 60)
    for result in test_results:
        print(result)
    
    passed_tests = sum(1 for result in test_results if result.startswith("✓"))
    total_tests = len(test_results)
    
    print(f"\nPassed: {passed_tests}/{total_tests} tests")
    
    if passed_tests == total_tests:
        print("🎉 ALL TESTS PASSED!")
    else:
        print("⚠ Some tests failed or had warnings (expected due to missing optional dependencies)")

if __name__ == "__main__":
    run_all_tests()
```
