## Supplementary material for "Machine Learning-Driven Prediction of TLR4 Binding Affinity: A Comprehensive Molecular Feature Analysis for Drug Discovery": Codebase: test_ml_models.py

```
"""
Comprehensive tests for traditional ML models baseline.

This module tests all traditional ML models including Random Forest,
SVR, XGBoost, and LightGBM with hyperparameter optimization,
feature importance extraction, and performance evaluation.
"""

import unittest
import sys
import tempfile
import shutil
from pathlib import Path
import pandas as pd
import numpy as np
from unittest.mock import patch, MagicMock

# Add src to path
sys.path.insert(0, str(Path(__file__).parent.parent / "src"))

from tlr4_binding.ml_components.trainer import (
    ModelTrainerInterface,
    RandomForestTrainer,
    SVRTrainer,
    XGBoostTrainer,
    LightGBMTrainer,
    MLModelTrainer
)


class TestModelTrainerInterface(unittest.TestCase):
    """Test the abstract ModelTrainerInterface."""
    
    def test_interface_abstract_methods(self):
        """Test that interface defines required abstract methods."""
        # Check that interface has required abstract methods
        abstract_methods = ModelTrainerInterface.__abstractmethods__
        self.assertIn('train', abstract_methods)
        self.assertIn('predict', abstract_methods)
        self.assertIn('get_feature_importance', abstract_methods)


class TestRandomForestTrainer(unittest.TestCase):
    """Test RandomForestTrainer functionality."""
    
    def setUp(self):
        """Set up test data."""
        np.random.seed(42)
        self.n_samples = 100
        self.n_features = 10
        
        # Create synthetic data
        X = np.random.randn(self.n_samples, self.n_features)
        y = np.random.randn(self.n_samples)
        
        self.X_train = pd.DataFrame(X, columns=[f'feature_{i}' for i in range(self.n_features)])
        self.y_train = pd.Series(y)
        
        self.trainer = RandomForestTrainer(n_jobs=1, random_state=42)
    
    def test_initialization(self):
        """Test RandomForestTrainer initialization."""
        self.assertEqual(self.trainer.n_jobs, 1)
        self.assertEqual(self.trainer.random_state, 42)
        self.assertIsInstance(self.trainer.param_grid, dict)
        self.assertIn('n_estimators', self.trainer.param_grid)
        self.assertIn('max_depth', self.trainer.param_grid)
    
    def test_train(self):
        """Test Random Forest model training."""
        model = self.trainer.train(self.X_train, self.y_train)
        
        # Check model is trained
        self.assertIsNotNone(model)
        self.assertTrue(hasattr(model, 'predict'))
        self.assertTrue(hasattr(model, 'feature_importances_'))
        
        # Check predictions are reasonable
        predictions = self.trainer.predict(model, self.X_train)
        self.assertEqual(len(predictions), len(self.y_train))
        self.assertIsInstance(predictions, np.ndarray)
    
    def test_predict(self):
        """Test prediction functionality."""
        model = self.trainer.train(self.X_train, self.y_train)
        predictions = self.trainer.predict(model, self.X_train)
        
        self.assertEqual(len(predictions), len(self.y_train))
        self.assertIsInstance(predictions, np.ndarray)
        self.assertFalse(np.any(np.isnan(predictions)))
    
    def test_get_feature_importance(self):
        """Test feature importance extraction."""
        model = self.trainer.train(self.X_train, self.y_train)
        importance = self.trainer.get_feature_importance(model)
        
        self.assertIsInstance(importance, dict)
        self.assertEqual(len(importance), self.n_features)
        
        # Check importance values are non-negative
        for feature, imp in importance.items():
            self.assertGreaterEqual(imp, 0)
            self.assertIn(feature, self.X_train.columns)


class TestSVRTrainer(unittest.TestCase):
    """Test SVRTrainer functionality."""
    
    def setUp(self):
        """Set up test data."""
        np.random.seed(42)
        self.n_samples = 100
        self.n_features = 10
        
        # Create synthetic data
        X = np.random.randn(self.n_samples, self.n_features)
        y = np.random.randn(self.n_samples)
        
        self.X_train = pd.DataFrame(X, columns=[f'feature_{i}' for i in range(self.n_features)])
        self.y_train = pd.Series(y)
        
        self.trainer = SVRTrainer(random_state=42)
    
    def test_initialization(self):
        """Test SVRTrainer initialization."""
        self.assertEqual(self.trainer.random_state, 42)
        self.assertIsInstance(self.trainer.param_grid, dict)
        self.assertIn('C', self.trainer.param_grid)
        self.assertIn('gamma', self.trainer.param_grid)
        self.assertIn('kernel', self.trainer.param_grid)
    
    def test_train(self):
        """Test SVR model training."""
        model = self.trainer.train(self.X_train, self.y_train)
        
        # Check model is trained
        self.assertIsNotNone(model)
        self.assertTrue(hasattr(model, 'predict'))
        
        # Check predictions are reasonable
        predictions = self.trainer.predict(model, self.X_train)
        self.assertEqual(len(predictions), len(self.y_train))
        self.assertIsInstance(predictions, np.ndarray)
    
    def test_predict(self):
        """Test prediction functionality."""
        model = self.trainer.train(self.X_train, self.y_train)
        predictions = self.trainer.predict(model, self.X_train)
        
        self.assertEqual(len(predictions), len(self.y_train))
        self.assertIsInstance(predictions, np.ndarray)
        self.assertFalse(np.any(np.isnan(predictions)))
    
    def test_get_feature_importance(self):
        """Test feature importance extraction (SVR returns empty dict)."""
        model = self.trainer.train(self.X_train, self.y_train)
        importance = self.trainer.get_feature_importance(model)
        
        # SVR doesn't have direct feature importance
        self.assertIsInstance(importance, dict)
        self.assertEqual(len(importance), 0)


class TestXGBoostTrainer(unittest.TestCase):
    """Test XGBoostTrainer functionality."""
    
    def setUp(self):
        """Set up test data."""
        np.random.seed(42)
        self.n_samples = 100
        self.n_features = 10
        
        # Create synthetic data
        X = np.random.randn(self.n_samples, self.n_features)
        y = np.random.randn(self.n_samples)
        
        self.X_train = pd.DataFrame(X, columns=[f'feature_{i}' for i in range(self.n_features)])
        self.y_train = pd.Series(y)
        
        self.trainer = XGBoostTrainer(random_state=42)
    
    def test_initialization(self):
        """Test XGBoostTrainer initialization."""
        self.assertEqual(self.trainer.random_state, 42)
        self.assertIsInstance(self.trainer.param_grid, dict)
        self.assertIn('n_estimators', self.trainer.param_grid)
        self.assertIn('max_depth', self.trainer.param_grid)
        self.assertIn('learning_rate', self.trainer.param_grid)
    
    def test_train(self):
        """Test XGBoost model training."""
        model = self.trainer.train(self.X_train, self.y_train)
        
        # Check model is trained
        self.assertIsNotNone(model)
        self.assertTrue(hasattr(model, 'predict'))
        self.assertTrue(hasattr(model, 'feature_importances_'))
        
        # Check predictions are reasonable
        predictions = self.trainer.predict(model, self.X_train)
        self.assertEqual(len(predictions), len(self.y_train))
        self.assertIsInstance(predictions, np.ndarray)
    
    def test_predict(self):
        """Test prediction functionality."""
        model = self.trainer.train(self.X_train, self.y_train)
        predictions = self.trainer.predict(model, self.X_train)
        
        self.assertEqual(len(predictions), len(self.y_train))
        self.assertIsInstance(predictions, np.ndarray)
        self.assertFalse(np.any(np.isnan(predictions)))
    
    def test_get_feature_importance(self):
        """Test feature importance extraction."""
        model = self.trainer.train(self.X_train, self.y_train)
        importance = self.trainer.get_feature_importance(model)
        
        self.assertIsInstance(importance, dict)
        self.assertEqual(len(importance), self.n_features)
        
        # Check importance values are non-negative
        for feature, imp in importance.items():
            self.assertGreaterEqual(imp, 0)
            self.assertIn(feature, self.X_train.columns)


class TestLightGBMTrainer(unittest.TestCase):
    """Test LightGBMTrainer functionality."""
    
    def setUp(self):
        """Set up test data."""
        np.random.seed(42)
        self.n_samples = 100
        self.n_features = 10
        
        # Create synthetic data
        X = np.random.randn(self.n_samples, self.n_features)
        y = np.random.randn(self.n_samples)
        
        self.X_train = pd.DataFrame(X, columns=[f'feature_{i}' for i in range(self.n_features)])
        self.y_train = pd.Series(y)
        
        self.trainer = LightGBMTrainer(random_state=42)
    
    def test_initialization(self):
        """Test LightGBMTrainer initialization."""
        self.assertEqual(self.trainer.random_state, 42)
        self.assertIsInstance(self.trainer.param_grid, dict)
        self.assertIn('n_estimators', self.trainer.param_grid)
        self.assertIn('max_depth', self.trainer.param_grid)
        self.assertIn('learning_rate', self.trainer.param_grid)
        self.assertIn('num_leaves', self.trainer.param_grid)
    
    def test_train(self):
        """Test LightGBM model training."""
        model = self.trainer.train(self.X_train, self.y_train)
        
        # Check model is trained
        self.assertIsNotNone(model)
        self.assertTrue(hasattr(model, 'predict'))
        self.assertTrue(hasattr(model, 'feature_importances_'))
        
        # Check predictions are reasonable
        predictions = self.trainer.predict(model, self.X_train)
        self.assertEqual(len(predictions), len(self.y_train))
        self.assertIsInstance(predictions, np.ndarray)
    
    def test_predict(self):
        """Test prediction functionality."""
        model = self.trainer.train(self.X_train, self.y_train)
        predictions = self.trainer.predict(model, self.X_train)
        
        self.assertEqual(len(predictions), len(self.y_train))
        self.assertIsInstance(predictions, np.ndarray)
        self.assertFalse(np.any(np.isnan(predictions)))
    
    def test_get_feature_importance(self):
        """Test feature importance extraction."""
        model = self.trainer.train(self.X_train, self.y_train)
        importance = self.trainer.get_feature_importance(model)
        
        self.assertIsInstance(importance, dict)
        self.assertEqual(len(importance), self.n_features)
        
        # Check importance values are non-negative
        for feature, imp in importance.items():
            self.assertGreaterEqual(imp, 0)
            self.assertIn(feature, self.X_train.columns)


class TestMLModelTrainer(unittest.TestCase):
    """Test MLModelTrainer coordinator functionality."""
    
    def setUp(self):
        """Set up test data and temporary directory."""
        np.random.seed(42)
        self.n_samples = 100
        self.n_features = 10
        
        # Create synthetic data
        X = np.random.randn(self.n_samples, self.n_features)
        y = np.random.randn(self.n_samples)
        
        self.X_train = pd.DataFrame(X, columns=[f'feature_{i}' for i in range(self.n_features)])
        self.y_train = pd.Series(y)
        
        # Create temporary directory for models
        self.temp_dir = tempfile.mkdtemp()
        self.trainer = MLModelTrainer(models_dir=self.temp_dir)
    
    def tearDown(self):
        """Clean up temporary directory."""
        shutil.rmtree(self.temp_dir)
    
    def test_initialization(self):
        """Test MLModelTrainer initialization."""
        self.assertIsInstance(self.trainer.models_dir, Path)
        self.assertIsInstance(self.trainer.trainers, dict)
        self.assertIsInstance(self.trainer.trained_models, dict)
        self.assertIsInstance(self.trainer.training_results, dict)
    
    def test_train_models(self):
        """Test training all available models."""
        trained_models = self.trainer.train_models(self.X_train, self.y_train)
        
        # Check that some models were trained
        self.assertGreater(len(trained_models), 0)
        self.assertEqual(len(trained_models), len(self.trainer.trained_models))
        
        # Check each trained model
        for model_name, model in trained_models.items():
            self.assertIsNotNone(model)
            self.assertTrue(hasattr(model, 'predict'))
    
    def test_evaluate_models(self):
        """Test model evaluation on test set."""
        # First train models
        self.trainer.train_models(self.X_train, self.y_train)
        
        # Create test data
        X_test = self.X_train.iloc[:20]  # Use subset as test
        y_test = self.y_train.iloc[:20]
        
        # Evaluate models
        results = self.trainer.evaluate_models(X_test, y_test)
        
        # Check evaluation results
        self.assertIsInstance(results, dict)
        self.assertEqual(len(results), len(self.trainer.trained_models))
        
        for model_name, result in results.items():
            self.assertIn('predictions', result)
            self.assertIn('metrics', result)
            self.assertIn('feature_importance', result)
            
            # Check predictions
            predictions = result['predictions']
            self.assertEqual(len(predictions), len(y_test))
            self.assertIsInstance(predictions, np.ndarray)
            
            # Check metrics
            metrics = result['metrics']
            self.assertIn('mse', metrics)
            self.assertIn('rmse', metrics)
            self.assertIn('mae', metrics)
            self.assertIn('r2', metrics)
    
    def test_get_best_model(self):
        """Test getting best model based on metric."""
        # First train models
        self.trainer.train_models(self.X_train, self.y_train)
        
        # Get best model
        best_name, best_model = self.trainer.get_best_model('r2')
        
        self.assertIsInstance(best_name, str)
        self.assertIsNotNone(best_model)
        self.assertIn(best_name, self.trainer.trained_models)
    
    def test_calculate_metrics(self):
        """Test metrics calculation."""
        y_true = pd.Series([1, 2, 3, 4, 5])
        y_pred = np.array([1.1, 2.1, 2.9, 4.1, 4.9])
        
        metrics = self.trainer._calculate_metrics(y_true, y_pred)
        
        self.assertIn('mse', metrics)
        self.assertIn('rmse', metrics)
        self.assertIn('mae', metrics)
        self.assertIn('r2', metrics)
        
        # Check metric values are reasonable
        self.assertGreater(metrics['r2'], 0.9)  # Should be high correlation
        self.assertGreater(metrics['mse'], 0)  # Should be positive
    
    def test_save_and_load_model(self):
        """Test model saving and loading."""
        # Train a model first
        self.trainer.train_models(self.X_train, self.y_train)
        
        if self.trainer.trained_models:
            model_name = list(self.trainer.trained_models.keys())[0]
            original_model = self.trainer.trained_models[model_name]
            
            # Load the model
            loaded_model = self.trainer.load_model(model_name)
            
            # Check that loaded model works
            predictions_original = original_model.predict(self.X_train.iloc[:5])
            predictions_loaded = loaded_model.predict(self.X_train.iloc[:5])
            
            np.testing.assert_array_almost_equal(predictions_original, predictions_loaded)
    
    def test_get_model_summary(self):
        """Test getting model summary."""
        # First train models
        self.trainer.train_models(self.X_train, self.y_train)
        
        # Get summary
        summary = self.trainer.get_model_summary()
        
        self.assertIsInstance(summary, pd.DataFrame)
        self.assertGreater(len(summary), 0)
        
        # Check summary columns
        expected_columns = ['model_name', 'mse', 'rmse', 'mae', 'r2']
        for col in expected_columns:
            self.assertIn(col, summary.columns)


class TestMLModelIntegration(unittest.TestCase):
    """Integration tests for ML model pipeline."""
    
    def setUp(self):
        """Set up test data."""
        np.random.seed(42)
        self.n_samples = 200
        self.n_features = 15
        
        # Create more realistic synthetic data
        X = np.random.randn(self.n_samples, self.n_features)
        # Create target with some structure
        y = (X[:, 0] * 2 + X[:, 1] * 1.5 + np.random.randn(self.n_samples) * 0.1)
        
        self.X = pd.DataFrame(X, columns=[f'feature_{i}' for i in range(self.n_features)])
        self.y = pd.Series(y)
        
        # Split data
        split_idx = int(0.7 * self.n_samples)
        self.X_train = self.X.iloc[:split_idx]
        self.y_train = self.y.iloc[:split_idx]
        self.X_test = self.X.iloc[split_idx:]
        self.y_test = self.y.iloc[split_idx:]
    
    def test_full_training_pipeline(self):
        """Test complete training and evaluation pipeline."""
        temp_dir = tempfile.mkdtemp()
        try:
            trainer = MLModelTrainer(models_dir=temp_dir)
            
            # Train models
            trained_models = trainer.train_models(self.X_train, self.y_train)
            self.assertGreater(len(trained_models), 0)
            
            # Evaluate models
            results = trainer.evaluate_models(self.X_test, self.y_test)
            self.assertEqual(len(results), len(trained_models))
            
            # Check that all models have reasonable performance
            for model_name, result in results.items():
                r2 = result['metrics']['r2']
                # Should have some predictive power
                self.assertGreater(r2, -1.0)  # Allow for some negative R²
                
        finally:
            shutil.rmtree(temp_dir)
    
    def test_model_comparison(self):
        """Test comparing different model performances."""
        temp_dir = tempfile.mkdtemp()
        try:
            trainer = MLModelTrainer(models_dir=temp_dir)
            
            # Train and evaluate
            trainer.train_models(self.X_train, self.y_train)
            results = trainer.evaluate_models(self.X_test, self.y_test)
            
            # Get model summary
            summary = trainer.get_model_summary()
            
            # Check that summary contains performance metrics
            self.assertIn('r2', summary.columns)
            self.assertIn('rmse', summary.columns)
            
            # Check that we can identify best model
            best_name, best_model = trainer.get_best_model('r2')
            self.assertIsNotNone(best_model)
            
        finally:
            shutil.rmtree(temp_dir)


if __name__ == '__main__':
    # Run tests with verbose output
    unittest.main(verbosity=2)
```
