## Supplementary material for "Machine Learning-Driven Prediction of TLR4 Binding Affinity: A Comprehensive Molecular Feature Analysis for Drug Discovery": Codebase: test_pdbqt_format.py

```
#!/usr/bin/env python3
"""
Test PDBQT format parsing.
"""

def create_proper_pdbqt():
    """Create a proper PDBQT file format."""
    content = """REMARK  VINA RESULT:    -6.5      0.000      0.000
ATOM      1  C   UNL     1      12.345   23.456   34.567  1.00  0.00     0.000 C
ATOM      2  N   UNL     1      13.345   24.456   35.567  1.00  0.00     0.000 N
ATOM      3  O   UNL     1      14.345   25.456   36.567  1.00  0.00     0.000 O
"""
    
    with open("test_proper.pdbqt", 'w') as f:
        f.write(content)
    
    # Test parsing
    with open("test_proper.pdbqt", 'r') as f:
        for i, line in enumerate(f):
            print(f"Line {i}: '{line.rstrip()}' (length: {len(line.rstrip())})")
            if line.startswith('ATOM'):
                print(f"  X: '{line[30:38]}' -> '{line[30:38].strip()}'")
                print(f"  Y: '{line[38:46]}' -> '{line[38:46].strip()}'")
                print(f"  Z: '{line[46:54]}' -> '{line[46:54].strip()}'")
                print(f"  Atom type: '{line[77:79]}' -> '{line[77:79].strip()}'")

if __name__ == "__main__":
    create_proper_pdbqt()
```
