## Supplementary material for "Machine Learning-Driven Prediction of TLR4 Binding Affinity: A Comprehensive Molecular Feature Analysis for Drug Discovery": Codebase: test_pipeline_orchestrator.py

```
#!/usr/bin/env python3
"""
Test suite for Research Pipeline Orchestrator

This script tests the pipeline orchestrator functionality including
MLflow integration, hyperparameter optimization, and report generation.
"""

import sys
import os
import unittest
import tempfile
import shutil
import numpy as np
import pandas as pd
from pathlib import Path

# Add src to path
sys.path.append(str(Path(__file__).parent.parent / "src"))

from tlr4_binding.ml_components.pipeline_orchestrator import (
    ResearchPipelineOrchestrator, 
    ExperimentConfig,
    create_experiment_config
)

class TestPipelineOrchestrator(unittest.TestCase):
    """Test cases for the Research Pipeline Orchestrator"""
    
    def setUp(self):
        """Set up test fixtures"""
        self.temp_dir = tempfile.mkdtemp()
        self.test_data = self._create_test_data()
        
    def tearDown(self):
        """Clean up test fixtures"""
        shutil.rmtree(self.temp_dir, ignore_errors=True)
    
    def _create_test_data(self, n_samples=50, n_features=10):
        """Create test data"""
        np.random.seed(42)
        X = pd.DataFrame(
            np.random.randn(n_samples, n_features),
            columns=[f'feature_{i}' for i in range(n_features)]
        )
        y = np.random.uniform(-10, 0, n_samples)
        return X, y
    
    def test_experiment_config_creation(self):
        """Test experiment configuration creation"""
        config = create_experiment_config(
            experiment_name="test_experiment",
            description="Test experiment",
            data_path="./data",
            output_path="./results"
        )
        
        self.assertEqual(config.experiment_name, "test_experiment")
        self.assertEqual(config.description, "Test experiment")
        self.assertTrue(config.enable_mlflow)
        self.assertTrue(config.enable_optuna)
    
    def test_orchestrator_initialization(self):
        """Test orchestrator initialization"""
        config = create_experiment_config(
            experiment_name="test_init",
            description="Test initialization",
            data_path="./data",
            output_path=self.temp_dir,
            enable_mlflow=False,  # Disable for testing
            enable_optuna=False
        )
        
        orchestrator = ResearchPipelineOrchestrator(config)
        self.assertIsNotNone(orchestrator)
        self.assertEqual(orchestrator.config.experiment_name, "test_init")
    
    def test_data_splitting(self):
        """Test data splitting functionality"""
        config = create_experiment_config(
            experiment_name="test_splitting",
            description="Test data splitting",
            data_path="./data",
            output_path=self.temp_dir,
            enable_mlflow=False,
            enable_optuna=False
        )
        
        orchestrator = ResearchPipelineOrchestrator(config)
        X, y = self.test_data
        
        splits = orchestrator._split_data(X, y)
        
        # Check that all splits exist
        expected_keys = ['X_train', 'X_val', 'X_test', 'y_train', 'y_val', 'y_test']
        for key in expected_keys:
            self.assertIn(key, splits)
        
        # Check data shapes
        total_samples = len(X)
        train_samples = len(splits['X_train'])
        val_samples = len(splits['X_val'])
        test_samples = len(splits['X_test'])
        
        self.assertEqual(train_samples + val_samples + test_samples, total_samples)
        self.assertGreater(train_samples, 0)
        self.assertGreater(val_samples, 0)
        self.assertGreater(test_samples, 0)
    
    def test_hyperparameter_sampling(self):
        """Test hyperparameter sampling for different models"""
        config = create_experiment_config(
            experiment_name="test_sampling",
            description="Test hyperparameter sampling",
            data_path="./data",
            output_path=self.temp_dir,
            enable_mlflow=False,
            enable_optuna=False
        )
        
        orchestrator = ResearchPipelineOrchestrator(config)
        
        # Test different model types
        models = ['random_forest', 'xgboost', 'neural_network']
        
        for model_name in models:
            # Create a mock trial object
            class MockTrial:
                def suggest_int(self, name, low, high):
                    return np.random.randint(low, high)
                
                def suggest_float(self, name, low, high):
                    return np.random.uniform(low, high)
                
                def suggest_categorical(self, name, choices):
                    # Handle the case where choices contains tuples (for hidden_layer_sizes)
                    if name == 'hidden_layer_sizes':
                        return choices[0]  # Return first choice
                    return np.random.choice(choices)
            
            trial = MockTrial()
            params = orchestrator._sample_hyperparameters(trial, model_name)
            
            self.assertIsInstance(params, dict)
            self.assertGreater(len(params), 0)
    
    def test_model_creation(self):
        """Test model creation for different types"""
        config = create_experiment_config(
            experiment_name="test_models",
            description="Test model creation",
            data_path="./data",
            output_path=self.temp_dir,
            enable_mlflow=False,
            enable_optuna=False
        )
        
        orchestrator = ResearchPipelineOrchestrator(config)
        
        # Test model creation
        models = ['random_forest', 'xgboost', 'neural_network']
        
        for model_name in models:
            try:
                model = orchestrator._create_model(model_name, {})
                self.assertIsNotNone(model)
            except Exception as e:
                # Some models might not be available in test environment
                print(f"Warning: Could not create {model_name}: {e}")
    
    def test_metrics_calculation(self):
        """Test metrics calculation"""
        config = create_experiment_config(
            experiment_name="test_metrics",
            description="Test metrics calculation",
            data_path="./data",
            output_path=self.temp_dir,
            enable_mlflow=False,
            enable_optuna=False
        )
        
        orchestrator = ResearchPipelineOrchestrator(config)
        
        # Create test data
        y_true = np.array([1, 2, 3, 4, 5])
        y_pred = np.array([1.1, 1.9, 3.1, 3.9, 5.1])
        
        metrics = orchestrator._calculate_metrics(y_true, y_pred)
        
        # Check that all expected metrics are present
        expected_metrics = ['r2_score', 'rmse', 'mae', 'spearman_corr', 'mse']
        for metric in expected_metrics:
            self.assertIn(metric, metrics)
            self.assertIsInstance(metrics[metric], (int, float))
    
    def test_config_save_load(self):
        """Test configuration save and load"""
        config = create_experiment_config(
            experiment_name="test_save_load",
            description="Test save/load",
            data_path="./data",
            output_path=self.temp_dir
        )
        
        # Save config
        config_path = os.path.join(self.temp_dir, "test_config.yaml")
        orchestrator = ResearchPipelineOrchestrator(config)
        orchestrator.save_config(config_path)
        
        # Load config
        loaded_orchestrator = ResearchPipelineOrchestrator.load_config(config_path)
        
        self.assertEqual(loaded_orchestrator.config.experiment_name, config.experiment_name)
        self.assertEqual(loaded_orchestrator.config.description, config.description)
    
    def test_markdown_report_generation(self):
        """Test markdown report generation"""
        config = create_experiment_config(
            experiment_name="test_report",
            description="Test report generation",
            data_path="./data",
            output_path=self.temp_dir,
            enable_mlflow=False,
            enable_optuna=False
        )
        
        orchestrator = ResearchPipelineOrchestrator(config)
        
        # Add some mock results
        orchestrator.results = {
            'model_evaluation': {
                'random_forest': {
                    'metrics': {
                        'r2_score': 0.85,
                        'rmse': 1.2,
                        'mae': 0.9,
                        'spearman_corr': 0.88
                    }
                }
            },
            'nested_cv': {
                'random_forest': {
                    'mean_score': 0.82,
                    'std_score': 0.05
                }
            }
        }
        
        report = orchestrator._create_markdown_report()
        
        # Check that report contains expected content
        self.assertIn("Research Report", report)
        self.assertIn("test_report", report)
        self.assertIn("random_forest", report)
        self.assertIn("0.85", report)  # R² score
    
    def test_pipeline_with_minimal_config(self):
        """Test pipeline with minimal configuration"""
        config = create_experiment_config(
            experiment_name="test_minimal",
            description="Test minimal pipeline",
            data_path="./data",
            output_path=self.temp_dir,
            enable_mlflow=False,
            enable_optuna=False,
            enable_nested_cv=False,
            enable_feature_ablation=False,
            enable_uncertainty_quantification=False,
            enable_interpretability=False,
            enable_ensemble=False,
            models_to_test=['random_forest']
        )
        
        orchestrator = ResearchPipelineOrchestrator(config)
        X, y = self.test_data
        
        try:
            results = orchestrator.run_complete_pipeline(X, y)
            self.assertIsInstance(results, dict)
        except Exception as e:
            # Some components might not be available in test environment
            print(f"Pipeline test failed (expected in some environments): {e}")

def run_integration_test():
    """Run integration test with real data"""
    print("Running integration test...")
    
    try:
        # Create test data
        np.random.seed(42)
        X = pd.DataFrame(
            np.random.randn(100, 10),
            columns=[f'feature_{i}' for i in range(10)]
        )
        y = np.random.uniform(-10, 0, 100)
        
        # Create configuration
        config = create_experiment_config(
            experiment_name="integration_test",
            description="Integration test with synthetic data",
            data_path="./data",
            output_path="./results/integration_test",
            n_trials=5,  # Minimal for testing
            cv_folds=3,
            models_to_test=['random_forest', 'neural_network'],
            enable_mlflow=False,  # Disable for testing
            enable_optuna=False
        )
        
        # Create orchestrator
        orchestrator = ResearchPipelineOrchestrator(config)
        
        # Run pipeline
        results = orchestrator.run_complete_pipeline(X, y)
        
        print("Integration test completed successfully!")
        print(f"Results keys: {list(results.keys())}")
        
        return True
        
    except Exception as e:
        print(f"Integration test failed: {e}")
        import traceback
        traceback.print_exc()
        return False

def main():
    """Run all tests"""
    print("Running Pipeline Orchestrator Tests")
    print("=" * 50)
    
    # Run unit tests
    unittest.main(argv=[''], exit=False, verbosity=2)
    
    # Run integration test
    print("\n" + "=" * 50)
    success = run_integration_test()
    
    if success:
        print("\nAll tests completed successfully!")
    else:
        print("\nSome tests failed - check output above")

if __name__ == "__main__":
    main()
```
