## Supplementary material for "Machine Learning-Driven Prediction of TLR4 Binding Affinity: A Comprehensive Molecular Feature Analysis for Drug Discovery": Codebase: test_uncertainty_minimal.py

```
#!/usr/bin/env python3
"""
Minimal test for uncertainty quantification methods.
This script tests only the core uncertainty quantification classes.
"""

import sys
import os
import numpy as np
from sklearn.ensemble import RandomForestRegressor
from sklearn.datasets import make_regression
from sklearn.model_selection import train_test_split
from sklearn.preprocessing import StandardScaler

# Add src to path
sys.path.append(os.path.join(os.path.dirname(__file__), 'src'))

def test_bootstrap_uncertainty_direct():
    """Test Bootstrap uncertainty quantification directly."""
    print("Testing Bootstrap Uncertainty Quantification...")
    
    try:
        # Import only the specific class
        from tlr4_binding.ml_components.uncertainty_quantification import BootstrapUncertainty
        
        # Generate sample data
        X, y = make_regression(n_samples=200, n_features=10, noise=0.1, random_state=42)
        X_train, X_test, y_train, y_test = train_test_split(X, y, test_size=0.2, random_state=42)
        
        # Scale features
        scaler = StandardScaler()
        X_train_scaled = scaler.fit_transform(X_train)
        X_test_scaled = scaler.transform(X_test)
        
        # Test Bootstrap Uncertainty
        bootstrap = BootstrapUncertainty(
            RandomForestRegressor,
            n_bootstrap=5,  # Reduced for faster testing
            random_state=42,
            n_estimators=5
        )
        
        # Fit
        bootstrap.fit(X_train_scaled, y_train)
        print(f"  ✓ Trained {len(bootstrap.bootstrap_models)} bootstrap models")
        
        # Predict
        result = bootstrap.predict_with_uncertainty(X_test_scaled)
        
        # Check results
        assert result.predictions.shape == (X_test_scaled.shape[0],)
        assert result.uncertainties.shape == (X_test_scaled.shape[0],)
        assert np.all(result.uncertainties >= 0)
        
        mse = np.mean((y_test - result.predictions) ** 2)
        print(f"  ✓ MSE: {mse:.4f}")
        print(f"  ✓ Mean Uncertainty: {np.mean(result.uncertainties):.4f}")
        
        return True
        
    except Exception as e:
        print(f"  ✗ Bootstrap test failed: {e}")
        import traceback
        traceback.print_exc()
        return False


def test_conformal_prediction_direct():
    """Test Conformal Prediction directly."""
    print("Testing Conformal Prediction...")
    
    try:
        # Import only the specific class
        from tlr4_binding.ml_components.uncertainty_quantification import ConformalPrediction
        
        # Generate sample data
        X, y = make_regression(n_samples=200, n_features=10, noise=0.1, random_state=42)
        X_train, X_test, y_train, y_test = train_test_split(X, y, test_size=0.2, random_state=42)
        
        # Scale features
        scaler = StandardScaler()
        X_train_scaled = scaler.fit_transform(X_train)
        X_test_scaled = scaler.transform(X_test)
        
        # Train base model
        base_model = RandomForestRegressor(n_estimators=5, random_state=42)
        base_model.fit(X_train_scaled, y_train)
        
        # Test Conformal Prediction
        conformal = ConformalPrediction(base_model, alpha=0.05, method='quantile')
        conformal.fit(X_train_scaled, y_train)
        
        # Predict
        result = conformal.predict_with_uncertainty(X_test_scaled)
        
        # Check results
        assert result.predictions.shape == (X_test_scaled.shape[0],)
        assert result.uncertainties.shape == (X_test_scaled.shape[0],)
        assert np.all(result.uncertainties >= 0)
        
        mse = np.mean((y_test - result.predictions) ** 2)
        print(f"  ✓ MSE: {mse:.4f}")
        print(f"  ✓ Mean Uncertainty: {np.mean(result.uncertainties):.4f}")
        
        return True
        
    except Exception as e:
        print(f"  ✗ Conformal prediction test failed: {e}")
        import traceback
        traceback.print_exc()
        return False


def test_ensemble_uncertainty_direct():
    """Test Ensemble uncertainty quantification directly."""
    print("Testing Ensemble Uncertainty...")
    
    try:
        # Import only the specific class
        from tlr4_binding.ml_components.uncertainty_quantification import EnsembleUncertainty
        
        # Generate sample data
        X, y = make_regression(n_samples=200, n_features=10, noise=0.1, random_state=42)
        X_train, X_test, y_train, y_test = train_test_split(X, y, test_size=0.2, random_state=42)
        
        # Scale features
        scaler = StandardScaler()
        X_train_scaled = scaler.fit_transform(X_train)
        X_test_scaled = scaler.transform(X_test)
        
        # Train multiple models
        models = []
        for i in range(3):
            model = RandomForestRegressor(n_estimators=5, random_state=42+i)
            model.fit(X_train_scaled, y_train)
            models.append(model)
        
        # Test Ensemble Uncertainty
        ensemble = EnsembleUncertainty(models)
        result = ensemble.predict_with_uncertainty(X_test_scaled)
        
        # Check results
        assert result.predictions.shape == (X_test_scaled.shape[0],)
        assert result.uncertainties.shape == (X_test_scaled.shape[0],)
        assert np.all(result.uncertainties >= 0)
        
        mse = np.mean((y_test - result.predictions) ** 2)
        print(f"  ✓ MSE: {mse:.4f}")
        print(f"  ✓ Mean Uncertainty: {np.mean(result.uncertainties):.4f}")
        
        return True
        
    except Exception as e:
        print(f"  ✗ Ensemble test failed: {e}")
        import traceback
        traceback.print_exc()
        return False


def test_uncertainty_calibration_direct():
    """Test uncertainty calibration analysis directly."""
    print("Testing Uncertainty Calibration...")
    
    try:
        # Import only the specific class
        from tlr4_binding.ml_components.uncertainty_quantification import UncertaintyCalibration
        
        # Generate sample data
        np.random.seed(42)
        n_samples = 100
        y_true = np.random.randn(n_samples)
        y_pred = y_true + 0.1 * np.random.randn(n_samples)
        uncertainties = np.abs(np.random.randn(n_samples)) + 0.1
        
        # Test calibration
        calibrator = UncertaintyCalibration()
        metrics = calibrator.calculate_calibration_metrics(y_true, y_pred, uncertainties)
        
        # Check metrics
        expected_metrics = ['calibration_error', 'coverage_probability', 'sharpness', 'negative_log_likelihood']
        for metric in expected_metrics:
            assert metric in metrics
            assert isinstance(metrics[metric], float)
        
        assert 0 <= metrics['coverage_probability'] <= 1
        assert metrics['sharpness'] >= 0
        
        print(f"  ✓ Calibration Error: {metrics['calibration_error']:.4f}")
        print(f"  ✓ Coverage Probability: {metrics['coverage_probability']:.4f}")
        print(f"  ✓ Sharpness: {metrics['sharpness']:.4f}")
        
        return True
        
    except Exception as e:
        print(f"  ✗ Calibration test failed: {e}")
        import traceback
        traceback.print_exc()
        return False


def main():
    """Run all tests."""
    print("=" * 60)
    print("UNCERTAINTY QUANTIFICATION MINIMAL TESTS")
    print("=" * 60)
    
    tests = [
        test_bootstrap_uncertainty_direct,
        test_conformal_prediction_direct,
        test_ensemble_uncertainty_direct,
        test_uncertainty_calibration_direct
    ]
    
    passed = 0
    total = len(tests)
    
    for test in tests:
        try:
            if test():
                passed += 1
        except Exception as e:
            print(f"  ✗ Test failed with exception: {e}")
        print()
    
    print("=" * 60)
    print(f"RESULTS: {passed}/{total} tests passed")
    print("=" * 60)
    
    if passed == total:
        print("🎉 All uncertainty quantification tests passed!")
        return True
    else:
        print("❌ Some tests failed.")
        return False


if __name__ == "__main__":
    success = main()
    sys.exit(0 if success else 1)
```
