## Supplementary material for "Machine Learning-Driven Prediction of TLR4 Binding Affinity: A Comprehensive Molecular Feature Analysis for Drug Discovery": Codebase: test_uncertainty_quantification.py

```
"""
Unit tests for uncertainty quantification methods.
"""

import pytest
import numpy as np
import torch
import torch.nn as nn
from sklearn.ensemble import RandomForestRegressor
from sklearn.datasets import make_regression
from sklearn.model_selection import train_test_split
from sklearn.preprocessing import StandardScaler

import sys
import os
sys.path.append(os.path.join(os.path.dirname(__file__), '..', 'src'))

from tlr4_binding.ml_components.uncertainty_quantification import (
    MonteCarloDropout,
    BootstrapUncertainty,
    ConformalPrediction,
    EnsembleUncertainty,
    UncertaintyQuantifier,
    UncertaintyCalibration,
    UncertaintyResult
)


class TestMonteCarloDropout:
    """Test Monte Carlo Dropout uncertainty quantification."""
    
    @pytest.fixture
    def sample_data(self):
        """Create sample data for testing."""
        X, y = make_regression(n_samples=100, n_features=10, noise=0.1, random_state=42)
        X_train, X_test, y_train, y_test = train_test_split(X, y, test_size=0.2, random_state=42)
        
        scaler = StandardScaler()
        X_train_scaled = scaler.fit_transform(X_train)
        X_test_scaled = scaler.transform(X_test)
        
        return X_train_scaled, X_test_scaled, y_train, y_test
    
    @pytest.fixture
    def simple_model(self, sample_data):
        """Create a simple neural network model."""
        X_train, _, _, _ = sample_data
        
        class SimpleModel(nn.Module):
            def __init__(self, input_size):
                super().__init__()
                self.network = nn.Sequential(
                    nn.Linear(input_size, 32),
                    nn.ReLU(),
                    nn.Dropout(0.2),
                    nn.Linear(32, 16),
                    nn.ReLU(),
                    nn.Dropout(0.2),
                    nn.Linear(16, 1)
                )
                
            def forward(self, x):
                return self.network(x).squeeze()
        
        model = SimpleModel(X_train.shape[1])
        
        # Train the model
        X_train_tensor = torch.FloatTensor(X_train)
        y_train_tensor = torch.FloatTensor(sample_data[2])
        
        criterion = nn.MSELoss()
        optimizer = torch.optim.Adam(model.parameters(), lr=0.01)
        
        model.train()
        for _ in range(50):
            optimizer.zero_grad()
            outputs = model(X_train_tensor)
            loss = criterion(outputs, y_train_tensor)
            loss.backward()
            optimizer.step()
        
        return model
    
    def test_mc_dropout_initialization(self):
        """Test Monte Carlo Dropout initialization."""
        model = nn.Linear(10, 1)
        mc_dropout = MonteCarloDropout(model, n_samples=50, dropout_rate=0.1)
        
        assert mc_dropout.model == model
        assert mc_dropout.n_samples == 50
        assert mc_dropout.dropout_rate == 0.1
    
    def test_mc_dropout_prediction(self, simple_model, sample_data):
        """Test Monte Carlo Dropout prediction with uncertainty."""
        _, X_test, _, _ = sample_data
        X_test_tensor = torch.FloatTensor(X_test)
        
        mc_dropout = MonteCarloDropout(simple_model, n_samples=20, dropout_rate=0.2)
        result = mc_dropout.predict_with_uncertainty(X_test_tensor)
        
        # Check result type
        assert isinstance(result, UncertaintyResult)
        
        # Check shapes
        assert result.predictions.shape == (X_test.shape[0],)
        assert result.uncertainties.shape == (X_test.shape[0],)
        assert result.confidence_intervals[0].shape == (X_test.shape[0],)
        assert result.confidence_intervals[1].shape == (X_test.shape[0],)
        
        # Check that uncertainties are non-negative
        assert np.all(result.uncertainties >= 0)
        
        # Check that confidence intervals are ordered correctly
        assert np.all(result.confidence_intervals[0] <= result.confidence_intervals[1])


class TestBootstrapUncertainty:
    """Test Bootstrap uncertainty quantification."""
    
    @pytest.fixture
    def sample_data(self):
        """Create sample data for testing."""
        X, y = make_regression(n_samples=200, n_features=10, noise=0.1, random_state=42)
        X_train, X_test, y_train, y_test = train_test_split(X, y, test_size=0.2, random_state=42)
        
        scaler = StandardScaler()
        X_train_scaled = scaler.fit_transform(X_train)
        X_test_scaled = scaler.transform(X_test)
        
        return X_train_scaled, X_test_scaled, y_train, y_test
    
    def test_bootstrap_initialization(self):
        """Test Bootstrap Uncertainty initialization."""
        bootstrap = BootstrapUncertainty(
            RandomForestRegressor, 
            n_bootstrap=20, 
            random_state=42
        )
        
        assert bootstrap.base_model_class == RandomForestRegressor
        assert bootstrap.n_bootstrap == 20
        assert bootstrap.random_state == 42
    
    def test_bootstrap_fit_predict(self, sample_data):
        """Test Bootstrap Uncertainty fit and predict."""
        X_train, X_test, y_train, y_test = sample_data
        
        bootstrap = BootstrapUncertainty(
            RandomForestRegressor, 
            n_bootstrap=10, 
            random_state=42,
            n_estimators=10
        )
        
        # Fit
        bootstrap.fit(X_train, y_train)
        assert len(bootstrap.bootstrap_models) == 10
        assert len(bootstrap.bootstrap_scores) == 10
        
        # Predict
        result = bootstrap.predict_with_uncertainty(X_test)
        
        # Check result type
        assert isinstance(result, UncertaintyResult)
        
        # Check shapes
        assert result.predictions.shape == (X_test.shape[0],)
        assert result.uncertainties.shape == (X_test.shape[0],)
        assert result.confidence_intervals[0].shape == (X_test.shape[0],)
        assert result.confidence_intervals[1].shape == (X_test.shape[0],)
        
        # Check that uncertainties are non-negative
        assert np.all(result.uncertainties >= 0)
        
        # Check that confidence intervals are ordered correctly
        assert np.all(result.confidence_intervals[0] <= result.confidence_intervals[1])


class TestConformalPrediction:
    """Test Conformal Prediction uncertainty quantification."""
    
    @pytest.fixture
    def sample_data(self):
        """Create sample data for testing."""
        X, y = make_regression(n_samples=200, n_features=10, noise=0.1, random_state=42)
        X_train, X_test, y_train, y_test = train_test_split(X, y, test_size=0.2, random_state=42)
        
        scaler = StandardScaler()
        X_train_scaled = scaler.fit_transform(X_train)
        X_test_scaled = scaler.transform(X_test)
        
        return X_train_scaled, X_test_scaled, y_train, y_test
    
    def test_conformal_initialization(self):
        """Test Conformal Prediction initialization."""
        model = RandomForestRegressor()
        conformal = ConformalPrediction(model, alpha=0.05, method='quantile')
        
        assert conformal.base_model == model
        assert conformal.alpha == 0.05
        assert conformal.method == 'quantile'
    
    def test_conformal_fit_predict(self, sample_data):
        """Test Conformal Prediction fit and predict."""
        X_train, X_test, y_train, y_test = sample_data
        
        model = RandomForestRegressor(n_estimators=10, random_state=42)
        conformal = ConformalPrediction(model, alpha=0.05, method='quantile')
        
        # Fit
        conformal.fit(X_train, y_train)
        assert conformal.conformity_scores is not None
        assert conformal.quantile_threshold is not None
        
        # Predict
        result = conformal.predict_with_uncertainty(X_test)
        
        # Check result type
        assert isinstance(result, UncertaintyResult)
        
        # Check shapes
        assert result.predictions.shape == (X_test.shape[0],)
        assert result.uncertainties.shape == (X_test.shape[0],)
        assert result.confidence_intervals[0].shape == (X_test.shape[0],)
        assert result.confidence_intervals[1].shape == (X_test.shape[0],)
        
        # Check that uncertainties are non-negative
        assert np.all(result.uncertainties >= 0)
        
        # Check that confidence intervals are ordered correctly
        assert np.all(result.confidence_intervals[0] <= result.confidence_intervals[1])


class TestEnsembleUncertainty:
    """Test Ensemble uncertainty quantification."""
    
    @pytest.fixture
    def sample_data(self):
        """Create sample data for testing."""
        X, y = make_regression(n_samples=200, n_features=10, noise=0.1, random_state=42)
        X_train, X_test, y_train, y_test = train_test_split(X, y, test_size=0.2, random_state=42)
        
        scaler = StandardScaler()
        X_train_scaled = scaler.fit_transform(X_train)
        X_test_scaled = scaler.transform(X_test)
        
        return X_train_scaled, X_test_scaled, y_train, y_test
    
    @pytest.fixture
    def ensemble_models(self, sample_data):
        """Create ensemble of models."""
        X_train, _, y_train, _ = sample_data
        
        models = []
        
        # Random Forest 1
        rf1 = RandomForestRegressor(n_estimators=10, random_state=42)
        rf1.fit(X_train, y_train)
        models.append(rf1)
        
        # Random Forest 2
        rf2 = RandomForestRegressor(n_estimators=10, random_state=43)
        rf2.fit(X_train, y_train)
        models.append(rf2)
        
        # Random Forest 3
        rf3 = RandomForestRegressor(n_estimators=10, random_state=44)
        rf3.fit(X_train, y_train)
        models.append(rf3)
        
        return models
    
    def test_ensemble_initialization(self, ensemble_models):
        """Test Ensemble Uncertainty initialization."""
        ensemble = EnsembleUncertainty(ensemble_models)
        
        assert len(ensemble.models) == 3
        assert len(ensemble.weights) == 3
        assert np.isclose(np.sum(ensemble.weights), 1.0)
    
    def test_ensemble_prediction(self, ensemble_models, sample_data):
        """Test Ensemble Uncertainty prediction."""
        _, X_test, _, _ = sample_data
        
        ensemble = EnsembleUncertainty(ensemble_models)
        result = ensemble.predict_with_uncertainty(X_test)
        
        # Check result type
        assert isinstance(result, UncertaintyResult)
        
        # Check shapes
        assert result.predictions.shape == (X_test.shape[0],)
        assert result.uncertainties.shape == (X_test.shape[0],)
        assert result.confidence_intervals[0].shape == (X_test.shape[0],)
        assert result.confidence_intervals[1].shape == (X_test.shape[0],)
        
        # Check that uncertainties are non-negative
        assert np.all(result.uncertainties >= 0)
        
        # Check that confidence intervals are ordered correctly
        assert np.all(result.confidence_intervals[0] <= result.confidence_intervals[1])


class TestUncertaintyCalibration:
    """Test uncertainty calibration analysis."""
    
    @pytest.fixture
    def sample_data(self):
        """Create sample data for testing."""
        np.random.seed(42)
        n_samples = 100
        
        y_true = np.random.randn(n_samples)
        y_pred = y_true + 0.1 * np.random.randn(n_samples)
        uncertainties = np.abs(np.random.randn(n_samples)) + 0.1
        
        return y_true, y_pred, uncertainties
    
    def test_calibration_metrics(self, sample_data):
        """Test calibration metrics calculation."""
        y_true, y_pred, uncertainties = sample_data
        
        calibrator = UncertaintyCalibration()
        metrics = calibrator.calculate_calibration_metrics(y_true, y_pred, uncertainties)
        
        # Check that all metrics are present
        expected_metrics = ['calibration_error', 'coverage_probability', 'sharpness', 'negative_log_likelihood']
        for metric in expected_metrics:
            assert metric in metrics
            assert isinstance(metrics[metric], float)
        
        # Check that coverage probability is between 0 and 1
        assert 0 <= metrics['coverage_probability'] <= 1
        
        # Check that sharpness is non-negative
        assert metrics['sharpness'] >= 0


class TestUncertaintyQuantifier:
    """Test main Uncertainty Quantifier class."""
    
    @pytest.fixture
    def sample_data(self):
        """Create sample data for testing."""
        X, y = make_regression(n_samples=200, n_features=10, noise=0.1, random_state=42)
        X_train, X_test, y_train, y_test = train_test_split(X, y, test_size=0.2, random_state=42)
        
        scaler = StandardScaler()
        X_train_scaled = scaler.fit_transform(X_train)
        X_test_scaled = scaler.transform(X_test)
        
        return X_train_scaled, X_test_scaled, y_train, y_test
    
    def test_quantifier_bootstrap(self, sample_data):
        """Test Uncertainty Quantifier with bootstrap method."""
        X_train, X_test, y_train, y_test = sample_data
        
        # Create base model
        base_model = RandomForestRegressor(n_estimators=10, random_state=42)
        base_model.fit(X_train, y_train)
        
        # Test bootstrap quantifier
        quantifier = UncertaintyQuantifier(method='bootstrap', n_bootstrap=10, random_state=42)
        quantifier.fit(base_model.__class__, X_train, y_train)
        
        result = quantifier.predict_with_uncertainty(X_test)
        
        assert isinstance(result, UncertaintyResult)
        assert result.predictions.shape == (X_test.shape[0],)
        assert result.uncertainties.shape == (X_test.shape[0],)
    
    def test_quantifier_conformal(self, sample_data):
        """Test Uncertainty Quantifier with conformal method."""
        X_train, X_test, y_train, y_test = sample_data
        
        # Create base model
        base_model = RandomForestRegressor(n_estimators=10, random_state=42)
        base_model.fit(X_train, y_train)
        
        # Test conformal quantifier
        quantifier = UncertaintyQuantifier(method='conformal', alpha=0.05)
        quantifier.fit(base_model, X_train, y_train)
        
        result = quantifier.predict_with_uncertainty(X_test)
        
        assert isinstance(result, UncertaintyResult)
        assert result.predictions.shape == (X_test.shape[0],)
        assert result.uncertainties.shape == (X_test.shape[0],)
    
    def test_quantifier_ensemble(self, sample_data):
        """Test Uncertainty Quantifier with ensemble method."""
        X_train, X_test, y_train, y_test = sample_data
        
        # Create ensemble models
        models = []
        for i in range(3):
            model = RandomForestRegressor(n_estimators=10, random_state=42+i)
            model.fit(X_train, y_train)
            models.append(model)
        
        # Test ensemble quantifier
        quantifier = UncertaintyQuantifier(method='ensemble')
        quantifier.fit(models, X_train, y_train)
        
        result = quantifier.predict_with_uncertainty(X_test)
        
        assert isinstance(result, UncertaintyResult)
        assert result.predictions.shape == (X_test.shape[0],)
        assert result.uncertainties.shape == (X_test.shape[0],)


if __name__ == "__main__":
    pytest.main([__file__])
```
