## Supplementary material for "Machine Learning-Driven Prediction of TLR4 Binding Affinity: A Comprehensive Molecular Feature Analysis for Drug Discovery": Codebase: test_uncertainty_standalone.py

```
#!/usr/bin/env python3
"""
Standalone test for uncertainty quantification methods.
This script tests the uncertainty quantification classes directly without importing the full module.
"""

import sys
import os
import numpy as np
from sklearn.ensemble import RandomForestRegressor
from sklearn.datasets import make_regression
from sklearn.model_selection import train_test_split
from sklearn.preprocessing import StandardScaler
from typing import Dict, List, Tuple, Optional, Union, Any
from dataclasses import dataclass
import logging

# Set up logging
logging.basicConfig(level=logging.INFO)
logger = logging.getLogger(__name__)


@dataclass
class UncertaintyResult:
    """Container for uncertainty quantification results"""
    predictions: np.ndarray
    uncertainties: np.ndarray
    confidence_intervals: Tuple[np.ndarray, np.ndarray]  # (lower, upper)
    prediction_intervals: Tuple[np.ndarray, np.ndarray]  # (lower, upper)
    epistemic_uncertainty: Optional[np.ndarray] = None
    aleatoric_uncertainty: Optional[np.ndarray] = None


class BootstrapUncertainty:
    """
    Bootstrap aggregating for uncertainty quantification in traditional ML models.
    """
    
    def __init__(self, 
                 base_model_class,
                 n_bootstrap: int = 100,
                 bootstrap_ratio: float = 0.8,
                 random_state: int = 42,
                 **model_kwargs):
        self.base_model_class = base_model_class
        self.n_bootstrap = n_bootstrap
        self.bootstrap_ratio = bootstrap_ratio
        self.random_state = random_state
        self.model_kwargs = model_kwargs
        self.bootstrap_models = []
        self.bootstrap_scores = []
        
    def fit(self, X: np.ndarray, y: np.ndarray) -> 'BootstrapUncertainty':
        """Fit bootstrap models."""
        n_samples = int(len(X) * self.bootstrap_ratio)
        rng = np.random.RandomState(self.random_state)
        
        logger.info(f"Training {self.n_bootstrap} bootstrap models...")
        
        for i in range(self.n_bootstrap):
            # Create bootstrap sample
            bootstrap_indices = rng.choice(len(X), size=n_samples, replace=True)
            X_bootstrap = X[bootstrap_indices]
            y_bootstrap = y[bootstrap_indices]
            
            # Train model
            model = self.base_model_class(**self.model_kwargs)
            model.fit(X_bootstrap, y_bootstrap)
            self.bootstrap_models.append(model)
            
            # Calculate bootstrap score
            y_pred = model.predict(X_bootstrap)
            score = np.mean((y_bootstrap - y_pred) ** 2)
            self.bootstrap_scores.append(score)
            
            if (i + 1) % 10 == 0:
                logger.info(f"Trained {i + 1}/{self.n_bootstrap} bootstrap models")
                
        return self
        
    def predict_with_uncertainty(self, X: np.ndarray) -> UncertaintyResult:
        """Make predictions with uncertainty quantification."""
        if not self.bootstrap_models:
            raise ValueError("Must call fit() before predict_with_uncertainty()")
            
        predictions = []
        
        for model in self.bootstrap_models:
            pred = model.predict(X)
            predictions.append(pred)
            
        predictions = np.array(predictions)
        
        # Calculate statistics
        mean_predictions = np.mean(predictions, axis=0)
        epistemic_uncertainty = np.var(predictions, axis=0)
        
        # Estimate aleatoric uncertainty from bootstrap scores
        aleatoric_uncertainty = np.full_like(mean_predictions, np.mean(self.bootstrap_scores))
        
        # Calculate confidence intervals (95%)
        lower_ci = np.percentile(predictions, 2.5, axis=0)
        upper_ci = np.percentile(predictions, 97.5, axis=0)
        
        # Calculate prediction intervals (includes both epistemic and aleatoric uncertainty)
        total_uncertainty = epistemic_uncertainty + aleatoric_uncertainty
        std_total = np.sqrt(total_uncertainty)
        lower_pi = mean_predictions - 1.96 * std_total
        upper_pi = mean_predictions + 1.96 * std_total
        
        return UncertaintyResult(
            predictions=mean_predictions,
            uncertainties=epistemic_uncertainty,
            confidence_intervals=(lower_ci, upper_ci),
            prediction_intervals=(lower_pi, upper_pi),
            epistemic_uncertainty=epistemic_uncertainty,
            aleatoric_uncertainty=aleatoric_uncertainty
        )


class ConformalPrediction:
    """
    Conformal Prediction for uncertainty quantification.
    """
    
    def __init__(self, 
                 base_model,
                 alpha: float = 0.05,
                 method: str = 'quantile'):
        self.base_model = base_model
        self.alpha = alpha
        self.method = method
        self.conformity_scores = None
        self.quantile_threshold = None
        
    def fit(self, X: np.ndarray, y: np.ndarray) -> 'ConformalPrediction':
        """Fit conformal prediction model."""
        # Fit base model
        self.base_model.fit(X, y)
        
        # Calculate conformity scores on training data
        y_pred = self.base_model.predict(X)
        residuals = np.abs(y - y_pred)
        
        if self.method == 'quantile':
            self.conformity_scores = residuals
            self.quantile_threshold = np.quantile(self.conformity_scores, 1 - self.alpha)
        elif self.method == 'normalized':
            # Normalize residuals by prediction magnitude
            normalized_residuals = residuals / (np.abs(y_pred) + 1e-8)
            self.conformity_scores = normalized_residuals
            self.quantile_threshold = np.quantile(self.conformity_scores, 1 - self.alpha)
        else:
            raise ValueError(f"Unknown method: {self.method}")
            
        return self
        
    def predict_with_uncertainty(self, X: np.ndarray) -> UncertaintyResult:
        """Make predictions with conformal prediction intervals."""
        if self.conformity_scores is None:
            raise ValueError("Must call fit() before predict_with_uncertainty()")
            
        # Get point predictions
        predictions = self.base_model.predict(X)
        
        # Calculate prediction intervals
        if self.method == 'quantile':
            interval_width = self.quantile_threshold
        elif self.method == 'normalized':
            interval_width = self.quantile_threshold * (np.abs(predictions) + 1e-8)
        else:
            raise ValueError(f"Unknown method: {self.method}")
            
        lower_pi = predictions - interval_width
        upper_pi = predictions + interval_width
        
        # For conformal prediction, epistemic uncertainty is the interval width
        epistemic_uncertainty = np.full_like(predictions, interval_width)
        
        return UncertaintyResult(
            predictions=predictions,
            uncertainties=epistemic_uncertainty,
            confidence_intervals=(lower_pi, upper_pi),
            prediction_intervals=(lower_pi, upper_pi),
            epistemic_uncertainty=epistemic_uncertainty
        )


class EnsembleUncertainty:
    """
    Ensemble-based uncertainty quantification.
    """
    
    def __init__(self, models: List[Any], weights: Optional[List[float]] = None):
        self.models = models
        self.weights = weights or [1.0 / len(models)] * len(models)
        
        if len(self.weights) != len(self.models):
            raise ValueError("Number of weights must match number of models")
            
    def predict_with_uncertainty(self, X: np.ndarray) -> UncertaintyResult:
        """Make predictions with ensemble uncertainty quantification."""
        predictions = []
        
        for model in self.models:
            if hasattr(model, 'predict'):
                pred = model.predict(X)
            else:
                raise ValueError(f"Model {type(model)} does not have predict method")
                
            predictions.append(pred)
            
        predictions = np.array(predictions)
        
        # Weighted ensemble predictions
        weighted_predictions = np.average(predictions, axis=0, weights=self.weights)
        
        # Calculate ensemble uncertainty
        # Epistemic uncertainty: variance across models
        epistemic_uncertainty = np.var(predictions, axis=0)
        
        # Total uncertainty: includes both epistemic and aleatoric
        # For ensemble, we estimate aleatoric as the mean prediction error
        mean_abs_error = np.mean(np.abs(predictions - weighted_predictions), axis=0)
        total_uncertainty = epistemic_uncertainty + mean_abs_error
        
        # Calculate confidence intervals (95%)
        lower_ci = np.percentile(predictions, 2.5, axis=0)
        upper_ci = np.percentile(predictions, 97.5, axis=0)
        
        # Calculate prediction intervals
        std_total = np.sqrt(total_uncertainty)
        lower_pi = weighted_predictions - 1.96 * std_total
        upper_pi = weighted_predictions + 1.96 * std_total
        
        return UncertaintyResult(
            predictions=weighted_predictions,
            uncertainties=epistemic_uncertainty,
            confidence_intervals=(lower_ci, upper_ci),
            prediction_intervals=(lower_pi, upper_pi),
            epistemic_uncertainty=epistemic_uncertainty,
            aleatoric_uncertainty=mean_abs_error
        )


class UncertaintyCalibration:
    """
    Uncertainty calibration analysis and reliability diagrams.
    """
    
    def __init__(self, n_bins: int = 10):
        self.n_bins = n_bins
        
    def calculate_calibration_metrics(self, 
                                    y_true: np.ndarray,
                                    y_pred: np.ndarray,
                                    uncertainties: np.ndarray) -> Dict[str, float]:
        """Calculate calibration metrics."""
        # Calculate residuals
        residuals = np.abs(y_true - y_pred)
        
        # Normalize residuals by uncertainty
        normalized_residuals = residuals / (uncertainties + 1e-8)
        
        # Calculate calibration metrics
        calibration_error = np.mean(np.abs(normalized_residuals - 1.0))
        
        # Calculate coverage probability
        coverage_prob = np.mean(normalized_residuals <= 1.0)
        
        # Calculate sharpness (average uncertainty)
        sharpness = np.mean(uncertainties)
        
        # Calculate negative log likelihood
        nll = np.mean(0.5 * np.log(2 * np.pi * uncertainties**2) + 
                     0.5 * (residuals**2) / (uncertainties**2))
        
        return {
            'calibration_error': calibration_error,
            'coverage_probability': coverage_prob,
            'sharpness': sharpness,
            'negative_log_likelihood': nll
        }


def test_bootstrap_uncertainty():
    """Test Bootstrap uncertainty quantification."""
    print("Testing Bootstrap Uncertainty Quantification...")
    
    try:
        # Generate sample data
        X, y = make_regression(n_samples=200, n_features=10, noise=0.1, random_state=42)
        X_train, X_test, y_train, y_test = train_test_split(X, y, test_size=0.2, random_state=42)
        
        # Scale features
        scaler = StandardScaler()
        X_train_scaled = scaler.fit_transform(X_train)
        X_test_scaled = scaler.transform(X_test)
        
        # Test Bootstrap Uncertainty
        bootstrap = BootstrapUncertainty(
            RandomForestRegressor,
            n_bootstrap=10,
            random_state=42,
            n_estimators=10
        )
        
        # Fit
        bootstrap.fit(X_train_scaled, y_train)
        print(f"  ✓ Trained {len(bootstrap.bootstrap_models)} bootstrap models")
        
        # Predict
        result = bootstrap.predict_with_uncertainty(X_test_scaled)
        
        # Check results
        assert result.predictions.shape == (X_test_scaled.shape[0],)
        assert result.uncertainties.shape == (X_test_scaled.shape[0],)
        assert np.all(result.uncertainties >= 0)
        
        mse = np.mean((y_test - result.predictions) ** 2)
        print(f"  ✓ MSE: {mse:.4f}")
        print(f"  ✓ Mean Uncertainty: {np.mean(result.uncertainties):.4f}")
        
        return True
        
    except Exception as e:
        print(f"  ✗ Bootstrap test failed: {e}")
        import traceback
        traceback.print_exc()
        return False


def test_conformal_prediction():
    """Test Conformal Prediction uncertainty quantification."""
    print("Testing Conformal Prediction...")
    
    try:
        # Generate sample data
        X, y = make_regression(n_samples=200, n_features=10, noise=0.1, random_state=42)
        X_train, X_test, y_train, y_test = train_test_split(X, y, test_size=0.2, random_state=42)
        
        # Scale features
        scaler = StandardScaler()
        X_train_scaled = scaler.fit_transform(X_train)
        X_test_scaled = scaler.transform(X_test)
        
        # Train base model
        base_model = RandomForestRegressor(n_estimators=10, random_state=42)
        base_model.fit(X_train_scaled, y_train)
        
        # Test Conformal Prediction
        conformal = ConformalPrediction(base_model, alpha=0.05, method='quantile')
        conformal.fit(X_train_scaled, y_train)
        
        # Predict
        result = conformal.predict_with_uncertainty(X_test_scaled)
        
        # Check results
        assert result.predictions.shape == (X_test_scaled.shape[0],)
        assert result.uncertainties.shape == (X_test_scaled.shape[0],)
        assert np.all(result.uncertainties >= 0)
        
        mse = np.mean((y_test - result.predictions) ** 2)
        print(f"  ✓ MSE: {mse:.4f}")
        print(f"  ✓ Mean Uncertainty: {np.mean(result.uncertainties):.4f}")
        
        return True
        
    except Exception as e:
        print(f"  ✗ Conformal prediction test failed: {e}")
        import traceback
        traceback.print_exc()
        return False


def test_ensemble_uncertainty():
    """Test Ensemble uncertainty quantification."""
    print("Testing Ensemble Uncertainty...")
    
    try:
        # Generate sample data
        X, y = make_regression(n_samples=200, n_features=10, noise=0.1, random_state=42)
        X_train, X_test, y_train, y_test = train_test_split(X, y, test_size=0.2, random_state=42)
        
        # Scale features
        scaler = StandardScaler()
        X_train_scaled = scaler.fit_transform(X_train)
        X_test_scaled = scaler.transform(X_test)
        
        # Train multiple models
        models = []
        for i in range(3):
            model = RandomForestRegressor(n_estimators=10, random_state=42+i)
            model.fit(X_train_scaled, y_train)
            models.append(model)
        
        # Test Ensemble Uncertainty
        ensemble = EnsembleUncertainty(models)
        result = ensemble.predict_with_uncertainty(X_test_scaled)
        
        # Check results
        assert result.predictions.shape == (X_test_scaled.shape[0],)
        assert result.uncertainties.shape == (X_test_scaled.shape[0],)
        assert np.all(result.uncertainties >= 0)
        
        mse = np.mean((y_test - result.predictions) ** 2)
        print(f"  ✓ MSE: {mse:.4f}")
        print(f"  ✓ Mean Uncertainty: {np.mean(result.uncertainties):.4f}")
        
        return True
        
    except Exception as e:
        print(f"  ✗ Ensemble test failed: {e}")
        import traceback
        traceback.print_exc()
        return False


def test_uncertainty_calibration():
    """Test uncertainty calibration analysis."""
    print("Testing Uncertainty Calibration...")
    
    try:
        # Generate sample data
        np.random.seed(42)
        n_samples = 100
        y_true = np.random.randn(n_samples)
        y_pred = y_true + 0.1 * np.random.randn(n_samples)
        uncertainties = np.abs(np.random.randn(n_samples)) + 0.1
        
        # Test calibration
        calibrator = UncertaintyCalibration()
        metrics = calibrator.calculate_calibration_metrics(y_true, y_pred, uncertainties)
        
        # Check metrics
        expected_metrics = ['calibration_error', 'coverage_probability', 'sharpness', 'negative_log_likelihood']
        for metric in expected_metrics:
            assert metric in metrics
            assert isinstance(metrics[metric], float)
        
        assert 0 <= metrics['coverage_probability'] <= 1
        assert metrics['sharpness'] >= 0
        
        print(f"  ✓ Calibration Error: {metrics['calibration_error']:.4f}")
        print(f"  ✓ Coverage Probability: {metrics['coverage_probability']:.4f}")
        print(f"  ✓ Sharpness: {metrics['sharpness']:.4f}")
        
        return True
        
    except Exception as e:
        print(f"  ✗ Calibration test failed: {e}")
        import traceback
        traceback.print_exc()
        return False


def main():
    """Run all tests."""
    print("=" * 60)
    print("UNCERTAINTY QUANTIFICATION STANDALONE TESTS")
    print("=" * 60)
    
    tests = [
        test_bootstrap_uncertainty,
        test_conformal_prediction,
        test_ensemble_uncertainty,
        test_uncertainty_calibration
    ]
    
    passed = 0
    total = len(tests)
    
    for test in tests:
        try:
            if test():
                passed += 1
        except Exception as e:
            print(f"  ✗ Test failed with exception: {e}")
        print()
    
    print("=" * 60)
    print(f"RESULTS: {passed}/{total} tests passed")
    print("=" * 60)
    
    if passed == total:
        print("🎉 All uncertainty quantification tests passed!")
        return True
    else:
        print("❌ Some tests failed.")
        return False


if __name__ == "__main__":
    success = main()
    sys.exit(0 if success else 1)
```
