## Supplementary material for "Machine Learning-Driven Prediction of TLR4 Binding Affinity: A Comprehensive Molecular Feature Analysis for Drug Discovery": Codebase: test_validation_benchmarking.py

```
import numpy as np
import pandas as pd
from sklearn.linear_model import LinearRegression

from src.tlr4_binding.ml_components.validation_benchmarking import (
    evaluate_leave_one_out,
    evaluate_temporal_split,
    evaluate_external,
    benchmark_models,
    recommend_sample_size_for_rmse_effect,
)


def make_tiny_regression(n=30, p=5, seed=0):
    rng = np.random.default_rng(seed)
    X = rng.normal(size=(n, p))
    coef = rng.normal(size=p)
    y = X @ coef + rng.normal(scale=0.5, size=n)
    return pd.DataFrame(X), pd.Series(y)


def test_evaluate_leave_one_out_verbose(capfd):
    X, y = make_tiny_regression(n=25, p=4)
    model = LinearRegression()
    res = evaluate_leave_one_out(model, X, y)
    assert "r2" in res and "rmse" in res and "mae" in res
    # Print verbose summary
    print("LOO RMSE mean:", res["rmse"]["mean"])  # noqa: T201
    out, _ = capfd.readouterr()
    assert "LOO RMSE mean:" in out


def test_evaluate_temporal_split():
    X, y = make_tiny_regression(n=40, p=3)
    timestamps = pd.date_range("2024-01-01", periods=len(X), freq="D")
    model = LinearRegression()
    res = evaluate_temporal_split(model, X, y, pd.Series(timestamps))
    assert res["n_train"] + res["n_test"] == len(X)
    assert set(["r2", "rmse", "mae"]) <= set(res["metrics"].keys())


def test_evaluate_external():
    X, y = make_tiny_regression(n=50, p=4)
    # External set: last 10
    X_train, y_train = X.iloc[:40], y.iloc[:40]
    X_ext, y_ext = X.iloc[40:], y.iloc[40:]
    model = LinearRegression()
    res = evaluate_external(model, X_train, y_train, X_ext, y_ext)
    assert res["n_external"] == 10
    assert "rmse" in res["metrics"]


def test_benchmark_models_verbose(capfd):
    X, y = make_tiny_regression(n=40, p=3)
    models = {
        "lr": LinearRegression(),
    }
    df = benchmark_models(models, X, y, cv_folds=3)
    assert not df.empty and "r2_mean" in df.columns
    print(df.to_string())  # noqa: T201
    out, _ = capfd.readouterr()
    assert "r2_mean" in out


def test_power_recommendation_runs():
    X, y = make_tiny_regression(n=60, p=4)
    a = LinearRegression()
    b = LinearRegression()
    res = recommend_sample_size_for_rmse_effect(
        a, b, X, y, target_rmse_improvement=0.1, power_target=0.5, max_n=40
    )
    assert res.required_n <= 40
```
