## Supplementary material for "Machine Learning-Driven Prediction of TLR4 Binding Affinity: A Comprehensive Molecular Feature Analysis for Drug Discovery": Codebase: standalone_pdbqt_parser.py

```
"""
Standalone PDBQT file parser for testing.

This is a copy of the parser module without external dependencies
for isolated testing of PDBQT parsing functionality.
"""

from abc import ABC, abstractmethod
from typing import Dict, List, Optional, Tuple, Union
import os
import logging
from pathlib import Path

logger = logging.getLogger(__name__)


class PDBQTParserInterface(ABC):
    """Abstract interface for PDBQT file parsing."""
    
    @abstractmethod
    def parse_file(self, file_path: str) -> Dict[str, Union[str, List[Dict]]]:
        """Parse a single PDBQT file and return structured data."""
        pass
    
    @abstractmethod
    def validate_file(self, file_path: str) -> Tuple[bool, List[str]]:
        """Validate PDBQT file format and return validation results."""
        pass
    
    @abstractmethod
    def extract_ligand_data(self, file_path: str) -> Dict[str, Union[str, float]]:
        """Extract ligand-specific data from PDBQT file."""
        pass


class PDBQTParser(PDBQTParserInterface):
    """
    PDBQT file parser with comprehensive validation and error handling.
    
    Handles parsing of AutoDock Vina output files in PDBQT format,
    including ligand structures, binding poses, and scoring information.
    """
    
    def __init__(self, strict_validation: bool = True):
        """
        Initialize parser with validation settings.
        
        Args:
            strict_validation: If True, enforce strict PDBQT format validation
        """
        self.strict_validation = strict_validation
        self.supported_atoms = {'C', 'N', 'O', 'S', 'P', 'F', 'Cl', 'Br', 'I', 'H'}
        self.required_sections = ['ATOM']  # Only ATOM/HETATM is truly required
    
    def parse_file(self, file_path: str) -> Dict[str, Union[str, List[Dict]]]:
        """
        Parse a single PDBQT file and return structured data.
        
        Args:
            file_path: Path to PDBQT file
            
        Returns:
            Dictionary containing parsed file data with sections:
            - header: File header information
            - atoms: List of atom records
            - models: List of binding poses/models
            - footer: File footer information
            
        Raises:
            FileNotFoundError: If file doesn't exist
            ValueError: If file format is invalid
        """
        if not os.path.exists(file_path):
            raise FileNotFoundError(f"PDBQT file not found: {file_path}")
        
        try:
            with open(file_path, 'r') as f:
                lines = f.readlines()
            
            parsed_data = {
                'file_path': file_path,
                'header': [],
                'atoms': [],
                'models': [],
                'footer': [],
                'metadata': {}
            }
            
            current_model = None
            in_atom_section = False
            
            for line_num, line in enumerate(lines, 1):
                line = line.strip()
                
                if not line:
                    continue
                
                # Parse different sections
                if line.startswith('REMARK'):
                    parsed_data['header'].append(line)
                    self._parse_remark(line, parsed_data['metadata'])
                
                elif line.startswith('ATOM') or line.startswith('HETATM'):
                    in_atom_section = True
                    atom_data = self._parse_atom_line(line, line_num)
                    parsed_data['atoms'].append(atom_data)
                    
                    # Add to current model if in model section
                    if current_model is not None:
                        current_model['atoms'].append(atom_data)
                
                elif line.startswith('MODEL'):
                    current_model = {
                        'model_number': self._extract_model_number(line),
                        'atoms': []
                    }
                    parsed_data['models'].append(current_model)
                
                elif line.startswith('ENDMDL'):
                    current_model = None
                    in_atom_section = False
                
                elif line == 'END':
                    parsed_data['footer'].append(line)
                    break
            
            # Validate parsed data
            if self.strict_validation:
                validation_errors = self._validate_parsed_data(parsed_data)
                if validation_errors:
                    raise ValueError(f"PDBQT validation failed: {validation_errors}")
            
            return parsed_data
            
        except Exception as e:
            logger.error(f"Error parsing PDBQT file {file_path}: {str(e)}")
            raise ValueError(f"Failed to parse PDBQT file: {str(e)}")
    
    def validate_file(self, file_path: str) -> Tuple[bool, List[str]]:
        """
        Validate PDBQT file format and return validation results.
        
        Args:
            file_path: Path to PDBQT file
            
        Returns:
            Tuple of (is_valid, error_messages)
        """
        errors = []
        
        if not os.path.exists(file_path):
            return False, [f"File not found: {file_path}"]
        
        try:
            with open(file_path, 'r') as f:
                lines = f.readlines()
            
            # Check file structure
            has_atoms = any(line.startswith(('ATOM', 'HETATM')) for line in lines)
            has_end = any(line.startswith('END') for line in lines)
            
            if not has_atoms:
                errors.append("No ATOM or HETATM records found")
            
            if not has_end:
                errors.append("No END record found")
            
            # Check atom format
            atom_lines = [line for line in lines if line.startswith(('ATOM', 'HETATM'))]
            for i, line in enumerate(atom_lines):
                if len(line) < 80:
                    errors.append(f"Atom line {i+1} too short: {line}")
                
                # Check atom name format
                atom_name = line[12:16].strip()
                if not atom_name:
                    errors.append(f"Atom line {i+1} missing atom name")
            
            # Check for required sections
            sections_found = set()
            for line in lines:
                if line.startswith('REMARK'):
                    sections_found.add('REMARK')
                elif line.startswith(('ATOM', 'HETATM')):
                    sections_found.add('ATOM')
                elif line.startswith('END'):
                    sections_found.add('END')
            
            missing_sections = set(self.required_sections) - sections_found
            if missing_sections:
                errors.append(f"Missing required sections: {missing_sections}")
            
            return len(errors) == 0, errors
            
        except Exception as e:
            return False, [f"Error reading file: {str(e)}"]
    
    def extract_ligand_data(self, file_path: str) -> Dict[str, Union[str, float]]:
        """
        Extract ligand-specific data from PDBQT file.
        
        Args:
            file_path: Path to PDBQT file
            
        Returns:
            Dictionary with ligand information including:
            - compound_name: Name of the ligand
            - atom_count: Number of atoms
            - molecular_formula: Chemical formula
            - binding_poses: Number of binding poses
        """
        parsed_data = self.parse_file(file_path)
        
        # Extract compound name from file path or REMARK lines
        compound_name = Path(file_path).stem
        
        # Count atoms and extract formula
        atoms = parsed_data['atoms']
        atom_count = len(atoms)
        
        # Count unique elements
        elements = {}
        for atom in atoms:
            element = atom.get('element', 'Unknown')
            elements[element] = elements.get(element, 0) + 1
        
        # Create molecular formula
        formula_parts = []
        for element in sorted(elements.keys()):
            count = elements[element]
            if count > 1:
                formula_parts.append(f"{element}{count}")
            else:
                formula_parts.append(element)
        molecular_formula = ''.join(formula_parts)
        
        # Count binding poses
        binding_poses = len(parsed_data['models'])
        
        return {
            'compound_name': compound_name,
            'atom_count': atom_count,
            'molecular_formula': molecular_formula,
            'binding_poses': binding_poses,
            'elements': elements,
            'file_path': file_path
        }
    
    def _parse_remark(self, line: str, metadata: Dict) -> None:
        """Parse REMARK line and extract metadata."""
        if 'VINA' in line.upper():
            # Extract Vina version or scoring information
            if 'VINA RESULT' in line.upper():
                parts = line.split()
                if len(parts) >= 4:
                    try:
                        metadata['vina_score'] = float(parts[3])
                    except ValueError:
                        pass
    
    def _parse_atom_line(self, line: str, line_num: int) -> Dict[str, Union[str, float, int]]:
        """Parse individual ATOM/HETATM line."""
        try:
            atom_data = {
                'line_number': line_num,
                'record_type': line[0:6].strip(),
                'atom_serial': int(line[6:11]) if line[6:11].strip() else 0,
                'atom_name': line[12:16].strip(),
                'residue_name': line[17:20].strip(),
                'chain_id': line[21:22].strip(),
                'residue_number': int(line[22:26]) if line[22:26].strip() else 0,
                'x': float(line[30:38]) if line[30:38].strip() else 0.0,
                'y': float(line[38:46]) if line[38:46].strip() else 0.0,
                'z': float(line[46:54]) if line[46:54].strip() else 0.0,
                'occupancy': float(line[54:60]) if line[54:60].strip() else 1.0,
                'temperature_factor': float(line[60:66]) if line[60:66].strip() else 0.0,
                'element': line[76:78].strip() if len(line) > 76 else '',
                'autodock_type': line[78:80].strip() if len(line) > 78 else '',
                'charge': line[66:76].strip() if len(line) > 76 else ''
            }
            
            # Extract element from atom name if not specified
            if not atom_data['element']:
                atom_name = atom_data['atom_name']
                # Try to extract element from atom name
                element = ''.join([c for c in atom_name if c.isalpha()])
                if element:
                    # Common element patterns in atom names
                    if element.startswith('C'):
                        atom_data['element'] = 'C'
                    elif element.startswith('N'):
                        atom_data['element'] = 'N'
                    elif element.startswith('O'):
                        atom_data['element'] = 'O'
                    elif element.startswith('S'):
                        atom_data['element'] = 'S'
                    elif element.startswith('P'):
                        atom_data['element'] = 'P'
                    elif element.startswith('H'):
                        atom_data['element'] = 'H'
                    elif element.startswith('F'):
                        atom_data['element'] = 'F'
                    elif element.startswith('Cl'):
                        atom_data['element'] = 'Cl'
                    elif element.startswith('Br'):
                        atom_data['element'] = 'Br'
                    elif element.startswith('I'):
                        atom_data['element'] = 'I'
                    else:
                        atom_data['element'] = element[:2] if len(element) > 1 else element
                else:
                    atom_data['element'] = 'Unknown'
            
            # Handle AutoDock atom types - extract element from autodock type
            if atom_data['autodock_type'] and not atom_data['element']:
                autodock_type = atom_data['autodock_type']
                if autodock_type.startswith('C'):
                    atom_data['element'] = 'C'
                elif autodock_type.startswith('N'):
                    atom_data['element'] = 'N'
                elif autodock_type.startswith('O'):
                    atom_data['element'] = 'O'
                elif autodock_type.startswith('S'):
                    atom_data['element'] = 'S'
                elif autodock_type.startswith('P'):
                    atom_data['element'] = 'P'
                elif autodock_type.startswith('H'):
                    atom_data['element'] = 'H'
                elif autodock_type.startswith('F'):
                    atom_data['element'] = 'F'
                elif autodock_type.startswith('Cl'):
                    atom_data['element'] = 'Cl'
                elif autodock_type.startswith('Br'):
                    atom_data['element'] = 'Br'
                elif autodock_type.startswith('I'):
                    atom_data['element'] = 'I'
            
            return atom_data
            
        except (ValueError, IndexError) as e:
            logger.warning(f"Error parsing atom line {line_num}: {str(e)}")
            return {
                'line_number': line_num,
                'record_type': 'UNKNOWN',
                'atom_serial': 0,
                'atom_name': 'UNK',
                'residue_name': 'UNK',
                'chain_id': '',
                'residue_number': 0,
                'x': 0.0,
                'y': 0.0,
                'z': 0.0,
                'occupancy': 1.0,
                'temperature_factor': 0.0,
                'element': 'Unknown',
                'charge': ''
            }
    
    def _extract_model_number(self, line: str) -> int:
        """Extract model number from MODEL line."""
        try:
            return int(line.split()[1])
        except (ValueError, IndexError):
            return 1
    
    def _validate_parsed_data(self, parsed_data: Dict) -> List[str]:
        """Validate parsed PDBQT data structure."""
        errors = []
        
        # Check for required data
        if not parsed_data['atoms']:
            errors.append("No atoms found in file")
        
        # Note: Models are optional in PDBQT files - some files don't have MODEL/ENDMDL sections
        
        # Validate atom data
        for atom in parsed_data['atoms']:
            if not atom.get('element'):
                errors.append(f"Atom {atom.get('atom_serial', 'unknown')} missing element")
            
            # Only validate against supported atoms if element is not empty
            if atom.get('element') and atom.get('element') not in self.supported_atoms:
                errors.append(f"Unsupported element: {atom.get('element')}")
        
        return errors
```
