## Supplementary material for "Machine Learning-Driven Prediction of TLR4 Binding Affinity: A Comprehensive Molecular Feature Analysis for Drug Discovery": Codebase: test_3d_features.py

```
#!/usr/bin/env python3
"""
Test script for 3D structural feature extraction.

This script tests the StructuralFeatureExtractor with sample PDBQT files
to verify that the implementation works correctly.
"""

import sys
import os
import logging
from pathlib import Path

# Add src to path
sys.path.insert(0, str(Path(__file__).parent.parent.parent / "src"))

from tlr4_binding.molecular_analysis.structure import StructuralFeatureExtractor, PYMOL_AVAILABLE

# Set up logging
logging.basicConfig(level=logging.INFO)
logger = logging.getLogger(__name__)


def test_3d_feature_extraction():
    """Test 3D feature extraction with sample PDBQT files."""
    
    print("=" * 60)
    print("Testing 3D Structural Feature Extraction")
    print("=" * 60)
    
    # Check PyMOL availability
    if not PYMOL_AVAILABLE:
        print("⚠️  PyMOL is not available. Testing fallback mode.")
    else:
        print("✅ PyMOL is available.")
    
    # Initialize extractor
    extractor = StructuralFeatureExtractor()
    
    # Get available features
    features = extractor.get_available_features()
    print(f"\n📊 Available 3D features ({len(features)}):")
    for i, feature in enumerate(features, 1):
        print(f"  {i:2d}. {feature}")
    
    # Look for sample PDBQT files
    sample_dirs = [
        "binding-data/raw/pdbqt",
        "data/raw/pdbqt", 
        "samples/pdbqt"
    ]
    
    sample_files = []
    for sample_dir in sample_dirs:
        if os.path.exists(sample_dir):
            pdbqt_files = list(Path(sample_dir).glob("*.pdbqt"))
            sample_files.extend(pdbqt_files[:3])  # Take first 3 files
            break
    
    if not sample_files:
        print("\n⚠️  No sample PDBQT files found. Creating a test file...")
        
        # Create a simple test PDBQT file
        test_content = """REMARK  Generated by AutoDock Vina
ATOM      1  C   LIG A   1      20.154  30.231  25.123  1.00  0.00      A    C
ATOM      2  N   LIG A   1      21.234  31.345  26.234  1.00  0.00      A    N
ATOM      3  O   LIG A   1      22.345  32.456  27.345  1.00  0.00      A    O
ATOM      4  H   LIG A   1      19.123  29.456  24.789  1.00  0.00      A    H
ATOM      5  H   LIG A   1      20.789  30.123  25.456  1.00  0.00      A    H
ENDMDL
"""
        
        test_file = Path("test_molecule.pdbqt")
        with open(test_file, 'w') as f:
            f.write(test_content)
        sample_files = [test_file]
        print(f"✅ Created test file: {test_file}")
    
    # Test feature extraction
    print(f"\n🧪 Testing feature extraction on {len(sample_files)} files:")
    
    for i, pdbqt_file in enumerate(sample_files, 1):
        print(f"\n--- File {i}: {pdbqt_file.name} ---")
        
        try:
            # Extract features
            features_dict = extractor.extract_features(str(pdbqt_file))
            
            print(f"✅ Successfully extracted {len(features_dict)} features")
            
            # Show some key features
            key_features = [
                'radius_of_gyration', 'molecular_volume', 'surface_area',
                'asphericity', 'eccentricity', 'spherocity_index'
            ]
            
            print("📈 Key 3D features:")
            for feature in key_features:
                if feature in features_dict:
                    value = features_dict[feature]
                    if isinstance(value, float) and not (value != value):  # Check for NaN
                        print(f"  {feature:20s}: {value:8.3f}")
                    else:
                        print(f"  {feature:20s}: {value}")
            
            # Show feature statistics
            valid_features = [v for v in features_dict.values() 
                            if isinstance(v, (int, float)) and not (v != v)]
            
            if valid_features:
                print(f"\n📊 Feature statistics:")
                print(f"  Valid features: {len(valid_features)}/{len(features_dict)}")
                print(f"  Min value: {min(valid_features):.3f}")
                print(f"  Max value: {max(valid_features):.3f}")
                print(f"  Mean value: {sum(valid_features)/len(valid_features):.3f}")
            
        except Exception as e:
            print(f"❌ Error extracting features: {str(e)}")
            logger.error(f"Error processing {pdbqt_file}: {str(e)}")
    
    # Cleanup
    try:
        extractor.cleanup_session()
        print("\n🧹 PyMOL session cleaned up")
    except Exception as e:
        print(f"\n⚠️  Warning during cleanup: {str(e)}")
    
    # Clean up test file if created
    test_file = Path("test_molecule.pdbqt")
    if test_file.exists():
        test_file.unlink()
        print("🗑️  Test file cleaned up")
    
    print("\n" + "=" * 60)
    print("3D Feature Extraction Test Complete")
    print("=" * 60)


if __name__ == "__main__":
    test_3d_feature_extraction()
```
