## Supplementary material for "Machine Learning-Driven Prediction of TLR4 Binding Affinity: A Comprehensive Molecular Feature Analysis for Drug Discovery": Codebase: test_3d_standalone.py

```
#!/usr/bin/env python3
"""
Standalone test script for 3D structural feature extraction.

This script directly tests the StructuralFeatureExtractor by importing
only the specific module without going through the package __init__.py
"""

import sys
import os
import logging
from pathlib import Path

# Add src to path
sys.path.insert(0, str(Path(__file__).parent.parent.parent / "src"))

# Direct import of the specific module
import importlib.util
spec = importlib.util.spec_from_file_location(
    "structure", 
    Path(__file__).parent / "src" / "tlr4_binding" / "molecular_analysis" / "structure.py"
)
structure_module = importlib.util.module_from_spec(spec)
spec.loader.exec_module(structure_module)

StructuralFeatureExtractor = structure_module.StructuralFeatureExtractor
PYMOL_AVAILABLE = structure_module.PYMOL_AVAILABLE

# Set up logging
logging.basicConfig(level=logging.INFO)
logger = logging.getLogger(__name__)


def test_3d_feature_extraction():
    """Test 3D feature extraction with sample PDBQT files."""
    
    print("=" * 60)
    print("Testing 3D Structural Feature Extraction")
    print("=" * 60)
    
    # Check PyMOL availability
    if not PYMOL_AVAILABLE:
        print("⚠️  PyMOL is not available. Testing fallback mode.")
    else:
        print("✅ PyMOL is available.")
    
    # Initialize extractor
    extractor = StructuralFeatureExtractor()
    
    # Get available features
    features = extractor.get_available_features()
    print(f"\n📊 Available 3D features ({len(features)}):")
    for i, feature in enumerate(features, 1):
        print(f"  {i:2d}. {feature}")
    
    # Create a simple test PDBQT file
    test_content = """REMARK  Generated by AutoDock Vina
ATOM      1  C   LIG A   1      20.154  30.231  25.123  1.00  0.00      A    C
ATOM      2  N   LIG A   1      21.234  31.345  26.234  1.00  0.00      A    N
ATOM      3  O   LIG A   1      22.345  32.456  27.345  1.00  0.00      A    O
ATOM      4  H   LIG A   1      19.123  29.456  24.789  1.00  0.00      A    H
ATOM      5  H   LIG A   1      20.789  30.123  25.456  1.00  0.00      A    H
ENDMDL
"""
    
    test_file = Path("test_molecule.pdbqt")
    with open(test_file, 'w') as f:
        f.write(test_content)
    
    print(f"\n🧪 Testing feature extraction on test file: {test_file}")
    
    try:
        # Extract features
        features_dict = extractor.extract_features(str(test_file))
        
        print(f"✅ Successfully extracted {len(features_dict)} features")
        
        # Show some key features
        key_features = [
            'radius_of_gyration', 'molecular_volume', 'surface_area',
            'asphericity', 'eccentricity', 'spherocity_index'
        ]
        
        print("\n📈 Key 3D features:")
        for feature in key_features:
            if feature in features_dict:
                value = features_dict[feature]
                if isinstance(value, float) and not (value != value):  # Check for NaN
                    print(f"  {feature:20s}: {value:8.3f}")
                else:
                    print(f"  {feature:20s}: {value}")
        
        # Show feature statistics
        valid_features = [v for v in features_dict.values() 
                        if isinstance(v, (int, float)) and not (v != v)]
        
        if valid_features:
            print(f"\n📊 Feature statistics:")
            print(f"  Valid features: {len(valid_features)}/{len(features_dict)}")
            print(f"  Min value: {min(valid_features):.3f}")
            print(f"  Max value: {max(valid_features):.3f}")
            print(f"  Mean value: {sum(valid_features)/len(valid_features):.3f}")
        else:
            print("\n⚠️  No valid numeric features extracted (expected in fallback mode)")
        
    except Exception as e:
        print(f"❌ Error extracting features: {str(e)}")
        logger.error(f"Error processing {test_file}: {str(e)}")
    
    # Cleanup
    try:
        extractor.cleanup_session()
        print("\n🧹 PyMOL session cleaned up")
    except Exception as e:
        print(f"\n⚠️  Warning during cleanup: {str(e)}")
    
    # Clean up test file
    if test_file.exists():
        test_file.unlink()
        print("🗑️  Test file cleaned up")
    
    print("\n" + "=" * 60)
    print("3D Feature Extraction Test Complete")
    print("=" * 60)


if __name__ == "__main__":
    test_3d_feature_extraction()
```
