## Supplementary material for "Machine Learning-Driven Prediction of TLR4 Binding Affinity: A Comprehensive Molecular Feature Analysis for Drug Discovery": Codebase: test_compound_matcher.py

```
"""
Unit tests for CompoundMatcher class.

Tests fuzzy string matching functionality for compound name alignment
between different datasets.
"""

import pytest
import pandas as pd
import numpy as np
from unittest.mock import patch, MagicMock
import sys
import os

# Add src to path for imports
sys.path.insert(0, os.path.join(os.path.dirname(__file__), '..', '..', 'src'))

from tlr4_binding.data_processing.preprocessor import CompoundMatcher


class TestCompoundMatcher:
    """Test cases for CompoundMatcher class."""
    
    def setup_method(self):
        """Set up test fixtures."""
        self.matcher = CompoundMatcher(threshold=80.0, use_partial_ratio=True)
        
        # Sample compound names for testing
        self.names1 = [
            "Andrographolide",
            "Curcumin",
            "Resveratrol",
            "Quercetin",
            "Epigallocatechin",
            "Capsaicin",
            "Gingerol",
            "Caffeic acid",
            "Chlorogenic acid",
            "Ferulic acid"
        ]
        
        self.names2 = [
            "andrographolide",  # Case difference
            "curcumin_compound",  # Suffix difference
            "resveratrol_molecule",  # Suffix difference
            "quercetin_drug",  # Suffix difference
            "epigallocatechin_gallate",  # Additional word
            "capsaicin_ligand",  # Suffix difference
            "gingerol_bound",  # Suffix difference
            "caffeic_acid",  # Underscore instead of space
            "chlorogenic_acid_compound",  # Multiple differences
            "ferulic_acid_molecule"  # Multiple differences
        ]
        
        self.names_no_match = [
            "completely_different_compound",
            "totally_unrelated_molecule",
            "xyz_abc_123"
        ]
    
    def test_initialization(self):
        """Test CompoundMatcher initialization."""
        matcher = CompoundMatcher(threshold=75.0, use_partial_ratio=False)
        assert matcher.threshold == 75.0
        assert matcher.use_partial_ratio == False
        assert matcher.match_cache == {}
    
    def test_initialization_defaults(self):
        """Test CompoundMatcher initialization with defaults."""
        matcher = CompoundMatcher()
        assert matcher.threshold == 80.0
        assert matcher.use_partial_ratio == True
        assert matcher.match_cache == {}
    
    def test_clean_name_basic(self):
        """Test basic name cleaning functionality."""
        test_cases = [
            ("Compound_123", "123"),  # Underscore removed, then prefix "compound" removed
            ("LIGAND_NAME", "name"),  # Underscore removed, then prefix "ligand" removed
            ("Molecule-ABC", "abc"),  # Hyphen removed, then prefix "molecule" removed
            ("Drug@Special#Chars", "specialchars"),  # Special chars removed, then prefix "drug" removed
            ("  Extra  Spaces  ", "extra spaces"),
            ("", ""),
            (None, "None")
        ]
        
        for input_name, expected in test_cases:
            result = self.matcher._clean_name(input_name)
            assert result == expected
    
    def test_clean_name_prefix_removal(self):
        """Test removal of common prefixes."""
        test_cases = [
            ("compound_andrographolide", "andrographolide"),
            ("ligand_curcumin", "curcumin"),
            ("molecule_resveratrol", "resveratrol"),
            ("drug_quercetin", "quercetin"),
            ("compound_compound_test", "compoundtest"),  # Only first prefix removed, underscore removed
        ]
        
        for input_name, expected in test_cases:
            result = self.matcher._clean_name(input_name)
            assert result == expected
    
    def test_clean_name_suffix_removal(self):
        """Test removal of common suffixes."""
        test_cases = [
            ("andrographolide_docked", "andrographolide"),
            ("curcumin_bound", "curcumin"),
            ("resveratrol_complex", "resveratrol"),
            ("test_docked_bound", "testdocked"),  # Only first suffix removed, underscore removed
        ]
        
        for input_name, expected in test_cases:
            result = self.matcher._clean_name(input_name)
            assert result == expected
    
    def test_clean_name_special_characters(self):
        """Test removal of special characters."""
        test_cases = [
            ("compound-123", "123"),  # Special chars removed, then prefix "compound" removed
            ("ligand@name", "name"),  # Special chars removed, then prefix "ligand" removed
            ("molecule#test", "test"),  # Special chars removed, then prefix "molecule" removed
            ("drug$special", "special"),  # Special chars removed, then prefix "drug" removed
            ("test%chars", "testchars"),  # Special chars removed
        ]
        
        for input_name, expected in test_cases:
            result = self.matcher._clean_name(input_name)
            assert result == expected
    
    def test_find_best_match_exact(self):
        """Test finding exact matches."""
        candidates = ["exact_match", "other_compound", "another_one"]
        target = "exact_match"
        
        result = self.matcher._find_best_match(target, candidates)
        assert result == "exact_match"
    
    def test_find_best_match_fuzzy(self):
        """Test finding fuzzy matches."""
        candidates = ["andrographolide", "curcumin", "resveratrol"]
        target = "andrographolide_compound"
        
        result = self.matcher._find_best_match(target, candidates)
        assert result == "andrographolide"
    
    def test_find_best_match_no_match(self):
        """Test when no match is found."""
        candidates = ["completely_different", "totally_unrelated"]
        target = "no_match_here"
        
        result = self.matcher._find_best_match(target, candidates)
        assert result is None
    
    def test_find_best_match_empty_candidates(self):
        """Test with empty candidate list."""
        candidates = []
        target = "any_target"
        
        result = self.matcher._find_best_match(target, candidates)
        assert result is None
    
    def test_find_best_match_threshold(self):
        """Test threshold-based matching."""
        # Create matcher with high threshold
        high_threshold_matcher = CompoundMatcher(threshold=95.0)
        
        candidates = ["andrographolide", "curcumin"]
        target = "andrographolide_compound"  # Should match but with lower score
        
        result = high_threshold_matcher._find_best_match(target, candidates)
        # Should still match because partial ratio should be high enough
        assert result == "andrographolide"
    
    def test_match_compounds_successful_matches(self):
        """Test successful compound matching."""
        matches = self.matcher.match_compounds(self.names1, self.names2)
        
        # Should have matches for most compounds
        assert len(matches) > 0
        
        # Check specific expected matches
        assert "Andrographolide" in matches
        assert "Curcumin" in matches
        assert "Resveratrol" in matches
    
    def test_match_compounds_no_matches(self):
        """Test matching with no possible matches."""
        matches = self.matcher.match_compounds(self.names_no_match, self.names2)
        
        # Should have no matches
        assert len(matches) == 0
    
    def test_match_compounds_empty_lists(self):
        """Test matching with empty lists."""
        matches = self.matcher.match_compounds([], self.names2)
        assert len(matches) == 0
        
        matches = self.matcher.match_compounds(self.names1, [])
        assert len(matches) == 0
    
    def test_match_compounds_caching(self):
        """Test that matches are cached."""
        # First call
        matches1 = self.matcher.match_compounds(self.names1[:3], self.names2)
        
        # Second call should use cache
        matches2 = self.matcher.match_compounds(self.names1[:3], self.names2)
        
        # Results should be identical
        assert matches1 == matches2
        
        # Cache should contain entries
        assert len(self.matcher.match_cache) > 0
    
    def test_get_match_confidence(self):
        """Test getting match confidence scores."""
        # Test with exact match
        confidence = self.matcher.get_match_confidence("test", "test")
        assert confidence == 100.0
        
        # Test with partial match
        confidence = self.matcher.get_match_confidence("andrographolide", "andrographolide_compound")
        assert confidence > 80.0  # Should be high due to partial ratio
        
        # Test with no match
        confidence = self.matcher.get_match_confidence("completely_different", "totally_unrelated")
        assert confidence < 50.0  # Should be low
    
    def test_get_match_confidence_partial_vs_ratio(self):
        """Test confidence calculation with different ratio methods."""
        matcher_partial = CompoundMatcher(use_partial_ratio=True)
        matcher_ratio = CompoundMatcher(use_partial_ratio=False)
        
        name1 = "andrographolide"
        name2 = "andrographolide_compound_extra_long_name"
        
        conf_partial = matcher_partial.get_match_confidence(name1, name2)
        conf_ratio = matcher_ratio.get_match_confidence(name1, name2)
        
        # Partial ratio should be higher for this case
        assert conf_partial > conf_ratio
    
    def test_match_compounds_with_special_characters(self):
        """Test matching with special characters in names."""
        names_with_special = [
            "Compound-123",
            "Ligand@Name",
            "Molecule#Test"
        ]
        
        clean_names = [
            "compound 123",
            "ligand name", 
            "molecule test"
        ]
        
        matches = self.matcher.match_compounds(names_with_special, clean_names)
        
        # Should find matches despite special characters
        assert len(matches) > 0
    
    def test_match_compounds_case_insensitive(self):
        """Test that matching is case insensitive."""
        upper_names = ["ANDROGRAPHOLIDE", "CURCUMIN", "RESVERATROL"]
        lower_names = ["andrographolide", "curcumin", "resveratrol"]
        
        matches = self.matcher.match_compounds(upper_names, lower_names)
        
        # Should find matches despite case differences
        assert len(matches) == len(upper_names)
        for name in upper_names:
            assert name in matches
    
    def test_match_compounds_duplicate_candidates(self):
        """Test matching with duplicate candidate names."""
        candidates_with_duplicates = ["andrographolide", "curcumin", "andrographolide", "resveratrol"]
        targets = ["Andrographolide", "Curcumin"]
        
        matches = self.matcher.match_compounds(targets, candidates_with_duplicates)
        
        # Should still work with duplicates
        assert len(matches) > 0
    
    def test_match_compounds_very_similar_names(self):
        """Test matching with very similar but different names."""
        similar_names = [
            "andrographolide",
            "andrographolide_derivative",
            "andrographolide_analog",
            "andrographolide_modified"
        ]
        
        target = "Andrographolide"
        
        matches = self.matcher.match_compounds([target], similar_names)
        
        # Should match the exact one
        assert target in matches
        assert matches[target] == "andrographolide"
    
    def test_match_compounds_performance(self):
        """Test matching performance with larger datasets."""
        # Create larger test datasets
        large_names1 = [f"compound_{i}" for i in range(100)]
        large_names2 = [f"compound_{i}_modified" for i in range(100)]
        
        matches = self.matcher.match_compounds(large_names1, large_names2)
        
        # Should find matches for most compounds
        assert len(matches) > 50  # At least half should match
    
    def test_error_handling_invalid_inputs(self):
        """Test error handling with invalid inputs."""
        # Test with None values - should return "None" string
        result = self.matcher._clean_name(None)
        assert result == "None"
        
        # Test with non-string inputs
        result = self.matcher._clean_name(123)
        assert result == "123"
        
        result = self.matcher._clean_name(123.45)
        assert result == "123.45"
    
    def test_match_compounds_with_whitespace(self):
        """Test matching with various whitespace patterns."""
        names_with_whitespace = [
            "  andrographolide  ",
            "\tcurcumin\t",
            "\nresveratrol\n",
            "  compound  with  spaces  "
        ]
        
        clean_names = [
            "andrographolide",
            "curcumin",
            "resveratrol",
            "compound with spaces"
        ]
        
        matches = self.matcher.match_compounds(names_with_whitespace, clean_names)
        
        # Should find matches despite whitespace
        assert len(matches) > 0


if __name__ == "__main__":
    pytest.main([__file__, "-v"])
```
