## Supplementary material for "Machine Learning-Driven Prediction of TLR4 Binding Affinity: A Comprehensive Molecular Feature Analysis for Drug Discovery": Codebase: test_data_integrator.py

```
"""
Unit tests for DataIntegrator class.

Tests data integration functionality for combining molecular features
with binding affinity data.
"""

import pytest
import pandas as pd
import numpy as np
from unittest.mock import patch, MagicMock
import sys
import os

# Add src to path for imports
sys.path.insert(0, os.path.join(os.path.dirname(__file__), '..', '..', 'src'))

from tlr4_binding.data_processing.preprocessor import DataIntegrator, CompoundMatcher


class TestDataIntegrator:
    """Test cases for DataIntegrator class."""
    
    def setup_method(self):
        """Set up test fixtures."""
        self.compound_matcher = CompoundMatcher(threshold=80.0)
        self.integrator = DataIntegrator(self.compound_matcher)
        
        # Sample molecular features data
        self.features_df = pd.DataFrame({
            'compound_name': [
                'Andrographolide',
                'Curcumin', 
                'Resveratrol',
                'Quercetin',
                'Epigallocatechin',
                'Capsaicin',
                'Gingerol',
                'Caffeic acid',
                'Chlorogenic acid',
                'Ferulic acid'
            ],
            'molecular_weight': [350.4, 368.4, 228.2, 302.2, 458.4, 305.4, 276.4, 180.2, 354.3, 194.2],
            'logp': [2.3, 2.5, 3.1, 1.8, 2.0, 3.8, 2.9, 1.2, 0.8, 1.4],
            'tpsa': [74.6, 93.1, 60.7, 131.4, 197.4, 58.6, 50.5, 77.8, 164.7, 66.8],
            'hbd': [3, 2, 3, 5, 8, 1, 1, 2, 4, 2],
            'hba': [4, 6, 3, 7, 11, 2, 2, 4, 7, 3]
        })
        
        # Sample binding data
        self.binding_df = pd.DataFrame({
            'ligand': [
                'andrographolide',  # Case difference
                'curcumin_compound',  # Suffix difference
                'resveratrol_molecule',  # Suffix difference
                'quercetin_drug',  # Suffix difference
                'epigallocatechin_gallate',  # Additional word
                'capsaicin_ligand',  # Suffix difference
                'gingerol_bound',  # Suffix difference
                'caffeic_acid',  # Underscore instead of space
                'chlorogenic_acid_compound',  # Multiple differences
                'ferulic_acid_molecule'  # Multiple differences
            ],
            'mode': [1, 1, 1, 1, 1, 1, 1, 1, 1, 1],
            'affinity': [-7.2, -6.8, -6.5, -6.9, -7.1, -6.3, -6.7, -5.8, -6.2, -5.9],
            'rmsd_lb': [0.0, 0.0, 0.0, 0.0, 0.0, 0.0, 0.0, 0.0, 0.0, 0.0],
            'rmsd_ub': [0.0, 0.0, 0.0, 0.0, 0.0, 0.0, 0.0, 0.0, 0.0, 0.0]
        })
        
        # Data with no matches
        self.no_match_features = pd.DataFrame({
            'compound_name': ['Unknown1', 'Unknown2', 'Unknown3'],
            'molecular_weight': [100.0, 200.0, 300.0],
            'logp': [1.0, 2.0, 3.0]
        })
        
        self.no_match_binding = pd.DataFrame({
            'ligand': ['Different1', 'Different2', 'Different3'],
            'mode': [1, 1, 1],
            'affinity': [-5.0, -6.0, -7.0],
            'rmsd_lb': [0.0, 0.0, 0.0],
            'rmsd_ub': [0.0, 0.0, 0.0]
        })
    
    def test_initialization(self):
        """Test DataIntegrator initialization."""
        integrator = DataIntegrator()
        assert integrator.compound_matcher is not None
        assert integrator.integration_stats == {}
    
    def test_initialization_with_matcher(self):
        """Test DataIntegrator initialization with custom matcher."""
        custom_matcher = CompoundMatcher(threshold=90.0)
        integrator = DataIntegrator(custom_matcher)
        assert integrator.compound_matcher == custom_matcher
    
    def test_integrate_datasets_successful(self):
        """Test successful dataset integration."""
        result_df = self.integrator.integrate_datasets(
            self.features_df, 
            self.binding_df
        )
        
        # Should have integrated records
        assert len(result_df) > 0
        
        # Should contain both feature and binding columns
        expected_columns = set(self.features_df.columns) | set(self.binding_df.columns)
        assert set(result_df.columns) >= expected_columns
        
        # Should have matched compound names
        assert 'matched_compound' in result_df.columns
    
    def test_integrate_datasets_no_matches(self):
        """Test integration with no matching compounds."""
        result_df = self.integrator.integrate_datasets(
            self.no_match_features,
            self.no_match_binding
        )
        
        # Should be empty due to no matches
        assert len(result_df) == 0
    
    def test_integrate_datasets_partial_matches(self):
        """Test integration with partial matches."""
        # Mix of matching and non-matching compounds
        partial_features = pd.concat([
            self.features_df.head(3),  # Some matches
            self.no_match_features.head(2)  # Some non-matches
        ]).reset_index(drop=True)
        
        result_df = self.integrator.integrate_datasets(
            partial_features,
            self.binding_df
        )
        
        # Should have some integrated records
        assert len(result_df) > 0
        assert len(result_df) < len(partial_features)  # Not all should match
    
    def test_integrate_datasets_custom_columns(self):
        """Test integration with custom column names."""
        # Rename columns
        features_renamed = self.features_df.rename(columns={'compound_name': 'molecule_name'})
        binding_renamed = self.binding_df.rename(columns={'ligand': 'compound_id'})
        
        result_df = self.integrator.integrate_datasets(
            features_renamed,
            binding_renamed,
            feature_compound_col='molecule_name',
            binding_compound_col='compound_id'
        )
        
        # Should still work with custom column names
        assert len(result_df) > 0
        assert 'molecule_name' in result_df.columns
        assert 'compound_id' in result_df.columns
    
    def test_integrate_datasets_empty_dataframes(self):
        """Test integration with empty DataFrames."""
        empty_features = pd.DataFrame(columns=['compound_name', 'molecular_weight'])
        empty_binding = pd.DataFrame(columns=['ligand', 'affinity', 'mode', 'rmsd_lb', 'rmsd_ub'])
        
        result_df = self.integrator.integrate_datasets(empty_features, empty_binding)
        
        # Should return empty DataFrame
        assert len(result_df) == 0
    
    def test_integrate_datasets_missing_columns(self):
        """Test integration with missing required columns."""
        # Missing compound name column in features
        features_no_compound = self.features_df.drop(columns=['compound_name'])
        
        with pytest.raises(KeyError):
            self.integrator.integrate_datasets(features_no_compound, self.binding_df)
        
        # Missing ligand column in binding data
        binding_no_ligand = self.binding_df.drop(columns=['ligand'])
        
        with pytest.raises(KeyError):
            self.integrator.integrate_datasets(self.features_df, binding_no_ligand)
    
    def test_integration_stats(self):
        """Test integration statistics tracking."""
        # Perform integration
        result_df = self.integrator.integrate_datasets(self.features_df, self.binding_df)
        
        # Get statistics
        stats = self.integrator.get_integration_stats()
        
        # Check that statistics are populated
        assert 'total_features' in stats
        assert 'total_binding' in stats
        assert 'successful_matches' in stats
        assert 'integrated_records' in stats
        assert 'match_rate' in stats
        
        # Check that values are reasonable
        assert stats['total_features'] == len(self.features_df)
        assert stats['total_binding'] == len(self.binding_df)
        assert stats['integrated_records'] == len(result_df)
        assert 0 <= stats['match_rate'] <= 1
    
    def test_integration_stats_no_matches(self):
        """Test integration statistics with no matches."""
        result_df = self.integrator.integrate_datasets(
            self.no_match_features,
            self.no_match_binding
        )
        
        stats = self.integrator.get_integration_stats()
        
        # Should have zero matches
        assert stats['successful_matches'] == 0
        assert stats['integrated_records'] == 0
        assert stats['match_rate'] == 0.0
    
    def test_integration_stats_multiple_calls(self):
        """Test that statistics are updated with multiple integration calls."""
        # First integration
        self.integrator.integrate_datasets(self.features_df, self.binding_df)
        stats1 = self.integrator.get_integration_stats()
        
        # Second integration with different data
        self.integrator.integrate_datasets(self.no_match_features, self.no_match_binding)
        stats2 = self.integrator.get_integration_stats()
        
        # Statistics should be updated
        assert stats1['total_features'] != stats2['total_features']
        assert stats1['integrated_records'] != stats2['integrated_records']
    
    def test_integration_preserves_data_types(self):
        """Test that integration preserves data types."""
        result_df = self.integrator.integrate_datasets(self.features_df, self.binding_df)
        
        # Check that numeric columns maintain their types
        assert pd.api.types.is_numeric_dtype(result_df['molecular_weight'])
        assert pd.api.types.is_numeric_dtype(result_df['logp'])
        assert pd.api.types.is_numeric_dtype(result_df['affinity'])
        
        # Check that string columns maintain their types
        assert pd.api.types.is_object_dtype(result_df['compound_name'])
        assert pd.api.types.is_object_dtype(result_df['ligand'])
    
    def test_integration_handles_duplicates(self):
        """Test integration with duplicate compound names."""
        # Add duplicate compound to features
        features_with_duplicates = pd.concat([
            self.features_df,
            self.features_df.iloc[[0]]  # Duplicate first row
        ]).reset_index(drop=True)
        
        result_df = self.integrator.integrate_datasets(
            features_with_duplicates,
            self.binding_df
        )
        
        # Should handle duplicates gracefully
        assert len(result_df) > 0
        
        # Check for duplicate compound names in result
        duplicate_compounds = result_df['compound_name'].duplicated().sum()
        assert duplicate_compounds >= 0  # May or may not have duplicates depending on matching
    
    def test_integration_handles_missing_values(self):
        """Test integration with missing values in data."""
        # Add missing values to features
        features_with_nulls = self.features_df.copy()
        features_with_nulls.loc[0, 'molecular_weight'] = np.nan
        features_with_nulls.loc[1, 'logp'] = np.nan
        
        # Add missing values to binding data
        binding_with_nulls = self.binding_df.copy()
        binding_with_nulls.loc[0, 'affinity'] = np.nan
        
        result_df = self.integrator.integrate_datasets(
            features_with_nulls,
            binding_with_nulls
        )
        
        # Should still work with missing values
        assert len(result_df) > 0
    
    def test_integration_with_large_datasets(self):
        """Test integration performance with larger datasets."""
        # Create larger test datasets
        large_features = pd.concat([self.features_df] * 10).reset_index(drop=True)
        large_binding = pd.concat([self.binding_df] * 10).reset_index(drop=True)
        
        # Modify names to avoid exact duplicates
        large_features['compound_name'] = [
            f"{name}_{i//10}" for i, name in enumerate(large_features['compound_name'])
        ]
        large_binding['ligand'] = [
            f"{name}_{i//10}" for i, name in enumerate(large_binding['ligand'])
        ]
        
        result_df = self.integrator.integrate_datasets(large_features, large_binding)
        
        # Should handle larger datasets
        assert len(result_df) > 0
        assert len(result_df) <= len(large_features)
    
    def test_integration_column_name_conflicts(self):
        """Test integration with conflicting column names."""
        # Create data with conflicting column names
        features_conflict = self.features_df.copy()
        features_conflict['affinity'] = 0.0  # Same column name as binding data
        
        result_df = self.integrator.integrate_datasets(features_conflict, self.binding_df)
        
        # Should handle column name conflicts
        assert len(result_df) > 0
        
        # Should have both affinity columns (with suffixes)
        affinity_columns = [col for col in result_df.columns if 'affinity' in col]
        assert len(affinity_columns) >= 1
    
    def test_integration_with_different_data_types(self):
        """Test integration with different data types in compound names."""
        # Features with string compound names
        features_str = self.features_df.copy()
        features_str['compound_name'] = features_str['compound_name'].astype(str)
        
        # Binding data with numeric compound names (converted to string)
        binding_numeric = self.binding_df.copy()
        binding_numeric['ligand'] = [f"compound_{i}" for i in range(len(binding_numeric))]
        
        result_df = self.integrator.integrate_datasets(features_str, binding_numeric)
        
        # Should handle different data types
        assert len(result_df) >= 0  # May or may not have matches
    
    def test_integration_merge_strategy(self):
        """Test that integration uses inner join strategy."""
        # Create data where only some compounds match
        partial_features = self.features_df.head(3)
        partial_binding = self.binding_df.head(5)  # More binding records than features
        
        result_df = self.integrator.integrate_datasets(partial_features, partial_binding)
        
        # Should only include compounds that exist in both datasets
        assert len(result_df) <= min(len(partial_features), len(partial_binding))
    
    def test_integration_preserves_index(self):
        """Test that integration preserves meaningful index information."""
        # Set custom index
        features_indexed = self.features_df.set_index('compound_name')
        binding_indexed = self.binding_df.set_index('ligand')
        
        result_df = self.integrator.integrate_datasets(
            features_indexed.reset_index(),
            binding_indexed.reset_index()
        )
        
        # Should work with custom indices
        assert len(result_df) > 0
    
    def test_error_handling_invalid_inputs(self):
        """Test error handling with invalid inputs."""
        # Test with None inputs
        with pytest.raises((AttributeError, TypeError)):
            self.integrator.integrate_datasets(None, self.binding_df)
        
        with pytest.raises((AttributeError, TypeError)):
            self.integrator.integrate_datasets(self.features_df, None)
        
        # Test with non-DataFrame inputs
        with pytest.raises((AttributeError, TypeError)):
            self.integrator.integrate_datasets([], self.binding_df)
    
    def test_integration_with_unicode_names(self):
        """Test integration with unicode compound names."""
        unicode_features = pd.DataFrame({
            'compound_name': ['α-andrographolide', 'β-curcumin', 'γ-resveratrol'],
            'molecular_weight': [350.4, 368.4, 228.2],
            'logp': [2.3, 2.5, 3.1]
        })
        
        unicode_binding = pd.DataFrame({
            'ligand': ['α-andrographolide', 'β-curcumin', 'γ-resveratrol'],
            'mode': [1, 1, 1],
            'affinity': [-7.2, -6.8, -6.5]
        })
        
        result_df = self.integrator.integrate_datasets(unicode_features, unicode_binding)
        
        # Should handle unicode names
        assert len(result_df) > 0


if __name__ == "__main__":
    pytest.main([__file__, "-v"])
```
