## Supplementary material for "Machine Learning-Driven Prediction of TLR4 Binding Affinity: A Comprehensive Molecular Feature Analysis for Drug Discovery": Codebase: test_extractor_simple.py

```
#!/usr/bin/env python3
"""
Simple test script for the molecular feature extractor without external dependencies.

This script tests the core functionality of the MolecularFeatureExtractor
by importing only the necessary modules directly.
"""

import sys
import logging
from pathlib import Path

# Add src to path
sys.path.insert(0, str(Path(__file__).parent.parent.parent / "src"))

# Set up logging
logging.basicConfig(
    level=logging.INFO,
    format='%(asctime)s - %(levelname)s - %(message)s'
)
logger = logging.getLogger(__name__)


def test_extractor_import():
    """Test that the extractor can be imported."""
    logger.info("=== Testing Extractor Import ===")
    
    try:
        # Import only the extractor module directly
        from tlr4_binding.molecular_analysis.extractor import MolecularFeatureExtractor
        logger.info("✓ MolecularFeatureExtractor imported successfully")
        
        # Test instantiation
        extractor = MolecularFeatureExtractor()
        logger.info("✓ MolecularFeatureExtractor instantiated successfully")
        
        # Test basic methods exist
        assert hasattr(extractor, 'extract_features')
        assert hasattr(extractor, 'batch_extract')
        assert hasattr(extractor, 'extract_features_from_list')
        assert hasattr(extractor, 'get_extraction_stats')
        assert hasattr(extractor, 'get_performance_report')
        assert hasattr(extractor, 'validate_features')
        
        logger.info("✓ All expected methods are present")
        
        # Test statistics structure
        stats = extractor.get_extraction_stats()
        expected_keys = [
            'total_files_processed', 'successful_extractions', 'failed_extractions',
            'extraction_errors', 'processing_times', 'feature_extraction_times',
            'memory_usage', 'start_time', 'end_time'
        ]
        
        for key in expected_keys:
            assert key in stats, f"Missing key in stats: {key}"
        
        logger.info("✓ Statistics structure is correct")
        
        return True
        
    except Exception as e:
        logger.error(f"✗ Import test failed: {str(e)}")
        return False


def test_extractor_initialization():
    """Test different initialization modes."""
    logger.info("=== Testing Extractor Initialization ===")
    
    try:
        from tlr4_binding.molecular_analysis.extractor import MolecularFeatureExtractor
        
        # Test default initialization
        extractor1 = MolecularFeatureExtractor()
        assert extractor1.include_2d_features is True
        assert extractor1.include_3d_features is True
        assert extractor1.include_advanced_features is True
        logger.info("✓ Default initialization works")
        
        # Test custom initialization
        extractor2 = MolecularFeatureExtractor(
            include_2d_features=False,
            include_3d_features=True,
            include_advanced_features=False
        )
        assert extractor2.include_2d_features is False
        assert extractor2.include_3d_features is True
        assert extractor2.include_advanced_features is False
        logger.info("✓ Custom initialization works")
        
        # Test minimal initialization
        extractor3 = MolecularFeatureExtractor(
            include_2d_features=False,
            include_3d_features=False,
            include_advanced_features=False
        )
        assert extractor3.include_2d_features is False
        assert extractor3.include_3d_features is False
        assert extractor3.include_advanced_features is False
        logger.info("✓ Minimal initialization works")
        
        return True
        
    except Exception as e:
        logger.error(f"✗ Initialization test failed: {str(e)}")
        return False


def test_error_handling():
    """Test error handling with invalid inputs."""
    logger.info("=== Testing Error Handling ===")
    
    try:
        from tlr4_binding.molecular_analysis.extractor import MolecularFeatureExtractor
        
        extractor = MolecularFeatureExtractor()
        
        # Test with non-existent file
        try:
            extractor.extract_features("non_existent_file.pdbqt")
            logger.error("✗ Should have raised ValueError for non-existent file")
            return False
        except ValueError:
            logger.info("✓ Correctly raised ValueError for non-existent file")
        
        # Test with non-existent directory
        try:
            extractor.batch_extract("non_existent_directory")
            logger.error("✗ Should have raised ValueError for non-existent directory")
            return False
        except ValueError:
            logger.info("✓ Correctly raised ValueError for non-existent directory")
        
        return True
        
    except Exception as e:
        logger.error(f"✗ Error handling test failed: {str(e)}")
        return False


def test_statistics_tracking():
    """Test statistics tracking functionality."""
    logger.info("=== Testing Statistics Tracking ===")
    
    try:
        from tlr4_binding.molecular_analysis.extractor import MolecularFeatureExtractor
        
        extractor = MolecularFeatureExtractor()
        
        # Test reset_stats
        extractor.reset_stats()
        stats = extractor.get_extraction_stats()
        assert stats['total_files_processed'] == 0
        assert stats['successful_extractions'] == 0
        assert stats['failed_extractions'] == 0
        assert len(stats['extraction_errors']) == 0
        logger.info("✓ Statistics reset correctly")
        
        # Test performance report generation (empty)
        performance_report = extractor.get_performance_report()
        assert isinstance(performance_report, dict)
        assert 'batch_summary' in performance_report
        logger.info("✓ Performance report generation works")
        
        # Test efficiency calculation (empty)
        efficiency = extractor.get_feature_extraction_efficiency()
        assert isinstance(efficiency, dict)
        logger.info("✓ Efficiency calculation works")
        
        return True
        
    except Exception as e:
        logger.error(f"✗ Statistics tracking test failed: {str(e)}")
        return False


def test_feature_validation_structure():
    """Test feature validation structure without actual data."""
    logger.info("=== Testing Feature Validation Structure ===")
    
    try:
        from tlr4_binding.molecular_analysis.extractor import MolecularFeatureExtractor
        import pandas as pd
        import numpy as np
        
        extractor = MolecularFeatureExtractor()
        
        # Create mock feature DataFrame
        mock_data = {
            'compound_name': ['test1', 'test2'],
            'molecular_weight': [100.0, np.nan],
            'logp': [1.5, 2.0],
            'tpsa': [50.0, np.inf],
            'rotatable_bonds': [3, 5]
        }
        mock_df = pd.DataFrame(mock_data)
        
        # Test validation
        validation_results = extractor.validate_features(mock_df)
        
        # Check structure
        expected_keys = ['missing_values', 'infinite_values', 'outliers', 'invalid_ranges']
        for key in expected_keys:
            assert key in validation_results
            assert isinstance(validation_results[key], list)
        
        logger.info("✓ Feature validation structure is correct")
        
        # Test feature summary
        summary_df = extractor.get_feature_summary(mock_df)
        assert isinstance(summary_df, pd.DataFrame)
        assert 'feature' in summary_df.columns
        assert 'count' in summary_df.columns
        
        logger.info("✓ Feature summary generation works")
        
        return True
        
    except Exception as e:
        logger.error(f"✗ Feature validation structure test failed: {str(e)}")
        return False


def main():
    """Run all simple tests."""
    logger.info("Starting Simple MolecularFeatureExtractor Tests")
    
    tests = [
        test_extractor_import,
        test_extractor_initialization,
        test_error_handling,
        test_statistics_tracking,
        test_feature_validation_structure
    ]
    
    passed = 0
    failed = 0
    
    for test_func in tests:
        try:
            if test_func():
                passed += 1
                logger.info(f"✓ {test_func.__name__} PASSED")
            else:
                failed += 1
                logger.error(f"✗ {test_func.__name__} FAILED")
        except Exception as e:
            failed += 1
            logger.error(f"✗ {test_func.__name__} FAILED with exception: {str(e)}")
        
        logger.info("-" * 60)
    
    # Summary
    logger.info(f"Simple Tests Summary:")
    logger.info(f"  Passed: {passed}")
    logger.info(f"  Failed: {failed}")
    logger.info(f"  Total: {passed + failed}")
    
    if failed == 0:
        logger.info("🎉 All simple tests passed!")
        return True
    else:
        logger.error(f"❌ {failed} simple tests failed!")
        return False


if __name__ == "__main__":
    success = main()
    sys.exit(0 if success else 1)
```
