## Supplementary material for "Machine Learning-Driven Prediction of TLR4 Binding Affinity: A Comprehensive Molecular Feature Analysis for Drug Discovery": Codebase: test_extractor_standalone.py

```
#!/usr/bin/env python3
"""
Standalone test for the molecular feature extractor.

This script tests the MolecularFeatureExtractor by importing it directly
without going through the full package initialization.
"""

import sys
import logging
from pathlib import Path

# Set up logging
logging.basicConfig(
    level=logging.INFO,
    format='%(asctime)s - %(levelname)s - %(message)s'
)
logger = logging.getLogger(__name__)


def test_direct_import():
    """Test direct import of the extractor module."""
    logger.info("=== Testing Direct Import ===")
    
    try:
        # Import directly from the module file
        sys.path.insert(0, str(Path(__file__).parent.parent.parent / "src" / "tlr4_binding" / "molecular_analysis"))
        
        from extractor import MolecularFeatureExtractor
        logger.info("✓ MolecularFeatureExtractor imported successfully")
        
        # Test instantiation
        extractor = MolecularFeatureExtractor()
        logger.info("✓ MolecularFeatureExtractor instantiated successfully")
        
        # Test basic methods exist
        methods = [
            'extract_features', 'batch_extract', 'extract_features_from_list',
            'get_extraction_stats', 'get_performance_report', 'validate_features',
            'get_feature_summary', 'reset_stats'
        ]
        
        for method in methods:
            assert hasattr(extractor, method), f"Missing method: {method}"
        
        logger.info("✓ All expected methods are present")
        
        # Test statistics structure
        stats = extractor.get_extraction_stats()
        expected_keys = [
            'total_files_processed', 'successful_extractions', 'failed_extractions',
            'extraction_errors', 'processing_times', 'feature_extraction_times',
            'memory_usage', 'start_time', 'end_time'
        ]
        
        for key in expected_keys:
            assert key in stats, f"Missing key in stats: {key}"
        
        logger.info("✓ Statistics structure is correct")
        
        return True
        
    except Exception as e:
        logger.error(f"✗ Direct import test failed: {str(e)}")
        return False


def test_initialization_modes():
    """Test different initialization modes."""
    logger.info("=== Testing Initialization Modes ===")
    
    try:
        sys.path.insert(0, str(Path(__file__).parent.parent.parent / "src" / "tlr4_binding" / "molecular_analysis"))
        from extractor import MolecularFeatureExtractor
        
        # Test default initialization
        extractor1 = MolecularFeatureExtractor()
        assert extractor1.include_2d_features is True
        assert extractor1.include_3d_features is True
        assert extractor1.include_advanced_features is True
        logger.info("✓ Default initialization works")
        
        # Test custom initialization
        extractor2 = MolecularFeatureExtractor(
            include_2d_features=False,
            include_3d_features=True,
            include_advanced_features=False
        )
        assert extractor2.include_2d_features is False
        assert extractor2.include_3d_features is True
        assert extractor2.include_advanced_features is False
        logger.info("✓ Custom initialization works")
        
        return True
        
    except Exception as e:
        logger.error(f"✗ Initialization test failed: {str(e)}")
        return False


def test_error_handling():
    """Test error handling with invalid inputs."""
    logger.info("=== Testing Error Handling ===")
    
    try:
        sys.path.insert(0, str(Path(__file__).parent.parent.parent / "src" / "tlr4_binding" / "molecular_analysis"))
        from extractor import MolecularFeatureExtractor
        
        extractor = MolecularFeatureExtractor()
        
        # Test with non-existent file
        try:
            extractor.extract_features("non_existent_file.pdbqt")
            logger.error("✗ Should have raised ValueError for non-existent file")
            return False
        except ValueError:
            logger.info("✓ Correctly raised ValueError for non-existent file")
        
        # Test with non-existent directory
        try:
            extractor.batch_extract("non_existent_directory")
            logger.error("✗ Should have raised ValueError for non-existent directory")
            return False
        except ValueError:
            logger.info("✓ Correctly raised ValueError for non-existent directory")
        
        return True
        
    except Exception as e:
        logger.error(f"✗ Error handling test failed: {str(e)}")
        return False


def test_statistics_functionality():
    """Test statistics and performance tracking."""
    logger.info("=== Testing Statistics Functionality ===")
    
    try:
        sys.path.insert(0, str(Path(__file__).parent.parent.parent / "src" / "tlr4_binding" / "molecular_analysis"))
        from extractor import MolecularFeatureExtractor
        
        extractor = MolecularFeatureExtractor()
        
        # Test reset_stats
        extractor.reset_stats()
        stats = extractor.get_extraction_stats()
        assert stats['total_files_processed'] == 0
        assert stats['successful_extractions'] == 0
        assert stats['failed_extractions'] == 0
        assert len(stats['extraction_errors']) == 0
        logger.info("✓ Statistics reset correctly")
        
        # Test performance report generation
        performance_report = extractor.get_performance_report()
        assert isinstance(performance_report, dict)
        assert 'batch_summary' in performance_report
        logger.info("✓ Performance report generation works")
        
        # Test efficiency calculation
        efficiency = extractor.get_feature_extraction_efficiency()
        assert isinstance(efficiency, dict)
        logger.info("✓ Efficiency calculation works")
        
        return True
        
    except Exception as e:
        logger.error(f"✗ Statistics test failed: {str(e)}")
        return False


def test_feature_validation_with_mock_data():
    """Test feature validation with mock data."""
    logger.info("=== Testing Feature Validation with Mock Data ===")
    
    try:
        sys.path.insert(0, str(Path(__file__).parent.parent.parent / "src" / "tlr4_binding" / "molecular_analysis"))
        from extractor import MolecularFeatureExtractor
        import pandas as pd
        import numpy as np
        
        extractor = MolecularFeatureExtractor()
        
        # Create comprehensive mock feature DataFrame
        mock_data = {
            'compound_name': ['test1', 'test2', 'test3'],
            'molecular_weight': [100.0, np.nan, 150.0],
            'logp': [1.5, 2.0, np.inf],
            'tpsa': [50.0, 75.0, 100.0],
            'rotatable_bonds': [3, 5, 2],
            'hbd': [2, 1, 3],
            'hba': [4, 5, 6],
            'formal_charge': [0, 1, -1],
            'radius_of_gyration': [3.5, 4.0, 3.2],
            'molecular_volume': [200.0, 250.0, 180.0],
            'surface_area': [150.0, 200.0, 120.0],
            'asphericity': [0.1, 0.2, 0.05],
            'ring_count': [1, 2, 0],
            'aromatic_rings': [1, 1, 0],
            'branching_index': [0.3, 0.4, 0.2],
            'dipole_moment': [1.5, 2.0, 1.0],
            'polarizability': [20.0, 25.0, 15.0]
        }
        mock_df = pd.DataFrame(mock_data)
        
        # Test validation
        validation_results = extractor.validate_features(mock_df)
        
        # Check structure
        expected_keys = ['missing_values', 'infinite_values', 'outliers', 'invalid_ranges']
        for key in expected_keys:
            assert key in validation_results
            assert isinstance(validation_results[key], list)
        
        logger.info("✓ Feature validation structure is correct")
        
        # Test feature summary
        summary_df = extractor.get_feature_summary(mock_df)
        assert isinstance(summary_df, pd.DataFrame)
        assert 'feature' in summary_df.columns
        assert 'count' in summary_df.columns
        
        logger.info("✓ Feature summary generation works")
        
        return True
        
    except Exception as e:
        logger.error(f"✗ Feature validation test failed: {str(e)}")
        return False


def test_performance_reporting():
    """Test performance reporting functionality."""
    logger.info("=== Testing Performance Reporting ===")
    
    try:
        sys.path.insert(0, str(Path(__file__).parent.parent.parent / "src" / "tlr4_binding" / "molecular_analysis"))
        from extractor import MolecularFeatureExtractor
        
        extractor = MolecularFeatureExtractor()
        
        # Test empty performance report
        report = extractor.get_performance_report()
        assert isinstance(report, dict)
        assert 'batch_summary' in report
        
        batch_summary = report['batch_summary']
        assert 'total_files' in batch_summary
        assert 'successful' in batch_summary
        assert 'failed' in batch_summary
        assert 'success_rate' in batch_summary
        
        logger.info("✓ Performance report structure is correct")
        
        # Test efficiency calculation
        efficiency = extractor.get_feature_extraction_efficiency()
        assert isinstance(efficiency, dict)
        
        logger.info("✓ Efficiency calculation works")
        
        return True
        
    except Exception as e:
        logger.error(f"✗ Performance reporting test failed: {str(e)}")
        return False


def main():
    """Run all standalone tests."""
    logger.info("Starting Standalone MolecularFeatureExtractor Tests")
    
    tests = [
        test_direct_import,
        test_initialization_modes,
        test_error_handling,
        test_statistics_functionality,
        test_feature_validation_with_mock_data,
        test_performance_reporting
    ]
    
    passed = 0
    failed = 0
    
    for test_func in tests:
        try:
            if test_func():
                passed += 1
                logger.info(f"✓ {test_func.__name__} PASSED")
            else:
                failed += 1
                logger.error(f"✗ {test_func.__name__} FAILED")
        except Exception as e:
            failed += 1
            logger.error(f"✗ {test_func.__name__} FAILED with exception: {str(e)}")
        
        logger.info("-" * 60)
    
    # Summary
    logger.info(f"Standalone Tests Summary:")
    logger.info(f"  Passed: {passed}")
    logger.info(f"  Failed: {failed}")
    logger.info(f"  Total: {passed + failed}")
    
    if failed == 0:
        logger.info("🎉 All standalone tests passed!")
        return True
    else:
        logger.error(f"❌ {failed} standalone tests failed!")
        return False


if __name__ == "__main__":
    success = main()
    sys.exit(0 if success else 1)
```
