## Supplementary material for "Machine Learning-Driven Prediction of TLR4 Binding Affinity: A Comprehensive Molecular Feature Analysis for Drug Discovery": Codebase: test_molecular_descriptors.py

```
"""
Unit tests for molecular descriptor calculation.

This module tests the MolecularDescriptorCalculator class with known compounds
to ensure accurate calculation of 2D molecular properties.
"""

import unittest
import sys
from pathlib import Path
import numpy as np
import pandas as pd

# Add src to path
sys.path.insert(0, str(Path(__file__).parent.parent.parent / "src"))

from tlr4_binding.molecular_analysis.descriptors import MolecularDescriptorCalculator


class TestMolecularDescriptorCalculator(unittest.TestCase):
    """Test MolecularDescriptorCalculator functionality."""
    
    def setUp(self):
        """Set up test fixtures."""
        self.calculator = MolecularDescriptorCalculator(include_advanced=True)
        
        # Known test compounds with expected properties
        self.test_compounds = {
            'aspirin': {
                'smiles': 'CC(=O)OC1=CC=CC=C1C(=O)O',
                'expected_mw': 180.16,  # Approximate
                'expected_logp': 1.19,  # Approximate
                'expected_hbd': 1,
                'expected_hba': 4,
                'expected_rotatable_bonds': 2
            },
            'caffeine': {
                'smiles': 'CN1C=NC2=C1C(=O)N(C(=O)N2C)C',
                'expected_mw': 194.19,  # Approximate
                'expected_logp': -0.07,  # Approximate
                'expected_hbd': 0,
                'expected_hba': 6,
                'expected_rotatable_bonds': 0
            },
            'benzene': {
                'smiles': 'C1=CC=CC=C1',
                'expected_mw': 78.11,
                'expected_logp': 2.13,  # Approximate
                'expected_hbd': 0,
                'expected_hba': 0,
                'expected_rotatable_bonds': 0
            }
        }
    
    def test_calculator_initialization(self):
        """Test calculator initialization."""
        self.assertIsNotNone(self.calculator)
        self.assertTrue(self.calculator.include_advanced)
        self.assertIsInstance(self.calculator.descriptor_functions, dict)
        self.assertGreater(len(self.calculator.descriptor_functions), 0)
    
    def test_get_available_descriptors(self):
        """Test getting list of available descriptors."""
        descriptors = self.calculator.get_available_descriptors()
        self.assertIsInstance(descriptors, list)
        self.assertGreater(len(descriptors), 0)
        
        # Check for key descriptors
        expected_descriptors = [
            'molecular_weight', 'logp', 'tpsa', 'hbd', 'hba', 
            'rotatable_bonds', 'ring_count', 'aromatic_rings'
        ]
        for desc in expected_descriptors:
            self.assertIn(desc, descriptors)
    
    def test_calculate_descriptors_aspirin(self):
        """Test descriptor calculation for aspirin."""
        aspirin_smiles = self.test_compounds['aspirin']['smiles']
        descriptors = self.calculator.calculate_descriptors(aspirin_smiles)
        
        self.assertIsInstance(descriptors, dict)
        self.assertGreater(len(descriptors), 0)
        
        # Check basic properties
        self.assertIn('molecular_weight', descriptors)
        self.assertIn('logp', descriptors)
        self.assertIn('hbd', descriptors)
        self.assertIn('hba', descriptors)
        self.assertIn('rotatable_bonds', descriptors)
        
        # Validate molecular weight (allow 5% tolerance)
        mw = descriptors['molecular_weight']
        expected_mw = self.test_compounds['aspirin']['expected_mw']
        self.assertAlmostEqual(mw, expected_mw, delta=expected_mw * 0.05)
        
        # Validate H-bond donors/acceptors (exact match expected)
        self.assertEqual(descriptors['hbd'], self.test_compounds['aspirin']['expected_hbd'])
        self.assertEqual(descriptors['hba'], self.test_compounds['aspirin']['expected_hba'])
        self.assertEqual(descriptors['rotatable_bonds'], self.test_compounds['aspirin']['expected_rotatable_bonds'])
    
    def test_calculate_descriptors_caffeine(self):
        """Test descriptor calculation for caffeine."""
        caffeine_smiles = self.test_compounds['caffeine']['smiles']
        descriptors = self.calculator.calculate_descriptors(caffeine_smiles)
        
        self.assertIsInstance(descriptors, dict)
        
        # Check that caffeine has no H-bond donors (caffeine is not acidic)
        self.assertEqual(descriptors['hbd'], 0)
        
        # Check that caffeine has multiple H-bond acceptors
        self.assertGreaterEqual(descriptors['hba'], 4)
        
        # Check that caffeine has no rotatable bonds (rigid structure)
        self.assertEqual(descriptors['rotatable_bonds'], 0)
    
    def test_calculate_descriptors_benzene(self):
        """Test descriptor calculation for benzene."""
        benzene_smiles = self.test_compounds['benzene']['smiles']
        descriptors = self.calculator.calculate_descriptors(benzene_smiles)
        
        self.assertIsInstance(descriptors, dict)
        
        # Benzene should have no H-bond donors or acceptors
        self.assertEqual(descriptors['hbd'], 0)
        self.assertEqual(descriptors['hba'], 0)
        
        # Benzene should have no rotatable bonds
        self.assertEqual(descriptors['rotatable_bonds'], 0)
        
        # Benzene should have 1 aromatic ring
        self.assertEqual(descriptors['aromatic_rings'], 1)
    
    def test_calculate_descriptor_subset(self):
        """Test calculating only specific descriptors."""
        aspirin_smiles = self.test_compounds['aspirin']['smiles']
        subset_names = ['molecular_weight', 'logp', 'hbd', 'hba']
        
        descriptors = self.calculator.calculate_descriptor_subset(
            aspirin_smiles, subset_names
        )
        
        self.assertIsInstance(descriptors, dict)
        self.assertEqual(len(descriptors), len(subset_names))
        
        for name in subset_names:
            self.assertIn(name, descriptors)
    
    def test_invalid_smiles_input(self):
        """Test handling of invalid SMILES input."""
        invalid_smiles = "invalid_smiles_string"
        
        with self.assertRaises(ValueError):
            self.calculator.calculate_descriptors(invalid_smiles)
    
    def test_none_input(self):
        """Test handling of None input."""
        with self.assertRaises(ValueError):
            self.calculator.calculate_descriptors(None)
    
    def test_empty_string_input(self):
        """Test handling of empty string input."""
        with self.assertRaises(ValueError):
            self.calculator.calculate_descriptors("")
    
    def test_descriptor_validation(self):
        """Test descriptor validation functionality."""
        aspirin_smiles = self.test_compounds['aspirin']['smiles']
        descriptors = self.calculator.calculate_descriptors(aspirin_smiles)
        
        validation_results = self.calculator.validate_descriptors(descriptors)
        
        self.assertIsInstance(validation_results, dict)
        self.assertIn('invalid_values', validation_results)
        self.assertIn('out_of_range', validation_results)
        self.assertIn('missing_values', validation_results)
        
        # Should have no invalid values for valid compound
        self.assertEqual(len(validation_results['invalid_values']), 0)
    
    def test_descriptor_validation_with_nan(self):
        """Test descriptor validation with NaN values."""
        # Create descriptors with NaN values
        descriptors_with_nan = {
            'molecular_weight': 180.16,
            'logp': np.nan,
            'tpsa': 50.0,
            'hbd': 1,
            'hba': 4
        }
        
        validation_results = self.calculator.validate_descriptors(descriptors_with_nan)
        
        # Should detect NaN values
        self.assertGreater(len(validation_results['invalid_values']), 0)
        self.assertIn('logp', str(validation_results['invalid_values']))
    
    def test_descriptor_validation_out_of_range(self):
        """Test descriptor validation with out-of-range values."""
        # Create descriptors with out-of-range values
        descriptors_out_of_range = {
            'molecular_weight': 5000.0,  # Too high
            'logp': 15.0,  # Too high
            'tpsa': 50.0,
            'hbd': -1,  # Negative H-bond donors
            'hba': 4
        }
        
        validation_results = self.calculator.validate_descriptors(descriptors_out_of_range)
        
        # Should detect out-of-range values
        self.assertGreater(len(validation_results['out_of_range']), 0)
    
    def test_advanced_descriptors(self):
        """Test calculation of advanced descriptors."""
        aspirin_smiles = self.test_compounds['aspirin']['smiles']
        descriptors = self.calculator.calculate_descriptors(aspirin_smiles)
        
        # Check for advanced descriptors
        advanced_descriptors = [
            'morgan_fingerprint_density',
            'maccs_keys_density',
            'molecular_flexibility',
            'aromatic_ratio',
            'heteroatom_ratio'
        ]
        
        for desc in advanced_descriptors:
            if self.calculator.include_advanced:
                self.assertIn(desc, descriptors)
                self.assertIsInstance(descriptors[desc], (int, float))
                self.assertFalse(np.isnan(descriptors[desc]))
    
    def test_descriptor_consistency(self):
        """Test that descriptors are consistent across multiple calculations."""
        aspirin_smiles = self.test_compounds['aspirin']['smiles']
        
        # Calculate descriptors multiple times
        desc1 = self.calculator.calculate_descriptors(aspirin_smiles)
        desc2 = self.calculator.calculate_descriptors(aspirin_smiles)
        
        # Should be identical
        for key in desc1:
            if key in desc2:
                if np.isnan(desc1[key]) and np.isnan(desc2[key]):
                    continue  # Both NaN is fine
                self.assertEqual(desc1[key], desc2[key])
    
    def test_descriptor_types(self):
        """Test that descriptors have correct data types."""
        aspirin_smiles = self.test_compounds['aspirin']['smiles']
        descriptors = self.calculator.calculate_descriptors(aspirin_smiles)
        
        for name, value in descriptors.items():
            self.assertIsInstance(value, (int, float, np.number))
            if not np.isnan(value):
                self.assertIsInstance(value, (int, float))
    
    def test_lipinski_rule_of_five_descriptors(self):
        """Test Lipinski's Rule of Five descriptors specifically."""
        aspirin_smiles = self.test_compounds['aspirin']['smiles']
        descriptors = self.calculator.calculate_descriptors(aspirin_smiles)
        
        # Check Lipinski descriptors
        lipinski_descriptors = ['molecular_weight', 'logp', 'hbd', 'hba']
        for desc in lipinski_descriptors:
            self.assertIn(desc, descriptors)
            self.assertIsInstance(descriptors[desc], (int, float))
        
        # Check that values are reasonable
        self.assertLess(descriptors['molecular_weight'], 1000)  # Should be < 500 for drug-like
        self.assertLess(descriptors['hbd'], 10)  # Should be <= 5 for drug-like
        self.assertLess(descriptors['hba'], 15)  # Should be <= 10 for drug-like


class TestMolecularDescriptorCalculatorEdgeCases(unittest.TestCase):
    """Test edge cases and error conditions."""
    
    def setUp(self):
        """Set up test fixtures."""
        self.calculator = MolecularDescriptorCalculator()
    
    def test_very_small_molecule(self):
        """Test descriptor calculation for very small molecules (e.g., water)."""
        water_smiles = "O"
        descriptors = self.calculator.calculate_descriptors(water_smiles)
        
        self.assertIsInstance(descriptors, dict)
        self.assertIn('molecular_weight', descriptors)
        self.assertLess(descriptors['molecular_weight'], 20)  # Water MW ~18
    
    def test_very_large_molecule(self):
        """Test descriptor calculation for large molecules."""
        # Create a large molecule SMILES (simplified)
        large_smiles = "C" * 100  # Very long carbon chain
        try:
            descriptors = self.calculator.calculate_descriptors(large_smiles)
            self.assertIsInstance(descriptors, dict)
        except ValueError:
            # This is acceptable - very large molecules might not be processable
            pass
    
    def test_charged_molecules(self):
        """Test descriptor calculation for charged molecules."""
        # Ammonium ion
        ammonium_smiles = "[NH4+]"
        try:
            descriptors = self.calculator.calculate_descriptors(ammonium_smiles)
            self.assertIsInstance(descriptors, dict)
            self.assertIn('formal_charge', descriptors)
            self.assertEqual(descriptors['formal_charge'], 1)
        except ValueError:
            # Charged molecules might not be processable in all cases
            pass
    
    def test_aromatic_heterocycles(self):
        """Test descriptor calculation for aromatic heterocycles."""
        # Pyridine
        pyridine_smiles = "C1=CC=NC=C1"
        try:
            descriptors = self.calculator.calculate_descriptors(pyridine_smiles)
            self.assertIsInstance(descriptors, dict)
            self.assertIn('aromatic_rings', descriptors)
            self.assertEqual(descriptors['aromatic_rings'], 1)
        except ValueError:
            pass


if __name__ == '__main__':
    # Run tests with verbose output
    unittest.main(verbosity=2)
```
