## Supplementary material for "Machine Learning-Driven Prediction of TLR4 Binding Affinity: A Comprehensive Molecular Feature Analysis for Drug Discovery": Codebase: test_molecular_descriptors_minimal.py

```
"""
Minimal unit tests for molecular descriptor calculation.

This module tests the MolecularDescriptorCalculator class with basic functionality
without requiring RDKit or other heavy dependencies.
"""

import unittest
import sys
from pathlib import Path
import numpy as np

# Add src to path
sys.path.insert(0, str(Path(__file__).parent.parent.parent / "src"))

from tlr4_binding.molecular_analysis.descriptors import MolecularDescriptorCalculator


class TestMolecularDescriptorCalculatorMinimal(unittest.TestCase):
    """Test MolecularDescriptorCalculator basic functionality."""
    
    def setUp(self):
        """Set up test fixtures."""
        self.calculator = MolecularDescriptorCalculator(include_advanced=False)
    
    def test_calculator_initialization(self):
        """Test calculator initialization."""
        self.assertIsNotNone(self.calculator)
        self.assertIsInstance(self.calculator.descriptor_functions, dict)
        self.assertGreater(len(self.calculator.descriptor_functions), 0)
    
    def test_get_available_descriptors(self):
        """Test getting list of available descriptors."""
        descriptors = self.calculator.get_available_descriptors()
        self.assertIsInstance(descriptors, list)
        self.assertGreater(len(descriptors), 0)
        
        # Check for key descriptors
        expected_descriptors = [
            'molecular_weight', 'logp', 'tpsa', 'hbd', 'hba', 
            'rotatable_bonds', 'ring_count', 'aromatic_rings'
        ]
        for desc in expected_descriptors:
            self.assertIn(desc, descriptors)
    
    def test_invalid_smiles_input(self):
        """Test handling of invalid SMILES input."""
        invalid_smiles = "invalid_smiles_string"
        
        with self.assertRaises(ValueError):
            self.calculator.calculate_descriptors(invalid_smiles)
    
    def test_none_input(self):
        """Test handling of None input."""
        with self.assertRaises(ValueError):
            self.calculator.calculate_descriptors(None)
    
    def test_empty_string_input(self):
        """Test handling of empty string input."""
        with self.assertRaises(ValueError):
            self.calculator.calculate_descriptors("")
    
    def test_descriptor_validation_with_nan(self):
        """Test descriptor validation with NaN values."""
        # Create descriptors with NaN values
        descriptors_with_nan = {
            'molecular_weight': 180.16,
            'logp': np.nan,
            'tpsa': 50.0,
            'hbd': 1,
            'hba': 4
        }
        
        validation_results = self.calculator.validate_descriptors(descriptors_with_nan)
        
        # Should detect NaN values
        self.assertGreater(len(validation_results['invalid_values']), 0)
        self.assertIn('logp', str(validation_results['invalid_values']))
    
    def test_descriptor_validation_out_of_range(self):
        """Test descriptor validation with out-of-range values."""
        # Create descriptors with out-of-range values
        descriptors_out_of_range = {
            'molecular_weight': 5000.0,  # Too high
            'logp': 15.0,  # Too high
            'tpsa': 50.0,
            'hbd': -1,  # Negative H-bond donors
            'hba': 4
        }
        
        validation_results = self.calculator.validate_descriptors(descriptors_out_of_range)
        
        # Should detect out-of-range values
        self.assertGreater(len(validation_results['out_of_range']), 0)
    
    def test_descriptor_subset_calculation(self):
        """Test calculating only specific descriptors."""
        # Test with invalid input to ensure error handling works
        subset_names = ['molecular_weight', 'logp', 'hbd', 'hba']
        
        with self.assertRaises(ValueError):
            self.calculator.calculate_descriptor_subset("invalid", subset_names)
    
    def test_fallback_descriptors(self):
        """Test fallback descriptor calculation when RDKit is not available."""
        # This should work even without RDKit
        descriptors = self.calculator._calculate_fallback_descriptors("test")
        self.assertIsInstance(descriptors, dict)
        
        # Should have all expected descriptors
        expected_descriptors = [
            'molecular_weight', 'logp', 'tpsa', 'formal_charge',
            'hbd', 'hba', 'rotatable_bonds', 'ring_count',
            'aromatic_rings', 'heavy_atoms', 'dipole_moment',
            'polarizability', 'molecular_volume', 'surface_area',
            'radius_of_gyration', 'asphericity'
        ]
        
        for desc in expected_descriptors:
            self.assertIn(desc, descriptors)
            self.assertTrue(np.isnan(descriptors[desc]))  # Should be NaN in fallback mode


if __name__ == '__main__':
    # Run tests with verbose output
    unittest.main(verbosity=2)
```
