## Supplementary material for "Machine Learning-Driven Prediction of TLR4 Binding Affinity: A Comprehensive Molecular Feature Analysis for Drug Discovery": Codebase: test_pdbqt_parser.py

```
"""
Unit tests for PDBQT file parser and validation.

Tests the PDBQTParser class with various file formats, validation scenarios,
and error handling cases to ensure robust parsing functionality.
"""

import pytest
import tempfile
import os
from pathlib import Path
from unittest.mock import patch, mock_open
from typing import Dict, List

from src.tlr4_binding.molecular_analysis.parser import PDBQTParser, PDBQTParserInterface


class TestPDBQTParserInterface:
    """Test the abstract interface implementation."""
    
    def test_interface_implementation(self):
        """Test that PDBQTParser implements the interface correctly."""
        parser = PDBQTParser()
        assert isinstance(parser, PDBQTParserInterface)
        
        # Test that all abstract methods are implemented
        assert hasattr(parser, 'parse_file')
        assert hasattr(parser, 'validate_file')
        assert hasattr(parser, 'extract_ligand_data')


class TestPDBQTParserInitialization:
    """Test parser initialization and configuration."""
    
    def test_default_initialization(self):
        """Test parser with default settings."""
        parser = PDBQTParser()
        assert parser.strict_validation is True
        assert 'C' in parser.supported_atoms
        assert 'N' in parser.supported_atoms
        assert 'O' in parser.supported_atoms
        assert 'REMARK' in parser.required_sections
        assert 'ATOM' in parser.required_sections
    
    def test_custom_initialization(self):
        """Test parser with custom validation settings."""
        parser = PDBQTParser(strict_validation=False)
        assert parser.strict_validation is False


class TestPDBQTParserValidation:
    """Test PDBQT file validation functionality."""
    
    def test_validate_nonexistent_file(self):
        """Test validation of non-existent file."""
        parser = PDBQTParser()
        is_valid, errors = parser.validate_file("/nonexistent/file.pdbqt")
        
        assert is_valid is False
        assert len(errors) == 1
        assert "File not found" in errors[0]
    
    def test_validate_empty_file(self):
        """Test validation of empty file."""
        parser = PDBQTParser()
        
        with tempfile.NamedTemporaryFile(mode='w', suffix='.pdbqt', delete=False) as f:
            f.write("")
            temp_file = f.name
        
        try:
            is_valid, errors = parser.validate_file(temp_file)
            assert is_valid is False
            assert "No ATOM or HETATM records found" in errors
            assert "No END record found" in errors
        finally:
            os.unlink(temp_file)
    
    def test_validate_valid_pdbqt_structure(self):
        """Test validation of valid PDBQT structure."""
        parser = PDBQTParser()
        
        valid_pdbqt_content = """REMARK Test file
HETATM    1  C1          0       0.000   0.000   0.000  1.00  0.00     0.000 C 
HETATM    2  N1          0       1.000   1.000   1.000  1.00  0.00     0.000 N 
END
"""
        
        with tempfile.NamedTemporaryFile(mode='w', suffix='.pdbqt', delete=False) as f:
            f.write(valid_pdbqt_content)
            temp_file = f.name
        
        try:
            is_valid, errors = parser.validate_file(temp_file)
            assert is_valid is True
            assert len(errors) == 0
        finally:
            os.unlink(temp_file)
    
    def test_validate_missing_atoms(self):
        """Test validation of file without atom records."""
        parser = PDBQTParser()
        
        invalid_content = """REMARK Test file
END
"""
        
        with tempfile.NamedTemporaryFile(mode='w', suffix='.pdbqt', delete=False) as f:
            f.write(invalid_content)
            temp_file = f.name
        
        try:
            is_valid, errors = parser.validate_file(temp_file)
            assert is_valid is False
            assert "No ATOM or HETATM records found" in errors
        finally:
            os.unlink(temp_file)
    
    def test_validate_missing_end(self):
        """Test validation of file without END record."""
        parser = PDBQTParser()
        
        invalid_content = """REMARK Test file
HETATM    1  C1          0       0.000   0.000   0.000  1.00  0.00     0.000 C 
"""
        
        with tempfile.NamedTemporaryFile(mode='w', suffix='.pdbqt', delete=False) as f:
            f.write(invalid_content)
            temp_file = f.name
        
        try:
            is_valid, errors = parser.validate_file(temp_file)
            assert is_valid is False
            assert "No END record found" in errors
        finally:
            os.unlink(temp_file)
    
    def test_validate_short_atom_line(self):
        """Test validation of atom line that's too short."""
        parser = PDBQTParser()
        
        invalid_content = """REMARK Test file
HETATM    1  C1          0       0.000   0.000   0.000  1.00  0.00     0.000 C 
HETATM    2  N1          0       1.000   1.000   1.000
END
"""
        
        with tempfile.NamedTemporaryFile(mode='w', suffix='.pdbqt', delete=False) as f:
            f.write(invalid_content)
            temp_file = f.name
        
        try:
            is_valid, errors = parser.validate_file(temp_file)
            assert is_valid is False
            assert any("too short" in error for error in errors)
        finally:
            os.unlink(temp_file)
    
    def test_validate_missing_atom_name(self):
        """Test validation of atom line missing atom name."""
        parser = PDBQTParser()
        
        invalid_content = """REMARK Test file
HETATM    1               0       0.000   0.000   0.000  1.00  0.00     0.000 C 
END
"""
        
        with tempfile.NamedTemporaryFile(mode='w', suffix='.pdbqt', delete=False) as f:
            f.write(invalid_content)
            temp_file = f.name
        
        try:
            is_valid, errors = parser.validate_file(temp_file)
            assert is_valid is False
            assert any("missing atom name" in error for error in errors)
        finally:
            os.unlink(temp_file)


class TestPDBQTParserParsing:
    """Test PDBQT file parsing functionality."""
    
    def test_parse_nonexistent_file(self):
        """Test parsing of non-existent file."""
        parser = PDBQTParser()
        
        with pytest.raises(FileNotFoundError):
            parser.parse_file("/nonexistent/file.pdbqt")
    
    def test_parse_simple_pdbqt_file(self):
        """Test parsing of simple PDBQT file."""
        parser = PDBQTParser()
        
        pdbqt_content = """REMARK Test file
HETATM    1  C1          0       0.000   0.000   0.000  1.00  0.00     0.000 C 
HETATM    2  N1          0       1.000   1.000   1.000  1.00  0.00     0.000 N 
END
"""
        
        with tempfile.NamedTemporaryFile(mode='w', suffix='.pdbqt', delete=False) as f:
            f.write(pdbqt_content)
            temp_file = f.name
        
        try:
            result = parser.parse_file(temp_file)
            
            assert result['file_path'] == temp_file
            assert len(result['header']) == 1
            assert len(result['atoms']) == 2
            assert len(result['models']) == 0
            assert len(result['footer']) == 1
            
            # Check first atom
            atom1 = result['atoms'][0]
            assert atom1['record_type'] == 'HETATM'
            assert atom1['atom_serial'] == 1
            assert atom1['atom_name'] == 'C1'
            assert atom1['element'] == 'C'
            assert atom1['x'] == 0.0
            assert atom1['y'] == 0.0
            assert atom1['z'] == 0.0
            
            # Check second atom
            atom2 = result['atoms'][1]
            assert atom2['atom_serial'] == 2
            assert atom2['atom_name'] == 'N1'
            assert atom2['element'] == 'N'
            assert atom2['x'] == 1.0
            assert atom2['y'] == 1.0
            assert atom2['z'] == 1.0
            
        finally:
            os.unlink(temp_file)
    
    def test_parse_pdbqt_with_models(self):
        """Test parsing of PDBQT file with multiple models."""
        parser = PDBQTParser()
        
        pdbqt_content = """REMARK Test file with models
MODEL 1
HETATM    1  C1          0       0.000   0.000   0.000  1.00  0.00     0.000 C 
ENDMDL
MODEL 2
HETATM    1  C1          0       1.000   1.000   1.000  1.00  0.00     0.000 C 
ENDMDL
END
"""
        
        with tempfile.NamedTemporaryFile(mode='w', suffix='.pdbqt', delete=False) as f:
            f.write(pdbqt_content)
            temp_file = f.name
        
        try:
            result = parser.parse_file(temp_file)
            
            assert len(result['models']) == 2
            assert result['models'][0]['model_number'] == 1
            assert result['models'][1]['model_number'] == 2
            
            # Each model should have its own atoms
            assert len(result['models'][0]['atoms']) == 1
            assert len(result['models'][1]['atoms']) == 1
            
        finally:
            os.unlink(temp_file)
    
    def test_parse_atom_line_with_missing_data(self):
        """Test parsing of atom line with missing or invalid data."""
        parser = PDBQTParser()
        
        pdbqt_content = """REMARK Test file
HETATM           C1          0       0.000   0.000   0.000  1.00  0.00     0.000 C 
HETATM    2  N1          0       1.000   1.000   1.000  1.00  0.00     0.000 N 
END
"""
        
        with tempfile.NamedTemporaryFile(mode='w', suffix='.pdbqt', delete=False) as f:
            f.write(pdbqt_content)
            temp_file = f.name
        
        try:
            result = parser.parse_file(temp_file)
            
            # First atom should have default values for missing serial
            atom1 = result['atoms'][0]
            assert atom1['atom_serial'] == 0  # Default value for missing serial
            assert atom1['atom_name'] == 'C1'
            
            # Second atom should parse normally
            atom2 = result['atoms'][1]
            assert atom2['atom_serial'] == 2
            assert atom2['atom_name'] == 'N1'
            
        finally:
            os.unlink(temp_file)
    
    def test_parse_vina_remark(self):
        """Test parsing of Vina scoring information in REMARK lines."""
        parser = PDBQTParser()
        
        pdbqt_content = """REMARK VINA RESULT: -5.2 0.000 0.000
HETATM    1  C1          0       0.000   0.000   0.000  1.00  0.00     0.000 C 
END
"""
        
        with tempfile.NamedTemporaryFile(mode='w', suffix='.pdbqt', delete=False) as f:
            f.write(pdbqt_content)
            temp_file = f.name
        
        try:
            result = parser.parse_file(temp_file)
            
            assert 'vina_score' in result['metadata']
            assert result['metadata']['vina_score'] == -5.2
            
        finally:
            os.unlink(temp_file)
    
    def test_parse_element_extraction(self):
        """Test extraction of element from atom name when not specified."""
        parser = PDBQTParser()
        
        pdbqt_content = """REMARK Test file
HETATM    1  CA          0       0.000   0.000   0.000  1.00  0.00     0.000
HETATM    2  N1          0       1.000   1.000   1.000  1.00  0.00     0.000
END
"""
        
        with tempfile.NamedTemporaryFile(mode='w', suffix='.pdbqt', delete=False) as f:
            f.write(pdbqt_content)
            temp_file = f.name
        
        try:
            result = parser.parse_file(temp_file)
            
            # First atom should extract 'C' from 'CA'
            atom1 = result['atoms'][0]
            assert atom1['element'] == 'C'
            
            # Second atom should extract 'N' from 'N1'
            atom2 = result['atoms'][1]
            assert atom2['element'] == 'N'
            
        finally:
            os.unlink(temp_file)
    
    def test_strict_validation_enabled(self):
        """Test parsing with strict validation enabled."""
        parser = PDBQTParser(strict_validation=True)
        
        # Create file with unsupported element
        pdbqt_content = """REMARK Test file
HETATM    1  X1          0       0.000   0.000   0.000  1.00  0.00     0.000 X 
END
"""
        
        with tempfile.NamedTemporaryFile(mode='w', suffix='.pdbqt', delete=False) as f:
            f.write(pdbqt_content)
            temp_file = f.name
        
        try:
            with pytest.raises(ValueError, match="PDBQT validation failed"):
                parser.parse_file(temp_file)
        finally:
            os.unlink(temp_file)
    
    def test_strict_validation_disabled(self):
        """Test parsing with strict validation disabled."""
        parser = PDBQTParser(strict_validation=False)
        
        # Create file with unsupported element
        pdbqt_content = """REMARK Test file
HETATM    1  X1          0       0.000   0.000   0.000  1.00  0.00     0.000 X 
END
"""
        
        with tempfile.NamedTemporaryFile(mode='w', suffix='.pdbqt', delete=False) as f:
            f.write(pdbqt_content)
            temp_file = f.name
        
        try:
            # Should not raise exception with strict validation disabled
            result = parser.parse_file(temp_file)
            assert len(result['atoms']) == 1
            assert result['atoms'][0]['element'] == 'X'
        finally:
            os.unlink(temp_file)


class TestPDBQTLigandDataExtraction:
    """Test ligand-specific data extraction functionality."""
    
    def test_extract_ligand_data_simple(self):
        """Test extraction of ligand data from simple PDBQT file."""
        parser = PDBQTParser()
        
        pdbqt_content = """REMARK Test file
HETATM    1  C1          0       0.000   0.000   0.000  1.00  0.00     0.000 C 
HETATM    2  N1          0       1.000   1.000   1.000  1.00  0.00     0.000 N 
HETATM    3  O1          0       2.000   2.000   2.000  1.00  0.00     0.000 O 
END
"""
        
        with tempfile.NamedTemporaryFile(mode='w', suffix='.pdbqt', delete=False) as f:
            f.write(pdbqt_content)
            temp_file = f.name
        
        try:
            ligand_data = parser.extract_ligand_data(temp_file)
            
            assert ligand_data['compound_name'] == Path(temp_file).stem
            assert ligand_data['atom_count'] == 3
            assert ligand_data['molecular_formula'] == 'CNO'
            assert ligand_data['binding_poses'] == 0  # No models in this file
            assert ligand_data['elements'] == {'C': 1, 'N': 1, 'O': 1}
            assert ligand_data['file_path'] == temp_file
            
        finally:
            os.unlink(temp_file)
    
    def test_extract_ligand_data_with_models(self):
        """Test extraction of ligand data from PDBQT file with multiple models."""
        parser = PDBQTParser()
        
        pdbqt_content = """REMARK Test file
MODEL 1
HETATM    1  C1          0       0.000   0.000   0.000  1.00  0.00     0.000 C 
ENDMDL
MODEL 2
HETATM    1  C1          0       1.000   1.000   1.000  1.00  0.00     0.000 C 
ENDMDL
END
"""
        
        with tempfile.NamedTemporaryFile(mode='w', suffix='.pdbqt', delete=False) as f:
            f.write(pdbqt_content)
            temp_file = f.name
        
        try:
            ligand_data = parser.extract_ligand_data(temp_file)
            
            assert ligand_data['atom_count'] == 2  # Total atoms across all models
            assert ligand_data['binding_poses'] == 2  # Two models
            assert ligand_data['elements'] == {'C': 2}
            
        finally:
            os.unlink(temp_file)
    
    def test_extract_ligand_data_molecular_formula(self):
        """Test molecular formula generation with multiple atoms of same element."""
        parser = PDBQTParser()
        
        pdbqt_content = """REMARK Test file
HETATM    1  C1          0       0.000   0.000   0.000  1.00  0.00     0.000 C 
HETATM    2  C2          0       1.000   1.000   1.000  1.00  0.00     0.000 C 
HETATM    3  C3          0       2.000   2.000   2.000  1.00  0.00     0.000 C 
HETATM    4  N1          0       3.000   3.000   3.000  1.00  0.00     0.000 N 
END
"""
        
        with tempfile.NamedTemporaryFile(mode='w', suffix='.pdbqt', delete=False) as f:
            f.write(pdbqt_content)
            temp_file = f.name
        
        try:
            ligand_data = parser.extract_ligand_data(temp_file)
            
            # Should group carbons together: C3N
            assert ligand_data['molecular_formula'] == 'C3N'
            assert ligand_data['elements'] == {'C': 3, 'N': 1}
            
        finally:
            os.unlink(temp_file)


class TestPDBQTParserErrorHandling:
    """Test error handling in PDBQT parser."""
    
    def test_parse_file_io_error(self):
        """Test handling of file I/O errors during parsing."""
        parser = PDBQTParser()
        
        with patch("builtins.open", mock_open()) as mock_file:
            mock_file.side_effect = IOError("Permission denied")
            
            with pytest.raises(ValueError, match="Failed to parse PDBQT file"):
                parser.parse_file("/some/file.pdbqt")
    
    def test_parse_file_validation_error(self):
        """Test handling of validation errors during parsing."""
        parser = PDBQTParser(strict_validation=True)
        
        # Create file with no atoms (validation should fail)
        pdbqt_content = """REMARK Test file
END
"""
        
        with tempfile.NamedTemporaryFile(mode='w', suffix='.pdbqt', delete=False) as f:
            f.write(pdbqt_content)
            temp_file = f.name
        
        try:
            with pytest.raises(ValueError, match="PDBQT validation failed"):
                parser.parse_file(temp_file)
        finally:
            os.unlink(temp_file)
    
    def test_extract_ligand_data_file_error(self):
        """Test error handling in ligand data extraction."""
        parser = PDBQTParser()
        
        with pytest.raises(FileNotFoundError):
            parser.extract_ligand_data("/nonexistent/file.pdbqt")


class TestPDBQTParserIntegration:
    """Integration tests with actual sample PDBQT files."""
    
    def test_parse_real_andrographolide_file(self):
        """Test parsing of real Andrographolide.pdbqt file."""
        parser = PDBQTParser()
        sample_file = "/home/brand/ember-pm/binding-data/raw/pdbqt/Andrographolide.pdbqt"
        
        if os.path.exists(sample_file):
            result = parser.parse_file(sample_file)
            
            # Basic structure validation
            assert result['file_path'] == sample_file
            assert len(result['atoms']) > 0
            assert len(result['header']) > 0
            
            # Check for typical PDBQT elements
            elements = {atom['element'] for atom in result['atoms']}
            assert 'C' in elements
            assert 'O' in elements
            
            # Extract ligand data
            ligand_data = parser.extract_ligand_data(sample_file)
            assert ligand_data['compound_name'] == 'Andrographolide'
            assert ligand_data['atom_count'] > 0
            assert 'C' in ligand_data['elements']
    
    def test_validate_real_files(self):
        """Test validation of real PDBQT files from the dataset."""
        parser = PDBQTParser()
        pdbqt_dir = "/home/brand/ember-pm/binding-data/raw/pdbqt"
        
        if os.path.exists(pdbqt_dir):
            sample_files = [
                "Andrographolide.pdbqt",
                "Apigenin.pdbqt", 
                "Artemisinin.pdbqt"
            ]
            
            for filename in sample_files:
                file_path = os.path.join(pdbqt_dir, filename)
                if os.path.exists(file_path):
                    is_valid, errors = parser.validate_file(file_path)
                    assert is_valid, f"Validation failed for {filename}: {errors}"
    
    def test_batch_processing_sample_files(self):
        """Test batch processing of multiple sample PDBQT files."""
        parser = PDBQTParser()
        pdbqt_dir = "/home/brand/ember-pm/binding-data/raw/pdbqt"
        
        if os.path.exists(pdbqt_dir):
            # Process first few files
            sample_files = [
                "Andrographolide.pdbqt",
                "Apigenin.pdbqt",
                "Artemisinin.pdbqt"
            ]
            
            results = []
            for filename in sample_files:
                file_path = os.path.join(pdbqt_dir, filename)
                if os.path.exists(file_path):
                    try:
                        ligand_data = parser.extract_ligand_data(file_path)
                        results.append(ligand_data)
                    except Exception as e:
                        pytest.fail(f"Failed to process {filename}: {e}")
            
            # Verify all files were processed successfully
            assert len(results) > 0
            for result in results:
                assert 'compound_name' in result
                assert 'atom_count' in result
                assert 'molecular_formula' in result


if __name__ == "__main__":
    pytest.main([__file__, "-v"])
```
