## Supplementary material for "Machine Learning-Driven Prediction of TLR4 Binding Affinity: A Comprehensive Molecular Feature Analysis for Drug Discovery": Codebase: test_structural_features.py

```
"""
Unit tests for 3D structural feature extraction using PyMOL.

Tests the StructuralFeatureExtractor class and its methods for calculating
3D molecular descriptors from PDBQT files.
"""

import pytest
import numpy as np
import tempfile
import os
from pathlib import Path
from unittest.mock import Mock, patch, MagicMock

from src.tlr4_binding.molecular_analysis.structure import (
    StructuralFeatureExtractor,
    StructuralFeatureExtractorInterface,
    PYMOL_AVAILABLE
)


class TestStructuralFeatureExtractorInterface:
    """Test the abstract interface."""
    
    def test_interface_abstract(self):
        """Test that interface cannot be instantiated."""
        with pytest.raises(TypeError):
            StructuralFeatureExtractorInterface()


class TestStructuralFeatureExtractor:
    """Test the StructuralFeatureExtractor class."""
    
    def setup_method(self):
        """Set up test fixtures."""
        self.extractor = StructuralFeatureExtractor()
        
        # Create a mock PDBQT file content
        self.mock_pdbqt_content = """REMARK  Generated by AutoDock Vina
ATOM      1  C   LIG A   1      20.154  30.231  25.123  1.00  0.00      A    C
ATOM      2  N   LIG A   1      21.234  31.345  26.234  1.00  0.00      A    N
ATOM      3  O   LIG A   1      22.345  32.456  27.345  1.00  0.00      A    O
ENDMDL
"""
    
    def test_initialization(self):
        """Test extractor initialization."""
        extractor = StructuralFeatureExtractor("test_session")
        assert extractor.session_name == "test_session"
        assert not extractor.session_initialized
    
    def test_get_available_features(self):
        """Test getting list of available features."""
        features = self.extractor.get_available_features()
        assert isinstance(features, list)
        assert len(features) > 0
        
        # Check for key features
        expected_features = [
            'radius_of_gyration', 'molecular_volume', 'surface_area',
            'asphericity', 'eccentricity', 'spherocity_index'
        ]
        for feature in expected_features:
            assert feature in features
    
    @patch('src.tlr4_binding.molecular_analysis.structure.PYMOL_AVAILABLE', False)
    def test_extract_features_fallback(self):
        """Test feature extraction when PyMOL is not available."""
        extractor = StructuralFeatureExtractor()
        
        with tempfile.NamedTemporaryFile(mode='w', suffix='.pdbqt', delete=False) as f:
            f.write(self.mock_pdbqt_content)
            temp_path = f.name
        
        try:
            features = extractor.extract_features(temp_path)
            assert isinstance(features, dict)
            
            # All features should be NaN when PyMOL is not available
            for feature_name in extractor.get_available_features():
                assert feature_name in features
                assert np.isnan(features[feature_name])
        finally:
            os.unlink(temp_path)
    
    @patch('src.tlr4_binding.molecular_analysis.structure.PYMOL_AVAILABLE', True)
    @patch('src.tlr4_binding.molecular_analysis.structure.cmd')
    def test_extract_features_with_pymol(self, mock_cmd):
        """Test feature extraction with PyMOL available."""
        # Mock PyMOL commands
        mock_cmd.load.return_value = None
        mock_cmd.delete.return_value = None
        mock_cmd.get_radius_of_gyration.return_value = 5.2
        mock_cmd.get.return_value = 5.2
        mock_cmd.measure_volume.return_value = None
        mock_cmd.measure_sasa.return_value = None
        mock_cmd.get_principal_moments.return_value = [10.0, 8.0, 6.0]
        mock_cmd.get_molecular_weight.return_value = 200.0
        mock_cmd.show.return_value = None
        
        with tempfile.NamedTemporaryFile(mode='w', suffix='.pdbqt', delete=False) as f:
            f.write(self.mock_pdbqt_content)
            temp_path = f.name
        
        try:
            features = extractor.extract_features(temp_path)
            assert isinstance(features, dict)
            
            # Check that key features are present
            expected_features = [
                'radius_of_gyration', 'molecular_volume', 'surface_area',
                'asphericity', 'eccentricity', 'spherocity_index'
            ]
            for feature in expected_features:
                assert feature in features
                
        finally:
            os.unlink(temp_path)
    
    def test_extract_features_invalid_file(self):
        """Test feature extraction with invalid file path."""
        with pytest.raises(ValueError):
            self.extractor.extract_features("nonexistent_file.pdbqt")
    
    @patch('src.tlr4_binding.molecular_analysis.structure.PYMOL_AVAILABLE', True)
    @patch('src.tlr4_binding.molecular_analysis.structure.cmd')
    def test_geometric_features_calculation(self, mock_cmd):
        """Test geometric feature calculations."""
        # Mock PyMOL responses
        mock_cmd.get_radius_of_gyration.return_value = 5.2
        mock_cmd.get.return_value = 5.2
        mock_cmd.measure_volume.return_value = None
        mock_cmd.measure_sasa.return_value = None
        mock_cmd.get_principal_moments.return_value = [10.0, 8.0, 6.0]
        
        features = self.extractor._calculate_geometric_features("test_mol")
        
        assert 'radius_of_gyration' in features
        assert 'molecular_volume' in features
        assert 'surface_area' in features
        assert 'asphericity' in features
        assert 'eccentricity' in features
        assert 'spherocity_index' in features
    
    @patch('src.tlr4_binding.molecular_analysis.structure.PYMOL_AVAILABLE', True)
    @patch('src.tlr4_binding.molecular_analysis.structure.cmd')
    def test_shape_features_calculation(self, mock_cmd):
        """Test shape feature calculations."""
        # Mock PyMOL responses
        mock_cmd.get_principal_moments.return_value = [10.0, 8.0, 6.0]
        mock_cmd.measure_volume.return_value = None
        mock_cmd.measure_sasa.return_value = None
        mock_cmd.get.return_value = 100.0
        
        features = self.extractor._calculate_shape_features("test_mol")
        
        assert 'elongation' in features
        assert 'flatness' in features
        assert 'compactness' in features
        assert 'convexity' in features
        assert 'concavity_index' in features
        assert 'roughness_index' in features
    
    @patch('src.tlr4_binding.molecular_analysis.structure.PYMOL_AVAILABLE', True)
    @patch('src.tlr4_binding.molecular_analysis.structure.cmd')
    def test_surface_features_calculation(self, mock_cmd):
        """Test surface feature calculations."""
        # Mock PyMOL responses
        mock_cmd.measure_sasa.return_value = None
        mock_cmd.get.return_value = 200.0
        mock_cmd.show.return_value = None
        
        features = self.extractor._calculate_surface_features("test_mol")
        
        assert 'polar_surface_area' in features
        assert 'hydrophobic_surface_area' in features
        assert 'positive_surface_area' in features
        assert 'negative_surface_area' in features
        assert 'surface_charge_density' in features
    
    @patch('src.tlr4_binding.molecular_analysis.structure.PYMOL_AVAILABLE', True)
    @patch('src.tlr4_binding.molecular_analysis.structure.cmd')
    def test_conformational_features_calculation(self, mock_cmd):
        """Test conformational feature calculations."""
        # Mock PyMOL responses
        mock_cmd.get_molecular_weight.return_value = 200.0
        mock_cmd.get.return_value = 200.0
        mock_cmd.get_principal_moments.return_value = [10.0, 8.0, 6.0]
        
        features = self.extractor._calculate_conformational_features("test_mol")
        
        assert 'flexibility_index' in features
        assert 'rigidity_index' in features
        assert 'planarity' in features
        assert 'torsional_angle_variance' in features
        assert 'bond_angle_variance' in features
    
    def test_radius_of_gyration_calculation(self):
        """Test radius of gyration calculation."""
        with patch('src.tlr4_binding.molecular_analysis.structure.cmd') as mock_cmd:
            mock_cmd.center.return_value = None
            mock_cmd.get_radius_of_gyration.return_value = 5.2
            mock_cmd.get.return_value = 5.2
            
            rg = self.extractor._calculate_radius_of_gyration("test_mol")
            assert rg == 5.2
    
    def test_radius_of_gyration_invalid(self):
        """Test radius of gyration with invalid result."""
        with patch('src.tlr4_binding.molecular_analysis.structure.cmd') as mock_cmd:
            mock_cmd.center.return_value = None
            mock_cmd.get_radius_of_gyration.return_value = -1.0
            mock_cmd.get.return_value = -1.0
            
            rg = self.extractor._calculate_radius_of_gyration("test_mol")
            assert np.isnan(rg)
    
    def test_molecular_volume_calculation(self):
        """Test molecular volume calculation."""
        with patch('src.tlr4_binding.molecular_analysis.structure.cmd') as mock_cmd:
            mock_cmd.measure_volume.return_value = None
            mock_cmd.get.return_value = 150.0
            
            volume = self.extractor._calculate_molecular_volume("test_mol")
            assert volume == 150.0
    
    def test_surface_area_calculation(self):
        """Test surface area calculation."""
        with patch('src.tlr4_binding.molecular_analysis.structure.cmd') as mock_cmd:
            mock_cmd.measure_sasa.return_value = None
            mock_cmd.get.return_value = 200.0
            
            sasa = self.extractor._calculate_surface_area("test_mol")
            assert sasa == 200.0
    
    def test_asphericity_calculation(self):
        """Test asphericity calculation."""
        with patch('src.tlr4_binding.molecular_analysis.structure.cmd') as mock_cmd:
            mock_cmd.iterate.return_value = None
            mock_cmd.get.return_value = [10.0, 8.0, 6.0]
            
            asphericity = self.extractor._calculate_asphericity("test_mol")
            expected = (10.0 - 8.0) / (10.0 + 8.0 + 6.0)  # 2/24 = 0.083
            assert abs(asphericity - expected) < 1e-6
    
    def test_eccentricity_calculation(self):
        """Test eccentricity calculation."""
        with patch('src.tlr4_binding.molecular_analysis.structure.cmd') as mock_cmd:
            mock_cmd.iterate.return_value = None
            mock_cmd.get.return_value = [10.0, 8.0, 6.0]
            
            eccentricity = self.extractor._calculate_eccentricity("test_mol")
            expected = np.sqrt(1 - (8.0 / 10.0) ** 2)
            assert abs(eccentricity - expected) < 1e-6
    
    def test_spherocity_calculation(self):
        """Test spherocity calculation."""
        with patch('src.tlr4_binding.molecular_analysis.structure.cmd') as mock_cmd:
            mock_cmd.iterate.return_value = None
            mock_cmd.get.return_value = [10.0, 8.0, 6.0]
            
            spherocity = self.extractor._calculate_spherocity_index("test_mol")
            expected = (3 * 6.0) / (10.0 + 8.0 + 6.0)  # 18/24 = 0.75
            assert abs(spherocity - expected) < 1e-6
    
    def test_elongation_calculation(self):
        """Test elongation calculation."""
        with patch('src.tlr4_binding.molecular_analysis.structure.cmd') as mock_cmd:
            mock_cmd.iterate.return_value = None
            mock_cmd.get.return_value = [10.0, 8.0, 6.0]
            
            elongation = self.extractor._calculate_elongation("test_mol")
            expected = np.sqrt(10.0 / 8.0)
            assert abs(elongation - expected) < 1e-6
    
    def test_flatness_calculation(self):
        """Test flatness calculation."""
        with patch('src.tlr4_binding.molecular_analysis.structure.cmd') as mock_cmd:
            mock_cmd.iterate.return_value = None
            mock_cmd.get.return_value = [10.0, 8.0, 6.0]
            
            flatness = self.extractor._calculate_flatness("test_mol")
            expected = np.sqrt(8.0 / 6.0)
            assert abs(flatness - expected) < 1e-6
    
    def test_cleanup_session(self):
        """Test PyMOL session cleanup."""
        with patch('src.tlr4_binding.molecular_analysis.structure.PYMOL_AVAILABLE', True):
            with patch('src.tlr4_binding.molecular_analysis.structure.cmd') as mock_cmd:
                self.extractor.session_initialized = True
                self.extractor.cleanup_session()
                mock_cmd.quit.assert_called_once()
                assert not self.extractor.session_initialized
    
    def test_destructor_cleanup(self):
        """Test that destructor calls cleanup."""
        with patch.object(self.extractor, 'cleanup_session') as mock_cleanup:
            del self.extractor
            mock_cleanup.assert_called_once()


if __name__ == "__main__":
    pytest.main([__file__])
```
