## Supplementary material for "Machine Learning-Driven Prediction of TLR4 Binding Affinity: A Comprehensive Molecular Feature Analysis for Drug Discovery": Codebase: test_structural_standalone.py

```
#!/usr/bin/env python3
"""
Standalone unit tests for 3D structural feature extraction.

This script tests the StructuralFeatureExtractor by importing
only the specific module without going through the package __init__.py
"""

import sys
import os
import logging
import tempfile
import numpy as np
from pathlib import Path
from unittest.mock import Mock, patch, MagicMock

# Add src to path
sys.path.insert(0, str(Path(__file__).parent.parent.parent / "src"))

# Direct import of the specific module
import importlib.util
spec = importlib.util.spec_from_file_location(
    "structure", 
    Path(__file__).parent / "src" / "tlr4_binding" / "molecular_analysis" / "structure.py"
)
structure_module = importlib.util.module_from_spec(spec)
spec.loader.exec_module(structure_module)

StructuralFeatureExtractor = structure_module.StructuralFeatureExtractor
StructuralFeatureExtractorInterface = structure_module.StructuralFeatureExtractorInterface
PYMOL_AVAILABLE = structure_module.PYMOL_AVAILABLE

# Set up logging
logging.basicConfig(level=logging.INFO)
logger = logging.getLogger(__name__)


def test_interface_abstract():
    """Test that interface cannot be instantiated."""
    try:
        StructuralFeatureExtractorInterface()
        assert False, "Should not be able to instantiate abstract interface"
    except TypeError:
        pass  # Expected


def test_initialization():
    """Test extractor initialization."""
    extractor = StructuralFeatureExtractor("test_session")
    assert extractor.session_name == "test_session"
    assert not extractor.session_initialized


def test_get_available_features():
    """Test getting list of available features."""
    extractor = StructuralFeatureExtractor()
    features = extractor.get_available_features()
    assert isinstance(features, list)
    assert len(features) > 0
    
    # Check for key features
    expected_features = [
        'radius_of_gyration', 'molecular_volume', 'surface_area',
        'asphericity', 'eccentricity', 'spherocity_index'
    ]
    for feature in expected_features:
        assert feature in features


def test_extract_features_fallback():
    """Test feature extraction when PyMOL is not available."""
    with patch.object(structure_module, 'PYMOL_AVAILABLE', False):
        extractor = StructuralFeatureExtractor()
        
        # Create a mock PDBQT file content
        mock_pdbqt_content = """REMARK  Generated by AutoDock Vina
ATOM      1  C   LIG A   1      20.154  30.231  25.123  1.00  0.00      A    C
ATOM      2  N   LIG A   1      21.234  31.345  26.234  1.00  0.00      A    N
ATOM      3  O   LIG A   1      22.345  32.456  27.345  1.00  0.00      A    O
ENDMDL
"""
        
        with tempfile.NamedTemporaryFile(mode='w', suffix='.pdbqt', delete=False) as f:
            f.write(mock_pdbqt_content)
            temp_path = f.name
        
        try:
            features = extractor.extract_features(temp_path)
            assert isinstance(features, dict)
            
            # All features should be NaN when PyMOL is not available
            for feature_name in extractor.get_available_features():
                assert feature_name in features
                assert np.isnan(features[feature_name])
        finally:
            os.unlink(temp_path)


def test_extract_features_invalid_file():
    """Test feature extraction with invalid file path."""
    extractor = StructuralFeatureExtractor()
    try:
        extractor.extract_features("nonexistent_file.pdbqt")
        assert False, "Should raise ValueError for nonexistent file"
    except ValueError:
        pass  # Expected


def test_radius_of_gyration_calculation():
    """Test radius of gyration calculation."""
    extractor = StructuralFeatureExtractor()
    
    with patch.object(structure_module, 'cmd') as mock_cmd:
        mock_cmd.center.return_value = None
        mock_cmd.get_radius_of_gyration.return_value = 5.2
        mock_cmd.get.return_value = 5.2
        
        rg = extractor._calculate_radius_of_gyration("test_mol")
        assert rg == 5.2


def test_radius_of_gyration_invalid():
    """Test radius of gyration with invalid result."""
    extractor = StructuralFeatureExtractor()
    
    with patch.object(structure_module, 'cmd') as mock_cmd:
        mock_cmd.center.return_value = None
        mock_cmd.get_radius_of_gyration.return_value = -1.0
        mock_cmd.get.return_value = -1.0
        
        rg = extractor._calculate_radius_of_gyration("test_mol")
        assert np.isnan(rg)


def test_molecular_volume_calculation():
    """Test molecular volume calculation."""
    extractor = StructuralFeatureExtractor()
    
    with patch.object(structure_module, 'cmd') as mock_cmd:
        mock_cmd.measure_volume.return_value = None
        mock_cmd.get.return_value = 150.0
        
        volume = extractor._calculate_molecular_volume("test_mol")
        assert volume == 150.0


def test_surface_area_calculation():
    """Test surface area calculation."""
    extractor = StructuralFeatureExtractor()
    
    with patch.object(structure_module, 'cmd') as mock_cmd:
        mock_cmd.measure_sasa.return_value = None
        mock_cmd.get.return_value = 200.0
        
        sasa = extractor._calculate_surface_area("test_mol")
        assert sasa == 200.0


def test_asphericity_calculation():
    """Test asphericity calculation."""
    extractor = StructuralFeatureExtractor()
    
    with patch.object(structure_module, 'cmd') as mock_cmd:
        mock_cmd.iterate.return_value = None
        mock_cmd.get.return_value = [10.0, 8.0, 6.0]
        
        asphericity = extractor._calculate_asphericity("test_mol")
        expected = (10.0 - 8.0) / (10.0 + 8.0 + 6.0)  # 2/24 = 0.083
        assert abs(asphericity - expected) < 1e-6


def test_eccentricity_calculation():
    """Test eccentricity calculation."""
    extractor = StructuralFeatureExtractor()
    
    with patch.object(structure_module, 'cmd') as mock_cmd:
        mock_cmd.iterate.return_value = None
        mock_cmd.get.return_value = [10.0, 8.0, 6.0]
        
        eccentricity = extractor._calculate_eccentricity("test_mol")
        expected = np.sqrt(1 - (8.0 / 10.0) ** 2)
        assert abs(eccentricity - expected) < 1e-6


def test_spherocity_calculation():
    """Test spherocity calculation."""
    extractor = StructuralFeatureExtractor()
    
    with patch.object(structure_module, 'cmd') as mock_cmd:
        mock_cmd.iterate.return_value = None
        mock_cmd.get.return_value = [10.0, 8.0, 6.0]
        
        spherocity = extractor._calculate_spherocity_index("test_mol")
        expected = (3 * 6.0) / (10.0 + 8.0 + 6.0)  # 18/24 = 0.75
        assert abs(spherocity - expected) < 1e-6


def test_elongation_calculation():
    """Test elongation calculation."""
    extractor = StructuralFeatureExtractor()
    
    with patch.object(structure_module, 'cmd') as mock_cmd:
        mock_cmd.iterate.return_value = None
        mock_cmd.get.return_value = [10.0, 8.0, 6.0]
        
        elongation = extractor._calculate_elongation("test_mol")
        expected = np.sqrt(10.0 / 8.0)
        assert abs(elongation - expected) < 1e-6


def test_flatness_calculation():
    """Test flatness calculation."""
    extractor = StructuralFeatureExtractor()
    
    with patch.object(structure_module, 'cmd') as mock_cmd:
        mock_cmd.iterate.return_value = None
        mock_cmd.get.return_value = [10.0, 8.0, 6.0]
        
        flatness = extractor._calculate_flatness("test_mol")
        expected = np.sqrt(8.0 / 6.0)
        assert abs(flatness - expected) < 1e-6


def test_cleanup_session():
    """Test PyMOL session cleanup."""
    with patch.object(structure_module, 'PYMOL_AVAILABLE', True):
        with patch.object(structure_module, 'cmd') as mock_cmd:
            extractor = StructuralFeatureExtractor()
            extractor.session_initialized = True
            extractor.cleanup_session()
            mock_cmd.quit.assert_called_once()
            assert not extractor.session_initialized


def run_all_tests():
    """Run all tests."""
    tests = [
        test_interface_abstract,
        test_initialization,
        test_get_available_features,
        test_extract_features_fallback,
        test_extract_features_invalid_file,
        test_radius_of_gyration_calculation,
        test_radius_of_gyration_invalid,
        test_molecular_volume_calculation,
        test_surface_area_calculation,
        test_asphericity_calculation,
        test_eccentricity_calculation,
        test_spherocity_calculation,
        test_elongation_calculation,
        test_flatness_calculation,
        test_cleanup_session,
    ]
    
    passed = 0
    failed = 0
    
    for test in tests:
        try:
            test()
            print(f"✅ {test.__name__}")
            passed += 1
        except Exception as e:
            print(f"❌ {test.__name__}: {str(e)}")
            failed += 1
    
    print(f"\n📊 Test Results: {passed} passed, {failed} failed")
    return failed == 0


if __name__ == "__main__":
    print("=" * 60)
    print("Running 3D Structural Feature Extraction Tests")
    print("=" * 60)
    
    success = run_all_tests()
    
    print("=" * 60)
    if success:
        print("🎉 All tests passed!")
    else:
        print("❌ Some tests failed!")
    print("=" * 60)
    
    sys.exit(0 if success else 1)
```
