## Supplementary material for "Machine Learning-Driven Prediction of TLR4 Binding Affinity: A Comprehensive Molecular Feature Analysis for Drug Discovery": Codebase: __init__.py

```
"""
TLR4 Binding Affinity Prediction System

A machine learning pipeline for predicting binding free energies of small molecules
to the TLR4 receptor using molecular structure analysis and feature extraction.
"""

__version__ = "0.1.0"
__author__ = "Brandon Yee & Maximilian Rutowski"
__email__ = ""

from .molecular_analysis.features import MolecularFeatures, BindingData, PredictionResult
from .molecular_analysis.extractor import MolecularFeatureExtractor
from .data_processing.preprocessor import DataPreprocessor
from .ml_components.trainer import MLModelTrainer

__all__ = [
    "MolecularFeatures",
    "BindingData", 
    "PredictionResult",
    "MolecularFeatureExtractor",
    "DataPreprocessor",
    "MLModelTrainer"
]
```
