## Supplementary material for "Machine Learning-Driven Prediction of TLR4 Binding Affinity: A Comprehensive Molecular Feature Analysis for Drug Discovery": Codebase: __init__.py

```
"""
Configuration Management Module

Provides configuration management for file paths, model parameters,
and system settings for TLR4 binding prediction.
"""

from .settings import Config, ModelConfig, DataConfig, PathConfig

__all__ = [
    "Config",
    "ModelConfig", 
    "DataConfig",
    "PathConfig"
]
```
