## Supplementary material for "Machine Learning-Driven Prediction of TLR4 Binding Affinity: A Comprehensive Molecular Feature Analysis for Drug Discovery": Codebase: settings.py

```
"""
Configuration settings and management.

This module provides comprehensive configuration management for
the TLR4 binding prediction system, including file paths, model
parameters, and system settings.
"""

import os
import yaml
import json
from pathlib import Path
from typing import Dict, Any, Optional, Union
from dataclasses import dataclass, field
import logging

logger = logging.getLogger(__name__)


@dataclass
class PathConfig:
    """Configuration for file and directory paths."""
    
    # Base directories
    project_root: str = "/home/brand/ember-pm"
    data_dir: str = "binding-data"
    models_dir: str = "models"
    results_dir: str = "results"
    logs_dir: str = "logs"
    
    # Data subdirectories
    raw_data_dir: str = "binding-data/raw"
    processed_data_dir: str = "binding-data/processed"
    pdbqt_dir: str = "binding-data/raw/pdbqt"
    binding_csv: str = "binding-data/processed/processed_logs.csv"
    
    # Model directories
    trained_models_dir: str = "models/trained"
    model_artifacts_dir: str = "models/artifacts"
    
    # Results directories
    figures_dir: str = "results/figures"
    tables_dir: str = "results/tables"
    reports_dir: str = "results/reports"
    
    def __post_init__(self):
        """Convert string paths to Path objects and create directories."""
        # Convert to Path objects
        for field_name in self.__dataclass_fields__:
            field_value = getattr(self, field_name)
            if isinstance(field_value, str):
                setattr(self, field_name, Path(field_value))
        
        # Create directories if they don't exist
        self._create_directories()
    
    def _create_directories(self) -> None:
        """Create necessary directories."""
        directories = [
            self.data_dir,
            self.models_dir,
            self.results_dir,
            self.logs_dir,
            self.raw_data_dir,
            self.processed_data_dir,
            self.pdbqt_dir,
            self.trained_models_dir,
            self.model_artifacts_dir,
            self.figures_dir,
            self.tables_dir,
            self.reports_dir
        ]
        
        for directory in directories:
            directory.mkdir(parents=True, exist_ok=True)


@dataclass
class DataConfig:
    """Configuration for data processing parameters."""
    
    # Data validation
    min_affinity: float = -20.0  # kcal/mol
    max_affinity: float = 5.0    # kcal/mol
    affinity_column: str = "affinity"
    ligand_column: str = "ligand"
    mode_column: str = "mode"
    
    # Data cleaning
    outlier_method: str = "iqr"  # 'iqr', 'zscore', 'isolation_forest'
    outlier_threshold: float = 1.5
    missing_value_strategy: str = "median"  # 'mean', 'median', 'drop'
    
    # Feature engineering
    correlation_threshold: float = 0.95
    feature_selection_method: str = "mutual_info"  # 'mutual_info', 'f_test', 'chi2'
    n_features_select: int = 50
    
    # Data splitting
    train_ratio: float = 0.7
    val_ratio: float = 0.15
    test_ratio: float = 0.15
    random_state: int = 42
    stratify: bool = False
    
    # Compound matching
    fuzzy_threshold: float = 80.0
    use_partial_ratio: bool = True


@dataclass
class ModelConfig:
    """Configuration for machine learning models."""
    
    # General training parameters
    random_state: int = 42
    n_jobs: int = -1
    cv_folds: int = 5
    scoring_metric: str = "neg_mean_squared_error"
    
    # Random Forest parameters
    rf_n_estimators: list = field(default_factory=lambda: [100, 200, 300])
    rf_max_depth: list = field(default_factory=lambda: [10, 20, 30, None])
    rf_min_samples_split: list = field(default_factory=lambda: [2, 5, 10])
    rf_min_samples_leaf: list = field(default_factory=lambda: [1, 2, 4])
    rf_max_features: list = field(default_factory=lambda: ['sqrt', 'log2', None])
    
    # SVR parameters
    svr_C: list = field(default_factory=lambda: [0.1, 1, 10, 100])
    svr_gamma: list = field(default_factory=lambda: ['scale', 'auto', 0.001, 0.01, 0.1, 1])
    svr_kernel: list = field(default_factory=lambda: ['rbf', 'poly', 'linear'])
    svr_epsilon: list = field(default_factory=lambda: [0.01, 0.1, 0.2, 0.5])
    
    # XGBoost parameters
    xgb_n_estimators: list = field(default_factory=lambda: [100, 200, 300])
    xgb_max_depth: list = field(default_factory=lambda: [3, 6, 9, 12])
    xgb_learning_rate: list = field(default_factory=lambda: [0.01, 0.1, 0.2])
    xgb_subsample: list = field(default_factory=lambda: [0.8, 0.9, 1.0])
    xgb_colsample_bytree: list = field(default_factory=lambda: [0.8, 0.9, 1.0])
    
    # LightGBM parameters
    lgb_n_estimators: list = field(default_factory=lambda: [100, 200, 300])
    lgb_max_depth: list = field(default_factory=lambda: [3, 6, 9, 12])
    lgb_learning_rate: list = field(default_factory=lambda: [0.01, 0.1, 0.2])
    lgb_subsample: list = field(default_factory=lambda: [0.8, 0.9, 1.0])
    lgb_colsample_bytree: list = field(default_factory=lambda: [0.8, 0.9, 1.0])
    lgb_num_leaves: list = field(default_factory=lambda: [31, 50, 100])
    
    # Model selection
    models_to_train: list = field(default_factory=lambda: ['random_forest', 'svr', 'xgboost', 'lightgbm'])
    best_model_metric: str = "r2"
    
    # Prediction settings
    confidence_level: float = 0.95
    uncertainty_method: str = "bootstrap"  # 'bootstrap', 'dropout', 'ensemble'


@dataclass
class Config:
    """Main configuration class combining all settings."""
    
    # Sub-configurations
    paths: PathConfig = field(default_factory=PathConfig)
    data: DataConfig = field(default_factory=DataConfig)
    model: ModelConfig = field(default_factory=ModelConfig)
    
    # System settings
    log_level: str = "INFO"
    log_format: str = "%(asctime)s - %(name)s - %(levelname)s - %(message)s"
    log_file: str = "logs/tlr4_binding.log"
    
    # Experiment tracking
    experiment_name: str = "tlr4_binding_prediction"
    experiment_version: str = "0.1.0"
    track_experiments: bool = True
    
    def __post_init__(self):
        """Initialize configuration after creation."""
        # Set up logging
        self._setup_logging()
        
        # Validate configuration
        self._validate_config()
    
    def _setup_logging(self) -> None:
        """Set up logging configuration."""
        # Create logs directory
        self.paths.logs_dir.mkdir(parents=True, exist_ok=True)
        
        # Configure logging
        logging.basicConfig(
            level=getattr(logging, self.log_level.upper()),
            format=self.log_format,
            handlers=[
                logging.FileHandler(self.paths.logs_dir / "tlr4_binding.log"),
                logging.StreamHandler()
            ]
        )
        
        logger.info("Logging configured successfully")
    
    def _validate_config(self) -> None:
        """Validate configuration parameters."""
        # Validate data ratios
        total_ratio = self.data.train_ratio + self.data.val_ratio + self.data.test_ratio
        if abs(total_ratio - 1.0) > 1e-6:
            raise ValueError(f"Data split ratios must sum to 1.0, got {total_ratio}")
        
        # Validate affinity range
        if self.data.min_affinity >= self.data.max_affinity:
            raise ValueError("min_affinity must be less than max_affinity")
        
        # Validate correlation threshold
        if not 0 <= self.data.correlation_threshold <= 1:
            raise ValueError("correlation_threshold must be between 0 and 1")
        
        # Validate fuzzy threshold
        if not 0 <= self.data.fuzzy_threshold <= 100:
            raise ValueError("fuzzy_threshold must be between 0 and 100")
        
        logger.info("Configuration validation completed successfully")
    
    def save_config(self, file_path: Union[str, Path]) -> None:
        """Save configuration to YAML file."""
        file_path = Path(file_path)
        
        # Convert dataclasses to dictionaries
        config_dict = {
            'paths': self._dataclass_to_dict(self.paths),
            'data': self._dataclass_to_dict(self.data),
            'model': self._dataclass_to_dict(self.model),
            'log_level': self.log_level,
            'log_format': self.log_format,
            'experiment_name': self.experiment_name,
            'experiment_version': self.experiment_version,
            'track_experiments': self.track_experiments
        }
        
        # Save to YAML
        with open(file_path, 'w') as f:
            yaml.dump(config_dict, f, default_flow_style=False, indent=2)
        
        logger.info(f"Configuration saved to {file_path}")
    
    @classmethod
    def load_config(cls, file_path: Union[str, Path]) -> 'Config':
        """Load configuration from YAML file."""
        file_path = Path(file_path)
        
        if not file_path.exists():
            raise FileNotFoundError(f"Configuration file not found: {file_path}")
        
        with open(file_path, 'r') as f:
            config_dict = yaml.safe_load(f)
        
        # Create configuration object
        config = cls()
        
        # Update with loaded values
        if 'paths' in config_dict:
            config.paths = PathConfig(**config_dict['paths'])
        
        if 'data' in config_dict:
            config.data = DataConfig(**config_dict['data'])
        
        if 'model' in config_dict:
            config.model = ModelConfig(**config_dict['model'])
        
        # Update other attributes
        for key, value in config_dict.items():
            if key not in ['paths', 'data', 'model']:
                setattr(config, key, value)
        
        logger.info(f"Configuration loaded from {file_path}")
        return config
    
    def _dataclass_to_dict(self, obj) -> Dict[str, Any]:
        """Convert dataclass to dictionary."""
        if hasattr(obj, '__dataclass_fields__'):
            result = {}
            for field_name, field_info in obj.__dataclass_fields__.items():
                value = getattr(obj, field_name)
                if hasattr(value, '__dataclass_fields__'):
                    result[field_name] = self._dataclass_to_dict(value)
                elif isinstance(value, Path):
                    result[field_name] = str(value)
                else:
                    result[field_name] = value
            return result
        return obj
    
    def get_model_param_grid(self, model_name: str) -> Dict[str, list]:
        """Get parameter grid for specific model."""
        if model_name == 'random_forest':
            return {
                'n_estimators': self.model.rf_n_estimators,
                'max_depth': self.model.rf_max_depth,
                'min_samples_split': self.model.rf_min_samples_split,
                'min_samples_leaf': self.model.rf_min_samples_leaf,
                'max_features': self.model.rf_max_features
            }
        elif model_name == 'svr':
            return {
                'C': self.model.svr_C,
                'gamma': self.model.svr_gamma,
                'kernel': self.model.svr_kernel,
                'epsilon': self.model.svr_epsilon
            }
        elif model_name == 'xgboost':
            return {
                'n_estimators': self.model.xgb_n_estimators,
                'max_depth': self.model.xgb_max_depth,
                'learning_rate': self.model.xgb_learning_rate,
                'subsample': self.model.xgb_subsample,
                'colsample_bytree': self.model.xgb_colsample_bytree
            }
        elif model_name == 'lightgbm':
            return {
                'n_estimators': self.model.lgb_n_estimators,
                'max_depth': self.model.lgb_max_depth,
                'learning_rate': self.model.lgb_learning_rate,
                'subsample': self.model.lgb_subsample,
                'colsample_bytree': self.model.lgb_colsample_bytree,
                'num_leaves': self.model.lgb_num_leaves
            }
        else:
            raise ValueError(f"Unknown model: {model_name}")
    
    def update_from_dict(self, updates: Dict[str, Any]) -> None:
        """Update configuration from dictionary."""
        for key, value in updates.items():
            if hasattr(self, key):
                setattr(self, key, value)
            elif hasattr(self.paths, key):
                setattr(self.paths, key, value)
            elif hasattr(self.data, key):
                setattr(self.data, key, value)
            elif hasattr(self.model, key):
                setattr(self.model, key, value)
            else:
                logger.warning(f"Unknown configuration key: {key}")
    
    def to_dict(self) -> Dict[str, Any]:
        """Convert entire configuration to dictionary."""
        return {
            'paths': self._dataclass_to_dict(self.paths),
            'data': self._dataclass_to_dict(self.data),
            'model': self._dataclass_to_dict(self.model),
            'log_level': self.log_level,
            'log_format': self.log_format,
            'log_file': self.log_file,
            'experiment_name': self.experiment_name,
            'experiment_version': self.experiment_version,
            'track_experiments': self.track_experiments
        }


# Global configuration instance
config = Config()

# Convenience functions
def get_config() -> Config:
    """Get global configuration instance."""
    return config

def update_config(updates: Dict[str, Any]) -> None:
    """Update global configuration."""
    config.update_from_dict(updates)

def save_config(file_path: Union[str, Path]) -> None:
    """Save global configuration."""
    config.save_config(file_path)

def load_config(file_path: Union[str, Path]) -> Config:
    """Load configuration and update global instance."""
    global config
    config = Config.load_config(file_path)
    return config
```
