## Supplementary material for "Machine Learning-Driven Prediction of TLR4 Binding Affinity: A Comprehensive Molecular Feature Analysis for Drug Discovery": Codebase: __init__.py

```
"""
Data Processing Module

Contains classes for data loading, preprocessing, integration,
and feature engineering for TLR4 binding prediction.
"""

from .preprocessor import DataPreprocessor, BindingDataLoader, CompoundMatcher, DataIntegrator
from .feature_engineering import FeatureEngineeringPipeline, FeatureScaler, FeatureSelector
from .validation import DataValidator, OutlierDetector

__all__ = [
    "DataPreprocessor",
    "BindingDataLoader", 
    "CompoundMatcher",
    "DataIntegrator",
    "FeatureEngineeringPipeline",
    "FeatureScaler",
    "FeatureSelector",
    "DataValidator",
    "OutlierDetector"
]
```
