## Supplementary material for "Machine Learning-Driven Prediction of TLR4 Binding Affinity: A Comprehensive Molecular Feature Analysis for Drug Discovery": Codebase: feature_engineering.py

```
"""
Feature engineering and preprocessing utilities.

This module provides comprehensive feature engineering functionality
for molecular descriptors including scaling, selection, and transformation.
"""

from abc import ABC, abstractmethod
from typing import Dict, List, Optional, Union, Tuple, Any
import pandas as pd
import numpy as np
import logging
from sklearn.preprocessing import StandardScaler, RobustScaler, MinMaxScaler
from sklearn.feature_selection import SelectKBest, mutual_info_regression, f_regression
from sklearn.decomposition import PCA
from sklearn.pipeline import Pipeline

logger = logging.getLogger(__name__)


class FeatureScalerInterface(ABC):
    """Abstract interface for feature scaling."""
    
    @abstractmethod
    def fit(self, X: pd.DataFrame) -> 'FeatureScalerInterface':
        """Fit scaler to data."""
        pass
    
    @abstractmethod
    def transform(self, X: pd.DataFrame) -> pd.DataFrame:
        """Transform data using fitted scaler."""
        pass
    
    @abstractmethod
    def fit_transform(self, X: pd.DataFrame) -> pd.DataFrame:
        """Fit scaler and transform data."""
        pass


class FeatureScaler(FeatureScalerInterface):
    """Comprehensive feature scaling with multiple scaling methods."""
    
    def __init__(self, method: str = 'standard', **kwargs):
        """
        Initialize feature scaler.
        
        Args:
            method: Scaling method ('standard', 'robust', 'minmax')
            **kwargs: Additional parameters for scaler
        """
        self.method = method
        self.kwargs = kwargs
        self.scaler = None
        self.feature_names = None
        
        # Initialize appropriate scaler
        if method == 'standard':
            self.scaler = StandardScaler(**kwargs)
        elif method == 'robust':
            self.scaler = RobustScaler(**kwargs)
        elif method == 'minmax':
            self.scaler = MinMaxScaler(**kwargs)
        else:
            raise ValueError(f"Unknown scaling method: {method}")
    
    def fit(self, X: pd.DataFrame) -> 'FeatureScaler':
        """Fit scaler to data."""
        self.feature_names = X.columns.tolist()
        self.scaler.fit(X)
        return self
    
    def transform(self, X: pd.DataFrame) -> pd.DataFrame:
        """Transform data using fitted scaler."""
        if self.scaler is None:
            raise ValueError("Scaler must be fitted before transform")
        
        # Ensure same columns as training data
        X_scaled = self.scaler.transform(X)
        return pd.DataFrame(X_scaled, columns=self.feature_names, index=X.index)
    
    def fit_transform(self, X: pd.DataFrame) -> pd.DataFrame:
        """Fit scaler and transform data."""
        return self.fit(X).transform(X)


class FeatureSelectorInterface(ABC):
    """Abstract interface for feature selection."""
    
    @abstractmethod
    def fit(self, X: pd.DataFrame, y: pd.Series) -> 'FeatureSelectorInterface':
        """Fit selector to data."""
        pass
    
    @abstractmethod
    def transform(self, X: pd.DataFrame) -> pd.DataFrame:
        """Transform data using fitted selector."""
        pass
    
    @abstractmethod
    def get_feature_importance(self) -> Dict[str, float]:
        """Get feature importance scores."""
        pass


class FeatureSelector(FeatureSelectorInterface):
    """Comprehensive feature selection with multiple methods."""
    
    def __init__(self, method: str = 'mutual_info', k: int = 50, **kwargs):
        """
        Initialize feature selector.
        
        Args:
            method: Selection method ('mutual_info', 'f_test', 'pca')
            k: Number of features to select
            **kwargs: Additional parameters for selector
        """
        self.method = method
        self.k = k
        self.kwargs = kwargs
        self.selector = None
        self.selected_features = None
        self.feature_scores = None
        
        # Initialize appropriate selector
        if method == 'mutual_info':
            self.selector = SelectKBest(score_func=mutual_info_regression, k=k)
        elif method == 'f_test':
            self.selector = SelectKBest(score_func=f_regression, k=k)
        elif method == 'pca':
            self.selector = PCA(n_components=k, **kwargs)
        else:
            raise ValueError(f"Unknown selection method: {method}")
    
    def fit(self, X: pd.DataFrame, y: pd.Series) -> 'FeatureSelector':
        """Fit selector to data."""
        self.selector.fit(X, y)
        
        if hasattr(self.selector, 'get_support'):
            # For SelectKBest
            self.selected_features = X.columns[self.selector.get_support()].tolist()
            self.feature_scores = dict(zip(X.columns, self.selector.scores_))
        else:
            # For PCA
            self.selected_features = [f'PC{i+1}' for i in range(self.k)]
            self.feature_scores = dict(zip(X.columns, self.selector.explained_variance_ratio_))
        
        return self
    
    def transform(self, X: pd.DataFrame) -> pd.DataFrame:
        """Transform data using fitted selector."""
        if self.selector is None:
            raise ValueError("Selector must be fitted before transform")
        
        X_selected = self.selector.transform(X)
        
        if hasattr(self.selector, 'get_support'):
            return pd.DataFrame(X_selected, columns=self.selected_features, index=X.index)
        else:
            return pd.DataFrame(X_selected, columns=self.selected_features, index=X.index)
    
    def get_feature_importance(self) -> Dict[str, float]:
        """Get feature importance scores."""
        if self.feature_scores is None:
            raise ValueError("Selector must be fitted before getting importance")
        return self.feature_scores


class FeatureEngineeringPipeline:
    """Complete feature engineering pipeline."""
    
    def __init__(self, scaler_method: str = 'standard', 
                 selector_method: str = 'mutual_info',
                 n_features: int = 50,
                 correlation_threshold: float = 0.95):
        """
        Initialize feature engineering pipeline.
        
        Args:
            scaler_method: Scaling method
            selector_method: Feature selection method
            n_features: Number of features to select
            correlation_threshold: Threshold for removing correlated features
        """
        self.scaler_method = scaler_method
        self.selector_method = selector_method
        self.n_features = n_features
        self.correlation_threshold = correlation_threshold
        
        # Initialize components
        self.scaler = FeatureScaler(method=scaler_method)
        self.selector = FeatureSelector(method=selector_method, k=n_features)
        self.pipeline = None
        
        # Statistics
        self.correlation_removed = 0
        self.original_features = 0
        self.final_features = 0
    
    def fit_transform(self, X: pd.DataFrame, y: pd.Series) -> pd.DataFrame:
        """
        Fit pipeline and transform features.
        
        Args:
            X: Feature matrix
            y: Target vector
            
        Returns:
            Transformed feature matrix
        """
        logger.info("Starting feature engineering pipeline")
        
        # Store original feature count
        self.original_features = X.shape[1]
        
        # Remove highly correlated features
        X_cleaned = self._remove_correlated_features(X)
        
        # Scale features
        X_scaled = self.scaler.fit_transform(X_cleaned)
        
        # Select features
        X_selected = self.selector.fit(X_scaled, y).transform(X_scaled)
        
        # Store final feature count
        self.final_features = X_selected.shape[1]
        
        logger.info(f"Feature engineering completed: {self.original_features} -> {self.final_features} features")
        logger.info(f"Removed {self.correlation_removed} highly correlated features")
        
        return X_selected
    
    def transform(self, X: pd.DataFrame) -> pd.DataFrame:
        """
        Transform new data using fitted pipeline.
        
        Args:
            X: Feature matrix
            
        Returns:
            Transformed feature matrix
        """
        if self.scaler is None or self.selector is None:
            raise ValueError("Pipeline must be fitted before transform")
        
        # Remove correlated features (using same features as training)
        X_cleaned = self._remove_correlated_features(X)
        
        # Scale features
        X_scaled = self.scaler.transform(X_cleaned)
        
        # Select features
        X_selected = self.selector.transform(X_scaled)
        
        return X_selected
    
    def _remove_correlated_features(self, X: pd.DataFrame) -> pd.DataFrame:
        """Remove highly correlated features."""
        # Calculate correlation matrix
        corr_matrix = X.corr().abs()
        
        # Find pairs above threshold
        upper_tri = corr_matrix.where(
            np.triu(np.ones(corr_matrix.shape), k=1).astype(bool)
        )
        
        # Find features to drop
        to_drop = [column for column in upper_tri.columns 
                  if any(upper_tri[column] > self.correlation_threshold)]
        
        # Count removed features
        self.correlation_removed = len(to_drop)
        
        # Remove correlated features
        X_cleaned = X.drop(columns=to_drop)
        
        if to_drop:
            logger.info(f"Removed {len(to_drop)} highly correlated features: {to_drop}")
        
        return X_cleaned
    
    def get_feature_importance(self) -> Dict[str, float]:
        """Get feature importance from selector."""
        return self.selector.get_feature_importance()
    
    def get_pipeline_stats(self) -> Dict[str, Any]:
        """Get pipeline statistics."""
        return {
            'original_features': self.original_features,
            'final_features': self.final_features,
            'correlation_removed': self.correlation_removed,
            'scaler_method': self.scaler_method,
            'selector_method': self.selector_method,
            'correlation_threshold': self.correlation_threshold
        }
```
