## Supplementary material for "Machine Learning-Driven Prediction of TLR4 Binding Affinity: A Comprehensive Molecular Feature Analysis for Drug Discovery": Codebase: imputation.py

```
"""
Advanced data imputation utilities for molecular features.

This module provides sophisticated imputation methods for handling
missing values in molecular feature datasets.
"""

import pandas as pd
import numpy as np
import logging
from typing import Dict, List, Optional, Tuple, Any
from sklearn.impute import SimpleImputer, KNNImputer
try:
    from sklearn.experimental import enable_iterative_imputer
    from sklearn.impute import IterativeImputer
    ITERATIVE_AVAILABLE = True
except ImportError:
    ITERATIVE_AVAILABLE = False
from sklearn.preprocessing import StandardScaler
from sklearn.ensemble import RandomForestRegressor
import warnings
warnings.filterwarnings('ignore')

logger = logging.getLogger(__name__)


class MolecularFeatureImputer:
    """
    Advanced imputation for molecular features with domain knowledge.
    
    Handles missing values in molecular descriptors using multiple
    imputation strategies tailored for chemical data.
    """
    
    def __init__(self, strategy: str = 'adaptive', n_neighbors: int = 5):
        """
        Initialize molecular feature imputer.
        
        Args:
            strategy: Imputation strategy ('simple', 'knn', 'iterative', 'adaptive')
            n_neighbors: Number of neighbors for KNN imputation
        """
        self.strategy = strategy
        self.n_neighbors = n_neighbors
        self.imputers = {}
        self.scalers = {}
        self.feature_groups = self._define_feature_groups()
        
    def _define_feature_groups(self) -> Dict[str, List[str]]:
        """Define groups of related molecular features."""
        return {
            'basic_properties': [
                'molecular_weight', 'heavy_atoms', 'heteroatoms',
                'formal_charge', 'rotatable_bonds'
            ],
            'lipophilicity': [
                'logp', 'molar_refractivity', 'tpsa'
            ],
            'hydrogen_bonding': [
                'hbd', 'hba', 'tpsa'
            ],
            'ring_features': [
                'ring_count', 'aromatic_rings', 'aliphatic_rings',
                'saturated_rings', 'aromatic_ratio'
            ],
            'shape_descriptors': [
                'radius_of_gyration', 'asphericity', 'eccentricity',
                'spherocity_index', 'elongation', 'flatness'
            ],
            'surface_properties': [
                'surface_area', 'polar_surface_area', 'hydrophobic_surface_area',
                'molecular_volume', 'compactness'
            ],
            'connectivity': [
                'balaban_j', 'bertz_ct', 'chi0v', 'chi1v', 'chi2v', 'chi3v', 'chi4v'
            ],
            'coordinate_features': [
                'coord_radius_of_gyration', 'coord_max_distance', 'coord_mean_distance',
                'coord_length', 'coord_width', 'coord_height', 'coord_volume',
                'coord_elongation', 'coord_flatness', 'coord_total_atoms'
            ]
        }
    
    def fit_transform(self, df: pd.DataFrame) -> pd.DataFrame:
        """
        Fit imputers and transform data.
        
        Args:
            df: DataFrame with molecular features
            
        Returns:
            DataFrame with imputed values
        """
        logger.info(f"Fitting imputers using {self.strategy} strategy")
        
        # Separate numeric and non-numeric columns
        numeric_cols = df.select_dtypes(include=[np.number]).columns.tolist()
        non_numeric_cols = df.select_dtypes(exclude=[np.number]).columns.tolist()
        
        # Remove metadata columns from imputation
        metadata_cols = ['compound_name', 'pdbqt_file', 'smiles', 'inchi', 'ligand', 'matched_compound']
        numeric_cols = [col for col in numeric_cols if col not in metadata_cols]
        
        if not numeric_cols:
            logger.warning("No numeric columns found for imputation")
            return df
        
        # Count missing values
        missing_before = df[numeric_cols].isna().sum().sum()
        logger.info(f"Missing values before imputation: {missing_before}")
        
        # Apply imputation strategy
        df_imputed = df.copy()
        
        if self.strategy == 'adaptive':
            df_imputed = self._adaptive_imputation(df_imputed, numeric_cols)
        elif self.strategy == 'knn':
            df_imputed = self._knn_imputation(df_imputed, numeric_cols)
        elif self.strategy == 'iterative':
            df_imputed = self._iterative_imputation(df_imputed, numeric_cols)
        else:  # simple
            df_imputed = self._simple_imputation(df_imputed, numeric_cols)
        
        # Count missing values after imputation
        missing_after = df_imputed[numeric_cols].isna().sum().sum()
        logger.info(f"Missing values after imputation: {missing_after}")
        logger.info(f"Imputed {missing_before - missing_after} missing values")
        
        # Final fallback: fill any remaining NaN values with 0
        if missing_after > 0:
            logger.warning(f"Still have {missing_after} missing values after imputation. Filling with zeros as final fallback.")
            df_imputed[numeric_cols] = df_imputed[numeric_cols].fillna(0.0)
            final_missing = df_imputed[numeric_cols].isna().sum().sum()
            logger.info(f"Missing values after final fallback: {final_missing}")
        
        return df_imputed
    
    def transform(self, df: pd.DataFrame) -> pd.DataFrame:
        """
        Transform new data using fitted imputers.
        
        Args:
            df: DataFrame to transform
            
        Returns:
            DataFrame with imputed values
        """
        if not self.imputers:
            raise ValueError("Imputers not fitted. Call fit_transform first.")
        
        df_transformed = df.copy()
        
        for group_name, imputer in self.imputers.items():
            group_features = self.feature_groups.get(group_name, [])
            available_features = [col for col in group_features if col in df.columns]
            
            if available_features:
                # Apply scaling if available
                if group_name in self.scalers:
                    scaled_data = self.scalers[group_name].transform(df[available_features])
                    imputed_data = imputer.transform(scaled_data)
                    # Inverse transform
                    df_transformed[available_features] = self.scalers[group_name].inverse_transform(imputed_data)
                else:
                    df_transformed[available_features] = imputer.transform(df[available_features])
        
        return df_transformed
    
    def _adaptive_imputation(self, df: pd.DataFrame, numeric_cols: List[str]) -> pd.DataFrame:
        """
        Adaptive imputation using different strategies for different feature groups.
        
        Args:
            df: DataFrame to impute
            numeric_cols: List of numeric columns
            
        Returns:
            DataFrame with imputed values
        """
        logger.info("Applying adaptive imputation strategy")
        
        for group_name, group_features in self.feature_groups.items():
            # Get available features in this group
            available_features = [col for col in group_features if col in numeric_cols]
            
            if not available_features:
                continue
            
            logger.info(f"Imputing {group_name}: {len(available_features)} features")
            
            # Check if all values are NaN for this group
            group_data = df[available_features]
            if group_data.isna().all().all():
                logger.warning(f"All values are NaN for {group_name}, filling with zeros")
                df[available_features] = 0.0
                continue
            
            # Choose imputation method based on feature group
            if group_name in ['coordinate_features', 'shape_descriptors']:
                # Use KNN for geometric features (they're often correlated)
                imputer = KNNImputer(n_neighbors=min(5, len(df)//2))
            elif group_name in ['connectivity', 'ring_features'] and ITERATIVE_AVAILABLE:
                # Use iterative imputation for complex molecular descriptors
                imputer = IterativeImputer(
                    estimator=RandomForestRegressor(n_estimators=10, random_state=42),
                    max_iter=5, random_state=42
                )
            else:
                # Use median for basic properties
                imputer = SimpleImputer(strategy='median')
            
            try:
                # Simple imputation for all features in this group
                imputed_data = imputer.fit_transform(df[available_features])
                if imputed_data.shape[1] > 0:  # Check if we have data
                    df[available_features] = pd.DataFrame(imputed_data, columns=available_features, index=df.index)
                
                self.imputers[group_name] = imputer
                
            except Exception as e:
                logger.warning(f"Imputation failed for {group_name}: {e}. Using median fallback.")
                # Fallback to simple median imputation
                try:
                    fallback_imputer = SimpleImputer(strategy='median')
                    imputed_data = fallback_imputer.fit_transform(df[available_features])
                    df[available_features] = pd.DataFrame(imputed_data, columns=available_features, index=df.index)
                    self.imputers[group_name] = fallback_imputer
                except Exception as e2:
                    logger.warning(f"Fallback imputation also failed for {group_name}: {e2}. Filling with zeros.")
                    df[available_features] = df[available_features].fillna(0.0)
        
        return df
    
    def _knn_imputation(self, df: pd.DataFrame, numeric_cols: List[str]) -> pd.DataFrame:
        """KNN imputation for all features."""
        logger.info("Applying KNN imputation")
        
        imputer = KNNImputer(n_neighbors=self.n_neighbors)
        df[numeric_cols] = imputer.fit_transform(df[numeric_cols])
        self.imputers['knn'] = imputer
        
        return df
    
    def _iterative_imputation(self, df: pd.DataFrame, numeric_cols: List[str]) -> pd.DataFrame:
        """Iterative imputation using RandomForest."""
        if not ITERATIVE_AVAILABLE:
            logger.warning("IterativeImputer not available, falling back to KNN imputation")
            return self._knn_imputation(df, numeric_cols)
            
        logger.info("Applying iterative imputation")
        
        imputer = IterativeImputer(
            estimator=RandomForestRegressor(n_estimators=10, random_state=42),
            max_iter=10, random_state=42
        )
        df[numeric_cols] = imputer.fit_transform(df[numeric_cols])
        self.imputers['iterative'] = imputer
        
        return df
    
    def _simple_imputation(self, df: pd.DataFrame, numeric_cols: List[str]) -> pd.DataFrame:
        """Simple median imputation."""
        logger.info("Applying simple median imputation")
        
        imputer = SimpleImputer(strategy='median')
        df[numeric_cols] = imputer.fit_transform(df[numeric_cols])
        self.imputers['simple'] = imputer
        
        return df
    
    def get_imputation_summary(self, df_before: pd.DataFrame, df_after: pd.DataFrame) -> Dict[str, Any]:
        """
        Get summary of imputation results.
        
        Args:
            df_before: DataFrame before imputation
            df_after: DataFrame after imputation
            
        Returns:
            Dictionary with imputation summary
        """
        numeric_cols = df_before.select_dtypes(include=[np.number]).columns.tolist()
        metadata_cols = ['compound_name', 'pdbqt_file', 'smiles', 'inchi', 'ligand', 'matched_compound']
        numeric_cols = [col for col in numeric_cols if col not in metadata_cols]
        
        summary = {
            'total_features': len(numeric_cols),
            'missing_before': df_before[numeric_cols].isna().sum().sum(),
            'missing_after': df_after[numeric_cols].isna().sum().sum(),
            'imputation_rate': 0.0,
            'feature_summary': []
        }
        
        if summary['missing_before'] > 0:
            summary['imputation_rate'] = (summary['missing_before'] - summary['missing_after']) / summary['missing_before']
        
        # Per-feature summary
        for col in numeric_cols:
            missing_before = df_before[col].isna().sum()
            missing_after = df_after[col].isna().sum()
            
            if missing_before > 0:
                summary['feature_summary'].append({
                    'feature': col,
                    'missing_before': missing_before,
                    'missing_after': missing_after,
                    'imputed': missing_before - missing_after,
                    'imputation_rate': (missing_before - missing_after) / missing_before
                })
        
        return summary


def create_robust_dataset(features_df: pd.DataFrame, 
                         binding_df: Optional[pd.DataFrame] = None,
                         imputation_strategy: str = 'adaptive') -> pd.DataFrame:
    """
    Create a robust dataset with comprehensive preprocessing.
    
    Args:
        features_df: DataFrame with molecular features
        binding_df: Optional DataFrame with binding data
        imputation_strategy: Imputation strategy to use
        
    Returns:
        Preprocessed DataFrame ready for ML
    """
    logger.info("Creating robust dataset with comprehensive preprocessing")
    
    # Initialize imputer
    imputer = MolecularFeatureImputer(strategy=imputation_strategy)
    
    # Apply imputation
    df_imputed = imputer.fit_transform(features_df)
    
    # Merge with binding data if provided
    if binding_df is not None:
        # Assume binding_df has been preprocessed to get best affinities
        df_final = pd.merge(df_imputed, binding_df, 
                           left_on='compound_name', right_on='ligand', 
                           how='inner')
    else:
        df_final = df_imputed
    
    # Validate final dataset
    numeric_cols = df_final.select_dtypes(include=[np.number]).columns.tolist()
    metadata_cols = ['compound_name', 'pdbqt_file', 'smiles', 'inchi', 'ligand', 'matched_compound']
    feature_cols = [col for col in numeric_cols if col not in metadata_cols]
    
    remaining_missing = df_final[feature_cols].isna().sum().sum()
    if remaining_missing > 0:
        logger.warning(f"Still have {remaining_missing} missing values after imputation")
    
    logger.info(f"Final dataset: {len(df_final)} samples, {len(feature_cols)} features")
    
    return df_final
```
