## Supplementary material for "Machine Learning-Driven Prediction of TLR4 Binding Affinity: A Comprehensive Molecular Feature Analysis for Drug Discovery": Codebase: molecular_splitting.py

```
"""
Molecular-aware data splitting strategies for drug discovery.

This module implements proper data splitting strategies that prevent data leakage
in molecular datasets by considering chemical similarity and scaffold diversity.
"""

import pandas as pd
import numpy as np
import logging
from typing import Dict, List, Optional, Tuple, Union
from dataclasses import dataclass
from sklearn.model_selection import train_test_split
import warnings

logger = logging.getLogger(__name__)

# RDKit imports with error handling
try:
    from rdkit import Chem
    from rdkit.Chem import Descriptors
    from rdkit.Chem.Scaffolds import MurckoScaffold
    from rdkit import DataStructs
    from rdkit.Chem import rdMolDescriptors
    RDKIT_AVAILABLE = True
except ImportError:
    logger.warning("RDKit not available. Molecular splitting will use fallback methods.")
    RDKIT_AVAILABLE = False


@dataclass
class SplitResult:
    """Container for data splitting results."""
    train_indices: List[int]
    val_indices: List[int]
    test_indices: List[int]
    split_info: Dict[str, any]


class MolecularScaffoldSplitter:
    """
    Molecular scaffold-based data splitting for drug discovery.
    
    Implements Bemis-Murcko scaffold splitting to ensure that compounds
    with similar scaffolds don't appear in both training and test sets,
    preventing data leakage in molecular property prediction.
    """
    
    def __init__(self, include_chirality: bool = False):
        """
        Initialize scaffold splitter.
        
        Args:
            include_chirality: Whether to include chirality in scaffold generation
        """
        self.include_chirality = include_chirality
        
    def split(self, 
              smiles_list: List[str], 
              test_size: float = 0.2,
              val_size: float = 0.1,
              random_state: int = 42) -> SplitResult:
        """
        Split data based on molecular scaffolds.
        
        Args:
            smiles_list: List of SMILES strings
            test_size: Fraction of data for test set
            val_size: Fraction of data for validation set
            random_state: Random state for reproducibility
            
        Returns:
            SplitResult with train/val/test indices and split information
        """
        if not RDKIT_AVAILABLE:
            logger.warning("RDKit not available. Using random split as fallback.")
            return self._fallback_random_split(smiles_list, test_size, val_size, random_state)
        
        logger.info("Performing scaffold-based molecular splitting")
        
        # Generate scaffolds for all molecules
        scaffolds = self._generate_scaffolds(smiles_list)
        
        # Group molecules by scaffold
        scaffold_to_indices = {}
        for idx, scaffold in enumerate(scaffolds):
            if scaffold not in scaffold_to_indices:
                scaffold_to_indices[scaffold] = []
            scaffold_to_indices[scaffold].append(idx)
        
        # Sort scaffolds by size (largest first) for balanced splitting
        sorted_scaffolds = sorted(scaffold_to_indices.items(), 
                                key=lambda x: len(x[1]), reverse=True)
        
        # Check if we have enough scaffolds for meaningful splitting
        if len(sorted_scaffolds) < 3:
            logger.warning(f"Only {len(sorted_scaffolds)} unique scaffolds found. Using random split as fallback.")
            return self._fallback_random_split(smiles_list, test_size, val_size, random_state)
        
        # Allocate scaffolds to splits
        train_indices, val_indices, test_indices = self._allocate_scaffolds(
            sorted_scaffolds, len(smiles_list), test_size, val_size, random_state
        )
        
        # Calculate split statistics
        split_info = self._calculate_split_info(
            scaffolds, scaffold_to_indices, train_indices, val_indices, test_indices
        )
        
        logger.info(f"Scaffold split completed:")
        logger.info(f"  Train: {len(train_indices)} compounds ({len(train_indices)/len(smiles_list)*100:.1f}%)")
        logger.info(f"  Val: {len(val_indices)} compounds ({len(val_indices)/len(smiles_list)*100:.1f}%)")
        logger.info(f"  Test: {len(test_indices)} compounds ({len(test_indices)/len(smiles_list)*100:.1f}%)")
        logger.info(f"  Unique scaffolds: {split_info['total_scaffolds']}")
        logger.info(f"  Scaffold overlap: {split_info['scaffold_overlap']:.1%}")
        
        return SplitResult(
            train_indices=train_indices,
            val_indices=val_indices,
            test_indices=test_indices,
            split_info=split_info
        )
    
    def _generate_scaffolds(self, smiles_list: List[str]) -> List[str]:
        """Generate Bemis-Murcko scaffolds for molecules."""
        scaffolds = []
        
        for i, smiles in enumerate(smiles_list):
            try:
                # Handle None or empty SMILES
                if smiles is None or smiles == '' or pd.isna(smiles):
                    # Create unique scaffold for each invalid SMILES to enable proper splitting
                    scaffolds.append(f"invalid_scaffold_{i}")
                    continue
                    
                mol = Chem.MolFromSmiles(smiles)
                if mol is not None:
                    scaffold = MurckoScaffold.GetScaffoldForMol(mol)
                    scaffold_smiles = Chem.MolToSmiles(scaffold, 
                                                     isomericSmiles=self.include_chirality)
                    scaffolds.append(scaffold_smiles)
                else:
                    # Create unique scaffold for invalid SMILES
                    scaffolds.append(f"invalid_scaffold_{i}")
            except Exception as e:
                logger.warning(f"Error generating scaffold for {smiles}: {e}")
                # Create unique scaffold for problematic SMILES
                scaffolds.append(f"invalid_scaffold_{i}")
        
        return scaffolds
    
    def _allocate_scaffolds(self, 
                          sorted_scaffolds: List[Tuple[str, List[int]]], 
                          total_size: int,
                          test_size: float, 
                          val_size: float,
                          random_state: int) -> Tuple[List[int], List[int], List[int]]:
        """Allocate scaffolds to train/val/test splits."""
        np.random.seed(random_state)
        
        target_test_size = int(total_size * test_size)
        target_val_size = int(total_size * val_size)
        
        train_indices = []
        val_indices = []
        test_indices = []
        
        # Shuffle scaffolds to randomize allocation order
        scaffolds_copy = sorted_scaffolds.copy()
        np.random.shuffle(scaffolds_copy)
        
        for scaffold, indices in scaffolds_copy:
            # Decide which split to add this scaffold to
            current_test_size = len(test_indices)
            current_val_size = len(val_indices)
            
            if current_test_size < target_test_size:
                test_indices.extend(indices)
            elif current_val_size < target_val_size:
                val_indices.extend(indices)
            else:
                train_indices.extend(indices)
        
        return train_indices, val_indices, test_indices
    
    def _calculate_split_info(self, 
                            scaffolds: List[str],
                            scaffold_to_indices: Dict[str, List[int]],
                            train_indices: List[int], 
                            val_indices: List[int], 
                            test_indices: List[int]) -> Dict[str, any]:
        """Calculate statistics about the scaffold split."""
        
        # Get scaffolds in each split
        train_scaffolds = set(scaffolds[i] for i in train_indices)
        val_scaffolds = set(scaffolds[i] for i in val_indices)
        test_scaffolds = set(scaffolds[i] for i in test_indices)
        
        # Calculate overlap
        all_scaffolds = train_scaffolds | val_scaffolds | test_scaffolds
        overlapping_scaffolds = (train_scaffolds & val_scaffolds) | \
                              (train_scaffolds & test_scaffolds) | \
                              (val_scaffolds & test_scaffolds)
        
        scaffold_overlap = len(overlapping_scaffolds) / len(all_scaffolds) if all_scaffolds else 0
        
        return {
            'total_scaffolds': len(all_scaffolds),
            'train_scaffolds': len(train_scaffolds),
            'val_scaffolds': len(val_scaffolds),
            'test_scaffolds': len(test_scaffolds),
            'scaffold_overlap': scaffold_overlap,
            'overlapping_scaffolds': list(overlapping_scaffolds),
            'scaffold_sizes': {scaffold: len(indices) 
                             for scaffold, indices in scaffold_to_indices.items()}
        }
    
    def _fallback_random_split(self, 
                             smiles_list: List[str], 
                             test_size: float, 
                             val_size: float,
                             random_state: int) -> SplitResult:
        """Fallback to random splitting when RDKit is not available."""
        indices = list(range(len(smiles_list)))
        
        # First split: separate test set
        train_val_indices, test_indices = train_test_split(
            indices, test_size=test_size, random_state=random_state
        )
        
        # Second split: separate validation from training
        if val_size > 0:
            val_size_adjusted = val_size / (1 - test_size)  # Adjust for remaining data
            train_indices, val_indices = train_test_split(
                train_val_indices, test_size=val_size_adjusted, random_state=random_state
            )
        else:
            train_indices = train_val_indices
            val_indices = []
        
        split_info = {
            'method': 'random_fallback',
            'total_scaffolds': len(smiles_list),  # Each molecule is its own "scaffold"
            'scaffold_overlap': 0.0,
            'warning': 'RDKit not available, used random split instead of scaffold split'
        }
        
        return SplitResult(
            train_indices=train_indices,
            val_indices=val_indices,
            test_indices=test_indices,
            split_info=split_info
        )


class TemporalSplitter:
    """
    Temporal data splitting for time-series molecular data.
    
    Splits data based on temporal information to simulate real-world
    deployment where models are trained on historical data and tested
    on future data.
    """
    
    def __init__(self):
        """Initialize temporal splitter."""
        pass
    
    def split(self, 
              timestamps: List[Union[str, pd.Timestamp]], 
              test_size: float = 0.2,
              val_size: float = 0.1) -> SplitResult:
        """
        Split data based on temporal order.
        
        Args:
            timestamps: List of timestamps or date strings
            test_size: Fraction of most recent data for test set
            val_size: Fraction of data for validation set
            
        Returns:
            SplitResult with chronologically ordered splits
        """
        logger.info("Performing temporal data splitting")
        
        # Convert to pandas timestamps if needed
        if isinstance(timestamps[0], str):
            timestamps = pd.to_datetime(timestamps)
        
        # Sort by timestamp
        sorted_indices = np.argsort(timestamps)
        
        # Calculate split points
        n_total = len(timestamps)
        n_test = int(n_total * test_size)
        n_val = int(n_total * val_size)
        n_train = n_total - n_test - n_val
        
        # Assign indices (chronological order)
        train_indices = sorted_indices[:n_train].tolist()
        val_indices = sorted_indices[n_train:n_train + n_val].tolist()
        test_indices = sorted_indices[n_train + n_val:].tolist()
        
        # Calculate split info
        train_period = (timestamps[sorted_indices[0]], timestamps[sorted_indices[n_train-1]])
        val_period = (timestamps[sorted_indices[n_train]], 
                     timestamps[sorted_indices[n_train + n_val - 1]]) if n_val > 0 else None
        test_period = (timestamps[sorted_indices[n_train + n_val]], 
                      timestamps[sorted_indices[-1]])
        
        split_info = {
            'method': 'temporal',
            'train_period': train_period,
            'val_period': val_period,
            'test_period': test_period,
            'temporal_gap': (test_period[0] - train_period[1]).days if val_size == 0 else 
                           (val_period[0] - train_period[1]).days
        }
        
        logger.info(f"Temporal split completed:")
        logger.info(f"  Train period: {train_period[0]} to {train_period[1]}")
        if val_period:
            logger.info(f"  Val period: {val_period[0]} to {val_period[1]}")
        logger.info(f"  Test period: {test_period[0]} to {test_period[1]}")
        
        return SplitResult(
            train_indices=train_indices,
            val_indices=val_indices,
            test_indices=test_indices,
            split_info=split_info
        )


class ClusterBasedSplitter:
    """
    Cluster-based data splitting using molecular similarity.
    
    Groups molecules by similarity and ensures that similar molecules
    don't appear in both training and test sets.
    """
    
    def __init__(self, n_clusters: Optional[int] = None, similarity_threshold: float = 0.7):
        """
        Initialize cluster-based splitter.
        
        Args:
            n_clusters: Number of clusters (auto-determined if None)
            similarity_threshold: Tanimoto similarity threshold for clustering
        """
        self.n_clusters = n_clusters
        self.similarity_threshold = similarity_threshold
    
    def split(self, 
              smiles_list: List[str], 
              test_size: float = 0.2,
              val_size: float = 0.1,
              random_state: int = 42) -> SplitResult:
        """
        Split data based on molecular similarity clusters.
        
        Args:
            smiles_list: List of SMILES strings
            test_size: Fraction of data for test set
            val_size: Fraction of data for validation set
            random_state: Random state for reproducibility
            
        Returns:
            SplitResult with cluster-based splits
        """
        if not RDKIT_AVAILABLE:
            logger.warning("RDKit not available. Using random split as fallback.")
            return MolecularScaffoldSplitter()._fallback_random_split(
                smiles_list, test_size, val_size, random_state
            )
        
        logger.info("Performing cluster-based molecular splitting")
        
        # Generate molecular fingerprints
        fingerprints = self._generate_fingerprints(smiles_list)
        
        # Perform clustering
        clusters = self._cluster_molecules(fingerprints, random_state)
        
        # Allocate clusters to splits
        train_indices, val_indices, test_indices = self._allocate_clusters(
            clusters, len(smiles_list), test_size, val_size, random_state
        )
        
        split_info = {
            'method': 'cluster_based',
            'n_clusters': len(set(clusters)),
            'similarity_threshold': self.similarity_threshold,
            'cluster_sizes': {i: clusters.count(i) for i in set(clusters)}
        }
        
        logger.info(f"Cluster-based split completed:")
        logger.info(f"  Clusters: {len(set(clusters))}")
        logger.info(f"  Train: {len(train_indices)} compounds")
        logger.info(f"  Val: {len(val_indices)} compounds")
        logger.info(f"  Test: {len(test_indices)} compounds")
        
        return SplitResult(
            train_indices=train_indices,
            val_indices=val_indices,
            test_indices=test_indices,
            split_info=split_info
        )
    
    def _generate_fingerprints(self, smiles_list: List[str]) -> List:
        """Generate molecular fingerprints for clustering."""
        fingerprints = []
        
        for smiles in smiles_list:
            try:
                mol = Chem.MolFromSmiles(smiles)
                if mol is not None:
                    fp = rdMolDescriptors.GetMorganFingerprintAsBitVect(mol, 2, nBits=2048)
                    fingerprints.append(fp)
                else:
                    # Create dummy fingerprint for invalid SMILES
                    fingerprints.append(None)
            except Exception as e:
                logger.warning(f"Error generating fingerprint for {smiles}: {e}")
                fingerprints.append(None)
        
        return fingerprints
    
    def _cluster_molecules(self, fingerprints: List, random_state: int) -> List[int]:
        """Cluster molecules based on fingerprint similarity."""
        # Simple clustering based on similarity threshold
        clusters = []
        cluster_representatives = []
        
        for i, fp in enumerate(fingerprints):
            if fp is None:
                clusters.append(-1)  # Invalid molecules get their own cluster
                continue
            
            # Find best matching cluster
            best_cluster = -1
            best_similarity = 0
            
            for j, rep_fp in enumerate(cluster_representatives):
                if rep_fp is not None:
                    similarity = DataStructs.TanimotoSimilarity(fp, rep_fp)
                    if similarity > best_similarity and similarity >= self.similarity_threshold:
                        best_similarity = similarity
                        best_cluster = j
            
            if best_cluster == -1:
                # Create new cluster
                clusters.append(len(cluster_representatives))
                cluster_representatives.append(fp)
            else:
                clusters.append(best_cluster)
        
        return clusters
    
    def _allocate_clusters(self, 
                         clusters: List[int], 
                         total_size: int,
                         test_size: float, 
                         val_size: float,
                         random_state: int) -> Tuple[List[int], List[int], List[int]]:
        """Allocate clusters to train/val/test splits."""
        np.random.seed(random_state)
        
        # Group indices by cluster
        cluster_to_indices = {}
        for idx, cluster in enumerate(clusters):
            if cluster not in cluster_to_indices:
                cluster_to_indices[cluster] = []
            cluster_to_indices[cluster].append(idx)
        
        # Sort clusters by size and shuffle
        cluster_items = list(cluster_to_indices.items())
        np.random.shuffle(cluster_items)
        
        # Allocate clusters to splits
        target_test_size = int(total_size * test_size)
        target_val_size = int(total_size * val_size)
        
        train_indices = []
        val_indices = []
        test_indices = []
        
        for cluster_id, indices in cluster_items:
            current_test_size = len(test_indices)
            current_val_size = len(val_indices)
            
            if current_test_size < target_test_size:
                test_indices.extend(indices)
            elif current_val_size < target_val_size:
                val_indices.extend(indices)
            else:
                train_indices.extend(indices)
        
        return train_indices, val_indices, test_indices


def create_molecular_splits(df: pd.DataFrame,
                          smiles_column: str = 'smiles',
                          method: str = 'scaffold',
                          test_size: float = 0.2,
                          val_size: float = 0.1,
                          random_state: int = 42,
                          **kwargs) -> Tuple[pd.DataFrame, pd.DataFrame, pd.DataFrame]:
    """
    Create molecular-aware data splits.
    
    Args:
        df: DataFrame with molecular data
        smiles_column: Name of column containing SMILES strings
        method: Splitting method ('scaffold', 'temporal', 'cluster', 'random')
        test_size: Fraction of data for test set
        val_size: Fraction of data for validation set
        random_state: Random state for reproducibility
        **kwargs: Additional arguments for specific splitters
        
    Returns:
        Tuple of (train_df, val_df, test_df)
    """
    logger.info(f"Creating molecular splits using {method} method")
    
    if smiles_column not in df.columns:
        logger.warning(f"SMILES column '{smiles_column}' not found. Using random split.")
        method = 'random'
    
    if method == 'scaffold':
        splitter = MolecularScaffoldSplitter(**kwargs)
        split_result = splitter.split(df[smiles_column].tolist(), test_size, val_size, random_state)
    
    elif method == 'temporal':
        if 'timestamp_column' not in kwargs:
            raise ValueError("Temporal splitting requires 'timestamp_column' parameter")
        timestamp_col = kwargs['timestamp_column']
        splitter = TemporalSplitter()
        split_result = splitter.split(df[timestamp_col].tolist(), test_size, val_size)
    
    elif method == 'cluster':
        splitter = ClusterBasedSplitter(**kwargs)
        split_result = splitter.split(df[smiles_column].tolist(), test_size, val_size, random_state)
    
    elif method == 'random':
        # Standard random split
        indices = list(range(len(df)))
        train_val_indices, test_indices = train_test_split(
            indices, test_size=test_size, random_state=random_state
        )
        
        if val_size > 0:
            val_size_adjusted = val_size / (1 - test_size)
            train_indices, val_indices = train_test_split(
                train_val_indices, test_size=val_size_adjusted, random_state=random_state
            )
        else:
            train_indices = train_val_indices
            val_indices = []
        
        split_result = SplitResult(
            train_indices=train_indices,
            val_indices=val_indices,
            test_indices=test_indices,
            split_info={'method': 'random'}
        )
    
    else:
        raise ValueError(f"Unknown splitting method: {method}")
    
    # Create DataFrames
    train_df = df.iloc[split_result.train_indices].copy()
    val_df = df.iloc[split_result.val_indices].copy() if split_result.val_indices else pd.DataFrame()
    test_df = df.iloc[split_result.test_indices].copy()
    
    logger.info(f"Split completed: Train={len(train_df)}, Val={len(val_df)}, Test={len(test_df)}")
    
    return train_df, val_df, test_df
```
