## Supplementary material for "Machine Learning-Driven Prediction of TLR4 Binding Affinity: A Comprehensive Molecular Feature Analysis for Drug Discovery": Codebase: preprocessor.py

```
"""
Data preprocessing and integration utilities.

This module provides comprehensive data preprocessing functionality
for TLR4 binding prediction, including data loading, cleaning,
and integration of molecular features with binding affinities.
"""

from abc import ABC, abstractmethod
from typing import Dict, List, Optional, Union, Tuple, Any
import pandas as pd
import numpy as np
import logging
from pathlib import Path
try:
    from fuzzywuzzy import fuzz, process  # type: ignore
    _FUZZY_AVAILABLE = True
except Exception:
    # Graceful fallback using difflib
    import difflib

    _FUZZY_AVAILABLE = False

    class _FuzzCompat:
        @staticmethod
        def ratio(a: str, b: str) -> int:
            return int(round(difflib.SequenceMatcher(None, a, b).ratio() * 100))

        @staticmethod
        def partial_ratio(a: str, b: str) -> int:
            # Simple heuristic: compare shorter against best matching window in longer
            a = a or ""
            b = b or ""
            if len(a) == 0 or len(b) == 0:
                return 0
            if len(a) > len(b):
                a, b = b, a
            best = 0.0
            for i in range(0, len(b) - len(a) + 1):
                s = difflib.SequenceMatcher(None, a, b[i:i+len(a)]).ratio()
                if s > best:
                    best = s
            return int(round(best * 100))

    class _ProcessCompat:
        @staticmethod
        def extractOne(query: str, choices: list[str]):
            if not choices:
                return None
            scores = [(c, _FuzzCompat.ratio(query, c)) for c in choices]
            return max(scores, key=lambda x: x[1])

    fuzz = _FuzzCompat()  # type: ignore
    process = _ProcessCompat()  # type: ignore
import re

from ..molecular_analysis.features import MolecularFeatures, BindingData, FeatureSet

logger = logging.getLogger(__name__)


class DataPreprocessorInterface(ABC):
    """Abstract interface for data preprocessing operations."""
    
    @abstractmethod
    def load_binding_data(self, csv_path: str) -> pd.DataFrame:
        """Load and validate binding affinity data."""
        pass
    
    @abstractmethod
    def get_best_affinities(self, binding_df: pd.DataFrame) -> pd.DataFrame:
        """Extract best binding mode for each compound."""
        pass
    
    @abstractmethod
    def integrate_datasets(self, features_df: pd.DataFrame, 
                          binding_df: pd.DataFrame) -> pd.DataFrame:
        """Combine molecular features with binding data."""
        pass


class BindingDataLoader:
    """
    Loader for AutoDock Vina binding affinity data.
    
    Handles reading, validation, and basic preprocessing of binding
    affinity CSV files generated from AutoDock Vina results.
    """
    
    def __init__(self, affinity_column: str = 'affinity', 
                 ligand_column: str = 'ligand',
                 mode_column: str = 'mode'):
        """
        Initialize binding data loader.
        
        Args:
            affinity_column: Name of affinity column in CSV
            ligand_column: Name of ligand column in CSV
            mode_column: Name of mode column in CSV
        """
        self.affinity_column = affinity_column
        self.ligand_column = ligand_column
        self.mode_column = mode_column
        self.required_columns = [affinity_column, ligand_column, mode_column]
    
    def load_csv(self, csv_path: str) -> pd.DataFrame:
        """
        Load binding data from CSV file.
        
        Args:
            csv_path: Path to CSV file
            
        Returns:
            DataFrame with binding data
            
        Raises:
            FileNotFoundError: If file doesn't exist
            ValueError: If required columns are missing
        """
        if not Path(csv_path).exists():
            raise FileNotFoundError(f"Binding data file not found: {csv_path}")
        
        try:
            df = pd.read_csv(csv_path)
            
            # Validate required columns
            missing_cols = set(self.required_columns) - set(df.columns)
            if missing_cols:
                raise ValueError(f"Missing required columns: {missing_cols}")
            
            # Convert affinity to numeric
            df[self.affinity_column] = pd.to_numeric(df[self.affinity_column], errors='coerce')
            
            # Remove rows with missing affinity values
            initial_count = len(df)
            df = df.dropna(subset=[self.affinity_column])
            removed_count = initial_count - len(df)
            
            if removed_count > 0:
                logger.warning(f"Removed {removed_count} rows with missing affinity values")
            
            logger.info(f"Loaded {len(df)} binding records from {csv_path}")
            return df
            
        except Exception as e:
            logger.error(f"Error loading binding data from {csv_path}: {str(e)}")
            raise ValueError(f"Failed to load binding data: {str(e)}")
    
    def validate_binding_data(self, df: pd.DataFrame) -> Dict[str, List[str]]:
        """
        Validate binding data quality with comprehensive outlier detection.
        
        Args:
            df: DataFrame with binding data
            
        Returns:
            Dictionary with validation results and issues found
        """
        issues = {
            'missing_values': [],
            'invalid_affinities': [],
            'duplicate_entries': [],
            'outliers': [],
            'statistical_anomalies': []
        }
        
        # Check for missing values
        for col in self.required_columns:
            missing_count = df[col].isna().sum()
            if missing_count > 0:
                issues['missing_values'].append(f"{col}: {missing_count} missing values")
        
        # Check for invalid affinity values
        invalid_affinities = df[~np.isfinite(df[self.affinity_column])]
        if len(invalid_affinities) > 0:
            issues['invalid_affinities'].append(f"{len(invalid_affinities)} invalid affinity values")
        
        # Check for duplicate ligand-mode combinations
        duplicates = df.duplicated(subset=[self.ligand_column, self.mode_column])
        if duplicates.any():
            issues['duplicate_entries'].append(f"{duplicates.sum()} duplicate ligand-mode entries")
        
        # Enhanced outlier detection for binding affinities
        if len(df) > 0:
            outlier_info = self._detect_affinity_outliers(df)
            if outlier_info['outliers']:
                issues['outliers'].extend(outlier_info['outliers'])
            if outlier_info['statistical_anomalies']:
                issues['statistical_anomalies'].extend(outlier_info['statistical_anomalies'])
        
        return issues
    
    def _detect_affinity_outliers(self, df: pd.DataFrame) -> Dict[str, List[str]]:
        """
        Detect outliers in binding affinity data using multiple methods.
        
        Args:
            df: DataFrame with binding data
            
        Returns:
            Dictionary with outlier detection results
        """
        outliers = []
        statistical_anomalies = []
        affinity_col = self.affinity_column
        
        if len(df) < 4:  # Need at least 4 points for outlier detection
            return {'outliers': outliers, 'statistical_anomalies': statistical_anomalies}
        
        # Method 1: IQR-based outlier detection
        Q1 = df[affinity_col].quantile(0.25)
        Q3 = df[affinity_col].quantile(0.75)
        IQR = Q3 - Q1
        lower_bound = Q1 - 1.5 * IQR
        upper_bound = Q3 + 1.5 * IQR
        
        iqr_outliers = df[(df[affinity_col] < lower_bound) | 
                         (df[affinity_col] > upper_bound)]
        if len(iqr_outliers) > 0:
            outliers.append(f"IQR method: {len(iqr_outliers)} outliers detected (bounds: {lower_bound:.3f} to {upper_bound:.3f})")
        
        # Method 2: Z-score based outlier detection
        z_scores = np.abs((df[affinity_col] - df[affinity_col].mean()) / df[affinity_col].std())
        z_outliers = df[z_scores > 3]
        if len(z_outliers) > 0:
            outliers.append(f"Z-score method: {len(z_outliers)} outliers detected (|z| > 3)")
        
        # Method 3: Modified Z-score using median absolute deviation (MAD)
        median = df[affinity_col].median()
        mad = np.median(np.abs(df[affinity_col] - median))
        modified_z_scores = 0.6745 * (df[affinity_col] - median) / mad
        mad_outliers = df[np.abs(modified_z_scores) > 3.5]
        if len(mad_outliers) > 0:
            outliers.append(f"MAD method: {len(mad_outliers)} outliers detected")
        
        # Statistical anomaly detection
        affinity_stats = df[affinity_col].describe()
        
        # Check for unusual distribution characteristics
        if affinity_stats['std'] < 0.1:
            statistical_anomalies.append("Very low standard deviation - data may be too uniform")
        
        if affinity_stats['max'] - affinity_stats['min'] < 1.0:
            statistical_anomalies.append("Very narrow affinity range - limited binding diversity")
        
        # Check for binding affinity reasonableness (typical range: -15 to +5 kcal/mol)
        if affinity_stats['min'] > 5:
            statistical_anomalies.append("All affinities are positive - unusual for binding data")
        elif affinity_stats['max'] < -20:
            statistical_anomalies.append("Very low affinities detected - verify data quality")
        
        # Check for mode distribution anomalies
        mode_counts = df[self.mode_column].value_counts()
        if len(mode_counts) > 0 and mode_counts.max() / mode_counts.sum() > 0.8:
            statistical_anomalies.append("Highly skewed mode distribution - one mode dominates")
        
        return {'outliers': outliers, 'statistical_anomalies': statistical_anomalies}
    
    def clean_binding_data(self, df: pd.DataFrame, 
                          outlier_method: str = 'iqr',
                          remove_outliers: bool = False) -> pd.DataFrame:
        """
        Clean binding data by handling outliers and anomalies.
        
        Args:
            df: DataFrame with binding data
            outlier_method: Method for outlier detection ('iqr', 'zscore', 'mad')
            remove_outliers: Whether to remove outliers or cap them
            
        Returns:
            Cleaned DataFrame
        """
        logger.info(f"Cleaning binding data using {outlier_method} method")
        
        cleaned_df = df.copy()
        original_count = len(cleaned_df)
        
        if outlier_method == 'iqr':
            # IQR-based cleaning
            Q1 = cleaned_df[self.affinity_column].quantile(0.25)
            Q3 = cleaned_df[self.affinity_column].quantile(0.75)
            IQR = Q3 - Q1
            lower_bound = Q1 - 1.5 * IQR
            upper_bound = Q3 + 1.5 * IQR
            
            if remove_outliers:
                cleaned_df = cleaned_df[(cleaned_df[self.affinity_column] >= lower_bound) & 
                                      (cleaned_df[self.affinity_column] <= upper_bound)]
            else:
                # Cap outliers instead of removing
                cleaned_df[self.affinity_column] = cleaned_df[self.affinity_column].clip(
                    lower=lower_bound, upper=upper_bound
                )
        
        elif outlier_method == 'zscore':
            # Z-score based cleaning
            z_scores = np.abs((cleaned_df[self.affinity_column] - cleaned_df[self.affinity_column].mean()) / 
                             cleaned_df[self.affinity_column].std())
            
            if remove_outliers:
                cleaned_df = cleaned_df[z_scores <= 3]
            else:
                # Cap outliers at 3 standard deviations
                mean = cleaned_df[self.affinity_column].mean()
                std = cleaned_df[self.affinity_column].std()
                cleaned_df[self.affinity_column] = cleaned_df[self.affinity_column].clip(
                    lower=mean - 3*std, upper=mean + 3*std
                )
        
        elif outlier_method == 'mad':
            # MAD-based cleaning
            median = cleaned_df[self.affinity_column].median()
            mad = np.median(np.abs(cleaned_df[self.affinity_column] - median))
            modified_z_scores = 0.6745 * (cleaned_df[self.affinity_column] - median) / mad
            
            if remove_outliers:
                cleaned_df = cleaned_df[np.abs(modified_z_scores) <= 3.5]
            else:
                # Cap outliers at 3.5 MAD
                threshold = 3.5 * mad / 0.6745
                cleaned_df[self.affinity_column] = cleaned_df[self.affinity_column].clip(
                    lower=median - threshold, upper=median + threshold
                )
        
        removed_count = original_count - len(cleaned_df)
        if removed_count > 0:
            logger.info(f"Cleaning complete: {removed_count} records {'removed' if remove_outliers else 'capped'}")
        
        return cleaned_df


class CompoundMatcher:
    """
    Fuzzy string matching for compound name alignment.
    
    Handles matching compound names between different datasets
    using fuzzy string matching algorithms.
    """
    
    def __init__(self, threshold: float = 80.0, 
                 use_partial_ratio: bool = True):
        """
        Initialize compound matcher.
        
        Args:
            threshold: Minimum similarity score for matches (0-100)
            use_partial_ratio: Whether to use partial ratio for matching
        """
        self.threshold = threshold
        self.use_partial_ratio = use_partial_ratio
        self.match_cache = {}
    
    def match_compounds(self, names1: List[str], names2: List[str]) -> Dict[str, str]:
        """
        Match compound names between two lists.
        
        Args:
            names1: First list of compound names
            names2: Second list of compound names
            
        Returns:
            Dictionary mapping names1 to best matches in names2
        """
        matches = {}
        
        for name1 in names1:
            if name1 in self.match_cache:
                matches[name1] = self.match_cache[name1]
                continue
            
            best_match = self._find_best_match(name1, names2)
            if best_match:
                matches[name1] = best_match
                self.match_cache[name1] = best_match
            else:
                logger.warning(f"No match found for compound: {name1}")
        
        return matches
    
    def _find_best_match(self, target_name: str, candidate_names: List[str]) -> Optional[str]:
        """Find best match for a single compound name."""
        if not candidate_names:
            return None
        
        # Clean and normalize names
        target_clean = self._clean_name(target_name)
        
        best_match = None
        best_score = 0
        
        for candidate in candidate_names:
            candidate_clean = self._clean_name(candidate)
            
            # Calculate similarity scores
            if self.use_partial_ratio:
                score = fuzz.partial_ratio(target_clean, candidate_clean)
            else:
                score = fuzz.ratio(target_clean, candidate_clean)
            
            if score > best_score and score >= self.threshold:
                best_score = score
                best_match = candidate
        
        return best_match
    
    def _clean_name(self, name: str) -> str:
        """Clean and normalize compound name for matching."""
        if not isinstance(name, str):
            return str(name)
        
        # Convert to lowercase
        cleaned = name.lower().strip()
        
        # Remove common prefixes/suffixes
        prefixes = ['compound', 'ligand', 'molecule', 'drug']
        suffixes = ['_docked', '_bound', '_complex']
        
        for prefix in prefixes:
            if cleaned.startswith(prefix):
                cleaned = cleaned[len(prefix):].strip()
        
        for suffix in suffixes:
            if cleaned.endswith(suffix):
                cleaned = cleaned[:-len(suffix)].strip()
        
        # Remove special characters except alphanumeric and spaces
        cleaned = re.sub(r'[^a-zA-Z0-9\s]', '', cleaned)
        
        # Remove extra spaces
        cleaned = ' '.join(cleaned.split())
        
        return cleaned
    
    def get_match_confidence(self, name1: str, name2: str) -> float:
        """Get confidence score for a specific name pair."""
        clean1 = self._clean_name(name1)
        clean2 = self._clean_name(name2)
        
        if self.use_partial_ratio:
            return fuzz.partial_ratio(clean1, clean2)
        else:
            return fuzz.ratio(clean1, clean2)


class DataIntegrator:
    """
    Integrates molecular features with binding affinity data.
    
    Combines datasets from different sources and handles missing
    data, outliers, and data quality issues.
    """
    
    def __init__(self, compound_matcher: Optional[CompoundMatcher] = None):
        """
        Initialize data integrator.
        
        Args:
            compound_matcher: Optional compound matcher for name alignment
        """
        self.compound_matcher = compound_matcher or CompoundMatcher()
        self.integration_stats = {}
    
    def integrate_datasets(self, features_df: pd.DataFrame, 
                          binding_df: pd.DataFrame,
                          feature_compound_col: str = 'compound_name',
                          binding_compound_col: str = 'ligand') -> pd.DataFrame:
        """
        Integrate molecular features with binding data.
        
        Args:
            features_df: DataFrame with molecular features
            binding_df: DataFrame with binding affinities
            feature_compound_col: Column name for compound names in features
            binding_compound_col: Column name for compound names in binding data
            
        Returns:
            Integrated DataFrame with features and binding data
        """
        logger.info("Starting dataset integration")
        
        # Get compound names from both datasets
        feature_compounds = features_df[feature_compound_col].unique().tolist()
        binding_compounds = binding_df[binding_compound_col].unique().tolist()
        
        # Match compound names
        matches = self.compound_matcher.match_compounds(feature_compounds, binding_compounds)
        
        # Create mapping for binding data
        binding_mapping = {v: k for k, v in matches.items()}
        
        # Add matched compound names to binding data
        binding_df_matched = binding_df.copy()
        binding_df_matched['matched_compound'] = binding_df_matched[binding_compound_col].map(binding_mapping)
        
        # Merge datasets
        integrated_df = pd.merge(
            features_df,
            binding_df_matched,
            left_on=feature_compound_col,
            right_on='matched_compound',
            how='inner'
        )
        
        # Calculate integration statistics
        self.integration_stats = {
            'total_features': len(features_df),
            'total_binding': len(binding_df),
            'successful_matches': len(matches),
            'integrated_records': len(integrated_df),
            'match_rate': len(matches) / len(feature_compounds) if feature_compounds else 0
        }
        
        logger.info(f"Integration complete: {self.integration_stats['integrated_records']} records")
        logger.info(f"Match rate: {self.integration_stats['match_rate']:.2%}")
        
        return integrated_df
    
    def get_integration_stats(self) -> Dict[str, Any]:
        """Get integration statistics."""
        return self.integration_stats.copy()


class DataPreprocessor(DataPreprocessorInterface):
    """
    Main data preprocessing coordinator.
    
    Orchestrates the complete data preprocessing pipeline including
    loading, cleaning, integration, and validation.
    """
    
    def __init__(self, binding_loader: Optional[BindingDataLoader] = None,
                 compound_matcher: Optional[CompoundMatcher] = None,
                 data_integrator: Optional[DataIntegrator] = None):
        """
        Initialize data preprocessor.
        
        Args:
            binding_loader: Optional binding data loader
            compound_matcher: Optional compound matcher
            data_integrator: Optional data integrator
        """
        self.binding_loader = binding_loader or BindingDataLoader()
        self.compound_matcher = compound_matcher or CompoundMatcher()
        self.data_integrator = data_integrator or DataIntegrator(self.compound_matcher)
    
    def load_binding_data(self, csv_path: str) -> pd.DataFrame:
        """Load and validate binding affinity data."""
        return self.binding_loader.load_csv(csv_path)
    
    def get_best_affinities(self, binding_df: pd.DataFrame) -> pd.DataFrame:
        """
        Extract best binding mode (lowest/most negative affinity) per compound for strongest TLR4 binding.
        
        This method identifies the strongest binding interaction for each compound by selecting
        the mode with the lowest (most negative) affinity value, which represents the strongest
        binding interaction in AutoDock Vina scoring.
        
        Args:
            binding_df: DataFrame with binding data
            
        Returns:
            DataFrame with best binding mode per compound
            
        Raises:
            ValueError: If affinity values are not properly formatted or contain invalid data
        """
        logger.info("Extracting best binding affinities (strongest TLR4 binding)")
        
        # Validate affinity data before processing
        self._validate_affinity_data(binding_df)
        
        # Group by ligand and find minimum affinity (strongest binding)
        # Lower (more negative) affinity values indicate stronger binding
        best_affinities = binding_df.groupby(self.binding_loader.ligand_column).agg({
            self.binding_loader.affinity_column: 'min',  # Most negative = strongest binding
            self.binding_loader.mode_column: 'first'     # Take first mode with min affinity
        }).reset_index()
        
        # Add additional information from original data
        best_affinities = best_affinities.merge(
            binding_df,
            on=[self.binding_loader.ligand_column, self.binding_loader.affinity_column],
            how='left'
        )
        
        # Remove duplicates
        best_affinities = best_affinities.drop_duplicates(
            subset=[self.binding_loader.ligand_column]
        )
        
        # Validate that we have the strongest binding modes
        self._validate_best_affinities(best_affinities, binding_df)
        
        logger.info(f"Extracted {len(best_affinities)} best binding modes")
        logger.info(f"Affinity range: {best_affinities[self.binding_loader.affinity_column].min():.3f} to {best_affinities[self.binding_loader.affinity_column].max():.3f} kcal/mol")
        
        return best_affinities
    
    def _validate_affinity_data(self, binding_df: pd.DataFrame) -> None:
        """
        Validate that affinity data represents binding strength correctly.
        
        Args:
            binding_df: DataFrame with binding data
            
        Raises:
            ValueError: If affinity data is invalid or doesn't represent binding strength
        """
        affinity_col = self.binding_loader.affinity_column
        
        # Check for non-numeric values
        if not pd.api.types.is_numeric_dtype(binding_df[affinity_col]):
            raise ValueError(f"Affinity column '{affinity_col}' must contain numeric values")
        
        # Check for infinite values
        if not np.isfinite(binding_df[affinity_col]).all():
            infinite_count = (~np.isfinite(binding_df[affinity_col])).sum()
            raise ValueError(f"Found {infinite_count} non-finite affinity values")
        
        # Validate that lower values represent stronger binding
        # In AutoDock Vina, more negative values indicate stronger binding
        affinity_stats = binding_df[affinity_col].describe()
        logger.info(f"Affinity statistics: min={affinity_stats['min']:.3f}, max={affinity_stats['max']:.3f}, mean={affinity_stats['mean']:.3f}")
        
        # Check if we have reasonable binding affinity range (typically -15 to +5 kcal/mol)
        if affinity_stats['min'] > 0:
            logger.warning("All affinity values are positive - this may indicate data processing issues")
        elif affinity_stats['max'] < -20:
            logger.warning("Very low affinity values detected - verify data quality")
    
    def _validate_best_affinities(self, best_affinities: pd.DataFrame, original_df: pd.DataFrame) -> None:
        """
        Validate that best affinities are correctly extracted.
        
        Args:
            best_affinities: DataFrame with best binding modes
            original_df: Original binding data DataFrame
        """
        affinity_col = self.binding_loader.affinity_column
        ligand_col = self.binding_loader.ligand_column
        
        # Verify that best affinities are indeed the minimum for each ligand
        for _, row in best_affinities.iterrows():
            ligand = row[ligand_col]
            best_affinity = row[affinity_col]
            
            # Get all affinities for this ligand
            ligand_affinities = original_df[original_df[ligand_col] == ligand][affinity_col]
            
            # Verify this is indeed the minimum (strongest binding)
            if not np.isclose(best_affinity, ligand_affinities.min(), rtol=1e-6):
                logger.warning(f"Best affinity for {ligand} may not be correctly extracted")
        
        logger.info("Best affinity validation completed successfully")
    
    def integrate_datasets(self, features_df: pd.DataFrame, 
                          binding_df: pd.DataFrame) -> pd.DataFrame:
        """Combine molecular features with binding data."""
        return self.data_integrator.integrate_datasets(features_df, binding_df)
    
    def preprocess_pipeline(self, features_df: pd.DataFrame, 
                           binding_csv_path: str) -> pd.DataFrame:
        """
        Complete preprocessing pipeline.
        
        Args:
            features_df: DataFrame with molecular features
            binding_csv_path: Path to binding data CSV
            
        Returns:
            Preprocessed and integrated DataFrame
        """
        logger.info("Starting complete preprocessing pipeline")
        
        # Load binding data
        binding_df = self.load_binding_data(binding_csv_path)
        
        # Get best affinities
        best_affinities = self.get_best_affinities(binding_df)
        
        # Integrate datasets
        integrated_df = self.integrate_datasets(features_df, best_affinities)
        
        # Validate integrated data
        validation_results = self._validate_integrated_data(integrated_df)
        
        if validation_results['issues']:
            logger.warning(f"Data validation issues found: {validation_results['issues']}")
        
        logger.info("Preprocessing pipeline completed successfully")
        return integrated_df
    
    def _validate_integrated_data(self, df: pd.DataFrame) -> Dict[str, Any]:
        """Validate integrated dataset."""
        issues = []
        
        # Check for empty dataset
        if len(df) == 0:
            issues.append("Empty integrated dataset")
        
        # Check for missing target values
        if self.binding_loader.affinity_column in df.columns:
            missing_targets = df[self.binding_loader.affinity_column].isna().sum()
            if missing_targets > 0:
                issues.append(f"{missing_targets} missing target values")
        
        # Check for duplicate compounds
        if 'compound_name' in df.columns:
            duplicates = df['compound_name'].duplicated().sum()
            if duplicates > 0:
                issues.append(f"{duplicates} duplicate compounds")
        
        return {
            'valid': len(issues) == 0,
            'issues': issues,
            'record_count': len(df)
        }
```
