## Supplementary material for "Machine Learning-Driven Prediction of TLR4 Binding Affinity: A Comprehensive Molecular Feature Analysis for Drug Discovery": Codebase: validation.py

```
"""
Data validation and quality assessment utilities.

This module provides comprehensive data validation functionality
for molecular features and binding data quality assessment.
"""

from abc import ABC, abstractmethod
from typing import Dict, List, Optional, Union, Tuple, Any
import pandas as pd
import numpy as np
import logging
from sklearn.ensemble import IsolationForest
from sklearn.preprocessing import StandardScaler
from scipy import stats

logger = logging.getLogger(__name__)


class DataValidatorInterface(ABC):
    """Abstract interface for data validation."""
    
    @abstractmethod
    def validate(self, data: pd.DataFrame) -> Dict[str, Any]:
        """Validate data quality."""
        pass
    
    @abstractmethod
    def get_validation_report(self) -> Dict[str, Any]:
        """Get comprehensive validation report."""
        pass


class DataValidator(DataValidatorInterface):
    """Comprehensive data validation for molecular features and binding data."""
    
    def __init__(self, strict_validation: bool = True):
        """
        Initialize data validator.
        
        Args:
            strict_validation: If True, enforce strict validation rules
        """
        self.strict_validation = strict_validation
        self.validation_results = {}
        self.validation_history = []
    
    def validate(self, data: pd.DataFrame) -> Dict[str, Any]:
        """
        Validate data quality comprehensively.
        
        Args:
            data: DataFrame to validate
            
        Returns:
            Dictionary with validation results
        """
        logger.info(f"Validating data with {len(data)} records and {len(data.columns)} features")
        
        validation_results = {
            'basic_stats': self._validate_basic_stats(data),
            'missing_values': self._validate_missing_values(data),
            'data_types': self._validate_data_types(data),
            'outliers': self._validate_outliers(data),
            'correlations': self._validate_correlations(data),
            'distributions': self._validate_distributions(data),
            'consistency': self._validate_consistency(data)
        }
        
        # Calculate overall validation score
        validation_results['overall_score'] = self._calculate_overall_score(validation_results)
        validation_results['is_valid'] = validation_results['overall_score'] >= 0.8
        
        # Store results
        self.validation_results = validation_results
        self.validation_history.append({
            'timestamp': pd.Timestamp.now(),
            'data_shape': data.shape,
            'results': validation_results
        })
        
        logger.info(f"Validation completed. Overall score: {validation_results['overall_score']:.2f}")
        
        return validation_results
    
    def _validate_basic_stats(self, data: pd.DataFrame) -> Dict[str, Any]:
        """Validate basic data statistics."""
        return {
            'record_count': len(data),
            'feature_count': len(data.columns),
            'memory_usage': data.memory_usage(deep=True).sum(),
            'duplicate_rows': data.duplicated().sum(),
            'empty_rows': data.isnull().all(axis=1).sum()
        }
    
    def _validate_missing_values(self, data: pd.DataFrame) -> Dict[str, Any]:
        """Validate missing value patterns."""
        missing_stats = data.isnull().sum()
        missing_percentages = (missing_stats / len(data)) * 100
        
        return {
            'total_missing': missing_stats.sum(),
            'missing_percentage': (missing_stats.sum() / (len(data) * len(data.columns))) * 100,
            'columns_with_missing': missing_stats[missing_stats > 0].to_dict(),
            'missing_percentages': missing_percentages[missing_percentages > 0].to_dict(),
            'high_missing_columns': missing_percentages[missing_percentages > 50].to_dict()
        }
    
    def _validate_data_types(self, data: pd.DataFrame) -> Dict[str, Any]:
        """Validate data types and consistency."""
        type_counts = data.dtypes.value_counts()
        numeric_columns = data.select_dtypes(include=[np.number]).columns.tolist()
        categorical_columns = data.select_dtypes(include=['object', 'category']).columns.tolist()
        
        return {
            'type_distribution': type_counts.to_dict(),
            'numeric_columns': numeric_columns,
            'categorical_columns': categorical_columns,
            'mixed_type_columns': self._find_mixed_type_columns(data)
        }
    
    def _validate_outliers(self, data: pd.DataFrame) -> Dict[str, Any]:
        """Validate outlier patterns."""
        numeric_data = data.select_dtypes(include=[np.number])
        
        if numeric_data.empty:
            return {'outlier_analysis': 'No numeric columns for outlier analysis'}
        
        outlier_results = {}
        
        # IQR method
        iqr_outliers = self._detect_iqr_outliers(numeric_data)
        outlier_results['iqr_outliers'] = iqr_outliers
        
        # Z-score method
        zscore_outliers = self._detect_zscore_outliers(numeric_data)
        outlier_results['zscore_outliers'] = zscore_outliers
        
        # Isolation Forest method
        isolation_outliers = self._detect_isolation_forest_outliers(numeric_data)
        outlier_results['isolation_forest_outliers'] = isolation_outliers
        
        return outlier_results
    
    def _validate_correlations(self, data: pd.DataFrame) -> Dict[str, Any]:
        """Validate correlation patterns."""
        numeric_data = data.select_dtypes(include=[np.number])
        
        if numeric_data.empty:
            return {'correlation_analysis': 'No numeric columns for correlation analysis'}
        
        corr_matrix = numeric_data.corr()
        
        # Find highly correlated pairs
        high_corr_pairs = []
        for i in range(len(corr_matrix.columns)):
            for j in range(i+1, len(corr_matrix.columns)):
                corr_val = corr_matrix.iloc[i, j]
                if abs(corr_val) > 0.95:
                    high_corr_pairs.append({
                        'feature_1': corr_matrix.columns[i],
                        'feature_2': corr_matrix.columns[j],
                        'correlation': corr_val
                    })
        
        return {
            'correlation_matrix': corr_matrix,
            'high_correlation_pairs': high_corr_pairs,
            'max_correlation': corr_matrix.abs().max().max(),
            'mean_correlation': corr_matrix.abs().mean().mean()
        }
    
    def _validate_distributions(self, data: pd.DataFrame) -> Dict[str, Any]:
        """Validate data distributions."""
        numeric_data = data.select_dtypes(include=[np.number])
        
        if numeric_data.empty:
            return {'distribution_analysis': 'No numeric columns for distribution analysis'}
        
        distribution_results = {}
        
        for column in numeric_data.columns:
            values = numeric_data[column].dropna()
            if len(values) > 0:
                # Shapiro-Wilk test for normality
                if len(values) <= 5000:  # Shapiro-Wilk works best with smaller samples
                    shapiro_stat, shapiro_p = stats.shapiro(values)
                    distribution_results[column] = {
                        'shapiro_statistic': shapiro_stat,
                        'shapiro_p_value': shapiro_p,
                        'is_normal': shapiro_p > 0.05,
                        'skewness': stats.skew(values),
                        'kurtosis': stats.kurtosis(values)
                    }
                else:
                    # For large samples, use Kolmogorov-Smirnov test
                    ks_stat, ks_p = stats.kstest(values, 'norm')
                    distribution_results[column] = {
                        'ks_statistic': ks_stat,
                        'ks_p_value': ks_p,
                        'is_normal': ks_p > 0.05,
                        'skewness': stats.skew(values),
                        'kurtosis': stats.kurtosis(values)
                    }
        
        return distribution_results
    
    def _validate_consistency(self, data: pd.DataFrame) -> Dict[str, Any]:
        """Validate data consistency and business rules."""
        consistency_issues = []
        
        # Check for negative values in positive-only columns
        positive_columns = ['molecular_weight', 'tpsa', 'surface_area', 'molecular_volume']
        for col in positive_columns:
            if col in data.columns:
                negative_count = (data[col] < 0).sum()
                if negative_count > 0:
                    consistency_issues.append(f"{col}: {negative_count} negative values")
        
        # Check for unrealistic values
        if 'molecular_weight' in data.columns:
            unrealistic_mw = ((data['molecular_weight'] < 10) | (data['molecular_weight'] > 2000)).sum()
            if unrealistic_mw > 0:
                consistency_issues.append(f"molecular_weight: {unrealistic_mw} unrealistic values")
        
        # Check for missing compound names
        if 'compound_name' in data.columns:
            missing_names = data['compound_name'].isnull().sum()
            if missing_names > 0:
                consistency_issues.append(f"compound_name: {missing_names} missing values")
        
        return {
            'consistency_issues': consistency_issues,
            'issue_count': len(consistency_issues)
        }
    
    def _find_mixed_type_columns(self, data: pd.DataFrame) -> List[str]:
        """Find columns with mixed data types."""
        mixed_columns = []
        
        for column in data.columns:
            if data[column].dtype == 'object':
                # Check if column contains mixed types
                non_null_values = data[column].dropna()
                if len(non_null_values) > 0:
                    type_counts = non_null_values.apply(type).value_counts()
                    if len(type_counts) > 1:
                        mixed_columns.append(column)
        
        return mixed_columns
    
    def _detect_iqr_outliers(self, data: pd.DataFrame) -> Dict[str, int]:
        """Detect outliers using IQR method."""
        outlier_counts = {}
        
        for column in data.columns:
            Q1 = data[column].quantile(0.25)
            Q3 = data[column].quantile(0.75)
            IQR = Q3 - Q1
            lower_bound = Q1 - 1.5 * IQR
            upper_bound = Q3 + 1.5 * IQR
            outliers = ((data[column] < lower_bound) | (data[column] > upper_bound)).sum()
            outlier_counts[column] = int(outliers)
        
        return outlier_counts
    
    def _detect_zscore_outliers(self, data: pd.DataFrame, threshold: float = 3.0) -> Dict[str, int]:
        """Detect outliers using Z-score method."""
        outlier_counts = {}
        
        for column in data.columns:
            z_scores = np.abs(stats.zscore(data[column].dropna()))
            outliers = (z_scores > threshold).sum()
            outlier_counts[column] = int(outliers)
        
        return outlier_counts
    
    def _detect_isolation_forest_outliers(self, data: pd.DataFrame) -> Dict[str, int]:
        """Detect outliers using Isolation Forest."""
        try:
            # Fill missing values for Isolation Forest
            data_filled = data.fillna(data.median())
            
            # Fit Isolation Forest
            iso_forest = IsolationForest(contamination=0.1, random_state=42)
            outlier_labels = iso_forest.fit_predict(data_filled)
            
            # Count outliers per column
            outlier_counts = {}
            for i, column in enumerate(data.columns):
                column_outliers = (outlier_labels == -1).sum()
                outlier_counts[column] = int(column_outliers)
            
            return outlier_counts
            
        except Exception as e:
            logger.warning(f"Isolation Forest outlier detection failed: {str(e)}")
            return {column: 0 for column in data.columns}
    
    def _calculate_overall_score(self, validation_results: Dict[str, Any]) -> float:
        """Calculate overall validation score."""
        scores = []
        
        # Missing values score
        missing_pct = validation_results['missing_values']['missing_percentage']
        missing_score = max(0, 1 - missing_pct / 100)
        scores.append(missing_score)
        
        # Outlier score
        outlier_results = validation_results['outliers']
        if 'iqr_outliers' in outlier_results:
            total_outliers = sum(outlier_results['iqr_outliers'].values())
            outlier_score = max(0, 1 - total_outliers / (len(validation_results['basic_stats']['record_count']) * 10))
            scores.append(outlier_score)
        
        # Consistency score
        consistency_issues = validation_results['consistency']['issue_count']
        consistency_score = max(0, 1 - consistency_issues / 10)
        scores.append(consistency_score)
        
        # Correlation score
        if 'high_correlation_pairs' in validation_results['correlations']:
            high_corr_count = len(validation_results['correlations']['high_correlation_pairs'])
            correlation_score = max(0, 1 - high_corr_count / 50)
            scores.append(correlation_score)
        
        return np.mean(scores) if scores else 0.0
    
    def get_validation_report(self) -> Dict[str, Any]:
        """Get comprehensive validation report."""
        if not self.validation_results:
            return {'error': 'No validation results available'}
        
        return {
            'validation_summary': {
                'overall_score': self.validation_results['overall_score'],
                'is_valid': self.validation_results['is_valid'],
                'record_count': self.validation_results['basic_stats']['record_count'],
                'feature_count': self.validation_results['basic_stats']['feature_count']
            },
            'detailed_results': self.validation_results,
            'validation_history': self.validation_history
        }


class OutlierDetector:
    """Specialized outlier detection for molecular data."""
    
    def __init__(self, method: str = 'iqr', threshold: float = 1.5):
        """
        Initialize outlier detector.
        
        Args:
            method: Detection method ('iqr', 'zscore', 'isolation_forest')
            threshold: Threshold for outlier detection
        """
        self.method = method
        self.threshold = threshold
        self.outlier_mask = None
    
    def detect_outliers(self, data: pd.DataFrame) -> pd.DataFrame:
        """
        Detect outliers in molecular data.
        
        Args:
            data: DataFrame with molecular features
            
        Returns:
            DataFrame with outlier flags
        """
        if self.method == 'iqr':
            return self._detect_iqr_outliers(data)
        elif self.method == 'zscore':
            return self._detect_zscore_outliers(data)
        elif self.method == 'isolation_forest':
            return self._detect_isolation_forest_outliers(data)
        else:
            raise ValueError(f"Unknown outlier detection method: {self.method}")
    
    def _detect_iqr_outliers(self, data: pd.DataFrame) -> pd.DataFrame:
        """Detect outliers using IQR method."""
        outlier_flags = pd.DataFrame(index=data.index, columns=data.columns, dtype=bool)
        
        for column in data.select_dtypes(include=[np.number]).columns:
            Q1 = data[column].quantile(0.25)
            Q3 = data[column].quantile(0.75)
            IQR = Q3 - Q1
            lower_bound = Q1 - self.threshold * IQR
            upper_bound = Q3 + self.threshold * IQR
            outlier_flags[column] = (data[column] < lower_bound) | (data[column] > upper_bound)
        
        return outlier_flags
    
    def _detect_zscore_outliers(self, data: pd.DataFrame) -> pd.DataFrame:
        """Detect outliers using Z-score method."""
        outlier_flags = pd.DataFrame(index=data.index, columns=data.columns, dtype=bool)
        
        for column in data.select_dtypes(include=[np.number]).columns:
            z_scores = np.abs(stats.zscore(data[column].dropna()))
            outlier_flags[column] = z_scores > self.threshold
        
        return outlier_flags
    
    def _detect_isolation_forest_outliers(self, data: pd.DataFrame) -> pd.DataFrame:
        """Detect outliers using Isolation Forest."""
        try:
            # Fill missing values
            data_filled = data.select_dtypes(include=[np.number]).fillna(data.median())
            
            # Fit Isolation Forest
            iso_forest = IsolationForest(contamination=0.1, random_state=42)
            outlier_labels = iso_forest.fit_predict(data_filled)
            
            # Create outlier flags
            outlier_flags = pd.DataFrame(index=data.index, columns=data.columns, dtype=bool)
            outlier_flags.loc[data_filled.index, data_filled.columns] = outlier_labels == -1
            
            return outlier_flags
            
        except Exception as e:
            logger.warning(f"Isolation Forest outlier detection failed: {str(e)}")
            return pd.DataFrame(index=data.index, columns=data.columns, dtype=bool)
```
