## Supplementary material for "Machine Learning-Driven Prediction of TLR4 Binding Affinity: A Comprehensive Molecular Feature Analysis for Drug Discovery": Codebase: __init__.py

```
"""
Machine Learning Components Module

Contains classes for model training, evaluation, and prediction
for TLR4 binding affinity prediction using various ML algorithms.
"""

from .trainer import MLModelTrainer, ModelTrainerInterface
from .evaluator import ModelEvaluator, PerformanceMetrics
from .predictor import BindingPredictor, UncertaintyEstimatorInterface
from .data_splitting import (
    DataSplitter, 
    CrossValidationSetup, 
    DataValidationFramework,
    DataQualityReporter,
    DataSplitConfig,
    CrossValidationConfig
)
from .ablation_study import (
    AblationStudyFramework,
    FeatureAblationStudy,
    DataSizeAblationStudy,
    HyperparameterAblationStudy,
    ArchitectureAblationStudy,
    AblationConfig,
    AblationResult
)
from .uncertainty_quantification import (
    MonteCarloDropout,
    BootstrapUncertainty,
    ConformalPrediction,
    EnsembleUncertainty,
    UncertaintyCalibration,
    UncertaintyQuantifier,
    UncertaintyResult
)
from .compound_analysis import CompoundAnalysis

__all__ = [
    "MLModelTrainer",
    "ModelTrainerInterface",
    "ModelEvaluator",
    "PerformanceMetrics",
    "BindingPredictor",
    "UncertaintyEstimatorInterface",
    "DataSplitter",
    "CrossValidationSetup",
    "DataValidationFramework",
    "DataQualityReporter",
    "DataSplitConfig",
    "CrossValidationConfig",
    "AblationStudyFramework",
    "FeatureAblationStudy",
    "DataSizeAblationStudy",
    "HyperparameterAblationStudy",
    "ArchitectureAblationStudy",
    "AblationConfig",
    "AblationResult",
    "MonteCarloDropout",
    "BootstrapUncertainty",
    "ConformalPrediction",
    "EnsembleUncertainty",
    "UncertaintyCalibration",
    "UncertaintyQuantifier",
    "UncertaintyResult",
    
    "CompoundAnalysis"
]
```
