## Supplementary material for "Machine Learning-Driven Prediction of TLR4 Binding Affinity: A Comprehensive Molecular Feature Analysis for Drug Discovery": Codebase: ablation_study.py

```
"""
Ablation Study Framework for TLR4 Binding Prediction

This module implements comprehensive ablation studies to analyze the contribution
of different components to model performance, including:
- Feature ablation studies
- Model architecture ablation
- Data size ablation
- Hyperparameter sensitivity analysis
"""

import numpy as np
import pandas as pd
import matplotlib.pyplot as plt
import seaborn as sns
from typing import Dict, List, Tuple, Any, Optional, Union
from sklearn.model_selection import cross_val_score, KFold
from sklearn.metrics import r2_score, mean_squared_error, mean_absolute_error
from scipy import stats
import warnings
from pathlib import Path
import json
import logging
from dataclasses import dataclass, asdict
from concurrent.futures import ThreadPoolExecutor, ProcessPoolExecutor
import time

from ..utils.logger import get_logger
from .evaluator import ModelEvaluator
from .trainer import MLModelTrainer
from .feature_engineering import FeatureEngineeringPipeline

logger = get_logger(__name__)

@dataclass
class AblationResult:
    """Container for ablation study results"""
    component_name: str
    baseline_score: float
    ablated_score: float
    score_difference: float
    relative_change: float
    p_value: Optional[float] = None
    confidence_interval: Optional[Tuple[float, float]] = None
    metadata: Optional[Dict[str, Any]] = None

@dataclass
class AblationConfig:
    """Configuration for ablation studies"""
    cv_folds: int = 5
    n_iterations: int = 10
    random_state: int = 42
    confidence_level: float = 0.95
    min_effect_size: float = 0.01
    parallel_jobs: int = -1
    save_models: bool = False
    verbose: bool = True

class FeatureAblationStudy:
    """Feature ablation study to identify critical molecular descriptors"""
    
    def __init__(self, config: AblationConfig = None):
        self.config = config or AblationConfig()
        self.evaluator = ModelEvaluator()
        self.results: List[AblationResult] = []
        
    def run_feature_ablation(self, 
                           X: pd.DataFrame, 
                           y: pd.Series, 
                           model_trainer: MLModelTrainer,
                           feature_groups: Optional[Dict[str, List[str]]] = None) -> List[AblationResult]:
        """
        Run feature ablation study by removing groups of features
        
        Args:
            X: Feature matrix
            y: Target values
            model_trainer: Trained model for evaluation
            feature_groups: Optional grouping of features for ablation
                           If None, ablates individual features
        """
        logger.info("Starting feature ablation study")
        
        # Get baseline performance
        baseline_scores = self._get_baseline_performance(X, y, model_trainer)
        
        if feature_groups is None:
            # Ablate individual features
            feature_groups = {f"feature_{col}": [col] for col in X.columns}
        
        results = []
        for group_name, features in feature_groups.items():
            logger.info(f"Ablating feature group: {group_name}")
            
            # Remove features
            X_ablated = X.drop(columns=features)
            
            if X_ablated.empty:
                logger.warning(f"Skipping {group_name} - would remove all features")
                continue
                
            # Get ablated performance
            ablated_scores = self._get_ablated_performance(X_ablated, y, model_trainer)
            
            # Calculate statistics
            result = self._calculate_ablation_stats(
                group_name, baseline_scores, ablated_scores, 
                {"removed_features": features, "n_features": len(features)}
            )
            results.append(result)
            
        self.results.extend(results)
        return results
    
    def run_sequential_feature_ablation(self, 
                                      X: pd.DataFrame, 
                                      y: pd.Series, 
                                      model_trainer: MLModelTrainer,
                                      method: str = "forward") -> List[AblationResult]:
        """
        Run sequential feature ablation (forward/backward selection)
        
        Args:
            X: Feature matrix
            y: Target values
            model_trainer: Trained model for evaluation
            method: "forward" or "backward"
        """
        logger.info(f"Starting sequential feature ablation ({method})")
        
        if method == "forward":
            return self._forward_feature_selection(X, y, model_trainer)
        else:
            return self._backward_feature_elimination(X, y, model_trainer)
    
    def _forward_feature_selection(self, X: pd.DataFrame, y: pd.Series, 
                                 model_trainer: MLModelTrainer) -> List[AblationResult]:
        """Forward feature selection"""
        selected_features = []
        remaining_features = list(X.columns)
        results = []
        
        while remaining_features:
            best_score = -np.inf
            best_feature = None
            
            for feature in remaining_features:
                candidate_features = selected_features + [feature]
                X_candidate = X[candidate_features]
                scores = self._get_baseline_performance(X_candidate, y, model_trainer)
                avg_score = np.mean(scores)
                
                if avg_score > best_score:
                    best_score = avg_score
                    best_feature = feature
            
            if best_feature:
                selected_features.append(best_feature)
                remaining_features.remove(best_feature)
                
                result = AblationResult(
                    component_name=f"forward_step_{len(selected_features)}",
                    baseline_score=best_score,
                    ablated_score=0.0,
                    score_difference=best_score,
                    relative_change=0.0,
                    metadata={"selected_features": selected_features.copy()}
                )
                results.append(result)
                
                logger.info(f"Added feature {best_feature}, score: {best_score:.4f}")
        
        return results
    
    def _backward_feature_elimination(self, X: pd.DataFrame, y: pd.Series,
                                    model_trainer: MLModelTrainer) -> List[AblationResult]:
        """Backward feature elimination"""
        remaining_features = list(X.columns)
        results = []
        baseline_scores = self._get_baseline_performance(X, y, model_trainer)
        
        while len(remaining_features) > 1:
            worst_score = np.inf
            worst_feature = None
            
            for feature in remaining_features:
                candidate_features = [f for f in remaining_features if f != feature]
                X_candidate = X[candidate_features]
                scores = self._get_ablated_performance(X_candidate, y, model_trainer)
                avg_score = np.mean(scores)
                
                if avg_score < worst_score:
                    worst_score = avg_score
                    worst_feature = feature
            
            if worst_feature:
                remaining_features.remove(worst_feature)
                
                result = self._calculate_ablation_stats(
                    f"backward_step_{len(remaining_features)}",
                    baseline_scores, 
                    self._get_ablated_performance(X[remaining_features], y, model_trainer),
                    {"remaining_features": remaining_features.copy()}
                )
                results.append(result)
                
                logger.info(f"Removed feature {worst_feature}, remaining: {len(remaining_features)}")
        
        return results
    
    def _get_baseline_performance(self, X: pd.DataFrame, y: pd.Series,
                                model_trainer: MLModelTrainer) -> List[float]:
        """Get baseline model performance using cross-validation"""
        kf = KFold(n_splits=self.config.cv_folds, shuffle=True, random_state=self.config.random_state)
        
        scores = []
        for train_idx, val_idx in kf.split(X):
            X_train, X_val = X.iloc[train_idx], X.iloc[val_idx]
            y_train, y_val = y.iloc[train_idx], y.iloc[val_idx]
            
            # Train model
            model = model_trainer.train_single_model(X_train, y_train)
            
            # Evaluate
            y_pred = model.predict(X_val)
            score = r2_score(y_val, y_pred)
            scores.append(score)
        
        return scores
    
    def _get_ablated_performance(self, X: pd.DataFrame, y: pd.Series,
                               model_trainer: MLModelTrainer) -> List[float]:
        """Get ablated model performance using cross-validation"""
        return self._get_baseline_performance(X, y, model_trainer)
    
    def _calculate_ablation_stats(self, component_name: str, baseline_scores: List[float],
                                ablated_scores: List[float], metadata: Dict[str, Any]) -> AblationResult:
        """Calculate statistical measures for ablation study"""
        baseline_mean = np.mean(baseline_scores)
        ablated_mean = np.mean(ablated_scores)
        
        score_diff = baseline_mean - ablated_mean
        relative_change = score_diff / abs(baseline_mean) if baseline_mean != 0 else 0
        
        # Statistical significance test
        t_stat, p_value = stats.ttest_rel(baseline_scores, ablated_scores)
        
        # Confidence interval for difference
        diff_scores = np.array(baseline_scores) - np.array(ablated_scores)
        mean_diff = np.mean(diff_scores)
        std_diff = np.std(diff_scores, ddof=1)
        se_diff = std_diff / np.sqrt(len(diff_scores))
        
        alpha = 1 - self.config.confidence_level
        t_critical = stats.t.ppf(1 - alpha/2, len(diff_scores) - 1)
        ci_lower = mean_diff - t_critical * se_diff
        ci_upper = mean_diff + t_critical * se_diff
        
        return AblationResult(
            component_name=component_name,
            baseline_score=baseline_mean,
            ablated_score=ablated_mean,
            score_difference=score_diff,
            relative_change=relative_change,
            p_value=p_value,
            confidence_interval=(ci_lower, ci_upper),
            metadata=metadata
        )

class ArchitectureAblationStudy:
    """Architecture ablation study for deep learning models"""
    
    def __init__(self, config: AblationConfig = None):
        self.config = config or AblationConfig()
        self.evaluator = ModelEvaluator()
        self.results: List[AblationResult] = []
    
    def run_architecture_ablation(self, 
                                X_train: pd.DataFrame, 
                                y_train: pd.Series,
                                X_val: pd.DataFrame, 
                                y_val: pd.Series,
                                base_model_config: Dict[str, Any]) -> List[AblationResult]:
        """
        Run architecture ablation study for deep learning models
        
        Args:
            X_train, y_train: Training data
            X_val, y_val: Validation data
            base_model_config: Base model configuration to modify
        """
        logger.info("Starting architecture ablation study")
        
        # Define ablation variants
        ablation_variants = self._generate_architecture_variants(base_model_config)
        
        results = []
        for variant_name, variant_config in ablation_variants.items():
            logger.info(f"Testing architecture variant: {variant_name}")
            
            # Train and evaluate variant
            variant_scores = self._evaluate_architecture_variant(
                X_train, y_train, X_val, y_val, variant_config
            )
            
            result = AblationResult(
                component_name=variant_name,
                baseline_score=variant_scores["baseline"],
                ablated_score=variant_scores["variant"],
                score_difference=variant_scores["difference"],
                relative_change=variant_scores["relative_change"],
                metadata={"config": variant_config}
            )
            results.append(result)
        
        self.results.extend(results)
        return results
    
    def _generate_architecture_variants(self, base_config: Dict[str, Any]) -> Dict[str, Dict[str, Any]]:
        """Generate architecture variants for ablation study"""
        variants = {}
        
        # Layer depth ablation
        for n_layers in [1, 2, 3, 4, 6]:
            config = base_config.copy()
            config["n_layers"] = n_layers
            variants[f"layers_{n_layers}"] = config
        
        # Layer width ablation
        for hidden_size in [32, 64, 128, 256, 512]:
            config = base_config.copy()
            config["hidden_size"] = hidden_size
            variants[f"width_{hidden_size}"] = config
        
        # Dropout ablation
        for dropout in [0.0, 0.1, 0.2, 0.3, 0.5]:
            config = base_config.copy()
            config["dropout"] = dropout
            variants[f"dropout_{dropout}"] = config
        
        # Activation function ablation
        for activation in ["relu", "tanh", "gelu", "swish"]:
            config = base_config.copy()
            config["activation"] = activation
            variants[f"activation_{activation}"] = config
        
        return variants
    
    def _evaluate_architecture_variant(self, X_train: pd.DataFrame, y_train: pd.Series,
                                     X_val: pd.DataFrame, y_val: pd.Series,
                                     config: Dict[str, Any]) -> Dict[str, float]:
        """Evaluate a single architecture variant"""
        # This would integrate with your deep learning trainer
        # For now, return mock results
        baseline_score = np.random.uniform(0.6, 0.8)
        variant_score = baseline_score + np.random.uniform(-0.1, 0.1)
        
        return {
            "baseline": baseline_score,
            "variant": variant_score,
            "difference": baseline_score - variant_score,
            "relative_change": (baseline_score - variant_score) / abs(baseline_score)
        }

class DataSizeAblationStudy:
    """Data size ablation study to understand learning curves"""
    
    def __init__(self, config: AblationConfig = None):
        self.config = config or AblationConfig()
        self.results: List[AblationResult] = []
    
    def run_data_size_ablation(self, 
                             X: pd.DataFrame, 
                             y: pd.Series,
                             model_trainer: MLModelTrainer,
                             sample_sizes: Optional[List[int]] = None) -> List[AblationResult]:
        """
        Run data size ablation study
        
        Args:
            X: Feature matrix
            y: Target values
            model_trainer: Model trainer
            sample_sizes: List of sample sizes to test
        """
        logger.info("Starting data size ablation study")
        
        if sample_sizes is None:
            n_samples = len(X)
            sample_sizes = [
                int(n_samples * 0.1),
                int(n_samples * 0.2),
                int(n_samples * 0.3),
                int(n_samples * 0.5),
                int(n_samples * 0.7),
                int(n_samples * 0.9),
                n_samples
            ]
        
        results = []
        full_scores = self._get_baseline_performance(X, y, model_trainer)
        
        for size in sample_sizes:
            logger.info(f"Testing with {size} samples")
            
            # Sample data
            indices = np.random.choice(len(X), size=size, replace=False)
            X_sample = X.iloc[indices]
            y_sample = y.iloc[indices]
            
            # Get performance
            sample_scores = self._get_baseline_performance(X_sample, y_sample, model_trainer)
            
            result = AblationResult(
                component_name=f"data_size_{size}",
                baseline_score=np.mean(full_scores),
                ablated_score=np.mean(sample_scores),
                score_difference=np.mean(full_scores) - np.mean(sample_scores),
                relative_change=(np.mean(full_scores) - np.mean(sample_scores)) / abs(np.mean(full_scores)),
                metadata={"sample_size": size, "n_features": X.shape[1]}
            )
            results.append(result)
        
        self.results.extend(results)
        return results
    
    def _get_baseline_performance(self, X: pd.DataFrame, y: pd.Series,
                                model_trainer: MLModelTrainer) -> List[float]:
        """Get model performance using cross-validation"""
        kf = KFold(n_splits=min(self.config.cv_folds, len(X)//2), shuffle=True, 
                  random_state=self.config.random_state)
        
        scores = []
        for train_idx, val_idx in kf.split(X):
            X_train, X_val = X.iloc[train_idx], X.iloc[val_idx]
            y_train, y_val = y.iloc[train_idx], y.iloc[val_idx]
            
            model = model_trainer.train_single_model(X_train, y_train)
            y_pred = model.predict(X_val)
            score = r2_score(y_val, y_pred)
            scores.append(score)
        
        return scores

class HyperparameterAblationStudy:
    """Hyperparameter sensitivity analysis"""
    
    def __init__(self, config: AblationConfig = None):
        self.config = config or AblationConfig()
        self.results: List[AblationResult] = []
    
    def run_hyperparameter_ablation(self, 
                                  X: pd.DataFrame, 
                                  y: pd.Series,
                                  model_trainer: MLModelTrainer,
                                  param_grids: Dict[str, List[Any]]) -> List[AblationResult]:
        """
        Run hyperparameter ablation study
        
        Args:
            X: Feature matrix
            y: Target values
            model_trainer: Model trainer
            param_grids: Dictionary of parameter grids to test
        """
        logger.info("Starting hyperparameter ablation study")
        
        # Get baseline with default parameters
        baseline_scores = self._get_baseline_performance(X, y, model_trainer, {})
        
        results = []
        for param_name, param_values in param_grids.items():
            logger.info(f"Testing hyperparameter: {param_name}")
            
            for value in param_values:
                param_config = {param_name: value}
                variant_scores = self._get_baseline_performance(X, y, model_trainer, param_config)
                
                result = AblationResult(
                    component_name=f"{param_name}_{value}",
                    baseline_score=np.mean(baseline_scores),
                    ablated_score=np.mean(variant_scores),
                    score_difference=np.mean(baseline_scores) - np.mean(variant_scores),
                    relative_change=(np.mean(baseline_scores) - np.mean(variant_scores)) / abs(np.mean(baseline_scores)),
                    metadata={"parameter": param_name, "value": value}
                )
                results.append(result)
        
        self.results.extend(results)
        return results
    
    def _get_baseline_performance(self, X: pd.DataFrame, y: pd.Series,
                                model_trainer: MLModelTrainer, 
                                param_config: Dict[str, Any]) -> List[float]:
        """Get model performance with specific parameters"""
        kf = KFold(n_splits=self.config.cv_folds, shuffle=True, random_state=self.config.random_state)
        
        scores = []
        for train_idx, val_idx in kf.split(X):
            X_train, X_val = X.iloc[train_idx], X.iloc[val_idx]
            y_train, y_val = y.iloc[train_idx], y.iloc[val_idx]
            
            # Train with specific parameters
            model = model_trainer.train_single_model(X_train, y_train, **param_config)
            y_pred = model.predict(X_val)
            score = r2_score(y_val, y_pred)
            scores.append(score)
        
        return scores

class AblationStudyFramework:
    """Main framework for conducting comprehensive ablation studies"""
    
    def __init__(self, config: AblationConfig = None):
        self.config = config or AblationConfig()
        self.feature_study = FeatureAblationStudy(config)
        self.architecture_study = ArchitectureAblationStudy(config)
        self.data_size_study = DataSizeAblationStudy(config)
        self.hyperparameter_study = HyperparameterAblationStudy(config)
        
        self.all_results: List[AblationResult] = []
    
    def run_comprehensive_ablation(self, 
                                 X: pd.DataFrame, 
                                 y: pd.Series,
                                 model_trainer: MLModelTrainer,
                                 study_types: List[str] = None) -> Dict[str, List[AblationResult]]:
        """
        Run comprehensive ablation studies
        
        Args:
            X: Feature matrix
            y: Target values
            model_trainer: Model trainer
            study_types: List of study types to run
        """
        if study_types is None:
            study_types = ["feature", "data_size", "hyperparameter"]
        
        results = {}
        
        if "feature" in study_types:
            logger.info("Running feature ablation study")
            results["feature"] = self.feature_study.run_feature_ablation(X, y, model_trainer)
            self.all_results.extend(results["feature"])
        
        if "data_size" in study_types:
            logger.info("Running data size ablation study")
            results["data_size"] = self.data_size_study.run_data_size_ablation(X, y, model_trainer)
            self.all_results.extend(results["data_size"])
        
        if "hyperparameter" in study_types:
            logger.info("Running hyperparameter ablation study")
            # Define parameter grids for common algorithms
            param_grids = {
                "n_estimators": [50, 100, 200, 500],
                "max_depth": [3, 5, 10, None],
                "min_samples_split": [2, 5, 10],
                "learning_rate": [0.01, 0.1, 0.3]
            }
            results["hyperparameter"] = self.hyperparameter_study.run_hyperparameter_ablation(
                X, y, model_trainer, param_grids
            )
            self.all_results.extend(results["hyperparameter"])
        
        return results
    
    def generate_ablation_report(self, 
                               results: Dict[str, List[AblationResult]],
                               save_path: Optional[str] = None) -> Dict[str, Any]:
        """
        Generate comprehensive ablation study report
        
        Args:
            results: Dictionary of ablation study results
            save_path: Optional path to save report
        """
        logger.info("Generating ablation study report")
        
        report = {
            "summary": self._generate_summary(results),
            "feature_analysis": self._analyze_feature_importance(results.get("feature", [])),
            "data_size_analysis": self._analyze_data_size_impact(results.get("data_size", [])),
            "hyperparameter_analysis": self._analyze_hyperparameter_sensitivity(results.get("hyperparameter", [])),
            "statistical_significance": self._analyze_statistical_significance(),
            "recommendations": self._generate_recommendations(results)
        }
        
        if save_path:
            self._save_report(report, save_path)
        
        return report
    
    def _generate_summary(self, results: Dict[str, List[AblationResult]]) -> Dict[str, Any]:
        """Generate summary statistics for ablation studies"""
        summary = {
            "total_studies": len(results),
            "total_experiments": sum(len(r) for r in results.values()),
            "significant_effects": len([r for r in self.all_results 
                                     if r.p_value and r.p_value < 0.05]),
            "largest_effect": max(self.all_results, key=lambda x: abs(x.score_difference)) if self.all_results else None,
            "most_sensitive_component": self._find_most_sensitive_component()
        }
        return summary
    
    def _analyze_feature_importance(self, feature_results: List[AblationResult]) -> Dict[str, Any]:
        """Analyze feature importance from ablation results"""
        if not feature_results:
            return {}
        
        # Sort by impact
        sorted_features = sorted(feature_results, key=lambda x: abs(x.score_difference), reverse=True)
        
        return {
            "most_important_features": sorted_features[:10],
            "least_important_features": sorted_features[-5:],
            "feature_impact_distribution": {
                "mean_impact": np.mean([abs(r.score_difference) for r in feature_results]),
                "std_impact": np.std([abs(r.score_difference) for r in feature_results])
            }
        }
    
    def _analyze_data_size_impact(self, data_size_results: List[AblationResult]) -> Dict[str, Any]:
        """Analyze data size impact from ablation results"""
        if not data_size_results:
            return {}
        
        # Extract sample sizes and scores
        sample_sizes = [r.metadata["sample_size"] for r in data_size_results]
        scores = [r.ablated_score for r in data_size_results]
        
        return {
            "learning_curve": list(zip(sample_sizes, scores)),
            "data_efficiency": self._calculate_data_efficiency(sample_sizes, scores),
            "minimum_data_requirement": self._estimate_minimum_data_requirement(sample_sizes, scores)
        }
    
    def _analyze_hyperparameter_sensitivity(self, hyperparameter_results: List[AblationResult]) -> Dict[str, Any]:
        """Analyze hyperparameter sensitivity from ablation results"""
        if not hyperparameter_results:
            return {}
        
        # Group by parameter
        param_groups = {}
        for result in hyperparameter_results:
            param_name = result.metadata["parameter"]
            if param_name not in param_groups:
                param_groups[param_name] = []
            param_groups[param_name].append(result)
        
        sensitivity_analysis = {}
        for param_name, param_results in param_groups.items():
            scores = [r.ablated_score for r in param_results]
            sensitivity_analysis[param_name] = {
                "score_range": max(scores) - min(scores),
                "score_std": np.std(scores),
                "optimal_value": param_results[np.argmax(scores)].metadata["value"]
            }
        
        return sensitivity_analysis
    
    def _analyze_statistical_significance(self) -> Dict[str, Any]:
        """Analyze statistical significance of ablation results"""
        significant_results = [r for r in self.all_results if r.p_value and r.p_value < 0.05]
        
        return {
            "significant_results_count": len(significant_results),
            "total_results_count": len(self.all_results),
            "significance_rate": len(significant_results) / len(self.all_results) if self.all_results else 0,
            "significant_results": significant_results
        }
    
    def _generate_recommendations(self, results: Dict[str, List[AblationResult]]) -> List[str]:
        """Generate recommendations based on ablation study results"""
        recommendations = []
        
        # Feature recommendations
        if "feature" in results:
            important_features = [r for r in results["feature"] 
                                if abs(r.score_difference) > self.config.min_effect_size]
            if important_features:
                recommendations.append(
                    f"Focus on {len(important_features)} most important features for optimal performance"
                )
        
        # Data size recommendations
        if "data_size" in results:
            recommendations.append(
                "Consider data augmentation or collection if performance plateaus with current dataset size"
            )
        
        # Hyperparameter recommendations
        if "hyperparameter" in results:
            recommendations.append(
                "Hyperparameter optimization shows significant impact - invest in automated tuning"
            )
        
        return recommendations
    
    def _find_most_sensitive_component(self) -> Optional[AblationResult]:
        """Find the component with highest sensitivity to ablation"""
        if not self.all_results:
            return None
        
        return max(self.all_results, key=lambda x: abs(x.relative_change))
    
    def _calculate_data_efficiency(self, sample_sizes: List[int], scores: List[float]) -> float:
        """Calculate data efficiency metric"""
        if len(sample_sizes) < 2:
            return 0.0
        
        # Calculate improvement per additional sample
        improvements = []
        for i in range(1, len(sample_sizes)):
            score_improvement = scores[i] - scores[i-1]
            sample_increase = sample_sizes[i] - sample_sizes[i-1]
            if sample_increase > 0:
                improvements.append(score_improvement / sample_increase)
        
        return np.mean(improvements) if improvements else 0.0
    
    def _estimate_minimum_data_requirement(self, sample_sizes: List[int], scores: List[float]) -> int:
        """Estimate minimum data requirement for reasonable performance"""
        if len(sample_sizes) < 2:
            return sample_sizes[0] if sample_sizes else 0
        
        # Find point where performance improvement becomes minimal
        max_score = max(scores)
        threshold = max_score * 0.95  # 95% of maximum performance
        
        for i, score in enumerate(scores):
            if score >= threshold:
                return sample_sizes[i]
        
        return sample_sizes[-1]
    
    def _save_report(self, report: Dict[str, Any], save_path: str):
        """Save ablation study report"""
        save_path = Path(save_path)
        save_path.mkdir(parents=True, exist_ok=True)
        
        # Save JSON report
        with open(save_path / "ablation_report.json", "w") as f:
            json.dump(report, f, indent=2, default=str)
        
        # Save summary CSV
        if self.all_results:
            df = pd.DataFrame([asdict(r) for r in self.all_results])
            df.to_csv(save_path / "ablation_results.csv", index=False)
        
        logger.info(f"Ablation study report saved to {save_path}")
    
    def plot_ablation_results(self, 
                            results: Dict[str, List[AblationResult]],
                            save_path: Optional[str] = None):
        """Create visualization plots for ablation study results"""
        logger.info("Creating ablation study visualizations")
        
        n_studies = len(results)
        if n_studies == 0:
            return
        
        fig, axes = plt.subplots(2, 2, figsize=(15, 12))
        axes = axes.flatten()
        
        plot_idx = 0
        
        # Feature ablation plot
        if "feature" in results and results["feature"]:
            self._plot_feature_ablation(results["feature"], axes[plot_idx])
            plot_idx += 1
        
        # Data size ablation plot
        if "data_size" in results and results["data_size"]:
            self._plot_data_size_ablation(results["data_size"], axes[plot_idx])
            plot_idx += 1
        
        # Hyperparameter ablation plot
        if "hyperparameter" in results and results["hyperparameter"]:
            self._plot_hyperparameter_ablation(results["hyperparameter"], axes[plot_idx])
            plot_idx += 1
        
        # Summary plot
        self._plot_ablation_summary(results, axes[plot_idx])
        
        plt.tight_layout()
        
        if save_path:
            plt.savefig(save_path, dpi=300, bbox_inches="tight")
        
        plt.show()
    
    def _plot_feature_ablation(self, results: List[AblationResult], ax):
        """Plot feature ablation results"""
        features = [r.component_name for r in results]
        impacts = [abs(r.score_difference) for r in results]
        
        # Sort by impact
        sorted_data = sorted(zip(features, impacts), key=lambda x: x[1], reverse=True)
        features, impacts = zip(*sorted_data[:10])  # Top 10
        
        ax.barh(features, impacts)
        ax.set_xlabel("Performance Impact (|ΔR²|)")
        ax.set_title("Feature Importance from Ablation Study")
        ax.grid(True, alpha=0.3)
    
    def _plot_data_size_ablation(self, results: List[AblationResult], ax):
        """Plot data size ablation results"""
        sample_sizes = [r.metadata["sample_size"] for r in results]
        scores = [r.ablated_score for r in results]
        
        ax.plot(sample_sizes, scores, "o-", linewidth=2, markersize=8)
        ax.set_xlabel("Sample Size")
        ax.set_ylabel("R² Score")
        ax.set_title("Learning Curve from Data Size Ablation")
        ax.grid(True, alpha=0.3)
    
    def _plot_hyperparameter_ablation(self, results: List[AblationResult], ax):
        """Plot hyperparameter ablation results"""
        # Group by parameter
        param_groups = {}
        for result in results:
            param_name = result.metadata["parameter"]
            if param_name not in param_groups:
                param_groups[param_name] = []
            param_groups[param_name].append(result)
        
        # Plot top 3 most sensitive parameters
        sorted_params = sorted(param_groups.items(), 
                             key=lambda x: np.std([r.ablated_score for r in x[1]]), 
                             reverse=True)[:3]
        
        for param_name, param_results in sorted_params:
            values = [r.metadata["value"] for r in param_results]
            scores = [r.ablated_score for r in param_results]
            ax.plot(values, scores, "o-", label=param_name, linewidth=2)
        
        ax.set_xlabel("Parameter Value")
        ax.set_ylabel("R² Score")
        ax.set_title("Hyperparameter Sensitivity")
        ax.legend()
        ax.grid(True, alpha=0.3)
    
    def _plot_ablation_summary(self, results: Dict[str, List[AblationResult]], ax):
        """Plot ablation study summary"""
        study_names = list(results.keys())
        study_impacts = []
        
        for study_name, study_results in results.items():
            if study_results:
                max_impact = max(abs(r.score_difference) for r in study_results)
                study_impacts.append(max_impact)
            else:
                study_impacts.append(0)
        
        bars = ax.bar(study_names, study_impacts)
        ax.set_ylabel("Maximum Performance Impact")
        ax.set_title("Ablation Study Summary")
        ax.grid(True, alpha=0.3)
        
        # Add value labels on bars
        for bar, impact in zip(bars, study_impacts):
            ax.text(bar.get_x() + bar.get_width()/2, bar.get_height() + 0.001,
                   f"{impact:.3f}", ha="center", va="bottom")
```
