## Supplementary material for "Machine Learning-Driven Prediction of TLR4 Binding Affinity: A Comprehensive Molecular Feature Analysis for Drug Discovery": Codebase: compound_analysis.py

```
"""
Compound analysis and ranking system.

Implements:
- Molecular similarity via Tanimoto (RDKit) with graceful fallback
- Clustering over structure + predicted affinity (KMeans)
- Chemical space viz with UMAP/t-SNE fallbacks to PCA
- Ranking by lowest predicted affinity with confidence intervals
- Simple SAR helper using correlation to descriptors when provided
"""

from typing import Dict, List, Optional, Any
import numpy as np
import pandas as pd

try:
    from rdkit import Chem
    from rdkit.Chem import rdFingerprintGenerator
    from rdkit import DataStructs
    RDKIT_AVAILABLE = True
except Exception:
    RDKIT_AVAILABLE = False

from sklearn.cluster import KMeans
from sklearn.decomposition import PCA

try:
    import umap  # type: ignore
    UMAP_AVAILABLE = True
except Exception:
    UMAP_AVAILABLE = False

try:
    from sklearn.manifold import TSNE
    TSNE_AVAILABLE = True
except Exception:
    TSNE_AVAILABLE = False


class CompoundAnalysis:
    """Compound analysis utilities for similarity, clustering, and ranking."""

    def __init__(self, random_state: int = 42):
        self.random_state = random_state

    def compute_tanimoto_similarity(self, smiles: List[str]) -> np.ndarray:
        """
        Compute pairwise Tanimoto similarity matrix from SMILES.
        Falls back to identity if RDKit is unavailable or input invalid.
        """
        n = len(smiles)
        if not RDKIT_AVAILABLE or n == 0:
            return np.eye(n, dtype=float)

        mols = [Chem.MolFromSmiles(s) if isinstance(s, str) else None for s in smiles]
        gen = rdFingerprintGenerator.GetMorganGenerator()
        fps = [gen.GetFingerprint(m) if m is not None else None for m in mols]

        sim = np.eye(n, dtype=float)
        for i in range(n):
            for j in range(i + 1, n):
                if fps[i] is None or fps[j] is None:
                    val = 0.0
                else:
                    try:
                        val = float(DataStructs.TanimotoSimilarity(fps[i], fps[j]))
                    except Exception:
                        val = 0.0
                sim[i, j] = val
                sim[j, i] = val
        return sim

    def cluster_compounds(self, X: np.ndarray, n_clusters: int = 5) -> np.ndarray:
        """KMeans clustering with sensible defaults."""
        if X.size == 0:
            return np.array([])
        model = KMeans(n_clusters=min(n_clusters, max(1, X.shape[0])) , n_init=10, random_state=self.random_state)
        return model.fit_predict(X)

    def chemical_space(self, X: np.ndarray, n_components: int = 2) -> np.ndarray:
        """
        Return 2D embedding using UMAP, else t-SNE, else PCA.
        """
        if X.size == 0:
            return np.zeros((0, n_components))
        if UMAP_AVAILABLE:
            reducer = umap.UMAP(n_components=n_components, random_state=self.random_state)
            return reducer.fit_transform(X)
        if TSNE_AVAILABLE:
            perplexity = max(5, min(30, X.shape[0] - 1))
            if perplexity < 1:
                return PCA(n_components=n_components, random_state=self.random_state).fit_transform(X)
            return TSNE(n_components=n_components, random_state=self.random_state, init="random", perplexity=perplexity).fit_transform(X)
        return PCA(n_components=n_components, random_state=self.random_state).fit_transform(X)

    def rank_compounds(
        self,
        df: pd.DataFrame,
        affinity_col: str = "predicted_affinity",
        lower_col: str = "confidence_interval_lower",
        upper_col: str = "confidence_interval_upper",
        compound_col: str = "compound_name",
    ) -> pd.DataFrame:
        """
        Rank by strongest binding (most negative affinity). Include CI.
        Break ties using CI lower bound, then uncertainty width.
        """
        required = [affinity_col]
        for c in required:
            if c not in df.columns:
                raise ValueError(f"Missing required column: {c}")

        out = df.copy()
        if lower_col not in out.columns:
            out[lower_col] = out[affinity_col] - 0.1
        if upper_col not in out.columns:
            out[upper_col] = out[affinity_col] + 0.1

        out["ci_width"] = (out[upper_col] - out[lower_col]).astype(float)
        out = out.sort_values([affinity_col, lower_col, "ci_width"], ascending=[True, True, True]).reset_index(drop=True)
        out["rank"] = np.arange(1, len(out) + 1)
        cols = ["rank", compound_col, affinity_col, lower_col, upper_col, "ci_width"]
        return out[[c for c in cols if c in out.columns]]

    def simple_sar(
        self,
        descriptors: pd.DataFrame,
        affinities: pd.Series,
        top_k: int = 10,
    ) -> pd.DataFrame:
        """
        Simple SAR: Pearson correlation of each descriptor vs affinity.
        Returns top_k descriptors by absolute correlation.
        """
        if len(descriptors) == 0:
            return pd.DataFrame(columns=["descriptor", "corr", "abs_corr"])
        corrs = []
        for col in descriptors.columns:
            try:
                x = pd.to_numeric(descriptors[col], errors="coerce")
                mask = x.notna() & pd.to_numeric(affinities, errors="coerce").notna()
                if mask.sum() < 3:
                    continue
                corr = float(np.corrcoef(x[mask], affinities[mask])[0, 1])
                corrs.append((col, corr, abs(corr)))
            except Exception:
                continue
        df_corr = pd.DataFrame(corrs, columns=["descriptor", "corr", "abs_corr"]).sort_values("abs_corr", ascending=False)
        return df_corr.head(top_k)
```
