## Supplementary material for "Machine Learning-Driven Prediction of TLR4 Binding Affinity: A Comprehensive Molecular Feature Analysis for Drug Discovery": Codebase: comprehensive_evaluator.py

```
"""
Comprehensive model evaluation framework for TLR4 binding prediction.

This module provides a complete evaluation framework that integrates with
all model types (traditional ML, deep learning, GNN, ensemble) and provides
comprehensive performance analysis, statistical testing, and visualization.
"""

import os
import json
import logging
from typing import Dict, List, Optional, Any, Tuple, Union
from pathlib import Path
import pandas as pd
import numpy as np
import matplotlib.pyplot as plt
import seaborn as sns
from sklearn.model_selection import KFold, cross_val_score
from sklearn.metrics import make_scorer
import warnings

from .evaluator import ModelEvaluator, PerformanceMetrics
from .trainer import MLModelTrainer
from .deep_learning_trainer import DeepLearningTrainer
from .ensemble_models import EnsembleTrainer

logger = logging.getLogger(__name__)


class ComprehensiveEvaluator:
    """
    Comprehensive evaluation framework for all model types in TLR4 binding prediction.
    
    This class provides a unified interface for evaluating traditional ML models,
    deep learning models, graph neural networks, and ensemble models with
    consistent metrics, statistical testing, and visualization.
    """
    
    def __init__(self, output_dir: str = "results/evaluation", 
                 confidence_level: float = 0.95):
        """
        Initialize comprehensive evaluator.
        
        Args:
            output_dir: Directory to save evaluation results and plots
            confidence_level: Confidence level for statistical tests
        """
        self.output_dir = Path(output_dir)
        self.output_dir.mkdir(parents=True, exist_ok=True)
        self.confidence_level = confidence_level
        
        # Initialize base evaluator
        self.base_evaluator = ModelEvaluator(confidence_level=confidence_level)
        
        # Store evaluation results
        self.evaluation_results = {}
        self.cross_validation_results = {}
        self.learning_curves = {}
        self.validation_curves = {}
        
        # Set up plotting style
        plt.style.use('default')
        sns.set_palette("husl")
        
        logger.info(f"Comprehensive evaluator initialized with output directory: {self.output_dir}")
    
    def evaluate_all_models(self, X_train: np.ndarray, y_train: np.ndarray,
                           X_val: np.ndarray, y_val: np.ndarray,
                           X_test: np.ndarray, y_test: np.ndarray,
                           feature_names: Optional[List[str]] = None,
                           model_configs: Optional[Dict[str, Dict]] = None) -> Dict[str, Any]:
        """
        Evaluate all model types comprehensively.
        
        Args:
            X_train: Training features
            y_train: Training targets
            X_val: Validation features
            y_val: Validation targets
            X_test: Test features
            y_test: Test targets
            feature_names: Names of features
            model_configs: Configuration for different model types
            
        Returns:
            Comprehensive evaluation results
        """
        logger.info("Starting comprehensive evaluation of all model types")
        
        if model_configs is None:
            model_configs = self._get_default_model_configs()
        
        results = {
            'traditional_ml': {},
            'deep_learning': {},
            'gnn': {},
            'ensemble': {},
            'cross_validation': {},
            'learning_curves': {},
            'validation_curves': {},
            'statistical_tests': {},
            'summary': {}
        }
        
        # Evaluate traditional ML models
        logger.info("Evaluating traditional ML models")
        results['traditional_ml'] = self._evaluate_traditional_ml_models(
            X_train, y_train, X_val, y_val, X_test, y_test, model_configs.get('traditional_ml', {})
        )
        
        # Evaluate deep learning models
        logger.info("Evaluating deep learning models")
        results['deep_learning'] = self._evaluate_deep_learning_models(
            X_train, y_train, X_val, y_val, X_test, y_test, model_configs.get('deep_learning', {})
        )
        
        # Evaluate ensemble models
        logger.info("Evaluating ensemble models")
        results['ensemble'] = self._evaluate_ensemble_models(
            X_train, y_train, X_val, y_val, X_test, y_test, model_configs.get('ensemble', {})
        )
        
        # Perform cross-validation evaluation
        logger.info("Performing cross-validation evaluation")
        results['cross_validation'] = self._perform_cross_validation_evaluation(
            X_train, y_train, results
        )
        
        # Generate learning curves for best models
        logger.info("Generating learning curves")
        results['learning_curves'] = self._generate_learning_curves(
            X_train, y_train, results
        )
        
        # Perform statistical significance testing
        logger.info("Performing statistical significance testing")
        results['statistical_tests'] = self._perform_statistical_tests(results)
        
        # Generate comprehensive summary
        results['summary'] = self._generate_evaluation_summary(results)
        
        # Save results
        self._save_evaluation_results(results)
        
        # Generate comprehensive plots
        self._generate_comprehensive_plots(results)
        
        logger.info("Comprehensive evaluation completed")
        return results
    
    def _evaluate_traditional_ml_models(self, X_train: np.ndarray, y_train: np.ndarray,
                                       X_val: np.ndarray, y_val: np.ndarray,
                                       X_test: np.ndarray, y_test: np.ndarray,
                                       config: Dict) -> Dict[str, Any]:
        """Evaluate traditional ML models."""
        results = {}
        
        try:
            trainer = MLModelTrainer()
            models = trainer.train_all_models(X_train, y_train, X_val, y_val)
            
            for model_name, model in models.items():
                # Test set evaluation
                y_pred = model.predict(X_test)
                metrics = self.base_evaluator.evaluate(y_test, y_pred, model_name)
                results[model_name] = {
                    'model': model,
                    'metrics': metrics,
                    'predictions': y_pred
                }
                
                logger.info(f"Traditional ML model {model_name} evaluated: R² = {metrics.metrics['r2']:.4f}")
        
        except Exception as e:
            logger.error(f"Error evaluating traditional ML models: {e}")
            results['error'] = str(e)
        
        return results
    
    def _evaluate_deep_learning_models(self, X_train: np.ndarray, y_train: np.ndarray,
                                      X_val: np.ndarray, y_val: np.ndarray,
                                      X_test: np.ndarray, y_test: np.ndarray,
                                      config: Dict) -> Dict[str, Any]:
        """Evaluate deep learning models."""
        results = {}
        
        try:
            trainer = DeepLearningTrainer()
            models = trainer.train_all_models(X_train, y_train, X_val, y_val)
            
            for model_name, model in models.items():
                # Test set evaluation
                y_pred = model.predict(X_test)
                metrics = self.base_evaluator.evaluate(y_test, y_pred, model_name)
                results[model_name] = {
                    'model': model,
                    'metrics': metrics,
                    'predictions': y_pred
                }
                
                logger.info(f"Deep learning model {model_name} evaluated: R² = {metrics.metrics['r2']:.4f}")
        
        except Exception as e:
            logger.error(f"Error evaluating deep learning models: {e}")
            results['error'] = str(e)
        
        return results
    
    def _evaluate_ensemble_models(self, X_train: np.ndarray, y_train: np.ndarray,
                                 X_val: np.ndarray, y_val: np.ndarray,
                                 X_test: np.ndarray, y_test: np.ndarray,
                                 config: Dict) -> Dict[str, Any]:
        """Evaluate ensemble models."""
        results = {}
        
        try:
            trainer = EnsembleTrainer()
            models = trainer.train_all_models(X_train, y_train, X_val, y_val)
            
            for model_name, model in models.items():
                # Test set evaluation
                y_pred = model.predict(X_test)
                metrics = self.base_evaluator.evaluate(y_test, y_pred, model_name)
                results[model_name] = {
                    'model': model,
                    'metrics': metrics,
                    'predictions': y_pred
                }
                
                logger.info(f"Ensemble model {model_name} evaluated: R² = {metrics.metrics['r2']:.4f}")
        
        except Exception as e:
            logger.error(f"Error evaluating ensemble models: {e}")
            results['error'] = str(e)
        
        return results
    
    def _perform_cross_validation_evaluation(self, X: np.ndarray, y: np.ndarray,
                                           all_results: Dict) -> Dict[str, Any]:
        """Perform cross-validation evaluation for all models."""
        cv_results = {}
        
        # Get all trained models
        all_models = {}
        for category in ['traditional_ml', 'deep_learning', 'ensemble']:
            if category in all_results and 'error' not in all_results[category]:
                for model_name, model_data in all_results[category].items():
                    if isinstance(model_data, dict) and 'model' in model_data:
                        all_models[f"{category}_{model_name}"] = model_data['model']
        
        # Perform cross-validation for each model
        for model_name, model in all_models.items():
            try:
                cv_result = self.base_evaluator.cross_validate_model(
                    model, X, y, model_name, cv=5
                )
                cv_results[model_name] = cv_result
                logger.info(f"Cross-validation completed for {model_name}")
            except Exception as e:
                logger.warning(f"Cross-validation failed for {model_name}: {e}")
                cv_results[model_name] = {'error': str(e)}
        
        return cv_results
    
    def _generate_learning_curves(self, X: np.ndarray, y: np.ndarray,
                                 all_results: Dict) -> Dict[str, Any]:
        """Generate learning curves for best performing models."""
        learning_curves = {}
        
        # Find best models from each category
        best_models = self._get_best_models(all_results)
        
        for category, model_info in best_models.items():
            try:
                model_name = model_info['name']
                model = model_info['model']
                
                # Generate learning curves
                curve_data = self.base_evaluator.generate_learning_curves(
                    model, X, y, model_name, save_path=str(self.output_dir / f"{model_name}_learning_curves.png")
                )
                learning_curves[model_name] = curve_data
                
            except Exception as e:
                logger.warning(f"Failed to generate learning curves for {category}: {e}")
        
        return learning_curves
    
    def _perform_statistical_tests(self, all_results: Dict) -> Dict[str, Any]:
        """Perform statistical significance tests between models."""
        statistical_tests = {}
        
        # Collect all metrics for comparison
        all_metrics = {}
        for category in ['traditional_ml', 'deep_learning', 'ensemble']:
            if category in all_results and 'error' not in all_results[category]:
                for model_name, model_data in all_results[category].items():
                    if isinstance(model_data, dict) and 'metrics' in model_data:
                        all_metrics[f"{category}_{model_name}"] = model_data['metrics']
        
        # Perform pairwise statistical tests
        model_names = list(all_metrics.keys())
        for i in range(len(model_names)):
            for j in range(i + 1, len(model_names)):
                model1_name, model2_name = model_names[i], model_names[j]
                test_name = f"{model1_name}_vs_{model2_name}"
                
                try:
                    test_result = self.base_evaluator.statistical_significance_test(
                        all_metrics[model1_name], all_metrics[model2_name]
                    )
                    statistical_tests[test_name] = test_result
                except Exception as e:
                    logger.warning(f"Statistical test failed for {test_name}: {e}")
                    statistical_tests[test_name] = {'error': str(e)}
        
        return statistical_tests
    
    def _get_best_models(self, all_results: Dict) -> Dict[str, Dict]:
        """Get best performing model from each category."""
        best_models = {}
        
        for category in ['traditional_ml', 'deep_learning', 'ensemble']:
            if category in all_results and 'error' not in all_results[category]:
                best_r2 = -float('inf')
                best_model = None
                
                for model_name, model_data in all_results[category].items():
                    if isinstance(model_data, dict) and 'metrics' in model_data:
                        r2_score = model_data['metrics'].metrics['r2']
                        if r2_score > best_r2:
                            best_r2 = r2_score
                            best_model = {
                                'name': model_name,
                                'model': model_data['model'],
                                'r2_score': r2_score
                            }
                
                if best_model:
                    best_models[category] = best_model
        
        return best_models
    
    def _generate_evaluation_summary(self, all_results: Dict) -> Dict[str, Any]:
        """Generate comprehensive evaluation summary."""
        summary = {
            'total_models_evaluated': 0,
            'best_overall_model': None,
            'best_r2_score': -float('inf'),
            'category_performance': {},
            'cross_validation_summary': {},
            'statistical_significance_summary': {}
        }
        
        # Collect all models and their performance
        all_models_performance = []
        
        for category in ['traditional_ml', 'deep_learning', 'ensemble']:
            if category in all_results and 'error' not in all_results[category]:
                category_models = []
                for model_name, model_data in all_results[category].items():
                    if isinstance(model_data, dict) and 'metrics' in model_data:
                        r2_score = model_data['metrics'].metrics['r2']
                        rmse_score = model_data['metrics'].metrics['rmse']
                        
                        model_info = {
                            'category': category,
                            'name': model_name,
                            'r2_score': r2_score,
                            'rmse_score': rmse_score
                        }
                        
                        all_models_performance.append(model_info)
                        category_models.append(model_info)
                        
                        # Track best overall model
                        if r2_score > summary['best_r2_score']:
                            summary['best_r2_score'] = r2_score
                            summary['best_overall_model'] = f"{category}_{model_name}"
                
                # Category summary
                if category_models:
                    category_df = pd.DataFrame(category_models)
                    summary['category_performance'][category] = {
                        'best_r2': category_df['r2_score'].max(),
                        'best_model': category_df.loc[category_df['r2_score'].idxmax(), 'name'],
                        'mean_r2': category_df['r2_score'].mean(),
                        'model_count': len(category_models)
                    }
        
        summary['total_models_evaluated'] = len(all_models_performance)
        
        # Cross-validation summary
        if 'cross_validation' in all_results:
            cv_summary = {}
            for model_name, cv_result in all_results['cross_validation'].items():
                if 'error' not in cv_result and 'r2' in cv_result:
                    cv_summary[model_name] = {
                        'mean_r2': cv_result['r2']['mean'],
                        'std_r2': cv_result['r2']['std']
                    }
            summary['cross_validation_summary'] = cv_summary
        
        return summary
    
    def _save_evaluation_results(self, results: Dict[str, Any]) -> None:
        """Save evaluation results to files."""
        # Save summary as JSON
        summary_path = self.output_dir / "evaluation_summary.json"
        with open(summary_path, 'w') as f:
            # Convert numpy types to Python types for JSON serialization
            summary_serializable = self._make_serializable(results['summary'])
            json.dump(summary_serializable, f, indent=2)
        
        # Save detailed results as pickle
        import pickle
        detailed_path = self.output_dir / "detailed_evaluation_results.pkl"
        with open(detailed_path, 'wb') as f:
            pickle.dump(results, f)
        
        logger.info(f"Evaluation results saved to {self.output_dir}")
    
    def _make_serializable(self, obj: Any) -> Any:
        """Convert numpy types to Python types for JSON serialization."""
        if isinstance(obj, dict):
            return {key: self._make_serializable(value) for key, value in obj.items()}
        elif isinstance(obj, list):
            return [self._make_serializable(item) for item in obj]
        elif isinstance(obj, np.integer):
            return int(obj)
        elif isinstance(obj, np.floating):
            return float(obj)
        elif isinstance(obj, np.ndarray):
            return obj.tolist()
        else:
            return obj
    
    def _generate_comprehensive_plots(self, results: Dict[str, Any]) -> None:
        """Generate comprehensive visualization plots."""
        logger.info("Generating comprehensive evaluation plots")
        
        # Model comparison plot
        self._plot_model_comparison(results)
        
        # Cross-validation comparison plot
        self._plot_cross_validation_comparison(results)
        
        # Statistical significance heatmap
        self._plot_statistical_significance_heatmap(results)
        
        # Performance distribution plot
        self._plot_performance_distribution(results)
    
    def _plot_model_comparison(self, results: Dict[str, Any]) -> None:
        """Plot comprehensive model comparison."""
        fig, axes = plt.subplots(2, 2, figsize=(16, 12))
        fig.suptitle('Comprehensive Model Performance Comparison', fontsize=16, fontweight='bold')
        
        # Collect all model performances
        model_data = []
        for category in ['traditional_ml', 'deep_learning', 'ensemble']:
            if category in results and 'error' not in results[category]:
                for model_name, model_result in results[category].items():
                    if isinstance(model_result, dict) and 'metrics' in model_result:
                        metrics = model_result['metrics']
                        model_data.append({
                            'model': f"{category}\n{model_name}",
                            'r2': metrics.metrics['r2'],
                            'rmse': metrics.metrics['rmse'],
                            'mae': metrics.metrics['mae'],
                            'pearson_r': metrics.metrics['pearson_r'],
                            'category': category
                        })
        
        if not model_data:
            logger.warning("No model data available for comparison plot")
            return
        
        df = pd.DataFrame(model_data)
        
        # R² scores
        sns.barplot(data=df, x='model', y='r2', hue='category', ax=axes[0, 0])
        axes[0, 0].set_title('R² Score Comparison')
        axes[0, 0].set_ylabel('R² Score')
        axes[0, 0].tick_params(axis='x', rotation=45)
        
        # RMSE scores
        sns.barplot(data=df, x='model', y='rmse', hue='category', ax=axes[0, 1])
        axes[0, 1].set_title('RMSE Comparison')
        axes[0, 1].set_ylabel('RMSE')
        axes[0, 1].tick_params(axis='x', rotation=45)
        
        # MAE scores
        sns.barplot(data=df, x='model', y='mae', hue='category', ax=axes[1, 0])
        axes[1, 0].set_title('MAE Comparison')
        axes[1, 0].set_ylabel('MAE')
        axes[1, 0].tick_params(axis='x', rotation=45)
        
        # Pearson correlation
        sns.barplot(data=df, x='model', y='pearson_r', hue='category', ax=axes[1, 1])
        axes[1, 1].set_title('Pearson Correlation Comparison')
        axes[1, 1].set_ylabel('Pearson Correlation')
        axes[1, 1].tick_params(axis='x', rotation=45)
        
        plt.tight_layout()
        plt.savefig(self.output_dir / "comprehensive_model_comparison.png", 
                   dpi=300, bbox_inches='tight')
        plt.show()
    
    def _plot_cross_validation_comparison(self, results: Dict[str, Any]) -> None:
        """Plot cross-validation comparison."""
        if 'cross_validation' not in results:
            return
        
        cv_data = []
        for model_name, cv_result in results['cross_validation'].items():
            if 'error' not in cv_result and 'r2' in cv_result:
                cv_data.append({
                    'model': model_name,
                    'mean_r2': cv_result['r2']['mean'],
                    'std_r2': cv_result['r2']['std'],
                    'mean_rmse': cv_result.get('rmse', {}).get('mean', 0),
                    'std_rmse': cv_result.get('rmse', {}).get('std', 0)
                })
        
        if not cv_data:
            return
        
        df = pd.DataFrame(cv_data)
        
        fig, axes = plt.subplots(1, 2, figsize=(15, 6))
        fig.suptitle('Cross-Validation Performance Comparison', fontsize=14, fontweight='bold')
        
        # R² with error bars
        axes[0].bar(df['model'], df['mean_r2'], yerr=df['std_r2'], 
                   capsize=5, alpha=0.7)
        axes[0].set_title('R² Score (Cross-Validation)')
        axes[0].set_ylabel('R² Score')
        axes[0].tick_params(axis='x', rotation=45)
        
        # RMSE with error bars
        axes[1].bar(df['model'], df['mean_rmse'], yerr=df['std_rmse'], 
                   capsize=5, alpha=0.7)
        axes[1].set_title('RMSE (Cross-Validation)')
        axes[1].set_ylabel('RMSE')
        axes[1].tick_params(axis='x', rotation=45)
        
        plt.tight_layout()
        plt.savefig(self.output_dir / "cross_validation_comparison.png", 
                   dpi=300, bbox_inches='tight')
        plt.show()
    
    def _plot_statistical_significance_heatmap(self, results: Dict[str, Any]) -> None:
        """Plot statistical significance heatmap."""
        if 'statistical_tests' not in results:
            return
        
        # Create significance matrix
        model_names = set()
        for test_name in results['statistical_tests'].keys():
            models = test_name.split('_vs_')
            model_names.update(models)
        
        model_names = sorted(list(model_names))
        n_models = len(model_names)
        
        significance_matrix = np.zeros((n_models, n_models))
        
        for test_name, test_result in results['statistical_tests'].items():
            if 'error' not in test_result:
                models = test_name.split('_vs_')
                if len(models) == 2:
                    i = model_names.index(models[0])
                    j = model_names.index(models[1])
                    
                    # Use p-value from paired t-test
                    if 'paired_t_test' in test_result:
                        p_value = test_result['paired_t_test']['p_value']
                        significance_matrix[i, j] = -np.log10(p_value) if p_value > 0 else 10
                        significance_matrix[j, i] = -np.log10(p_value) if p_value > 0 else 10
        
        # Plot heatmap
        plt.figure(figsize=(12, 10))
        sns.heatmap(significance_matrix, 
                   xticklabels=model_names, 
                   yticklabels=model_names,
                   annot=True, 
                   fmt='.2f',
                   cmap='RdYlBu_r',
                   center=1.3)  # -log10(0.05) ≈ 1.3
        
        plt.title('Statistical Significance Heatmap\n(-log10(p-value), red=significant)', 
                 fontsize=14, fontweight='bold')
        plt.tight_layout()
        plt.savefig(self.output_dir / "statistical_significance_heatmap.png", 
                   dpi=300, bbox_inches='tight')
        plt.show()
    
    def _plot_performance_distribution(self, results: Dict[str, Any]) -> None:
        """Plot performance distribution across models."""
        # Collect all R² scores
        r2_scores = []
        categories = []
        
        for category in ['traditional_ml', 'deep_learning', 'ensemble']:
            if category in results and 'error' not in results[category]:
                for model_name, model_result in results[category].items():
                    if isinstance(model_result, dict) and 'metrics' in model_result:
                        r2_scores.append(model_result['metrics'].metrics['r2'])
                        categories.append(category)
        
        if not r2_scores:
            return
        
        # Create distribution plot
        fig, axes = plt.subplots(1, 2, figsize=(15, 6))
        fig.suptitle('Model Performance Distribution', fontsize=14, fontweight='bold')
        
        # Box plot by category
        df = pd.DataFrame({'r2_score': r2_scores, 'category': categories})
        sns.boxplot(data=df, x='category', y='r2_score', ax=axes[0])
        axes[0].set_title('R² Score Distribution by Category')
        axes[0].set_ylabel('R² Score')
        
        # Histogram of all R² scores
        axes[1].hist(r2_scores, bins=15, alpha=0.7, edgecolor='black')
        axes[1].set_title('Overall R² Score Distribution')
        axes[1].set_xlabel('R² Score')
        axes[1].set_ylabel('Frequency')
        axes[1].axvline(np.mean(r2_scores), color='red', linestyle='--', 
                       label=f'Mean: {np.mean(r2_scores):.3f}')
        axes[1].legend()
        
        plt.tight_layout()
        plt.savefig(self.output_dir / "performance_distribution.png", 
                   dpi=300, bbox_inches='tight')
        plt.show()
    
    def _get_default_model_configs(self) -> Dict[str, Dict]:
        """Get default model configurations."""
        return {
            'traditional_ml': {
                'random_forest': {'n_estimators': 100, 'max_depth': 10},
                'xgboost': {'n_estimators': 100, 'max_depth': 6},
                'svr': {'C': 1.0, 'gamma': 'scale'},
                'lightgbm': {'n_estimators': 100, 'max_depth': 6}
            },
            'deep_learning': {
                'mlp': {'hidden_layers': [128, 64], 'dropout': 0.2},
                'cnn': {'filters': [32, 64], 'kernel_size': 3},
                'transformer': {'d_model': 64, 'nhead': 4}
            },
            'ensemble': {
                'stacked_ensemble': {'base_models': ['random_forest', 'xgboost', 'lightgbm']},
                'weighted_ensemble': {'weights_method': 'cv_based'}
            }
        }
    
    def generate_evaluation_report(self, results: Dict[str, Any]) -> str:
        """
        Generate a comprehensive evaluation report.
        
        Args:
            results: Evaluation results from evaluate_all_models
            
        Returns:
            Formatted evaluation report string
        """
        report = []
        report.append("=" * 80)
        report.append("COMPREHENSIVE MODEL EVALUATION REPORT")
        report.append("TLR4 Binding Affinity Prediction")
        report.append("=" * 80)
        report.append("")
        
        # Summary section
        summary = results.get('summary', {})
        report.append("EXECUTIVE SUMMARY")
        report.append("-" * 40)
        report.append(f"Total Models Evaluated: {summary.get('total_models_evaluated', 0)}")
        report.append(f"Best Overall Model: {summary.get('best_overall_model', 'N/A')}")
        report.append(f"Best R² Score: {summary.get('best_r2_score', 0):.4f}")
        report.append("")
        
        # Category performance
        report.append("CATEGORY PERFORMANCE")
        report.append("-" * 40)
        category_perf = summary.get('category_performance', {})
        for category, perf in category_perf.items():
            report.append(f"{category.upper()}:")
            report.append(f"  Best Model: {perf.get('best_model', 'N/A')}")
            report.append(f"  Best R²: {perf.get('best_r2', 0):.4f}")
            report.append(f"  Mean R²: {perf.get('mean_r2', 0):.4f}")
            report.append(f"  Model Count: {perf.get('model_count', 0)}")
            report.append("")
        
        # Detailed model performance
        report.append("DETAILED MODEL PERFORMANCE")
        report.append("-" * 40)
        
        for category in ['traditional_ml', 'deep_learning', 'ensemble']:
            if category in results and 'error' not in results[category]:
                report.append(f"\n{category.upper()} MODELS:")
                for model_name, model_result in results[category].items():
                    if isinstance(model_result, dict) and 'metrics' in model_result:
                        metrics = model_result['metrics']
                        report.append(f"  {model_name}:")
                        report.append(f"    R² Score: {metrics.metrics['r2']:.4f}")
                        report.append(f"    RMSE: {metrics.metrics['rmse']:.4f}")
                        report.append(f"    MAE: {metrics.metrics['mae']:.4f}")
                        report.append(f"    Pearson r: {metrics.metrics['pearson_r']:.4f}")
                        report.append(f"    Spearman ρ: {metrics.metrics['spearman_r']:.4f}")
        
        # Cross-validation summary
        report.append("\nCROSS-VALIDATION RESULTS")
        report.append("-" * 40)
        cv_summary = summary.get('cross_validation_summary', {})
        for model_name, cv_result in cv_summary.items():
            report.append(f"{model_name}:")
            report.append(f"  Mean R²: {cv_result.get('mean_r2', 0):.4f} ± {cv_result.get('std_r2', 0):.4f}")
        
        # Statistical significance
        report.append("\nSTATISTICAL SIGNIFICANCE TESTS")
        report.append("-" * 40)
        stat_tests = results.get('statistical_tests', {})
        significant_tests = []
        
        for test_name, test_result in stat_tests.items():
            if 'error' not in test_result and 'paired_t_test' in test_result:
                p_value = test_result['paired_t_test']['p_value']
                significant = test_result['paired_t_test']['significant']
                
                if significant:
                    significant_tests.append(f"{test_name}: p = {p_value:.4f}")
        
        if significant_tests:
            report.append("Significant differences found:")
            for test in significant_tests:
                report.append(f"  {test}")
        else:
            report.append("No significant differences found between models.")
        
        report.append("\n" + "=" * 80)
        report.append("End of Report")
        report.append("=" * 80)
        
        return "\n".join(report)
```
