## Supplementary material for "Machine Learning-Driven Prediction of TLR4 Binding Affinity: A Comprehensive Molecular Feature Analysis for Drug Discovery": Codebase: data_splitting.py

```
"""
Data Splitting and Validation Framework for TLR4 Binding Prediction

This module provides comprehensive data splitting and validation capabilities
including stratified train/validation/test splits and k-fold cross-validation
for molecular binding affinity prediction.
"""

import numpy as np
import pandas as pd
from typing import Dict, List, Tuple, Optional, Any, Union
from sklearn.model_selection import (
    train_test_split, 
    StratifiedKFold, 
    KFold,
    cross_val_score,
    LeaveOneOut
)
from sklearn.preprocessing import KBinsDiscretizer
from dataclasses import dataclass
import logging
import warnings

logger = logging.getLogger(__name__)


@dataclass
class DataSplitConfig:
    """Configuration for data splitting parameters"""
    test_size: float = 0.15
    validation_size: float = 0.15
    train_size: float = 0.70
    random_state: int = 42
    stratify: bool = True
    n_bins: int = 5  # For stratified splitting based on continuous targets
    shuffle: bool = True


@dataclass
class CrossValidationConfig:
    """Configuration for cross-validation parameters"""
    n_folds: int = 5
    random_state: int = 42
    shuffle: bool = True
    stratify: bool = True
    cv_type: str = 'kfold'  # 'kfold', 'stratified_kfold', 'loo'


class DataQualityReporter:
    """
    Generates comprehensive data quality reports and statistics
    for molecular binding affinity datasets.
    """
    
    def __init__(self):
        self.quality_metrics = {}
        
    def generate_report(self, 
                       X: pd.DataFrame, 
                       y: pd.Series, 
                       split_info: Optional[Dict] = None) -> Dict[str, Any]:
        """
        Generate comprehensive data quality report.
        
        Args:
            X: Features DataFrame
            y: Target Series (binding affinities)
            split_info: Optional split information for detailed analysis
            
        Returns:
            Dictionary containing comprehensive quality metrics
        """
        logger.info("Generating data quality report")
        
        report = {
            "dataset_overview": self._get_dataset_overview(X, y),
            "feature_analysis": self._analyze_features(X),
            "target_analysis": self._analyze_target(y),
            "missing_data": self._analyze_missing_data(X, y),
            "outlier_analysis": self._analyze_outliers(X, y),
            "distribution_analysis": self._analyze_distributions(X, y),
            "correlation_analysis": self._analyze_correlations(X, y),
            "split_quality": split_info if split_info else {}
        }
        
        self.quality_metrics = report
        return report
        
    def _get_dataset_overview(self, X: pd.DataFrame, y: pd.Series) -> Dict[str, Any]:
        """Get basic dataset overview statistics."""
        return {
            "n_samples": len(X),
            "n_features": X.shape[1],
            "memory_usage_mb": X.memory_usage(deep=True).sum() / 1024**2,
            "target_range": (y.min(), y.max()),
            "target_mean": y.mean(),
            "target_std": y.std(),
            "feature_types": {
                "numeric": X.select_dtypes(include=[np.number]).shape[1],
                "categorical": X.select_dtypes(include=['object', 'category']).shape[1]
            }
        }
        
    def _analyze_features(self, X: pd.DataFrame) -> Dict[str, Any]:
        """Analyze feature characteristics."""
        numeric_features = X.select_dtypes(include=[np.number])
        
        if numeric_features.empty:
            return {"numeric_features": 0, "feature_stats": {}}
            
        feature_stats = {
            "mean": numeric_features.mean().to_dict(),
            "std": numeric_features.std().to_dict(),
            "min": numeric_features.min().to_dict(),
            "max": numeric_features.max().to_dict(),
            "zero_variance_features": (numeric_features.var() == 0).sum(),
            "high_variance_features": (numeric_features.var() > 100).sum()
        }
        
        return {
            "numeric_features": numeric_features.shape[1],
            "feature_stats": feature_stats,
            "feature_names": numeric_features.columns.tolist()
        }
        
    def _analyze_target(self, y: pd.Series) -> Dict[str, Any]:
        """Analyze target variable characteristics."""
        return {
            "count": y.count(),
            "mean": y.mean(),
            "std": y.std(),
            "min": y.min(),
            "max": y.max(),
            "median": y.median(),
            "q25": y.quantile(0.25),
            "q75": y.quantile(0.75),
            "skewness": y.skew(),
            "kurtosis": y.kurtosis(),
            "missing_count": y.isnull().sum(),
            "unique_values": y.nunique(),
            "value_counts": y.value_counts().head(10).to_dict()
        }
        
    def _analyze_missing_data(self, X: pd.DataFrame, y: pd.Series) -> Dict[str, Any]:
        """Analyze missing data patterns."""
        missing_X = X.isnull().sum()
        missing_y = y.isnull().sum()
        
        return {
            "features_with_missing": (missing_X > 0).sum(),
            "samples_with_missing": X.isnull().any(axis=1).sum(),
            "missing_by_feature": missing_X[missing_X > 0].to_dict(),
            "target_missing": missing_y,
            "missing_percentage": {
                "features": (missing_X > 0).mean() * 100,
                "samples": X.isnull().any(axis=1).mean() * 100,
                "target": (missing_y / len(y)) * 100
            }
        }
        
    def _analyze_outliers(self, X: pd.DataFrame, y: pd.Series) -> Dict[str, Any]:
        """Analyze outliers using IQR method."""
        numeric_features = X.select_dtypes(include=[np.number])
        
        if numeric_features.empty:
            return {"outlier_analysis": "No numeric features available"}
            
        outlier_info = {}
        for col in numeric_features.columns:
            Q1 = numeric_features[col].quantile(0.25)
            Q3 = numeric_features[col].quantile(0.75)
            IQR = Q3 - Q1
            lower_bound = Q1 - 1.5 * IQR
            upper_bound = Q3 + 1.5 * IQR
            
            outliers = ((numeric_features[col] < lower_bound) | 
                       (numeric_features[col] > upper_bound)).sum()
            
            outlier_info[col] = {
                "outlier_count": outliers,
                "outlier_percentage": (outliers / len(numeric_features)) * 100,
                "lower_bound": lower_bound,
                "upper_bound": upper_bound
            }
        
        # Target outliers
        Q1_y = y.quantile(0.25)
        Q3_y = y.quantile(0.75)
        IQR_y = Q3_y - Q1_y
        lower_bound_y = Q1_y - 1.5 * IQR_y
        upper_bound_y = Q3_y + 1.5 * IQR_y
        
        target_outliers = ((y < lower_bound_y) | (y > upper_bound_y)).sum()
        
        return {
            "feature_outliers": outlier_info,
            "target_outliers": {
                "count": target_outliers,
                "percentage": (target_outliers / len(y)) * 100,
                "lower_bound": lower_bound_y,
                "upper_bound": upper_bound_y
            }
        }
        
    def _analyze_distributions(self, X: pd.DataFrame, y: pd.Series) -> Dict[str, Any]:
        """Analyze data distributions."""
        numeric_features = X.select_dtypes(include=[np.number])
        
        distribution_info = {}
        for col in numeric_features.columns:
            distribution_info[col] = {
                "skewness": numeric_features[col].skew(),
                "kurtosis": numeric_features[col].kurtosis(),
                "normality_test": self._shapiro_wilk_test(numeric_features[col])
            }
            
        return {
            "feature_distributions": distribution_info,
            "target_distribution": {
                "skewness": y.skew(),
                "kurtosis": y.kurtosis(),
                "normality_test": self._shapiro_wilk_test(y)
            }
        }
        
    def _analyze_correlations(self, X: pd.DataFrame, y: pd.Series) -> Dict[str, Any]:
        """Analyze feature correlations and feature-target correlations."""
        numeric_features = X.select_dtypes(include=[np.number])
        
        if numeric_features.empty:
            return {"correlation_analysis": "No numeric features available"}
            
        # Feature-target correlations
        feature_target_corr = numeric_features.corrwith(y).abs().sort_values(ascending=False)
        
        # Feature-feature correlations
        feature_corr_matrix = numeric_features.corr().abs()
        high_corr_pairs = []
        
        for i in range(len(feature_corr_matrix.columns)):
            for j in range(i + 1, len(feature_corr_matrix.columns)):
                corr_val = feature_corr_matrix.iloc[i, j]
                if corr_val > 0.8:  # High correlation threshold
                    high_corr_pairs.append({
                        "feature1": feature_corr_matrix.columns[i],
                        "feature2": feature_corr_matrix.columns[j],
                        "correlation": corr_val
                    })
        
        # Calculate max correlation, handling edge case of single feature
        if len(feature_corr_matrix.columns) <= 1:
            max_correlation = 0.0
        else:
            upper_triangle = feature_corr_matrix.values[np.triu_indices_from(feature_corr_matrix.values, k=1)]
            max_correlation = upper_triangle.max() if len(upper_triangle) > 0 else 0.0
        
        return {
            "feature_target_correlations": feature_target_corr.to_dict(),
            "high_feature_correlations": high_corr_pairs,
            "max_feature_correlation": max_correlation
        }
        
    def _shapiro_wilk_test(self, data: pd.Series) -> Dict[str, float]:
        """Perform Shapiro-Wilk test for normality."""
        try:
            from scipy import stats
            statistic, p_value = stats.shapiro(data.dropna())
            return {
                "statistic": statistic,
                "p_value": p_value,
                "is_normal": p_value > 0.05
            }
        except ImportError:
            return {"error": "scipy not available for normality testing"}
        except Exception as e:
            return {"error": str(e)}
    
    def print_summary(self, report: Optional[Dict] = None) -> None:
        """Print a formatted summary of the data quality report."""
        if report is None:
            report = self.quality_metrics
            
        if not report:
            logger.warning("No quality report available. Generate report first.")
            return
            
        print("\n" + "="*60)
        print("DATA QUALITY REPORT")
        print("="*60)
        
        # Dataset Overview
        overview = report["dataset_overview"]
        print(f"\nDataset Overview:")
        print(f"  Samples: {overview['n_samples']}")
        print(f"  Features: {overview['n_features']}")
        print(f"  Memory Usage: {overview['memory_usage_mb']:.2f} MB")
        print(f"  Target Range: [{overview['target_range'][0]:.3f}, {overview['target_range'][1]:.3f}]")
        
        # Missing Data
        missing = report["missing_data"]
        print(f"\nMissing Data:")
        print(f"  Features with missing: {missing['features_with_missing']}")
        print(f"  Samples with missing: {missing['samples_with_missing']}")
        print(f"  Target missing: {missing['target_missing']}")
        
        # Outliers
        outliers = report["outlier_analysis"]
        if "target_outliers" in outliers:
            print(f"\nOutliers:")
            print(f"  Target outliers: {outliers['target_outliers']['count']} ({outliers['target_outliers']['percentage']:.1f}%)")
        
        # High correlations
        correlations = report["correlation_analysis"]
        if "high_feature_correlations" in correlations:
            print(f"\nHigh Feature Correlations (>0.8): {len(correlations['high_feature_correlations'])}")
        
        print("="*60)


class DataSplitter:
    """
    Handles train/validation/test data splitting with stratification support.
    
    Provides stratified splitting for continuous targets by binning the target
    variable and ensuring balanced representation across splits.
    """
    
    def __init__(self, config: Optional[DataSplitConfig] = None):
        """
        Initialize the data splitter.
        
        Args:
            config: Configuration object for splitting parameters
        """
        self.config = config or DataSplitConfig()
        self.split_info = {}
        self.discretizer = None
        
    def split_data(self, 
                   X: pd.DataFrame, 
                   y: pd.Series,
                   compound_names: Optional[pd.Series] = None) -> Tuple[pd.DataFrame, pd.DataFrame, pd.DataFrame, 
                                                                       pd.Series, pd.Series, pd.Series]:
        """
        Split data into train/validation/test sets.
        
        Args:
            X: Features DataFrame
            y: Target Series (binding affinities)
            compound_names: Optional compound names for tracking
            
        Returns:
            Tuple of (X_train, X_val, X_test, y_train, y_val, y_test)
        """
        logger.info(f"Splitting data: {self.config.train_size:.1%} train, "
                   f"{self.config.validation_size:.1%} val, "
                   f"{self.config.test_size:.1%} test")
        
        if len(X) != len(y):
            raise ValueError("X and y must have the same length")
            
        # Calculate actual split sizes
        n_samples = len(X)
        n_test = int(n_samples * self.config.test_size)
        n_val = int(n_samples * self.config.validation_size)
        n_train = n_samples - n_test - n_val
        
        # Create stratification labels if requested
        stratify_labels = None
        if self.config.stratify:
            stratify_labels = self._create_stratification_labels(y)
            
        # First split: separate test set
        X_temp, X_test, y_temp, y_test = train_test_split(
            X, y,
            test_size=n_test,
            random_state=self.config.random_state,
            stratify=stratify_labels,
            shuffle=self.config.shuffle
        )
        
        # Second split: separate train and validation from remaining data
        if stratify_labels is not None:
            # Recalculate stratification for remaining data
            stratify_temp = stratify_labels[X_temp.index]
        else:
            stratify_temp = None
            
        X_train, X_val, y_train, y_val = train_test_split(
            X_temp, y_temp,
            test_size=n_val,
            random_state=self.config.random_state + 1,
            stratify=stratify_temp,
            shuffle=self.config.shuffle
        )
        
        # Store split information
        self._store_split_info(X_train, X_val, X_test, y_train, y_val, y_test, compound_names)
        
        logger.info(f"Split completed: {len(X_train)} train, {len(X_val)} val, {len(X_test)} test")
        
        return X_train, X_val, X_test, y_train, y_val, y_test
        
    def _create_stratification_labels(self, y: pd.Series) -> np.ndarray:
        """
        Create stratification labels for continuous target using binning.
        
        Args:
            y: Target Series
            
        Returns:
            Array of stratification labels
        """
        # Use KBinsDiscretizer to create bins for stratification
        self.discretizer = KBinsDiscretizer(
            n_bins=self.config.n_bins,
            encode='ordinal',
            strategy='quantile'
        )
        
        # Fit and transform target to create bins
        stratify_labels = self.discretizer.fit_transform(y.values.reshape(-1, 1)).flatten()
        
        logger.info(f"Created {self.config.n_bins} stratification bins for continuous target")
        return stratify_labels
        
    def _store_split_info(self, 
                         X_train: pd.DataFrame, X_val: pd.DataFrame, X_test: pd.DataFrame,
                         y_train: pd.Series, y_val: pd.Series, y_test: pd.Series,
                         compound_names: Optional[pd.Series] = None) -> None:
        """Store detailed information about the data splits."""
        
        self.split_info = {
            "split_sizes": {
                "train": len(X_train),
                "validation": len(X_val),
                "test": len(X_test),
                "total": len(X_train) + len(X_val) + len(X_test)
            },
            "split_ratios": {
                "train": len(X_train) / (len(X_train) + len(X_val) + len(X_test)),
                "validation": len(X_val) / (len(X_train) + len(X_val) + len(X_test)),
                "test": len(X_test) / (len(X_train) + len(X_val) + len(X_test))
            },
            "target_statistics": {
                "train": {
                    "mean": y_train.mean(),
                    "std": y_train.std(),
                    "min": y_train.min(),
                    "max": y_train.max()
                },
                "validation": {
                    "mean": y_val.mean(),
                    "std": y_val.std(),
                    "min": y_val.min(),
                    "max": y_val.max()
                },
                "test": {
                    "mean": y_test.mean(),
                    "std": y_test.std(),
                    "min": y_test.min(),
                    "max": y_test.max()
                }
            },
            "config": {
                "test_size": self.config.test_size,
                "validation_size": self.config.validation_size,
                "train_size": self.config.train_size,
                "random_state": self.config.random_state,
                "stratify": self.config.stratify
            }
        }
        
        # Add compound names if provided
        if compound_names is not None:
            self.split_info["compound_names"] = {
                "train": compound_names[X_train.index].tolist(),
                "validation": compound_names[X_val.index].tolist(),
                "test": compound_names[X_test.index].tolist()
            }
    
    def get_split_info(self) -> Dict[str, Any]:
        """Get detailed information about the data splits."""
        return self.split_info.copy()
        
    def validate_splits(self, 
                       X_train: pd.DataFrame, X_val: pd.DataFrame, X_test: pd.DataFrame,
                       y_train: pd.Series, y_val: pd.Series, y_test: pd.Series) -> Dict[str, Any]:
        """
        Validate that data splits meet quality criteria.
        
        Args:
            X_train, X_val, X_test: Feature DataFrames
            y_train, y_val, y_test: Target Series
            
        Returns:
            Dictionary with validation results
        """
        validation_results = {
            "size_validation": self._validate_split_sizes(X_train, X_val, X_test),
            "distribution_validation": self._validate_target_distributions(y_train, y_val, y_test),
            "feature_validation": self._validate_feature_consistency(X_train, X_val, X_test),
            "overlap_validation": self._validate_no_overlap(X_train, X_val, X_test)
        }
        
        # Overall validation status
        all_valid = all(
            result.get("status", False) for result in validation_results.values()
        )
        validation_results["overall_valid"] = all_valid
        
        return validation_results
        
    def _validate_split_sizes(self, X_train: pd.DataFrame, X_val: pd.DataFrame, X_test: pd.DataFrame) -> Dict[str, Any]:
        """Validate that split sizes are reasonable."""
        total_size = len(X_train) + len(X_val) + len(X_test)
        train_ratio = len(X_train) / total_size
        val_ratio = len(X_val) / total_size
        test_ratio = len(X_test) / total_size
        
        # Check if ratios are within acceptable ranges
        expected_train = self.config.train_size
        expected_val = self.config.validation_size
        expected_test = self.config.test_size
        
        tolerance = 0.05  # 5% tolerance
        
        train_valid = abs(train_ratio - expected_train) <= tolerance
        val_valid = abs(val_ratio - expected_val) <= tolerance
        test_valid = abs(test_ratio - expected_test) <= tolerance
        
        return {
            "status": train_valid and val_valid and test_valid,
            "actual_ratios": {"train": train_ratio, "validation": val_ratio, "test": test_ratio},
            "expected_ratios": {"train": expected_train, "validation": expected_val, "test": expected_test},
            "tolerance": tolerance
        }
        
    def _validate_target_distributions(self, y_train: pd.Series, y_val: pd.Series, y_test: pd.Series) -> Dict[str, Any]:
        """Validate that target distributions are similar across splits."""
        # Calculate distribution statistics
        train_stats = {"mean": y_train.mean(), "std": y_train.std()}
        val_stats = {"mean": y_val.mean(), "std": y_val.std()}
        test_stats = {"mean": y_test.mean(), "std": y_test.std()}
        
        # Check if means and stds are within reasonable ranges
        mean_tolerance = 0.2  # 20% of overall std
        std_tolerance = 0.3   # 30% difference allowed
        
        overall_mean = (y_train.mean() + y_val.mean() + y_test.mean()) / 3
        overall_std = (y_train.std() + y_val.std() + y_test.std()) / 3
        
        mean_valid = all(
            abs(stats["mean"] - overall_mean) <= mean_tolerance * overall_std
            for stats in [train_stats, val_stats, test_stats]
        )
        
        std_valid = all(
            abs(stats["std"] - overall_std) <= std_tolerance * overall_std
            for stats in [train_stats, val_stats, test_stats]
        )
        
        return {
            "status": mean_valid and std_valid,
            "statistics": {"train": train_stats, "validation": val_stats, "test": test_stats},
            "overall_mean": overall_mean,
            "overall_std": overall_std
        }
        
    def _validate_feature_consistency(self, X_train: pd.DataFrame, X_val: pd.DataFrame, X_test: pd.DataFrame) -> Dict[str, Any]:
        """Validate that features are consistent across splits."""
        # Check feature names consistency
        train_features = set(X_train.columns)
        val_features = set(X_val.columns)
        test_features = set(X_test.columns)
        
        features_consistent = train_features == val_features == test_features
        
        return {
            "status": features_consistent,
            "train_features": len(train_features),
            "val_features": len(val_features),
            "test_features": len(test_features),
            "feature_names_match": features_consistent
        }
        
    def _validate_no_overlap(self, X_train: pd.DataFrame, X_val: pd.DataFrame, X_test: pd.DataFrame) -> Dict[str, Any]:
        """Validate that there's no overlap between splits."""
        train_indices = set(X_train.index)
        val_indices = set(X_val.index)
        test_indices = set(X_test.index)
        
        train_val_overlap = len(train_indices.intersection(val_indices))
        train_test_overlap = len(train_indices.intersection(test_indices))
        val_test_overlap = len(val_indices.intersection(test_indices))
        
        no_overlap = train_val_overlap == 0 and train_test_overlap == 0 and val_test_overlap == 0
        
        return {
            "status": no_overlap,
            "train_val_overlap": train_val_overlap,
            "train_test_overlap": train_test_overlap,
            "val_test_overlap": val_test_overlap
        }


class CrossValidationSetup:
    """
    Sets up and manages k-fold cross-validation for model evaluation.
    
    Supports multiple CV strategies including standard k-fold, stratified k-fold,
    and leave-one-out cross-validation.
    """
    
    def __init__(self, config: Optional[CrossValidationConfig] = None):
        """
        Initialize cross-validation setup.
        
        Args:
            config: Configuration object for CV parameters
        """
        self.config = config or CrossValidationConfig()
        self.cv_splits = None
        self.cv_info = {}
        
    def setup_cv(self, X: pd.DataFrame, y: pd.Series) -> Any:
        """
        Set up cross-validation splits.
        
        Args:
            X: Features DataFrame
            y: Target Series
            
        Returns:
            Cross-validation splitter object
        """
        logger.info(f"Setting up {self.config.cv_type} cross-validation with {self.config.n_folds} folds")
        
        if self.config.cv_type == 'kfold':
            cv = KFold(
                n_splits=self.config.n_folds,
                shuffle=self.config.shuffle,
                random_state=self.config.random_state
            )
            
        elif self.config.cv_type == 'stratified_kfold':
            # Create stratification labels for continuous target
            if self.config.stratify:
                discretizer = KBinsDiscretizer(
                    n_bins=min(5, len(y) // 4),  # Adaptive number of bins
                    encode='ordinal',
                    strategy='quantile'
                )
                stratify_labels = discretizer.fit_transform(y.values.reshape(-1, 1)).flatten()
                
                # Store discretizer for later use
                self._stratify_discretizer = discretizer
                
                cv = StratifiedKFold(
                    n_splits=self.config.n_folds,
                    shuffle=self.config.shuffle,
                    random_state=self.config.random_state
                )
            else:
                # If stratification is disabled, use regular KFold
                cv = KFold(
                    n_splits=self.config.n_folds,
                    shuffle=self.config.shuffle,
                    random_state=self.config.random_state
                )
                stratify_labels = None
                self._stratify_discretizer = None
            
        elif self.config.cv_type == 'loo':
            cv = LeaveOneOut()
            
        else:
            raise ValueError(f"Unknown CV type: {self.config.cv_type}")
            
        self.cv_splits = cv
        self._store_cv_info(X, y, cv)
        
        return cv
        
    def _store_cv_info(self, X: pd.DataFrame, y: pd.Series, cv: Any) -> None:
        """Store information about the CV setup."""
        self.cv_info = {
            "cv_type": self.config.cv_type,
            "n_folds": self.config.n_folds if hasattr(cv, 'n_splits') else len(X),
            "n_samples": len(X),
            "n_features": X.shape[1],
            "shuffle": self.config.shuffle,
            "stratify": self.config.stratify,
            "random_state": self.config.random_state,
            "target_stats": {
                "mean": y.mean(),
                "std": y.std(),
                "min": y.min(),
                "max": y.max()
            }
        }
        
    def get_cv_splits(self, X: pd.DataFrame, y: pd.Series) -> List[Tuple[np.ndarray, np.ndarray]]:
        """
        Get cross-validation splits as list of (train_idx, val_idx) tuples.
        
        Args:
            X: Features DataFrame
            y: Target Series
            
        Returns:
            List of (train_indices, validation_indices) tuples
        """
        if self.cv_splits is None:
            self.setup_cv(X, y)
            
        splits = []
        
        # Handle stratified splits with discretized labels
        if (self.config.cv_type == 'stratified_kfold' and 
            self.config.stratify and 
            hasattr(self, '_stratify_discretizer')):
            
            # Recreate stratification labels for the current data
            stratify_labels = self._stratify_discretizer.transform(y.values.reshape(-1, 1)).flatten()
            
            for train_idx, val_idx in self.cv_splits.split(X, stratify_labels):
                splits.append((train_idx, val_idx))
        else:
            # Regular splits
            for train_idx, val_idx in self.cv_splits.split(X, y):
                splits.append((train_idx, val_idx))
            
        return splits
        
    def validate_cv_setup(self, X: pd.DataFrame, y: pd.Series) -> Dict[str, Any]:
        """
        Validate that CV setup is appropriate for the dataset.
        
        Args:
            X: Features DataFrame
            y: Target Series
            
        Returns:
            Dictionary with validation results
        """
        validation_results = {
            "dataset_size_check": self._check_dataset_size(X, y),
            "cv_type_appropriateness": self._check_cv_type_appropriateness(X, y),
            "fold_size_validation": self._check_fold_sizes(X, y),
            "stratification_check": self._check_stratification(X, y)
        }
        
        all_valid = all(result.get("status", True) for result in validation_results.values())
        validation_results["overall_valid"] = all_valid
        
        return validation_results
        
    def _check_dataset_size(self, X: pd.DataFrame, y: pd.Series) -> Dict[str, Any]:
        """Check if dataset size is appropriate for CV."""
        n_samples = len(X)
        
        # Minimum samples per fold
        min_samples_per_fold = 2
        min_total_samples = self.config.n_folds * min_samples_per_fold
        
        size_adequate = n_samples >= min_total_samples
        
        return {
            "status": size_adequate,
            "n_samples": n_samples,
            "min_required": min_total_samples,
            "recommendation": "Consider reducing n_folds" if not size_adequate else "Dataset size adequate"
        }
        
    def _check_cv_type_appropriateness(self, X: pd.DataFrame, y: pd.Series) -> Dict[str, Any]:
        """Check if CV type is appropriate for the dataset."""
        n_samples = len(X)
        
        if self.config.cv_type == 'loo' and n_samples > 100:
            return {
                "status": False,
                "issue": "LOO CV with large dataset may be computationally expensive",
                "recommendation": "Consider using k-fold CV instead"
            }
        elif self.config.cv_type == 'stratified_kfold' and n_samples < 20:
            return {
                "status": False,
                "issue": "Stratified CV may not work well with small datasets",
                "recommendation": "Consider using standard k-fold CV"
            }
        else:
            return {
                "status": True,
                "message": "CV type is appropriate for dataset size"
            }
            
    def _check_fold_sizes(self, X: pd.DataFrame, y: pd.Series) -> Dict[str, Any]:
        """Check if fold sizes are reasonable."""
        n_samples = len(X)
        
        if self.config.cv_type == 'loo':
            return {"status": True, "fold_size": 1}
            
        # Calculate fold sizes for k-fold
        fold_size = n_samples // self.config.n_folds
        remainder = n_samples % self.config.n_folds
        
        # Check if fold sizes are too small
        min_fold_size = 2
        fold_size_adequate = fold_size >= min_fold_size
        
        return {
            "status": fold_size_adequate,
            "average_fold_size": fold_size,
            "size_variation": remainder,
            "min_fold_size": min_fold_size
        }
        
    def _check_stratification(self, X: pd.DataFrame, y: pd.Series) -> Dict[str, Any]:
        """Check stratification effectiveness."""
        if not self.config.stratify or self.config.cv_type not in ['stratified_kfold']:
            return {"status": True, "message": "Stratification not applicable"}
            
        # Check target distribution
        target_std = y.std()
        target_range = y.max() - y.min()
        
        # If target has low variance, stratification may not be effective
        cv_effective = target_std > 0.1 * target_range
        
        return {
            "status": cv_effective,
            "target_std": target_std,
            "target_range": target_range,
            "recommendation": "Consider disabling stratification" if not cv_effective else "Stratification should be effective"
        }
        
    def get_cv_info(self) -> Dict[str, Any]:
        """Get information about the CV setup."""
        return self.cv_info.copy()


class DataValidationFramework:
    """
    Comprehensive framework for data splitting, validation, and quality assessment.
    
    Combines data splitting, cross-validation, and quality reporting into a single
    cohesive framework for molecular binding affinity prediction.
    """
    
    def __init__(self, 
                 split_config: Optional[DataSplitConfig] = None,
                 cv_config: Optional[CrossValidationConfig] = None):
        """
        Initialize the validation framework.
        
        Args:
            split_config: Configuration for data splitting
            cv_config: Configuration for cross-validation
        """
        self.splitter = DataSplitter(split_config)
        self.cv_setup = CrossValidationSetup(cv_config)
        self.quality_reporter = DataQualityReporter()
        self.framework_info = {}
        
    def process_dataset(self, 
                       X: pd.DataFrame, 
                       y: pd.Series,
                       compound_names: Optional[pd.Series] = None,
                       generate_quality_report: bool = True) -> Dict[str, Any]:
        """
        Process dataset through complete validation framework.
        
        Args:
            X: Features DataFrame
            y: Target Series
            compound_names: Optional compound names for tracking
            generate_quality_report: Whether to generate quality report
            
        Returns:
            Dictionary containing all processing results
        """
        logger.info("Processing dataset through validation framework")
        
        results = {}
        
        # Generate quality report if requested
        if generate_quality_report:
            logger.info("Generating data quality report")
            quality_report = self.quality_reporter.generate_report(X, y)
            results["quality_report"] = quality_report
            
        # Split data
        logger.info("Splitting data into train/validation/test sets")
        X_train, X_val, X_test, y_train, y_val, y_test = self.splitter.split_data(
            X, y, compound_names
        )
        
        # Validate splits
        split_validation = self.splitter.validate_splits(
            X_train, X_val, X_test, y_train, y_val, y_test
        )
        
        # Setup cross-validation
        logger.info("Setting up cross-validation")
        cv = self.cv_setup.setup_cv(X_train, y_train)
        
        # Validate CV setup
        cv_validation = self.cv_setup.validate_cv_setup(X_train, y_train)
        
        # Store results
        results.update({
            "data_splits": {
                "X_train": X_train,
                "X_val": X_val,
                "X_test": X_test,
                "y_train": y_train,
                "y_val": y_val,
                "y_test": y_test
            },
            "split_info": self.splitter.get_split_info(),
            "split_validation": split_validation,
            "cv_setup": cv,
            "cv_info": self.cv_setup.get_cv_info(),
            "cv_validation": cv_validation
        })
        
        # Store framework info
        self.framework_info = results
        
        logger.info("Dataset processing completed successfully")
        
        return results
        
    def get_framework_summary(self) -> Dict[str, Any]:
        """Get comprehensive summary of the validation framework."""
        if not self.framework_info:
            return {"error": "No framework processing completed yet"}
            
        summary = {
            "dataset_overview": self.framework_info.get("quality_report", {}).get("dataset_overview", {}),
            "split_summary": self.framework_info.get("split_info", {}),
            "cv_summary": self.framework_info.get("cv_info", {}),
            "validation_status": {
                "split_valid": self.framework_info.get("split_validation", {}).get("overall_valid", False),
                "cv_valid": self.framework_info.get("cv_validation", {}).get("overall_valid", False)
            }
        }
        
        return summary
        
    def print_framework_summary(self) -> None:
        """Print a formatted summary of the validation framework."""
        summary = self.get_framework_summary()
        
        if "error" in summary:
            logger.warning(summary["error"])
            return
            
        print("\n" + "="*60)
        print("DATA VALIDATION FRAMEWORK SUMMARY")
        print("="*60)
        
        # Dataset overview
        overview = summary.get("dataset_overview", {})
        print(f"\nDataset: {overview.get('n_samples', 'N/A')} samples, {overview.get('n_features', 'N/A')} features")
        
        # Split summary
        split_summary = summary.get("split_summary", {})
        if "split_sizes" in split_summary:
            sizes = split_summary["split_sizes"]
            print(f"\nData Splits:")
            print(f"  Train: {sizes.get('train', 'N/A')} samples")
            print(f"  Validation: {sizes.get('validation', 'N/A')} samples")
            print(f"  Test: {sizes.get('test', 'N/A')} samples")
        
        # Validation status
        validation = summary.get("validation_status", {})
        print(f"\nValidation Status:")
        print(f"  Split Validation: {'✓' if validation.get('split_valid') else '✗'}")
        print(f"  CV Validation: {'✓' if validation.get('cv_valid') else '✗'}")
        
        print("="*60)
```
