## Supplementary material for "Machine Learning-Driven Prediction of TLR4 Binding Affinity: A Comprehensive Molecular Feature Analysis for Drug Discovery": Codebase: deep_learning_models.py

```
"""
Deep learning models for molecular binding prediction.

This module implements various deep learning architectures including
CNN for 3D molecular voxel representations, molecular transformers
for SMILES-based learning, and multi-task neural networks.
"""

from abc import ABC, abstractmethod
from typing import Dict, List, Optional, Union, Any, Tuple, NamedTuple
import pandas as pd
import numpy as np
import logging
from pathlib import Path
import joblib
from datetime import datetime

logger = logging.getLogger(__name__)

# PyTorch imports with error handling
try:
    import torch
    import torch.nn as nn
    import torch.nn.functional as F
    from torch.utils.data import Dataset, DataLoader
    from torch.optim.lr_scheduler import ReduceLROnPlateau, CosineAnnealingLR
    TORCH_AVAILABLE = True
except ImportError:
    logger.warning("PyTorch not available. Deep learning models will be limited.")
    TORCH_AVAILABLE = False
    # Define dummy classes for when PyTorch is not available
    class torch:
        class Tensor:
            pass
        class device:
            def __init__(self, device_str):
                self.device_str = device_str
        class float:
            pass
        class long:
            pass
        @staticmethod
        def tensor(data, dtype=None):
            return data
        @staticmethod
        def is_available():
            return False
        @staticmethod
        def cuda():
            return False
    class nn:
        class Module:
            def __init__(self):
                pass
            def train(self):
                pass
            def eval(self):
                pass
            def to(self, device):
                return self
        class Linear:
            def __init__(self, in_features, out_features):
                pass
        class Conv3d:
            def __init__(self, *args, **kwargs):
                pass
        class BatchNorm3d:
            def __init__(self, num_features):
                pass
        class MaxPool3d:
            def __init__(self, *args, **kwargs):
                pass
        class AdaptiveAvgPool3d:
            def __init__(self, *args, **kwargs):
                pass
        class Dropout:
            def __init__(self, p):
                pass
        class Embedding:
            def __init__(self, *args, **kwargs):
                pass
        class TransformerEncoder:
            def __init__(self, *args, **kwargs):
                pass
        class TransformerEncoderLayer:
            def __init__(self, *args, **kwargs):
                pass
        class LayerNorm:
            def __init__(self, *args, **kwargs):
                pass
        class MultiheadAttention:
            def __init__(self, *args, **kwargs):
                pass
        class ModuleList:
            def __init__(self, modules=None):
                self.modules = modules or []
            def append(self, module):
                self.modules.append(module)
            def __len__(self):
                return len(self.modules)
            def __getitem__(self, idx):
                return self.modules[idx]
    class F:
        @staticmethod
        def mse_loss(pred, target):
            return 0.0
        @staticmethod
        def relu(x):
            return x
        @staticmethod
        def dropout(x, p, training):
            return x
        @staticmethod
        def gelu(x):
            return x
    class Dataset:
        pass
    class DataLoader:
        def __init__(self, dataset, batch_size=1, shuffle=False, num_workers=0):
            self.dataset = dataset
            self.batch_size = batch_size
            self.shuffle = shuffle
            self.num_workers = num_workers
        def __iter__(self):
            return iter([])
        def __len__(self):
            return 0

# RDKit for molecular processing
try:
    from rdkit import Chem
    from rdkit.Chem import rdMolDescriptors, Descriptors
    from rdkit.Chem.rdMolDescriptors import GetMorganFingerprintAsBitVect
    RDKIT_AVAILABLE = True
except ImportError:
    logger.warning("RDKit not available. Molecular processing will be limited.")
    RDKIT_AVAILABLE = False


class MolecularVoxelData(NamedTuple):
    """Container for 3D molecular voxel data."""
    voxel_grid: np.ndarray  # 3D voxel grid
    compound_name: str
    binding_affinity: float


class SMILESData(NamedTuple):
    """Container for SMILES molecular data."""
    smiles: str
    compound_name: str
    binding_affinity: float
    token_ids: Optional[np.ndarray] = None


class VoxelDataset(Dataset):
    """PyTorch Dataset for 3D molecular voxel data."""
    
    def __init__(self, voxel_data: List[MolecularVoxelData]):
        """
        Initialize voxel dataset.
        
        Args:
            voxel_data: List of molecular voxel data
        """
        self.voxel_data = voxel_data
        
    def __len__(self) -> int:
        return len(self.voxel_data)
    
    def __getitem__(self, idx: int) -> Tuple[torch.Tensor, float]:
        """Get voxel grid and target at index."""
        data = self.voxel_data[idx]
        voxel_tensor = torch.tensor(data.voxel_grid, dtype=torch.float32).unsqueeze(0)  # Add channel dimension
        return voxel_tensor, data.binding_affinity


class SMILESDataset(Dataset):
    """PyTorch Dataset for SMILES molecular data."""
    
    def __init__(self, smiles_data: List[SMILESData], vocab_size: int = 1000, max_length: int = 128):
        """
        Initialize SMILES dataset.
        
        Args:
            smiles_data: List of SMILES data
            vocab_size: Vocabulary size for tokenization
            max_length: Maximum sequence length
        """
        self.smiles_data = smiles_data
        self.vocab_size = vocab_size
        self.max_length = max_length
        self.tokenizer = self._create_tokenizer()
        
    def __len__(self) -> int:
        return len(self.smiles_data)
    
    def __getitem__(self, idx: int) -> Tuple[torch.Tensor, float]:
        """Get tokenized SMILES and target at index."""
        data = self.smiles_data[idx]
        if data.token_ids is not None:
            token_ids = data.token_ids
        else:
            token_ids = self._tokenize_smiles(data.smiles)
        
        # Pad or truncate to max_length
        if len(token_ids) > self.max_length:
            token_ids = token_ids[:self.max_length]
        else:
            token_ids = np.pad(token_ids, (0, self.max_length - len(token_ids)), 'constant')
        
        return torch.tensor(token_ids, dtype=torch.long), data.binding_affinity
    
    def _create_tokenizer(self) -> Dict[str, int]:
        """Create SMILES tokenizer."""
        # Basic SMILES vocabulary
        vocab = {
            '<PAD>': 0, '<UNK>': 1, '<START>': 2, '<END>': 3,
            'C': 4, 'N': 5, 'O': 6, 'S': 7, 'P': 8, 'F': 9, 'Cl': 10, 'Br': 11, 'I': 12,
            'H': 13, 'c': 14, 'n': 15, 'o': 16, 's': 17, 'p': 18,
            '=': 19, '#': 20, '(': 21, ')': 22, '[': 23, ']': 24,
            '+': 25, '-': 26, '@': 27, '\\': 28, '/': 29, '1': 30, '2': 31, '3': 32, '4': 33
        }
        return vocab
    
    def _tokenize_smiles(self, smiles: str) -> np.ndarray:
        """Tokenize SMILES string."""
        tokens = []
        i = 0
        while i < len(smiles):
            # Check for two-character tokens first
            if i + 1 < len(smiles) and smiles[i:i+2] in self.tokenizer:
                tokens.append(self.tokenizer[smiles[i:i+2]])
                i += 2
            elif smiles[i] in self.tokenizer:
                tokens.append(self.tokenizer[smiles[i]])
                i += 1
            else:
                tokens.append(self.tokenizer['<UNK>'])
                i += 1
        
        return np.array(tokens)


class MolecularVoxelizer:
    """Converts molecular structures to 3D voxel grids."""
    
    def __init__(self, grid_size: int = 32, resolution: float = 1.0):
        """
        Initialize molecular voxelizer.
        
        Args:
            grid_size: Size of the 3D grid (grid_size x grid_size x grid_size)
            resolution: Resolution in Angstroms per voxel
        """
        self.grid_size = grid_size
        self.resolution = resolution
        
    def voxelize_pdbqt(self, pdbqt_path: str) -> Optional[MolecularVoxelData]:
        """
        Convert PDBQT file to 3D voxel grid.
        
        Args:
            pdbqt_path: Path to PDBQT file
            
        Returns:
            MolecularVoxelData object or None if conversion fails
        """
        try:
            # Parse PDBQT file
            coords, atom_types = self._parse_pdbqt(pdbqt_path)
            
            if coords is None or len(coords) == 0:
                logger.warning(f"No valid coordinates found in {pdbqt_path}")
                return None
            
            # Create voxel grid
            voxel_grid = self._create_voxel_grid(coords, atom_types)
            
            return MolecularVoxelData(
                voxel_grid=voxel_grid,
                compound_name=Path(pdbqt_path).stem,
                binding_affinity=0.0  # Will be set later
            )
            
        except Exception as e:
            logger.error(f"Error voxelizing {pdbqt_path}: {str(e)}")
            return None
    
    def _parse_pdbqt(self, pdbqt_path: str) -> Tuple[Optional[np.ndarray], Optional[List[int]]]:
        """Parse PDBQT file to extract coordinates and atom types."""
        coords = []
        atom_types = []
        
        try:
            with open(pdbqt_path, 'r') as f:
                for line in f:
                    if line.startswith('ATOM') or line.startswith('HETATM'):
                        # Extract coordinates using more robust parsing
                        try:
                            # Try column-based parsing first
                            x = float(line[30:38].strip())
                            y = float(line[38:46].strip())
                            z = float(line[46:54].strip())
                        except (ValueError, IndexError):
                            # Fall back to whitespace splitting
                            parts = line.split()
                            if len(parts) >= 8:
                                x = float(parts[5])
                                y = float(parts[6])
                                z = float(parts[7])
                            else:
                                logger.warning(f"Skipping malformed line: {line.strip()}")
                                continue
                        
                        coords.append([x, y, z])
                        
                        # Extract atom type - use the atom name from column 12-16
                        atom_name = line[12:16].strip()
                        atom_type = atom_name[0] if atom_name else 'C'
                        atom_types.append(self._atom_type_to_number(atom_type))
            
            return np.array(coords), atom_types
            
        except Exception as e:
            logger.error(f"Error parsing PDBQT file {pdbqt_path}: {str(e)}")
            return None, None
    
    def _atom_type_to_number(self, atom_type: str) -> int:
        """Convert atom type string to atomic number."""
        atom_type_map = {
            'C': 6, 'N': 7, 'O': 8, 'S': 16, 'P': 15,
            'H': 1, 'F': 9, 'Cl': 17, 'Br': 35, 'I': 53
        }
        return atom_type_map.get(atom_type, 6)
    
    def _create_voxel_grid(self, coords: np.ndarray, atom_types: List[int]) -> np.ndarray:
        """Create 3D voxel grid from coordinates and atom types."""
        # Center coordinates
        center = np.mean(coords, axis=0)
        coords_centered = coords - center
        
        # Scale to grid
        coords_scaled = coords_centered / self.resolution + self.grid_size // 2
        
        # Create voxel grid
        voxel_grid = np.zeros((self.grid_size, self.grid_size, self.grid_size), dtype=np.float32)
        
        for coord, atom_type in zip(coords_scaled, atom_types):
            x, y, z = coord.astype(int)
            
            # Check bounds
            if 0 <= x < self.grid_size and 0 <= y < self.grid_size and 0 <= z < self.grid_size:
                # Use atomic number as voxel value
                voxel_grid[x, y, z] = float(atom_type)
        
        return voxel_grid


class MolecularCNN3D(nn.Module):
    """3D CNN for molecular voxel grid analysis."""
    
    def __init__(self, input_channels: int = 1, num_classes: int = 1, 
                 dropout: float = 0.1, use_batch_norm: bool = True):
        """
        Initialize 3D CNN model.
        
        Args:
            input_channels: Number of input channels
            num_classes: Number of output classes (1 for regression)
            dropout: Dropout rate
            use_batch_norm: Whether to use batch normalization
        """
        super().__init__()
        
        if not TORCH_AVAILABLE:
            raise RuntimeError("PyTorch not available for CNN model")
        
        self.input_channels = input_channels
        self.num_classes = num_classes
        self.dropout = dropout
        self.use_batch_norm = use_batch_norm
        
        # 3D Convolutional layers
        self.conv_layers = nn.ModuleList([
            # First block
            nn.Conv3d(input_channels, 32, kernel_size=3, padding=1),
            nn.BatchNorm3d(32) if use_batch_norm else nn.Identity(),
            nn.ReLU(),
            nn.MaxPool3d(2),
            
            # Second block
            nn.Conv3d(32, 64, kernel_size=3, padding=1),
            nn.BatchNorm3d(64) if use_batch_norm else nn.Identity(),
            nn.ReLU(),
            nn.MaxPool3d(2),
            
            # Third block
            nn.Conv3d(64, 128, kernel_size=3, padding=1),
            nn.BatchNorm3d(128) if use_batch_norm else nn.Identity(),
            nn.ReLU(),
            nn.MaxPool3d(2),
            
            # Fourth block
            nn.Conv3d(128, 256, kernel_size=3, padding=1),
            nn.BatchNorm3d(256) if use_batch_norm else nn.Identity(),
            nn.ReLU(),
            nn.AdaptiveAvgPool3d(1)
        ])
        
        # Global average pooling
        self.global_pool = nn.AdaptiveAvgPool3d(1)
        
        # Fully connected layers
        self.fc_layers = nn.ModuleList([
            nn.Linear(256, 512),
            nn.ReLU(),
            nn.Dropout(dropout),
            nn.Linear(512, 256),
            nn.ReLU(),
            nn.Dropout(dropout),
            nn.Linear(256, num_classes)
        ])
        
    def forward(self, x: torch.Tensor) -> torch.Tensor:
        """Forward pass through the model."""
        # Apply convolutional layers
        for layer in self.conv_layers:
            x = layer(x)
        
        # Global pooling
        x = self.global_pool(x)
        x = x.view(x.size(0), -1)  # Flatten
        
        # Apply fully connected layers
        for layer in self.fc_layers:
            x = layer(x)
        
        return x


class MolecularTransformer(nn.Module):
    """Transformer model for SMILES-based molecular property prediction."""
    
    def __init__(self, vocab_size: int = 1000, d_model: int = 256, nhead: int = 8,
                 num_layers: int = 6, dim_feedforward: int = 1024, 
                 dropout: float = 0.1, max_length: int = 128):
        """
        Initialize molecular transformer.
        
        Args:
            vocab_size: Size of the vocabulary
            d_model: Model dimension
            nhead: Number of attention heads
            num_layers: Number of transformer layers
            dim_feedforward: Feedforward dimension
            dropout: Dropout rate
            max_length: Maximum sequence length
        """
        super().__init__()
        
        if not TORCH_AVAILABLE:
            raise RuntimeError("PyTorch not available for Transformer model")
        
        self.vocab_size = vocab_size
        self.d_model = d_model
        self.max_length = max_length
        
        # Embedding layers
        self.token_embedding = nn.Embedding(vocab_size, d_model)
        self.position_embedding = nn.Embedding(max_length, d_model)
        
        # Transformer encoder
        encoder_layer = nn.TransformerEncoderLayer(
            d_model=d_model,
            nhead=nhead,
            dim_feedforward=dim_feedforward,
            dropout=dropout,
            activation='gelu',
            batch_first=True
        )
        self.transformer = nn.TransformerEncoder(encoder_layer, num_layers=num_layers)
        
        # Output layers
        self.layer_norm = nn.LayerNorm(d_model)
        self.dropout = nn.Dropout(dropout)
        self.output_projection = nn.Linear(d_model, 1)
        
    def forward(self, x: torch.Tensor) -> torch.Tensor:
        """Forward pass through the model."""
        batch_size, seq_len = x.size()
        
        # Create position indices
        positions = torch.arange(seq_len, device=x.device).unsqueeze(0).expand(batch_size, -1)
        
        # Embeddings
        token_emb = self.token_embedding(x)
        pos_emb = self.position_embedding(positions)
        x = token_emb + pos_emb
        
        # Apply dropout
        x = self.dropout(x)
        
        # Transformer encoder
        x = self.transformer(x)
        
        # Global average pooling
        x = x.mean(dim=1)  # Average over sequence length
        
        # Layer normalization and output
        x = self.layer_norm(x)
        x = self.output_projection(x)
        
        return x


class MultiTaskNeuralNetwork(nn.Module):
    """Multi-task neural network for related molecular properties."""
    
    def __init__(self, input_dim: int, hidden_dims: List[int] = [512, 256, 128],
                 num_tasks: int = 3, dropout: float = 0.1, use_batch_norm: bool = True):
        """
        Initialize multi-task neural network.
        
        Args:
            input_dim: Input feature dimension
            hidden_dims: List of hidden layer dimensions
            num_tasks: Number of tasks to predict
            dropout: Dropout rate
            use_batch_norm: Whether to use batch normalization
        """
        super().__init__()
        
        if not TORCH_AVAILABLE:
            raise RuntimeError("PyTorch not available for Multi-task model")
        
        self.input_dim = input_dim
        self.hidden_dims = hidden_dims
        self.num_tasks = num_tasks
        self.dropout = dropout
        self.use_batch_norm = use_batch_norm
        
        # Shared layers
        self.shared_layers = nn.ModuleList()
        prev_dim = input_dim
        
        for hidden_dim in hidden_dims:
            self.shared_layers.append(nn.Linear(prev_dim, hidden_dim))
            if use_batch_norm:
                self.shared_layers.append(nn.BatchNorm1d(hidden_dim))
            self.shared_layers.append(nn.ReLU())
            self.shared_layers.append(nn.Dropout(dropout))
            prev_dim = hidden_dim
        
        # Task-specific heads
        self.task_heads = nn.ModuleList()
        for _ in range(num_tasks):
            self.task_heads.append(nn.Linear(prev_dim, 1))
        
        # Task names (can be customized)
        self.task_names = [
            'binding_affinity',
            'molecular_weight',
            'logp'
        ]
    
    def forward(self, x: torch.Tensor) -> List[torch.Tensor]:
        """Forward pass through the model."""
        # Shared layers
        for layer in self.shared_layers:
            x = layer(x)
        
        # Task-specific predictions
        predictions = []
        for head in self.task_heads:
            predictions.append(head(x))
        
        return predictions
    
    def predict_single_task(self, x: torch.Tensor, task_idx: int) -> torch.Tensor:
        """Predict for a single task."""
        # Shared layers
        for layer in self.shared_layers:
            x = layer(x)
        
        # Specific task head
        return self.task_heads[task_idx](x)


class DeepLearningTrainer:
    """Trainer for deep learning models."""
    
    def __init__(self, model_type: str = "cnn3d", device: str = "auto"):
        """
        Initialize deep learning trainer.
        
        Args:
            model_type: Type of model ("cnn3d", "transformer", "multitask")
            device: Device to use for training ("auto", "cpu", "cuda")
        """
        if not TORCH_AVAILABLE:
            raise RuntimeError("PyTorch not available for deep learning training")
        
        self.model_type = model_type
        self.device = self._get_device(device)
        self.model = None
        self.optimizer = None
        self.scheduler = None
        
    def _get_device(self, device: str) -> torch.device:
        """Get appropriate device for training."""
        if device == "auto":
            if torch.cuda.is_available():
                return torch.device("cuda")
            else:
                return torch.device("cpu")
        else:
            return torch.device(device)
    
    def create_model(self, **kwargs) -> nn.Module:
        """Create deep learning model based on type."""
        if self.model_type == "cnn3d":
            return MolecularCNN3D(**kwargs)
        elif self.model_type == "transformer":
            return MolecularTransformer(**kwargs)
        elif self.model_type == "multitask":
            return MultiTaskNeuralNetwork(**kwargs)
        else:
            raise ValueError(f"Unknown model type: {self.model_type}")
    
    def train(self, train_loader: DataLoader, val_loader: Optional[DataLoader] = None,
              epochs: int = 100, lr: float = 0.001, weight_decay: float = 1e-4,
              patience: int = 20, scheduler_type: str = "plateau") -> Dict[str, List[float]]:
        """
        Train deep learning model.
        
        Args:
            train_loader: Training data loader
            val_loader: Validation data loader (optional)
            epochs: Number of training epochs
            lr: Learning rate
            weight_decay: Weight decay for regularization
            patience: Early stopping patience
            scheduler_type: Type of learning rate scheduler
            
        Returns:
            Dictionary of training history
        """
        if self.model is None:
            raise ValueError("Model not created. Call create_model() first.")
        
        self.model.to(self.device)
        self.optimizer = torch.optim.Adam(self.model.parameters(), lr=lr, weight_decay=weight_decay)
        
        # Learning rate scheduler
        if scheduler_type == "plateau" and val_loader is not None:
            self.scheduler = ReduceLROnPlateau(
                self.optimizer, mode='min', factor=0.5, patience=10, verbose=True
            )
        elif scheduler_type == "cosine":
            self.scheduler = CosineAnnealingLR(self.optimizer, T_max=epochs)
        else:
            self.scheduler = None
        
        history = {'train_loss': [], 'val_loss': []}
        best_val_loss = float('inf')
        patience_counter = 0
        
        for epoch in range(epochs):
            # Training
            train_loss = self._train_epoch(train_loader)
            history['train_loss'].append(train_loss)
            
            # Validation
            if val_loader is not None:
                val_loss = self._validate_epoch(val_loader)
                history['val_loss'].append(val_loss)
                
                # Early stopping
                if val_loss < best_val_loss:
                    best_val_loss = val_loss
                    patience_counter = 0
                else:
                    patience_counter += 1
                
                if patience_counter >= patience:
                    logger.info(f"Early stopping at epoch {epoch}")
                    break
                
                # Learning rate scheduling
                if self.scheduler is not None:
                    if scheduler_type == "plateau":
                        self.scheduler.step(val_loss)
                    else:
                        self.scheduler.step()
            else:
                history['val_loss'].append(train_loss)
            
            if epoch % 10 == 0:
                logger.info(f"Epoch {epoch}: Train Loss = {train_loss:.4f}, "
                          f"Val Loss = {history['val_loss'][-1]:.4f}")
        
        return history
    
    def _train_epoch(self, train_loader: DataLoader) -> float:
        """Train for one epoch."""
        self.model.train()
        total_loss = 0.0
        
        for batch in train_loader:
            if isinstance(batch, (list, tuple)) and len(batch) == 2:
                x, y = batch
                x, y = x.to(self.device, dtype=torch.float32), y.to(self.device, dtype=torch.float32)
                
                self.optimizer.zero_grad()
                pred = self.model(x)
                
                if self.model_type == "multitask":
                    # Multi-task loss (sum of individual task losses)
                    loss = 0.0
                    for i, task_pred in enumerate(pred):
                        loss += F.mse_loss(task_pred.squeeze(), y)
                else:
                    loss = F.mse_loss(pred.squeeze(), y)
                
                loss.backward()
                self.optimizer.step()
                
                total_loss += loss.item()
        
        return total_loss / len(train_loader)
    
    def _validate_epoch(self, val_loader: DataLoader) -> float:
        """Validate for one epoch."""
        self.model.eval()
        total_loss = 0.0
        
        with torch.no_grad():
            for batch in val_loader:
                if isinstance(batch, (list, tuple)) and len(batch) == 2:
                    x, y = batch
                    x, y = x.to(self.device, dtype=torch.float32), y.to(self.device, dtype=torch.float32)
                    
                    pred = self.model(x)
                    
                    if self.model_type == "multitask":
                        # Multi-task loss (sum of individual task losses)
                        loss = 0.0
                        for i, task_pred in enumerate(pred):
                            loss += F.mse_loss(task_pred.squeeze(), y)
                    else:
                        loss = F.mse_loss(pred.squeeze(), y)
                    
                    total_loss += loss.item()
        
        return total_loss / len(val_loader)
    
    def predict(self, data_loader: DataLoader) -> np.ndarray:
        """Make predictions on data loader."""
        if self.model is None:
            raise ValueError("Model not trained. Call train() first.")
        
        self.model.eval()
        predictions = []
        
        with torch.no_grad():
            for batch in data_loader:
                if isinstance(batch, (list, tuple)) and len(batch) == 2:
                    x, _ = batch
                    x = x.to(self.device)
                    pred = self.model(x)
                    
                    if self.model_type == "multitask":
                        # For multi-task, return first task prediction
                        pred = pred[0]
                    
                    predictions.append(pred.cpu().numpy())
        
        return np.concatenate(predictions, axis=0)
    
    def save_model(self, path: str) -> None:
        """Save trained model."""
        if self.model is None:
            raise ValueError("No model to save")
        
        torch.save({
            'model_state_dict': self.model.state_dict(),
            'model_type': self.model_type,
            'optimizer_state_dict': self.optimizer.state_dict() if self.optimizer else None
        }, path)
        
        logger.info(f"Model saved to {path}")
    
    def load_model(self, path: str, **kwargs) -> None:
        """Load trained model."""
        checkpoint = torch.load(path, map_location=self.device)
        
        self.model = self.create_model(**kwargs)
        self.model.load_state_dict(checkpoint['model_state_dict'])
        self.model.to(self.device)
        
        logger.info(f"Model loaded from {path}")
```
