## Supplementary material for "Machine Learning-Driven Prediction of TLR4 Binding Affinity: A Comprehensive Molecular Feature Analysis for Drug Discovery": Codebase: deep_learning_trainer.py

```
"""
Deep learning model trainer coordinator.

This module provides comprehensive deep learning model training functionality
for TLR4 binding prediction using CNN, Transformer, and Multi-task approaches.
"""

from typing import Dict, List, Optional, Union, Any, Tuple
import pandas as pd
import numpy as np
import logging
from pathlib import Path
import joblib
from datetime import datetime

# PyTorch imports with error handling
try:
    import torch
    TORCH_AVAILABLE = True
except ImportError:
    TORCH_AVAILABLE = False
    # Define dummy torch for type hints
    class torch:
        class Tensor:
            pass

try:
    from .deep_learning_models import (
        MolecularVoxelizer, VoxelDataset, SMILESDataset, SMILESData,
        DeepLearningTrainer, MolecularCNN3D, MolecularTransformer, MultiTaskNeuralNetwork
    )
except ImportError:
    from deep_learning_models import (
        MolecularVoxelizer, VoxelDataset, SMILESDataset, SMILESData,
        DeepLearningTrainer, MolecularCNN3D, MolecularTransformer, MultiTaskNeuralNetwork
    )

logger = logging.getLogger(__name__)

# RDKit for SMILES processing
try:
    from rdkit import Chem
    from rdkit.Chem import rdMolDescriptors, Descriptors
    RDKIT_AVAILABLE = True
except ImportError:
    logger.warning("RDKit not available. SMILES processing will be limited.")
    RDKIT_AVAILABLE = False


class DeepLearningModelTrainer:
    """
    Main deep learning model trainer coordinator.
    
    Orchestrates training of CNN, Transformer, and Multi-task models
    for molecular binding prediction.
    """
    
    def __init__(self, models_dir: str = "models/deep_learning"):
        """
        Initialize deep learning model trainer.
        
        Args:
            models_dir: Directory to save trained models
        """
        self.models_dir = Path(models_dir)
        self.models_dir.mkdir(parents=True, exist_ok=True)
        
        # Initialize components
        self.voxelizer = MolecularVoxelizer()
        self.trained_models = {}
        self.training_results = {}
        
        # Model configurations
        self.model_configs = {
            'cnn3d': {
                'input_channels': 1,
                'num_classes': 1,
                'dropout': 0.1,
                'use_batch_norm': True
            },
            'transformer': {
                'vocab_size': 1000,
                'd_model': 256,
                'nhead': 8,
                'num_layers': 6,
                'dim_feedforward': 1024,
                'dropout': 0.1,
                'max_length': 128
            },
            'multitask': {
                'input_dim': 100,  # Will be set based on feature data
                'hidden_dims': [512, 256, 128],
                'num_tasks': 3,
                'dropout': 0.1,
                'use_batch_norm': True
            }
        }
    
    def prepare_voxel_data(self, pdbqt_files: List[str], 
                          binding_affinities: Optional[List[float]] = None) -> VoxelDataset:
        """
        Prepare voxel dataset from PDBQT files.
        
        Args:
            pdbqt_files: List of PDBQT file paths
            binding_affinities: Optional list of binding affinities
            
        Returns:
            VoxelDataset object
        """
        logger.info(f"Preparing voxel data from {len(pdbqt_files)} PDBQT files")
        
        voxel_data = []
        targets = binding_affinities if binding_affinities is not None else [0.0] * len(pdbqt_files)
        
        for i, pdbqt_file in enumerate(pdbqt_files):
            voxel_data_item = self.voxelizer.voxelize_pdbqt(pdbqt_file)
            if voxel_data_item is not None:
                # Set binding affinity
                voxel_data_item = voxel_data_item._replace(binding_affinity=targets[i])
                voxel_data.append(voxel_data_item)
            else:
                logger.warning(f"Failed to voxelize {pdbqt_file}")
        
        logger.info(f"Successfully prepared {len(voxel_data)} voxel samples")
        return VoxelDataset(voxel_data)
    
    def prepare_smiles_data(self, smiles_list: List[str], compound_names: List[str],
                           binding_affinities: Optional[List[float]] = None,
                           vocab_size: int = 1000, max_length: int = 128) -> SMILESDataset:
        """
        Prepare SMILES dataset from SMILES strings.
        
        Args:
            smiles_list: List of SMILES strings
            compound_names: List of compound names
            binding_affinities: Optional list of binding affinities
            vocab_size: Vocabulary size for tokenization
            max_length: Maximum sequence length
            
        Returns:
            SMILESDataset object
        """
        logger.info(f"Preparing SMILES data from {len(smiles_list)} SMILES strings")
        
        targets = binding_affinities if binding_affinities is not None else [0.0] * len(smiles_list)
        
        smiles_data = []
        for smiles, name, affinity in zip(smiles_list, compound_names, targets):
            smiles_data.append(SMILESData(
                smiles=smiles,
                compound_name=name,
                binding_affinity=affinity
            ))
        
        logger.info(f"Successfully prepared {len(smiles_data)} SMILES samples")
        return SMILESDataset(smiles_data, vocab_size=vocab_size, max_length=max_length)
    
    def prepare_multitask_data(self, features_df: pd.DataFrame, 
                              binding_affinities: pd.Series,
                              additional_targets: Optional[Dict[str, pd.Series]] = None) -> Tuple[torch.Tensor, List[torch.Tensor]]:
        """
        Prepare multi-task data from feature DataFrame.
        
        Args:
            features_df: DataFrame with molecular features
            binding_affinities: Series with binding affinities
            additional_targets: Optional dict with additional target properties
            
        Returns:
            Tuple of (features_tensor, targets_list)
        """
        logger.info("Preparing multi-task data from feature DataFrame")
        
        # Convert features to tensor
        features_tensor = torch.tensor(features_df.values, dtype=torch.float)
        
        # Prepare targets
        targets = [binding_affinities.values]
        
        if additional_targets is not None:
            for target_name, target_values in additional_targets.items():
                targets.append(target_values.values)
        
        # Convert to tensors
        targets_tensors = [torch.tensor(target, dtype=torch.float) for target in targets]
        
        logger.info(f"Prepared multi-task data with {len(targets_tensors)} tasks")
        return features_tensor, targets_tensors
    
    def train_cnn3d_model(self, train_dataset: VoxelDataset, val_dataset: Optional[VoxelDataset] = None,
                         **kwargs) -> Dict[str, Any]:
        """
        Train 3D CNN model on voxel data.
        
        Args:
            train_dataset: Training voxel dataset
            val_dataset: Validation voxel dataset (optional)
            **kwargs: Additional training parameters
            
        Returns:
            Dictionary with training results
        """
        logger.info("Training 3D CNN model on voxel data")
        
        # Create trainer
        trainer = DeepLearningTrainer(model_type="cnn3d", device=kwargs.get('device', 'auto'))
        trainer.model = trainer.create_model(**self.model_configs['cnn3d'])
        
        # Create data loaders
        train_loader = DataLoader(train_dataset, batch_size=kwargs.get('batch_size', 16), 
                                 shuffle=True, num_workers=0)
        
        val_loader = None
        if val_dataset is not None:
            val_loader = DataLoader(val_dataset, batch_size=kwargs.get('batch_size', 16), 
                                   shuffle=False, num_workers=0)
        
        # Train model
        history = trainer.train(train_loader, val_loader, **kwargs)
        
        # Store results
        self.trained_models['cnn3d'] = trainer
        self.training_results['cnn3d'] = {
            'trainer': trainer,
            'history': history,
            'model_type': 'cnn3d'
        }
        
        # Save model
        model_path = self.models_dir / "cnn3d_model.pth"
        trainer.save_model(str(model_path))
        
        logger.info("3D CNN model training completed successfully")
        return self.training_results['cnn3d']
    
    def train_transformer_model(self, train_dataset: SMILESDataset, 
                               val_dataset: Optional[SMILESDataset] = None,
                               **kwargs) -> Dict[str, Any]:
        """
        Train Transformer model on SMILES data.
        
        Args:
            train_dataset: Training SMILES dataset
            val_dataset: Validation SMILES dataset (optional)
            **kwargs: Additional training parameters
            
        Returns:
            Dictionary with training results
        """
        logger.info("Training Transformer model on SMILES data")
        
        # Create trainer
        trainer = DeepLearningTrainer(model_type="transformer", device=kwargs.get('device', 'auto'))
        trainer.model = trainer.create_model(**self.model_configs['transformer'])
        
        # Create data loaders
        train_loader = DataLoader(train_dataset, batch_size=kwargs.get('batch_size', 32), 
                                 shuffle=True, num_workers=0)
        
        val_loader = None
        if val_dataset is not None:
            val_loader = DataLoader(val_dataset, batch_size=kwargs.get('batch_size', 32), 
                                   shuffle=False, num_workers=0)
        
        # Train model
        history = trainer.train(train_loader, val_loader, **kwargs)
        
        # Store results
        self.trained_models['transformer'] = trainer
        self.training_results['transformer'] = {
            'trainer': trainer,
            'history': history,
            'model_type': 'transformer'
        }
        
        # Save model
        model_path = self.models_dir / "transformer_model.pth"
        trainer.save_model(str(model_path))
        
        logger.info("Transformer model training completed successfully")
        return self.training_results['transformer']
    
    def train_multitask_model(self, features_tensor: torch.Tensor, targets_tensors: List[torch.Tensor],
                             train_indices: List[int], val_indices: Optional[List[int]] = None,
                             **kwargs) -> Dict[str, Any]:
        """
        Train Multi-task model on feature data.
        
        Args:
            features_tensor: Feature tensor
            targets_tensors: List of target tensors
            train_indices: Training indices
            val_indices: Validation indices (optional)
            **kwargs: Additional training parameters
            
        Returns:
            Dictionary with training results
        """
        logger.info("Training Multi-task model on feature data")
        
        # Update input dimension in config
        self.model_configs['multitask']['input_dim'] = features_tensor.shape[1]
        self.model_configs['multitask']['num_tasks'] = len(targets_tensors)
        
        # Create trainer
        trainer = DeepLearningTrainer(model_type="multitask", device=kwargs.get('device', 'auto'))
        trainer.model = trainer.create_model(**self.model_configs['multitask'])
        
        # Prepare training data
        train_features = features_tensor[train_indices]
        train_targets = [target[train_indices] for target in targets_tensors]
        
        # Create custom dataset for multi-task
        class MultiTaskDataset(Dataset):
            def __init__(self, features, targets):
                self.features = features
                self.targets = targets[0]  # Use first target for DataLoader compatibility
            
            def __len__(self):
                return len(self.features)
            
            def __getitem__(self, idx):
                return self.features[idx], self.targets[idx]
        
        train_dataset = MultiTaskDataset(train_features, train_targets)
        train_loader = DataLoader(train_dataset, batch_size=kwargs.get('batch_size', 32), 
                                 shuffle=True, num_workers=0)
        
        # Prepare validation data if provided
        val_loader = None
        if val_indices is not None:
            val_features = features_tensor[val_indices]
            val_targets = [target[val_indices] for target in targets_tensors]
            val_dataset = MultiTaskDataset(val_features, val_targets)
            val_loader = DataLoader(val_dataset, batch_size=kwargs.get('batch_size', 32), 
                                   shuffle=False, num_workers=0)
        
        # Train model
        history = trainer.train(train_loader, val_loader, **kwargs)
        
        # Store results
        self.trained_models['multitask'] = trainer
        self.training_results['multitask'] = {
            'trainer': trainer,
            'history': history,
            'model_type': 'multitask'
        }
        
        # Save model
        model_path = self.models_dir / "multitask_model.pth"
        trainer.save_model(str(model_path))
        
        logger.info("Multi-task model training completed successfully")
        return self.training_results['multitask']
    
    def train_all_models(self, pdbqt_files: List[str], binding_affinities: List[float],
                        train_indices: List[int], features_df: Optional[pd.DataFrame] = None,
                        smiles_list: Optional[List[str]] = None,
                        compound_names: Optional[List[str]] = None,
                        val_indices: Optional[List[int]] = None,
                        model_types: List[str] = None, **kwargs) -> Dict[str, Any]:
        """
        Train all deep learning models.
        
        Args:
            pdbqt_files: List of PDBQT file paths
            binding_affinities: List of binding affinities
            features_df: Optional DataFrame with molecular features
            smiles_list: Optional list of SMILES strings
            compound_names: Optional list of compound names
            train_indices: Training indices
            val_indices: Validation indices (optional)
            model_types: List of model types to train
            **kwargs: Additional training parameters
            
        Returns:
            Dictionary of training results
        """
        if model_types is None:
            model_types = ['cnn3d', 'transformer', 'multitask']
        
        logger.info(f"Training {len(model_types)} deep learning models")
        
        results = {}
        
        # Train CNN3D model
        if 'cnn3d' in model_types:
            try:
                # Prepare voxel data
                train_pdbqt_files = [pdbqt_files[i] for i in train_indices]
                train_affinities = [binding_affinities[i] for i in train_indices]
                train_voxel_dataset = self.prepare_voxel_data(train_pdbqt_files, train_affinities)
                
                val_voxel_dataset = None
                if val_indices is not None:
                    val_pdbqt_files = [pdbqt_files[i] for i in val_indices]
                    val_affinities = [binding_affinities[i] for i in val_indices]
                    val_voxel_dataset = self.prepare_voxel_data(val_pdbqt_files, val_affinities)
                
                results['cnn3d'] = self.train_cnn3d_model(train_voxel_dataset, val_voxel_dataset, **kwargs)
                
            except Exception as e:
                logger.error(f"Error training CNN3D model: {str(e)}")
                results['cnn3d'] = None
        
        # Train Transformer model
        if 'transformer' in model_types and smiles_list is not None and compound_names is not None:
            try:
                # Prepare SMILES data
                train_smiles = [smiles_list[i] for i in train_indices]
                train_names = [compound_names[i] for i in train_indices]
                train_affinities = [binding_affinities[i] for i in train_indices]
                train_smiles_dataset = self.prepare_smiles_data(train_smiles, train_names, train_affinities)
                
                val_smiles_dataset = None
                if val_indices is not None:
                    val_smiles = [smiles_list[i] for i in val_indices]
                    val_names = [compound_names[i] for i in val_indices]
                    val_affinities = [binding_affinities[i] for i in val_indices]
                    val_smiles_dataset = self.prepare_smiles_data(val_smiles, val_names, val_affinities)
                
                results['transformer'] = self.train_transformer_model(train_smiles_dataset, val_smiles_dataset, **kwargs)
                
            except Exception as e:
                logger.error(f"Error training Transformer model: {str(e)}")
                results['transformer'] = None
        
        # Train Multi-task model
        if 'multitask' in model_types and features_df is not None:
            try:
                # Prepare multi-task data
                features_tensor, targets_tensors = self.prepare_multitask_data(
                    features_df, pd.Series(binding_affinities)
                )
                
                results['multitask'] = self.train_multitask_model(
                    features_tensor, targets_tensors, train_indices, val_indices, **kwargs
                )
                
            except Exception as e:
                logger.error(f"Error training Multi-task model: {str(e)}")
                results['multitask'] = None
        
        logger.info(f"Successfully trained {len([r for r in results.values() if r is not None])} deep learning models")
        return results
    
    def evaluate_models(self, test_pdbqt_files: List[str], test_binding_affinities: List[float],
                       test_indices: List[int], test_features_df: Optional[pd.DataFrame] = None,
                       test_smiles_list: Optional[List[str]] = None,
                       test_compound_names: Optional[List[str]] = None) -> Dict[str, Dict]:
        """
        Evaluate all trained models on test set.
        
        Args:
            test_pdbqt_files: List of test PDBQT file paths
            test_binding_affinities: List of test binding affinities
            test_features_df: Optional DataFrame with test molecular features
            test_smiles_list: Optional list of test SMILES strings
            test_compound_names: Optional list of test compound names
            test_indices: Test indices
            
        Returns:
            Dictionary of evaluation results
        """
        logger.info("Evaluating deep learning models on test set")
        
        evaluation_results = {}
        
        # Evaluate CNN3D model
        if 'cnn3d' in self.trained_models:
            try:
                test_pdbqt_files_subset = [test_pdbqt_files[i] for i in test_indices]
                test_affinities_subset = [test_binding_affinities[i] for i in test_indices]
                test_voxel_dataset = self.prepare_voxel_data(test_pdbqt_files_subset, test_affinities_subset)
                
                test_loader = DataLoader(test_voxel_dataset, batch_size=16, shuffle=False, num_workers=0)
                trainer = self.trained_models['cnn3d']
                predictions = trainer.predict(test_loader)
                
                metrics = self._calculate_metrics(test_affinities_subset, predictions.flatten())
                evaluation_results['cnn3d'] = {
                    'predictions': predictions,
                    'metrics': metrics
                }
                
            except Exception as e:
                logger.error(f"Error evaluating CNN3D model: {str(e)}")
                evaluation_results['cnn3d'] = None
        
        # Evaluate Transformer model
        if 'transformer' in self.trained_models and test_smiles_list is not None:
            try:
                test_smiles_subset = [test_smiles_list[i] for i in test_indices]
                test_names_subset = [test_compound_names[i] for i in test_indices]
                test_affinities_subset = [test_binding_affinities[i] for i in test_indices]
                test_smiles_dataset = self.prepare_smiles_data(test_smiles_subset, test_names_subset, test_affinities_subset)
                
                test_loader = DataLoader(test_smiles_dataset, batch_size=32, shuffle=False, num_workers=0)
                trainer = self.trained_models['transformer']
                predictions = trainer.predict(test_loader)
                
                metrics = self._calculate_metrics(test_affinities_subset, predictions.flatten())
                evaluation_results['transformer'] = {
                    'predictions': predictions,
                    'metrics': metrics
                }
                
            except Exception as e:
                logger.error(f"Error evaluating Transformer model: {str(e)}")
                evaluation_results['transformer'] = None
        
        # Evaluate Multi-task model
        if 'multitask' in self.trained_models and test_features_df is not None:
            try:
                test_features_subset = test_features_df.iloc[test_indices]
                test_affinities_subset = [test_binding_affinities[i] for i in test_indices]
                
                # Create test dataset
                test_features_tensor = torch.tensor(test_features_subset.values, dtype=torch.float)
                test_targets = torch.tensor(test_affinities_subset, dtype=torch.float)
                
                class TestDataset(Dataset):
                    def __init__(self, features, targets):
                        self.features = features
                        self.targets = targets
                    
                    def __len__(self):
                        return len(self.features)
                    
                    def __getitem__(self, idx):
                        return self.features[idx], self.targets[idx]
                
                test_dataset = TestDataset(test_features_tensor, test_targets)
                test_loader = DataLoader(test_dataset, batch_size=32, shuffle=False, num_workers=0)
                
                trainer = self.trained_models['multitask']
                predictions = trainer.predict(test_loader)
                
                metrics = self._calculate_metrics(test_affinities_subset, predictions.flatten())
                evaluation_results['multitask'] = {
                    'predictions': predictions,
                    'metrics': metrics
                }
                
            except Exception as e:
                logger.error(f"Error evaluating Multi-task model: {str(e)}")
                evaluation_results['multitask'] = None
        
        logger.info(f"Successfully evaluated {len([r for r in evaluation_results.values() if r is not None])} deep learning models")
        return evaluation_results
    
    def _calculate_metrics(self, y_true: List[float], y_pred: np.ndarray) -> Dict[str, float]:
        """Calculate regression metrics."""
        y_true = np.array(y_true)
        y_pred = np.array(y_pred)
        
        mse = np.mean((y_true - y_pred) ** 2)
        rmse = np.sqrt(mse)
        mae = np.mean(np.abs(y_true - y_pred))
        
        # R² score
        ss_res = np.sum((y_true - y_pred) ** 2)
        ss_tot = np.sum((y_true - np.mean(y_true)) ** 2)
        r2 = 1 - (ss_res / ss_tot) if ss_tot != 0 else 0
        
        return {
            'mse': mse,
            'rmse': rmse,
            'mae': mae,
            'r2': r2
        }
    
    def get_best_model(self, metric: str = 'r2') -> Tuple[str, Any]:
        """
        Get the best performing model based on specified metric.
        
        Args:
            metric: Metric to use for model selection
            
        Returns:
            Tuple of (model_name, trainer)
        """
        if not self.training_results:
            raise ValueError("No models have been trained yet")
        
        best_model_name = None
        best_score = float('-inf') if metric == 'r2' else float('inf')
        
        for model_name, results in self.training_results.items():
            if results is None:
                continue
                
            # Get validation score from history
            if 'val_loss' in results['history']:
                val_losses = results['history']['val_loss']
                if val_losses:
                    score = -min(val_losses)  # Convert loss to score
                    if metric == 'r2':
                        if score > best_score:
                            best_score = score
                            best_model_name = model_name
                    else:  # For loss-based metrics
                        if score < best_score:
                            best_score = score
                            best_model_name = model_name
        
        if best_model_name is None:
            raise ValueError(f"No model found with metric {metric}")
        
        return best_model_name, self.trained_models[best_model_name]
    
    def get_model_summary(self) -> pd.DataFrame:
        """Get summary of all trained models."""
        if not self.training_results:
            return pd.DataFrame()
        
        summary_data = []
        for model_name, results in self.training_results.items():
            if results is None:
                continue
                
            model_type = "Deep Learning"
            summary_data.append({
                'model_name': model_name,
                'model_type': model_type,
                'final_train_loss': results['history']['train_loss'][-1] if results['history']['train_loss'] else 0.0,
                'final_val_loss': results['history']['val_loss'][-1] if results['history']['val_loss'] else 0.0,
                'best_val_loss': min(results['history']['val_loss']) if results['history']['val_loss'] else 0.0
            })
        
        return pd.DataFrame(summary_data)
```
