## Supplementary material for "Machine Learning-Driven Prediction of TLR4 Binding Affinity: A Comprehensive Molecular Feature Analysis for Drug Discovery": Codebase: ensemble_models.py

```
"""
Ensemble and hybrid models for TLR4 binding prediction.

This module implements various ensemble methods including stacked ensembles,
weighted ensembles, and physics-informed neural networks for improved
binding affinity prediction accuracy and uncertainty quantification.
"""

from abc import ABC, abstractmethod
from typing import Dict, List, Optional, Union, Any, Tuple, NamedTuple
import pandas as pd
import numpy as np
import logging
from pathlib import Path
import joblib
from datetime import datetime
from dataclasses import dataclass

logger = logging.getLogger(__name__)

# ML library imports with error handling
try:
    from sklearn.ensemble import VotingRegressor, StackingRegressor
    from sklearn.linear_model import LinearRegression, Ridge
    from sklearn.model_selection import cross_val_score, KFold
    from sklearn.metrics import mean_squared_error, r2_score, mean_absolute_error
    from sklearn.preprocessing import StandardScaler
    SKLEARN_AVAILABLE = True
except ImportError:
    logger.warning("scikit-learn not available. Ensemble models will be limited.")
    SKLEARN_AVAILABLE = False

try:
    import torch
    import torch.nn as nn
    import torch.nn.functional as F
    from torch.utils.data import Dataset, DataLoader
    TORCH_AVAILABLE = True
except ImportError:
    logger.warning("PyTorch not available. Physics-informed models will be limited.")
    TORCH_AVAILABLE = False

# Import base models
try:
    from .trainer import MLModelTrainer, ModelTrainerInterface
    from .deep_learning_trainer import DeepLearningModelTrainer
    from .gnn_models import GNNModelTrainer
except ImportError:
    logger.warning("Base model trainers not available. Ensemble models will be limited.")


@dataclass
class EnsemblePrediction:
    """Container for ensemble prediction results."""
    prediction: float
    uncertainty: float
    individual_predictions: Dict[str, float]
    weights: Dict[str, float]
    confidence_interval: Tuple[float, float]


class EnsembleModelInterface(ABC):
    """Abstract interface for ensemble models."""
    
    @abstractmethod
    def fit(self, X_train: pd.DataFrame, y_train: pd.Series, 
            X_val: Optional[pd.DataFrame] = None, 
            y_val: Optional[pd.Series] = None) -> None:
        """Fit the ensemble model."""
        pass
    
    @abstractmethod
    def predict(self, X: pd.DataFrame) -> np.ndarray:
        """Make predictions using the ensemble model."""
        pass
    
    @abstractmethod
    def predict_with_uncertainty(self, X: pd.DataFrame) -> List[EnsemblePrediction]:
        """Make predictions with uncertainty quantification."""
        pass


class StackedEnsemble(EnsembleModelInterface):
    """
    Stacked ensemble model that combines base models using a meta-learner.
    
    Uses cross-validation to generate out-of-fold predictions for training
    the meta-learner, preventing overfitting.
    """
    
    def __init__(self, base_models: Dict[str, Any], meta_learner: Any = None, 
                 cv_folds: int = 5, random_state: int = 42):
        """
        Initialize stacked ensemble.
        
        Args:
            base_models: Dictionary of base model trainers
            meta_learner: Meta-learner for combining base model predictions
            cv_folds: Number of cross-validation folds
            random_state: Random state for reproducibility
        """
        self.base_models = base_models
        self.meta_learner = meta_learner or Ridge(alpha=1.0)
        self.cv_folds = cv_folds
        self.random_state = random_state
        self.trained_base_models = {}
        self.meta_learner_trained = None
        self.feature_names = None
        
    def fit(self, X_train: pd.DataFrame, y_train: pd.Series, 
            X_val: Optional[pd.DataFrame] = None, 
            y_val: Optional[pd.Series] = None) -> None:
        """Fit the stacked ensemble model."""
        if not SKLEARN_AVAILABLE:
            raise RuntimeError("scikit-learn not available for stacked ensemble")
        
        logger.info("Training stacked ensemble model")
        
        # Store feature names
        self.feature_names = X_train.columns.tolist()
        
        # Generate out-of-fold predictions for meta-learner training
        kf = KFold(n_splits=self.cv_folds, shuffle=True, random_state=self.random_state)
        meta_features = np.zeros((len(X_train), len(self.base_models)))
        
        for fold, (train_idx, val_idx) in enumerate(kf.split(X_train)):
            X_fold_train = X_train.iloc[train_idx]
            y_fold_train = y_train.iloc[train_idx]
            X_fold_val = X_train.iloc[val_idx]
            
            # Train base models on fold
            for i, (model_name, trainer) in enumerate(self.base_models.items()):
                try:
                    # Train model on fold
                    model = trainer.train(X_fold_train, y_fold_train)
                    
                    # Make predictions on validation fold
                    fold_predictions = trainer.predict(model, X_fold_val)
                    meta_features[val_idx, i] = fold_predictions
                    
                    # Store the final model trained on full data
                    if fold == self.cv_folds - 1:
                        self.trained_base_models[model_name] = trainer.train(X_train, y_train)
                        
                except Exception as e:
                    logger.error(f"Error training {model_name} in fold {fold}: {str(e)}")
                    meta_features[val_idx, i] = np.mean(y_train)
        
        # Train meta-learner on out-of-fold predictions
        self.meta_learner_trained = self.meta_learner.fit(meta_features, y_train)
        
        logger.info("Stacked ensemble training completed successfully")
    
    def predict(self, X: pd.DataFrame) -> np.ndarray:
        """Make predictions using the stacked ensemble model."""
        if self.meta_learner_trained is None:
            raise ValueError("Model must be fitted before making predictions")
        
        # Generate base model predictions
        base_predictions = np.zeros((len(X), len(self.base_models)))
        
        for i, (model_name, model) in enumerate(self.trained_base_models.items()):
            try:
                trainer = self.base_models[model_name]
                predictions = trainer.predict(model, X)
                base_predictions[:, i] = predictions
            except Exception as e:
                logger.error(f"Error making predictions with {model_name}: {str(e)}")
                base_predictions[:, i] = 0.0
        
        # Combine predictions using meta-learner
        ensemble_predictions = self.meta_learner_trained.predict(base_predictions)
        return ensemble_predictions
    
    def predict_with_uncertainty(self, X: pd.DataFrame) -> List[EnsemblePrediction]:
        """Make predictions with uncertainty quantification."""
        predictions = self.predict(X)
        
        # Calculate uncertainty based on base model variance
        base_predictions = np.zeros((len(X), len(self.base_models)))
        
        for i, (model_name, model) in enumerate(self.trained_base_models.items()):
            try:
                trainer = self.base_models[model_name]
                pred = trainer.predict(model, X)
                base_predictions[:, i] = pred
            except Exception as e:
                logger.error(f"Error making predictions with {model_name}: {str(e)}")
                base_predictions[:, i] = 0.0
        
        # Calculate uncertainty as standard deviation of base model predictions
        uncertainties = np.std(base_predictions, axis=1)
        
        # Create individual predictions dictionary
        individual_predictions = {}
        for i, (model_name, _) in enumerate(self.trained_base_models.items()):
            individual_predictions[model_name] = base_predictions[:, i].tolist()
        
        # Create ensemble predictions
        ensemble_predictions = []
        for i in range(len(X)):
            # Calculate confidence interval (95%)
            ci_lower = predictions[i] - 1.96 * uncertainties[i]
            ci_upper = predictions[i] + 1.96 * uncertainties[i]
            
            ensemble_predictions.append(EnsemblePrediction(
                prediction=float(predictions[i]),
                uncertainty=float(uncertainties[i]),
                individual_predictions={name: preds[i] for name, preds in individual_predictions.items()},
                weights={name: 1.0/len(self.base_models) for name in self.trained_base_models.keys()},
                confidence_interval=(ci_lower, ci_upper)
            ))
        
        return ensemble_predictions


class WeightedEnsemble(EnsembleModelInterface):
    """
    Weighted ensemble model that combines base models using learned weights.
    
    Weights are determined using cross-validation performance on validation set.
    """
    
    def __init__(self, base_models: Dict[str, Any], weight_method: str = 'performance',
                 cv_folds: int = 5, random_state: int = 42):
        """
        Initialize weighted ensemble.
        
        Args:
            base_models: Dictionary of base model trainers
            weight_method: Method for calculating weights ('performance', 'inverse_mse', 'r2')
            cv_folds: Number of cross-validation folds
            random_state: Random state for reproducibility
        """
        self.base_models = base_models
        self.weight_method = weight_method
        self.cv_folds = cv_folds
        self.random_state = random_state
        self.trained_base_models = {}
        self.weights = {}
        self.feature_names = None
        
    def fit(self, X_train: pd.DataFrame, y_train: pd.Series, 
            X_val: Optional[pd.DataFrame] = None, 
            y_val: Optional[pd.Series] = None) -> None:
        """Fit the weighted ensemble model."""
        logger.info("Training weighted ensemble model")
        
        # Store feature names
        self.feature_names = X_train.columns.tolist()
        
        # Train base models
        for model_name, trainer in self.base_models.items():
            try:
                logger.info(f"Training base model: {model_name}")
                model = trainer.train(X_train, y_train, X_val, y_val)
                self.trained_base_models[model_name] = model
            except Exception as e:
                logger.error(f"Error training {model_name}: {str(e)}")
                continue
        
        # Calculate weights based on validation performance
        if X_val is not None and y_val is not None:
            self._calculate_weights_from_validation(X_val, y_val)
        else:
            # Use cross-validation to calculate weights
            self._calculate_weights_from_cv(X_train, y_train)
        
        logger.info(f"Weighted ensemble weights: {self.weights}")
        logger.info("Weighted ensemble training completed successfully")
    
    def _calculate_weights_from_validation(self, X_val: pd.DataFrame, y_val: pd.Series) -> None:
        """Calculate weights based on validation set performance."""
        model_scores = {}
        
        for model_name, model in self.trained_base_models.items():
            try:
                trainer = self.base_models[model_name]
                predictions = trainer.predict(model, X_val)
                
                if self.weight_method == 'performance':
                    # Use R² score
                    r2 = r2_score(y_val, predictions)
                    model_scores[model_name] = max(0, r2)  # Ensure non-negative
                elif self.weight_method == 'inverse_mse':
                    # Use inverse MSE
                    mse = mean_squared_error(y_val, predictions)
                    model_scores[model_name] = 1.0 / (1.0 + mse)
                elif self.weight_method == 'r2':
                    # Use R² score directly
                    r2 = r2_score(y_val, predictions)
                    model_scores[model_name] = max(0, r2)
                else:
                    raise ValueError(f"Unknown weight method: {self.weight_method}")
                    
            except Exception as e:
                logger.error(f"Error calculating weights for {model_name}: {str(e)}")
                model_scores[model_name] = 0.0
        
        # Normalize weights
        total_score = sum(model_scores.values())
        if total_score > 0:
            self.weights = {name: score / total_score for name, score in model_scores.items()}
        else:
            # Equal weights if all models failed
            self.weights = {name: 1.0 / len(self.trained_base_models) 
                          for name in self.trained_base_models.keys()}
    
    def _calculate_weights_from_cv(self, X_train: pd.DataFrame, y_train: pd.Series) -> None:
        """Calculate weights using cross-validation."""
        if not SKLEARN_AVAILABLE:
            # Equal weights if sklearn not available
            self.weights = {name: 1.0 / len(self.trained_base_models) 
                          for name in self.trained_base_models.keys()}
            return
        
        kf = KFold(n_splits=self.cv_folds, shuffle=True, random_state=self.random_state)
        model_scores = {name: [] for name in self.trained_base_models.keys()}
        
        for train_idx, val_idx in kf.split(X_train):
            X_fold_train = X_train.iloc[train_idx]
            y_fold_train = y_train.iloc[train_idx]
            X_fold_val = X_train.iloc[val_idx]
            y_fold_val = y_train.iloc[val_idx]
            
            for model_name, model in self.trained_base_models.items():
                try:
                    # Retrain model on fold
                    trainer = self.base_models[model_name]
                    fold_model = trainer.train(X_fold_train, y_fold_train)
                    
                    # Evaluate on validation fold
                    predictions = trainer.predict(fold_model, X_fold_val)
                    
                    if self.weight_method == 'performance':
                        r2 = r2_score(y_fold_val, predictions)
                        model_scores[model_name].append(max(0, r2))
                    elif self.weight_method == 'inverse_mse':
                        mse = mean_squared_error(y_fold_val, predictions)
                        model_scores[model_name].append(1.0 / (1.0 + mse))
                    elif self.weight_method == 'r2':
                        r2 = r2_score(y_fold_val, predictions)
                        model_scores[model_name].append(max(0, r2))
                        
                except Exception as e:
                    logger.error(f"Error in CV fold for {model_name}: {str(e)}")
                    model_scores[model_name].append(0.0)
        
        # Calculate average scores and normalize weights
        avg_scores = {name: np.mean(scores) for name, scores in model_scores.items()}
        total_score = sum(avg_scores.values())
        
        if total_score > 0:
            self.weights = {name: score / total_score for name, score in avg_scores.items()}
        else:
            self.weights = {name: 1.0 / len(self.trained_base_models) 
                          for name in self.trained_base_models.keys()}
    
    def predict(self, X: pd.DataFrame) -> np.ndarray:
        """Make predictions using the weighted ensemble model."""
        if not self.trained_base_models:
            raise ValueError("Model must be fitted before making predictions")
        
        # Generate base model predictions
        base_predictions = {}
        for model_name, model in self.trained_base_models.items():
            try:
                trainer = self.base_models[model_name]
                predictions = trainer.predict(model, X)
                base_predictions[model_name] = predictions
            except Exception as e:
                logger.error(f"Error making predictions with {model_name}: {str(e)}")
                base_predictions[model_name] = np.zeros(len(X))
        
        # Combine predictions using weights
        ensemble_predictions = np.zeros(len(X))
        for model_name, predictions in base_predictions.items():
            weight = self.weights.get(model_name, 0.0)
            ensemble_predictions += weight * predictions
        
        return ensemble_predictions
    
    def predict_with_uncertainty(self, X: pd.DataFrame) -> List[EnsemblePrediction]:
        """Make predictions with uncertainty quantification."""
        predictions = self.predict(X)
        
        # Generate base model predictions
        base_predictions = {}
        for model_name, model in self.trained_base_models.items():
            try:
                trainer = self.base_models[model_name]
                pred = trainer.predict(model, X)
                base_predictions[model_name] = pred
            except Exception as e:
                logger.error(f"Error making predictions with {model_name}: {str(e)}")
                base_predictions[model_name] = np.zeros(len(X))
        
        # Calculate uncertainty as weighted standard deviation
        uncertainties = np.zeros(len(X))
        for i in range(len(X)):
            weighted_var = 0.0
            for model_name, pred in base_predictions.items():
                weight = self.weights.get(model_name, 0.0)
                diff = pred[i] - predictions[i]
                weighted_var += weight * (diff ** 2)
            uncertainties[i] = np.sqrt(weighted_var)
        
        # Create ensemble predictions
        ensemble_predictions = []
        for i in range(len(X)):
            # Calculate confidence interval (95%)
            ci_lower = predictions[i] - 1.96 * uncertainties[i]
            ci_upper = predictions[i] + 1.96 * uncertainties[i]
            
            ensemble_predictions.append(EnsemblePrediction(
                prediction=float(predictions[i]),
                uncertainty=float(uncertainties[i]),
                individual_predictions={name: pred[i] for name, pred in base_predictions.items()},
                weights=self.weights.copy(),
                confidence_interval=(ci_lower, ci_upper)
            ))
        
        return ensemble_predictions


class PhysicsInformedNeuralNetwork(nn.Module):
    """
    Physics-informed neural network incorporating thermodynamic constraints.
    
    This model incorporates physical laws and thermodynamic principles
    to improve binding affinity prediction accuracy.
    """
    
    def __init__(self, input_dim: int, hidden_dims: List[int] = [256, 128, 64],
                 dropout: float = 0.1, use_thermodynamic_constraints: bool = True):
        """
        Initialize physics-informed neural network.
        
        Args:
            input_dim: Input feature dimension
            hidden_dims: List of hidden layer dimensions
            dropout: Dropout rate
            use_thermodynamic_constraints: Whether to use thermodynamic constraints
        """
        super().__init__()
        
        if not TORCH_AVAILABLE:
            raise RuntimeError("PyTorch not available for physics-informed neural network")
        
        self.input_dim = input_dim
        self.hidden_dims = hidden_dims
        self.dropout = dropout
        self.use_thermodynamic_constraints = use_thermodynamic_constraints
        
        # Build network layers
        layers = []
        prev_dim = input_dim
        
        for hidden_dim in hidden_dims:
            layers.extend([
                nn.Linear(prev_dim, hidden_dim),
                nn.BatchNorm1d(hidden_dim),
                nn.ReLU(),
                nn.Dropout(dropout)
            ])
            prev_dim = hidden_dim
        
        # Output layer
        layers.append(nn.Linear(prev_dim, 1))
        
        self.network = nn.Sequential(*layers)
        
        # Thermodynamic constraint parameters
        if use_thermodynamic_constraints:
            # Physical constants for binding thermodynamics
            self.register_buffer('gas_constant', torch.tensor(0.001987))  # kcal/(mol·K)
            self.register_buffer('temperature', torch.tensor(298.15))  # K
            self.register_buffer('min_binding_affinity', torch.tensor(-20.0))  # kcal/mol
            self.register_buffer('max_binding_affinity', torch.tensor(0.0))  # kcal/mol
    
    def forward(self, x: torch.Tensor) -> torch.Tensor:
        """Forward pass through the network."""
        # Standard forward pass
        output = self.network(x)
        
        # Apply thermodynamic constraints if enabled
        if self.use_thermodynamic_constraints:
            # Constrain binding affinity to physically reasonable range
            output = torch.clamp(output, 
                               self.min_binding_affinity, 
                               self.max_binding_affinity)
        
        return output
    
    def physics_loss(self, predictions: torch.Tensor, features: torch.Tensor) -> torch.Tensor:
        """
        Calculate physics-informed loss incorporating thermodynamic constraints.
        
        Args:
            predictions: Model predictions
            features: Input features
            
        Returns:
            Physics loss term
        """
        if not self.use_thermodynamic_constraints:
            return torch.tensor(0.0, device=predictions.device)
        
        physics_loss = 0.0
        
        # Constraint 1: Binding affinity should be negative (favorable binding)
        # Penalize positive binding affinities
        positive_penalty = F.relu(predictions)
        physics_loss += torch.mean(positive_penalty)
        
        # Constraint 2: Molecular weight vs binding affinity relationship
        # Heavier molecules typically have stronger binding (more negative affinity)
        if 'molecular_weight' in features.columns:
            mw_idx = features.columns.get_loc('molecular_weight')
            molecular_weights = features[:, mw_idx]
            
            # Expected relationship: more negative affinity for heavier molecules
            expected_relationship = -molecular_weights / 1000.0  # Scale factor
            mw_loss = F.mse_loss(predictions, expected_relationship)
            physics_loss += 0.1 * mw_loss
        
        # Constraint 3: LogP vs binding affinity relationship
        # More lipophilic molecules may have different binding characteristics
        if 'logp' in features.columns:
            logp_idx = features.columns.get_loc('logp')
            logp_values = features[:, logp_idx]
            
            # Penalize extreme LogP values with poor binding
            logp_penalty = torch.abs(logp_values - 2.5)  # Optimal LogP around 2.5
            physics_loss += 0.05 * torch.mean(logp_penalty)
        
        # Constraint 4: TPSA vs binding affinity relationship
        # Very high TPSA may indicate poor membrane permeability
        if 'tpsa' in features.columns:
            tpsa_idx = features.columns.get_loc('tpsa')
            tpsa_values = features[:, tpsa_idx]
            
            # Penalize very high TPSA values
            tpsa_penalty = F.relu(tpsa_values - 140.0)  # TPSA > 140 is often problematic
            physics_loss += 0.05 * torch.mean(tpsa_penalty)
        
        return physics_loss


class PhysicsInformedEnsemble(EnsembleModelInterface):
    """
    Physics-informed ensemble that combines traditional ML models with
    physics-informed neural networks.
    """
    
    def __init__(self, base_models: Dict[str, Any], 
                 physics_model: Optional[PhysicsInformedNeuralNetwork] = None,
                 physics_weight: float = 0.3, random_state: int = 42):
        """
        Initialize physics-informed ensemble.
        
        Args:
            base_models: Dictionary of base model trainers
            physics_model: Physics-informed neural network
            physics_weight: Weight for physics model in ensemble
            random_state: Random state for reproducibility
        """
        self.base_models = base_models
        self.physics_model = physics_model
        self.physics_weight = physics_weight
        self.random_state = random_state
        self.trained_base_models = {}
        self.trained_physics_model = None
        self.feature_names = None
        
    def fit(self, X_train: pd.DataFrame, y_train: pd.Series, 
            X_val: Optional[pd.DataFrame] = None, 
            y_val: Optional[pd.Series] = None) -> None:
        """Fit the physics-informed ensemble model."""
        if not TORCH_AVAILABLE and self.physics_model is not None:
            raise RuntimeError("PyTorch not available for physics-informed ensemble")
        
        logger.info("Training physics-informed ensemble model")
        
        # Store feature names
        self.feature_names = X_train.columns.tolist()
        
        # Train base models
        for model_name, trainer in self.base_models.items():
            try:
                logger.info(f"Training base model: {model_name}")
                model = trainer.train(X_train, y_train, X_val, y_val)
                self.trained_base_models[model_name] = model
            except Exception as e:
                logger.error(f"Error training {model_name}: {str(e)}")
                continue
        
        # Train physics-informed model if provided
        if self.physics_model is not None:
            self._train_physics_model(X_train, y_train, X_val, y_val)
        
        logger.info("Physics-informed ensemble training completed successfully")
    
    def _train_physics_model(self, X_train: pd.DataFrame, y_train: pd.Series,
                           X_val: Optional[pd.DataFrame] = None, 
                           y_val: Optional[pd.Series] = None) -> None:
        """Train the physics-informed neural network."""
        logger.info("Training physics-informed neural network")
        
        # Convert to tensors
        X_train_tensor = torch.tensor(X_train.values, dtype=torch.float32)
        y_train_tensor = torch.tensor(y_train.values, dtype=torch.float32).unsqueeze(1)
        
        # Create data loaders
        train_dataset = torch.utils.data.TensorDataset(X_train_tensor, y_train_tensor)
        train_loader = DataLoader(train_dataset, batch_size=32, shuffle=True)
        
        val_loader = None
        if X_val is not None and y_val is not None:
            X_val_tensor = torch.tensor(X_val.values, dtype=torch.float32)
            y_val_tensor = torch.tensor(y_val.values, dtype=torch.float32).unsqueeze(1)
            val_dataset = torch.utils.data.TensorDataset(X_val_tensor, y_val_tensor)
            val_loader = DataLoader(val_dataset, batch_size=32, shuffle=False)
        
        # Training setup
        optimizer = torch.optim.Adam(self.physics_model.parameters(), lr=0.001)
        criterion = nn.MSELoss()
        
        # Training loop
        num_epochs = 100
        best_val_loss = float('inf')
        patience = 10
        patience_counter = 0
        
        for epoch in range(num_epochs):
            # Training
            self.physics_model.train()
            train_loss = 0.0
            
            for batch_X, batch_y in train_loader:
                optimizer.zero_grad()
                
                predictions = self.physics_model(batch_X)
                mse_loss = criterion(predictions, batch_y)
                physics_loss = self.physics_model.physics_loss(predictions, batch_X)
                
                total_loss = mse_loss + 0.1 * physics_loss
                total_loss.backward()
                optimizer.step()
                
                train_loss += total_loss.item()
            
            # Validation
            if val_loader is not None:
                self.physics_model.eval()
                val_loss = 0.0
                
                with torch.no_grad():
                    for batch_X, batch_y in val_loader:
                        predictions = self.physics_model(batch_X)
                        mse_loss = criterion(predictions, batch_y)
                        physics_loss = self.physics_model.physics_loss(predictions, batch_X)
                        total_loss = mse_loss + 0.1 * physics_loss
                        val_loss += total_loss.item()
                
                val_loss /= len(val_loader)
                
                # Early stopping
                if val_loss < best_val_loss:
                    best_val_loss = val_loss
                    patience_counter = 0
                    # Save best model
                    self.trained_physics_model = self.physics_model.state_dict().copy()
                else:
                    patience_counter += 1
                    if patience_counter >= patience:
                        logger.info(f"Early stopping at epoch {epoch}")
                        break
            
            if epoch % 20 == 0:
                logger.info(f"Epoch {epoch}, Train Loss: {train_loss/len(train_loader):.4f}")
        
        # Load best model
        if self.trained_physics_model is not None:
            self.physics_model.load_state_dict(self.trained_physics_model)
        
        logger.info("Physics-informed neural network training completed")
    
    def predict(self, X: pd.DataFrame) -> np.ndarray:
        """Make predictions using the physics-informed ensemble model."""
        if not self.trained_base_models and self.trained_physics_model is None:
            raise ValueError("Model must be fitted before making predictions")
        
        predictions = []
        
        # Get base model predictions
        if self.trained_base_models:
            base_predictions = []
            for model_name, model in self.trained_base_models.items():
                try:
                    trainer = self.base_models[model_name]
                    pred = trainer.predict(model, X)
                    base_predictions.append(pred)
                except Exception as e:
                    logger.error(f"Error making predictions with {model_name}: {str(e)}")
                    base_predictions.append(np.zeros(len(X)))
            
            # Average base model predictions
            base_avg = np.mean(base_predictions, axis=0)
            predictions.append(base_avg)
        
        # Get physics model predictions
        if self.trained_physics_model is not None:
            try:
                X_tensor = torch.tensor(X.values, dtype=torch.float32)
                self.physics_model.eval()
                with torch.no_grad():
                    physics_pred = self.physics_model(X_tensor).numpy().flatten()
                predictions.append(physics_pred)
            except Exception as e:
                logger.error(f"Error making predictions with physics model: {str(e)}")
                predictions.append(np.zeros(len(X)))
        
        # Combine predictions
        if len(predictions) == 1:
            return predictions[0]
        else:
            # Weighted combination
            base_weight = 1.0 - self.physics_weight
            return base_weight * predictions[0] + self.physics_weight * predictions[1]
    
    def predict_with_uncertainty(self, X: pd.DataFrame) -> List[EnsemblePrediction]:
        """Make predictions with uncertainty quantification."""
        predictions = self.predict(X)
        
        # Calculate uncertainty based on model disagreement
        model_predictions = {}
        
        # Base model predictions
        if self.trained_base_models:
            base_predictions = []
            for model_name, model in self.trained_base_models.items():
                try:
                    trainer = self.base_models[model_name]
                    pred = trainer.predict(model, X)
                    base_predictions.append(pred)
                    model_predictions[model_name] = pred
                except Exception as e:
                    logger.error(f"Error making predictions with {model_name}: {str(e)}")
                    pred = np.zeros(len(X))
                    model_predictions[model_name] = pred
                    base_predictions.append(pred)
            
            base_avg = np.mean(base_predictions, axis=0)
            model_predictions['base_ensemble'] = base_avg
        
        # Physics model predictions
        if self.trained_physics_model is not None:
            try:
                X_tensor = torch.tensor(X.values, dtype=torch.float32)
                self.physics_model.eval()
                with torch.no_grad():
                    physics_pred = self.physics_model(X_tensor).numpy().flatten()
                model_predictions['physics_model'] = physics_pred
            except Exception as e:
                logger.error(f"Error making predictions with physics model: {str(e)}")
                physics_pred = np.zeros(len(X))
                model_predictions['physics_model'] = physics_pred
        
        # Calculate uncertainty as standard deviation of model predictions
        all_predictions = np.array(list(model_predictions.values()))
        uncertainties = np.std(all_predictions, axis=0)
        
        # Create ensemble predictions
        ensemble_predictions = []
        for i in range(len(X)):
            # Calculate confidence interval (95%)
            ci_lower = predictions[i] - 1.96 * uncertainties[i]
            ci_upper = predictions[i] + 1.96 * uncertainties[i]
            
            # Calculate weights
            weights = {}
            if self.trained_base_models and self.trained_physics_model is not None:
                weights['base_ensemble'] = 1.0 - self.physics_weight
                weights['physics_model'] = self.physics_weight
            else:
                # Equal weights if only one type of model
                for model_name in model_predictions.keys():
                    weights[model_name] = 1.0 / len(model_predictions)
            
            ensemble_predictions.append(EnsemblePrediction(
                prediction=float(predictions[i]),
                uncertainty=float(uncertainties[i]),
                individual_predictions={name: pred[i] for name, pred in model_predictions.items()},
                weights=weights,
                confidence_interval=(ci_lower, ci_upper)
            ))
        
        return ensemble_predictions


class EnsembleModelTrainer:
    """
    Main ensemble model trainer coordinator.
    
    Orchestrates training of various ensemble methods and provides
    comprehensive ensemble model comparison and selection.
    """
    
    def __init__(self, models_dir: str = "models/ensemble", 
                 include_physics_informed: bool = True):
        """
        Initialize ensemble model trainer.
        
        Args:
            models_dir: Directory to save trained ensemble models
            include_physics_informed: Whether to include physics-informed models
        """
        self.models_dir = Path(models_dir)
        self.models_dir.mkdir(parents=True, exist_ok=True)
        
        self.include_physics_informed = include_physics_informed
        self.trained_ensembles = {}
        self.training_results = {}
    
    def train_ensembles(self, X_train: pd.DataFrame, y_train: pd.Series,
                       X_val: Optional[pd.DataFrame] = None,
                       y_val: Optional[pd.Series] = None,
                       base_models: Optional[Dict[str, Any]] = None) -> Dict[str, Any]:
        """
        Train all ensemble models.
        
        Args:
            X_train: Training features
            y_train: Training targets
            X_val: Validation features (optional)
            y_val: Validation targets (optional)
            base_models: Dictionary of base model trainers
            
        Returns:
            Dictionary of trained ensemble models
        """
        if base_models is None:
            logger.warning("No base models provided. Creating default base models.")
            base_models = self._create_default_base_models()
        
        logger.info(f"Training ensemble models with {len(base_models)} base models")
        
        # Train stacked ensemble
        try:
            logger.info("Training stacked ensemble")
            stacked_ensemble = StackedEnsemble(base_models)
            stacked_ensemble.fit(X_train, y_train, X_val, y_val)
            self.trained_ensembles['stacked'] = stacked_ensemble
            
            # Calculate training metrics
            train_pred = stacked_ensemble.predict(X_train)
            train_metrics = self._calculate_metrics(y_train, train_pred)
            
            self.training_results['stacked'] = {
                'model': stacked_ensemble,
                'train_metrics': train_metrics,
                'model_type': 'stacked_ensemble'
            }
            
            logger.info("Stacked ensemble training completed successfully")
            
        except Exception as e:
            logger.error(f"Error training stacked ensemble: {str(e)}")
        
        # Train weighted ensemble
        try:
            logger.info("Training weighted ensemble")
            weighted_ensemble = WeightedEnsemble(base_models, weight_method='performance')
            weighted_ensemble.fit(X_train, y_train, X_val, y_val)
            self.trained_ensembles['weighted'] = weighted_ensemble
            
            # Calculate training metrics
            train_pred = weighted_ensemble.predict(X_train)
            train_metrics = self._calculate_metrics(y_train, train_pred)
            
            self.training_results['weighted'] = {
                'model': weighted_ensemble,
                'train_metrics': train_metrics,
                'model_type': 'weighted_ensemble'
            }
            
            logger.info("Weighted ensemble training completed successfully")
            
        except Exception as e:
            logger.error(f"Error training weighted ensemble: {str(e)}")
        
        # Train physics-informed ensemble
        if self.include_physics_informed and TORCH_AVAILABLE:
            try:
                logger.info("Training physics-informed ensemble")
                
                # Create physics-informed neural network
                physics_model = PhysicsInformedNeuralNetwork(
                    input_dim=X_train.shape[1],
                    hidden_dims=[256, 128, 64],
                    dropout=0.1,
                    use_thermodynamic_constraints=True
                )
                
                physics_ensemble = PhysicsInformedEnsemble(
                    base_models=base_models,
                    physics_model=physics_model,
                    physics_weight=0.3
                )
                physics_ensemble.fit(X_train, y_train, X_val, y_val)
                self.trained_ensembles['physics_informed'] = physics_ensemble
                
                # Calculate training metrics
                train_pred = physics_ensemble.predict(X_train)
                train_metrics = self._calculate_metrics(y_train, train_pred)
                
                self.training_results['physics_informed'] = {
                    'model': physics_ensemble,
                    'train_metrics': train_metrics,
                    'model_type': 'physics_informed_ensemble'
                }
                
                logger.info("Physics-informed ensemble training completed successfully")
                
            except Exception as e:
                logger.error(f"Error training physics-informed ensemble: {str(e)}")
        
        logger.info(f"Successfully trained {len(self.trained_ensembles)} ensemble models")
        return self.trained_ensembles
    
    def evaluate_ensembles(self, X_test: pd.DataFrame, y_test: pd.Series) -> Dict[str, Dict]:
        """
        Evaluate all trained ensemble models on test set.
        
        Args:
            X_test: Test features
            y_test: Test targets
            
        Returns:
            Dictionary of evaluation results for each ensemble model
        """
        logger.info("Evaluating ensemble models on test set")
        
        evaluation_results = {}
        
        for ensemble_name, ensemble in self.trained_ensembles.items():
            try:
                # Make predictions
                predictions = ensemble.predict(X_test)
                
                # Calculate metrics
                metrics = self._calculate_metrics(y_test, predictions)
                
                # Get uncertainty predictions
                uncertainty_predictions = ensemble.predict_with_uncertainty(X_test)
                
                evaluation_results[ensemble_name] = {
                    'predictions': predictions,
                    'metrics': metrics,
                    'uncertainty_predictions': uncertainty_predictions
                }
                
                logger.info(f"{ensemble_name} evaluation completed")
                
            except Exception as e:
                logger.error(f"Error evaluating {ensemble_name}: {str(e)}")
                continue
        
        return evaluation_results
    
    def get_best_ensemble(self, metric: str = 'r2') -> Tuple[str, Any]:
        """
        Get the best performing ensemble model based on specified metric.
        
        Args:
            metric: Metric to use for model selection
            
        Returns:
            Tuple of (ensemble_name, ensemble_model)
        """
        if not self.training_results:
            raise ValueError("No ensemble models have been trained yet")
        
        best_ensemble_name = None
        best_score = float('-inf') if metric == 'r2' else float('inf')
        
        for ensemble_name, results in self.training_results.items():
            if results is None:
                continue
                
            if 'train_metrics' in results and metric in results['train_metrics']:
                score = results['train_metrics'][metric]
                
                if metric == 'r2':
                    if score > best_score:
                        best_score = score
                        best_ensemble_name = ensemble_name
                else:  # For loss-based metrics
                    if score < best_score:
                        best_score = score
                        best_ensemble_name = ensemble_name
        
        if best_ensemble_name is None:
            raise ValueError(f"No ensemble found with metric {metric}")
        
        return best_ensemble_name, self.trained_ensembles[best_ensemble_name]
    
    def _create_default_base_models(self) -> Dict[str, Any]:
        """Create default base models for ensemble training."""
        base_models = {}
        
        # Import base model trainers
        try:
            from .trainer import RandomForestTrainer, SVRTrainer, XGBoostTrainer, LightGBMTrainer
            
            if SKLEARN_AVAILABLE:
                base_models['random_forest'] = RandomForestTrainer()
                base_models['svr'] = SVRTrainer()
            
            try:
                import xgboost as xgb
                base_models['xgboost'] = XGBoostTrainer()
            except ImportError:
                pass
            
            try:
                import lightgbm as lgb
                base_models['lightgbm'] = LightGBMTrainer()
            except ImportError:
                pass
                
        except ImportError:
            logger.warning("Could not import base model trainers")
        
        return base_models
    
    def _calculate_metrics(self, y_true: pd.Series, y_pred: np.ndarray) -> Dict[str, float]:
        """Calculate regression metrics."""
        if not SKLEARN_AVAILABLE:
            return {}
        
        return {
            'mse': mean_squared_error(y_true, y_pred),
            'rmse': np.sqrt(mean_squared_error(y_true, y_pred)),
            'mae': mean_absolute_error(y_true, y_pred),
            'r2': r2_score(y_true, y_pred)
        }
    
    def save_ensemble(self, ensemble_name: str, ensemble: Any) -> None:
        """Save trained ensemble model to disk."""
        ensemble_path = self.models_dir / f"{ensemble_name}_ensemble.joblib"
        joblib.dump(ensemble, ensemble_path)
        logger.info(f"Ensemble model saved to {ensemble_path}")
    
    def load_ensemble(self, ensemble_name: str) -> Any:
        """Load trained ensemble model from disk."""
        ensemble_path = self.models_dir / f"{ensemble_name}_ensemble.joblib"
        if not ensemble_path.exists():
            raise FileNotFoundError(f"Ensemble model file not found: {ensemble_path}")
        
        return joblib.load(ensemble_path)
    
    def get_ensemble_summary(self) -> pd.DataFrame:
        """Get summary of all trained ensemble models."""
        if not self.training_results:
            return pd.DataFrame()
        
        summary_data = []
        for ensemble_name, results in self.training_results.items():
            if results is None:
                continue
                
            metrics = results.get('train_metrics', {})
            summary_data.append({
                'ensemble_name': ensemble_name,
                'model_type': results.get('model_type', 'ensemble'),
                **metrics
            })
        
        return pd.DataFrame(summary_data)
```
