## Supplementary material for "Machine Learning-Driven Prediction of TLR4 Binding Affinity: A Comprehensive Molecular Feature Analysis for Drug Discovery": Codebase: evaluator.py

```
"""
Model evaluation and performance metrics.

This module provides comprehensive model evaluation functionality
for TLR4 binding prediction including various regression metrics
and statistical analysis.
"""

from abc import ABC, abstractmethod
from typing import Dict, List, Optional, Union, Tuple, Any
import pandas as pd
import numpy as np
import logging
from sklearn.metrics import (
    mean_squared_error, mean_absolute_error, r2_score,
    explained_variance_score, max_error
)
from sklearn.model_selection import (
    learning_curve, validation_curve, cross_val_score,
    KFold, StratifiedKFold
)
from scipy import stats
import matplotlib.pyplot as plt
import seaborn as sns
from sklearn.base import BaseEstimator

logger = logging.getLogger(__name__)


class PerformanceMetrics:
    """Container for performance metrics and statistical analysis."""
    
    def __init__(self, y_true: np.ndarray, y_pred: np.ndarray, model_name: str = ""):
        """
        Initialize performance metrics.
        
        Args:
            y_true: True target values
            y_pred: Predicted target values
            model_name: Name of the model
        """
        self.y_true = y_true
        self.y_pred = y_pred
        self.model_name = model_name
        self.metrics = self._calculate_metrics()
    
    def _calculate_metrics(self) -> Dict[str, float]:
        """Calculate comprehensive regression metrics."""
        # Basic regression metrics
        mse = mean_squared_error(self.y_true, self.y_pred)
        rmse = np.sqrt(mse)
        mae = mean_absolute_error(self.y_true, self.y_pred)
        r2 = r2_score(self.y_true, self.y_pred)
        evs = explained_variance_score(self.y_true, self.y_pred)
        max_err = max_error(self.y_true, self.y_pred)
        
        # Additional metrics
        mape = self._calculate_mape(self.y_true, self.y_pred)
        smape = self._calculate_smape(self.y_true, self.y_pred)
        pearson_r, pearson_p = stats.pearsonr(self.y_true, self.y_pred)
        spearman_r, spearman_p = stats.spearmanr(self.y_true, self.y_pred)
        
        # Residual analysis
        residuals = self.y_true - self.y_pred
        residual_std = np.std(residuals)
        residual_mean = np.mean(residuals)
        
        return {
            'mse': mse,
            'rmse': rmse,
            'mae': mae,
            'r2': r2,
            'explained_variance': evs,
            'max_error': max_err,
            'mape': mape,
            'smape': smape,
            'pearson_r': pearson_r,
            'pearson_p': pearson_p,
            'spearman_r': spearman_r,
            'spearman_p': spearman_p,
            'residual_mean': residual_mean,
            'residual_std': residual_std,
            'residual_rmse': np.sqrt(np.mean(residuals**2))
        }
    
    def _calculate_mape(self, y_true: np.ndarray, y_pred: np.ndarray) -> float:
        """Calculate Mean Absolute Percentage Error."""
        mask = y_true != 0
        if not np.any(mask):
            return np.nan
        return np.mean(np.abs((y_true[mask] - y_pred[mask]) / y_true[mask])) * 100
    
    def _calculate_smape(self, y_true: np.ndarray, y_pred: np.ndarray) -> float:
        """Calculate Symmetric Mean Absolute Percentage Error."""
        denominator = (np.abs(y_true) + np.abs(y_pred)) / 2
        mask = denominator != 0
        if not np.any(mask):
            return np.nan
        return np.mean(np.abs(y_true[mask] - y_pred[mask]) / denominator[mask]) * 100
    
    def get_summary(self) -> Dict[str, Any]:
        """Get summary of key metrics."""
        return {
            'model_name': self.model_name,
            'r2_score': self.metrics['r2'],
            'rmse': self.metrics['rmse'],
            'mae': self.metrics['mae'],
            'pearson_correlation': self.metrics['pearson_r'],
            'spearman_correlation': self.metrics['spearman_r']
        }
    
    def to_dataframe(self) -> pd.DataFrame:
        """Convert metrics to DataFrame."""
        return pd.DataFrame([self.metrics])


class ModelEvaluatorInterface(ABC):
    """Abstract interface for model evaluation."""
    
    @abstractmethod
    def evaluate(self, y_true: np.ndarray, y_pred: np.ndarray, 
                model_name: str = "") -> PerformanceMetrics:
        """Evaluate model performance."""
        pass
    
    @abstractmethod
    def compare_models(self, results: Dict[str, PerformanceMetrics]) -> pd.DataFrame:
        """Compare multiple model performances."""
        pass


class ModelEvaluator(ModelEvaluatorInterface):
    """Comprehensive model evaluation with statistical analysis."""
    
    def __init__(self, confidence_level: float = 0.95):
        """
        Initialize model evaluator.
        
        Args:
            confidence_level: Confidence level for statistical tests
        """
        self.confidence_level = confidence_level
        self.evaluation_history = []
    
    def evaluate(self, y_true: np.ndarray, y_pred: np.ndarray, 
                model_name: str = "") -> PerformanceMetrics:
        """
        Evaluate model performance comprehensively.
        
        Args:
            y_true: True target values
            y_pred: Predicted target values
            model_name: Name of the model
            
        Returns:
            PerformanceMetrics object with all calculated metrics
        """
        logger.info(f"Evaluating model: {model_name}")
        
        # Create performance metrics
        metrics = PerformanceMetrics(y_true, y_pred, model_name)
        
        # Store evaluation history
        self.evaluation_history.append({
            'timestamp': pd.Timestamp.now(),
            'model_name': model_name,
            'metrics': metrics
        })
        
        logger.info(f"Model {model_name} evaluation completed. R² = {metrics.metrics['r2']:.4f}")
        
        return metrics
    
    def compare_models(self, results: Dict[str, PerformanceMetrics]) -> pd.DataFrame:
        """
        Compare multiple model performances.
        
        Args:
            results: Dictionary of model names and PerformanceMetrics objects
            
        Returns:
            DataFrame with comparison of all models
        """
        if not results:
            return pd.DataFrame()
        
        comparison_data = []
        for model_name, metrics in results.items():
            summary = metrics.get_summary()
            comparison_data.append(summary)
        
        comparison_df = pd.DataFrame(comparison_data)
        
        # Sort by R² score (descending)
        comparison_df = comparison_df.sort_values('r2_score', ascending=False)
        
        logger.info(f"Model comparison completed for {len(results)} models")
        
        return comparison_df
    
    def statistical_significance_test(self, metrics1: PerformanceMetrics, 
                                   metrics2: PerformanceMetrics) -> Dict[str, Any]:
        """
        Perform statistical significance test between two models.
        
        Args:
            metrics1: Performance metrics for first model
            metrics2: Performance metrics for second model
            
        Returns:
            Dictionary with statistical test results
        """
        # Paired t-test on residuals
        residuals1 = metrics1.y_true - metrics1.y_pred
        residuals2 = metrics2.y_true - metrics2.y_pred
        
        if len(residuals1) != len(residuals2):
            raise ValueError("Models must have same number of predictions for comparison")
        
        # Paired t-test
        t_stat, t_p = stats.ttest_rel(residuals1, residuals2)
        
        # Wilcoxon signed-rank test (non-parametric alternative)
        try:
            w_stat, w_p = stats.wilcoxon(residuals1, residuals2)
        except ValueError:
            w_stat, w_p = np.nan, np.nan
        
        # McNemar's test for classification accuracy (if applicable)
        # This is a placeholder for future classification metrics
        
        return {
            'paired_t_test': {
                'statistic': t_stat,
                'p_value': t_p,
                'significant': t_p < (1 - self.confidence_level)
            },
            'wilcoxon_test': {
                'statistic': w_stat,
                'p_value': w_p,
                'significant': w_p < (1 - self.confidence_level) if not np.isnan(w_p) else False
            },
            'model1_r2': metrics1.metrics['r2'],
            'model2_r2': metrics2.metrics['r2'],
            'r2_difference': metrics1.metrics['r2'] - metrics2.metrics['r2']
        }
    
    def generate_evaluation_report(self, results: Dict[str, PerformanceMetrics]) -> Dict[str, Any]:
        """
        Generate comprehensive evaluation report.
        
        Args:
            results: Dictionary of model names and PerformanceMetrics objects
            
        Returns:
            Dictionary with comprehensive evaluation report
        """
        if not results:
            return {'error': 'No evaluation results available'}
        
        # Model comparison
        comparison_df = self.compare_models(results)
        
        # Best model identification
        best_model = comparison_df.iloc[0]['model_name']
        best_r2 = comparison_df.iloc[0]['r2_score']
        
        # Statistical significance tests
        significance_tests = {}
        model_names = list(results.keys())
        for i in range(len(model_names)):
            for j in range(i + 1, len(model_names)):
                model1, model2 = model_names[i], model_names[j]
                test_name = f"{model1}_vs_{model2}"
                significance_tests[test_name] = self.statistical_significance_test(
                    results[model1], results[model2]
                )
        
        # Summary statistics
        r2_scores = [metrics.metrics['r2'] for metrics in results.values()]
        rmse_scores = [metrics.metrics['rmse'] for metrics in results.values()]
        
        report = {
            'summary': {
                'total_models': len(results),
                'best_model': best_model,
                'best_r2_score': best_r2,
                'r2_range': [min(r2_scores), max(r2_scores)],
                'rmse_range': [min(rmse_scores), max(rmse_scores)]
            },
            'model_comparison': comparison_df.to_dict('records'),
            'statistical_tests': significance_tests,
            'evaluation_timestamp': pd.Timestamp.now().isoformat()
        }
        
        return report
    
    def plot_residuals(self, metrics: PerformanceMetrics, 
                      save_path: Optional[str] = None) -> None:
        """
        Plot residual analysis for a model.
        
        Args:
            metrics: PerformanceMetrics object
            save_path: Optional path to save plot
        """
        fig, axes = plt.subplots(2, 2, figsize=(12, 10))
        fig.suptitle(f'Residual Analysis - {metrics.model_name}', fontsize=16)
        
        residuals = metrics.y_true - metrics.y_pred
        
        # Residuals vs Predicted
        axes[0, 0].scatter(metrics.y_pred, residuals, alpha=0.6)
        axes[0, 0].axhline(y=0, color='r', linestyle='--')
        axes[0, 0].set_xlabel('Predicted Values')
        axes[0, 0].set_ylabel('Residuals')
        axes[0, 0].set_title('Residuals vs Predicted')
        
        # Q-Q plot
        stats.probplot(residuals, dist="norm", plot=axes[0, 1])
        axes[0, 1].set_title('Q-Q Plot of Residuals')
        
        # Histogram of residuals
        axes[1, 0].hist(residuals, bins=30, alpha=0.7, edgecolor='black')
        axes[1, 0].set_xlabel('Residuals')
        axes[1, 0].set_ylabel('Frequency')
        axes[1, 0].set_title('Distribution of Residuals')
        
        # Actual vs Predicted
        axes[1, 1].scatter(metrics.y_true, metrics.y_pred, alpha=0.6)
        min_val = min(metrics.y_true.min(), metrics.y_pred.min())
        max_val = max(metrics.y_true.max(), metrics.y_pred.max())
        axes[1, 1].plot([min_val, max_val], [min_val, max_val], 'r--', lw=2)
        axes[1, 1].set_xlabel('Actual Values')
        axes[1, 1].set_ylabel('Predicted Values')
        axes[1, 1].set_title('Actual vs Predicted')
        
        plt.tight_layout()
        
        if save_path:
            plt.savefig(save_path, dpi=300, bbox_inches='tight')
            logger.info(f"Residual plot saved to {save_path}")
        
        plt.show()
    
    def plot_model_comparison(self, results: Dict[str, PerformanceMetrics],
                            save_path: Optional[str] = None) -> None:
        """
        Plot comparison of multiple models.
        
        Args:
            results: Dictionary of model names and PerformanceMetrics objects
            save_path: Optional path to save plot
        """
        if not results:
            logger.warning("No results to plot")
            return
        
        fig, axes = plt.subplots(2, 2, figsize=(15, 12))
        fig.suptitle('Model Performance Comparison', fontsize=16)
        
        # R² scores
        model_names = list(results.keys())
        r2_scores = [results[name].metrics['r2'] for name in model_names]
        axes[0, 0].bar(model_names, r2_scores)
        axes[0, 0].set_ylabel('R² Score')
        axes[0, 0].set_title('R² Score Comparison')
        axes[0, 0].tick_params(axis='x', rotation=45)
        
        # RMSE scores
        rmse_scores = [results[name].metrics['rmse'] for name in model_names]
        axes[0, 1].bar(model_names, rmse_scores)
        axes[0, 1].set_ylabel('RMSE')
        axes[0, 1].set_title('RMSE Comparison')
        axes[0, 1].tick_params(axis='x', rotation=45)
        
        # MAE scores
        mae_scores = [results[name].metrics['mae'] for name in model_names]
        axes[1, 0].bar(model_names, mae_scores)
        axes[1, 0].set_ylabel('MAE')
        axes[1, 0].set_title('MAE Comparison')
        axes[1, 0].tick_params(axis='x', rotation=45)
        
        # Pearson correlation
        pearson_scores = [results[name].metrics['pearson_r'] for name in model_names]
        axes[1, 1].bar(model_names, pearson_scores)
        axes[1, 1].set_ylabel('Pearson Correlation')
        axes[1, 1].set_title('Pearson Correlation Comparison')
        axes[1, 1].tick_params(axis='x', rotation=45)
        
        plt.tight_layout()
        
        if save_path:
            plt.savefig(save_path, dpi=300, bbox_inches='tight')
            logger.info(f"Model comparison plot saved to {save_path}")
        
        plt.show()
    
    def generate_learning_curves(self, model: BaseEstimator, X: np.ndarray, y: np.ndarray,
                                model_name: str = "", cv: int = 5, 
                                train_sizes: Optional[List[float]] = None,
                                save_path: Optional[str] = None) -> Dict[str, Any]:
        """
        Generate learning curves for a model.
        
        Args:
            model: Trained model object
            X: Feature matrix
            y: Target values
            model_name: Name of the model
            cv: Number of cross-validation folds
            train_sizes: Training set sizes to evaluate
            save_path: Optional path to save plot
            
        Returns:
            Dictionary with learning curve data
        """
        if train_sizes is None:
            train_sizes = np.linspace(0.1, 1.0, 10)
        
        logger.info(f"Generating learning curves for {model_name}")
        
        # Generate learning curves
        train_sizes_abs, train_scores, val_scores = learning_curve(
            model, X, y, cv=cv, train_sizes=train_sizes,
            scoring='neg_mean_squared_error', n_jobs=-1
        )
        
        # Convert to positive RMSE
        train_rmse = np.sqrt(-train_scores)
        val_rmse = np.sqrt(-val_scores)
        
        # Calculate means and standard deviations
        train_mean = np.mean(train_rmse, axis=1)
        train_std = np.std(train_rmse, axis=1)
        val_mean = np.mean(val_rmse, axis=1)
        val_std = np.std(val_rmse, axis=1)
        
        # Plot learning curves
        plt.figure(figsize=(10, 6))
        plt.plot(train_sizes_abs, train_mean, 'o-', label='Training RMSE', color='blue')
        plt.fill_between(train_sizes_abs, train_mean - train_std, 
                        train_mean + train_std, alpha=0.1, color='blue')
        
        plt.plot(train_sizes_abs, val_mean, 'o-', label='Validation RMSE', color='red')
        plt.fill_between(train_sizes_abs, val_mean - val_std, 
                        val_mean + val_std, alpha=0.1, color='red')
        
        plt.xlabel('Training Set Size')
        plt.ylabel('RMSE')
        plt.title(f'Learning Curves - {model_name}')
        plt.legend()
        plt.grid(True, alpha=0.3)
        
        if save_path:
            plt.savefig(save_path, dpi=300, bbox_inches='tight')
            logger.info(f"Learning curves saved to {save_path}")
        
        plt.show()
        
        return {
            'train_sizes': train_sizes_abs,
            'train_mean': train_mean,
            'train_std': train_std,
            'val_mean': val_mean,
            'val_std': val_std,
            'model_name': model_name
        }
    
    def generate_validation_curves(self, model: BaseEstimator, X: np.ndarray, y: np.ndarray,
                                  param_name: str, param_range: List[Any],
                                  model_name: str = "", cv: int = 5,
                                  save_path: Optional[str] = None) -> Dict[str, Any]:
        """
        Generate validation curves for a model parameter.
        
        Args:
            model: Model object (will be cloned for each parameter value)
            X: Feature matrix
            y: Target values
            param_name: Name of parameter to vary
            param_range: Range of parameter values to test
            model_name: Name of the model
            cv: Number of cross-validation folds
            save_path: Optional path to save plot
            
        Returns:
            Dictionary with validation curve data
        """
        logger.info(f"Generating validation curves for {model_name} parameter: {param_name}")
        
        # Generate validation curves
        train_scores, val_scores = validation_curve(
            model, X, y, param_name=param_name, param_range=param_range,
            cv=cv, scoring='neg_mean_squared_error', n_jobs=-1
        )
        
        # Convert to positive RMSE
        train_rmse = np.sqrt(-train_scores)
        val_rmse = np.sqrt(-val_scores)
        
        # Calculate means and standard deviations
        train_mean = np.mean(train_rmse, axis=1)
        train_std = np.std(train_rmse, axis=1)
        val_mean = np.mean(val_rmse, axis=1)
        val_std = np.std(val_rmse, axis=1)
        
        # Plot validation curves
        plt.figure(figsize=(10, 6))
        plt.plot(param_range, train_mean, 'o-', label='Training RMSE', color='blue')
        plt.fill_between(param_range, train_mean - train_std, 
                        train_mean + train_std, alpha=0.1, color='blue')
        
        plt.plot(param_range, val_mean, 'o-', label='Validation RMSE', color='red')
        plt.fill_between(param_range, val_mean - val_std, 
                        val_mean + val_std, alpha=0.1, color='red')
        
        plt.xlabel(param_name)
        plt.ylabel('RMSE')
        plt.title(f'Validation Curves - {model_name} ({param_name})')
        plt.legend()
        plt.grid(True, alpha=0.3)
        
        if save_path:
            plt.savefig(save_path, dpi=300, bbox_inches='tight')
            logger.info(f"Validation curves saved to {save_path}")
        
        plt.show()
        
        return {
            'param_range': param_range,
            'train_mean': train_mean,
            'train_std': train_std,
            'val_mean': val_mean,
            'val_std': val_std,
            'param_name': param_name,
            'model_name': model_name
        }
    
    def cross_validate_model(self, model: BaseEstimator, X: np.ndarray, y: np.ndarray,
                           model_name: str = "", cv: int = 5, 
                           scoring: List[str] = None) -> Dict[str, Any]:
        """
        Perform comprehensive cross-validation evaluation.
        
        Args:
            model: Model object to evaluate
            X: Feature matrix
            y: Target values
            model_name: Name of the model
            cv: Number of cross-validation folds
            scoring: List of scoring metrics
            
        Returns:
            Dictionary with cross-validation results
        """
        if scoring is None:
            scoring = ['neg_mean_squared_error', 'neg_mean_absolute_error', 'r2']
        
        logger.info(f"Performing {cv}-fold cross-validation for {model_name}")
        
        cv_results = {}
        cv_fold = KFold(n_splits=cv, shuffle=True, random_state=42)
        
        for metric in scoring:
            scores = cross_val_score(model, X, y, cv=cv_fold, scoring=metric, n_jobs=-1)
            cv_results[metric] = {
                'scores': scores,
                'mean': np.mean(scores),
                'std': np.std(scores),
                'min': np.min(scores),
                'max': np.max(scores)
            }
        
        # Convert MSE and MAE to positive values for easier interpretation
        if 'neg_mean_squared_error' in cv_results:
            cv_results['rmse'] = {
                'scores': np.sqrt(-cv_results['neg_mean_squared_error']['scores']),
                'mean': np.sqrt(-cv_results['neg_mean_squared_error']['mean']),
                'std': cv_results['neg_mean_squared_error']['std'] / (2 * np.sqrt(-cv_results['neg_mean_squared_error']['mean'])),
                'min': np.sqrt(-cv_results['neg_mean_squared_error']['max']),
                'max': np.sqrt(-cv_results['neg_mean_squared_error']['min'])
            }
        
        if 'neg_mean_absolute_error' in cv_results:
            cv_results['mae'] = {
                'scores': -cv_results['neg_mean_absolute_error']['scores'],
                'mean': -cv_results['neg_mean_absolute_error']['mean'],
                'std': cv_results['neg_mean_absolute_error']['std'],
                'min': -cv_results['neg_mean_absolute_error']['max'],
                'max': -cv_results['neg_mean_absolute_error']['min']
            }
        
        logger.info(f"Cross-validation completed for {model_name}")
        return cv_results
```
