## Supplementary material for "Machine Learning-Driven Prediction of TLR4 Binding Affinity: A Comprehensive Molecular Feature Analysis for Drug Discovery": Codebase: feature_engineering.py

```
"""
Feature Engineering Pipeline for TLR4 Binding Prediction

This module provides comprehensive feature engineering capabilities including
scaling, correlation analysis, and feature selection for molecular descriptors.
"""

import numpy as np
import pandas as pd
from typing import Dict, List, Tuple, Optional, Any
from sklearn.preprocessing import StandardScaler, RobustScaler
from sklearn.feature_selection import mutual_info_regression, SelectKBest, f_regression
from sklearn.decomposition import PCA
import logging
from dataclasses import dataclass

logger = logging.getLogger(__name__)


@dataclass
class FeatureEngineeringConfig:
    """Configuration for feature engineering pipeline"""
    correlation_threshold: float = 0.95
    mutual_info_k: int = 20
    use_robust_scaling: bool = False
    apply_pca: bool = False
    pca_components: Optional[int] = None
    min_variance: float = 1e-6


class FeatureScaler:
    """
    Handles feature scaling and normalization for molecular descriptors.
    
    Supports both standard scaling and robust scaling to handle outliers
    commonly found in molecular descriptor datasets.
    """
    
    def __init__(self, use_robust: bool = False):
        """
        Initialize the feature scaler.
        
        Args:
            use_robust: If True, use RobustScaler (median/IQR) instead of StandardScaler
        """
        self.use_robust = use_robust
        self.scaler = RobustScaler() if use_robust else StandardScaler()
        self.is_fitted = False
        self.feature_names = None
        
    def fit(self, X: pd.DataFrame) -> 'FeatureScaler':
        """
        Fit the scaler to the training data.
        
        Args:
            X: Training features DataFrame
            
        Returns:
            Self for method chaining
        """
        if X.empty:
            raise ValueError("Cannot fit scaler on empty DataFrame")
            
        # Store feature names for consistency
        self.feature_names = X.columns.tolist()
        
        # Fit the scaler
        self.scaler.fit(X)
        self.is_fitted = True
        
        logger.info(f"Fitted {type(self.scaler).__name__} on {X.shape[1]} features")
        return self
        
    def transform(self, X: pd.DataFrame) -> pd.DataFrame:
        """
        Transform features using fitted scaler.
        
        Args:
            X: Features to transform
            
        Returns:
            Scaled features DataFrame
        """
        if not self.is_fitted:
            raise ValueError("Scaler must be fitted before transform")
            
        if X.empty:
            return X
            
        # Ensure same features as training
        missing_features = set(self.feature_names) - set(X.columns)
        if missing_features:
            raise ValueError(f"Missing features: {missing_features}")
            
        # Transform and return as DataFrame
        X_scaled = self.scaler.transform(X[self.feature_names])
        return pd.DataFrame(X_scaled, columns=self.feature_names, index=X.index)
        
    def fit_transform(self, X: pd.DataFrame) -> pd.DataFrame:
        """Fit scaler and transform features in one step."""
        return self.fit(X).transform(X)


class CorrelationAnalyzer:
    """
    Analyzes and removes highly correlated features to reduce redundancy.
    
    Identifies feature pairs with correlation above threshold and removes
    one feature from each highly correlated pair.
    """
    
    def __init__(self, threshold: float = 0.95):
        """
        Initialize correlation analyzer.
        
        Args:
            threshold: Correlation threshold above which features are considered redundant
        """
        self.threshold = threshold
        self.correlated_pairs = []
        self.features_to_remove = set()
        
    def find_correlated_features(self, X: pd.DataFrame) -> Dict[str, Any]:
        """
        Find highly correlated feature pairs.
        
        Args:
            X: Features DataFrame
            
        Returns:
            Dictionary with correlation analysis results
        """
        if X.empty:
            return {"correlated_pairs": [], "features_to_remove": [], "correlation_matrix": None}
            
        # Calculate correlation matrix
        corr_matrix = X.corr().abs()
        
        # Find pairs above threshold (excluding diagonal)
        # Get highly correlated pairs
        correlated_pairs = []
        for i in range(len(corr_matrix.columns)):
            for j in range(i + 1, len(corr_matrix.columns)):
                correlation = corr_matrix.iloc[i, j]
                if not pd.isna(correlation) and correlation > self.threshold:
                    feature1 = corr_matrix.columns[i]
                    feature2 = corr_matrix.columns[j]
                    correlated_pairs.append({
                        'feature1': feature1,
                        'feature2': feature2,
                        'correlation': correlation
                    })
        
        self.correlated_pairs = correlated_pairs
        
        # Select features to remove (keep the one with higher variance)
        features_to_remove = []
        for pair in correlated_pairs:
            var1 = X[pair['feature1']].var()
            var2 = X[pair['feature2']].var()
            
            # Remove the feature with lower variance
            if var1 < var2:
                features_to_remove.append(pair['feature1'])
            else:
                features_to_remove.append(pair['feature2'])
        
        self.features_to_remove = set(features_to_remove)
        
        logger.info(f"Found {len(correlated_pairs)} highly correlated pairs")
        logger.info(f"Removing {len(features_to_remove)} redundant features")
        
        return {
            "correlated_pairs": correlated_pairs,
            "features_to_remove": features_to_remove,
            "correlation_matrix": corr_matrix
        }
        
    def remove_correlated_features(self, X: pd.DataFrame) -> pd.DataFrame:
        """
        Remove highly correlated features from DataFrame.
        
        Args:
            X: Features DataFrame
            
        Returns:
            DataFrame with correlated features removed
        """
        if not self.features_to_remove:
            self.find_correlated_features(X)
            
        # Remove features
        features_to_keep = [col for col in X.columns if col not in self.features_to_remove]
        X_reduced = X[features_to_keep].copy()
        
        logger.info(f"Removed {len(self.features_to_remove)} correlated features")
        logger.info(f"Remaining features: {len(features_to_keep)}")
        
        return X_reduced


class FeatureSelector:
    """
    Performs feature selection using mutual information and statistical tests.
    
    Identifies the most informative features for binding affinity prediction
    using multiple selection criteria.
    """
    
    def __init__(self, k: int = 20, method: str = 'mutual_info'):
        """
        Initialize feature selector.
        
        Args:
            k: Number of top features to select
            method: Selection method ('mutual_info', 'f_regression', 'variance')
        """
        self.k = k
        self.method = method
        self.selected_features = []
        self.feature_scores = {}
        self.selector = None
        
    def fit(self, X: pd.DataFrame, y: pd.Series) -> 'FeatureSelector':
        """
        Fit feature selector to training data.
        
        Args:
            X: Training features
            y: Training targets
            
        Returns:
            Self for method chaining
        """
        if X.empty or y.empty:
            raise ValueError("Cannot fit selector on empty data")
            
        if len(X) != len(y):
            raise ValueError("X and y must have same length")
            
        # Remove features with zero variance
        X_clean = X.loc[:, X.var() > 1e-6]
        
        if self.method == 'mutual_info':
            # Use mutual information for feature selection
            scores = mutual_info_regression(X_clean, y, random_state=42)
            self.selector = SelectKBest(score_func=lambda X, y: (scores, None), k=min(self.k, len(X_clean.columns)))
            
        elif self.method == 'f_regression':
            # Use F-statistic for feature selection
            self.selector = SelectKBest(score_func=f_regression, k=min(self.k, len(X_clean.columns)))
            
        else:
            raise ValueError(f"Unknown selection method: {self.method}")
            
        # Fit selector
        self.selector.fit(X_clean, y)
        
        # Get selected features
        self.selected_features = X_clean.columns[self.selector.get_support()].tolist()
        
        # Store feature scores
        if hasattr(self.selector, 'scores_'):
            self.feature_scores = dict(zip(X_clean.columns, self.selector.scores_))
        
        logger.info(f"Selected {len(self.selected_features)} features using {self.method}")
        
        return self
        
    def transform(self, X: pd.DataFrame) -> pd.DataFrame:
        """
        Transform features using fitted selector.
        
        Args:
            X: Features to transform
            
        Returns:
            DataFrame with selected features only
        """
        if not self.selected_features:
            raise ValueError("Selector must be fitted before transform")
            
        # Ensure all selected features are present
        missing_features = set(self.selected_features) - set(X.columns)
        if missing_features:
            logger.warning(f"Missing selected features: {missing_features}")
            available_features = [f for f in self.selected_features if f in X.columns]
            return X[available_features]
            
        return X[self.selected_features]
        
    def fit_transform(self, X: pd.DataFrame, y: pd.Series) -> pd.DataFrame:
        """Fit selector and transform features in one step."""
        return self.fit(X, y).transform(X)
        
    def get_feature_importance(self) -> pd.DataFrame:
        """
        Get feature importance scores.
        
        Returns:
            DataFrame with features and their importance scores
        """
        if not self.feature_scores:
            return pd.DataFrame()
            
        importance_df = pd.DataFrame([
            {'feature': feature, 'score': score}
            for feature, score in self.feature_scores.items()
            if feature in self.selected_features
        ]).sort_values('score', ascending=False)
        
        return importance_df


class FeatureEngineeringPipeline:
    """
    Main feature engineering pipeline that orchestrates all preprocessing steps.
    
    Combines scaling, correlation removal, and feature selection into a single
    pipeline for consistent preprocessing of molecular descriptor data.
    """
    
    def __init__(self, config: Optional[FeatureEngineeringConfig] = None):
        """
        Initialize the feature engineering pipeline.
        
        Args:
            config: Configuration object for pipeline parameters
        """
        self.config = config or FeatureEngineeringConfig()
        self.scaler = FeatureScaler(use_robust=self.config.use_robust_scaling)
        self.correlation_analyzer = CorrelationAnalyzer(threshold=self.config.correlation_threshold)
        self.feature_selector = FeatureSelector(k=self.config.mutual_info_k)
        self.pca = None
        self.is_fitted = False
        
    def fit(self, X: pd.DataFrame, y: pd.Series) -> 'FeatureEngineeringPipeline':
        """
        Fit the entire pipeline to training data.
        
        Args:
            X: Training features
            y: Training targets
            
        Returns:
            Self for method chaining
        """
        logger.info("Starting feature engineering pipeline fitting")
        
        # Step 1: Remove highly correlated features
        logger.info("Step 1: Removing highly correlated features")
        X_reduced = self.correlation_analyzer.remove_correlated_features(X)
        
        # Step 2: Scale features
        logger.info("Step 2: Scaling features")
        X_scaled = self.scaler.fit_transform(X_reduced)
        
        # Step 3: Select most informative features
        logger.info("Step 3: Selecting most informative features")
        X_selected = self.feature_selector.fit_transform(X_scaled, y)
        
        # Step 4: Apply PCA if requested
        if self.config.apply_pca:
            logger.info("Step 4: Applying PCA")
            n_components = self.config.pca_components or min(X_selected.shape[1], X_selected.shape[0] - 1)
            self.pca = PCA(n_components=n_components)
            X_final = self.pca.fit_transform(X_selected)
            self.final_features = [f"PC_{i+1}" for i in range(n_components)]
        else:
            X_final = X_selected
            self.final_features = X_selected.columns.tolist()
        
        self.is_fitted = True
        
        logger.info(f"Pipeline fitted successfully. Final features: {len(self.final_features)}")
        
        return self
        
    def transform(self, X: pd.DataFrame) -> pd.DataFrame:
        """
        Transform features using fitted pipeline.
        
        Args:
            X: Features to transform
            
        Returns:
            Transformed features DataFrame
        """
        if not self.is_fitted:
            raise ValueError("Pipeline must be fitted before transform")
            
        logger.info("Transforming features through pipeline")
        
        # Step 1: Remove correlated features (using same features as training)
        X_reduced = X[[col for col in X.columns if col not in self.correlation_analyzer.features_to_remove]]
        
        # Step 2: Scale features
        X_scaled = self.scaler.transform(X_reduced)
        
        # Step 3: Select features
        X_selected = self.feature_selector.transform(X_scaled)
        
        # Step 4: Apply PCA if used
        if self.pca is not None:
            X_final = self.pca.transform(X_selected)
            return pd.DataFrame(X_final, columns=self.final_features, index=X.index)
        else:
            return X_selected
        
    def fit_transform(self, X: pd.DataFrame, y: pd.Series) -> pd.DataFrame:
        """Fit pipeline and transform features in one step."""
        return self.fit(X, y).transform(X)
        
    def get_feature_importance(self) -> pd.DataFrame:
        """
        Get feature importance from the feature selector.
        
        Returns:
            DataFrame with feature importance scores
        """
        if not self.is_fitted:
            raise ValueError("Pipeline must be fitted before getting feature importance")
            
        return self.feature_selector.get_feature_importance()
        
    def get_correlation_analysis(self) -> Dict[str, Any]:
        """
        Get correlation analysis results.
        
        Returns:
            Dictionary with correlation analysis details
        """
        if not self.is_fitted:
            raise ValueError("Pipeline must be fitted before getting correlation analysis")
            
        return {
            "correlated_pairs": self.correlation_analyzer.correlated_pairs,
            "features_removed": list(self.correlation_analyzer.features_to_remove),
            "removal_threshold": self.correlation_analyzer.threshold
        }
        
    def get_pipeline_summary(self) -> Dict[str, Any]:
        """
        Get comprehensive summary of the fitted pipeline.
        
        Returns:
            Dictionary with pipeline configuration and results
        """
        if not self.is_fitted:
            raise ValueError("Pipeline must be fitted before getting summary")
            
        return {
            "config": {
                "correlation_threshold": self.config.correlation_threshold,
                "mutual_info_k": self.config.mutual_info_k,
                "use_robust_scaling": self.config.use_robust_scaling,
                "apply_pca": self.config.apply_pca,
                "pca_components": self.config.pca_components
            },
            "correlation_analysis": self.get_correlation_analysis(),
            "feature_importance": self.get_feature_importance().to_dict('records'),
            "final_features": self.final_features,
            "total_features_removed": len(self.correlation_analyzer.features_to_remove)
        }
```
