## Supplementary material for "Machine Learning-Driven Prediction of TLR4 Binding Affinity: A Comprehensive Molecular Feature Analysis for Drug Discovery": Codebase: gnn_models.py

```
"""
Graph Neural Network models for molecular binding prediction.

This module implements various GNN architectures for predicting
TLR4 binding affinities from molecular graph representations.
"""

from abc import ABC, abstractmethod
from typing import Dict, List, Optional, Union, Any, Tuple, NamedTuple
import pandas as pd
import numpy as np
import logging
from pathlib import Path
import joblib
from datetime import datetime

logger = logging.getLogger(__name__)

# PyTorch and PyTorch Geometric imports with error handling
try:
    import torch
    import torch.nn as nn
    import torch.nn.functional as F
    from torch.utils.data import Dataset, DataLoader
    TORCH_AVAILABLE = True
except ImportError:
    logger.warning("PyTorch not available. GNN models will be limited.")
    TORCH_AVAILABLE = False
    # Define dummy classes for when PyTorch is not available
    class torch:
        class Tensor:
            pass
        class device:
            def __init__(self, device_str):
                self.device_str = device_str
            def type(self):
                return self.device_str
        class float:
            pass
        class long:
            pass
        @staticmethod
        def tensor(data, dtype=None):
            return data
        @staticmethod
        def is_available():
            return False
        @staticmethod
        def cuda():
            return False
    class nn:
        class Module:
            def __init__(self):
                pass
            def train(self):
                pass
            def eval(self):
                pass
            def to(self, device):
                return self
        class Linear:
            def __init__(self, in_features, out_features):
                pass
        class GRUCell:
            def __init__(self, input_size, hidden_size):
                pass
        class BatchNorm1d:
            def __init__(self, num_features):
                pass
        class ModuleList:
            def __init__(self, modules=None):
                self.modules = modules or []
            def append(self, module):
                self.modules.append(module)
            def __len__(self):
                return len(self.modules)
            def __getitem__(self, idx):
                return self.modules[idx]
    class F:
        @staticmethod
        def mse_loss(pred, target):
            return 0.0
        @staticmethod
        def relu(x):
            return x
        @staticmethod
        def dropout(x, p, training):
            return x
    class Dataset:
        pass
    class DataLoader:
        def __init__(self, dataset, batch_size=1, shuffle=False, num_workers=0):
            self.dataset = dataset
            self.batch_size = batch_size
            self.shuffle = shuffle
            self.num_workers = num_workers
        def __iter__(self):
            return iter([])
        def __len__(self):
            return 0

try:
    import torch_geometric
    from torch_geometric.data import Data, Batch
    from torch_geometric.nn import GCNConv, GATConv, SAGEConv, global_mean_pool, global_max_pool, global_add_pool
    from torch_geometric.nn import MessagePassing
    from torch_geometric.utils import to_networkx, from_networkx
    TORCH_GEOMETRIC_AVAILABLE = True
except ImportError:
    logger.warning("PyTorch Geometric not available. GNN models will be limited.")
    TORCH_GEOMETRIC_AVAILABLE = False
    # Define dummy classes for when PyTorch Geometric is not available
    class Data:
        def __init__(self, **kwargs):
            for key, value in kwargs.items():
                setattr(self, key, value)
    class Batch:
        pass
    class GCNConv:
        def __init__(self, *args, **kwargs):
            pass
    class GATConv:
        def __init__(self, *args, **kwargs):
            pass
    class SAGEConv:
        def __init__(self, *args, **kwargs):
            pass
    def global_mean_pool(x, batch):
        return x.mean(dim=0, keepdim=True)
    def global_max_pool(x, batch):
        return x.max(dim=0, keepdim=True)[0]
    def global_add_pool(x, batch):
        return x.sum(dim=0, keepdim=True)
    class MessagePassing:
        pass
    def to_networkx(data, to_undirected=True):
        return None
    def from_networkx(G):
        return None

# RDKit for molecular graph construction
try:
    from rdkit import Chem
    from rdkit.Chem import rdMolDescriptors, Descriptors
    from rdkit.Chem.rdMolDescriptors import GetMorganFingerprintAsBitVect
    RDKIT_AVAILABLE = True
except ImportError:
    logger.warning("RDKit not available. Molecular graph construction will be limited.")
    RDKIT_AVAILABLE = False


class MolecularGraph(NamedTuple):
    """Container for molecular graph data."""
    node_features: np.ndarray
    edge_index: np.ndarray
    edge_attr: Optional[np.ndarray]
    num_nodes: int
    num_edges: int
    compound_name: str


class GraphDataset(Dataset):
    """PyTorch Dataset for molecular graphs."""
    
    def __init__(self, graphs: List[MolecularGraph], targets: Optional[List[float]] = None):
        """
        Initialize graph dataset.
        
        Args:
            graphs: List of molecular graphs
            targets: Optional list of target values (binding affinities)
        """
        self.graphs = graphs
        self.targets = targets if targets is not None else [0.0] * len(graphs)
        
    def __len__(self) -> int:
        return len(self.graphs)
    
    def __getitem__(self, idx: int) -> Union[Data, Tuple[Data, float]]:
        """Get graph and target at index."""
        graph = self.graphs[idx]
        
        # Convert to PyTorch Geometric Data object
        data = Data(
            x=torch.tensor(graph.node_features, dtype=torch.float),
            edge_index=torch.tensor(graph.edge_index, dtype=torch.long),
            edge_attr=torch.tensor(graph.edge_attr, dtype=torch.float) if graph.edge_attr is not None else None,
            y=torch.tensor([self.targets[idx]], dtype=torch.float),
            num_nodes=graph.num_nodes
        )
        
        return data


class MolecularGraphBuilder:
    """Builds molecular graphs from PDBQT files and RDKit molecules."""
    
    def __init__(self, use_3d_coords: bool = True, max_atoms: int = 200):
        """
        Initialize molecular graph builder.
        
        Args:
            use_3d_coords: Whether to use 3D coordinates for node features
            max_atoms: Maximum number of atoms to include in graph
        """
        self.use_3d_coords = use_3d_coords
        self.max_atoms = max_atoms
        
    def build_graph_from_pdbqt(self, pdbqt_path: str) -> Optional[MolecularGraph]:
        """
        Build molecular graph from PDBQT file.
        
        Args:
            pdbqt_path: Path to PDBQT file
            
        Returns:
            MolecularGraph object or None if parsing fails
        """
        try:
            # Read PDBQT file and extract coordinates
            coords, atom_types = self._parse_pdbqt(pdbqt_path)
            
            if coords is None or len(coords) == 0:
                logger.warning(f"No valid coordinates found in {pdbqt_path}")
                return None
            
            # Limit number of atoms
            if len(coords) > self.max_atoms:
                logger.warning(f"Truncating {len(coords)} atoms to {self.max_atoms}")
                coords = coords[:self.max_atoms]
                atom_types = atom_types[:self.max_atoms]
            
            # Build molecular graph
            return self._build_graph_from_coords(coords, atom_types, Path(pdbqt_path).stem)
            
        except Exception as e:
            logger.error(f"Error building graph from {pdbqt_path}: {str(e)}")
            return None
    
    def build_graph_from_smiles(self, smiles: str, compound_name: str = "unknown") -> Optional[MolecularGraph]:
        """
        Build molecular graph from SMILES string.
        
        Args:
            smiles: SMILES string
            compound_name: Name of the compound
            
        Returns:
            MolecularGraph object or None if parsing fails
        """
        if not RDKIT_AVAILABLE:
            logger.error("RDKit not available for SMILES parsing")
            return None
            
        try:
            mol = Chem.MolFromSmiles(smiles)
            if mol is None:
                logger.warning(f"Failed to parse SMILES: {smiles}")
                return None
            
            # Generate 3D coordinates
            mol = Chem.AddHs(mol)
            from rdkit.Chem import AllChem
            AllChem.EmbedMolecule(mol)
            AllChem.MMFFOptimizeMolecule(mol)
            
            # Extract coordinates and atom types
            coords = mol.GetConformer().GetPositions()
            atom_types = [atom.GetAtomicNum() for atom in mol.GetAtoms()]
            
            return self._build_graph_from_coords(coords, atom_types, compound_name)
            
        except Exception as e:
            logger.error(f"Error building graph from SMILES {smiles}: {str(e)}")
            return None
    
    def _parse_pdbqt(self, pdbqt_path: str) -> Tuple[Optional[np.ndarray], Optional[List[int]]]:
        """Parse PDBQT file to extract coordinates and atom types."""
        coords = []
        atom_types = []
        
        try:
            with open(pdbqt_path, 'r') as f:
                for line in f:
                    if line.startswith('ATOM') or line.startswith('HETATM'):
                        # Extract coordinates (columns 30-54)
                        x = float(line[30:38].strip())
                        y = float(line[38:46].strip())
                        z = float(line[46:54].strip())
                        coords.append([x, y, z])
                        
                        # Extract atom type (column 77)
                        atom_type = line[77:79].strip()
                        atom_types.append(self._atom_type_to_number(atom_type))
            
            return np.array(coords), atom_types
            
        except Exception as e:
            logger.error(f"Error parsing PDBQT file {pdbqt_path}: {str(e)}")
            return None, None
    
    def _atom_type_to_number(self, atom_type: str) -> int:
        """Convert atom type string to atomic number."""
        atom_type_map = {
            'C': 6, 'N': 7, 'O': 8, 'S': 16, 'P': 15,
            'H': 1, 'F': 9, 'Cl': 17, 'Br': 35, 'I': 53
        }
        return atom_type_map.get(atom_type, 6)  # Default to carbon
    
    def _build_graph_from_coords(self, coords: np.ndarray, atom_types: List[int], 
                                compound_name: str) -> MolecularGraph:
        """Build molecular graph from coordinates and atom types."""
        num_atoms = len(coords)
        
        # Create node features
        node_features = self._create_node_features(coords, atom_types)
        
        # Create edge index and attributes
        edge_index, edge_attr = self._create_edges(coords, atom_types)
        
        return MolecularGraph(
            node_features=node_features,
            edge_index=edge_index,
            edge_attr=edge_attr,
            num_nodes=num_atoms,
            num_edges=edge_index.shape[1],
            compound_name=compound_name
        )
    
    def _create_node_features(self, coords: np.ndarray, atom_types: List[int]) -> np.ndarray:
        """Create node features from coordinates and atom types."""
        num_atoms = len(coords)
        feature_dim = 9  # atomic_num, x, y, z, degree, formal_charge, is_aromatic, is_ring, is_metal
        
        features = np.zeros((num_atoms, feature_dim))
        
        for i, (coord, atom_type) in enumerate(zip(coords, atom_types)):
            features[i, 0] = atom_type  # atomic number
            features[i, 1:4] = coord    # 3D coordinates
            features[i, 4] = 0  # degree (would need molecular graph to calculate)
            features[i, 5] = 0  # formal charge
            features[i, 6] = 0  # is_aromatic
            features[i, 7] = 0  # is_ring
            features[i, 8] = 1 if atom_type > 20 else 0  # is_metal (simplified)
        
        return features
    
    def _create_edges(self, coords: np.ndarray, atom_types: List[int]) -> Tuple[np.ndarray, np.ndarray]:
        """Create edges based on distance threshold."""
        num_atoms = len(coords)
        edge_threshold = 2.0  # Angstroms
        
        edges = []
        edge_attrs = []
        
        for i in range(num_atoms):
            for j in range(i + 1, num_atoms):
                dist = np.linalg.norm(coords[i] - coords[j])
                if dist < edge_threshold:
                    edges.append([i, j])
                    edges.append([j, i])  # Undirected graph
                    edge_attrs.append([dist, 1.0])  # distance and bond type
                    edge_attrs.append([dist, 1.0])
        
        if not edges:
            # If no edges found, create a fully connected graph
            for i in range(num_atoms):
                for j in range(i + 1, num_atoms):
                    dist = np.linalg.norm(coords[i] - coords[j])
                    edges.append([i, j])
                    edges.append([j, i])
                    edge_attrs.append([dist, 1.0])
                    edge_attrs.append([dist, 1.0])
        
        edge_index = np.array(edges).T if edges else np.array([[0], [0]])
        edge_attr = np.array(edge_attrs) if edge_attrs else np.array([[0.0, 1.0]])
        
        return edge_index, edge_attr


class GNNModelInterface(ABC):
    """Abstract interface for GNN models."""
    
    @abstractmethod
    def forward(self, data: Data) -> torch.Tensor:
        """Forward pass through the model."""
        pass
    
    @abstractmethod
    def get_embeddings(self, data: Data) -> torch.Tensor:
        """Get node or graph embeddings."""
        pass


class GraphConvModel(nn.Module, GNNModelInterface):
    """Graph Convolutional Network for molecular property prediction."""
    
    def __init__(self, input_dim: int, hidden_dim: int = 64, output_dim: int = 1, 
                 num_layers: int = 3, dropout: float = 0.1):
        """
        Initialize GraphConv model.
        
        Args:
            input_dim: Input feature dimension
            hidden_dim: Hidden layer dimension
            output_dim: Output dimension (1 for regression)
            num_layers: Number of GCN layers
            dropout: Dropout rate
        """
        super().__init__()
        
        if not TORCH_GEOMETRIC_AVAILABLE:
            raise RuntimeError("PyTorch Geometric not available for GraphConv model")
        
        self.input_dim = input_dim
        self.hidden_dim = hidden_dim
        self.output_dim = output_dim
        self.num_layers = num_layers
        self.dropout = dropout
        
        # GCN layers
        self.convs = nn.ModuleList()
        self.convs.append(GCNConv(input_dim, hidden_dim))
        
        for _ in range(num_layers - 2):
            self.convs.append(GCNConv(hidden_dim, hidden_dim))
        
        self.convs.append(GCNConv(hidden_dim, hidden_dim))
        
        # Output layer
        self.out = nn.Linear(hidden_dim, output_dim)
        
        # Batch normalization
        self.batch_norms = nn.ModuleList()
        for _ in range(num_layers):
            self.batch_norms.append(nn.BatchNorm1d(hidden_dim))
        
    def forward(self, data: Data) -> torch.Tensor:
        """Forward pass through the model."""
        x, edge_index, batch = data.x, data.edge_index, data.batch
        
        # Apply GCN layers
        for i, conv in enumerate(self.convs):
            x = conv(x, edge_index)
            x = self.batch_norms[i](x)
            x = F.relu(x)
            x = F.dropout(x, p=self.dropout, training=self.training)
        
        # Global pooling
        x = global_mean_pool(x, batch)
        
        # Output layer
        x = self.out(x)
        
        return x
    
    def get_embeddings(self, data: Data) -> torch.Tensor:
        """Get graph-level embeddings."""
        x, edge_index, batch = data.x, data.edge_index, data.batch
        
        # Apply GCN layers
        for i, conv in enumerate(self.convs):
            x = conv(x, edge_index)
            x = self.batch_norms[i](x)
            x = F.relu(x)
            x = F.dropout(x, p=self.dropout, training=self.training)
        
        # Global pooling
        x = global_mean_pool(x, batch)
        
        return x


class MPNNModel(nn.Module, GNNModelInterface):
    """Message Passing Neural Network for molecular property prediction."""
    
    def __init__(self, input_dim: int, hidden_dim: int = 64, output_dim: int = 1,
                 num_layers: int = 3, dropout: float = 0.1):
        """
        Initialize MPNN model.
        
        Args:
            input_dim: Input feature dimension
            hidden_dim: Hidden layer dimension
            output_dim: Output dimension (1 for regression)
            num_layers: Number of message passing layers
            dropout: Dropout rate
        """
        super().__init__()
        
        if not TORCH_GEOMETRIC_AVAILABLE:
            raise RuntimeError("PyTorch Geometric not available for MPNN model")
        
        self.input_dim = input_dim
        self.hidden_dim = hidden_dim
        self.output_dim = output_dim
        self.num_layers = num_layers
        self.dropout = dropout
        
        # Input projection
        self.input_proj = nn.Linear(input_dim, hidden_dim)
        
        # Message passing layers
        self.message_layers = nn.ModuleList()
        self.update_layers = nn.ModuleList()
        
        for _ in range(num_layers):
            self.message_layers.append(nn.Linear(hidden_dim * 2 + 2, hidden_dim))  # +2 for edge features
            self.update_layers.append(nn.GRUCell(hidden_dim, hidden_dim))
        
        # Output layer
        self.out = nn.Linear(hidden_dim, output_dim)
        
    def forward(self, data: Data) -> torch.Tensor:
        """Forward pass through the model."""
        x, edge_index, edge_attr, batch = data.x, data.edge_index, data.edge_attr, data.batch
        
        # Input projection
        x = self.input_proj(x)
        
        # Message passing
        for i in range(self.num_layers):
            # Create messages
            row, col = edge_index
            edge_features = torch.cat([x[row], x[col], edge_attr], dim=1)
            messages = self.message_layers[i](edge_features)
            
            # Aggregate messages
            message_aggr = torch.zeros_like(x)
            message_aggr.scatter_add_(0, col.unsqueeze(-1).expand_as(messages), messages)
            
            # Update node features
            x = self.update_layers[i](message_aggr, x)
            x = F.dropout(x, p=self.dropout, training=self.training)
        
        # Global pooling
        x = global_mean_pool(x, batch)
        
        # Output layer
        x = self.out(x)
        
        return x
    
    def get_embeddings(self, data: Data) -> torch.Tensor:
        """Get graph-level embeddings."""
        x, edge_index, edge_attr, batch = data.x, data.edge_index, data.edge_attr, data.batch
        
        # Input projection
        x = self.input_proj(x)
        
        # Message passing
        for i in range(self.num_layers):
            # Create messages
            row, col = edge_index
            edge_features = torch.cat([x[row], x[col], edge_attr], dim=1)
            messages = self.message_layers[i](edge_features)
            
            # Aggregate messages
            message_aggr = torch.zeros_like(x)
            message_aggr.scatter_add_(0, col.unsqueeze(-1).expand_as(messages), messages)
            
            # Update node features
            x = self.update_layers[i](message_aggr, x)
            x = F.dropout(x, p=self.dropout, training=self.training)
        
        # Global pooling
        x = global_mean_pool(x, batch)
        
        return x


class AttentiveFPModel(nn.Module, GNNModelInterface):
    """AttentiveFP model for molecular property prediction."""
    
    def __init__(self, input_dim: int, hidden_dim: int = 64, output_dim: int = 1,
                 num_layers: int = 3, dropout: float = 0.1):
        """
        Initialize AttentiveFP model.
        
        Args:
            input_dim: Input feature dimension
            hidden_dim: Hidden layer dimension
            output_dim: Output dimension (1 for regression)
            num_layers: Number of attention layers
            dropout: Dropout rate
        """
        super().__init__()
        
        if not TORCH_GEOMETRIC_AVAILABLE:
            raise RuntimeError("PyTorch Geometric not available for AttentiveFP model")
        
        self.input_dim = input_dim
        self.hidden_dim = hidden_dim
        self.output_dim = output_dim
        self.num_layers = num_layers
        self.dropout = dropout
        
        # Input projection
        self.input_proj = nn.Linear(input_dim, hidden_dim)
        
        # Attention layers
        self.attention_layers = nn.ModuleList()
        for _ in range(num_layers):
            self.attention_layers.append(GATConv(hidden_dim, hidden_dim, heads=4, 
                                               concat=False, dropout=dropout))
        
        # Output layer
        self.out = nn.Linear(hidden_dim, output_dim)
        
    def forward(self, data: Data) -> torch.Tensor:
        """Forward pass through the model."""
        x, edge_index, batch = data.x, data.edge_index, data.batch
        
        # Input projection
        x = self.input_proj(x)
        
        # Apply attention layers
        for attention in self.attention_layers:
            x = attention(x, edge_index)
            x = F.relu(x)
            x = F.dropout(x, p=self.dropout, training=self.training)
        
        # Global pooling
        x = global_mean_pool(x, batch)
        
        # Output layer
        x = self.out(x)
        
        return x
    
    def get_embeddings(self, data: Data) -> torch.Tensor:
        """Get graph-level embeddings."""
        x, edge_index, batch = data.x, data.edge_index, data.batch
        
        # Input projection
        x = self.input_proj(x)
        
        # Apply attention layers
        for attention in self.attention_layers:
            x = attention(x, edge_index)
            x = F.relu(x)
            x = F.dropout(x, p=self.dropout, training=self.training)
        
        # Global pooling
        x = global_mean_pool(x, batch)
        
        return x


class GNNTrainer:
    """Trainer for Graph Neural Network models."""
    
    def __init__(self, model_type: str = "graphconv", device: str = "auto"):
        """
        Initialize GNN trainer.
        
        Args:
            model_type: Type of GNN model ("graphconv", "mpnn", "attentivefp")
            device: Device to use for training ("auto", "cpu", "cuda")
        """
        if not TORCH_AVAILABLE:
            raise RuntimeError("PyTorch not available for GNN training")
        
        self.model_type = model_type
        self.device = self._get_device(device)
        self.model = None
        self.optimizer = None
        self.scheduler = None
        
    def _get_device(self, device: str) -> torch.device:
        """Get appropriate device for training."""
        if device == "auto":
            if torch.cuda.is_available():
                return torch.device("cuda")
            else:
                return torch.device("cpu")
        else:
            return torch.device(device)
    
    def create_model(self, input_dim: int, hidden_dim: int = 64, 
                    output_dim: int = 1, **kwargs) -> nn.Module:
        """Create GNN model based on type."""
        if self.model_type == "graphconv":
            return GraphConvModel(input_dim, hidden_dim, output_dim, **kwargs)
        elif self.model_type == "mpnn":
            return MPNNModel(input_dim, hidden_dim, output_dim, **kwargs)
        elif self.model_type == "attentivefp":
            return AttentiveFPModel(input_dim, hidden_dim, output_dim, **kwargs)
        else:
            raise ValueError(f"Unknown model type: {self.model_type}")
    
    def train(self, train_loader: DataLoader, val_loader: Optional[DataLoader] = None,
              epochs: int = 100, lr: float = 0.001, weight_decay: float = 1e-4,
              patience: int = 20) -> Dict[str, List[float]]:
        """
        Train GNN model.
        
        Args:
            train_loader: Training data loader
            val_loader: Validation data loader (optional)
            epochs: Number of training epochs
            lr: Learning rate
            weight_decay: Weight decay for regularization
            patience: Early stopping patience
            
        Returns:
            Dictionary of training history
        """
        if self.model is None:
            raise ValueError("Model not created. Call create_model() first.")
        
        self.model.to(self.device)
        self.optimizer = torch.optim.Adam(self.model.parameters(), lr=lr, weight_decay=weight_decay)
        self.scheduler = torch.optim.lr_scheduler.ReduceLROnPlateau(
            self.optimizer, mode='min', factor=0.5, patience=10, verbose=True
        )
        
        history = {'train_loss': [], 'val_loss': []}
        best_val_loss = float('inf')
        patience_counter = 0
        
        for epoch in range(epochs):
            # Training
            train_loss = self._train_epoch(train_loader)
            history['train_loss'].append(train_loss)
            
            # Validation
            if val_loader is not None:
                val_loss = self._validate_epoch(val_loader)
                history['val_loss'].append(val_loss)
                
                # Early stopping
                if val_loss < best_val_loss:
                    best_val_loss = val_loss
                    patience_counter = 0
                else:
                    patience_counter += 1
                
                if patience_counter >= patience:
                    logger.info(f"Early stopping at epoch {epoch}")
                    break
                
                # Learning rate scheduling
                self.scheduler.step(val_loss)
            else:
                history['val_loss'].append(train_loss)
            
            if epoch % 10 == 0:
                logger.info(f"Epoch {epoch}: Train Loss = {train_loss:.4f}, "
                          f"Val Loss = {history['val_loss'][-1]:.4f}")
        
        return history
    
    def _train_epoch(self, train_loader: DataLoader) -> float:
        """Train for one epoch."""
        self.model.train()
        total_loss = 0.0
        
        for batch in train_loader:
            batch = batch.to(self.device)
            
            self.optimizer.zero_grad()
            pred = self.model(batch)
            loss = F.mse_loss(pred.squeeze(), batch.y.squeeze())
            loss.backward()
            self.optimizer.step()
            
            total_loss += loss.item()
        
        return total_loss / len(train_loader)
    
    def _validate_epoch(self, val_loader: DataLoader) -> float:
        """Validate for one epoch."""
        self.model.eval()
        total_loss = 0.0
        
        with torch.no_grad():
            for batch in val_loader:
                batch = batch.to(self.device)
                pred = self.model(batch)
                loss = F.mse_loss(pred.squeeze(), batch.y.squeeze())
                total_loss += loss.item()
        
        return total_loss / len(val_loader)
    
    def predict(self, data_loader: DataLoader) -> np.ndarray:
        """Make predictions on data loader."""
        if self.model is None:
            raise ValueError("Model not trained. Call train() first.")
        
        self.model.eval()
        predictions = []
        
        with torch.no_grad():
            for batch in data_loader:
                batch = batch.to(self.device)
                pred = self.model(batch)
                predictions.append(pred.cpu().numpy())
        
        return np.concatenate(predictions, axis=0)
    
    def save_model(self, path: str) -> None:
        """Save trained model."""
        if self.model is None:
            raise ValueError("No model to save")
        
        torch.save({
            'model_state_dict': self.model.state_dict(),
            'model_type': self.model_type,
            'optimizer_state_dict': self.optimizer.state_dict() if self.optimizer else None
        }, path)
        
        logger.info(f"Model saved to {path}")
    
    def load_model(self, path: str, input_dim: int, **kwargs) -> None:
        """Load trained model."""
        checkpoint = torch.load(path, map_location=self.device)
        
        self.model = self.create_model(input_dim, **kwargs)
        self.model.load_state_dict(checkpoint['model_state_dict'])
        self.model.to(self.device)
        
        logger.info(f"Model loaded from {path}")


class GNNModelTrainer:
    """Main GNN model trainer coordinator."""
    
    def __init__(self, models_dir: str = "models/gnn"):
        """
        Initialize GNN model trainer.
        
        Args:
            models_dir: Directory to save trained models
        """
        self.models_dir = Path(models_dir)
        self.models_dir.mkdir(parents=True, exist_ok=True)
        
        self.graph_builder = MolecularGraphBuilder()
        self.trained_models = {}
        self.training_results = {}
    
    def prepare_graph_data(self, pdbqt_files: List[str], 
                          binding_affinities: Optional[List[float]] = None) -> GraphDataset:
        """
        Prepare graph dataset from PDBQT files.
        
        Args:
            pdbqt_files: List of PDBQT file paths
            binding_affinities: Optional list of binding affinities
            
        Returns:
            GraphDataset object
        """
        logger.info(f"Building graphs from {len(pdbqt_files)} PDBQT files")
        
        graphs = []
        targets = binding_affinities if binding_affinities is not None else [0.0] * len(pdbqt_files)
        
        for pdbqt_file in pdbqt_files:
            graph = self.graph_builder.build_graph_from_pdbqt(pdbqt_file)
            if graph is not None:
                graphs.append(graph)
            else:
                logger.warning(f"Failed to build graph from {pdbqt_file}")
        
        logger.info(f"Successfully built {len(graphs)} graphs")
        return GraphDataset(graphs, targets[:len(graphs)])
    
    def train_models(self, train_dataset: GraphDataset, val_dataset: Optional[GraphDataset] = None,
                    model_types: List[str] = None, **kwargs) -> Dict[str, Any]:
        """
        Train multiple GNN models.
        
        Args:
            train_dataset: Training dataset
            val_dataset: Validation dataset (optional)
            model_types: List of model types to train
            **kwargs: Additional training parameters
            
        Returns:
            Dictionary of trained models
        """
        if model_types is None:
            model_types = ["graphconv", "mpnn", "attentivefp"]
        
        logger.info(f"Training {len(model_types)} GNN models")
        
        # Create data loaders
        train_loader = DataLoader(train_dataset, batch_size=kwargs.get('batch_size', 32), 
                                 shuffle=True, num_workers=0)
        
        val_loader = None
        if val_dataset is not None:
            val_loader = DataLoader(val_dataset, batch_size=kwargs.get('batch_size', 32), 
                                   shuffle=False, num_workers=0)
        
        # Get input dimension from first graph
        if len(train_dataset) == 0:
            raise ValueError("Training dataset is empty")
        
        sample_graph = train_dataset[0]
        input_dim = sample_graph.x.shape[1]
        
        for model_type in model_types:
            try:
                logger.info(f"Training {model_type} model")
                
                # Create trainer
                trainer = GNNTrainer(model_type=model_type, device=kwargs.get('device', 'auto'))
                trainer.model = trainer.create_model(input_dim, **kwargs)
                
                # Train model
                history = trainer.train(train_loader, val_loader, **kwargs)
                
                # Store results
                self.trained_models[model_type] = trainer
                self.training_results[model_type] = {
                    'trainer': trainer,
                    'history': history,
                    'input_dim': input_dim
                }
                
                # Save model
                model_path = self.models_dir / f"{model_type}_model.pth"
                trainer.save_model(str(model_path))
                
                logger.info(f"{model_type} training completed successfully")
                
            except Exception as e:
                logger.error(f"Error training {model_type}: {str(e)}")
                continue
        
        logger.info(f"Successfully trained {len(self.trained_models)} GNN models")
        return self.trained_models
    
    def evaluate_models(self, test_dataset: GraphDataset) -> Dict[str, Dict]:
        """
        Evaluate all trained models on test dataset.
        
        Args:
            test_dataset: Test dataset
            
        Returns:
            Dictionary of evaluation results
        """
        logger.info("Evaluating GNN models on test set")
        
        test_loader = DataLoader(test_dataset, batch_size=32, shuffle=False, num_workers=0)
        evaluation_results = {}
        
        for model_name, trainer in self.trained_models.items():
            try:
                # Make predictions
                predictions = trainer.predict(test_loader)
                
                # Calculate metrics
                true_values = [target for target in test_dataset.targets]
                metrics = self._calculate_metrics(true_values, predictions.flatten())
                
                evaluation_results[model_name] = {
                    'predictions': predictions,
                    'metrics': metrics
                }
                
                logger.info(f"{model_name} evaluation completed")
                
            except Exception as e:
                logger.error(f"Error evaluating {model_name}: {str(e)}")
                continue
        
        return evaluation_results
    
    def _calculate_metrics(self, y_true: List[float], y_pred: np.ndarray) -> Dict[str, float]:
        """Calculate regression metrics."""
        y_true = np.array(y_true)
        y_pred = np.array(y_pred)
        
        mse = np.mean((y_true - y_pred) ** 2)
        rmse = np.sqrt(mse)
        mae = np.mean(np.abs(y_true - y_pred))
        
        # R² score
        ss_res = np.sum((y_true - y_pred) ** 2)
        ss_tot = np.sum((y_true - np.mean(y_true)) ** 2)
        r2 = 1 - (ss_res / ss_tot) if ss_tot != 0 else 0
        
        return {
            'mse': mse,
            'rmse': rmse,
            'mae': mae,
            'r2': r2
        }
    
    def get_best_model(self, metric: str = 'r2') -> Tuple[str, Any]:
        """
        Get the best performing model based on specified metric.
        
        Args:
            metric: Metric to use for model selection
            
        Returns:
            Tuple of (model_name, trainer)
        """
        if not self.training_results:
            raise ValueError("No models have been trained yet")
        
        best_model_name = None
        best_score = float('-inf') if metric == 'r2' else float('inf')
        
        for model_name, results in self.training_results.items():
            # Get validation score from history
            if 'val_loss' in results['history']:
                val_losses = results['history']['val_loss']
                if val_losses:
                    score = -min(val_losses)  # Convert loss to score
                    if metric == 'r2':
                        if score > best_score:
                            best_score = score
                            best_model_name = model_name
                    else:  # For loss-based metrics
                        if score < best_score:
                            best_score = score
                            best_model_name = model_name
        
        if best_model_name is None:
            raise ValueError(f"No model found with metric {metric}")
        
        return best_model_name, self.trained_models[best_model_name]
```
