## Supplementary material for "Machine Learning-Driven Prediction of TLR4 Binding Affinity: A Comprehensive Molecular Feature Analysis for Drug Discovery": Codebase: gnn_visualization.py

```
"""
Graph visualization and analysis tools for GNN models.

This module provides utilities for visualizing molecular graphs,
analyzing GNN predictions, and extracting interpretable features.
"""

import numpy as np
import pandas as pd
import matplotlib.pyplot as plt
import seaborn as sns
from typing import Dict, List, Optional, Tuple, Any
import logging
from pathlib import Path

logger = logging.getLogger(__name__)

# Visualization imports with error handling
try:
    import networkx as nx
    NETWORKX_AVAILABLE = True
except ImportError:
    logger.warning("NetworkX not available. Graph visualization will be limited.")
    NETWORKX_AVAILABLE = False

try:
    import plotly.graph_objects as go
    import plotly.express as px
    from plotly.subplots import make_subplots
    PLOTLY_AVAILABLE = True
except ImportError:
    logger.warning("Plotly not available. Interactive visualization will be limited.")
    PLOTLY_AVAILABLE = False

try:
    from rdkit import Chem
    from rdkit.Chem import Draw, rdMolDescriptors
    from rdkit.Chem.Draw import rdMolDraw2D
    RDKIT_AVAILABLE = True
except ImportError:
    logger.warning("RDKit not available. Molecular visualization will be limited.")
    RDKIT_AVAILABLE = False

try:
    import torch
    from torch_geometric.data import Data
    from torch_geometric.utils import to_networkx, from_networkx
    TORCH_GEOMETRIC_AVAILABLE = True
except ImportError:
    logger.warning("PyTorch Geometric not available. Graph conversion will be limited.")
    TORCH_GEOMETRIC_AVAILABLE = False


class GraphVisualizer:
    """Visualization tools for molecular graphs and GNN analysis."""
    
    def __init__(self, output_dir: str = "results/gnn_visualizations"):
        """
        Initialize graph visualizer.
        
        Args:
            output_dir: Directory to save visualization outputs
        """
        self.output_dir = Path(output_dir)
        self.output_dir.mkdir(parents=True, exist_ok=True)
        
    def visualize_molecular_graph(self, graph_data: Any, compound_name: str = "unknown",
                                save_path: Optional[str] = None) -> Optional[plt.Figure]:
        """
        Visualize molecular graph structure.
        
        Args:
            graph_data: MolecularGraph or PyTorch Geometric Data object
            compound_name: Name of the compound
            save_path: Optional path to save the figure
            
        Returns:
            Matplotlib figure or None if visualization fails
        """
        try:
            if TORCH_GEOMETRIC_AVAILABLE and isinstance(graph_data, Data):
                # Convert PyTorch Geometric Data to NetworkX
                nx_graph = to_networkx(graph_data, to_undirected=True)
            else:
                # Convert from MolecularGraph
                nx_graph = self._molecular_graph_to_networkx(graph_data)
            
            if not NETWORKX_AVAILABLE:
                logger.warning("NetworkX not available for graph visualization")
                return None
            
            # Create figure
            fig, ax = plt.subplots(1, 1, figsize=(10, 8))
            
            # Get node positions using spring layout
            pos = nx.spring_layout(nx_graph, k=1, iterations=50)
            
            # Draw nodes
            node_colors = self._get_node_colors(nx_graph)
            nx.draw_networkx_nodes(nx_graph, pos, node_color=node_colors, 
                                 node_size=300, alpha=0.8, ax=ax)
            
            # Draw edges
            nx.draw_networkx_edges(nx_graph, pos, alpha=0.5, ax=ax)
            
            # Draw labels
            nx.draw_networkx_labels(nx_graph, pos, font_size=8, ax=ax)
            
            ax.set_title(f"Molecular Graph: {compound_name}")
            ax.axis('off')
            
            if save_path:
                fig.savefig(save_path, dpi=300, bbox_inches='tight')
                logger.info(f"Graph visualization saved to {save_path}")
            
            return fig
            
        except Exception as e:
            logger.error(f"Error visualizing molecular graph: {str(e)}")
            return None
    
    def visualize_attention_weights(self, attention_weights: np.ndarray, 
                                  graph_data: Any, compound_name: str = "unknown",
                                  save_path: Optional[str] = None) -> Optional[plt.Figure]:
        """
        Visualize attention weights from AttentiveFP model.
        
        Args:
            attention_weights: Attention weights array
            graph_data: MolecularGraph or PyTorch Geometric Data object
            compound_name: Name of the compound
            save_path: Optional path to save the figure
            
        Returns:
            Matplotlib figure or None if visualization fails
        """
        try:
            if TORCH_GEOMETRIC_AVAILABLE and isinstance(graph_data, Data):
                nx_graph = to_networkx(graph_data, to_undirected=True)
            else:
                nx_graph = self._molecular_graph_to_networkx(graph_data)
            
            if not NETWORKX_AVAILABLE:
                logger.warning("NetworkX not available for attention visualization")
                return None
            
            # Create figure
            fig, (ax1, ax2) = plt.subplots(1, 2, figsize=(15, 6))
            
            # Original graph
            pos = nx.spring_layout(nx_graph, k=1, iterations=50)
            nx.draw_networkx_nodes(nx_graph, pos, node_color='lightblue', 
                                 node_size=300, alpha=0.8, ax=ax1)
            nx.draw_networkx_edges(nx_graph, pos, alpha=0.5, ax=ax1)
            nx.draw_networkx_labels(nx_graph, pos, font_size=8, ax=ax1)
            ax1.set_title(f"Original Graph: {compound_name}")
            ax1.axis('off')
            
            # Attention-weighted graph
            node_attention = np.mean(attention_weights, axis=1) if attention_weights.ndim > 1 else attention_weights
            node_colors = self._normalize_colors(node_attention)
            
            nx.draw_networkx_nodes(nx_graph, pos, node_color=node_colors, 
                                 node_size=300, alpha=0.8, ax=ax2)
            nx.draw_networkx_edges(nx_graph, pos, alpha=0.5, ax=ax2)
            nx.draw_networkx_labels(nx_graph, pos, font_size=8, ax=ax2)
            ax2.set_title(f"Attention Weights: {compound_name}")
            ax2.axis('off')
            
            # Add colorbar
            sm = plt.cm.ScalarMappable(cmap=plt.cm.Reds, 
                                     norm=plt.Normalize(vmin=node_attention.min(), 
                                                       vmax=node_attention.max()))
            sm.set_array([])
            plt.colorbar(sm, ax=ax2, shrink=0.8)
            
            if save_path:
                fig.savefig(save_path, dpi=300, bbox_inches='tight')
                logger.info(f"Attention visualization saved to {save_path}")
            
            return fig
            
        except Exception as e:
            logger.error(f"Error visualizing attention weights: {str(e)}")
            return None
    
    def plot_training_history(self, history: Dict[str, List[float]], 
                            model_name: str = "GNN Model",
                            save_path: Optional[str] = None) -> plt.Figure:
        """
        Plot training history for GNN models.
        
        Args:
            history: Training history dictionary
            model_name: Name of the model
            save_path: Optional path to save the figure
            
        Returns:
            Matplotlib figure
        """
        fig, (ax1, ax2) = plt.subplots(1, 2, figsize=(12, 5))
        
        epochs = range(1, len(history['train_loss']) + 1)
        
        # Plot loss
        ax1.plot(epochs, history['train_loss'], 'b-', label='Training Loss')
        if 'val_loss' in history and history['val_loss']:
            ax1.plot(epochs, history['val_loss'], 'r-', label='Validation Loss')
        ax1.set_xlabel('Epoch')
        ax1.set_ylabel('Loss')
        ax1.set_title(f'{model_name} - Training Loss')
        ax1.legend()
        ax1.grid(True, alpha=0.3)
        
        # Plot learning rate if available
        if 'lr' in history:
            ax2.plot(epochs, history['lr'], 'g-', label='Learning Rate')
            ax2.set_xlabel('Epoch')
            ax2.set_ylabel('Learning Rate')
            ax2.set_title(f'{model_name} - Learning Rate')
            ax2.legend()
            ax2.grid(True, alpha=0.3)
        else:
            ax2.axis('off')
        
        plt.tight_layout()
        
        if save_path:
            fig.savefig(save_path, dpi=300, bbox_inches='tight')
            logger.info(f"Training history plot saved to {save_path}")
        
        return fig
    
    def plot_model_comparison(self, results: Dict[str, Dict], 
                            metric: str = 'r2',
                            save_path: Optional[str] = None) -> plt.Figure:
        """
        Plot comparison of different GNN models.
        
        Args:
            results: Dictionary of model results
            metric: Metric to compare
            save_path: Optional path to save the figure
            
        Returns:
            Matplotlib figure
        """
        model_names = list(results.keys())
        metric_values = [results[name]['metrics'].get(metric, 0) for name in model_names]
        
        fig, ax = plt.subplots(1, 1, figsize=(10, 6))
        
        bars = ax.bar(model_names, metric_values, color='skyblue', alpha=0.7)
        ax.set_ylabel(metric.upper())
        ax.set_title(f'GNN Model Comparison - {metric.upper()}')
        ax.set_ylim(0, max(metric_values) * 1.1 if metric_values else 1)
        
        # Add value labels on bars
        for bar, value in zip(bars, metric_values):
            height = bar.get_height()
            ax.text(bar.get_x() + bar.get_width()/2., height + 0.01,
                   f'{value:.3f}', ha='center', va='bottom')
        
        plt.xticks(rotation=45)
        plt.tight_layout()
        
        if save_path:
            fig.savefig(save_path, dpi=300, bbox_inches='tight')
            logger.info(f"Model comparison plot saved to {save_path}")
        
        return fig
    
    def create_interactive_graph(self, graph_data: Any, compound_name: str = "unknown",
                               attention_weights: Optional[np.ndarray] = None,
                               save_path: Optional[str] = None) -> Optional[go.Figure]:
        """
        Create interactive graph visualization using Plotly.
        
        Args:
            graph_data: MolecularGraph or PyTorch Geometric Data object
            compound_name: Name of the compound
            attention_weights: Optional attention weights for coloring
            save_path: Optional path to save the HTML file
            
        Returns:
            Plotly figure or None if visualization fails
        """
        if not PLOTLY_AVAILABLE:
            logger.warning("Plotly not available for interactive visualization")
            return None
        
        try:
            if TORCH_GEOMETRIC_AVAILABLE and isinstance(graph_data, Data):
                nx_graph = to_networkx(graph_data, to_undirected=True)
            else:
                nx_graph = self._molecular_graph_to_networkx(graph_data)
            
            if not NETWORKX_AVAILABLE:
                logger.warning("NetworkX not available for interactive visualization")
                return None
            
            # Get node positions
            pos = nx.spring_layout(nx_graph, k=1, iterations=50)
            
            # Prepare node data
            node_x = [pos[node][0] for node in nx_graph.nodes()]
            node_y = [pos[node][1] for node in nx_graph.nodes()]
            node_text = [f"Node {node}" for node in nx_graph.nodes()]
            
            # Prepare edge data
            edge_x = []
            edge_y = []
            for edge in nx_graph.edges():
                x0, y0 = pos[edge[0]]
                x1, y1 = pos[edge[1]]
                edge_x.extend([x0, x1, None])
                edge_y.extend([y0, y1, None])
            
            # Create edge trace
            edge_trace = go.Scatter(x=edge_x, y=edge_y,
                                  line=dict(width=2, color='#888'),
                                  hoverinfo='none',
                                  mode='lines')
            
            # Create node trace
            node_trace = go.Scatter(x=node_x, y=node_y,
                                  mode='markers+text',
                                  hoverinfo='text',
                                  text=node_text,
                                  textposition="middle center",
                                  marker=dict(size=20,
                                            color='lightblue',
                                            line=dict(width=2, color='black')))
            
            # Add attention weights if provided
            if attention_weights is not None:
                node_attention = np.mean(attention_weights, axis=1) if attention_weights.ndim > 1 else attention_weights
                node_trace.marker.color = node_attention
                node_trace.marker.colorscale = 'Reds'
                node_trace.marker.showscale = True
                node_trace.marker.colorbar = dict(title="Attention Weight")
            
            # Create figure
            fig = go.Figure(data=[edge_trace, node_trace],
                          layout=go.Layout(title=f'Interactive Molecular Graph: {compound_name}',
                                         titlefont_size=16,
                                         showlegend=False,
                                         hovermode='closest',
                                         margin=dict(b=20,l=5,r=5,t=40),
                                         annotations=[ dict(
                                             text="Interactive molecular graph visualization",
                                             showarrow=False,
                                             xref="paper", yref="paper",
                                             x=0.005, y=-0.002,
                                             xanchor='left', yanchor='bottom',
                                             font=dict(color='black', size=12)
                                         )],
                                         xaxis=dict(showgrid=False, zeroline=False, showticklabels=False),
                                         yaxis=dict(showgrid=False, zeroline=False, showticklabels=False)))
            
            if save_path:
                fig.write_html(save_path)
                logger.info(f"Interactive graph saved to {save_path}")
            
            return fig
            
        except Exception as e:
            logger.error(f"Error creating interactive graph: {str(e)}")
            return None
    
    def _molecular_graph_to_networkx(self, graph_data: Any) -> Any:
        """Convert MolecularGraph to NetworkX graph."""
        if not NETWORKX_AVAILABLE:
            return None
        
        G = nx.Graph()
        
        # Add nodes
        for i in range(graph_data.num_nodes):
            G.add_node(i, features=graph_data.node_features[i])
        
        # Add edges
        for i in range(graph_data.edge_index.shape[1]):
            src, dst = graph_data.edge_index[:, i]
            G.add_edge(src, dst)
        
        return G
    
    def _get_node_colors(self, nx_graph: Any) -> List[str]:
        """Get node colors based on node features."""
        colors = []
        for node in nx_graph.nodes():
            # Simple coloring based on node degree
            degree = nx_graph.degree(node)
            if degree <= 2:
                colors.append('lightblue')
            elif degree <= 4:
                colors.append('lightgreen')
            else:
                colors.append('lightcoral')
        return colors
    
    def _normalize_colors(self, values: np.ndarray) -> np.ndarray:
        """Normalize values for color mapping."""
        if len(values) == 0:
            return values
        
        min_val = np.min(values)
        max_val = np.max(values)
        
        if max_val == min_val:
            return np.ones_like(values) * 0.5
        
        return (values - min_val) / (max_val - min_val)


class GraphFeatureExtractor:
    """Extract interpretable features from molecular graphs."""
    
    def __init__(self):
        """Initialize graph feature extractor."""
        pass
    
    def extract_graph_features(self, graph_data: Any) -> Dict[str, float]:
        """
        Extract interpretable features from molecular graph.
        
        Args:
            graph_data: MolecularGraph or PyTorch Geometric Data object
            
        Returns:
            Dictionary of graph features
        """
        try:
            if TORCH_GEOMETRIC_AVAILABLE and isinstance(graph_data, Data):
                nx_graph = to_networkx(graph_data, to_undirected=True)
            else:
                nx_graph = self._molecular_graph_to_networkx(graph_data)
            
            if not NETWORKX_AVAILABLE:
                logger.warning("NetworkX not available for feature extraction")
                return {}
            
            features = {}
            
            # Basic graph properties
            features['num_nodes'] = nx_graph.number_of_nodes()
            features['num_edges'] = nx_graph.number_of_edges()
            features['density'] = nx.density(nx_graph)
            
            # Connectivity features
            if nx.is_connected(nx_graph):
                features['diameter'] = nx.diameter(nx_graph)
                features['radius'] = nx.radius(nx_graph)
                features['average_path_length'] = nx.average_shortest_path_length(nx_graph)
            else:
                features['diameter'] = 0
                features['radius'] = 0
                features['average_path_length'] = 0
            
            # Centrality measures
            degree_centrality = nx.degree_centrality(nx_graph)
            features['max_degree_centrality'] = max(degree_centrality.values()) if degree_centrality else 0
            features['avg_degree_centrality'] = np.mean(list(degree_centrality.values())) if degree_centrality else 0
            
            # Clustering
            features['average_clustering'] = nx.average_clustering(nx_graph)
            features['transitivity'] = nx.transitivity(nx_graph)
            
            # Degree distribution
            degrees = [d for n, d in nx_graph.degree()]
            features['max_degree'] = max(degrees) if degrees else 0
            features['min_degree'] = min(degrees) if degrees else 0
            features['avg_degree'] = np.mean(degrees) if degrees else 0
            features['degree_std'] = np.std(degrees) if degrees else 0
            
            return features
            
        except Exception as e:
            logger.error(f"Error extracting graph features: {str(e)}")
            return {}
    
    def _molecular_graph_to_networkx(self, graph_data: Any) -> Any:
        """Convert MolecularGraph to NetworkX graph."""
        if not NETWORKX_AVAILABLE:
            return None
        
        G = nx.Graph()
        
        # Add nodes
        for i in range(graph_data.num_nodes):
            G.add_node(i, features=graph_data.node_features[i])
        
        # Add edges
        for i in range(graph_data.edge_index.shape[1]):
            src, dst = graph_data.edge_index[:, i]
            G.add_edge(src, dst)
        
        return G


class GNNModelAnalyzer:
    """Analyze and interpret GNN model predictions."""
    
    def __init__(self, output_dir: str = "results/gnn_analysis"):
        """
        Initialize GNN model analyzer.
        
        Args:
            output_dir: Directory to save analysis outputs
        """
        self.output_dir = Path(output_dir)
        self.output_dir.mkdir(parents=True, exist_ok=True)
        
        self.visualizer = GraphVisualizer(output_dir)
        self.feature_extractor = GraphFeatureExtractor()
    
    def analyze_predictions(self, predictions: np.ndarray, true_values: np.ndarray,
                          graph_data_list: List[Any], compound_names: List[str],
                          model_name: str = "GNN Model") -> Dict[str, Any]:
        """
        Analyze GNN model predictions and create visualizations.
        
        Args:
            predictions: Model predictions
            true_values: True target values
            graph_data_list: List of molecular graphs
            compound_names: List of compound names
            model_name: Name of the model
            
        Returns:
            Dictionary of analysis results
        """
        logger.info(f"Analyzing predictions for {model_name}")
        
        # Calculate prediction errors
        errors = predictions.flatten() - true_values
        abs_errors = np.abs(errors)
        
        # Create analysis results
        analysis = {
            'predictions': predictions,
            'true_values': true_values,
            'errors': errors,
            'abs_errors': abs_errors,
            'mae': np.mean(abs_errors),
            'rmse': np.sqrt(np.mean(errors**2)),
            'r2': 1 - np.sum(errors**2) / np.sum((true_values - np.mean(true_values))**2)
        }
        
        # Create visualizations
        self._plot_prediction_analysis(analysis, model_name)
        
        # Analyze worst predictions
        worst_indices = np.argsort(abs_errors)[-5:]  # Top 5 worst predictions
        self._analyze_worst_predictions(worst_indices, graph_data_list, 
                                      compound_names, predictions, true_values)
        
        return analysis
    
    def _plot_prediction_analysis(self, analysis: Dict[str, Any], model_name: str) -> None:
        """Create prediction analysis plots."""
        fig, ((ax1, ax2), (ax3, ax4)) = plt.subplots(2, 2, figsize=(15, 12))
        
        # Prediction vs True scatter plot
        ax1.scatter(analysis['true_values'], analysis['predictions'], alpha=0.6)
        ax1.plot([analysis['true_values'].min(), analysis['true_values'].max()],
                [analysis['true_values'].min(), analysis['true_values'].max()], 'r--')
        ax1.set_xlabel('True Values')
        ax1.set_ylabel('Predictions')
        ax1.set_title(f'{model_name} - Predictions vs True Values')
        ax1.grid(True, alpha=0.3)
        
        # Residuals plot
        ax2.scatter(analysis['predictions'], analysis['errors'], alpha=0.6)
        ax2.axhline(y=0, color='r', linestyle='--')
        ax2.set_xlabel('Predictions')
        ax2.set_ylabel('Residuals')
        ax2.set_title(f'{model_name} - Residuals Plot')
        ax2.grid(True, alpha=0.3)
        
        # Error distribution
        ax3.hist(analysis['errors'], bins=30, alpha=0.7, edgecolor='black')
        ax3.set_xlabel('Prediction Error')
        ax3.set_ylabel('Frequency')
        ax3.set_title(f'{model_name} - Error Distribution')
        ax3.grid(True, alpha=0.3)
        
        # Absolute error distribution
        ax4.hist(analysis['abs_errors'], bins=30, alpha=0.7, edgecolor='black')
        ax4.set_xlabel('Absolute Prediction Error')
        ax4.set_ylabel('Frequency')
        ax4.set_title(f'{model_name} - Absolute Error Distribution')
        ax4.grid(True, alpha=0.3)
        
        plt.tight_layout()
        
        save_path = self.output_dir / f"{model_name.lower().replace(' ', '_')}_prediction_analysis.png"
        fig.savefig(save_path, dpi=300, bbox_inches='tight')
        logger.info(f"Prediction analysis plot saved to {save_path}")
        
        plt.close(fig)
    
    def _analyze_worst_predictions(self, worst_indices: np.ndarray, 
                                 graph_data_list: List[Any], compound_names: List[str],
                                 predictions: np.ndarray, true_values: np.ndarray) -> None:
        """Analyze worst predictions and create visualizations."""
        logger.info("Analyzing worst predictions")
        
        for i, idx in enumerate(worst_indices):
            if idx < len(graph_data_list) and idx < len(compound_names):
                compound_name = compound_names[idx]
                graph_data = graph_data_list[idx]
                pred = predictions[idx]
                true = true_values[idx]
                
                # Visualize the molecular graph
                save_path = self.output_dir / f"worst_prediction_{i+1}_{compound_name}.png"
                self.visualizer.visualize_molecular_graph(
                    graph_data, compound_name, save_path=str(save_path)
                )
                
                # Extract graph features
                features = self.feature_extractor.extract_graph_features(graph_data)
                
                logger.info(f"Worst prediction {i+1}: {compound_name}")
                logger.info(f"  Predicted: {pred:.3f}, True: {true:.3f}, Error: {abs(pred-true):.3f}")
                logger.info(f"  Graph features: {features}")
    
    def create_model_report(self, model_results: Dict[str, Dict], 
                          output_path: Optional[str] = None) -> str:
        """
        Create comprehensive model analysis report.
        
        Args:
            model_results: Dictionary of model results
            output_path: Optional path to save the report
            
        Returns:
            Report content as string
        """
        report_lines = []
        report_lines.append("# GNN Model Analysis Report")
        report_lines.append(f"Generated on: {pd.Timestamp.now()}")
        report_lines.append("")
        
        # Model comparison table
        report_lines.append("## Model Performance Comparison")
        report_lines.append("")
        
        comparison_data = []
        for model_name, results in model_results.items():
            metrics = results.get('metrics', {})
            comparison_data.append({
                'Model': model_name,
                'R²': metrics.get('r2', 0),
                'RMSE': metrics.get('rmse', 0),
                'MAE': metrics.get('mae', 0)
            })
        
        df_comparison = pd.DataFrame(comparison_data)
        report_lines.append(df_comparison.to_string(index=False))
        report_lines.append("")
        
        # Best model analysis
        best_model = max(model_results.keys(), 
                        key=lambda x: model_results[x].get('metrics', {}).get('r2', 0))
        report_lines.append(f"## Best Model: {best_model}")
        report_lines.append("")
        
        best_metrics = model_results[best_model].get('metrics', {})
        for metric, value in best_metrics.items():
            report_lines.append(f"- {metric.upper()}: {value:.4f}")
        
        report_lines.append("")
        report_lines.append("## Analysis Summary")
        report_lines.append("")
        report_lines.append("This report provides a comprehensive analysis of Graph Neural Network")
        report_lines.append("models for TLR4 binding affinity prediction. The models were trained")
        report_lines.append("on molecular graph representations and evaluated using standard")
        report_lines.append("regression metrics.")
        
        report_content = "\n".join(report_lines)
        
        if output_path:
            with open(output_path, 'w') as f:
                f.write(report_content)
            logger.info(f"Model report saved to {output_path}")
        
        return report_content
```
