## Supplementary material for "Machine Learning-Driven Prediction of TLR4 Binding Affinity: A Comprehensive Molecular Feature Analysis for Drug Discovery": Codebase: interpretability.py

```
"""
Model Interpretability and Analysis Suite for TLR4 Binding Prediction

This module provides comprehensive interpretability tools for understanding
what molecular features drive strong TLR4 binding (low affinity values).
"""

import numpy as np
import pandas as pd
import matplotlib.pyplot as plt
import seaborn as sns
from typing import Dict, List, Tuple, Any, Optional, Union
import warnings
warnings.filterwarnings('ignore')

# SHAP for global and local feature importance
try:
    import shap
    SHAP_AVAILABLE = True
except ImportError:
    SHAP_AVAILABLE = False
    print("Warning: SHAP not available. Install with: pip install shap")

# LIME for local interpretability
try:
    import lime
    import lime.lime_tabular
    LIME_AVAILABLE = True
except ImportError:
    LIME_AVAILABLE = False
    print("Warning: LIME not available. Install with: pip install lime")

# RDKit for molecular analysis
try:
    from rdkit import Chem
    from rdkit.Chem import Descriptors, rdMolDescriptors, Draw
    from rdkit.Chem.Draw import rdMolDraw2D
    RDKIT_AVAILABLE = True
except ImportError:
    RDKIT_AVAILABLE = False
    print("Warning: RDKit not available. Install with: conda install -c conda-forge rdkit")

# PyTorch for attention visualization
try:
    import torch
    import torch.nn.functional as F
    TORCH_AVAILABLE = True
except ImportError:
    TORCH_AVAILABLE = False

from sklearn.ensemble import RandomForestRegressor
from sklearn.svm import SVR
from sklearn.linear_model import LinearRegression
from sklearn.preprocessing import StandardScaler
import joblib
import os
import json
from datetime import datetime


class ModelInterpretabilitySuite:
    """
    Comprehensive interpretability suite for TLR4 binding prediction models.
    
    Provides SHAP, LIME, molecular substructure analysis, and attention
    visualization to understand what drives strong TLR4 binding.
    """
    
    def __init__(self, models: Dict[str, Any], feature_names: List[str], 
                 X_train: pd.DataFrame, y_train: pd.Series,
                 X_test: pd.DataFrame, y_test: pd.Series,
                 output_dir: str = "results/interpretability"):
        """
        Initialize interpretability suite.
        
        Args:
            models: Dictionary of trained models {name: model}
            feature_names: List of feature names
            X_train, y_train: Training data
            X_test, y_test: Test data
            output_dir: Directory to save interpretability results
        """
        self.models = models
        self.feature_names = feature_names
        self.X_train = X_train
        self.y_train = y_train
        self.X_test = X_test
        self.y_test = y_test
        self.output_dir = output_dir
        
        # Create output directory
        os.makedirs(output_dir, exist_ok=True)
        
        # Initialize explainers
        self.shap_explainers = {}
        self.lime_explainers = {}
        
        # Results storage
        self.interpretability_results = {}
        
    def generate_shap_analysis(self, model_name: str, model: Any, 
                             sample_size: int = 100) -> Dict[str, Any]:
        """
        Generate SHAP analysis for a specific model.
        
        Args:
            model_name: Name of the model
            model: Trained model object
            sample_size: Number of samples to use for SHAP analysis
            
        Returns:
            Dictionary containing SHAP analysis results
        """
        if not SHAP_AVAILABLE:
            print(f"SHAP not available. Skipping SHAP analysis for {model_name}")
            return {}
            
        print(f"Generating SHAP analysis for {model_name}...")
        
        # Sample data for SHAP analysis
        sample_indices = np.random.choice(len(self.X_test), 
                                        min(sample_size, len(self.X_test)), 
                                        replace=False)
        X_sample = self.X_test.iloc[sample_indices]
        
        try:
            # Create SHAP explainer based on model type
            if hasattr(model, 'predict_proba'):
                # For models with probability prediction
                explainer = shap.Explainer(model, self.X_train)
            else:
                # For regression models
                if hasattr(model, 'feature_importances_'):
                    # Tree-based models
                    explainer = shap.TreeExplainer(model)
                else:
                    # Linear models
                    explainer = shap.LinearExplainer(model, self.X_train)
            
            # Calculate SHAP values
            shap_values = explainer.shap_values(X_sample)
            
            # Store explainer for later use
            self.shap_explainers[model_name] = explainer
            
            # Generate plots
            self._create_shap_plots(model_name, shap_values, X_sample)
            
            # Calculate feature importance
            feature_importance = self._calculate_shap_importance(shap_values)
            
            results = {
                'shap_values': shap_values,
                'feature_importance': feature_importance,
                'explainer': explainer,
                'sample_data': X_sample
            }
            
            self.interpretability_results[f"{model_name}_shap"] = results
            return results
            
        except Exception as e:
            print(f"Error generating SHAP analysis for {model_name}: {str(e)}")
            return {}
    
    def _create_shap_plots(self, model_name: str, shap_values: np.ndarray, 
                          X_sample: pd.DataFrame):
        """Create SHAP visualization plots."""
        try:
            # Summary plot
            plt.figure(figsize=(10, 8))
            shap.summary_plot(shap_values, X_sample, show=False, max_display=20)
            plt.title(f'SHAP Summary Plot - {model_name}')
            plt.tight_layout()
            plt.savefig(f'{self.output_dir}/shap_summary_{model_name}.png', 
                       dpi=300, bbox_inches='tight')
            plt.close()
            
            # Waterfall plot for first prediction
            if len(shap_values.shape) > 1:
                plt.figure(figsize=(10, 6))
                shap.waterfall_plot(shap_values[0], show=False)
                plt.title(f'SHAP Waterfall Plot - {model_name} (First Sample)')
                plt.tight_layout()
                plt.savefig(f'{self.output_dir}/shap_waterfall_{model_name}.png', 
                           dpi=300, bbox_inches='tight')
                plt.close()
            
            # Feature importance bar plot
            feature_importance = self._calculate_shap_importance(shap_values)
            self._plot_feature_importance(feature_importance, model_name, "SHAP")
            
        except Exception as e:
            print(f"Error creating SHAP plots for {model_name}: {str(e)}")
    
    def generate_lime_analysis(self, model_name: str, model: Any, 
                              num_samples: int = 5) -> Dict[str, Any]:
        """
        Generate LIME analysis for local interpretability.
        
        Args:
            model_name: Name of the model
            model: Trained model object
            num_samples: Number of samples to explain
            
        Returns:
            Dictionary containing LIME analysis results
        """
        if not LIME_AVAILABLE:
            print(f"LIME not available. Skipping LIME analysis for {model_name}")
            return {}
            
        print(f"Generating LIME analysis for {model_name}...")
        
        try:
            # Create LIME explainer
            explainer = lime.lime_tabular.LimeTabularExplainer(
                self.X_train.values,
                feature_names=self.feature_names,
                mode='regression',
                discretize_continuous=True
            )
            
            # Store explainer
            self.lime_explainers[model_name] = explainer
            
            # Generate explanations for sample predictions
            explanations = []
            sample_indices = np.random.choice(len(self.X_test), 
                                            min(num_samples, len(self.X_test)), 
                                            replace=False)
            
            for idx in sample_indices:
                X_instance = self.X_test.iloc[idx:idx+1]
                y_true = self.y_test.iloc[idx]
                y_pred = model.predict(X_instance)[0]
                
                # Generate LIME explanation
                explanation = explainer.explain_instance(
                    X_instance.values[0], 
                    model.predict, 
                    num_features=len(self.feature_names)
                )
                
                explanations.append({
                    'instance_idx': idx,
                    'true_value': y_true,
                    'predicted_value': y_pred,
                    'explanation': explanation,
                    'features': X_instance.iloc[0].to_dict()
                })
            
            # Create LIME plots
            self._create_lime_plots(model_name, explanations)
            
            results = {
                'explanations': explanations,
                'explainer': explainer
            }
            
            self.interpretability_results[f"{model_name}_lime"] = results
            return results
            
        except Exception as e:
            print(f"Error generating LIME analysis for {model_name}: {str(e)}")
            return {}
    
    def _create_lime_plots(self, model_name: str, explanations: List[Dict]):
        """Create LIME visualization plots."""
        try:
            for i, exp in enumerate(explanations):
                # Create explanation plot
                plt.figure(figsize=(10, 6))
                exp['explanation'].as_pyplot_figure()
                plt.title(f'LIME Explanation - {model_name} (Sample {i+1})\n'
                         f'True: {exp["true_value"]:.3f}, Predicted: {exp["predicted_value"]:.3f}')
                plt.tight_layout()
                plt.savefig(f'{self.output_dir}/lime_explanation_{model_name}_sample_{i+1}.png', 
                           dpi=300, bbox_inches='tight')
                plt.close()
                
        except Exception as e:
            print(f"Error creating LIME plots for {model_name}: {str(e)}")
    
    def analyze_molecular_substructures(self, compound_data: pd.DataFrame, 
                                      binding_affinities: pd.Series,
                                      threshold_percentile: float = 20) -> Dict[str, Any]:
        """
        Analyze molecular substructures that drive strong TLR4 binding.
        
        Args:
            compound_data: DataFrame with molecular features
            binding_affinities: Series of binding affinity values
            threshold_percentile: Percentile threshold for strong binders
            
        Returns:
            Dictionary containing substructure analysis results
        """
        if not RDKIT_AVAILABLE:
            print("RDKit not available. Skipping molecular substructure analysis.")
            return {}
            
        print("Analyzing molecular substructures for strong TLR4 binding...")
        
        # Identify strong binders (lowest affinity values)
        threshold_value = np.percentile(binding_affinities, threshold_percentile)
        strong_binders = binding_affinities <= threshold_value
        weak_binders = binding_affinities > threshold_value
        
        print(f"Strong binders threshold: {threshold_value:.3f}")
        print(f"Number of strong binders: {strong_binders.sum()}")
        print(f"Number of weak binders: {weak_binders.sum()}")
        
        # Analyze molecular descriptors for strong vs weak binders
        descriptor_analysis = self._analyze_molecular_descriptors(
            compound_data, strong_binders, weak_binders
        )
        
        # Analyze feature importance for binding strength
        feature_analysis = self._analyze_binding_features(
            compound_data, binding_affinities
        )
        
        results = {
            'threshold_value': threshold_value,
            'strong_binders_count': strong_binders.sum(),
            'weak_binders_count': weak_binders.sum(),
            'descriptor_analysis': descriptor_analysis,
            'feature_analysis': feature_analysis
        }
        
        # Create substructure analysis plots
        self._create_substructure_plots(results, compound_data, binding_affinities)
        
        self.interpretability_results['molecular_substructures'] = results
        return results
    
    def _analyze_molecular_descriptors(self, compound_data: pd.DataFrame,
                                     strong_binders: pd.Series,
                                     weak_binders: pd.Series) -> Dict[str, Any]:
        """Analyze molecular descriptors for strong vs weak binders."""
        
        # Get molecular descriptor columns (assuming they start with 'mol_')
        mol_columns = [col for col in compound_data.columns if col.startswith('mol_')]
        
        if not mol_columns:
            return {'error': 'No molecular descriptor columns found'}
        
        analysis = {}
        
        for col in mol_columns:
            strong_values = compound_data.loc[strong_binders, col].dropna()
            weak_values = compound_data.loc[weak_binders, col].dropna()
            
            if len(strong_values) > 0 and len(weak_values) > 0:
                # Statistical comparison
                from scipy import stats
                t_stat, p_value = stats.ttest_ind(strong_values, weak_values)
                
                analysis[col] = {
                    'strong_mean': strong_values.mean(),
                    'weak_mean': weak_values.mean(),
                    'strong_std': strong_values.std(),
                    'weak_std': weak_values.std(),
                    't_statistic': t_stat,
                    'p_value': p_value,
                    'effect_size': (strong_values.mean() - weak_values.mean()) / 
                                 np.sqrt((strong_values.var() + weak_values.var()) / 2)
                }
        
        return analysis
    
    def _analyze_binding_features(self, compound_data: pd.DataFrame,
                                binding_affinities: pd.Series) -> Dict[str, Any]:
        """Analyze which features are most important for binding strength."""
        
        # Calculate correlation between features and binding affinity
        correlations = compound_data.corrwith(binding_affinities).abs().sort_values(ascending=False)
        
        # Get top features
        top_features = correlations.head(20)
        
        return {
            'feature_correlations': correlations.to_dict(),
            'top_features': top_features.to_dict(),
            'correlation_analysis': {
                'positive_corr': correlations[correlations > 0.1].to_dict(),
                'negative_corr': correlations[correlations < -0.1].to_dict()
            }
        }
    
    def _create_substructure_plots(self, results: Dict, compound_data: pd.DataFrame,
                                 binding_affinities: pd.Series):
        """Create plots for molecular substructure analysis."""
        
        # Plot 1: Feature correlation with binding affinity
        correlations = results['feature_analysis']['feature_correlations']
        top_correlations = dict(list(correlations.items())[:15])
        
        plt.figure(figsize=(12, 8))
        features = list(top_correlations.keys())
        corr_values = list(top_correlations.values())
        
        plt.barh(features, corr_values)
        plt.xlabel('Absolute Correlation with Binding Affinity')
        plt.title('Top Features Correlated with TLR4 Binding Affinity')
        plt.tight_layout()
        plt.savefig(f'{self.output_dir}/feature_correlation_analysis.png', 
                   dpi=300, bbox_inches='tight')
        plt.close()
        
        # Plot 2: Molecular descriptor comparison
        descriptor_analysis = results['descriptor_analysis']
        if 'error' not in descriptor_analysis:
            self._plot_descriptor_comparison(descriptor_analysis)
    
    def _plot_descriptor_comparison(self, descriptor_analysis: Dict):
        """Plot comparison of molecular descriptors between strong and weak binders."""
        
        # Select top descriptors by effect size
        effect_sizes = {k: v['effect_size'] for k, v in descriptor_analysis.items() 
                       if 'effect_size' in v}
        top_descriptors = sorted(effect_sizes.items(), key=lambda x: abs(x[1]), reverse=True)[:10]
        
        if not top_descriptors:
            return
        
        fig, axes = plt.subplots(2, 5, figsize=(20, 10))
        axes = axes.flatten()
        
        for i, (desc, effect_size) in enumerate(top_descriptors):
            if i >= 10:
                break
                
            data = descriptor_analysis[desc]
            strong_mean = data['strong_mean']
            weak_mean = data['weak_mean']
            strong_std = data['strong_std']
            weak_std = data['weak_std']
            
            axes[i].bar(['Strong Binders', 'Weak Binders'], 
                       [strong_mean, weak_mean],
                       yerr=[strong_std, weak_std],
                       capsize=5, alpha=0.7)
            axes[i].set_title(f'{desc}\nEffect Size: {effect_size:.3f}')
            axes[i].set_ylabel('Value')
        
        plt.tight_layout()
        plt.savefig(f'{self.output_dir}/molecular_descriptor_comparison.png', 
                   dpi=300, bbox_inches='tight')
        plt.close()
    
    def visualize_attention_weights(self, model_name: str, model: Any, 
                                  sample_data: pd.DataFrame) -> Dict[str, Any]:
        """
        Visualize attention weights for transformer and GNN models.
        
        Args:
            model_name: Name of the model
            model: Trained model object
            sample_data: Sample data for visualization
            
        Returns:
            Dictionary containing attention visualization results
        """
        if not TORCH_AVAILABLE:
            print("PyTorch not available. Skipping attention visualization.")
            return {}
            
        print(f"Visualizing attention weights for {model_name}...")
        
        try:
            # Check if model has attention mechanisms
            if hasattr(model, 'attention_weights') or 'transformer' in model_name.lower():
                attention_results = self._extract_attention_weights(model, sample_data)
                
                # Create attention visualization plots
                self._create_attention_plots(model_name, attention_results)
                
                self.interpretability_results[f"{model_name}_attention"] = attention_results
                return attention_results
            else:
                print(f"Model {model_name} does not have attention mechanisms.")
                return {}
                
        except Exception as e:
            print(f"Error visualizing attention for {model_name}: {str(e)}")
            return {}
    
    def _extract_attention_weights(self, model: Any, sample_data: pd.DataFrame) -> Dict:
        """Extract attention weights from model."""
        # This is a placeholder - implementation depends on specific model architecture
        # In practice, you would extract attention weights from the model's forward pass
        
        attention_weights = {
            'layer_attention': [],
            'head_attention': [],
            'token_attention': []
        }
        
        # Placeholder implementation
        # Real implementation would depend on the specific model architecture
        
        return attention_weights
    
    def _create_attention_plots(self, model_name: str, attention_results: Dict):
        """Create attention visualization plots."""
        # Placeholder for attention plots
        # Implementation would depend on the specific attention mechanism
        pass
    
    def _calculate_shap_importance(self, shap_values: np.ndarray) -> Dict[str, float]:
        """Calculate feature importance from SHAP values."""
        if len(shap_values.shape) > 1:
            # Multi-dimensional SHAP values
            importance = np.mean(np.abs(shap_values), axis=0)
        else:
            # Single-dimensional SHAP values
            importance = np.abs(shap_values)
        
        return dict(zip(self.feature_names, importance))
    
    def _plot_feature_importance(self, importance_dict: Dict[str, float], 
                               model_name: str, method: str):
        """Plot feature importance."""
        # Sort features by importance
        sorted_features = sorted(importance_dict.items(), key=lambda x: x[1], reverse=True)
        top_features = sorted_features[:15]
        
        features, importance = zip(*top_features)
        
        plt.figure(figsize=(12, 8))
        plt.barh(features, importance)
        plt.xlabel(f'{method} Feature Importance')
        plt.title(f'Top Features - {model_name} ({method})')
        plt.tight_layout()
        plt.savefig(f'{self.output_dir}/feature_importance_{method.lower()}_{model_name}.png', 
                   dpi=300, bbox_inches='tight')
        plt.close()
    
    def generate_comprehensive_report(self) -> Dict[str, Any]:
        """
        Generate comprehensive interpretability report.
        
        Returns:
            Dictionary containing all interpretability results and analysis
        """
        print("Generating comprehensive interpretability report...")
        
        report = {
            'timestamp': datetime.now().isoformat(),
            'models_analyzed': list(self.models.keys()),
            'interpretability_results': self.interpretability_results,
            'summary': self._generate_summary()
        }
        
        # Save report
        report_path = f'{self.output_dir}/interpretability_report.json'
        with open(report_path, 'w') as f:
            json.dump(report, f, indent=2, default=str)
        
        # Generate markdown report
        self._generate_markdown_report(report)
        
        print(f"Interpretability report saved to {self.output_dir}/")
        return report
    
    def _generate_summary(self) -> Dict[str, Any]:
        """Generate summary of interpretability analysis."""
        summary = {
            'total_models': len(self.models),
            'shap_analyses': len([k for k in self.interpretability_results.keys() if 'shap' in k]),
            'lime_analyses': len([k for k in self.interpretability_results.keys() if 'lime' in k]),
            'attention_analyses': len([k for k in self.interpretability_results.keys() if 'attention' in k]),
            'substructure_analysis': 'molecular_substructures' in self.interpretability_results
        }
        
        return summary
    
    def _generate_markdown_report(self, report: Dict):
        """Generate markdown interpretability report."""
        md_content = f"""# Model Interpretability Report

Generated: {report['timestamp']}

## Summary
- Total models analyzed: {report['summary']['total_models']}
- SHAP analyses: {report['summary']['shap_analyses']}
- LIME analyses: {report['summary']['lime_analyses']}
- Attention analyses: {report['summary']['attention_analyses']}
- Substructure analysis: {report['summary']['substructure_analysis']}

## Key Findings

### Molecular Features Driving Strong TLR4 Binding
The analysis reveals the most important molecular features that predict strong TLR4 binding (low affinity values):

1. **Molecular Descriptors**: Key descriptors that distinguish strong from weak binders
2. **Feature Correlations**: Features most correlated with binding affinity
3. **Model Interpretability**: SHAP and LIME explanations for individual predictions

### Recommendations
Based on the interpretability analysis:
- Focus on molecular features with highest importance scores
- Consider structural modifications that enhance key descriptors
- Use attention visualizations to understand model decision-making

## Files Generated
- SHAP summary plots for each model
- LIME explanations for sample predictions
- Molecular substructure analysis plots
- Feature importance rankings
- Comprehensive JSON report

See individual plot files in the results directory for detailed visualizations.
"""
        
        with open(f'{self.output_dir}/interpretability_report.md', 'w') as f:
            f.write(md_content)
    
    def run_full_analysis(self, sample_size: int = 100, 
                         lime_samples: int = 5) -> Dict[str, Any]:
        """
        Run complete interpretability analysis for all models.
        
        Args:
            sample_size: Number of samples for SHAP analysis
            lime_samples: Number of samples for LIME analysis
            
        Returns:
            Complete interpretability analysis results
        """
        print("Running full interpretability analysis...")
        
        # Run SHAP analysis for all models
        for model_name, model in self.models.items():
            self.generate_shap_analysis(model_name, model, sample_size)
        
        # Run LIME analysis for all models
        for model_name, model in self.models.items():
            self.generate_lime_analysis(model_name, model, lime_samples)
        
        # Run molecular substructure analysis
        self.analyze_molecular_substructures(self.X_test, self.y_test)
        
        # Run attention visualization for applicable models
        for model_name, model in self.models.items():
            self.visualize_attention_weights(model_name, model, self.X_test.head(10))
        
        # Generate comprehensive report
        report = self.generate_comprehensive_report()
        
        print("Full interpretability analysis completed!")
        return report
```
