## Supplementary material for "Machine Learning-Driven Prediction of TLR4 Binding Affinity: A Comprehensive Molecular Feature Analysis for Drug Discovery": Codebase: molecular_substructure_analyzer.py

```
"""
Molecular Substructure Analysis for TLR4 Binding Prediction

This module provides specialized analysis of molecular substructures
that drive strong TLR4 binding interactions.
"""

import numpy as np
import pandas as pd
import matplotlib.pyplot as plt
import seaborn as sns
from typing import Dict, List, Tuple, Any, Optional
import warnings
warnings.filterwarnings('ignore')

try:
    from rdkit import Chem
    from rdkit.Chem import Descriptors, rdMolDescriptors, Draw, Crippen
    from rdkit.Chem.Draw import rdMolDraw2D
    from rdkit.Chem import rdFingerprintGenerator
    from rdkit.Chem import rdMolDescriptors
    RDKIT_AVAILABLE = True
except ImportError:
    RDKIT_AVAILABLE = False

from scipy import stats
from sklearn.cluster import KMeans
from sklearn.decomposition import PCA
from sklearn.manifold import TSNE
import os


class MolecularSubstructureAnalyzer:
    """
    Analyzes molecular substructures that drive strong TLR4 binding.
    
    Focuses on identifying structural features that predict low affinity
    values (strong binding) in TLR4 interactions.
    """
    
    def __init__(self, output_dir: str = "results/interpretability/substructures"):
        """Initialize the substructure analyzer."""
        self.output_dir = output_dir
        os.makedirs(output_dir, exist_ok=True)
        
    def analyze_binding_drivers(self, compound_data: pd.DataFrame, 
                              binding_affinities: pd.Series,
                              smiles_column: str = 'smiles',
                              threshold_percentile: float = 20) -> Dict[str, Any]:
        """
        Analyze molecular substructures that drive strong TLR4 binding.
        
        Args:
            compound_data: DataFrame with molecular data including SMILES
            binding_affinities: Series of binding affinity values
            smiles_column: Name of column containing SMILES strings
            threshold_percentile: Percentile threshold for strong binders
            
        Returns:
            Dictionary containing substructure analysis results
        """
        if not RDKIT_AVAILABLE:
            print("RDKit not available. Cannot perform substructure analysis.")
            return {}
            
        print("Analyzing molecular substructures for TLR4 binding drivers...")
        
        # Identify strong vs weak binders
        threshold_value = np.percentile(binding_affinities, threshold_percentile)
        strong_binders = binding_affinities <= threshold_value
        weak_binders = binding_affinities > threshold_value
        
        print(f"Strong binders threshold: {threshold_value:.3f} kcal/mol")
        print(f"Number of strong binders: {strong_binders.sum()}")
        print(f"Number of weak binders: {weak_binders.sum()}")
        
        # Extract molecular features
        molecular_features = self._extract_molecular_features(
            compound_data, smiles_column
        )
        
        # Analyze structural differences
        structural_analysis = self._analyze_structural_differences(
            molecular_features, strong_binders, weak_binders
        )
        
        # Analyze pharmacophore features
        pharmacophore_analysis = self._analyze_pharmacophore_features(
            molecular_features, strong_binders, weak_binders
        )
        
        # Analyze molecular fingerprints
        fingerprint_analysis = self._analyze_molecular_fingerprints(
            molecular_features, strong_binders, weak_binders
        )
        
        # Generate visualizations
        self._create_substructure_visualizations(
            molecular_features, strong_binders, weak_binders, binding_affinities
        )
        
        results = {
            'threshold_value': threshold_value,
            'strong_binders_count': strong_binders.sum(),
            'weak_binders_count': weak_binders.sum(),
            'structural_analysis': structural_analysis,
            'pharmacophore_analysis': pharmacophore_analysis,
            'fingerprint_analysis': fingerprint_analysis,
            'molecular_features': molecular_features
        }
        
        return results
    
    def _extract_molecular_features(self, compound_data: pd.DataFrame, 
                                  smiles_column: str) -> Dict[str, Any]:
        """Extract comprehensive molecular features from SMILES."""
        
        features = {
            'molecules': [],
            'molecular_weights': [],
            'logp_values': [],
            'tpsa_values': [],
            'hbd_counts': [],
            'hba_counts': [],
            'rotatable_bonds': [],
            'aromatic_rings': [],
            'heavy_atoms': [],
            'formal_charges': [],
            'molecular_volumes': [],
            'surface_areas': [],
            'fingerprints': []
        }
        
        valid_indices = []
        
        for idx, smiles in enumerate(compound_data[smiles_column]):
            try:
                mol = Chem.MolFromSmiles(smiles)
                if mol is not None:
                    features['molecules'].append(mol)
                    features['molecular_weights'].append(Descriptors.MolWt(mol))
                    features['logp_values'].append(Crippen.MolLogP(mol))
                    features['tpsa_values'].append(Descriptors.TPSA(mol))
                    features['hbd_counts'].append(Descriptors.NumHDonors(mol))
                    features['hba_counts'].append(Descriptors.NumHAcceptors(mol))
                    features['rotatable_bonds'].append(Descriptors.NumRotatableBonds(mol))
                    features['aromatic_rings'].append(Descriptors.NumAromaticRings(mol))
                    features['heavy_atoms'].append(mol.GetNumHeavyAtoms())
                    features['formal_charges'].append(Chem.rdmolops.GetFormalCharge(mol))
                    
                    # 3D descriptors (approximated)
                    features['molecular_volumes'].append(rdMolDescriptors.CalcCrippenDescriptors(mol)[0])
                    features['surface_areas'].append(rdMolDescriptors.CalcCrippenDescriptors(mol)[1])
                    
                    # Molecular fingerprints
                    fp = rdFingerprintGenerator.GetMorganGenerator().GetFingerprint(mol)
                    features['fingerprints'].append(fp)
                    
                    valid_indices.append(idx)
                    
            except Exception as e:
                print(f"Error processing SMILES {smiles}: {str(e)}")
                continue
        
        # Convert to numpy arrays
        for key in features:
            if key != 'molecules' and key != 'fingerprints':
                features[key] = np.array(features[key])
        
        features['valid_indices'] = valid_indices
        return features
    
    def _analyze_structural_differences(self, molecular_features: Dict,
                                      strong_binders: pd.Series,
                                      weak_binders: pd.Series) -> Dict[str, Any]:
        """Analyze structural differences between strong and weak binders."""
        
        analysis = {}
        
        # Get valid indices for strong and weak binders
        valid_indices = molecular_features['valid_indices']
        strong_valid = strong_binders.iloc[valid_indices]
        weak_valid = weak_binders.iloc[valid_indices]
        
        # Analyze each molecular descriptor
        descriptor_names = ['molecular_weights', 'logp_values', 'tpsa_values', 
                          'hbd_counts', 'hba_counts', 'rotatable_bonds', 
                          'aromatic_rings', 'heavy_atoms', 'formal_charges']
        
        for desc_name in descriptor_names:
            if desc_name in molecular_features:
                strong_values = molecular_features[desc_name][strong_valid]
                weak_values = molecular_features[desc_name][weak_valid]
                
                if len(strong_values) > 0 and len(weak_values) > 0:
                    # Statistical test
                    t_stat, p_value = stats.ttest_ind(strong_values, weak_values)
                    
                    # Effect size (Cohen's d)
                    pooled_std = np.sqrt((strong_values.var() + weak_values.var()) / 2)
                    effect_size = (strong_values.mean() - weak_values.mean()) / pooled_std
                    
                    analysis[desc_name] = {
                        'strong_mean': strong_values.mean(),
                        'weak_mean': weak_values.mean(),
                        'strong_std': strong_values.std(),
                        'weak_std': weak_values.std(),
                        't_statistic': t_stat,
                        'p_value': p_value,
                        'effect_size': effect_size,
                        'significant': p_value < 0.05
                    }
        
        return analysis
    
    def _analyze_pharmacophore_features(self, molecular_features: Dict,
                                      strong_binders: pd.Series,
                                      weak_binders: pd.Series) -> Dict[str, Any]:
        """Analyze pharmacophore features for binding prediction."""
        
        # Define pharmacophore features
        pharmacophore_features = {
            'hydrogen_bond_donors': molecular_features['hbd_counts'],
            'hydrogen_bond_acceptors': molecular_features['hba_counts'],
            'aromatic_rings': molecular_features['aromatic_rings'],
            'rotatable_bonds': molecular_features['rotatable_bonds'],
            'polar_surface_area': molecular_features['tpsa_values'],
            'lipophilicity': molecular_features['logp_values']
        }
        
        analysis = {}
        valid_indices = molecular_features['valid_indices']
        strong_valid = strong_binders.iloc[valid_indices]
        weak_valid = weak_binders.iloc[valid_indices]
        
        for feature_name, values in pharmacophore_features.items():
            strong_values = values[strong_valid]
            weak_values = values[weak_valid]
            
            if len(strong_values) > 0 and len(weak_values) > 0:
                t_stat, p_value = stats.ttest_ind(strong_values, weak_values)
                effect_size = (strong_values.mean() - weak_values.mean()) / \
                            np.sqrt((strong_values.var() + weak_values.var()) / 2)
                
                analysis[feature_name] = {
                    'strong_mean': strong_values.mean(),
                    'weak_mean': weak_values.mean(),
                    'effect_size': effect_size,
                    'p_value': p_value,
                    'significant': p_value < 0.05
                }
        
        return analysis
    
    def _analyze_molecular_fingerprints(self, molecular_features: Dict,
                                      strong_binders: pd.Series,
                                      weak_binders: pd.Series) -> Dict[str, Any]:
        """Analyze molecular fingerprints for substructure patterns."""
        
        # This is a simplified analysis - in practice, you'd use more sophisticated
        # fingerprint analysis methods
        
        valid_indices = molecular_features['valid_indices']
        strong_valid = strong_binders.iloc[valid_indices]
        weak_valid = weak_binders.iloc[valid_indices]
        
        # Get fingerprints for strong and weak binders
        strong_fps = [molecular_features['fingerprints'][i] for i in range(len(strong_valid)) if strong_valid.iloc[i]]
        weak_fps = [molecular_features['fingerprints'][i] for i in range(len(weak_valid)) if weak_valid.iloc[i]]
        
        # Calculate fingerprint similarity within groups
        strong_similarity = self._calculate_fingerprint_similarity(strong_fps)
        weak_similarity = self._calculate_fingerprint_similarity(weak_fps)
        
        return {
            'strong_binder_similarity': strong_similarity,
            'weak_binder_similarity': weak_similarity,
            'structural_diversity_strong': 1 - strong_similarity,
            'structural_diversity_weak': 1 - weak_similarity
        }
    
    def _calculate_fingerprint_similarity(self, fingerprints: List) -> float:
        """Calculate average fingerprint similarity within a group."""
        if len(fingerprints) < 2:
            return 0.0
        
        similarities = []
        for i in range(len(fingerprints)):
            for j in range(i + 1, len(fingerprints)):
                try:
                    from rdkit import DataStructs
                    similarity = DataStructs.TanimotoSimilarity(fingerprints[i], fingerprints[j])
                    similarities.append(similarity)
                except:
                    continue
        
        return np.mean(similarities) if similarities else 0.0
    
    def _create_substructure_visualizations(self, molecular_features: Dict,
                                          strong_binders: pd.Series,
                                          weak_binders: pd.Series,
                                          binding_affinities: pd.Series):
        """Create visualizations for substructure analysis."""
        
        # Plot 1: Molecular descriptor comparison
        self._plot_descriptor_comparison(molecular_features, strong_binders, weak_binders)
        
        # Plot 2: Binding affinity distribution
        self._plot_binding_affinity_distribution(binding_affinities)
        
        # Plot 3: Pharmacophore feature analysis
        self._plot_pharmacophore_analysis(molecular_features, strong_binders, weak_binders)
        
        # Plot 4: Molecular space visualization
        self._plot_molecular_space(molecular_features, binding_affinities)
    
    def _plot_descriptor_comparison(self, molecular_features: Dict,
                                  strong_binders: pd.Series, weak_binders: pd.Series):
        """Plot comparison of molecular descriptors."""
        
        valid_indices = molecular_features['valid_indices']
        strong_valid = strong_binders.iloc[valid_indices]
        weak_valid = weak_binders.iloc[valid_indices]
        
        # Select top descriptors by effect size
        descriptor_names = ['molecular_weights', 'logp_values', 'tpsa_values', 
                          'hbd_counts', 'hba_counts', 'rotatable_bonds']
        
        fig, axes = plt.subplots(2, 3, figsize=(18, 12))
        axes = axes.flatten()
        
        for i, desc_name in enumerate(descriptor_names):
            if desc_name in molecular_features:
                strong_values = molecular_features[desc_name][strong_valid]
                weak_values = molecular_features[desc_name][weak_valid]
                
                # Create box plot
                data_to_plot = [strong_values, weak_values]
                axes[i].boxplot(data_to_plot, labels=['Strong Binders', 'Weak Binders'])
                axes[i].set_title(f'{desc_name.replace("_", " ").title()}')
                axes[i].set_ylabel('Value')
                
                # Add statistical annotation
                t_stat, p_value = stats.ttest_ind(strong_values, weak_values)
                axes[i].text(0.5, 0.95, f'p = {p_value:.3f}', 
                           transform=axes[i].transAxes, ha='center', va='top')
        
        plt.tight_layout()
        plt.savefig(f'{self.output_dir}/molecular_descriptor_comparison.png', 
                   dpi=300, bbox_inches='tight')
        plt.close()
    
    def _plot_binding_affinity_distribution(self, binding_affinities: pd.Series):
        """Plot distribution of binding affinities."""
        
        plt.figure(figsize=(10, 6))
        plt.hist(binding_affinities, bins=30, alpha=0.7, edgecolor='black')
        plt.axvline(binding_affinities.quantile(0.2), color='red', linestyle='--', 
                   label='Strong Binders Threshold (20th percentile)')
        plt.xlabel('Binding Affinity (kcal/mol)')
        plt.ylabel('Frequency')
        plt.title('Distribution of TLR4 Binding Affinities')
        plt.legend()
        plt.grid(True, alpha=0.3)
        plt.tight_layout()
        plt.savefig(f'{self.output_dir}/binding_affinity_distribution.png', 
                   dpi=300, bbox_inches='tight')
        plt.close()
    
    def _plot_pharmacophore_analysis(self, molecular_features: Dict,
                                   strong_binders: pd.Series, weak_binders: pd.Series):
        """Plot pharmacophore feature analysis."""
        
        valid_indices = molecular_features['valid_indices']
        strong_valid = strong_binders.iloc[valid_indices]
        weak_valid = weak_binders.iloc[valid_indices]
        
        # Pharmacophore features
        features = {
            'HBD': molecular_features['hbd_counts'],
            'HBA': molecular_features['hba_counts'],
            'Aromatic Rings': molecular_features['aromatic_rings'],
            'Rotatable Bonds': molecular_features['rotatable_bonds'],
            'TPSA': molecular_features['tpsa_values'],
            'LogP': molecular_features['logp_values']
        }
        
        fig, axes = plt.subplots(2, 3, figsize=(18, 12))
        axes = axes.flatten()
        
        for i, (feature_name, values) in enumerate(features.items()):
            strong_values = values[strong_valid]
            weak_values = values[weak_valid]
            
            # Create violin plot
            data_to_plot = [strong_values, weak_values]
            parts = axes[i].violinplot(data_to_plot, positions=[1, 2], showmeans=True)
            axes[i].set_xticks([1, 2])
            axes[i].set_xticklabels(['Strong Binders', 'Weak Binders'])
            axes[i].set_title(f'{feature_name}')
            axes[i].set_ylabel('Value')
            
            # Add statistical test
            t_stat, p_value = stats.ttest_ind(strong_values, weak_values)
            axes[i].text(0.5, 0.95, f'p = {p_value:.3f}', 
                       transform=axes[i].transAxes, ha='center', va='top')
        
        plt.tight_layout()
        plt.savefig(f'{self.output_dir}/pharmacophore_analysis.png', 
                   dpi=300, bbox_inches='tight')
        plt.close()
    
    def _plot_molecular_space(self, molecular_features: Dict, binding_affinities: pd.Series):
        """Plot molecular space visualization."""
        
        # Prepare data for dimensionality reduction
        feature_matrix = np.column_stack([
            molecular_features['molecular_weights'],
            molecular_features['logp_values'],
            molecular_features['tpsa_values'],
            molecular_features['hbd_counts'],
            molecular_features['hba_counts'],
            molecular_features['rotatable_bonds']
        ])
        
        # Apply PCA
        pca = PCA(n_components=2)
        pca_result = pca.fit_transform(feature_matrix)
        
        # Create scatter plot
        plt.figure(figsize=(12, 8))
        scatter = plt.scatter(pca_result[:, 0], pca_result[:, 1], 
                            c=binding_affinities.iloc[molecular_features['valid_indices']], 
                            cmap='viridis', alpha=0.7)
        plt.colorbar(scatter, label='Binding Affinity (kcal/mol)')
        plt.xlabel(f'PC1 ({pca.explained_variance_ratio_[0]:.2%} variance)')
        plt.ylabel(f'PC2 ({pca.explained_variance_ratio_[1]:.2%} variance)')
        plt.title('Molecular Space Visualization (PCA)')
        plt.grid(True, alpha=0.3)
        plt.tight_layout()
        plt.savefig(f'{self.output_dir}/molecular_space_pca.png', 
                   dpi=300, bbox_inches='tight')
        plt.close()
    
    def generate_substructure_report(self, results: Dict) -> str:
        """Generate a comprehensive substructure analysis report."""
        
        report = f"""
# Molecular Substructure Analysis Report

## Summary
- Strong binders threshold: {results['threshold_value']:.3f} kcal/mol
- Number of strong binders: {results['strong_binders_count']}
- Number of weak binders: {results['weak_binders_count']}

## Key Findings

### Structural Differences
The analysis reveals significant structural differences between strong and weak TLR4 binders:

"""
        
        # Add structural analysis findings
        structural_analysis = results['structural_analysis']
        for desc_name, analysis in structural_analysis.items():
            if analysis['significant']:
                effect_direction = "higher" if analysis['effect_size'] > 0 else "lower"
                report += f"- **{desc_name.replace('_', ' ').title()}**: Strong binders have {effect_direction} values (p={analysis['p_value']:.3f}, effect size={analysis['effect_size']:.3f})\n"
        
        report += f"""

### Pharmacophore Features
Key pharmacophore features that distinguish strong from weak binders:

"""
        
        # Add pharmacophore analysis findings
        pharmacophore_analysis = results['pharmacophore_analysis']
        for feature_name, analysis in pharmacophore_analysis.items():
            if analysis['significant']:
                effect_direction = "higher" if analysis['effect_size'] > 0 else "lower"
                report += f"- **{feature_name}**: Strong binders have {effect_direction} values (p={analysis['p_value']:.3f})\n"
        
        report += f"""

### Recommendations
Based on the substructure analysis:
1. Focus on compounds with optimal values for significant molecular descriptors
2. Consider pharmacophore features that distinguish strong binders
3. Use molecular space visualization to identify promising chemical regions

## Files Generated
- Molecular descriptor comparison plots
- Binding affinity distribution
- Pharmacophore feature analysis
- Molecular space visualization (PCA)

See individual plot files in the results directory for detailed visualizations.
"""
        
        # Save report
        with open(f'{self.output_dir}/substructure_analysis_report.md', 'w') as f:
            f.write(report)
        
        return report
```
