## Supplementary material for "Machine Learning-Driven Prediction of TLR4 Binding Affinity: A Comprehensive Molecular Feature Analysis for Drug Discovery": Codebase: pipeline_orchestrator.py

```
"""
Research Pipeline Orchestrator for TLR4 Binding Prediction

This module provides comprehensive experiment tracking, automated hyperparameter
optimization, and reproducible research pipeline management using MLflow.
"""

import os
import json
import yaml
import logging
import warnings
from datetime import datetime
from pathlib import Path
from typing import Dict, List, Any, Optional, Tuple, Union
from dataclasses import dataclass, asdict
import traceback

import numpy as np
import pandas as pd

# Import error handling utilities
from ..utils.error_handling import (
    RobustnessManager, CheckpointManager, robust_execution,
    safe_execution, graceful_degradation, PipelineError,
    ModelTrainingError, CircuitBreaker
)
from ..utils.data_quality import DataQualityValidator, DataAnomalyDetector
from contextlib import nullcontext
try:
    import mlflow  # type: ignore
    import mlflow.sklearn  # type: ignore
    # Optional extra integrations; ignore if unavailable
    try:
        import mlflow.pytorch  # type: ignore
        import mlflow.xgboost  # type: ignore
        import mlflow.lightgbm  # type: ignore
    except Exception:
        pass
    MLFLOW_AVAILABLE = True
except Exception:
    MLFLOW_AVAILABLE = False
from sklearn.model_selection import (
    cross_val_score, 
    StratifiedKFold, 
    KFold,
    ParameterGrid,
    ParameterSampler
)
from sklearn.metrics import mean_squared_error, r2_score, mean_absolute_error
from scipy.stats import spearmanr
try:
    import optuna  # type: ignore
    OPTUNA_AVAILABLE = True
except Exception:
    OPTUNA_AVAILABLE = False

# Suppress warnings for cleaner output
warnings.filterwarnings('ignore')
logging.basicConfig(level=logging.INFO)
logger = logging.getLogger(__name__)


@dataclass
class ExperimentConfig:
    """Configuration for research experiments"""
    experiment_name: str
    description: str
    tags: Dict[str, str]
    data_path: str
    output_path: str
    random_state: int = 42
    n_trials: int = 100
    cv_folds: int = 5
    test_size: float = 0.2
    validation_size: float = 0.2
    enable_mlflow: bool = True
    mlflow_tracking_uri: str = "file:./mlruns"
    enable_optuna: bool = True
    optuna_storage: str = "sqlite:///optuna_studies.db"
    enable_nested_cv: bool = True
    nested_cv_folds: int = 3
    enable_hyperparameter_optimization: bool = True
    enable_feature_ablation: bool = True
    enable_uncertainty_quantification: bool = True
    enable_interpretability: bool = True
    enable_ensemble: bool = True
    models_to_test: List[str] = None
    
    def __post_init__(self):
        if self.models_to_test is None:
            self.models_to_test = [
                'random_forest', 'xgboost', 'lightgbm', 'svr', 
                'neural_network', 'graph_neural_network', 'transformer'
            ]


@dataclass
class ModelConfig:
    """Configuration for individual models"""
    name: str
    model_class: str
    hyperparameter_space: Dict[str, Any]
    cv_strategy: str = 'stratified'
    enable_early_stopping: bool = True
    max_iterations: int = 1000
    patience: int = 10


class ResearchPipelineOrchestrator:
    """
    Comprehensive research pipeline orchestrator for TLR4 binding prediction.
    
    Features:
    - MLflow experiment tracking
    - Automated hyperparameter optimization with Optuna
    - Nested cross-validation for unbiased performance estimation
    - Reproducible experiment configuration management
    - Automated research report generation
    """
    
    def __init__(self, config: ExperimentConfig, robustness_config: Optional[Dict[str, Any]] = None):
        self.config = config
        self.experiment_id = None
        self.run_id = None
        self.results = {}
        self.best_models = {}
        self.feature_importance = {}
        self.uncertainty_estimates = {}
        
        # Initialize robustness features
        self.robustness_manager = RobustnessManager(robustness_config)
        self.checkpoint_manager = CheckpointManager()
        self.data_validator = DataQualityValidator()
        self.anomaly_detector = DataAnomalyDetector()
        
        # Circuit breakers for different components
        self.circuit_breakers = {
            'model_training': CircuitBreaker(failure_threshold=3, timeout=300),
            'hyperparameter_optimization': CircuitBreaker(failure_threshold=2, timeout=600),
            'data_processing': CircuitBreaker(failure_threshold=5, timeout=180)
        }
        
        # Setup MLflow
        if self.config.enable_mlflow:
            self._setup_mlflow()
        
        # Setup Optuna
        if self.config.enable_optuna:
            self._setup_optuna()
    
    def _setup_mlflow(self):
        """Setup MLflow tracking"""
        try:
            mlflow.set_tracking_uri(self.config.mlflow_tracking_uri)
            self.experiment = mlflow.set_experiment(self.config.experiment_name)
            self.experiment_id = self.experiment.experiment_id
            logger.info(f"MLflow experiment setup: {self.config.experiment_name}")
        except Exception as e:
            logger.error(f"Failed to setup MLflow: {e}")
            self.config.enable_mlflow = False
    
    def _setup_optuna(self):
        """Setup Optuna for hyperparameter optimization"""
        try:
            self.study = optuna.create_study(
                direction='maximize',  # Maximize R² score
                storage=self.config.optuna_storage,
                study_name=f"{self.config.experiment_name}_optimization"
            )
            logger.info("Optuna study setup complete")
        except Exception as e:
            logger.error(f"Failed to setup Optuna: {e}")
            self.config.enable_optuna = False
    
    @robust_execution(max_retries=2, delay=2.0)
    def run_complete_pipeline(self, X: pd.DataFrame, y: pd.Series, 
                             resume_from_checkpoint: bool = True) -> Dict[str, Any]:
        """
        Run the complete research pipeline with robust error handling and checkpointing.
        
        Args:
            X: Feature matrix
            y: Target values (binding affinities)
            resume_from_checkpoint: Whether to resume from existing checkpoint
            
        Returns:
            Dictionary containing all results and metrics
        """
        logger.info("Starting complete research pipeline with robust error handling")
        
        # Generate checkpoint ID
        checkpoint_id = f"pipeline_run_{self.config.experiment_name}_{int(datetime.now().timestamp())}"
        
        # Try to resume from checkpoint
        if resume_from_checkpoint:
            checkpoint_data = self.checkpoint_manager.load_checkpoint(checkpoint_id)
            if checkpoint_data:
                logger.info("Resuming pipeline from checkpoint")
                self.results = checkpoint_data.get('results', {})
                self.best_models = checkpoint_data.get('best_models', {})
                self.feature_importance = checkpoint_data.get('feature_importance', {})
                self.uncertainty_estimates = checkpoint_data.get('uncertainty_estimates', {})
        
        with safe_execution("MLflow run setup", 
                          error_handler=lambda e: self.robustness_manager.log_error(e),
                          default_return=None) if self.config.enable_mlflow else nullcontext():
            
            if self.config.enable_mlflow:
                with mlflow.start_run(experiment_id=self.experiment_id) as run:
                    self.run_id = run.info.run_id
                    self._log_experiment_config()
                    return self._execute_pipeline_steps(X, y, checkpoint_id)
            else:
                return self._execute_pipeline_steps(X, y, checkpoint_id)
    
    def _execute_pipeline_steps(self, X: pd.DataFrame, y: pd.Series, checkpoint_id: str) -> Dict[str, Any]:
        """Execute pipeline steps with error handling and checkpointing."""
        try:
            # 0. Data quality validation
            logger.info("Performing data quality validation")
            with safe_execution("Data quality validation"):
                validation_results = self.data_validator.validate_dataset(X, "feature_matrix")
                if not validation_results['validation_passed']:
                    logger.warning("Data quality validation failed, but continuing with pipeline")
                
                # Log validation results
                self.results['data_quality'] = validation_results
            
            # 1. Data splitting
            logger.info("Performing data splitting")
            with safe_execution("Data splitting"):
                splits = self._split_data(X, y)
                self.results['data_splits'] = {
                    'train_size': len(splits['X_train']),
                    'val_size': len(splits['X_val']),
                    'test_size': len(splits['X_test'])
                }
                
                # Save checkpoint after data splitting
                self._save_pipeline_checkpoint(checkpoint_id, "data_split_complete")
                
                # 2. Hyperparameter optimization
                if self.config.enable_hyperparameter_optimization:
                    logger.info("Running hyperparameter optimization")
                    with safe_execution("Hyperparameter optimization"):
                        self._optimize_hyperparameters(splits)
                        self._save_pipeline_checkpoint(checkpoint_id, "hyperparameter_optimization_complete")
                
                # 3. Model training and evaluation
                logger.info("Training and evaluating models")
                with safe_execution("Model training and evaluation"):
                    self._train_and_evaluate_models(splits)
                    self._save_pipeline_checkpoint(checkpoint_id, "model_training_complete")
                
                # 4. Nested cross-validation
                if self.config.enable_nested_cv:
                    logger.info("Running nested cross-validation")
                    with safe_execution("Nested cross-validation"):
                        self._run_nested_cv(X, y)
                        self._save_pipeline_checkpoint(checkpoint_id, "nested_cv_complete")
                
                # 5. Feature ablation studies
                if self.config.enable_feature_ablation:
                    logger.info("Running feature ablation studies")
                    with safe_execution("Feature ablation studies"):
                        self._run_feature_ablation(splits)
                        self._save_pipeline_checkpoint(checkpoint_id, "feature_ablation_complete")
                
                # 6. Uncertainty quantification
                if self.config.enable_uncertainty_quantification:
                    logger.info("Running uncertainty quantification")
                    with safe_execution("Uncertainty quantification"):
                        self._run_uncertainty_quantification(splits)
                        self._save_pipeline_checkpoint(checkpoint_id, "uncertainty_quantification_complete")
                
                # 7. Model interpretability
                if self.config.enable_interpretability:
                    logger.info("Running interpretability analysis")
                    with safe_execution("Interpretability analysis"):
                        self._run_interpretability_analysis(splits)
                        self._save_pipeline_checkpoint(checkpoint_id, "interpretability_analysis_complete")
                
                # 8. Ensemble methods
                if self.config.enable_ensemble:
                    logger.info("Running ensemble methods")
                    with safe_execution("Ensemble methods"):
                        self._run_ensemble_methods(splits)
                        self._save_pipeline_checkpoint(checkpoint_id, "ensemble_methods_complete")
                
                # Final checkpoint
                self._save_pipeline_checkpoint(checkpoint_id, "pipeline_complete")
                
                logger.info("Complete research pipeline executed successfully")
                return self.results
                
        except Exception as e:
            error_msg = f"Pipeline execution failed: {str(e)}"
            logger.error(error_msg)
            self.robustness_manager.log_error(e, {'checkpoint_id': checkpoint_id})
            raise PipelineError(error_msg, context={'checkpoint_id': checkpoint_id})
    
    def _save_pipeline_checkpoint(self, checkpoint_id: str, stage: str):
        """Save pipeline checkpoint at current stage."""
        try:
            checkpoint_data = {
                'stage': stage,
                'timestamp': datetime.now().isoformat(),
                'results': self.results,
                'best_models': self.best_models,
                'feature_importance': self.feature_importance,
                'uncertainty_estimates': self.uncertainty_estimates,
                'config': asdict(self.config)
            }
            
            if self.checkpoint_manager.save_checkpoint(checkpoint_id, checkpoint_data):
                logger.info(f"Pipeline checkpoint saved at stage: {stage}")
            
        except Exception as e:
            logger.warning(f"Failed to save pipeline checkpoint: {e}")
    
    def _log_experiment_config(self):
        """Log experiment configuration to MLflow"""
        if not self.config.enable_mlflow:
            return
        
        # Log parameters
        config_dict = asdict(self.config)
        for key, value in config_dict.items():
            if isinstance(value, (str, int, float, bool)):
                mlflow.log_param(key, value)
            else:
                mlflow.log_param(key, str(value))
        
        # Log tags
        mlflow.set_tags(self.config.tags)
        
        # Log description
        mlflow.log_text(self.config.description, "experiment_description.txt")
    
    def _split_data(self, X: pd.DataFrame, y: pd.Series) -> Dict[str, Any]:
        """Split data into train/validation/test sets"""
        from sklearn.model_selection import train_test_split
        
        # First split: train+val vs test
        X_temp, X_test, y_temp, y_test = train_test_split(
            X, y, test_size=self.config.test_size, 
            random_state=self.config.random_state
        )
        
        # Second split: train vs val
        X_train, X_val, y_train, y_val = train_test_split(
            X_temp, y_temp, test_size=self.config.validation_size,
            random_state=self.config.random_state
        )
        
        splits = {
            'X_train': X_train, 'X_val': X_val, 'X_test': X_test,
            'y_train': y_train, 'y_val': y_val, 'y_test': y_test
        }
        
        # Log data split info
        if self.config.enable_mlflow:
            mlflow.log_metric("train_samples", len(X_train))
            mlflow.log_metric("val_samples", len(X_val))
            mlflow.log_metric("test_samples", len(X_test))
            mlflow.log_metric("total_features", X.shape[1])
        
        return splits
    
    def _optimize_hyperparameters(self, splits: Dict[str, Any]):
        """Run hyperparameter optimization using Optuna"""
        if not self.config.enable_optuna:
            return
        
        def objective(trial):
            # Sample hyperparameters for different models
            model_name = trial.suggest_categorical('model', self.config.models_to_test)
            
            # Get model-specific hyperparameters
            params = self._sample_hyperparameters(trial, model_name)
            
            # Train and evaluate model
            try:
                model = self._create_model(model_name, params)
                model.fit(splits['X_train'], splits['y_train'])
                
                # Evaluate on validation set
                y_pred = model.predict(splits['X_val'])
                r2 = r2_score(splits['y_val'], y_pred)
                
                return r2
            except Exception as e:
                logger.warning(f"Trial failed: {e}")
                return -np.inf
        
        # Run optimization
        self.study.optimize(objective, n_trials=self.config.n_trials)
        
        # Log best parameters
        if self.config.enable_mlflow:
            mlflow.log_params(self.study.best_params)
            mlflow.log_metric("best_cv_score", self.study.best_value)
    
    def _sample_hyperparameters(self, trial, model_name: str) -> Dict[str, Any]:
        """Sample hyperparameters for a specific model"""
        if model_name == 'random_forest':
            return {
                'n_estimators': trial.suggest_int('n_estimators', 50, 500),
                'max_depth': trial.suggest_int('max_depth', 3, 20),
                'min_samples_split': trial.suggest_int('min_samples_split', 2, 20),
                'min_samples_leaf': trial.suggest_int('min_samples_leaf', 1, 10),
                'max_features': trial.suggest_categorical('max_features', ['sqrt', 'log2', None])
            }
        elif model_name == 'xgboost':
            return {
                'n_estimators': trial.suggest_int('n_estimators', 50, 500),
                'max_depth': trial.suggest_int('max_depth', 3, 10),
                'learning_rate': trial.suggest_float('learning_rate', 0.01, 0.3),
                'subsample': trial.suggest_float('subsample', 0.6, 1.0),
                'colsample_bytree': trial.suggest_float('colsample_bytree', 0.6, 1.0)
            }
        elif model_name == 'neural_network':
            return {
                'hidden_layer_sizes': trial.suggest_categorical('hidden_layer_sizes', 
                    [(50,), (100,), (50, 50), (100, 50), (100, 100)]),
                'learning_rate_init': trial.suggest_float('learning_rate_init', 0.0001, 0.01),
                'alpha': trial.suggest_float('alpha', 0.0001, 0.1),
                'max_iter': trial.suggest_int('max_iter', 200, 1000)
            }
        else:
            return {}
    
    def _create_model(self, model_name: str, params: Dict[str, Any]):
        """Create model instance with given parameters"""
        if model_name == 'random_forest':
            from sklearn.ensemble import RandomForestRegressor
            return RandomForestRegressor(random_state=self.config.random_state, **params)
        elif model_name == 'xgboost':
            import xgboost as xgb
            return xgb.XGBRegressor(random_state=self.config.random_state, **params)
        elif model_name == 'neural_network':
            from sklearn.neural_network import MLPRegressor
            return MLPRegressor(random_state=self.config.random_state, **params)
        else:
            raise ValueError(f"Unknown model: {model_name}")
    
    def _train_and_evaluate_models(self, splits: Dict[str, Any]):
        """Train and evaluate all models"""
        results = {}
        
        for model_name in self.config.models_to_test:
            logger.info(f"Training {model_name}")
            
            try:
                # Get best parameters from optimization
                if self.config.enable_optuna and hasattr(self, 'study'):
                    best_params = {k: v for k, v in self.study.best_params.items() 
                                 if k != 'model' and self.study.best_params.get('model') == model_name}
                else:
                    best_params = {}
                
                # Create and train model
                model = self._create_model(model_name, best_params)
                model.fit(splits['X_train'], splits['y_train'])
                
                # Evaluate on test set
                y_pred = model.predict(splits['X_test'])
                
                # Calculate metrics
                metrics = self._calculate_metrics(splits['y_test'], y_pred)
                results[model_name] = {
                    'model': model,
                    'metrics': metrics,
                    'predictions': y_pred
                }
                
                # Log to MLflow
                if self.config.enable_mlflow:
                    with mlflow.start_run(nested=True):
                        mlflow.log_params(best_params)
                        for metric_name, value in metrics.items():
                            mlflow.log_metric(metric_name, value)
                        mlflow.sklearn.log_model(model, f"{model_name}_model")
                
                logger.info(f"{model_name} - R²: {metrics['r2_score']:.4f}, RMSE: {metrics['rmse']:.4f}")
                
            except Exception as e:
                logger.error(f"Failed to train {model_name}: {e}")
                results[model_name] = {'error': str(e)}
        
        self.results['model_evaluation'] = results
    
    def _calculate_metrics(self, y_true: np.ndarray, y_pred: np.ndarray) -> Dict[str, float]:
        """Calculate comprehensive evaluation metrics"""
        return {
            'r2_score': r2_score(y_true, y_pred),
            'rmse': np.sqrt(mean_squared_error(y_true, y_pred)),
            'mae': mean_absolute_error(y_true, y_pred),
            'spearman_corr': spearmanr(y_true, y_pred)[0],
            'mse': mean_squared_error(y_true, y_pred)
        }
    
    def _run_nested_cv(self, X: pd.DataFrame, y: pd.Series):
        """Run nested cross-validation for unbiased performance estimation"""
        logger.info("Running nested cross-validation")
        
        outer_cv = KFold(n_splits=self.config.nested_cv_folds, shuffle=True, 
                        random_state=self.config.random_state)
        inner_cv = KFold(n_splits=self.config.cv_folds, shuffle=True, 
                        random_state=self.config.random_state)
        
        nested_scores = {}
        
        for model_name in self.config.models_to_test:
            try:
                # Nested CV for each model
                scores = cross_val_score(
                    self._create_model(model_name, {}),
                    X, y,
                    cv=outer_cv,
                    scoring='r2',
                    n_jobs=-1
                )
                nested_scores[model_name] = {
                    'mean_score': scores.mean(),
                    'std_score': scores.std(),
                    'scores': scores.tolist()
                }
                
                logger.info(f"{model_name} nested CV: {scores.mean():.4f} ± {scores.std():.4f}")
                
            except Exception as e:
                logger.error(f"Nested CV failed for {model_name}: {e}")
                nested_scores[model_name] = {'error': str(e)}
        
        self.results['nested_cv'] = nested_scores
        
        # Log to MLflow
        if self.config.enable_mlflow:
            for model_name, scores in nested_scores.items():
                if 'error' not in scores:
                    mlflow.log_metric(f"{model_name}_nested_cv_mean", scores['mean_score'])
                    mlflow.log_metric(f"{model_name}_nested_cv_std", scores['std_score'])
    
    def _run_feature_ablation(self, splits: Dict[str, Any]):
        """Run feature ablation studies"""
        logger.info("Running feature ablation studies")
        
        # This would integrate with existing ablation study framework
        # For now, create a placeholder
        ablation_results = {
            'feature_importance': {},
            'ablation_scores': {}
        }
        
        self.results['feature_ablation'] = ablation_results
    
    def _run_uncertainty_quantification(self, splits: Dict[str, Any]):
        """Run uncertainty quantification analysis"""
        logger.info("Running uncertainty quantification")
        
        # This would integrate with existing uncertainty quantification framework
        uncertainty_results = {
            'prediction_intervals': {},
            'confidence_scores': {}
        }
        
        self.results['uncertainty_quantification'] = uncertainty_results
    
    def _run_interpretability_analysis(self, splits: Dict[str, Any]):
        """Run model interpretability analysis"""
        logger.info("Running interpretability analysis")
        
        # This would integrate with existing interpretability framework
        interpretability_results = {
            'feature_importance': {},
            'shap_values': {},
            'lime_explanations': {}
        }
        
        self.results['interpretability'] = interpretability_results
    
    def _train_ensemble_models(self, splits: Dict[str, Any]):
        """Train ensemble models"""
        logger.info("Training ensemble models")
        
        # This would integrate with existing ensemble framework
        ensemble_results = {
            'stacked_ensemble': {},
            'weighted_ensemble': {},
            'voting_ensemble': {}
        }
        
        self.results['ensemble_models'] = ensemble_results
    
    def _generate_research_report(self):
        """Generate comprehensive research report"""
        logger.info("Generating research report")
        
        report_path = Path(self.config.output_path) / "research_report.md"
        report_path.parent.mkdir(parents=True, exist_ok=True)
        
        # Generate markdown report
        report_content = self._create_markdown_report()
        
        with open(report_path, 'w') as f:
            f.write(report_content)
        
        # Log report to MLflow
        if self.config.enable_mlflow:
            mlflow.log_artifact(str(report_path))
        
        logger.info(f"Research report saved to: {report_path}")
    
    def _create_markdown_report(self) -> str:
        """Create comprehensive markdown research report"""
        report = f"""# Research Report: {self.config.experiment_name}

## Experiment Overview
- **Description**: {self.config.description}
- **Date**: {datetime.now().strftime('%Y-%m-%d %H:%M:%S')}
- **Random State**: {self.config.random_state}
- **CV Folds**: {self.config.cv_folds}

## Model Performance Results

### Individual Models
"""
        
        if 'model_evaluation' in self.results:
            for model_name, result in self.results['model_evaluation'].items():
                if 'error' not in result:
                    metrics = result['metrics']
                    report += f"""
#### {model_name.replace('_', ' ').title()}
- R² Score: {metrics['r2_score']:.4f}
- RMSE: {metrics['rmse']:.4f}
- MAE: {metrics['mae']:.4f}
- Spearman Correlation: {metrics['spearman_corr']:.4f}
"""
        
        if 'nested_cv' in self.results:
            report += "\n### Nested Cross-Validation Results\n"
            for model_name, scores in self.results['nested_cv'].items():
                if 'error' not in scores:
                    report += f"- **{model_name}**: {scores['mean_score']:.4f} ± {scores['std_score']:.4f}\n"
        
        report += f"""
## Configuration
- **Models Tested**: {', '.join(self.config.models_to_test)}
- **Hyperparameter Optimization**: {'Enabled' if self.config.enable_hyperparameter_optimization else 'Disabled'}
- **Nested CV**: {'Enabled' if self.config.enable_nested_cv else 'Disabled'}
- **Feature Ablation**: {'Enabled' if self.config.enable_feature_ablation else 'Disabled'}
- **Uncertainty Quantification**: {'Enabled' if self.config.enable_uncertainty_quantification else 'Disabled'}
- **Interpretability Analysis**: {'Enabled' if self.config.enable_interpretability else 'Disabled'}
- **Ensemble Methods**: {'Enabled' if self.config.enable_ensemble else 'Disabled'}

## Files and Artifacts
- **MLflow Experiment ID**: {self.experiment_id}
- **MLflow Run ID**: {self.run_id}
- **Output Directory**: {self.config.output_path}

---
*Report generated by TLR4 Binding Prediction Research Pipeline*
"""
        
        return report
    
    def save_config(self, filepath: str):
        """Save experiment configuration to file"""
        config_dict = asdict(self.config)
        
        with open(filepath, 'w') as f:
            yaml.dump(config_dict, f, default_flow_style=False)
        
        logger.info(f"Configuration saved to: {filepath}")
    
    @classmethod
    def load_config(cls, filepath: str) -> 'ResearchPipelineOrchestrator':
        """Load experiment configuration from file"""
        with open(filepath, 'r') as f:
            config_dict = yaml.safe_load(f)
        
        config = ExperimentConfig(**config_dict)
        return cls(config)


def create_experiment_config(
    experiment_name: str,
    description: str,
    data_path: str,
    output_path: str,
    **kwargs
) -> ExperimentConfig:
    """Create experiment configuration with sensible defaults"""
    
    default_tags = {
        'project': 'tlr4_binding_prediction',
        'version': '1.0',
        'framework': 'scikit-learn',
        'optimization': 'optuna'
    }
    
    return ExperimentConfig(
        experiment_name=experiment_name,
        description=description,
        tags=default_tags,
        data_path=data_path,
        output_path=output_path,
        **kwargs
    )


# Example usage and testing
if __name__ == "__main__":
    # Create sample configuration
    config = create_experiment_config(
        experiment_name="tlr4_binding_baseline",
        description="Baseline TLR4 binding affinity prediction experiment",
        data_path="./data",
        output_path="./results",
        n_trials=50,
        cv_folds=5
    )
    
    # Create orchestrator
    orchestrator = ResearchPipelineOrchestrator(config)
    
    # Save configuration
    orchestrator.save_config("./experiment_config.yaml")
    
    print("Research Pipeline Orchestrator created successfully!")
    print(f"Configuration saved to: ./experiment_config.yaml")
    print(f"MLflow tracking URI: {config.mlflow_tracking_uri}")
    print(f"Optuna storage: {config.optuna_storage}")
```
