## Supplementary material for "Machine Learning-Driven Prediction of TLR4 Binding Affinity: A Comprehensive Molecular Feature Analysis for Drug Discovery": Codebase: predictor.py

```
"""
Prediction interface and uncertainty quantification.

This module provides prediction functionality for trained models
including uncertainty estimation and confidence intervals.
"""

from abc import ABC, abstractmethod
from typing import Dict, List, Optional, Union, Tuple, Any
import pandas as pd
import numpy as np
import logging
from pathlib import Path
import joblib
from datetime import datetime

from ..molecular_analysis.features import PredictionResult, MolecularFeatures
from .evaluator import PerformanceMetrics

logger = logging.getLogger(__name__)


class UncertaintyEstimatorInterface(ABC):
    """Abstract interface for uncertainty estimation."""
    
    @abstractmethod
    def estimate_uncertainty(self, model: Any, X: pd.DataFrame) -> Dict[str, np.ndarray]:
        """Estimate prediction uncertainty."""
        pass


class BootstrapUncertaintyEstimator(UncertaintyEstimatorInterface):
    """Bootstrap-based uncertainty estimation."""
    
    def __init__(self, n_bootstrap: int = 100, random_state: int = 42):
        """
        Initialize bootstrap uncertainty estimator.
        
        Args:
            n_bootstrap: Number of bootstrap samples
            random_state: Random state for reproducibility
        """
        self.n_bootstrap = n_bootstrap
        self.random_state = random_state
    
    def estimate_uncertainty(self, model: Any, X: pd.DataFrame) -> Dict[str, np.ndarray]:
        """
        Estimate uncertainty using bootstrap sampling.
        
        Args:
            model: Trained model
            X: Feature matrix
            
        Returns:
            Dictionary with uncertainty estimates
        """
        np.random.seed(self.random_state)
        
        predictions = []
        
        for i in range(self.n_bootstrap):
            # Bootstrap sample
            bootstrap_indices = np.random.choice(
                len(X), size=len(X), replace=True
            )
            X_bootstrap = X.iloc[bootstrap_indices]
            
            # Make prediction
            pred = model.predict(X_bootstrap)
            predictions.append(pred)
        
        predictions = np.array(predictions)
        
        # Calculate statistics
        mean_predictions = np.mean(predictions, axis=0)
        std_predictions = np.std(predictions, axis=0)
        
        # Confidence intervals
        alpha = 0.05  # 95% confidence interval
        lower_percentile = (alpha / 2) * 100
        upper_percentile = (1 - alpha / 2) * 100
        
        lower_bound = np.percentile(predictions, lower_percentile, axis=0)
        upper_bound = np.percentile(predictions, upper_percentile, axis=0)
        
        return {
            'mean_predictions': mean_predictions,
            'std_predictions': std_predictions,
            'lower_bound': lower_bound,
            'upper_bound': upper_bound,
            'confidence_interval_width': upper_bound - lower_bound
        }


class DropoutUncertaintyEstimator(UncertaintyEstimatorInterface):
    """Monte Carlo Dropout uncertainty estimation for neural networks."""
    
    def __init__(self, n_samples: int = 100):
        """
        Initialize dropout uncertainty estimator.
        
        Args:
            n_samples: Number of Monte Carlo samples
        """
        self.n_samples = n_samples
    
    def estimate_uncertainty(self, model: Any, X: pd.DataFrame) -> Dict[str, np.ndarray]:
        """
        Estimate uncertainty using Monte Carlo Dropout.
        
        Args:
            model: Trained neural network model
            X: Feature matrix
            
        Returns:
            Dictionary with uncertainty estimates
        """
        # This is a placeholder for neural network dropout uncertainty
        # Implementation would depend on the specific neural network framework
        
        # For now, return dummy uncertainty estimates
        predictions = model.predict(X)
        
        return {
            'mean_predictions': predictions,
            'std_predictions': np.zeros_like(predictions),
            'lower_bound': predictions - 0.1,
            'upper_bound': predictions + 0.1,
            'confidence_interval_width': np.full_like(predictions, 0.2)
        }


class BindingPredictorInterface(ABC):
    """Abstract interface for binding prediction."""
    
    @abstractmethod
    def predict_single(self, features: MolecularFeatures) -> PredictionResult:
        """Predict binding affinity for single compound."""
        pass
    
    @abstractmethod
    def predict_batch(self, features_list: List[MolecularFeatures]) -> List[PredictionResult]:
        """Predict binding affinities for multiple compounds."""
        pass
    
    @abstractmethod
    def get_feature_importance(self) -> Dict[str, float]:
        """Get feature importance from best model."""
        pass


class BindingPredictor(BindingPredictorInterface):
    """
    Main binding prediction interface.
    
    Provides user-friendly interface for predicting binding affinities
    of new compounds using trained models.
    """
    
    def __init__(self, model_path: Optional[str] = None,
                 uncertainty_estimator: Optional[UncertaintyEstimatorInterface] = None):
        """
        Initialize binding predictor.
        
        Args:
            model_path: Path to trained model file
            uncertainty_estimator: Uncertainty estimation method
        """
        self.model_path = model_path
        self.model = None
        self.model_name = None
        self.feature_names = None
        self.uncertainty_estimator = uncertainty_estimator or BootstrapUncertaintyEstimator()
        
        if model_path:
            self.load_model(model_path)
    
    def load_model(self, model_path: str) -> None:
        """
        Load trained model from file.
        
        Args:
            model_path: Path to model file
        """
        model_path = Path(model_path)
        
        if not model_path.exists():
            raise FileNotFoundError(f"Model file not found: {model_path}")
        
        try:
            # Load model
            self.model = joblib.load(model_path)
            self.model_path = str(model_path)
            self.model_name = model_path.stem
            
            # Extract feature names if available
            if hasattr(self.model, 'feature_names_in_'):
                self.feature_names = self.model.feature_names_in_.tolist()
            
            logger.info(f"Model loaded successfully from {model_path}")
            
        except Exception as e:
            logger.error(f"Error loading model from {model_path}: {str(e)}")
            raise ValueError(f"Failed to load model: {str(e)}")
    
    def predict_single(self, features: MolecularFeatures) -> PredictionResult:
        """
        Predict binding affinity for single compound.
        
        Args:
            features: MolecularFeatures object
            
        Returns:
            PredictionResult with prediction and uncertainty
        """
        if self.model is None:
            raise ValueError("No model loaded. Call load_model() first.")
        
        # Convert features to DataFrame
        features_df = pd.DataFrame([features.to_dict()])
        
        # Select only model features if available
        if self.feature_names:
            available_features = [f for f in self.feature_names if f in features_df.columns]
            if not available_features:
                raise ValueError("No matching features found between model and input")
            features_df = features_df[available_features]
        
        # Make prediction
        prediction = self.model.predict(features_df)[0]
        
        # Estimate uncertainty
        uncertainty_results = self.uncertainty_estimator.estimate_uncertainty(
            self.model, features_df
        )
        
        # Create prediction result
        result = PredictionResult(
            compound_name=features.compound_name,
            predicted_affinity=float(prediction),
            confidence_interval_lower=float(uncertainty_results['lower_bound'][0]),
            confidence_interval_upper=float(uncertainty_results['upper_bound'][0]),
            model_used=self.model_name or "unknown",
            prediction_uncertainty=float(uncertainty_results['std_predictions'][0]),
            model_confidence=float(1.0 - uncertainty_results['confidence_interval_width'][0] / 10.0),
            prediction_timestamp=datetime.now().isoformat()
        )
        
        # Add feature contributions if available
        if hasattr(self.model, 'feature_importances_'):
            feature_importance = dict(zip(
                self.feature_names or features_df.columns,
                self.model.feature_importances_
            ))
            result.feature_contributions = feature_importance
        
        return result
    
    def predict_batch(self, features_list: List[MolecularFeatures]) -> List[PredictionResult]:
        """
        Predict binding affinities for multiple compounds.
        
        Args:
            features_list: List of MolecularFeatures objects
            
        Returns:
            List of PredictionResult objects
        """
        if self.model is None:
            raise ValueError("No model loaded. Call load_model() first.")
        
        if not features_list:
            return []
        
        # Convert features to DataFrame
        features_data = [features.to_dict() for features in features_list]
        features_df = pd.DataFrame(features_data)
        
        # Select only model features if available
        if self.feature_names:
            available_features = [f for f in self.feature_names if f in features_df.columns]
            if not available_features:
                raise ValueError("No matching features found between model and input")
            features_df = features_df[available_features]
        
        # Make predictions
        predictions = self.model.predict(features_df)
        
        # Estimate uncertainty
        uncertainty_results = self.uncertainty_estimator.estimate_uncertainty(
            self.model, features_df
        )
        
        # Create prediction results
        results = []
        for i, features in enumerate(features_list):
            result = PredictionResult(
                compound_name=features.compound_name,
                predicted_affinity=float(predictions[i]),
                confidence_interval_lower=float(uncertainty_results['lower_bound'][i]),
                confidence_interval_upper=float(uncertainty_results['upper_bound'][i]),
                model_used=self.model_name or "unknown",
                prediction_uncertainty=float(uncertainty_results['std_predictions'][i]),
                model_confidence=float(1.0 - uncertainty_results['confidence_interval_width'][i] / 10.0),
                prediction_timestamp=datetime.now().isoformat()
            )
            
            # Add feature contributions if available
            if hasattr(self.model, 'feature_importances_'):
                feature_importance = dict(zip(
                    self.feature_names or features_df.columns,
                    self.model.feature_importances_
                ))
                result.feature_contributions = feature_importance
            
            results.append(result)
        
        return results
    
    def get_feature_importance(self) -> Dict[str, float]:
        """Get feature importance from best model."""
        if self.model is None:
            raise ValueError("No model loaded. Call load_model() first.")
        
        if hasattr(self.model, 'feature_importances_'):
            if self.feature_names:
                return dict(zip(self.feature_names, self.model.feature_importances_))
            else:
                return dict(zip(range(len(self.model.feature_importances_)), 
                              self.model.feature_importances_))
        else:
            logger.warning("Model does not support feature importance")
            return {}
    
    def predict_from_dataframe(self, features_df: pd.DataFrame) -> pd.DataFrame:
        """
        Predict binding affinities from DataFrame.
        
        Args:
            features_df: DataFrame with molecular features
            
        Returns:
            DataFrame with predictions and uncertainty
        """
        if self.model is None:
            raise ValueError("No model loaded. Call load_model() first.")
        
        # Select only model features if available
        if self.feature_names:
            available_features = [f for f in self.feature_names if f in features_df.columns]
            if not available_features:
                raise ValueError("No matching features found between model and input")
            features_df = features_df[available_features]
        
        # Make predictions
        predictions = self.model.predict(features_df)
        
        # Estimate uncertainty
        uncertainty_results = self.uncertainty_estimator.estimate_uncertainty(
            self.model, features_df
        )
        
        # Create results DataFrame
        results_df = pd.DataFrame({
            'predicted_affinity': predictions,
            'confidence_interval_lower': uncertainty_results['lower_bound'],
            'confidence_interval_upper': uncertainty_results['upper_bound'],
            'prediction_uncertainty': uncertainty_results['std_predictions'],
            'model_confidence': 1.0 - uncertainty_results['confidence_interval_width'] / 10.0,
            'model_used': self.model_name or "unknown",
            'prediction_timestamp': datetime.now().isoformat()
        })
        
        # Add compound names if available
        if 'compound_name' in features_df.columns:
            results_df['compound_name'] = features_df['compound_name'].values
        
        return results_df
    
    def save_predictions(self, results: List[PredictionResult], 
                        output_path: str) -> None:
        """
        Save predictions to CSV file.
        
        Args:
            results: List of PredictionResult objects
            output_path: Path to save CSV file
        """
        if not results:
            logger.warning("No predictions to save")
            return
        
        # Convert to DataFrame
        results_data = [result.to_dict() for result in results]
        results_df = pd.DataFrame(results_data)
        
        # Save to CSV
        output_path = Path(output_path)
        output_path.parent.mkdir(parents=True, exist_ok=True)
        results_df.to_csv(output_path, index=False)
        
        logger.info(f"Predictions saved to {output_path}")
    
    def get_model_info(self) -> Dict[str, Any]:
        """Get information about loaded model."""
        if self.model is None:
            return {'error': 'No model loaded'}
        
        info = {
            'model_path': self.model_path,
            'model_name': self.model_name,
            'model_type': type(self.model).__name__,
            'feature_count': len(self.feature_names) if self.feature_names else 'unknown',
            'has_feature_importance': hasattr(self.model, 'feature_importances_'),
            'uncertainty_estimator': type(self.uncertainty_estimator).__name__
        }
        
        return info
```
