## Supplementary material for "Machine Learning-Driven Prediction of TLR4 Binding Affinity: A Comprehensive Molecular Feature Analysis for Drug Discovery": Codebase: statistical_validation.py

```
"""
Statistical validation framework for model comparison and significance testing.

This module provides comprehensive statistical validation tools for comparing
model performance and ensuring scientific rigor in molecular property prediction.
"""

import numpy as np
import pandas as pd
import logging
from typing import Dict, List, Optional, Tuple, Union, Any
from dataclasses import dataclass
from scipy import stats
from sklearn.model_selection import cross_val_score
from sklearn.metrics import mean_squared_error, r2_score, mean_absolute_error
import warnings

logger = logging.getLogger(__name__)


@dataclass
class StatisticalResult:
    """Container for statistical test results."""
    test_name: str
    statistic: float
    p_value: float
    effect_size: Optional[float] = None
    confidence_interval: Optional[Tuple[float, float]] = None
    interpretation: Optional[str] = None
    significant: Optional[bool] = None


@dataclass
class ModelComparisonResult:
    """Container for model comparison results."""
    model1_name: str
    model2_name: str
    model1_scores: np.ndarray
    model2_scores: np.ndarray
    statistical_tests: List[StatisticalResult]
    summary: Dict[str, Any]


class StatisticalValidator:
    """
    Comprehensive statistical validation framework for model comparison.
    
    Provides various statistical tests to validate model performance differences
    and ensure scientific rigor in machine learning experiments.
    """
    
    def __init__(self, alpha: float = 0.05, n_bootstrap: int = 1000):
        """
        Initialize statistical validator.
        
        Args:
            alpha: Significance level for statistical tests
            n_bootstrap: Number of bootstrap samples for confidence intervals
        """
        self.alpha = alpha
        self.n_bootstrap = n_bootstrap
    
    def compare_models(self, 
                      model1_scores: Union[np.ndarray, List[float]],
                      model2_scores: Union[np.ndarray, List[float]],
                      model1_name: str = "Model 1",
                      model2_name: str = "Model 2") -> ModelComparisonResult:
        """
        Comprehensive comparison of two models using multiple statistical tests.
        
        Args:
            model1_scores: Performance scores for first model
            model2_scores: Performance scores for second model
            model1_name: Name of first model
            model2_name: Name of second model
            
        Returns:
            ModelComparisonResult with comprehensive comparison
        """
        model1_scores = np.array(model1_scores)
        model2_scores = np.array(model2_scores)
        
        logger.info(f"Comparing {model1_name} vs {model2_name}")
        
        # Perform multiple statistical tests
        tests = []
        
        # 1. Paired t-test
        tests.append(self.paired_t_test(model1_scores, model2_scores))
        
        # 2. Wilcoxon signed-rank test (non-parametric)
        tests.append(self.wilcoxon_test(model1_scores, model2_scores))
        
        # 3. Bootstrap confidence interval for difference
        tests.append(self.bootstrap_difference_test(model1_scores, model2_scores))
        
        # 4. Effect size (Cohen's d)
        tests.append(self.cohens_d_test(model1_scores, model2_scores))
        
        # 5. Permutation test
        tests.append(self.permutation_test(model1_scores, model2_scores))
        
        # Create summary
        summary = self._create_comparison_summary(model1_scores, model2_scores, tests)
        
        return ModelComparisonResult(
            model1_name=model1_name,
            model2_name=model2_name,
            model1_scores=model1_scores,
            model2_scores=model2_scores,
            statistical_tests=tests,
            summary=summary
        )
    
    def paired_t_test(self, scores1: np.ndarray, scores2: np.ndarray) -> StatisticalResult:
        """
        Perform paired t-test for model comparison.
        
        Args:
            scores1: Scores for first model
            scores2: Scores for second model
            
        Returns:
            StatisticalResult with t-test results
        """
        if len(scores1) != len(scores2):
            raise ValueError("Score arrays must have the same length for paired t-test")
        
        # Calculate differences
        differences = scores1 - scores2
        
        # Perform t-test
        statistic, p_value = stats.ttest_rel(scores1, scores2)
        
        # Calculate confidence interval for mean difference
        mean_diff = np.mean(differences)
        se_diff = stats.sem(differences)
        ci = stats.t.interval(1 - self.alpha, len(differences) - 1, 
                             loc=mean_diff, scale=se_diff)
        
        # Interpretation
        significant = p_value < self.alpha
        if significant:
            if mean_diff > 0:
                interpretation = f"Model 1 significantly outperforms Model 2 (p={p_value:.4f})"
            else:
                interpretation = f"Model 2 significantly outperforms Model 1 (p={p_value:.4f})"
        else:
            interpretation = f"No significant difference between models (p={p_value:.4f})"
        
        return StatisticalResult(
            test_name="Paired t-test",
            statistic=statistic,
            p_value=p_value,
            confidence_interval=ci,
            interpretation=interpretation,
            significant=significant
        )
    
    def wilcoxon_test(self, scores1: np.ndarray, scores2: np.ndarray) -> StatisticalResult:
        """
        Perform Wilcoxon signed-rank test (non-parametric alternative to t-test).
        
        Args:
            scores1: Scores for first model
            scores2: Scores for second model
            
        Returns:
            StatisticalResult with Wilcoxon test results
        """
        if len(scores1) != len(scores2):
            raise ValueError("Score arrays must have the same length for Wilcoxon test")
        
        # Perform Wilcoxon test
        try:
            statistic, p_value = stats.wilcoxon(scores1, scores2, alternative='two-sided')
        except ValueError as e:
            # Handle case where all differences are zero
            return StatisticalResult(
                test_name="Wilcoxon signed-rank test",
                statistic=0.0,
                p_value=1.0,
                interpretation="All differences are zero - models perform identically",
                significant=False
            )
        
        # Interpretation
        significant = p_value < self.alpha
        median_diff = np.median(scores1 - scores2)
        
        if significant:
            if median_diff > 0:
                interpretation = f"Model 1 significantly outperforms Model 2 (Wilcoxon p={p_value:.4f})"
            else:
                interpretation = f"Model 2 significantly outperforms Model 1 (Wilcoxon p={p_value:.4f})"
        else:
            interpretation = f"No significant difference (Wilcoxon p={p_value:.4f})"
        
        return StatisticalResult(
            test_name="Wilcoxon signed-rank test",
            statistic=statistic,
            p_value=p_value,
            interpretation=interpretation,
            significant=significant
        )
    
    def bootstrap_difference_test(self, scores1: np.ndarray, scores2: np.ndarray) -> StatisticalResult:
        """
        Bootstrap confidence interval for the difference in means.
        
        Args:
            scores1: Scores for first model
            scores2: Scores for second model
            
        Returns:
            StatisticalResult with bootstrap results
        """
        differences = scores1 - scores2
        
        # Bootstrap sampling
        bootstrap_diffs = []
        for _ in range(self.n_bootstrap):
            bootstrap_sample = np.random.choice(differences, size=len(differences), replace=True)
            bootstrap_diffs.append(np.mean(bootstrap_sample))
        
        bootstrap_diffs = np.array(bootstrap_diffs)
        
        # Calculate confidence interval
        ci_lower = np.percentile(bootstrap_diffs, (self.alpha/2) * 100)
        ci_upper = np.percentile(bootstrap_diffs, (1 - self.alpha/2) * 100)
        
        # Check if CI contains zero
        significant = not (ci_lower <= 0 <= ci_upper)
        
        # Calculate p-value (proportion of bootstrap samples with opposite sign)
        mean_diff = np.mean(differences)
        if mean_diff >= 0:
            p_value = np.mean(bootstrap_diffs <= 0) * 2  # Two-tailed
        else:
            p_value = np.mean(bootstrap_diffs >= 0) * 2  # Two-tailed
        
        p_value = min(p_value, 1.0)  # Cap at 1.0
        
        interpretation = f"Bootstrap CI: [{ci_lower:.4f}, {ci_upper:.4f}]"
        if significant:
            interpretation += " (does not contain 0 - significant difference)"
        else:
            interpretation += " (contains 0 - no significant difference)"
        
        return StatisticalResult(
            test_name="Bootstrap difference test",
            statistic=mean_diff,
            p_value=p_value,
            confidence_interval=(ci_lower, ci_upper),
            interpretation=interpretation,
            significant=significant
        )
    
    def cohens_d_test(self, scores1: np.ndarray, scores2: np.ndarray) -> StatisticalResult:
        """
        Calculate Cohen's d effect size.
        
        Args:
            scores1: Scores for first model
            scores2: Scores for second model
            
        Returns:
            StatisticalResult with effect size
        """
        # Calculate Cohen's d
        mean1, mean2 = np.mean(scores1), np.mean(scores2)
        std1, std2 = np.std(scores1, ddof=1), np.std(scores2, ddof=1)
        
        # Pooled standard deviation
        n1, n2 = len(scores1), len(scores2)
        pooled_std = np.sqrt(((n1 - 1) * std1**2 + (n2 - 1) * std2**2) / (n1 + n2 - 2))
        
        cohens_d = (mean1 - mean2) / pooled_std
        
        # Interpret effect size
        abs_d = abs(cohens_d)
        if abs_d < 0.2:
            effect_interpretation = "negligible"
        elif abs_d < 0.5:
            effect_interpretation = "small"
        elif abs_d < 0.8:
            effect_interpretation = "medium"
        else:
            effect_interpretation = "large"
        
        interpretation = f"Cohen's d = {cohens_d:.3f} ({effect_interpretation} effect)"
        
        return StatisticalResult(
            test_name="Cohen's d effect size",
            statistic=cohens_d,
            p_value=None,  # Effect size doesn't have p-value
            effect_size=cohens_d,
            interpretation=interpretation,
            significant=abs_d >= 0.2  # Small effect or larger
        )
    
    def permutation_test(self, scores1: np.ndarray, scores2: np.ndarray, 
                        n_permutations: int = 10000) -> StatisticalResult:
        """
        Permutation test for difference in means.
        
        Args:
            scores1: Scores for first model
            scores2: Scores for second model
            n_permutations: Number of permutations
            
        Returns:
            StatisticalResult with permutation test results
        """
        # Observed difference
        observed_diff = np.mean(scores1) - np.mean(scores2)
        
        # Combine all scores
        all_scores = np.concatenate([scores1, scores2])
        n1 = len(scores1)
        
        # Permutation test
        permuted_diffs = []
        for _ in range(n_permutations):
            # Randomly permute the combined scores
            permuted = np.random.permutation(all_scores)
            perm_group1 = permuted[:n1]
            perm_group2 = permuted[n1:]
            
            perm_diff = np.mean(perm_group1) - np.mean(perm_group2)
            permuted_diffs.append(perm_diff)
        
        permuted_diffs = np.array(permuted_diffs)
        
        # Calculate p-value (two-tailed)
        p_value = np.mean(np.abs(permuted_diffs) >= np.abs(observed_diff))
        
        significant = p_value < self.alpha
        
        interpretation = f"Permutation test: observed difference = {observed_diff:.4f}, p = {p_value:.4f}"
        if significant:
            interpretation += " (significant)"
        else:
            interpretation += " (not significant)"
        
        return StatisticalResult(
            test_name="Permutation test",
            statistic=observed_diff,
            p_value=p_value,
            interpretation=interpretation,
            significant=significant
        )
    
    def _create_comparison_summary(self, scores1: np.ndarray, scores2: np.ndarray, 
                                 tests: List[StatisticalResult]) -> Dict[str, Any]:
        """Create summary of model comparison."""
        
        # Basic statistics
        summary = {
            'model1_stats': {
                'mean': np.mean(scores1),
                'std': np.std(scores1),
                'median': np.median(scores1),
                'min': np.min(scores1),
                'max': np.max(scores1)
            },
            'model2_stats': {
                'mean': np.mean(scores2),
                'std': np.std(scores2),
                'median': np.median(scores2),
                'min': np.min(scores2),
                'max': np.max(scores2)
            },
            'difference_stats': {
                'mean_difference': np.mean(scores1) - np.mean(scores2),
                'median_difference': np.median(scores1) - np.median(scores2),
                'std_difference': np.std(scores1 - scores2)
            }
        }
        
        # Test results summary
        significant_tests = [test for test in tests if test.significant]
        summary['statistical_summary'] = {
            'total_tests': len(tests),
            'significant_tests': len(significant_tests),
            'consensus': len(significant_tests) > len(tests) / 2,
            'test_names': [test.test_name for test in tests],
            'p_values': [test.p_value for test in tests if test.p_value is not None]
        }
        
        # Overall conclusion
        if summary['statistical_summary']['consensus']:
            if summary['difference_stats']['mean_difference'] > 0:
                summary['conclusion'] = "Model 1 significantly outperforms Model 2"
            else:
                summary['conclusion'] = "Model 2 significantly outperforms Model 1"
        else:
            summary['conclusion'] = "No significant difference between models"
        
        return summary
    
    def multiple_model_comparison(self, 
                                model_scores: Dict[str, np.ndarray],
                                correction_method: str = 'bonferroni') -> Dict[str, Any]:
        """
        Compare multiple models with multiple comparison correction.
        
        Args:
            model_scores: Dictionary mapping model names to score arrays
            correction_method: Method for multiple comparison correction
            
        Returns:
            Dictionary with pairwise comparison results
        """
        model_names = list(model_scores.keys())
        n_models = len(model_names)
        n_comparisons = n_models * (n_models - 1) // 2
        
        logger.info(f"Performing multiple model comparison ({n_comparisons} pairwise comparisons)")
        
        # Perform all pairwise comparisons
        pairwise_results = {}
        p_values = []
        
        for i in range(n_models):
            for j in range(i + 1, n_models):
                model1_name = model_names[i]
                model2_name = model_names[j]
                
                comparison = self.compare_models(
                    model_scores[model1_name],
                    model_scores[model2_name],
                    model1_name,
                    model2_name
                )
                
                pairwise_results[f"{model1_name}_vs_{model2_name}"] = comparison
                
                # Collect p-values for correction
                for test in comparison.statistical_tests:
                    if test.p_value is not None:
                        p_values.append(test.p_value)
        
        # Apply multiple comparison correction
        corrected_results = self._apply_multiple_comparison_correction(
            pairwise_results, correction_method
        )
        
        return {
            'pairwise_comparisons': pairwise_results,
            'corrected_results': corrected_results,
            'correction_method': correction_method,
            'n_comparisons': n_comparisons,
            'original_alpha': self.alpha,
            'corrected_alpha': self.alpha / n_comparisons if correction_method == 'bonferroni' else self.alpha
        }
    
    def _apply_multiple_comparison_correction(self, 
                                           pairwise_results: Dict[str, ModelComparisonResult],
                                           method: str) -> Dict[str, Any]:
        """Apply multiple comparison correction to p-values."""
        
        if method == 'bonferroni':
            n_comparisons = len(pairwise_results)
            corrected_alpha = self.alpha / n_comparisons
            
            corrected_results = {}
            for comparison_name, result in pairwise_results.items():
                corrected_tests = []
                for test in result.statistical_tests:
                    if test.p_value is not None:
                        corrected_test = StatisticalResult(
                            test_name=test.test_name + " (Bonferroni corrected)",
                            statistic=test.statistic,
                            p_value=test.p_value,
                            effect_size=test.effect_size,
                            confidence_interval=test.confidence_interval,
                            significant=test.p_value < corrected_alpha,
                            interpretation=test.interpretation + f" (corrected α={corrected_alpha:.4f})"
                        )
                        corrected_tests.append(corrected_test)
                
                corrected_results[comparison_name] = corrected_tests
            
            return {
                'method': 'bonferroni',
                'corrected_alpha': corrected_alpha,
                'corrected_tests': corrected_results
            }
        
        else:
            raise ValueError(f"Unknown correction method: {method}")


class BaselineComparator:
    """
    Compare models against established baselines.
    
    Provides comparison against simple baseline models to validate
    that complex models are actually providing value.
    """
    
    def __init__(self):
        """Initialize baseline comparator."""
        pass
    
    def create_baselines(self, y_train: np.ndarray, X_test: np.ndarray) -> Dict[str, np.ndarray]:
        """
        Create baseline predictions.
        
        Args:
            y_train: Training target values
            X_test: Test features
            
        Returns:
            Dictionary of baseline predictions
        """
        baselines = {}
        
        # Mean predictor
        baselines['mean_predictor'] = np.full(len(X_test), np.mean(y_train))
        
        # Median predictor
        baselines['median_predictor'] = np.full(len(X_test), np.median(y_train))
        
        # Random predictor (from training distribution)
        np.random.seed(42)
        baselines['random_predictor'] = np.random.choice(y_train, size=len(X_test))
        
        return baselines
    
    def compare_against_baselines(self, 
                                model_predictions: np.ndarray,
                                y_true: np.ndarray,
                                y_train: np.ndarray,
                                model_name: str = "Model") -> Dict[str, Any]:
        """
        Compare model against baseline predictors.
        
        Args:
            model_predictions: Model predictions
            y_true: True target values
            y_train: Training target values (for baseline creation)
            model_name: Name of the model
            
        Returns:
            Dictionary with baseline comparison results
        """
        # Create baselines
        baselines = self.create_baselines(y_train, np.zeros(len(y_true)))  # X_test not needed for simple baselines
        
        # Calculate metrics for model and baselines
        results = {}
        
        # Model metrics
        model_metrics = {
            'rmse': np.sqrt(mean_squared_error(y_true, model_predictions)),
            'mae': mean_absolute_error(y_true, model_predictions),
            'r2': r2_score(y_true, model_predictions)
        }
        results[model_name] = model_metrics
        
        # Baseline metrics
        for baseline_name, baseline_preds in baselines.items():
            baseline_metrics = {
                'rmse': np.sqrt(mean_squared_error(y_true, baseline_preds)),
                'mae': mean_absolute_error(y_true, baseline_preds),
                'r2': r2_score(y_true, baseline_preds)
            }
            results[baseline_name] = baseline_metrics
        
        # Calculate improvements
        improvements = {}
        for baseline_name in baselines.keys():
            improvements[baseline_name] = {
                'rmse_improvement': (results[baseline_name]['rmse'] - model_metrics['rmse']) / results[baseline_name]['rmse'],
                'mae_improvement': (results[baseline_name]['mae'] - model_metrics['mae']) / results[baseline_name]['mae'],
                'r2_improvement': model_metrics['r2'] - results[baseline_name]['r2']
            }
        
        return {
            'metrics': results,
            'improvements': improvements,
            'summary': {
                'beats_mean_predictor': model_metrics['r2'] > results['mean_predictor']['r2'],
                'beats_median_predictor': model_metrics['r2'] > results['median_predictor']['r2'],
                'beats_random_predictor': model_metrics['r2'] > results['random_predictor']['r2'],
                'best_baseline_r2': max(results[name]['r2'] for name in baselines.keys()),
                'model_r2': model_metrics['r2']
            }
        }


def validate_model_performance(model_scores: Dict[str, np.ndarray],
                             alpha: float = 0.05,
                             correction_method: str = 'bonferroni') -> Dict[str, Any]:
    """
    Comprehensive validation of model performance with statistical testing.
    
    Args:
        model_scores: Dictionary mapping model names to cross-validation scores
        alpha: Significance level
        correction_method: Multiple comparison correction method
        
    Returns:
        Dictionary with comprehensive validation results
    """
    validator = StatisticalValidator(alpha=alpha)
    
    # Multiple model comparison
    comparison_results = validator.multiple_model_comparison(model_scores, correction_method)
    
    # Find best performing model
    mean_scores = {name: np.mean(scores) for name, scores in model_scores.items()}
    best_model = max(mean_scores, key=mean_scores.get)
    
    # Summary statistics
    summary_stats = {}
    for name, scores in model_scores.items():
        summary_stats[name] = {
            'mean': np.mean(scores),
            'std': np.std(scores),
            'median': np.median(scores),
            'ci_lower': np.percentile(scores, 2.5),
            'ci_upper': np.percentile(scores, 97.5)
        }
    
    return {
        'comparison_results': comparison_results,
        'best_model': best_model,
        'summary_statistics': summary_stats,
        'validation_summary': {
            'n_models': len(model_scores),
            'n_comparisons': len(model_scores) * (len(model_scores) - 1) // 2,
            'best_model': best_model,
            'best_score': mean_scores[best_model],
            'score_range': max(mean_scores.values()) - min(mean_scores.values())
        }
    }
```
