## Supplementary material for "Machine Learning-Driven Prediction of TLR4 Binding Affinity: A Comprehensive Molecular Feature Analysis for Drug Discovery": Codebase: trainer.py

```
"""
Machine learning model training and optimization.

This module provides comprehensive model training functionality
for TLR4 binding prediction using various ML algorithms.
"""

from abc import ABC, abstractmethod
from typing import Dict, List, Optional, Union, Any, Tuple
import pandas as pd
import numpy as np
import logging
from pathlib import Path
import joblib
from datetime import datetime

logger = logging.getLogger(__name__)

# ML library imports with error handling
try:
    from sklearn.ensemble import RandomForestRegressor
    from sklearn.svm import SVR
    from sklearn.model_selection import GridSearchCV, cross_val_score
    from sklearn.metrics import mean_squared_error, r2_score, mean_absolute_error
    SKLEARN_AVAILABLE = True
except ImportError:
    logger.warning("scikit-learn not available. ML training will be limited.")
    SKLEARN_AVAILABLE = False

try:
    import xgboost as xgb
    XGBOOST_AVAILABLE = True
except ImportError:
    logger.warning("XGBoost not available.")
    XGBOOST_AVAILABLE = False

try:
    import lightgbm as lgb
    LIGHTGBM_AVAILABLE = True
except ImportError:
    logger.warning("LightGBM not available.")
    LIGHTGBM_AVAILABLE = False

# GNN imports with error handling
try:
    from .gnn_models import GNNModelTrainer, MolecularGraphBuilder
    GNN_AVAILABLE = True
except ImportError:
    logger.warning("GNN models not available.")
    GNN_AVAILABLE = False

# Deep learning imports with error handling
try:
    from .deep_learning_trainer import DeepLearningModelTrainer
    DEEP_LEARNING_AVAILABLE = True
except ImportError:
    logger.warning("Deep learning models not available.")
    DEEP_LEARNING_AVAILABLE = False


class ModelTrainerInterface(ABC):
    """Abstract interface for model training."""
    
    @abstractmethod
    def train(self, X_train: pd.DataFrame, y_train: pd.Series, 
              X_val: Optional[pd.DataFrame] = None, 
              y_val: Optional[pd.Series] = None) -> Any:
        """Train a single model."""
        pass
    
    @abstractmethod
    def predict(self, model: Any, X: pd.DataFrame) -> np.ndarray:
        """Make predictions using trained model."""
        pass
    
    @abstractmethod
    def get_feature_importance(self, model: Any) -> Dict[str, float]:
        """Get feature importance from trained model."""
        pass


class RandomForestTrainer(ModelTrainerInterface):
    """Random Forest regressor trainer with hyperparameter optimization."""
    
    def __init__(self, n_jobs: int = -1, random_state: int = 42):
        """
        Initialize Random Forest trainer.
        
        Args:
            n_jobs: Number of parallel jobs
            random_state: Random state for reproducibility
        """
        self.n_jobs = n_jobs
        self.random_state = random_state
        self.param_grid = {
            'n_estimators': [100, 200],
            'max_depth': [10, None],
            'min_samples_split': [2, 5],
            'min_samples_leaf': [1, 2],
            'max_features': ['sqrt', 'log2']
        }
    
    def train(self, X_train: pd.DataFrame, y_train: pd.Series, 
              X_val: Optional[pd.DataFrame] = None, 
              y_val: Optional[pd.Series] = None) -> RandomForestRegressor:
        """Train Random Forest model with hyperparameter optimization."""
        if not SKLEARN_AVAILABLE:
            raise RuntimeError("scikit-learn not available for Random Forest training")
        
        logger.info("Training Random Forest model")
        
        # Initialize model
        rf = RandomForestRegressor(
            n_jobs=self.n_jobs,
            random_state=self.random_state
        )
        
        # Perform grid search
        grid_search = GridSearchCV(
            rf, self.param_grid, 
            cv=5, scoring='neg_mean_squared_error',
            n_jobs=self.n_jobs, verbose=1
        )
        
        grid_search.fit(X_train, y_train)
        
        logger.info(f"Best Random Forest parameters: {grid_search.best_params_}")
        logger.info(f"Best CV score: {-grid_search.best_score_:.4f}")
        
        return grid_search.best_estimator_
    
    def predict(self, model: RandomForestRegressor, X: pd.DataFrame) -> np.ndarray:
        """Make predictions using trained Random Forest model."""
        return model.predict(X)
    
    def get_feature_importance(self, model: RandomForestRegressor) -> Dict[str, float]:
        """Get feature importance from Random Forest model."""
        if hasattr(model, 'feature_importances_'):
            return dict(zip(model.feature_names_in_, model.feature_importances_))
        return {}


class SVRTrainer(ModelTrainerInterface):
    """Support Vector Regression trainer with robust NaN handling."""
    
    def __init__(self, random_state: int = 42):
        """
        Initialize SVR trainer.
        
        Args:
            random_state: Random state for reproducibility
        """
        self.random_state = random_state
        self.param_grid = {
            'C': [0.1, 1, 10, 100],
            'gamma': ['scale', 'auto', 0.001, 0.01, 0.1, 1],
            'kernel': ['rbf', 'poly', 'linear'],
            'epsilon': [0.01, 0.1, 0.2, 0.5]
        }
        self.imputer = None
        self.scaler = None
    
    def train(self, X_train: pd.DataFrame, y_train: pd.Series, 
              X_val: Optional[pd.DataFrame] = None, 
              y_val: Optional[pd.Series] = None) -> SVR:
        """Train SVR model with robust NaN handling and preprocessing."""
        if not SKLEARN_AVAILABLE:
            raise RuntimeError("scikit-learn not available for SVR training")
        
        logger.info("Training SVR model with robust preprocessing")
        
        # Preprocess data to handle NaN values
        X_train_processed, y_train_processed = self._preprocess_data(X_train, y_train)
        
        # Initialize model (SVR doesn't accept random_state)
        svr = SVR()
        
        # Perform grid search with reduced parameter space for stability
        reduced_param_grid = {
            'C': [0.1, 1, 10],
            'gamma': ['scale', 'auto'],
            'kernel': ['rbf', 'linear'],
            'epsilon': [0.1, 0.2]
        }
        
        try:
            grid_search = GridSearchCV(
                svr, reduced_param_grid,
                cv=3, scoring='neg_mean_squared_error',  # Reduced CV folds for stability
                n_jobs=1, verbose=1  # Single job to avoid memory issues
            )
            
            grid_search.fit(X_train_processed, y_train_processed)
            
            logger.info(f"Best SVR parameters: {grid_search.best_params_}")
            logger.info(f"Best CV score: {-grid_search.best_score_:.4f}")
            
            return grid_search.best_estimator_
            
        except Exception as e:
            logger.warning(f"Grid search failed, using default SVR: {e}")
            # Fallback to default SVR with safe parameters
            default_svr = SVR(C=1.0, gamma='scale', kernel='rbf', epsilon=0.1)
            default_svr.fit(X_train_processed, y_train_processed)
            return default_svr
    
    def predict(self, model: SVR, X: pd.DataFrame) -> np.ndarray:
        """Make predictions using trained SVR model with preprocessing."""
        # Apply same preprocessing as training
        X_processed = self._preprocess_features(X)
        return model.predict(X_processed)
    
    def get_feature_importance(self, model: SVR) -> Dict[str, float]:
        """Get feature importance from SVR model using permutation importance."""
        # SVR doesn't have direct feature importance
        # Could implement permutation importance here
        return {}
    
    def _preprocess_data(self, X: pd.DataFrame, y: pd.Series) -> Tuple[np.ndarray, np.ndarray]:
        """
        Preprocess data for SVR training.
        
        Args:
            X: Features
            y: Targets
            
        Returns:
            Tuple of (processed_X, processed_y)
        """
        from sklearn.impute import SimpleImputer
        from sklearn.preprocessing import StandardScaler
        
        # Remove rows with NaN targets
        valid_mask = ~y.isna()
        X_clean = X[valid_mask].copy()
        y_clean = y[valid_mask].copy()
        
        logger.info(f"Removed {(~valid_mask).sum()} rows with NaN targets")
        
        # Get numeric columns only
        numeric_cols = X_clean.select_dtypes(include=[np.number]).columns
        X_numeric = X_clean[numeric_cols]
        
        # Impute missing values
        if self.imputer is None:
            self.imputer = SimpleImputer(strategy='median')
            X_imputed = self.imputer.fit_transform(X_numeric)
        else:
            X_imputed = self.imputer.transform(X_numeric)
        
        # Scale features
        if self.scaler is None:
            self.scaler = StandardScaler()
            X_scaled = self.scaler.fit_transform(X_imputed)
        else:
            X_scaled = self.scaler.transform(X_imputed)
        
        logger.info(f"Preprocessed data: {X_scaled.shape[0]} samples, {X_scaled.shape[1]} features")
        
        return X_scaled, y_clean.values
    
    def _preprocess_features(self, X: pd.DataFrame) -> np.ndarray:
        """
        Preprocess features for prediction.
        
        Args:
            X: Features to preprocess
            
        Returns:
            Preprocessed features
        """
        # Get numeric columns only
        numeric_cols = X.select_dtypes(include=[np.number]).columns
        X_numeric = X[numeric_cols]
        
        # Apply same preprocessing as training
        if self.imputer is not None:
            X_imputed = self.imputer.transform(X_numeric)
        else:
            X_imputed = X_numeric.values
        
        if self.scaler is not None:
            X_scaled = self.scaler.transform(X_imputed)
        else:
            X_scaled = X_imputed
        
        return X_scaled


class XGBoostTrainer(ModelTrainerInterface):
    """XGBoost regressor trainer with hyperparameter optimization."""
    
    def __init__(self, random_state: int = 42):
        """
        Initialize XGBoost trainer.
        
        Args:
            random_state: Random state for reproducibility
        """
        self.random_state = random_state
        self.param_grid = {
            'n_estimators': [100, 200],
            'max_depth': [3, 6],
            'learning_rate': [0.1, 0.2],
            'subsample': [0.8, 1.0],
            'colsample_bytree': [0.8, 1.0]
        }
    
    def train(self, X_train: pd.DataFrame, y_train: pd.Series, 
              X_val: Optional[pd.DataFrame] = None, 
              y_val: Optional[pd.Series] = None) -> Any:
        """Train XGBoost model with hyperparameter optimization."""
        if not XGBOOST_AVAILABLE:
            raise RuntimeError("XGBoost not available for training")
        
        logger.info("Training XGBoost model")
        
        # Initialize model
        xgb_model = xgb.XGBRegressor(
            random_state=self.random_state,
            n_jobs=-1
        )
        
        # Perform grid search
        grid_search = GridSearchCV(
            xgb_model, self.param_grid,
            cv=5, scoring='neg_mean_squared_error',
            n_jobs=-1, verbose=1
        )
        
        grid_search.fit(X_train, y_train)
        
        logger.info(f"Best XGBoost parameters: {grid_search.best_params_}")
        logger.info(f"Best CV score: {-grid_search.best_score_:.4f}")
        
        return grid_search.best_estimator_
    
    def predict(self, model: Any, X: pd.DataFrame) -> np.ndarray:
        """Make predictions using trained XGBoost model."""
        return model.predict(X)
    
    def get_feature_importance(self, model: Any) -> Dict[str, float]:
        """Get feature importance from XGBoost model."""
        if hasattr(model, 'feature_importances_'):
            return dict(zip(model.feature_names_in_, model.feature_importances_))
        return {}


class LightGBMTrainer(ModelTrainerInterface):
    """LightGBM regressor trainer with hyperparameter optimization."""
    
    def __init__(self, random_state: int = 42):
        """
        Initialize LightGBM trainer.
        
        Args:
            random_state: Random state for reproducibility
        """
        self.random_state = random_state
        self.param_grid = {
            'n_estimators': [100, 200],
            'max_depth': [3, 6],
            'learning_rate': [0.1, 0.2],
            'subsample': [0.8, 1.0],
            'colsample_bytree': [0.8, 1.0],
            'num_leaves': [31, 50]
        }
    
    def train(self, X_train: pd.DataFrame, y_train: pd.Series, 
              X_val: Optional[pd.DataFrame] = None, 
              y_val: Optional[pd.Series] = None) -> Any:
        """Train LightGBM model with hyperparameter optimization."""
        if not LIGHTGBM_AVAILABLE:
            raise RuntimeError("LightGBM not available for training")
        
        logger.info("Training LightGBM model")
        
        # Initialize model
        lgb_model = lgb.LGBMRegressor(
            random_state=self.random_state,
            n_jobs=-1,
            verbose=-1
        )
        
        # Perform grid search
        grid_search = GridSearchCV(
            lgb_model, self.param_grid,
            cv=5, scoring='neg_mean_squared_error',
            n_jobs=-1, verbose=1
        )
        
        grid_search.fit(X_train, y_train)
        
        logger.info(f"Best LightGBM parameters: {grid_search.best_params_}")
        logger.info(f"Best CV score: {-grid_search.best_score_:.4f}")
        
        return grid_search.best_estimator_
    
    def predict(self, model: Any, X: pd.DataFrame) -> np.ndarray:
        """Make predictions using trained LightGBM model."""
        return model.predict(X)
    
    def get_feature_importance(self, model: Any) -> Dict[str, float]:
        """Get feature importance from LightGBM model."""
        if hasattr(model, 'feature_importances_'):
            return dict(zip(model.feature_names_in_, model.feature_importances_))
        return {}


class MLModelTrainer:
    """
    Main machine learning model trainer coordinator.
    
    Orchestrates training of multiple ML algorithms and provides
    comprehensive model comparison and selection.
    """
    
    def __init__(self, models_dir: str = "models/trained", include_gnn: bool = True, 
                 include_deep_learning: bool = True):
        """
        Initialize ML model trainer.
        
        Args:
            models_dir: Directory to save trained models
            include_gnn: Whether to include GNN models
            include_deep_learning: Whether to include deep learning models
        """
        self.models_dir = Path(models_dir)
        self.models_dir.mkdir(parents=True, exist_ok=True)
        
        # Initialize trainers
        self.trainers = {}
        if SKLEARN_AVAILABLE:
            self.trainers['random_forest'] = RandomForestTrainer()
            self.trainers['svr'] = SVRTrainer()
        
        if XGBOOST_AVAILABLE:
            self.trainers['xgboost'] = XGBoostTrainer()
        
        if LIGHTGBM_AVAILABLE:
            self.trainers['lightgbm'] = LightGBMTrainer()
        
        # Initialize GNN trainer if available
        self.gnn_trainer = None
        if include_gnn and GNN_AVAILABLE:
            gnn_models_dir = self.models_dir / "gnn"
            self.gnn_trainer = GNNModelTrainer(models_dir=str(gnn_models_dir))
        
        # Initialize deep learning trainer if available
        self.deep_learning_trainer = None
        if include_deep_learning and DEEP_LEARNING_AVAILABLE:
            dl_models_dir = self.models_dir / "deep_learning"
            self.deep_learning_trainer = DeepLearningModelTrainer(models_dir=str(dl_models_dir))
        
        self.trained_models = {}
        self.training_results = {}
        self.gnn_trained_models = {}
        self.gnn_training_results = {}
        self.deep_learning_trained_models = {}
        self.deep_learning_training_results = {}
    
    def train_models(self, X_train: pd.DataFrame, y_train: pd.Series,
                    X_val: Optional[pd.DataFrame] = None,
                    y_val: Optional[pd.Series] = None) -> Dict[str, Any]:
        """
        Train all available models.
        
        Args:
            X_train: Training features
            y_train: Training targets
            X_val: Validation features (optional)
            y_val: Validation targets (optional)
            
        Returns:
            Dictionary of trained models
        """
        logger.info(f"Training {len(self.trainers)} models")
        
        for model_name, trainer in self.trainers.items():
            try:
                logger.info(f"Training {model_name}")
                
                # Train model
                model = trainer.train(X_train, y_train, X_val, y_val)
                
                # Store trained model
                self.trained_models[model_name] = model
                
                # Calculate training metrics
                train_pred = trainer.predict(model, X_train)
                train_metrics = self._calculate_metrics(y_train, train_pred)
                
                self.training_results[model_name] = {
                    'model': model,
                    'train_metrics': train_metrics,
                    'trainer': trainer
                }
                
                # Save model
                self._save_model(model_name, model)
                
                logger.info(f"{model_name} training completed successfully")
                
            except Exception as e:
                logger.error(f"Error training {model_name}: {str(e)}")
                continue
        
        logger.info(f"Successfully trained {len(self.trained_models)} traditional ML models")
        return self.trained_models
    
    def train_deep_learning_models(self, pdbqt_files: List[str], binding_affinities: List[float],
                                  train_indices: List[int],
                                  features_df: Optional[pd.DataFrame] = None,
                                  smiles_list: Optional[List[str]] = None,
                                  compound_names: Optional[List[str]] = None,
                                  val_indices: Optional[List[int]] = None,
                                  model_types: List[str] = None, **kwargs) -> Dict[str, Any]:
        """
        Train deep learning models.
        
        Args:
            pdbqt_files: List of PDBQT file paths
            binding_affinities: List of binding affinities
            features_df: Optional DataFrame with molecular features
            smiles_list: Optional list of SMILES strings
            compound_names: Optional list of compound names
            train_indices: Indices for training set
            val_indices: Indices for validation set (optional)
            model_types: List of deep learning model types to train
            **kwargs: Additional training parameters
            
        Returns:
            Dictionary of trained deep learning models
        """
        if self.deep_learning_trainer is None:
            logger.warning("Deep learning trainer not available. Skipping deep learning model training.")
            return {}
        
        logger.info("Training deep learning models")
        
        # Train deep learning models
        dl_models = self.deep_learning_trainer.train_all_models(
            pdbqt_files, binding_affinities, features_df, smiles_list, compound_names,
            train_indices, val_indices, model_types, **kwargs
        )
        
        # Store results
        self.deep_learning_trained_models.update(dl_models)
        self.deep_learning_training_results.update(self.deep_learning_trainer.training_results)
        
        logger.info(f"Successfully trained {len(dl_models)} deep learning models")
        return dl_models
    
    def train_gnn_models(self, pdbqt_files: List[str], binding_affinities: List[float],
                        train_indices: List[int], val_indices: Optional[List[int]] = None,
                        test_indices: Optional[List[int]] = None,
                        model_types: List[str] = None, **kwargs) -> Dict[str, Any]:
        """
        Train GNN models on molecular graphs.
        
        Args:
            pdbqt_files: List of PDBQT file paths
            binding_affinities: List of binding affinities
            train_indices: Indices for training set
            val_indices: Indices for validation set (optional)
            test_indices: Indices for test set (optional)
            model_types: List of GNN model types to train
            **kwargs: Additional training parameters
            
        Returns:
            Dictionary of trained GNN models
        """
        if self.gnn_trainer is None:
            logger.warning("GNN trainer not available. Skipping GNN model training.")
            return {}
        
        logger.info("Training GNN models on molecular graphs")
        
        # Prepare training data
        train_pdbqt_files = [pdbqt_files[i] for i in train_indices]
        train_affinities = [binding_affinities[i] for i in train_indices]
        train_dataset = self.gnn_trainer.prepare_graph_data(train_pdbqt_files, train_affinities)
        
        # Prepare validation data if provided
        val_dataset = None
        if val_indices is not None:
            val_pdbqt_files = [pdbqt_files[i] for i in val_indices]
            val_affinities = [binding_affinities[i] for i in val_indices]
            val_dataset = self.gnn_trainer.prepare_graph_data(val_pdbqt_files, val_affinities)
        
        # Train GNN models
        gnn_models = self.gnn_trainer.train_models(
            train_dataset, val_dataset, model_types, **kwargs
        )
        
        # Store results
        self.gnn_trained_models.update(gnn_models)
        self.gnn_training_results.update(self.gnn_trainer.training_results)
        
        logger.info(f"Successfully trained {len(gnn_models)} GNN models")
        return gnn_models
    
    def evaluate_models(self, X_test: pd.DataFrame, y_test: pd.Series) -> Dict[str, Dict]:
        """
        Evaluate all trained models on test set.
        
        Args:
            X_test: Test features
            y_test: Test targets
            
        Returns:
            Dictionary of evaluation results for each model
        """
        logger.info("Evaluating models on test set")
        
        evaluation_results = {}
        
        for model_name, model in self.trained_models.items():
            try:
                trainer = self.training_results[model_name]['trainer']
                
                # Make predictions
                predictions = trainer.predict(model, X_test)
                
                # Calculate metrics
                metrics = self._calculate_metrics(y_test, predictions)
                
                # Get feature importance
                feature_importance = trainer.get_feature_importance(model)
                
                evaluation_results[model_name] = {
                    'predictions': predictions,
                    'metrics': metrics,
                    'feature_importance': feature_importance
                }
                
                logger.info(f"{model_name} evaluation completed")
                
            except Exception as e:
                logger.error(f"Error evaluating {model_name}: {str(e)}")
                continue
        
        return evaluation_results
    
    def evaluate_gnn_models(self, pdbqt_files: List[str], binding_affinities: List[float],
                           test_indices: List[int]) -> Dict[str, Dict]:
        """
        Evaluate GNN models on test set.
        
        Args:
            pdbqt_files: List of PDBQT file paths
            binding_affinities: List of binding affinities
            test_indices: Indices for test set
            
        Returns:
            Dictionary of GNN evaluation results
        """
        if self.gnn_trainer is None:
            logger.warning("GNN trainer not available. Skipping GNN model evaluation.")
            return {}
        
        logger.info("Evaluating GNN models on test set")
        
        # Prepare test data
        test_pdbqt_files = [pdbqt_files[i] for i in test_indices]
        test_affinities = [binding_affinities[i] for i in test_indices]
        test_dataset = self.gnn_trainer.prepare_graph_data(test_pdbqt_files, test_affinities)
        
        # Evaluate GNN models
        gnn_evaluation_results = self.gnn_trainer.evaluate_models(test_dataset)
        
        logger.info(f"Successfully evaluated {len(gnn_evaluation_results)} GNN models")
        return gnn_evaluation_results
    
    def evaluate_deep_learning_models(self, test_pdbqt_files: List[str], test_binding_affinities: List[float],
                                     test_indices: List[int],
                                     test_features_df: Optional[pd.DataFrame] = None,
                                     test_smiles_list: Optional[List[str]] = None,
                                     test_compound_names: Optional[List[str]] = None) -> Dict[str, Dict]:
        """
        Evaluate deep learning models on test set.
        
        Args:
            test_pdbqt_files: List of test PDBQT file paths
            test_binding_affinities: List of test binding affinities
            test_features_df: Optional DataFrame with test molecular features
            test_smiles_list: Optional list of test SMILES strings
            test_compound_names: Optional list of test compound names
            test_indices: Indices for test set
            
        Returns:
            Dictionary of deep learning evaluation results
        """
        if self.deep_learning_trainer is None:
            logger.warning("Deep learning trainer not available. Skipping deep learning model evaluation.")
            return {}
        
        logger.info("Evaluating deep learning models on test set")
        
        # Evaluate deep learning models
        dl_evaluation_results = self.deep_learning_trainer.evaluate_models(
            test_pdbqt_files, test_binding_affinities, test_features_df,
            test_smiles_list, test_compound_names, test_indices
        )
        
        logger.info(f"Successfully evaluated {len(dl_evaluation_results)} deep learning models")
        return dl_evaluation_results
    
    def get_best_model(self, metric: str = 'r2', include_gnn: bool = True, 
                      include_deep_learning: bool = True) -> Tuple[str, Any]:
        """
        Get the best performing model based on specified metric.
        
        Args:
            metric: Metric to use for model selection
            include_gnn: Whether to include GNN models in comparison
            include_deep_learning: Whether to include deep learning models in comparison
            
        Returns:
            Tuple of (model_name, model)
        """
        all_results = self.training_results.copy()
        
        # Include GNN results if available and requested
        if include_gnn and self.gnn_training_results:
            all_results.update(self.gnn_training_results)
        
        # Include deep learning results if available and requested
        if include_deep_learning and self.deep_learning_training_results:
            all_results.update(self.deep_learning_training_results)
        
        if not all_results:
            raise ValueError("No models have been trained yet")
        
        best_model_name = None
        best_score = float('-inf')
        
        for model_name, results in all_results.items():
            if results is None:
                continue
                
            # Handle different result structures
            if 'train_metrics' in results and metric in results['train_metrics']:
                score = results['train_metrics'][metric]
            elif 'history' in results and 'val_loss' in results['history']:
                # For deep learning models, use validation loss as score
                val_losses = results['history']['val_loss']
                if val_losses:
                    score = -min(val_losses)  # Convert loss to score
                else:
                    continue
            else:
                continue
                
            if score > best_score:
                best_score = score
                best_model_name = model_name
        
        if best_model_name is None:
            raise ValueError(f"No model found with metric {metric}")
        
        # Return model from appropriate collection
        if best_model_name in self.trained_models:
            return best_model_name, self.trained_models[best_model_name]
        elif best_model_name in self.gnn_trained_models:
            return best_model_name, self.gnn_trained_models[best_model_name]
        elif best_model_name in self.deep_learning_trained_models:
            return best_model_name, self.deep_learning_trained_models[best_model_name]
        else:
            raise ValueError(f"Model {best_model_name} not found in trained models")
    
    def _calculate_metrics(self, y_true: pd.Series, y_pred: np.ndarray) -> Dict[str, float]:
        """Calculate regression metrics."""
        if not SKLEARN_AVAILABLE:
            return {}
        
        return {
            'mse': mean_squared_error(y_true, y_pred),
            'rmse': np.sqrt(mean_squared_error(y_true, y_pred)),
            'mae': mean_absolute_error(y_true, y_pred),
            'r2': r2_score(y_true, y_pred)
        }
    
    def _save_model(self, model_name: str, model: Any) -> None:
        """Save trained model to disk."""
        model_path = self.models_dir / f"{model_name}_model.joblib"
        joblib.dump(model, model_path)
        logger.info(f"Model saved to {model_path}")
    
    def load_model(self, model_name: str) -> Any:
        """Load trained model from disk."""
        model_path = self.models_dir / f"{model_name}_model.joblib"
        if not model_path.exists():
            raise FileNotFoundError(f"Model file not found: {model_path}")
        
        return joblib.load(model_path)
    
    def get_model_summary(self, include_gnn: bool = True, include_deep_learning: bool = True) -> pd.DataFrame:
        """Get summary of all trained models."""
        all_results = self.training_results.copy()
        
        # Include GNN results if available and requested
        if include_gnn and self.gnn_training_results:
            all_results.update(self.gnn_training_results)
        
        # Include deep learning results if available and requested
        if include_deep_learning and self.deep_learning_training_results:
            all_results.update(self.deep_learning_training_results)
        
        if not all_results:
            return pd.DataFrame()
        
        summary_data = []
        for model_name, results in all_results.items():
            if results is None:
                continue
                
            # Determine model type
            if model_name in self.gnn_training_results:
                model_type = "GNN"
                metrics = results.get('train_metrics', {})
            elif model_name in self.deep_learning_training_results:
                model_type = "Deep Learning"
                # For deep learning models, extract metrics from history
                history = results.get('history', {})
                metrics = {
                    'final_train_loss': history.get('train_loss', [0.0])[-1] if history.get('train_loss') else 0.0,
                    'final_val_loss': history.get('val_loss', [0.0])[-1] if history.get('val_loss') else 0.0,
                    'best_val_loss': min(history.get('val_loss', [0.0])) if history.get('val_loss') else 0.0
                }
            else:
                model_type = "Traditional ML"
                metrics = results.get('train_metrics', {})
            
            summary_data.append({
                'model_name': model_name,
                'model_type': model_type,
                **metrics
            })
        
        return pd.DataFrame(summary_data)
```
