## Supplementary material for "Machine Learning-Driven Prediction of TLR4 Binding Affinity: A Comprehensive Molecular Feature Analysis for Drug Discovery": Codebase: uncertainty_quantification.py

```
"""
Uncertainty Quantification Methods for TLR4 Binding Prediction

This module implements various uncertainty quantification techniques including:
- Monte Carlo Dropout for neural networks
- Bootstrap aggregating for traditional ML models
- Conformal prediction intervals
- Ensemble-based uncertainty quantification
- Calibration analysis and reliability diagrams
"""

import numpy as np
import torch
import torch.nn as nn
from typing import Dict, List, Tuple, Optional, Union, Any
from sklearn.ensemble import RandomForestRegressor, GradientBoostingRegressor
from sklearn.model_selection import KFold
from sklearn.metrics import mean_squared_error, mean_absolute_error
import matplotlib.pyplot as plt
import seaborn as sns
from scipy import stats
import warnings
from dataclasses import dataclass
import logging

logger = logging.getLogger(__name__)


@dataclass
class UncertaintyResult:
    """Container for uncertainty quantification results"""
    predictions: np.ndarray
    uncertainties: np.ndarray
    confidence_intervals: Tuple[np.ndarray, np.ndarray]  # (lower, upper)
    prediction_intervals: Tuple[np.ndarray, np.ndarray]  # (lower, upper)
    epistemic_uncertainty: Optional[np.ndarray] = None
    aleatoric_uncertainty: Optional[np.ndarray] = None


class MonteCarloDropout:
    """
    Monte Carlo Dropout for uncertainty quantification in neural networks.
    
    This class enables dropout during inference to estimate model uncertainty
    by sampling multiple predictions with different dropout masks.
    """
    
    def __init__(self, model: nn.Module, n_samples: int = 100, dropout_rate: float = 0.1):
        """
        Initialize Monte Carlo Dropout.
        
        Args:
            model: PyTorch neural network model
            n_samples: Number of Monte Carlo samples for uncertainty estimation
            dropout_rate: Dropout rate to use during inference
        """
        self.model = model
        self.n_samples = n_samples
        self.dropout_rate = dropout_rate
        self.original_training_state = model.training
        
    def enable_dropout(self):
        """Enable dropout in all dropout layers"""
        for module in self.model.modules():
            if isinstance(module, nn.Dropout):
                module.train()
                module.p = self.dropout_rate
                
    def disable_dropout(self):
        """Disable dropout in all dropout layers"""
        for module in self.model.modules():
            if isinstance(module, nn.Dropout):
                module.eval()
                
    def predict_with_uncertainty(self, X: torch.Tensor) -> UncertaintyResult:
        """
        Make predictions with uncertainty quantification using Monte Carlo dropout.
        
        Args:
            X: Input tensor
            
        Returns:
            UncertaintyResult containing predictions and uncertainty estimates
        """
        self.model.eval()
        self.enable_dropout()
        
        predictions = []
        
        with torch.no_grad():
            for _ in range(self.n_samples):
                pred = self.model(X)
                predictions.append(pred.cpu().numpy())
                
        predictions = np.array(predictions)
        
        # Calculate statistics
        mean_predictions = np.mean(predictions, axis=0)
        epistemic_uncertainty = np.var(predictions, axis=0)
        
        # Calculate confidence intervals (95%)
        lower_ci = np.percentile(predictions, 2.5, axis=0)
        upper_ci = np.percentile(predictions, 97.5, axis=0)
        
        # Reset model state
        self.disable_dropout()
        if self.original_training_state:
            self.model.train()
            
        return UncertaintyResult(
            predictions=mean_predictions,
            uncertainties=epistemic_uncertainty,
            confidence_intervals=(lower_ci, upper_ci),
            prediction_intervals=(lower_ci, upper_ci),  # Same as confidence for MC dropout
            epistemic_uncertainty=epistemic_uncertainty
        )


class BootstrapUncertainty:
    """
    Bootstrap aggregating for uncertainty quantification in traditional ML models.
    
    This class uses bootstrap sampling to estimate prediction uncertainty
    by training multiple models on bootstrap samples of the training data.
    """
    
    def __init__(self, 
                 base_model_class,
                 n_bootstrap: int = 100,
                 bootstrap_ratio: float = 0.8,
                 random_state: int = 42,
                 **model_kwargs):
        """
        Initialize Bootstrap Uncertainty Quantification.
        
        Args:
            base_model_class: Scikit-learn model class
            n_bootstrap: Number of bootstrap samples
            bootstrap_ratio: Fraction of data to use in each bootstrap sample
            random_state: Random state for reproducibility
            **model_kwargs: Additional arguments for the base model
        """
        self.base_model_class = base_model_class
        self.n_bootstrap = n_bootstrap
        self.bootstrap_ratio = bootstrap_ratio
        self.random_state = random_state
        self.model_kwargs = model_kwargs
        self.bootstrap_models = []
        self.bootstrap_scores = []
        
    def fit(self, X: np.ndarray, y: np.ndarray) -> 'BootstrapUncertainty':
        """
        Fit bootstrap models.
        
        Args:
            X: Training features
            y: Training targets
            
        Returns:
            Self for method chaining
        """
        n_samples = int(len(X) * self.bootstrap_ratio)
        rng = np.random.RandomState(self.random_state)
        
        logger.info(f"Training {self.n_bootstrap} bootstrap models...")
        
        for i in range(self.n_bootstrap):
            # Create bootstrap sample
            bootstrap_indices = rng.choice(len(X), size=n_samples, replace=True)
            X_bootstrap = X[bootstrap_indices]
            y_bootstrap = y[bootstrap_indices]
            
            # Train model
            model = self.base_model_class(**self.model_kwargs)
            model.fit(X_bootstrap, y_bootstrap)
            self.bootstrap_models.append(model)
            
            # Calculate bootstrap score
            y_pred = model.predict(X_bootstrap)
            score = mean_squared_error(y_bootstrap, y_pred)
            self.bootstrap_scores.append(score)
            
            if (i + 1) % 10 == 0:
                logger.info(f"Trained {i + 1}/{self.n_bootstrap} bootstrap models")
                
        return self
        
    def predict_with_uncertainty(self, X: np.ndarray) -> UncertaintyResult:
        """
        Make predictions with uncertainty quantification.
        
        Args:
            X: Input features
            
        Returns:
            UncertaintyResult containing predictions and uncertainty estimates
        """
        if not self.bootstrap_models:
            raise ValueError("Must call fit() before predict_with_uncertainty()")
            
        predictions = []
        
        for model in self.bootstrap_models:
            pred = model.predict(X)
            predictions.append(pred)
            
        predictions = np.array(predictions)
        
        # Calculate statistics
        mean_predictions = np.mean(predictions, axis=0)
        epistemic_uncertainty = np.var(predictions, axis=0)
        
        # Estimate aleatoric uncertainty from bootstrap scores
        aleatoric_uncertainty = np.full_like(mean_predictions, np.mean(self.bootstrap_scores))
        
        # Calculate confidence intervals (95%)
        lower_ci = np.percentile(predictions, 2.5, axis=0)
        upper_ci = np.percentile(predictions, 97.5, axis=0)
        
        # Calculate prediction intervals (includes both epistemic and aleatoric uncertainty)
        total_uncertainty = epistemic_uncertainty + aleatoric_uncertainty
        std_total = np.sqrt(total_uncertainty)
        lower_pi = mean_predictions - 1.96 * std_total
        upper_pi = mean_predictions + 1.96 * std_total
        
        return UncertaintyResult(
            predictions=mean_predictions,
            uncertainties=epistemic_uncertainty,
            confidence_intervals=(lower_ci, upper_ci),
            prediction_intervals=(lower_pi, upper_pi),
            epistemic_uncertainty=epistemic_uncertainty,
            aleatoric_uncertainty=aleatoric_uncertainty
        )


class ConformalPrediction:
    """
    Conformal Prediction for uncertainty quantification.
    
    This class implements conformal prediction to provide prediction intervals
    with guaranteed coverage probability.
    """
    
    def __init__(self, 
                 base_model,
                 alpha: float = 0.05,
                 method: str = 'quantile'):
        """
        Initialize Conformal Prediction.
        
        Args:
            base_model: Base model for predictions
            alpha: Significance level (1 - coverage probability)
            method: Method for conformal prediction ('quantile' or 'normalized')
        """
        self.base_model = base_model
        self.alpha = alpha
        self.method = method
        self.conformity_scores = None
        self.quantile_threshold = None
        
    def fit(self, X: np.ndarray, y: np.ndarray) -> 'ConformalPrediction':
        """
        Fit conformal prediction model.
        
        Args:
            X: Training features
            y: Training targets
            
        Returns:
            Self for method chaining
        """
        # Fit base model
        self.base_model.fit(X, y)
        
        # Calculate conformity scores on training data
        y_pred = self.base_model.predict(X)
        residuals = np.abs(y - y_pred)
        
        if self.method == 'quantile':
            self.conformity_scores = residuals
            self.quantile_threshold = np.quantile(self.conformity_scores, 1 - self.alpha)
        elif self.method == 'normalized':
            # Normalize residuals by prediction magnitude
            normalized_residuals = residuals / (np.abs(y_pred) + 1e-8)
            self.conformity_scores = normalized_residuals
            self.quantile_threshold = np.quantile(self.conformity_scores, 1 - self.alpha)
        else:
            raise ValueError(f"Unknown method: {self.method}")
            
        return self
        
    def predict_with_uncertainty(self, X: np.ndarray) -> UncertaintyResult:
        """
        Make predictions with conformal prediction intervals.
        
        Args:
            X: Input features
            
        Returns:
            UncertaintyResult containing predictions and conformal intervals
        """
        if self.conformity_scores is None:
            raise ValueError("Must call fit() before predict_with_uncertainty()")
            
        # Get point predictions
        predictions = self.base_model.predict(X)
        
        # Calculate prediction intervals
        if self.method == 'quantile':
            interval_width = self.quantile_threshold
        elif self.method == 'normalized':
            interval_width = self.quantile_threshold * (np.abs(predictions) + 1e-8)
        else:
            raise ValueError(f"Unknown method: {self.method}")
            
        lower_pi = predictions - interval_width
        upper_pi = predictions + interval_width
        
        # For conformal prediction, epistemic uncertainty is the interval width
        epistemic_uncertainty = interval_width
        
        return UncertaintyResult(
            predictions=predictions,
            uncertainties=epistemic_uncertainty,
            confidence_intervals=(lower_pi, upper_pi),
            prediction_intervals=(lower_pi, upper_pi),
            epistemic_uncertainty=epistemic_uncertainty
        )


class EnsembleUncertainty:
    """
    Ensemble-based uncertainty quantification.
    
    This class combines multiple models to estimate prediction uncertainty
    through ensemble variance and disagreement.
    """
    
    def __init__(self, models: List[Any], weights: Optional[List[float]] = None):
        """
        Initialize Ensemble Uncertainty Quantification.
        
        Args:
            models: List of trained models
            weights: Optional weights for model combination
        """
        self.models = models
        self.weights = weights or [1.0 / len(models)] * len(models)
        
        if len(self.weights) != len(self.models):
            raise ValueError("Number of weights must match number of models")
            
    def predict_with_uncertainty(self, X: np.ndarray) -> UncertaintyResult:
        """
        Make predictions with ensemble uncertainty quantification.
        
        Args:
            X: Input features
            
        Returns:
            UncertaintyResult containing predictions and uncertainty estimates
        """
        predictions = []
        
        for model in self.models:
            if hasattr(model, 'predict'):
                pred = model.predict(X)
            elif hasattr(model, 'forward'):
                with torch.no_grad():
                    if isinstance(X, np.ndarray):
                        X_tensor = torch.FloatTensor(X)
                    else:
                        X_tensor = X
                    pred = model(X_tensor).cpu().numpy()
            else:
                raise ValueError(f"Model {type(model)} does not have predict or forward method")
                
            predictions.append(pred)
            
        predictions = np.array(predictions)
        
        # Weighted ensemble predictions
        weighted_predictions = np.average(predictions, axis=0, weights=self.weights)
        
        # Calculate ensemble uncertainty
        # Epistemic uncertainty: variance across models
        epistemic_uncertainty = np.var(predictions, axis=0)
        
        # Total uncertainty: includes both epistemic and aleatoric
        # For ensemble, we estimate aleatoric as the mean prediction error
        mean_abs_error = np.mean(np.abs(predictions - weighted_predictions), axis=0)
        total_uncertainty = epistemic_uncertainty + mean_abs_error
        
        # Calculate confidence intervals (95%)
        lower_ci = np.percentile(predictions, 2.5, axis=0)
        upper_ci = np.percentile(predictions, 97.5, axis=0)
        
        # Calculate prediction intervals
        std_total = np.sqrt(total_uncertainty)
        lower_pi = weighted_predictions - 1.96 * std_total
        upper_pi = weighted_predictions + 1.96 * std_total
        
        return UncertaintyResult(
            predictions=weighted_predictions,
            uncertainties=epistemic_uncertainty,
            confidence_intervals=(lower_ci, upper_ci),
            prediction_intervals=(lower_pi, upper_pi),
            epistemic_uncertainty=epistemic_uncertainty,
            aleatoric_uncertainty=mean_abs_error
        )


class UncertaintyCalibration:
    """
    Uncertainty calibration analysis and reliability diagrams.
    
    This class evaluates the quality of uncertainty estimates through
    calibration analysis and generates reliability diagrams.
    """
    
    def __init__(self, n_bins: int = 10):
        """
        Initialize Uncertainty Calibration.
        
        Args:
            n_bins: Number of bins for calibration analysis
        """
        self.n_bins = n_bins
        
    def calculate_calibration_metrics(self, 
                                    y_true: np.ndarray,
                                    y_pred: np.ndarray,
                                    uncertainties: np.ndarray) -> Dict[str, float]:
        """
        Calculate calibration metrics.
        
        Args:
            y_true: True values
            y_pred: Predicted values
            uncertainties: Uncertainty estimates
            
        Returns:
            Dictionary of calibration metrics
        """
        # Calculate residuals
        residuals = np.abs(y_true - y_pred)
        
        # Normalize residuals by uncertainty
        normalized_residuals = residuals / (uncertainties + 1e-8)
        
        # Calculate calibration metrics
        calibration_error = np.mean(np.abs(normalized_residuals - 1.0))
        
        # Calculate coverage probability
        coverage_prob = np.mean(normalized_residuals <= 1.0)
        
        # Calculate sharpness (average uncertainty)
        sharpness = np.mean(uncertainties)
        
        # Calculate negative log likelihood
        nll = np.mean(0.5 * np.log(2 * np.pi * uncertainties**2) + 
                     0.5 * (residuals**2) / (uncertainties**2))
        
        return {
            'calibration_error': calibration_error,
            'coverage_probability': coverage_prob,
            'sharpness': sharpness,
            'negative_log_likelihood': nll
        }
        
    def plot_reliability_diagram(self, 
                                y_true: np.ndarray,
                                y_pred: np.ndarray,
                                uncertainties: np.ndarray,
                                save_path: Optional[str] = None) -> plt.Figure:
        """
        Plot reliability diagram for uncertainty calibration.
        
        Args:
            y_true: True values
            y_pred: Predicted values
            uncertainties: Uncertainty estimates
            save_path: Optional path to save the plot
            
        Returns:
            Matplotlib figure
        """
        # Calculate residuals and normalized residuals
        residuals = np.abs(y_true - y_pred)
        normalized_residuals = residuals / (uncertainties + 1e-8)
        
        # Create bins based on uncertainty quantiles
        uncertainty_quantiles = np.quantile(uncertainties, np.linspace(0, 1, self.n_bins + 1))
        
        bin_centers = []
        empirical_coverage = []
        predicted_coverage = []
        bin_counts = []
        
        for i in range(self.n_bins):
            # Get samples in this bin
            mask = (uncertainties >= uncertainty_quantiles[i]) & (uncertainties < uncertainty_quantiles[i + 1])
            if i == self.n_bins - 1:  # Include the last quantile
                mask = (uncertainties >= uncertainty_quantiles[i]) & (uncertainties <= uncertainty_quantiles[i + 1])
                
            if np.sum(mask) == 0:
                continue
                
            bin_uncertainties = uncertainties[mask]
            bin_residuals = residuals[mask]
            bin_normalized_residuals = normalized_residuals[mask]
            
            # Calculate coverage
            empirical_cov = np.mean(bin_normalized_residuals <= 1.0)
            predicted_cov = 0.68  # For 1-sigma intervals
            
            bin_centers.append(np.mean(bin_uncertainties))
            empirical_coverage.append(empirical_cov)
            predicted_coverage.append(predicted_cov)
            bin_counts.append(np.sum(mask))
            
        # Create the plot
        fig, (ax1, ax2) = plt.subplots(1, 2, figsize=(12, 5))
        
        # Plot 1: Coverage vs Uncertainty
        ax1.scatter(bin_centers, empirical_coverage, s=bin_counts, alpha=0.7, label='Empirical')
        ax1.axhline(y=0.68, color='r', linestyle='--', label='Perfect Calibration')
        ax1.set_xlabel('Uncertainty')
        ax1.set_ylabel('Coverage Probability')
        ax1.set_title('Reliability Diagram')
        ax1.legend()
        ax1.grid(True, alpha=0.3)
        
        # Plot 2: Residuals vs Uncertainty
        ax2.scatter(uncertainties, residuals, alpha=0.5)
        ax2.plot(uncertainties, uncertainties, 'r--', label='Perfect Calibration')
        ax2.set_xlabel('Predicted Uncertainty')
        ax2.set_ylabel('Actual Residuals')
        ax2.set_title('Residuals vs Uncertainty')
        ax2.legend()
        ax2.grid(True, alpha=0.3)
        
        plt.tight_layout()
        
        if save_path:
            plt.savefig(save_path, dpi=300, bbox_inches='tight')
            
        return fig
        
    def plot_uncertainty_distribution(self, 
                                    uncertainties: np.ndarray,
                                    save_path: Optional[str] = None) -> plt.Figure:
        """
        Plot uncertainty distribution.
        
        Args:
            uncertainties: Uncertainty estimates
            save_path: Optional path to save the plot
            
        Returns:
            Matplotlib figure
        """
        fig, ax = plt.subplots(figsize=(8, 6))
        
        ax.hist(uncertainties, bins=50, alpha=0.7, edgecolor='black')
        ax.axvline(np.mean(uncertainties), color='r', linestyle='--', 
                  label=f'Mean: {np.mean(uncertainties):.4f}')
        ax.axvline(np.median(uncertainties), color='g', linestyle='--', 
                  label=f'Median: {np.median(uncertainties):.4f}')
        
        ax.set_xlabel('Uncertainty')
        ax.set_ylabel('Frequency')
        ax.set_title('Uncertainty Distribution')
        ax.legend()
        ax.grid(True, alpha=0.3)
        
        if save_path:
            plt.savefig(save_path, dpi=300, bbox_inches='tight')
            
        return fig


class UncertaintyQuantifier:
    """
    Main class for uncertainty quantification that combines all methods.
    
    This class provides a unified interface for different uncertainty
    quantification methods and can be used with any model type.
    """
    
    def __init__(self, method: str = 'ensemble', **kwargs):
        """
        Initialize Uncertainty Quantifier.
        
        Args:
            method: Uncertainty quantification method ('mc_dropout', 'bootstrap', 
                   'conformal', 'ensemble')
            **kwargs: Additional arguments for the specific method
        """
        self.method = method
        self.kwargs = kwargs
        self.uncertainty_model = None
        
    def fit(self, model, X: np.ndarray, y: np.ndarray) -> 'UncertaintyQuantifier':
        """
        Fit uncertainty quantification model.
        
        Args:
            model: Trained model or list of models
            X: Training features
            y: Training targets
            
        Returns:
            Self for method chaining
        """
        if self.method == 'mc_dropout':
            if not isinstance(model, nn.Module):
                raise ValueError("MC Dropout requires a PyTorch model")
            self.uncertainty_model = MonteCarloDropout(model, **self.kwargs)
            
        elif self.method == 'bootstrap':
            if not hasattr(model, '__class__'):
                raise ValueError("Bootstrap requires a model class")
            self.uncertainty_model = BootstrapUncertainty(
                model.__class__, 
                **self.kwargs
            ).fit(X, y)
            
        elif self.method == 'conformal':
            self.uncertainty_model = ConformalPrediction(
                model, 
                **self.kwargs
            ).fit(X, y)
            
        elif self.method == 'ensemble':
            if not isinstance(model, list):
                raise ValueError("Ensemble method requires a list of models")
            self.uncertainty_model = EnsembleUncertainty(model, **self.kwargs)
            
        else:
            raise ValueError(f"Unknown method: {self.method}")
            
        return self
        
    def predict_with_uncertainty(self, X: np.ndarray) -> UncertaintyResult:
        """
        Make predictions with uncertainty quantification.
        
        Args:
            X: Input features
            
        Returns:
            UncertaintyResult containing predictions and uncertainty estimates
        """
        if self.uncertainty_model is None:
            raise ValueError("Must call fit() before predict_with_uncertainty()")
            
        return self.uncertainty_model.predict_with_uncertainty(X)
        
    def evaluate_calibration(self, 
                           y_true: np.ndarray,
                           y_pred: np.ndarray,
                           uncertainties: np.ndarray) -> Dict[str, float]:
        """
        Evaluate uncertainty calibration.
        
        Args:
            y_true: True values
            y_pred: Predicted values
            uncertainties: Uncertainty estimates
            
        Returns:
            Dictionary of calibration metrics
        """
        calibrator = UncertaintyCalibration()
        return calibrator.calculate_calibration_metrics(y_true, y_pred, uncertainties)
        
    def plot_calibration(self, 
                        y_true: np.ndarray,
                        y_pred: np.ndarray,
                        uncertainties: np.ndarray,
                        save_dir: Optional[str] = None) -> List[plt.Figure]:
        """
        Plot calibration analysis.
        
        Args:
            y_true: True values
            y_pred: Predicted values
            uncertainties: Uncertainty estimates
            save_dir: Optional directory to save plots
            
        Returns:
            List of matplotlib figures
        """
        calibrator = UncertaintyCalibration()
        
        figures = []
        
        # Reliability diagram
        fig1 = calibrator.plot_reliability_diagram(y_true, y_pred, uncertainties)
        figures.append(fig1)
        
        if save_dir:
            fig1.savefig(f"{save_dir}/reliability_diagram.png", dpi=300, bbox_inches='tight')
            
        # Uncertainty distribution
        fig2 = calibrator.plot_uncertainty_distribution(uncertainties)
        figures.append(fig2)
        
        if save_dir:
            fig2.savefig(f"{save_dir}/uncertainty_distribution.png", dpi=300, bbox_inches='tight')
            
        return figures
```
