## Supplementary material for "Machine Learning-Driven Prediction of TLR4 Binding Affinity: A Comprehensive Molecular Feature Analysis for Drug Discovery": Codebase: validation_benchmarking.py

```
"""
Validation and benchmarking utilities for Task 20.

Implements:
- Leave-one-out cross-validation (LOO-CV)
- Temporal validation (time-based train/test split)
- External validation against a held-out dataset
- Simple benchmarking across multiple models
- Bootstrap-based power analysis and sample size recommendation
"""

from __future__ import annotations

from dataclasses import dataclass
from typing import Dict, List, Optional, Tuple, Any

import logging
import numpy as np
import pandas as pd

from sklearn.model_selection import LeaveOneOut, cross_val_score
from sklearn.metrics import (
    mean_squared_error,
    mean_absolute_error,
    r2_score,
)
from sklearn.base import BaseEstimator, clone

from .evaluator import ModelEvaluator, PerformanceMetrics


logger = logging.getLogger(__name__)


def _calc_metrics(y_true: np.ndarray, y_pred: np.ndarray) -> Dict[str, float]:
    return {
        "r2": r2_score(y_true, y_pred),
        "rmse": float(np.sqrt(mean_squared_error(y_true, y_pred))),
        "mae": float(mean_absolute_error(y_true, y_pred)),
    }


def evaluate_leave_one_out(model: BaseEstimator, X: pd.DataFrame, y: pd.Series,
                           scoring: Optional[List[str]] = None) -> Dict[str, Any]:
    """
    Perform leave-one-out cross-validation for small datasets.
    Returns per-metric score arrays and summary stats.
    """
    if scoring is None:
        scoring = ["neg_mean_squared_error", "neg_mean_absolute_error", "r2"]

    logger.info("Running Leave-One-Out cross-validation")
    loo = LeaveOneOut()
    results: Dict[str, Any] = {}
    for metric in scoring:
        scores = cross_val_score(model, X, y, cv=loo, scoring=metric, n_jobs=-1)
        results[metric] = {
            "scores": scores,
            "mean": float(np.mean(scores)),
            "std": float(np.std(scores)),
            "min": float(np.min(scores)),
            "max": float(np.max(scores)),
        }

    if "neg_mean_squared_error" in results:
        rmse_scores = np.sqrt(-results["neg_mean_squared_error"]["scores"])  # type: ignore[index]
        results["rmse"] = {
            "scores": rmse_scores,
            "mean": float(np.mean(rmse_scores)),
            "std": float(np.std(rmse_scores)),
            "min": float(np.min(rmse_scores)),
            "max": float(np.max(rmse_scores)),
        }
    if "neg_mean_absolute_error" in results:
        mae_scores = -results["neg_mean_absolute_error"]["scores"]  # type: ignore[index]
        results["mae"] = {
            "scores": mae_scores,
            "mean": float(np.mean(mae_scores)),
            "std": float(np.std(mae_scores)),
            "min": float(np.min(mae_scores)),
            "max": float(np.max(mae_scores)),
        }
    return results


def evaluate_temporal_split(model: BaseEstimator,
                            X: pd.DataFrame,
                            y: pd.Series,
                            timestamps: pd.Series,
                            split_time: Optional[pd.Timestamp] = None) -> Dict[str, Any]:
    """
    Temporal validation: train on earlier timestamps, test on later timestamps.
    If split_time not provided, uses median timestamp.
    """
    if timestamps is None or len(timestamps) != len(X):
        raise ValueError("timestamps Series must align with X")

    ts = pd.to_datetime(timestamps)
    cutoff = split_time or ts.median()
    train_idx = ts <= cutoff
    test_idx = ts > cutoff
    if not test_idx.any() or not train_idx.any():
        raise ValueError("Temporal split produced empty train or test set")

    model_fitted = clone(model)
    model_fitted.fit(X.loc[train_idx], y.loc[train_idx])
    y_pred = model_fitted.predict(X.loc[test_idx])
    metrics = _calc_metrics(y.loc[test_idx].to_numpy(), np.asarray(y_pred))
    return {
        "cutoff": pd.Timestamp(cutoff),
        "n_train": int(train_idx.sum()),
        "n_test": int(test_idx.sum()),
        "metrics": metrics,
    }


def evaluate_external(model: BaseEstimator,
                      X_train: pd.DataFrame,
                      y_train: pd.Series,
                      X_external: pd.DataFrame,
                      y_external: pd.Series) -> Dict[str, Any]:
    """
    Train on internal data, evaluate on external dataset.
    """
    if X_external.shape[1] != X_train.shape[1]:
        raise ValueError("Feature dimension mismatch between train and external")

    model_fitted = clone(model)
    model_fitted.fit(X_train, y_train)
    y_pred = model_fitted.predict(X_external)
    metrics = _calc_metrics(y_external.to_numpy(), np.asarray(y_pred))
    return {
        "n_train": int(len(X_train)),
        "n_external": int(len(X_external)),
        "metrics": metrics,
    }


def benchmark_models(models: Dict[str, BaseEstimator],
                     X: pd.DataFrame,
                     y: pd.Series,
                     cv_folds: int = 5) -> pd.DataFrame:
    """
    Simple benchmark: cross-validated R2/RMSE/MAE for each model.
    Returns a DataFrame sorted by R2 desc.
    """
    evaluator = ModelEvaluator()
    rows: List[Dict[str, Any]] = []
    for name, model in models.items():
        logger.info("Benchmarking model: %s", name)
        cv = evaluate_leave_one_out(model, X, y) if cv_folds == -1 else evaluator.cross_validate_model(
            model, X, y, model_name=name, cv=cv_folds
        )
        r2_mean = float(cv["r2"]["mean"]) if "r2" in cv else float("nan")
        rmse_mean = float(cv.get("rmse", {}).get("mean", float("nan")))
        mae_mean = float(cv.get("mae", {}).get("mean", float("nan")))
        rows.append({
            "model": name,
            "r2_mean": r2_mean,
            "rmse_mean": rmse_mean,
            "mae_mean": mae_mean,
        })
    df = pd.DataFrame(rows)
    if not df.empty:
        df = df.sort_values("r2_mean", ascending=False).reset_index(drop=True)
    return df


@dataclass
class PowerAnalysisResult:
    target_effect_rmse: float
    alpha: float
    power: float
    required_n: int


def bootstrap_power_for_rmse_diff(y_true: np.ndarray,
                                  residuals_a: np.ndarray,
                                  residuals_b: np.ndarray,
                                  alpha: float = 0.05,
                                  n_boot: int = 2000,
                                  sample_size: Optional[int] = None) -> float:
    """
    Estimate power to detect RMSE difference between two models using bootstrap
    on residuals with paired t-test.
    If sample_size is provided, subsample with replacement to that size.
    Returns estimated power in [0,1].
    """
    rng = np.random.default_rng(42)
    n = len(y_true)
    if sample_size is None:
        sample_size = n
    indices = np.arange(n)
    rejections = 0
    for _ in range(n_boot):
        idx = rng.choice(indices, size=sample_size, replace=True)
        diff = (residuals_a[idx] ** 2) - (residuals_b[idx] ** 2)
        # Paired t-test on squared residuals (proxy for MSE)
        d_mean = diff.mean()
        d_std = diff.std(ddof=1) if sample_size > 1 else 0.0
        if d_std == 0.0:
            continue
        t_stat = d_mean / (d_std / np.sqrt(sample_size))
        # Two-sided critical t approx with normal (large-sample)
        from scipy.stats import norm
        crit = norm.ppf(1 - alpha / 2)
        if abs(t_stat) > crit:
            rejections += 1
    return rejections / float(n_boot)


def recommend_sample_size_for_rmse_effect(model_a: BaseEstimator,
                                          model_b: BaseEstimator,
                                          X: pd.DataFrame,
                                          y: pd.Series,
                                          target_rmse_improvement: float,
                                          alpha: float = 0.05,
                                          power_target: float = 0.8,
                                          max_n: Optional[int] = None) -> PowerAnalysisResult:
    """
    Recommend sample size to detect a target RMSE improvement between two models.
    Uses bootstrap power estimation over increasing sample sizes.
    """
    max_n = max_n or len(X)

    # Fit both models on full data to obtain residuals distribution proxy
    a = clone(model_a).fit(X, y)
    b = clone(model_b).fit(X, y)
    ra = y.to_numpy() - np.asarray(a.predict(X))
    rb = y.to_numpy() - np.asarray(b.predict(X))

    # Scan sample sizes
    for n in range(20, max_n + 1, max(5, max_n // 20)):
        power = bootstrap_power_for_rmse_diff(y.to_numpy(), ra, rb, alpha=alpha, sample_size=n)
        if power >= power_target:
            return PowerAnalysisResult(
                target_effect_rmse=float(target_rmse_improvement),
                alpha=alpha,
                power=float(power),
                required_n=n,
            )

    # Fallback: return max_n with achieved power
    final_power = bootstrap_power_for_rmse_diff(y.to_numpy(), ra, rb, alpha=alpha, sample_size=max_n)
    return PowerAnalysisResult(
        target_effect_rmse=float(target_rmse_improvement),
        alpha=alpha,
        power=float(final_power),
        required_n=max_n,
    )
```
