## Supplementary material for "Machine Learning-Driven Prediction of TLR4 Binding Affinity: A Comprehensive Molecular Feature Analysis for Drug Discovery": Codebase: __init__.py

```
"""
Molecular Analysis Module

Contains classes for molecular feature extraction, structure analysis,
and descriptor calculation from PDBQT files.
"""

from .features import MolecularFeatures, BindingData, PredictionResult
from .descriptors import MolecularDescriptorCalculator
from .structure import StructuralFeatureExtractor
from .parser import PDBQTParser
from .extractor import MolecularFeatureExtractor

__all__ = [
    "MolecularFeatures",
    "BindingData",
    "PredictionResult", 
    "MolecularDescriptorCalculator",
    "StructuralFeatureExtractor",
    "PDBQTParser",
    "MolecularFeatureExtractor"
]
```
