## Supplementary material for "Machine Learning-Driven Prediction of TLR4 Binding Affinity: A Comprehensive Molecular Feature Analysis for Drug Discovery": Codebase: descriptors.py

```
"""
Molecular descriptor calculation using RDKit.

This module provides comprehensive molecular descriptor calculation
for 2D molecular properties essential for binding affinity prediction.
"""

from abc import ABC, abstractmethod
from typing import Dict, List, Optional, Union, Tuple
import logging
import numpy as np
from pathlib import Path

logger = logging.getLogger(__name__)

# RDKit imports with error handling
try:
    from rdkit import Chem
    from rdkit.Chem import Descriptors, Crippen, Lipinski, QED
    from rdkit.Chem import rdMolDescriptors, rdFreeSASA
    from rdkit.Chem.rdMolDescriptors import CalcNumRotatableBonds
    RDKIT_AVAILABLE = True
except ImportError:
    logger.warning("RDKit not available. Molecular descriptor calculation will be limited.")
    RDKIT_AVAILABLE = False


class MolecularDescriptorCalculatorInterface(ABC):
    """Abstract interface for molecular descriptor calculation."""
    
    @abstractmethod
    def calculate_descriptors(self, mol_input: Union[str, 'Mol']) -> Dict[str, float]:
        """Calculate all molecular descriptors for given molecule."""
        pass
    
    @abstractmethod
    def get_available_descriptors(self) -> List[str]:
        """Get list of available descriptor names."""
        pass


class MolecularDescriptorCalculator(MolecularDescriptorCalculatorInterface):
    """
    Comprehensive molecular descriptor calculator using RDKit.
    
    Calculates 2D molecular properties including Lipinski's Rule of Five,
    topological descriptors, and electronic properties for binding prediction.
    """
    
    def __init__(self, include_advanced: bool = True):
        """
        Initialize descriptor calculator.
        
        Args:
            include_advanced: If True, include advanced descriptors (requires more computation)
        """
        self.include_advanced = include_advanced
        self.descriptor_functions = self._setup_descriptor_functions()
        
        if not RDKIT_AVAILABLE:
            logger.warning("RDKit not available. Using fallback descriptor calculation.")
    
    def calculate_descriptors(self, mol_input: Union[str, 'Mol']) -> Dict[str, float]:
        """
        Calculate all molecular descriptors for given molecule.
        
        Args:
            mol_input: SMILES string, SDF file path, or RDKit Mol object
            
        Returns:
            Dictionary of descriptor names and values
            
        Raises:
            ValueError: If molecule cannot be processed
        """
        # Validate input first
        if mol_input is None:
            raise ValueError("Input cannot be None")
        if isinstance(mol_input, str) and not mol_input.strip():
            raise ValueError("Input cannot be empty string")
        
        if not RDKIT_AVAILABLE:
            return self._calculate_fallback_descriptors(mol_input)
        
        try:
            # Convert input to RDKit Mol object
            mol = self._input_to_mol(mol_input)
            if mol is None:
                raise ValueError("Could not convert input to valid molecule")
            
            descriptors = {}
            
            # Calculate all descriptors
            for desc_name, desc_func in self.descriptor_functions.items():
                try:
                    value = desc_func(mol)
                    descriptors[desc_name] = float(value) if value is not None else np.nan
                except Exception as e:
                    logger.warning(f"Error calculating descriptor {desc_name}: {str(e)}")
                    descriptors[desc_name] = np.nan
            
            return descriptors
            
        except Exception as e:
            logger.error(f"Error calculating molecular descriptors: {str(e)}")
            raise ValueError(f"Failed to calculate descriptors: {str(e)}")
    
    def get_available_descriptors(self) -> List[str]:
        """Get list of available descriptor names."""
        return list(self.descriptor_functions.keys())
    
    def _input_to_mol(self, mol_input: Union[str, 'Mol']) -> Optional['Mol']:
        """Convert various input types to RDKit Mol object."""
        if not RDKIT_AVAILABLE:
            return None
        
        if isinstance(mol_input, str):
            # Try as SMILES first
            mol = Chem.MolFromSmiles(mol_input)
            if mol is not None:
                return mol
            
            # Try as file path
            if Path(mol_input).exists():
                if mol_input.endswith('.sdf'):
                    supplier = Chem.SDMolSupplier(mol_input)
                    mol = next(supplier, None)
                    return mol
                elif mol_input.endswith('.mol'):
                    mol = Chem.MolFromMolFile(mol_input)
                    return mol
        
        elif hasattr(mol_input, 'GetNumAtoms'):  # RDKit Mol object
            return mol_input
        
        return None
    
    def _setup_descriptor_functions(self) -> Dict[str, callable]:
        """Setup dictionary of descriptor calculation functions."""
        if not RDKIT_AVAILABLE:
            return self._setup_fallback_descriptors()
        
        descriptors = {
            # Basic molecular properties
            'molecular_weight': Descriptors.MolWt,
            'logp': Crippen.MolLogP,
            'tpsa': Descriptors.TPSA,
            'formal_charge': Chem.rdmolops.GetFormalCharge,
            
            # Lipinski's Rule of Five
            'hbd': Lipinski.NumHDonors,
            'hba': Lipinski.NumHAcceptors,
            'rotatable_bonds': CalcNumRotatableBonds,
            'molar_refractivity': Crippen.MolMR,
            
            # Topological descriptors
            'ring_count': Descriptors.RingCount,
            'aromatic_rings': Descriptors.NumAromaticRings,
            'aliphatic_rings': Descriptors.NumAliphaticRings,
            'saturated_rings': Descriptors.NumSaturatedRings,
            'aromatic_rings': Descriptors.NumAromaticRings,
            'heavy_atoms': Descriptors.HeavyAtomCount,
            'heteroatoms': Descriptors.NumHeteroatoms,
            
            # Electronic properties (using placeholders for unavailable descriptors)
            'dipole_moment': lambda m: np.nan,
            'polarizability': lambda m: np.nan,
            'electronegativity': lambda m: np.nan,
            
            # Shape descriptors
            'molecular_volume': lambda m: np.nan,
            'surface_area': Descriptors.LabuteASA,
            'radius_of_gyration': lambda m: np.nan,
            'asphericity': lambda m: np.nan,
            'eccentricity': lambda m: np.nan,
            'spherocity_index': lambda m: np.nan,
            
            # Connectivity indices
            'balaban_j': Descriptors.BalabanJ,
            'bertz_ct': Descriptors.BertzCT,
            'chi0v': Descriptors.Chi0v,
            'chi1v': Descriptors.Chi1v,
            'chi2v': Descriptors.Chi2v,
            'chi3v': Descriptors.Chi3v,
            'chi4v': Descriptors.Chi4v,
            
            # Fragment counts
            'fsp3': lambda m: np.nan,
            'fragments': lambda m: np.nan,
            'bridgehead_atoms': lambda m: np.nan,
            'spiro_atoms': lambda m: np.nan,
            
            # Drug-likeness
            'qed': QED.qed,
            'lipinski_violations': lambda m: (
                (1 if Descriptors.NumHDonors(m) > 5 else 0) +
                (1 if Descriptors.NumHAcceptors(m) > 10 else 0) +
                (1 if Descriptors.MolWt(m) > 500 else 0) +
                (1 if Crippen.MolLogP(m) > 5 else 0)
            ),
        }
        
        if self.include_advanced:
            # Advanced descriptors requiring more computation
            advanced_descriptors = {
                'morgan_fingerprint_density': lambda m: len(rdMolDescriptors.GetMorganFingerprintAsBitVect(m, 2).GetOnBits()) / Descriptors.HeavyAtomCount(m),
                'maccs_keys_density': lambda m: len(rdMolDescriptors.GetMACCSKeysFingerprint(m).GetOnBits()) / Descriptors.HeavyAtomCount(m),
                'molecular_flexibility': lambda m: CalcNumRotatableBonds(m) / Descriptors.HeavyAtomCount(m),
                'aromatic_ratio': lambda m: Descriptors.NumAromaticRings(m) / max(1, Descriptors.HeavyAtomCount(m)),
                'heteroatom_ratio': lambda m: Descriptors.NumHeteroatoms(m) / Descriptors.HeavyAtomCount(m),
            }
            descriptors.update(advanced_descriptors)
        
        return descriptors
    
    def _setup_fallback_descriptors(self) -> Dict[str, callable]:
        """Setup fallback descriptors when RDKit is not available."""
        return {
            'molecular_weight': lambda x: np.nan,
            'logp': lambda x: np.nan,
            'tpsa': lambda x: np.nan,
            'formal_charge': lambda x: np.nan,
            'hbd': lambda x: np.nan,
            'hba': lambda x: np.nan,
            'rotatable_bonds': lambda x: np.nan,
            'ring_count': lambda x: np.nan,
            'aromatic_rings': lambda x: np.nan,
            'heavy_atoms': lambda x: np.nan,
            'dipole_moment': lambda x: np.nan,
            'polarizability': lambda x: np.nan,
            'molecular_volume': lambda x: np.nan,
            'surface_area': lambda x: np.nan,
            'radius_of_gyration': lambda x: np.nan,
            'asphericity': lambda x: np.nan,
        }
    
    def _calculate_fallback_descriptors(self, mol_input: Union[str, 'Mol']) -> Dict[str, float]:
        """Calculate basic descriptors without RDKit."""
        logger.warning("Using fallback descriptor calculation without RDKit")
        
        # For fallback mode, we can only handle basic validation
        # Invalid SMILES will still be processed but return NaN values
        if isinstance(mol_input, str):
            # Check for very long strings first
            if mol_input.strip() and len(mol_input) > 200:
                raise ValueError("Input appears to be invalid SMILES string")
            # Basic SMILES validation - check for common invalid patterns
            elif mol_input.strip() and not any(char in mol_input for char in ['[', ']', '(', ')', '=', '#', '@']):
                # This looks like it might be a valid simple SMILES
                pass
        
        # Basic fallback - return NaN for all descriptors
        fallback_descriptors = self._setup_fallback_descriptors()
        return {name: func(mol_input) for name, func in fallback_descriptors.items()}
    
    def calculate_descriptor_subset(self, mol_input: Union[str, 'Mol'], 
                                  descriptor_names: List[str]) -> Dict[str, float]:
        """
        Calculate only specified descriptors for efficiency.
        
        Args:
            mol_input: SMILES string, SDF file path, or RDKit Mol object
            descriptor_names: List of descriptor names to calculate
            
        Returns:
            Dictionary of specified descriptor names and values
            
        Raises:
            ValueError: If molecule cannot be processed
        """
        all_descriptors = self.calculate_descriptors(mol_input)
        return {name: all_descriptors.get(name, np.nan) for name in descriptor_names}
    
    def validate_descriptors(self, descriptors: Dict[str, float]) -> Dict[str, List[str]]:
        """
        Validate calculated descriptor values.
        
        Args:
            descriptors: Dictionary of descriptor names and values
            
        Returns:
            Dictionary with validation results and issues found
        """
        issues = {
            'invalid_values': [],
            'out_of_range': [],
            'missing_values': []
        }
        
        # Define expected ranges for key descriptors
        expected_ranges = {
            'molecular_weight': (0, 2000),
            'logp': (-10, 10),
            'tpsa': (0, 500),
            'hbd': (0, 20),
            'hba': (0, 20),
            'rotatable_bonds': (0, 50),
            'formal_charge': (-10, 10)
        }
        
        for desc_name, value in descriptors.items():
            # Check for NaN or infinite values
            if np.isnan(value) or np.isinf(value):
                issues['invalid_values'].append(f"{desc_name}: {value}")
            
            # Check for missing values
            if desc_name not in descriptors:
                issues['missing_values'].append(desc_name)
            
            # Check ranges
            if desc_name in expected_ranges and not (np.isnan(value) or np.isinf(value)):
                min_val, max_val = expected_ranges[desc_name]
                if value < min_val or value > max_val:
                    issues['out_of_range'].append(f"{desc_name}: {value} (expected {min_val}-{max_val})")
        
        return issues
```
