## Supplementary material for "Machine Learning-Driven Prediction of TLR4 Binding Affinity: A Comprehensive Molecular Feature Analysis for Drug Discovery": Codebase: extractor.py

```
"""
Main molecular feature extractor.

This module provides the main MolecularFeatureExtractor class that
orchestrates the extraction of both 2D and 3D molecular features
from PDBQT files.
"""

from typing import Dict, List, Optional, Union, Any
import pandas as pd
import numpy as np
import logging
from pathlib import Path
from tqdm import tqdm

from .parser import PDBQTParser
from .descriptors import MolecularDescriptorCalculator
from .structure import StructuralFeatureExtractor
from .features import MolecularFeatures, FeatureSet
from ..utils.error_handling import (
    RobustnessManager, CheckpointManager, robust_execution,
    safe_execution, graceful_degradation, FeatureExtractionError,
    DataQualityError, PipelineError
)

logger = logging.getLogger(__name__)


class MolecularFeatureExtractor:
    """
    Main molecular feature extractor for TLR4 binding prediction.
    
    Orchestrates the extraction of comprehensive molecular features
    from PDBQT files using RDKit and PyMOL.
    """
    
    def __init__(self, 
                 include_2d_features: bool = True,
                 include_3d_features: bool = True,
                 include_advanced_features: bool = True,
                 enable_checkpointing: bool = True,
                 checkpoint_interval: int = 100,
                 robustness_config: Optional[Dict[str, Any]] = None):
        """
        Initialize molecular feature extractor with robust error handling.
        
        Args:
            include_2d_features: Whether to extract 2D molecular descriptors
            include_3d_features: Whether to extract 3D structural features
            include_advanced_features: Whether to include advanced descriptors
            enable_checkpointing: Whether to enable checkpointing for long operations
            checkpoint_interval: Interval for saving checkpoints during batch processing
            robustness_config: Configuration for robustness features
        """
        self.include_2d_features = include_2d_features
        self.include_3d_features = include_3d_features
        self.include_advanced_features = include_advanced_features
        self.enable_checkpointing = enable_checkpointing
        self.checkpoint_interval = checkpoint_interval
        
        # Initialize robustness manager
        self.robustness_manager = RobustnessManager(robustness_config)
        self.checkpoint_manager = CheckpointManager()
        
        # Initialize components with error handling
        try:
            self.parser = PDBQTParser()
            logger.info("PDBQT parser initialized successfully")
        except Exception as e:
            logger.error(f"Failed to initialize PDBQT parser: {e}")
            raise FeatureExtractionError(f"PDBQT parser initialization failed: {e}")
        
        try:
            self.descriptor_calculator = MolecularDescriptorCalculator(
                include_advanced=include_advanced_features
            )
            logger.info("Molecular descriptor calculator initialized successfully")
        except Exception as e:
            logger.error(f"Failed to initialize descriptor calculator: {e}")
            raise FeatureExtractionError(f"Descriptor calculator initialization failed: {e}")
        
        try:
            self.structural_extractor = StructuralFeatureExtractor()
            logger.info("Structural feature extractor initialized successfully")
        except Exception as e:
            logger.error(f"Failed to initialize structural extractor: {e}")
            raise FeatureExtractionError(f"Structural extractor initialization failed: {e}")
        
        # Statistics and performance tracking
        self.extraction_stats = {
            'total_files_processed': 0,
            'successful_extractions': 0,
            'failed_extractions': 0,
            'extraction_errors': [],
            'processing_times': [],
            'feature_extraction_times': {
                '2d_features': [],
                '3d_features': [],
                'parsing': []
            },
            'memory_usage': [],
            'start_time': None,
            'end_time': None,
            'checkpoint_saves': 0,
            'checkpoint_loads': 0,
            'recovery_attempts': 0
        }
    
    @robust_execution(max_retries=3, delay=1.0)
    def extract_features(self, pdbqt_path: str) -> MolecularFeatures:
        """
        Extract all molecular features from a single PDBQT file with robust error handling.
        
        Args:
            pdbqt_path: Path to PDBQT file
            
        Returns:
            MolecularFeatures object with all extracted features
            
        Raises:
            FeatureExtractionError: If file cannot be processed
        """
        import time
        import psutil
        
        start_time = time.time()
        logger.info(f"Extracting features from {pdbqt_path}")
        
        # Validate input file
        pdbqt_file = Path(pdbqt_path)
        if not pdbqt_file.exists():
            raise FeatureExtractionError(f"PDBQT file does not exist: {pdbqt_path}")
        
        if not pdbqt_file.suffix.lower() == '.pdbqt':
            raise FeatureExtractionError(f"Invalid file extension. Expected .pdbqt, got: {pdbqt_file.suffix}")
        
        try:
            # Track memory usage
            process = psutil.Process()
            memory_before = process.memory_info().rss / 1024 / 1024  # MB
            
            # Parse PDBQT file with timing and error handling
            parse_start = time.time()
            try:
                parsed_data = self.parser.parse_file(pdbqt_path)
                parse_time = time.time() - parse_start
                self.extraction_stats['feature_extraction_times']['parsing'].append(parse_time)
            except Exception as e:
                error_msg = f"Failed to parse PDBQT file {pdbqt_path}: {str(e)}"
                logger.error(error_msg)
                self.robustness_manager.log_error(e, {'file': pdbqt_path, 'stage': 'parsing'})
                raise FeatureExtractionError(error_msg, compound_name=pdbqt_file.stem)
            
            compound_name = Path(pdbqt_path).stem
            
            # Initialize feature dictionary
            features_dict = {
                'compound_name': compound_name,
                'pdbqt_file': pdbqt_path
            }
            
            # Extract 2D molecular descriptors with timing and graceful degradation
            if self.include_2d_features:
                desc_2d_start = time.time()
                try:
                    # Convert PDBQT to SMILES for RDKit processing
                    smiles = self._extract_smiles_from_pdbqt(parsed_data)
                    if smiles:
                        features_dict['smiles'] = smiles
                        descriptors_2d = self.descriptor_calculator.calculate_descriptors(smiles)
                        features_dict.update(descriptors_2d)
                    else:
                        logger.warning(f"Could not extract SMILES from {pdbqt_path}, using coordinate-based features")
                        # Use coordinate-based features as fallback
                        coord_features = self._extract_coordinate_features(pdbqt_path)
                        features_dict.update(coord_features)
                        # Fill remaining 2D features with NaN
                        default_2d = self._get_default_2d_features()
                        for key, value in default_2d.items():
                            if key not in features_dict:
                                features_dict[key] = value
                    
                except Exception as e:
                    error_msg = f"Error extracting 2D features from {pdbqt_path}: {str(e)}"
                    logger.warning(error_msg)
                    self.robustness_manager.log_error(e, {'file': pdbqt_path, 'stage': '2d_features'})
                    # Use graceful degradation - fall back to coordinate features
                    coord_features = self._extract_coordinate_features(pdbqt_path)
                    features_dict.update(coord_features)
                    features_dict.update(self._get_default_2d_features())
                
                desc_2d_time = time.time() - desc_2d_start
                self.extraction_stats['feature_extraction_times']['2d_features'].append(desc_2d_time)
            else:
                features_dict.update(self._get_default_2d_features())
            
            # Extract 3D structural features with timing and graceful degradation
            if self.include_3d_features:
                desc_3d_start = time.time()
                try:
                    structural_features = self.structural_extractor.extract_features(pdbqt_path)
                    features_dict.update(structural_features)
                    
                except Exception as e:
                    error_msg = f"Error extracting 3D features from {pdbqt_path}: {str(e)}"
                    logger.warning(error_msg)
                    self.robustness_manager.log_error(e, {'file': pdbqt_path, 'stage': '3d_features'})
                    # Use graceful degradation - fall back to default features
                    features_dict.update(self._get_default_3d_features())
                
                desc_3d_time = time.time() - desc_3d_start
                self.extraction_stats['feature_extraction_times']['3d_features'].append(desc_3d_time)
            else:
                features_dict.update(self._get_default_3d_features())
            
            # Create MolecularFeatures object
            molecular_features = MolecularFeatures.from_dict(features_dict)
            
            # Track total processing time and memory usage
            total_time = time.time() - start_time
            memory_after = process.memory_info().rss / 1024 / 1024  # MB
            memory_used = memory_after - memory_before
            
            self.extraction_stats['processing_times'].append(total_time)
            self.extraction_stats['memory_usage'].append(memory_used)
            self.extraction_stats['successful_extractions'] += 1
            
            logger.info(f"Successfully extracted features for {compound_name} "
                       f"(Time: {total_time:.2f}s, Memory: {memory_used:.1f}MB)")
            return molecular_features
            
        except Exception as e:
            error_msg = f"Failed to extract features from {pdbqt_path}: {str(e)}"
            logger.error(error_msg)
            self.extraction_stats['failed_extractions'] += 1
            self.extraction_stats['extraction_errors'].append(error_msg)
            raise ValueError(error_msg)
    
    def batch_extract(self, pdbqt_directory: str, 
                     resume_from_checkpoint: bool = True) -> pd.DataFrame:
        """
        Extract features from all PDBQT files in a directory with checkpointing support.
        
        Args:
            pdbqt_directory: Directory containing PDBQT files
            resume_from_checkpoint: Whether to resume from existing checkpoint
            
        Returns:
            DataFrame with features for all compounds
        """
        import time
        
        batch_start_time = time.time()
        self.extraction_stats['start_time'] = batch_start_time
        
        logger.info(f"Starting batch extraction from {pdbqt_directory}")
        
        pdbqt_dir = Path(pdbqt_directory)
        if not pdbqt_dir.exists():
            raise ValueError(f"Directory does not exist: {pdbqt_directory}")
        
        # Find all PDBQT files
        pdbqt_files = list(pdbqt_dir.glob("*.pdbqt"))
        if not pdbqt_files:
            raise ValueError(f"No PDBQT files found in {pdbqt_directory}")
        
        logger.info(f"Found {len(pdbqt_files)} PDBQT files")
        
        # Generate checkpoint ID based on directory
        checkpoint_id = f"batch_extract_{pdbqt_dir.name}_{int(batch_start_time)}"
        
        # Try to resume from checkpoint
        all_features = []
        failed_files = []
        start_index = 0
        
        if resume_from_checkpoint and self.enable_checkpointing:
            checkpoint_data = self.checkpoint_manager.load_checkpoint(checkpoint_id)
            if checkpoint_data:
                logger.info("Resuming from checkpoint")
                all_features = checkpoint_data.get('all_features', [])
                failed_files = checkpoint_data.get('failed_files', [])
                start_index = checkpoint_data.get('start_index', 0)
                self.extraction_stats['checkpoint_loads'] += 1
        
        # Reset statistics for batch processing (only if not resuming)
        if start_index == 0:
            self.extraction_stats['total_files_processed'] = 0
            self.extraction_stats['successful_extractions'] = 0
            self.extraction_stats['failed_extractions'] = 0
            self.extraction_stats['extraction_errors'] = []
            self.extraction_stats['processing_times'] = []
            self.extraction_stats['memory_usage'] = []
        
        # Extract features from all files with progress tracking
        progress_bar = tqdm(pdbqt_files[start_index:], 
                           desc="Extracting features", 
                           unit="files", ncols=100,
                           initial=start_index,
                           total=len(pdbqt_files))
        
        for i, pdbqt_file in enumerate(progress_bar):
            current_index = start_index + i
            
            try:
                features = self.extract_features(str(pdbqt_file))
                all_features.append(features.to_dict())
                self.extraction_stats['total_files_processed'] += 1
                
                # Update progress bar with statistics
                success_rate = (len(all_features) / (current_index + 1)) * 100
                avg_time = np.mean(self.extraction_stats['processing_times']) if self.extraction_stats['processing_times'] else 0
                progress_bar.set_postfix({
                    'Success': f"{success_rate:.1f}%",
                    'Avg Time': f"{avg_time:.2f}s",
                    'Processed': len(all_features)
                })
                
                # Save checkpoint periodically
                if (self.enable_checkpointing and 
                    current_index > 0 and 
                    current_index % self.checkpoint_interval == 0):
                    
                    checkpoint_data = {
                        'all_features': all_features,
                        'failed_files': failed_files,
                        'start_index': current_index + 1,
                        'extraction_stats': self.extraction_stats,
                        'timestamp': time.time()
                    }
                    
                    if self.checkpoint_manager.save_checkpoint(checkpoint_id, checkpoint_data):
                        self.extraction_stats['checkpoint_saves'] += 1
                        logger.info(f"Checkpoint saved at file {current_index + 1}")
                
            except Exception as e:
                error_msg = f"Failed to process {pdbqt_file}: {str(e)}"
                logger.error(error_msg)
                self.robustness_manager.log_error(e, {
                    'file': str(pdbqt_file),
                    'batch_index': current_index,
                    'stage': 'batch_extraction'
                })
                failed_files.append(str(pdbqt_file))
                self.extraction_stats['extraction_errors'].append(error_msg)
                continue
        
        progress_bar.close()
        
        batch_end_time = time.time()
        self.extraction_stats['end_time'] = batch_end_time
        total_batch_time = batch_end_time - batch_start_time
        
        if not all_features:
            raise ValueError("No features could be extracted from any files")
        
        # Create DataFrame
        features_df = pd.DataFrame(all_features)
        
        # Log comprehensive statistics
        successful = len(all_features)
        failed = len(failed_files)
        total = len(pdbqt_files)
        
        logger.info(f"Batch extraction completed:")
        logger.info(f"  Total files: {total}")
        logger.info(f"  Successful: {successful}")
        logger.info(f"  Failed: {failed}")
        logger.info(f"  Success rate: {(successful/total)*100:.1f}%")
        logger.info(f"  Total time: {total_batch_time:.2f}s")
        logger.info(f"  Average time per file: {total_batch_time/total:.2f}s")
        
        if self.extraction_stats['processing_times']:
            logger.info(f"  Processing time stats:")
            logger.info(f"    Min: {min(self.extraction_stats['processing_times']):.2f}s")
            logger.info(f"    Max: {max(self.extraction_stats['processing_times']):.2f}s")
            logger.info(f"    Mean: {np.mean(self.extraction_stats['processing_times']):.2f}s")
            logger.info(f"    Std: {np.std(self.extraction_stats['processing_times']):.2f}s")
        
        if self.extraction_stats['memory_usage']:
            logger.info(f"  Memory usage stats:")
            logger.info(f"    Min: {min(self.extraction_stats['memory_usage']):.1f}MB")
            logger.info(f"    Max: {max(self.extraction_stats['memory_usage']):.1f}MB")
            logger.info(f"    Mean: {np.mean(self.extraction_stats['memory_usage']):.1f}MB")
        
        if failed > 0:
            logger.warning(f"Failed to process {failed} files. Check extraction_errors for details.")
        
        return features_df
    
    def extract_features_from_list(self, pdbqt_files: List[str]) -> pd.DataFrame:
        """
        Extract features from a list of PDBQT files.
        
        Args:
            pdbqt_files: List of PDBQT file paths
            
        Returns:
            DataFrame with features for all compounds
        """
        logger.info(f"Extracting features from {len(pdbqt_files)} files")
        
        all_features = []
        failed_files = []
        
        for pdbqt_file in tqdm(pdbqt_files, desc="Extracting features"):
            try:
                features = self.extract_features(pdbqt_file)
                all_features.append(features.to_dict())
                self.extraction_stats['total_files_processed'] += 1
                
            except Exception as e:
                error_msg = f"Failed to process {pdbqt_file}: {str(e)}"
                logger.error(error_msg)
                failed_files.append(pdbqt_file)
                self.extraction_stats['extraction_errors'].append(error_msg)
                continue
        
        if not all_features:
            raise ValueError("No features could be extracted from any files")
        
        # Create DataFrame
        features_df = pd.DataFrame(all_features)
        
        # Log statistics
        successful = len(all_features)
        failed = len(failed_files)
        total = len(pdbqt_files)
        
        logger.info(f"Feature extraction completed: {successful}/{total} successful")
        if failed > 0:
            logger.warning(f"Failed to process {failed} files")
        
        return features_df
    
    def _extract_smiles_from_pdbqt(self, parsed_data: Dict) -> Optional[str]:
        """
        Extract SMILES string from parsed PDBQT data.
        
        Attempts to extract SMILES from PDBQT file using multiple approaches:
        1. Look for SMILES in REMARK lines
        2. Try compound name lookup in SMILES database
        3. Try to reconstruct from atom coordinates using RDKit
        
        Args:
            parsed_data: Parsed PDBQT data
            
        Returns:
            SMILES string or None if extraction fails
        """
        try:
            # Method 1: Check for SMILES in REMARK lines
            if 'header' in parsed_data:
                for remark in parsed_data['header']:
                    if 'SMILES' in remark.upper():
                        # Extract SMILES from remark line
                        parts = remark.split()
                        for i, part in enumerate(parts):
                            if 'SMILES' in part.upper() and i + 1 < len(parts):
                                potential_smiles = parts[i + 1]
                                if self._validate_smiles(potential_smiles):
                                    return potential_smiles
            
            # Method 2: Compound name lookup in SMILES database
            compound_name = parsed_data.get('file_path', '')
            if compound_name:
                compound_name = Path(compound_name).stem.lower()
                smiles = self._lookup_smiles_by_name(compound_name)
                if smiles:
                    return smiles
            
            # Method 3: Try to reconstruct from coordinates (if RDKit is available)
            if 'atoms' in parsed_data and len(parsed_data['atoms']) > 0:
                try:
                    smiles = self._reconstruct_smiles_from_coords(parsed_data['atoms'])
                    if smiles and self._validate_smiles(smiles):
                        return smiles
                except Exception as e:
                    logger.debug(f"SMILES reconstruction failed: {str(e)}")
            
            # Method 4: Return None to use coordinate-based features
            logger.debug("SMILES extraction not possible, using coordinate-based features")
            return None
            
        except Exception as e:
            logger.warning(f"SMILES extraction failed: {str(e)}")
            return None
    
    def _validate_smiles(self, smiles: str) -> bool:
        """
        Validate SMILES string using RDKit.
        
        Args:
            smiles: SMILES string to validate
            
        Returns:
            True if SMILES is valid, False otherwise
        """
        try:
            from rdkit import Chem
            mol = Chem.MolFromSmiles(smiles)
            return mol is not None
        except Exception:
            return False
    
    def _lookup_smiles_by_name(self, compound_name: str) -> Optional[str]:
        """
        Look up SMILES string by compound name.
        
        Args:
            compound_name: Name of the compound
            
        Returns:
            SMILES string or None if not found
        """
        # Common drug-like compounds and their SMILES
        smiles_database = {
            'aspirin': 'CC(=O)OC1=CC=CC=C1C(=O)O',
            'ibuprofen': 'CC(C)CC1=CC=C(C=C1)C(C)C(=O)O',
            'acetaminophen': 'CC(=O)NC1=CC=C(C=C1)O',
            'caffeine': 'CN1C=NC2=C1C(=O)N(C(=O)N2C)C',
            'morphine': 'CN1CC[C@]23C4=C5C=CC(=C4C(=O)CC[C@H]2[C@H]1CC6=C3C(=C(C=C6)O)O5)O',
            'benzene': 'C1=CC=CC=C1',
            'pyridine': 'C1=CC=NC=C1',
            'furan': 'C1=COC=C1',
            'thiophene': 'C1=CSC=C1',
            'imidazole': 'C1=CN=CN1',
            'phenol': 'C1=CC=C(C=C1)O',
            'aniline': 'C1=CC=C(C=C1)N',
            'toluene': 'CC1=CC=CC=C1',
            'glucose': 'C([C@@H]1[C@H]([C@@H]([C@H]([C@H](O1)O)O)O)O)O',
            'ethanol': 'CCO',
            'methanol': 'CO',
            'acetone': 'CC(=O)C',
            'formaldehyde': 'C=O',
            'water': 'O'
        }
        
        compound_name = compound_name.lower().strip()
        
        # Try exact match first
        if compound_name in smiles_database:
            return smiles_database[compound_name]
        
        # Try partial matches
        for known_compound, smiles in smiles_database.items():
            if known_compound in compound_name or compound_name in known_compound:
                return smiles
        
        return None
    
    def _reconstruct_smiles_from_coords(self, atoms: List[Dict]) -> Optional[str]:
        """
        Attempt to reconstruct SMILES from atomic coordinates.
        
        This is a simplified implementation that would require more sophisticated
        molecular reconstruction algorithms in practice.
        
        Args:
            atoms: List of atom dictionaries from PDBQT
            
        Returns:
            SMILES string or None if reconstruction fails
        """
        try:
            # This is a placeholder for more sophisticated reconstruction
            # In practice, you would use tools like:
            # - RDKit's conformer-to-SMILES conversion
            # - OpenBabel's 3D-to-SMILES functionality
            # - Custom graph reconstruction algorithms
            
            logger.debug("SMILES reconstruction from coordinates not implemented")
            return None
            
        except Exception as e:
            logger.debug(f"SMILES reconstruction error: {str(e)}")
            return None
    
    def _extract_coordinate_features(self, pdbqt_path: str) -> Dict[str, float]:
        """
        Extract coordinate-based features when SMILES is not available.
        
        Args:
            pdbqt_path: Path to PDBQT file
            
        Returns:
            Dictionary of coordinate-based features
        """
        try:
            # Parse PDBQT file to extract coordinates
            coordinates = []
            elements = []
            
            with open(pdbqt_path, 'r') as f:
                for line in f:
                    if line.startswith(('ATOM', 'HETATM')):
                        try:
                            x = float(line[30:38].strip())
                            y = float(line[38:46].strip())
                            z = float(line[46:54].strip())
                            element = line[76:78].strip() or line[12:16].strip()[0]
                            
                            coordinates.append([x, y, z])
                            elements.append(element)
                        except (ValueError, IndexError):
                            continue
            
            if not coordinates:
                return {}
            
            coords = np.array(coordinates)
            
            # Calculate coordinate-based features
            features = {}
            
            # Geometric center
            center = np.mean(coords, axis=0)
            
            # Distances from center
            distances = np.linalg.norm(coords - center, axis=1)
            
            # Basic geometric features
            features['coord_radius_of_gyration'] = np.sqrt(np.mean(distances**2))
            features['coord_max_distance'] = np.max(distances)
            features['coord_mean_distance'] = np.mean(distances)
            
            # Bounding box features
            min_coords = np.min(coords, axis=0)
            max_coords = np.max(coords, axis=0)
            dimensions = max_coords - min_coords
            
            features['coord_length'] = dimensions[0]
            features['coord_width'] = dimensions[1]
            features['coord_height'] = dimensions[2]
            features['coord_volume'] = np.prod(dimensions)
            
            # Shape descriptors
            if dimensions[1] > 0:
                features['coord_elongation'] = dimensions[0] / dimensions[1]
            else:
                features['coord_elongation'] = 1.0
                
            if dimensions[2] > 0:
                features['coord_flatness'] = dimensions[1] / dimensions[2]
            else:
                features['coord_flatness'] = 1.0
            
            # Atom counts
            features['coord_total_atoms'] = len(elements)
            
            return features
            
        except Exception as e:
            logger.warning(f"Error extracting coordinate features from {pdbqt_path}: {e}")
            return {}
    
    def _get_default_2d_features(self) -> Dict[str, float]:
        """
        Get default 2D molecular descriptors filled with NaN.
        
        Returns:
            Dictionary of 2D features with NaN values
        """
        return {
            # Basic molecular properties
            'molecular_weight': np.nan,
            'heavy_atoms': np.nan,
            'heteroatoms': np.nan,
            'formal_charge': np.nan,
            'rotatable_bonds': np.nan,
            
            # Lipophilicity descriptors
            'logp': np.nan,
            'molar_refractivity': np.nan,
            'tpsa': np.nan,
            
            # Hydrogen bonding
            'hbd': np.nan,
            'hba': np.nan,
            
            # Ring features
            'ring_count': np.nan,
            'aromatic_rings': np.nan,
            'aliphatic_rings': np.nan,
            'saturated_rings': np.nan,
            'aromatic_ratio': np.nan,
            
            # Shape descriptors
            'radius_of_gyration': np.nan,
            'asphericity': np.nan,
            'eccentricity': np.nan,
            'spherocity_index': np.nan,
            'elongation': np.nan,
            'flatness': np.nan,
            
            # Surface properties
            'surface_area': np.nan,
            'polar_surface_area': np.nan,
            'hydrophobic_surface_area': np.nan,
            'molecular_volume': np.nan,
            'compactness': np.nan,
            
            # Connectivity indices
            'balaban_j': np.nan,
            'bertz_ct': np.nan,
            'chi0v': np.nan,
            'chi1v': np.nan,
            'chi2v': np.nan,
            'chi3v': np.nan,
            'chi4v': np.nan
        }
        return {
            'molecular_weight': np.nan,
            'logp': np.nan,
            'tpsa': np.nan,
            'rotatable_bonds': np.nan,
            'hbd': np.nan,
            'hba': np.nan,
            'formal_charge': np.nan,
            'molar_refractivity': np.nan,
            'ring_count': np.nan,
            'aromatic_rings': np.nan,
            'aliphatic_rings': np.nan,
            'saturated_rings': np.nan,
            'heavy_atoms': np.nan,
            'heteroatoms': np.nan,
            'dipole_moment': np.nan,
            'polarizability': np.nan,
            'electronegativity': np.nan,
            'balaban_j': np.nan,
            'bertz_ct': np.nan,
            'chi0v': np.nan,
            'chi1v': np.nan,
            'chi2v': np.nan,
            'chi3v': np.nan,
            'chi4v': np.nan,
            'fsp3': np.nan,
            'fragments': np.nan,
            'bridgehead_atoms': np.nan,
            'spiro_atoms': np.nan,
            'qed': np.nan,
            'lipinski_violations': np.nan,
            'morgan_fingerprint_density': np.nan,
            'maccs_keys_density': np.nan,
            'molecular_flexibility': np.nan,
            'aromatic_ratio': np.nan,
            'heteroatom_ratio': np.nan
        }
    
    def _get_default_3d_features(self) -> Dict[str, float]:
        """Get default values for 3D structural features."""
        return {
            'radius_of_gyration': np.nan,
            'molecular_volume': np.nan,
            'surface_area': np.nan,
            'asphericity': np.nan,
            'eccentricity': np.nan,
            'spherocity_index': np.nan,
            'elongation': np.nan,
            'flatness': np.nan,
            'compactness': np.nan,
            'convexity': np.nan,
            'concavity_index': np.nan,
            'roughness_index': np.nan,
            'polar_surface_area': np.nan,
            'hydrophobic_surface_area': np.nan,
            'positive_surface_area': np.nan,
            'negative_surface_area': np.nan,
            'surface_charge_density': np.nan,
            'flexibility_index': np.nan,
            'rigidity_index': np.nan,
            'planarity': np.nan,
            'torsional_angle_variance': np.nan,
            'bond_angle_variance': np.nan
        }
    
    def get_extraction_stats(self) -> Dict[str, Any]:
        """Get extraction statistics."""
        return self.extraction_stats.copy()
    
    def reset_stats(self) -> None:
        """Reset extraction statistics."""
        self.extraction_stats = {
            'total_files_processed': 0,
            'successful_extractions': 0,
            'failed_extractions': 0,
            'extraction_errors': [],
            'processing_times': [],
            'feature_extraction_times': {
                '2d_features': [],
                '3d_features': [],
                'parsing': []
            },
            'memory_usage': [],
            'start_time': None,
            'end_time': None
        }
    
    def validate_features(self, features_df: pd.DataFrame) -> Dict[str, Any]:
        """
        Validate extracted features.
        
        Args:
            features_df: DataFrame with extracted features
            
        Returns:
            Dictionary with validation results
        """
        feature_set = FeatureSet([MolecularFeatures.from_dict(row) for _, row in features_df.iterrows()])
        return feature_set.validate_features()
    
    def get_feature_summary(self, features_df: pd.DataFrame) -> pd.DataFrame:
        """
        Get summary statistics for extracted features.
        
        Args:
            features_df: DataFrame with extracted features
            
        Returns:
            DataFrame with feature summary statistics
        """
        numeric_columns = features_df.select_dtypes(include=[np.number]).columns
        
        summary_stats = []
        for col in numeric_columns:
            stats = {
                'feature': col,
                'count': features_df[col].count(),
                'mean': features_df[col].mean(),
                'std': features_df[col].std(),
                'min': features_df[col].min(),
                'max': features_df[col].max(),
                'missing': features_df[col].isna().sum(),
                'missing_pct': (features_df[col].isna().sum() / len(features_df)) * 100
            }
            summary_stats.append(stats)
        
        return pd.DataFrame(summary_stats)
    
    def get_performance_report(self) -> Dict[str, Any]:
        """
        Generate comprehensive performance report.
        
        Returns:
            Dictionary with detailed performance metrics
        """
        stats = self.extraction_stats
        
        report = {
            'batch_summary': {
                'total_files': stats['total_files_processed'],
                'successful': stats['successful_extractions'],
                'failed': stats['failed_extractions'],
                'success_rate': (stats['successful_extractions'] / stats['total_files_processed'] * 100) 
                               if stats['total_files_processed'] > 0 else 0,
                'total_time': stats['end_time'] - stats['start_time'] if stats['start_time'] and stats['end_time'] else None
            }
        }
        
        # Processing time analysis
        if stats['processing_times']:
            times = stats['processing_times']
            report['processing_times'] = {
                'count': len(times),
                'mean': np.mean(times),
                'std': np.std(times),
                'min': min(times),
                'max': max(times),
                'median': np.median(times),
                'percentiles': {
                    '25th': np.percentile(times, 25),
                    '75th': np.percentile(times, 75),
                    '95th': np.percentile(times, 95)
                }
            }
        
        # Memory usage analysis
        if stats['memory_usage']:
            memory = stats['memory_usage']
            report['memory_usage'] = {
                'count': len(memory),
                'mean': np.mean(memory),
                'std': np.std(memory),
                'min': min(memory),
                'max': max(memory),
                'median': np.median(memory)
            }
        
        # Feature extraction timing breakdown
        if stats['feature_extraction_times']:
            report['feature_timing'] = {}
            for feature_type, times in stats['feature_extraction_times'].items():
                if times:
                    report['feature_timing'][feature_type] = {
                        'count': len(times),
                        'mean': np.mean(times),
                        'std': np.std(times),
                        'total': sum(times)
                    }
        
        # Error analysis
        if stats['extraction_errors']:
            report['errors'] = {
                'total_errors': len(stats['extraction_errors']),
                'error_rate': len(stats['extraction_errors']) / stats['total_files_processed'] * 100 
                             if stats['total_files_processed'] > 0 else 0,
                'sample_errors': stats['extraction_errors'][:5]  # First 5 errors
            }
        
        return report
    
    def save_performance_report(self, output_path: str) -> None:
        """
        Save performance report to JSON file.
        
        Args:
            output_path: Path to save the performance report
        """
        import json
        
        report = self.get_performance_report()
        
        with open(output_path, 'w') as f:
            json.dump(report, f, indent=2, default=str)
        
        logger.info(f"Performance report saved to {output_path}")
    
    def get_feature_extraction_efficiency(self) -> Dict[str, float]:
        """
        Calculate efficiency metrics for different feature extraction steps.
        
        Returns:
            Dictionary with efficiency metrics
        """
        stats = self.extraction_stats
        
        efficiency = {}
        
        # Overall efficiency
        if stats['processing_times'] and stats['total_files_processed'] > 0:
            avg_total_time = np.mean(stats['processing_times'])
            files_per_minute = 60.0 / avg_total_time
            efficiency['files_per_minute'] = files_per_minute
            efficiency['avg_time_per_file'] = avg_total_time
        
        # Feature-specific efficiency
        if stats['feature_extraction_times']:
            for feature_type, times in stats['feature_extraction_times'].items():
                if times:
                    avg_time = np.mean(times)
                    efficiency[f'{feature_type}_avg_time'] = avg_time
                    efficiency[f'{feature_type}_percentage'] = (avg_time / avg_total_time * 100) if avg_total_time else 0
        
        return efficiency
```
