## Supplementary material for "Machine Learning-Driven Prediction of TLR4 Binding Affinity: A Comprehensive Molecular Feature Analysis for Drug Discovery": Codebase: features.py

```
"""
Core data models and feature definitions for TLR4 binding prediction.

This module defines the fundamental data structures used throughout the pipeline
for representing molecular features, binding data, and prediction results.
"""

from dataclasses import dataclass, field
from typing import Dict, List, Optional, Union
import pandas as pd
import numpy as np


@dataclass
class MolecularFeatures:
    """
    Comprehensive molecular feature representation for binding prediction.
    
    Contains both 2D and 3D molecular descriptors extracted from PDBQT files
    and calculated using RDKit and PyMOL.
    """
    # Basic identification
    compound_name: str
    
    # 2D Molecular Descriptors (RDKit)
    molecular_weight: float = 0.0
    logp: float = 0.0  # Lipophilicity
    tpsa: float = 0.0  # Topological Polar Surface Area
    rotatable_bonds: int = 0
    hbd: int = 0  # H-bond donors
    hba: int = 0  # H-bond acceptors
    formal_charge: int = 0
    molar_refractivity: float = 0.0
    
    # Topological Features
    ring_count: int = 0
    aromatic_rings: int = 0
    aliphatic_rings: int = 0
    saturated_rings: int = 0
    heavy_atoms: int = 0
    heteroatoms: int = 0
    
    # Electronic Properties
    dipole_moment: float = 0.0
    polarizability: float = 0.0
    electronegativity: float = 0.0
    
    # Shape descriptors
    molecular_volume: float = 0.0
    surface_area: float = 0.0
    radius_of_gyration: float = 0.0
    asphericity: float = 0.0
    eccentricity: float = 0.0
    spherocity_index: float = 0.0
    
    # Shape descriptors (continued)
    elongation: float = 0.0
    flatness: float = 0.0
    compactness: float = 0.0
    convexity: float = 0.0
    concavity_index: float = 0.0
    roughness_index: float = 0.0
    
    # Surface properties
    polar_surface_area: float = 0.0
    hydrophobic_surface_area: float = 0.0
    positive_surface_area: float = 0.0
    negative_surface_area: float = 0.0
    surface_charge_density: float = 0.0
    
    # Conformational features
    flexibility_index: float = 0.0
    rigidity_index: float = 0.0
    planarity: float = 0.0
    torsional_angle_variance: float = 0.0
    bond_angle_variance: float = 0.0
    
    # Connectivity indices
    balaban_j: float = 0.0
    bertz_ct: float = 0.0
    chi0v: float = 0.0
    chi1v: float = 0.0
    chi2v: float = 0.0
    chi3v: float = 0.0
    chi4v: float = 0.0
    
    # Fragment counts
    fsp3: float = 0.0
    fragments: float = 0.0
    bridgehead_atoms: float = 0.0
    spiro_atoms: float = 0.0
    
    # Drug-likeness
    qed: float = 0.0
    lipinski_violations: float = 0.0
    
    # Advanced descriptors
    morgan_fingerprint_density: float = 0.0
    maccs_keys_density: float = 0.0
    molecular_flexibility: float = 0.0
    
    # Coordinate-based features (when SMILES not available)
    coord_radius_of_gyration: float = 0.0
    coord_max_distance: float = 0.0
    coord_mean_distance: float = 0.0
    coord_length: float = 0.0
    coord_width: float = 0.0
    coord_height: float = 0.0
    coord_volume: float = 0.0
    coord_elongation: float = 0.0
    coord_flatness: float = 0.0
    coord_total_atoms: float = 0.0
    aromatic_ratio: float = 0.0
    heteroatom_ratio: float = 0.0
    
    # Additional metadata
    pdbqt_file: Optional[str] = None
    smiles: Optional[str] = None
    inchi: Optional[str] = None
    
    def to_dict(self) -> Dict[str, Union[float, int, str]]:
        """Convert to dictionary for DataFrame creation."""
        result = {}
        for field_name, field_def in self.__dataclass_fields__.items():
            value = getattr(self, field_name)
            result[field_name] = value
        return result
    
    @classmethod
    def from_dict(cls, data: Dict[str, Union[float, int, str]]) -> 'MolecularFeatures':
        """Create instance from dictionary, using default values for missing fields."""
        # Filter data to only include fields that exist in the dataclass
        valid_fields = set(cls.__dataclass_fields__.keys())
        filtered_data = {k: v for k, v in data.items() if k in valid_fields}
        
        # Create instance with filtered data (missing fields will use defaults)
        return cls(**filtered_data)


@dataclass
class BindingData:
    """
    Binding affinity data from AutoDock Vina results.
    
    Represents the binding interaction between a ligand and TLR4 receptor
    with associated confidence metrics.
    """
    ligand: str
    mode: int
    affinity: float  # kcal/mol (lower values = stronger binding)
    rmsd_lb: float  # RMSD lower bound
    rmsd_ub: float  # RMSD upper bound
    
    # Additional binding metrics
    binding_energy: Optional[float] = None
    interaction_energy: Optional[float] = None
    vina_score: Optional[float] = None
    
    def is_strong_binding(self, threshold: float = -7.0) -> bool:
        """
        Check if binding is considered strong based on affinity threshold.
        
        Args:
            threshold: Affinity threshold in kcal/mol (default: -7.0)
            
        Returns:
            True if affinity is below threshold (stronger binding)
        """
        return self.affinity <= threshold
    
    def to_dict(self) -> Dict[str, Union[float, int, str]]:
        """Convert to dictionary for DataFrame creation."""
        return {
            'ligand': self.ligand,
            'mode': self.mode,
            'affinity': self.affinity,
            'rmsd_lb': self.rmsd_lb,
            'rmsd_ub': self.rmsd_ub,
            'binding_energy': self.binding_energy,
            'interaction_energy': self.interaction_energy,
            'vina_score': self.vina_score
        }


@dataclass
class PredictionResult:
    """
    Prediction result with uncertainty quantification.
    
    Contains the predicted binding affinity along with confidence intervals
    and feature importance information.
    """
    compound_name: str
    predicted_affinity: float  # kcal/mol
    confidence_interval_lower: float
    confidence_interval_upper: float
    model_used: str
    feature_contributions: Dict[str, float] = field(default_factory=dict)
    
    # Additional prediction metadata
    prediction_uncertainty: Optional[float] = None
    model_confidence: Optional[float] = None
    prediction_timestamp: Optional[str] = None
    
    def is_strong_binding(self, threshold: float = -7.0) -> bool:
        """
        Check if predicted binding is strong based on affinity threshold.
        
        Args:
            threshold: Affinity threshold in kcal/mol (default: -7.0)
            
        Returns:
            True if predicted affinity is below threshold
        """
        return self.predicted_affinity <= threshold
    
    def get_confidence_interval_width(self) -> float:
        """Get the width of the confidence interval."""
        return self.confidence_interval_upper - self.confidence_interval_lower
    
    def to_dict(self) -> Dict[str, Union[float, str, Dict]]:
        """Convert to dictionary for DataFrame creation."""
        return {
            'compound_name': self.compound_name,
            'predicted_affinity': self.predicted_affinity,
            'confidence_interval_lower': self.confidence_interval_lower,
            'confidence_interval_upper': self.confidence_interval_upper,
            'model_used': self.model_used,
            'feature_contributions': self.feature_contributions,
            'prediction_uncertainty': self.prediction_uncertainty,
            'model_confidence': self.model_confidence,
            'prediction_timestamp': self.prediction_timestamp
        }


class FeatureSet:
    """
    Container for molecular feature sets with validation and analysis methods.
    
    Provides utilities for feature validation, correlation analysis, and
    feature selection operations.
    """
    
    def __init__(self, features: List[MolecularFeatures]):
        """
        Initialize with list of molecular features.
        
        Args:
            features: List of MolecularFeatures objects
        """
        self.features = features
        self._dataframe = None
    
    @property
    def dataframe(self) -> pd.DataFrame:
        """Get features as pandas DataFrame."""
        if self._dataframe is None:
            self._dataframe = pd.DataFrame([f.to_dict() for f in self.features])
        return self._dataframe
    
    def get_numerical_features(self) -> List[str]:
        """Get list of numerical feature column names."""
        numerical_cols = []
        for col in self.dataframe.columns:
            if col not in ['compound_name', 'pdbqt_file', 'smiles', 'inchi']:
                if pd.api.types.is_numeric_dtype(self.dataframe[col]):
                    numerical_cols.append(col)
        return numerical_cols
    
    def validate_features(self) -> Dict[str, List[str]]:
        """
        Validate feature data quality.
        
        Returns:
            Dictionary with validation results and issues found
        """
        issues = {
            'missing_values': [],
            'infinite_values': [],
            'outliers': [],
            'invalid_ranges': []
        }
        
        df = self.dataframe
        numerical_cols = self.get_numerical_features()
        
        # Check for missing values
        for col in numerical_cols:
            missing_count = df[col].isna().sum()
            if missing_count > 0:
                issues['missing_values'].append(f"{col}: {missing_count} missing values")
        
        # Check for infinite values
        for col in numerical_cols:
            inf_count = np.isinf(df[col]).sum()
            if inf_count > 0:
                issues['infinite_values'].append(f"{col}: {inf_count} infinite values")
        
        # Check for outliers (using IQR method)
        for col in numerical_cols:
            Q1 = df[col].quantile(0.25)
            Q3 = df[col].quantile(0.75)
            IQR = Q3 - Q1
            lower_bound = Q1 - 1.5 * IQR
            upper_bound = Q3 + 1.5 * IQR
            outliers = df[(df[col] < lower_bound) | (df[col] > upper_bound)]
            if len(outliers) > 0:
                issues['outliers'].append(f"{col}: {len(outliers)} outliers detected")
        
        return issues
    
    def get_feature_correlations(self, threshold: float = 0.95) -> pd.DataFrame:
        """
        Find highly correlated feature pairs.
        
        Args:
            threshold: Correlation threshold for flagging high correlations
            
        Returns:
            DataFrame with highly correlated feature pairs
        """
        df = self.dataframe[self.get_numerical_features()]
        corr_matrix = df.corr().abs()
        
        # Find pairs above threshold
        high_corr_pairs = []
        for i in range(len(corr_matrix.columns)):
            for j in range(i+1, len(corr_matrix.columns)):
                corr_val = corr_matrix.iloc[i, j]
                if corr_val >= threshold:
                    high_corr_pairs.append({
                        'feature_1': corr_matrix.columns[i],
                        'feature_2': corr_matrix.columns[j],
                        'correlation': corr_val
                    })
        
        return pd.DataFrame(high_corr_pairs)
```
