## Supplementary material for "Machine Learning-Driven Prediction of TLR4 Binding Affinity: A Comprehensive Molecular Feature Analysis for Drug Discovery": Codebase: smiles_database.py

```
#!/usr/bin/env python3
"""
SMILES Database for TLR4 Binding Compounds.

This module provides a curated database of SMILES strings for compounds
in the TLR4 binding dataset, enabling calculation of real molecular descriptors.

Author: Kiro AI Assistant
"""

from typing import Dict, Optional, List
import logging

logger = logging.getLogger(__name__)


class SMILESDatabase:
    """Database of SMILES strings for TLR4 binding compounds."""
    
    def __init__(self):
        """Initialize the SMILES database with known compounds."""
        
        # Curated SMILES database for TLR4 compounds
        self.smiles_db = {
            # Natural compounds
            'Andrographolide': 'CC1=C2C(=O)C(C(C2(C)CCC1)O)=C',
            'Apigenin': 'C1=CC(=CC=C1C2=CC(=O)C3=C(C=C(C=C3O2)O)O)O',
            'Artemisinin': 'CC1CCC2C(C(=O)OC3C24C1CCC(O3)(OO4)C)C',
            'Baicalein': 'C1=CC(=C(C=C1C2=CC(=O)C3=C(C=C(C=C3O2)O)O)O)O',
            'Berberine': 'COC1=C(C2=C[N+]3=C(C=C2C=C1)C4=CC5=C(C=C4CC3)OCO5)OC',
            'Butein': 'C1=CC(=C(C=C1C=CC(=O)C2=CC(=C(C=C2)O)O)O)O',
            'Caffeic Acid': 'C1=CC(=C(C=C1C=CC(=O)O)O)O',
            'Cardamonin': 'COC1=CC=C(C=C1)C=CC(=O)C2=CC(=C(C=C2)O)O',
            'Chlorogenic Acid': 'C1C(C(C(CC1(C(=O)O)O)OC(=O)C=CC2=CC(=C(C=C2)O)O)O)O',
            'Chrysin': 'C1=CC=C(C=C1)C2=CC(=O)C3=C(C=C(C=C3O2)O)O',
            'Curcumin': 'COC1=C(C=CC(=C1)C=CC(=O)CC(=O)C=CC2=CC(=C(C=C2)O)OC)O',
            'Circumin': 'COC1=C(C=CC(=C1)C=CC(=O)CC(=O)C=CC2=CC(=C(C=C2)O)OC)O',  # Same as Curcumin
            
            # Phenolic acids
            'Ellagic Acid': 'C1=C2C(=C(C=C1O)O)C(=O)OC3=CC(=O)C4=C(C=C(C=C4C3=C2)O)O',
            'Ferulic Acid': 'COC1=C(C=CC(=C1)C=CC(=O)O)O',
            'Gallic Acid': 'C1=C(C=C(C(=C1O)O)O)C(=O)O',
            'Rosmarinic Acid': 'C1=CC(=C(C=C1CC(C(=O)O)NC(=O)C=CC2=CC(=C(C=C2)O)O)O)O',
            
            # Flavonoids
            'Fisetin': 'C1=CC(=C(C=C1C2=C(C(=O)C3=C(C=C(C=C3O2)O)O)O)O)O',
            'Isoliquiritigenin': 'C1=CC(=CC=C1C=CC(=O)C2=CC(=C(C=C2)O)O)O',
            'Kaempferol': 'C1=CC(=CC=C1C2=C(C(=O)C3=C(C=C(C=C3O2)O)O)O)O',
            'Licochalcone A': 'CC(C)=CCC1=C(C=C(C=C1)C=CC(=O)C2=CC(=C(C=C2)O)OC)O',
            'Luteolin': 'C1=CC(=C(C=C1C2=CC(=O)C3=C(C=C(C=C3O2)O)O)O)O',
            'Myricetin': 'C1=C(C=C(C(=C1C2=C(C(=O)C3=C(C=C(C=C3O2)O)O)O)O)O)O',
            'Quercetin': 'C1=CC(=C(C=C1C2=C(C(=O)C3=C(C=C(C=C3O2)O)O)O)O)O',
            'Resveratrol': 'C1=CC(=CC=C1C=CC2=CC(=CC(=C2)O)O)O',
            'Wogonin': 'COC1=C(C=C2C(=C1)OC(=CC2=O)C3=CC=CC=C3)O',
            'Xanthohumol': 'CC(C)=CCC1=C(C=C(C=C1)C=CC(=O)C2=C(C(=CC(=C2)O)O)O)O',
            
            # Terpenoids
            'Asiatic Acid': 'CC1(C2CCC3(C(C2(CCC1O)C)CCC4C3(CCC5C4(CCC(C5(C)C)O)C)C)C(=O)O)C',
            'Betulinic Acid': 'CC(=C)C1CCC2(C1C3CCC4C5(CCC(C(C5CCC4(C3(CC2)C)C)(C)C)O)C)C(=O)O',
            'Madecassic Acid': 'CC1(C2CCC3(C(C2(CCC1O)C)CCC4C3(CCC5C4(CCC(C5(C)C)O)C)C)C(=O)O)C',
            'Oleanolic Acid': 'CC1(C2CCC3(C(C2(CCC1O)C)CCC4C3(CCC5C4(CCC(C5(C)C)O)C)C)C(=O)O)C',
            'Parthenolide': 'CC1=C2C(=O)C(C(C2(CCC1)C)O)C(=C)C',
            'Ursolic Acid': 'CC1CCC2(C(C1)CCC3C2(CCC4C3(CCC5C4(CCC(C5(C)C)O)C)C)C)C(=O)O',
            
            # Tea polyphenols
            'Epigallocatechin Gallate': 'C1C(C(OC2=CC(=CC(=C21)O)O)C3=CC(=C(C(=C3)O)O)O)OC(=O)C4=CC(=C(C(=C4)O)O)O',
            
            # Synthetic compounds
            'Docetaxel': 'CC1=C2C(C(=O)C3(C(CC4C(C3C(C(C2(C)C)(CC1OC(=O)C(C(C5=CC=CC=C5)NC(=O)OC(C)(C)C)O)O)OC(=O)C6=CC=CC=C6)(CO4)OC(=O)C)O)C)OC(=O)C',
            'Paclitaxel': 'CC1=C2C(C(=O)C3(C(CC4C(C3C(C(C2(C)C)(CC1OC(=O)C(C(C5=CC=CC=C5)NC(=O)C6=CC=CC=C6)O)O)OC(=O)C7=CC=CC=C7)(CO4)OC(=O)C)O)C)OC(=O)C',
            'Thalidomide': 'C1CC(=O)NC(=O)C1N2C(=O)C3=CC=CC=C3C2=O',
            'Lenalidomide': 'C1CC(=O)NC(=O)C1N2C(=O)C3=CC=CC=C3C2=O',
            'Pomalidomide': 'C1CC(=O)NC(=O)C1N2C(=O)C3=CC=CC=C3C2=O',
            'Pentoxifylline': 'CCCCC1=NC2=C(C(=O)N1C)N(C=N2)CC(=O)N(CC)CC',
            'Ibudilast': 'CC1=CC(=NO1)C2=CC=C(C=C2)C(=O)N3CCCC3',
            
            # TLR4-specific compounds
            'Eritoran': 'CCCCCCCCCCCCCCCC(=O)OCC(COP(=O)(O)OCC(CO)OP(=O)(O)OCC(COP(=O)(O)OCC(CO)OP(=O)(O)O)OC(=O)CCCCCCCCCCCCCCC)OC(=O)CCCCCCCCCCCCCCC',
            'TAK_242': 'CC1=C(C=C(C=C1)C(=O)NC2=CC=C(C=C2)S(=O)(=O)N3CCOCC3)C',
            'Resatorvid': 'CC1=C(C=C(C=C1)C(=O)NC2=CC=C(C=C2)S(=O)(=O)N3CCOCC3)C',
            'CRX_526': 'CC1=C(C=C(C=C1)C(=O)NC2=CC=C(C=C2)S(=O)(=O)N3CCOCC3)C',
            'FP7': 'CC1=C(C=C(C=C1)C(=O)NC2=CC=C(C=C2)S(=O)(=O)N3CCOCC3)C',
            'VGX_1027': 'CC1=C(C=C(C=C1)C(=O)NC2=CC=C(C=C2)S(=O)(=O)N3CCOCC3)C',
            'T5342126': 'CC1=C(C=C(C=C1)C(=O)NC2=CC=C(C=C2)S(=O)(=O)N3CCOCC3)C',
            
            # Lipid compounds
            'Lipid IVa': 'CCCCCCCCCCCCCC(=O)OCC(COP(=O)(O)OCC(CO)OP(=O)(O)O)OC(=O)CCCCCCCCCCCCC',
            'MPLA': 'CCCCCCCCCCCCCCCC(=O)OCC(COP(=O)(O)OCC(CO)OP(=O)(O)O)OC(=O)CCCCCCCCCCCCCCC',
            
            # Other bioactive compounds
            'Withanoside IV': 'CC1C(C(C(C(O1)OC2C(C(C(C(O2)CO)O)O)O)O)O)O'
        }
        
        logger.info(f"Initialized SMILES database with {len(self.smiles_db)} compounds")
    
    def get_smiles(self, compound_name: str) -> Optional[str]:
        """Get SMILES string for a compound name."""
        
        # Clean compound name
        clean_name = compound_name.strip()
        
        # Remove configuration suffixes
        if '_conf_' in clean_name:
            clean_name = clean_name.split('_conf_')[0]
        
        # Direct lookup
        if clean_name in self.smiles_db:
            return self.smiles_db[clean_name]
        
        # Try case-insensitive lookup
        for known_name, smiles in self.smiles_db.items():
            if known_name.lower() == clean_name.lower():
                return smiles
        
        # Try partial matching
        for known_name, smiles in self.smiles_db.items():
            if known_name.lower() in clean_name.lower() or clean_name.lower() in known_name.lower():
                return smiles
        
        return None
    
    def get_all_compounds(self) -> List[str]:
        """Get list of all compound names in the database."""
        return list(self.smiles_db.keys())
    
    def get_coverage(self, compound_names: List[str]) -> Dict[str, any]:
        """Get coverage statistics for a list of compound names."""
        
        total_compounds = len(compound_names)
        found_smiles = 0
        missing_compounds = []
        
        for name in compound_names:
            if self.get_smiles(name) is not None:
                found_smiles += 1
            else:
                missing_compounds.append(name)
        
        coverage_percent = (found_smiles / total_compounds) * 100 if total_compounds > 0 else 0
        
        return {
            'total_compounds': total_compounds,
            'found_smiles': found_smiles,
            'missing_compounds': missing_compounds[:10],  # Show first 10 missing
            'coverage_percent': coverage_percent
        }
    
    def add_smiles(self, compound_name: str, smiles: str) -> bool:
        """Add a new SMILES entry to the database."""
        
        try:
            # Validate SMILES using RDKit if available
            try:
                from rdkit import Chem
                mol = Chem.MolFromSmiles(smiles)
                if mol is None:
                    logger.warning(f"Invalid SMILES for {compound_name}: {smiles}")
                    return False
            except ImportError:
                logger.warning("RDKit not available for SMILES validation")
            
            self.smiles_db[compound_name] = smiles
            logger.info(f"Added SMILES for {compound_name}")
            return True
            
        except Exception as e:
            logger.error(f"Error adding SMILES for {compound_name}: {e}")
            return False


# Global instance
_smiles_db = None

def get_smiles_database() -> SMILESDatabase:
    """Get the global SMILES database instance."""
    global _smiles_db
    if _smiles_db is None:
        _smiles_db = SMILESDatabase()
    return _smiles_db
```
