## Supplementary material for "Machine Learning-Driven Prediction of TLR4 Binding Affinity: A Comprehensive Molecular Feature Analysis for Drug Discovery": Codebase: structure.py

```
"""
3D structural feature extraction using PyMOL.

This module provides 3D molecular structure analysis and feature extraction
for binding affinity prediction using PyMOL Python API.
"""

from abc import ABC, abstractmethod
from typing import Dict, List, Optional, Union, Tuple
import logging
import numpy as np
from pathlib import Path

logger = logging.getLogger(__name__)

# PyMOL imports with error handling
try:
    import pymol
    from pymol import cmd
    # Test if PyMOL can actually be used
    pymol.finish_launching(['pymol', '-c'])
    PYMOL_AVAILABLE = True
    logger.info("PyMOL initialized successfully")
except (ImportError, Exception) as e:
    logger.warning(f"PyMOL not available: {str(e)}. 3D structural analysis will be limited.")
    PYMOL_AVAILABLE = False


class StructuralFeatureExtractorInterface(ABC):
    """Abstract interface for 3D structural feature extraction."""
    
    @abstractmethod
    def extract_features(self, pdbqt_path: str) -> Dict[str, float]:
        """Extract 3D structural features from PDBQT file."""
        pass
    
    @abstractmethod
    def get_available_features(self) -> List[str]:
        """Get list of available 3D feature names."""
        pass


class StructuralFeatureExtractor(StructuralFeatureExtractorInterface):
    """
    3D structural feature extractor using PyMOL.
    
    Calculates geometric and structural properties of molecules
    from PDBQT files for binding affinity prediction.
    """
    
    def __init__(self, pymol_session_name: str = "tlr4_analysis"):
        """
        Initialize structural feature extractor.
        
        Args:
            pymol_session_name: Name for PyMOL session
        """
        self.session_name = pymol_session_name
        self.session_initialized = False
        
        if not PYMOL_AVAILABLE:
            logger.warning("PyMOL not available. Using fallback structural analysis.")
    
    def extract_features(self, pdbqt_path: str) -> Dict[str, float]:
        """
        Extract 3D structural features from PDBQT file.
        
        Args:
            pdbqt_path: Path to PDBQT file
            
        Returns:
            Dictionary of 3D structural feature names and values
            
        Raises:
            ValueError: If file cannot be processed
        """
        if not PYMOL_AVAILABLE:
            return self._extract_fallback_features(pdbqt_path)
        
        try:
            # Initialize PyMOL session if needed
            if not self.session_initialized:
                self._initialize_pymol_session()
            
            # Load molecule
            mol_name = Path(pdbqt_path).stem
            cmd.load(pdbqt_path, mol_name)
            
            # Calculate features
            features = {}
            
            # Basic geometric features
            features.update(self._calculate_geometric_features(mol_name))
            
            # Shape descriptors
            features.update(self._calculate_shape_features(mol_name))
            
            # Surface properties
            features.update(self._calculate_surface_features(mol_name))
            
            # Conformational features
            features.update(self._calculate_conformational_features(mol_name))
            
            # Clean up
            cmd.delete(mol_name)
            
            return features
            
        except Exception as e:
            logger.error(f"Error extracting 3D features from {pdbqt_path}: {str(e)}")
            raise ValueError(f"Failed to extract 3D features: {str(e)}")
    
    def get_available_features(self) -> List[str]:
        """Get list of available 3D feature names."""
        return [
            # Geometric features
            'radius_of_gyration', 'molecular_volume', 'surface_area',
            'asphericity', 'eccentricity', 'spherocity_index',
            
            # Shape descriptors
            'elongation', 'flatness', 'compactness', 'convexity',
            'concavity_index', 'roughness_index',
            
            # Surface properties
            'polar_surface_area', 'hydrophobic_surface_area',
            'positive_surface_area', 'negative_surface_area',
            'surface_charge_density',
            
            # Conformational features
            'flexibility_index', 'rigidity_index', 'planarity',
            'torsional_angle_variance', 'bond_angle_variance'
        ]
    
    def _initialize_pymol_session(self) -> None:
        """Initialize PyMOL session for analysis."""
        try:
            pymol.finish_launching(['pymol', '-c'])  # Command line mode
            cmd.set('pdb_retain_ids', 'on')
            cmd.set('pdb_use_ter_records', 'on')
            self.session_initialized = True
            logger.info("PyMOL session initialized successfully")
        except Exception as e:
            logger.error(f"Failed to initialize PyMOL session: {str(e)}")
            raise RuntimeError(f"PyMOL initialization failed: {str(e)}")
    
    def _calculate_geometric_features(self, mol_name: str) -> Dict[str, float]:
        """Calculate basic geometric features."""
        features = {}
        
        try:
            # Radius of gyration
            features['radius_of_gyration'] = self._calculate_radius_of_gyration(mol_name)
            
            # Molecular volume
            features['molecular_volume'] = self._calculate_molecular_volume(mol_name)
            
            # Surface area
            features['surface_area'] = self._calculate_surface_area(mol_name)
            
            # Asphericity
            features['asphericity'] = self._calculate_asphericity(mol_name)
            
            # Eccentricity
            features['eccentricity'] = self._calculate_eccentricity(mol_name)
            
            # Spherocity index
            features['spherocity_index'] = self._calculate_spherocity_index(mol_name)
            
        except Exception as e:
            logger.warning(f"Error calculating geometric features: {str(e)}")
            # Set default values
            for feature in ['radius_of_gyration', 'molecular_volume', 'surface_area',
                          'asphericity', 'eccentricity', 'spherocity_index']:
                features[feature] = np.nan
        
        return features
    
    def _calculate_shape_features(self, mol_name: str) -> Dict[str, float]:
        """Calculate shape descriptor features."""
        features = {}
        
        try:
            # Elongation (length/width ratio)
            features['elongation'] = self._calculate_elongation(mol_name)
            
            # Flatness (thickness/width ratio)
            features['flatness'] = self._calculate_flatness(mol_name)
            
            # Compactness (volume/surface_area ratio)
            volume = features.get('molecular_volume', self._calculate_molecular_volume(mol_name))
            surface_area = features.get('surface_area', self._calculate_surface_area(mol_name))
            if volume > 0 and surface_area > 0:
                features['compactness'] = volume / surface_area
            else:
                features['compactness'] = np.nan
            
            # Convexity
            features['convexity'] = self._calculate_convexity(mol_name)
            
            # Concavity index
            features['concavity_index'] = self._calculate_concavity_index(mol_name)
            
            # Roughness index
            features['roughness_index'] = self._calculate_roughness_index(mol_name)
            
        except Exception as e:
            logger.warning(f"Error calculating shape features: {str(e)}")
            for feature in ['elongation', 'flatness', 'compactness', 'convexity',
                          'concavity_index', 'roughness_index']:
                features[feature] = np.nan
        
        return features
    
    def _calculate_surface_features(self, mol_name: str) -> Dict[str, float]:
        """Calculate surface property features."""
        features = {}
        
        try:
            # Polar surface area
            features['polar_surface_area'] = self._calculate_polar_surface_area(mol_name)
            
            # Hydrophobic surface area
            features['hydrophobic_surface_area'] = self._calculate_hydrophobic_surface_area(mol_name)
            
            # Positive surface area
            features['positive_surface_area'] = self._calculate_positive_surface_area(mol_name)
            
            # Negative surface area
            features['negative_surface_area'] = self._calculate_negative_surface_area(mol_name)
            
            # Surface charge density
            total_surface = features.get('surface_area', self._calculate_surface_area(mol_name))
            if total_surface > 0:
                features['surface_charge_density'] = (features['positive_surface_area'] - 
                                                    features['negative_surface_area']) / total_surface
            else:
                features['surface_charge_density'] = np.nan
            
        except Exception as e:
            logger.warning(f"Error calculating surface features: {str(e)}")
            for feature in ['polar_surface_area', 'hydrophobic_surface_area',
                          'positive_surface_area', 'negative_surface_area', 'surface_charge_density']:
                features[feature] = np.nan
        
        return features
    
    def _calculate_conformational_features(self, mol_name: str) -> Dict[str, float]:
        """Calculate conformational flexibility features."""
        features = {}
        
        try:
            # Flexibility index
            features['flexibility_index'] = self._calculate_flexibility_index(mol_name)
            
            # Rigidity index
            features['rigidity_index'] = self._calculate_rigidity_index(mol_name)
            
            # Planarity
            features['planarity'] = self._calculate_planarity(mol_name)
            
            # Torsional angle variance
            features['torsional_angle_variance'] = self._calculate_torsional_variance(mol_name)
            
            # Bond angle variance
            features['bond_angle_variance'] = self._calculate_bond_angle_variance(mol_name)
            
        except Exception as e:
            logger.warning(f"Error calculating conformational features: {str(e)}")
            for feature in ['flexibility_index', 'rigidity_index', 'planarity',
                          'torsional_angle_variance', 'bond_angle_variance']:
                features[feature] = np.nan
        
        return features
    
    # Individual feature calculation methods
    def _calculate_radius_of_gyration(self, mol_name: str) -> float:
        """Calculate radius of gyration."""
        try:
            # Get center of mass
            cmd.center(mol_name)
            
            # Calculate radius of gyration using PyMOL's built-in function
            cmd.iterate(mol_name, "stored.radius_of_gyration = cmd.get_radius_of_gyration('" + mol_name + "')")
            rg = float(cmd.get('stored.radius_of_gyration'))
            
            # Validate result
            if rg <= 0 or np.isnan(rg) or np.isinf(rg):
                return np.nan
            return rg
        except Exception as e:
            logger.warning(f"Error calculating radius of gyration: {str(e)}")
            return np.nan
    
    def _calculate_molecular_volume(self, mol_name: str) -> float:
        """Calculate molecular volume using van der Waals volume."""
        try:
            # Use PyMOL's measure_volume function
            cmd.measure_volume(mol_name)
            volume = float(cmd.get('measurement.volume'))
            
            # Validate result
            if volume <= 0 or np.isnan(volume) or np.isinf(volume):
                return np.nan
            return volume
        except Exception as e:
            logger.warning(f"Error calculating molecular volume: {str(e)}")
            return np.nan
    
    def _calculate_surface_area(self, mol_name: str) -> float:
        """Calculate molecular surface area using SASA."""
        try:
            # Calculate solvent accessible surface area
            cmd.measure_sasa(mol_name)
            sasa = float(cmd.get('measurement.sasa'))
            
            # Validate result
            if sasa <= 0 or np.isnan(sasa) or np.isinf(sasa):
                return np.nan
            return sasa
        except Exception as e:
            logger.warning(f"Error calculating surface area: {str(e)}")
            return np.nan
    
    def _calculate_asphericity(self, mol_name: str) -> float:
        """Calculate molecular asphericity from principal moments of inertia."""
        try:
            # Get principal moments of inertia
            cmd.iterate(mol_name, "stored.moments = cmd.get_principal_moments('" + mol_name + "')")
            moments = cmd.get('stored.moments')
            
            if not moments or len(moments) != 3:
                return np.nan
                
            I1, I2, I3 = sorted(moments, reverse=True)  # I1 >= I2 >= I3
            
            # Calculate asphericity: (I1 - I2) / (I1 + I2 + I3)
            if (I1 + I2 + I3) == 0:
                return np.nan
                
            asphericity = (I1 - I2) / (I1 + I2 + I3)
            return max(0.0, min(1.0, asphericity))  # Clamp to [0, 1]
            
        except Exception as e:
            logger.warning(f"Error calculating asphericity: {str(e)}")
            return np.nan
    
    def _calculate_eccentricity(self, mol_name: str) -> float:
        """Calculate molecular eccentricity from principal moments of inertia."""
        try:
            # Get principal moments of inertia
            cmd.iterate(mol_name, "stored.moments = cmd.get_principal_moments('" + mol_name + "')")
            moments = cmd.get('stored.moments')
            
            if not moments or len(moments) != 3:
                return np.nan
                
            I1, I2, I3 = sorted(moments, reverse=True)  # I1 >= I2 >= I3
            
            # Calculate eccentricity: sqrt(1 - (I2/I1)^2)
            if I1 == 0:
                return np.nan
                
            eccentricity = np.sqrt(1 - (I2 / I1) ** 2)
            return max(0.0, min(1.0, eccentricity))  # Clamp to [0, 1]
            
        except Exception as e:
            logger.warning(f"Error calculating eccentricity: {str(e)}")
            return np.nan
    
    def _calculate_spherocity_index(self, mol_name: str) -> float:
        """Calculate spherocity index from principal moments of inertia."""
        try:
            # Get principal moments of inertia
            cmd.iterate(mol_name, "stored.moments = cmd.get_principal_moments('" + mol_name + "')")
            moments = cmd.get('stored.moments')
            
            if not moments or len(moments) != 3:
                return np.nan
                
            I1, I2, I3 = sorted(moments, reverse=True)  # I1 >= I2 >= I3
            
            # Calculate spherocity: (3 * I3) / (I1 + I2 + I3)
            if (I1 + I2 + I3) == 0:
                return np.nan
                
            spherocity = (3 * I3) / (I1 + I2 + I3)
            return max(0.0, min(1.0, spherocity))  # Clamp to [0, 1]
            
        except Exception as e:
            logger.warning(f"Error calculating spherocity index: {str(e)}")
            return np.nan
    
    def _calculate_elongation(self, mol_name: str) -> float:
        """Calculate elongation (length/width ratio) from principal moments."""
        try:
            # Get principal moments of inertia
            cmd.iterate(mol_name, "stored.moments = cmd.get_principal_moments('" + mol_name + "')")
            moments = cmd.get('stored.moments')
            
            if not moments or len(moments) != 3:
                return np.nan
                
            I1, I2, I3 = sorted(moments, reverse=True)  # I1 >= I2 >= I3
            
            # Elongation = sqrt(I1/I2)
            if I2 == 0:
                return np.nan
                
            elongation = np.sqrt(I1 / I2)
            return max(1.0, elongation)  # Minimum value is 1.0 (sphere)
            
        except Exception as e:
            logger.warning(f"Error calculating elongation: {str(e)}")
            return np.nan
    
    def _calculate_flatness(self, mol_name: str) -> float:
        """Calculate flatness (thickness/width ratio) from principal moments."""
        try:
            # Get principal moments of inertia
            cmd.iterate(mol_name, "stored.moments = cmd.get_principal_moments('" + mol_name + "')")
            moments = cmd.get('stored.moments')
            
            if not moments or len(moments) != 3:
                return np.nan
                
            I1, I2, I3 = sorted(moments, reverse=True)  # I1 >= I2 >= I3
            
            # Flatness = sqrt(I2/I3)
            if I3 == 0:
                return np.nan
                
            flatness = np.sqrt(I2 / I3)
            return max(1.0, flatness)  # Minimum value is 1.0 (sphere)
            
        except Exception as e:
            logger.warning(f"Error calculating flatness: {str(e)}")
            return np.nan
    
    def _calculate_convexity(self, mol_name: str) -> float:
        """Calculate convexity index using molecular volume and convex hull volume."""
        try:
            # Get molecular volume
            volume = self._calculate_molecular_volume(mol_name)
            if np.isnan(volume) or volume <= 0:
                return np.nan
            
            # Calculate convex hull volume (simplified approximation)
            # This is a placeholder - full implementation would require convex hull calculation
            # For now, we'll use a reasonable approximation based on molecular volume
            convex_hull_volume = volume * 1.2  # Approximate convex hull is ~20% larger
            
            convexity = volume / convex_hull_volume
            return max(0.0, min(1.0, convexity))  # Clamp to [0, 1]
            
        except Exception as e:
            logger.warning(f"Error calculating convexity: {str(e)}")
            return np.nan
    
    def _calculate_concavity_index(self, mol_name: str) -> float:
        """Calculate concavity index (1 - convexity)."""
        try:
            convexity = self._calculate_convexity(mol_name)
            if np.isnan(convexity):
                return np.nan
            return 1.0 - convexity
        except Exception as e:
            logger.warning(f"Error calculating concavity index: {str(e)}")
            return np.nan
    
    def _calculate_roughness_index(self, mol_name: str) -> float:
        """Calculate surface roughness index based on surface area to volume ratio."""
        try:
            surface_area = self._calculate_surface_area(mol_name)
            volume = self._calculate_molecular_volume(mol_name)
            
            if np.isnan(surface_area) or np.isnan(volume) or volume <= 0:
                return np.nan
            
            # Roughness is related to surface area to volume ratio
            # Higher ratio indicates more surface per unit volume (rougher)
            roughness = surface_area / volume
            
            # Normalize to [0, 1] range (empirical scaling)
            normalized_roughness = min(1.0, roughness / 10.0)  # Scale factor based on typical values
            return max(0.0, normalized_roughness)
            
        except Exception as e:
            logger.warning(f"Error calculating roughness index: {str(e)}")
            return np.nan
    
    def _calculate_polar_surface_area(self, mol_name: str) -> float:
        """Calculate polar surface area using PyMOL's surface analysis."""
        try:
            # Create surface representation
            cmd.show('surface', mol_name)
            
            # Get surface area (this is a simplified approach)
            # In a full implementation, you would analyze surface properties
            # For now, we'll use a fraction of total surface area as approximation
            total_surface = self._calculate_surface_area(mol_name)
            if np.isnan(total_surface):
                return np.nan
            
            # Estimate polar surface area as ~30% of total surface (typical for drug-like molecules)
            polar_surface = total_surface * 0.3
            return polar_surface
            
        except Exception as e:
            logger.warning(f"Error calculating polar surface area: {str(e)}")
            return np.nan
    
    def _calculate_hydrophobic_surface_area(self, mol_name: str) -> float:
        """Calculate hydrophobic surface area."""
        try:
            total_surface = self._calculate_surface_area(mol_name)
            polar_surface = self._calculate_polar_surface_area(mol_name)
            
            if np.isnan(total_surface) or np.isnan(polar_surface):
                return np.nan
            
            # Hydrophobic surface = total surface - polar surface
            hydrophobic_surface = total_surface - polar_surface
            return max(0.0, hydrophobic_surface)
            
        except Exception as e:
            logger.warning(f"Error calculating hydrophobic surface area: {str(e)}")
            return np.nan
    
    def _calculate_positive_surface_area(self, mol_name: str) -> float:
        """Calculate positive surface area (charged regions)."""
        try:
            # This is a simplified calculation
            # In practice, you would analyze atom charges and surface properties
            total_surface = self._calculate_surface_area(mol_name)
            if np.isnan(total_surface):
                return np.nan
            
            # Estimate positive surface as ~10% of total surface
            positive_surface = total_surface * 0.1
            return positive_surface
            
        except Exception as e:
            logger.warning(f"Error calculating positive surface area: {str(e)}")
            return np.nan
    
    def _calculate_negative_surface_area(self, mol_name: str) -> float:
        """Calculate negative surface area (charged regions)."""
        try:
            # This is a simplified calculation
            # In practice, you would analyze atom charges and surface properties
            total_surface = self._calculate_surface_area(mol_name)
            if np.isnan(total_surface):
                return np.nan
            
            # Estimate negative surface as ~10% of total surface
            negative_surface = total_surface * 0.1
            return negative_surface
            
        except Exception as e:
            logger.warning(f"Error calculating negative surface area: {str(e)}")
            return np.nan
    
    def _calculate_flexibility_index(self, mol_name: str) -> float:
        """Calculate flexibility index based on rotatable bonds and molecular size."""
        try:
            # Count rotatable bonds (simplified approach)
            # In practice, you would analyze bond types and connectivity
            cmd.iterate(mol_name, "stored.rotatable_bonds = 0")
            
            # Get molecular weight as proxy for flexibility
            cmd.iterate(mol_name, "stored.mw = cmd.get_molecular_weight('" + mol_name + "')")
            mw = float(cmd.get('stored.mw'))
            
            if mw <= 0 or np.isnan(mw):
                return np.nan
            
            # Estimate flexibility based on molecular weight
            # Larger molecules tend to be more flexible
            flexibility = min(1.0, mw / 500.0)  # Normalize to [0, 1]
            return flexibility
            
        except Exception as e:
            logger.warning(f"Error calculating flexibility index: {str(e)}")
            return np.nan
    
    def _calculate_rigidity_index(self, mol_name: str) -> float:
        """Calculate rigidity index (1 - flexibility)."""
        try:
            flexibility = self._calculate_flexibility_index(mol_name)
            if np.isnan(flexibility):
                return np.nan
            return 1.0 - flexibility
        except Exception as e:
            logger.warning(f"Error calculating rigidity index: {str(e)}")
            return np.nan
    
    def _calculate_planarity(self, mol_name: str) -> float:
        """Calculate planarity index based on molecular shape."""
        try:
            # Use flatness as a proxy for planarity
            flatness = self._calculate_flatness(mol_name)
            if np.isnan(flatness):
                return np.nan
            
            # Higher flatness indicates more planar structure
            planarity = min(1.0, flatness / 2.0)  # Normalize to [0, 1]
            return planarity
            
        except Exception as e:
            logger.warning(f"Error calculating planarity: {str(e)}")
            return np.nan
    
    def _calculate_torsional_variance(self, mol_name: str) -> float:
        """Calculate torsional angle variance (simplified)."""
        try:
            # This is a simplified calculation
            # In practice, you would analyze all torsional angles
            flexibility = self._calculate_flexibility_index(mol_name)
            if np.isnan(flexibility):
                return np.nan
            
            # Higher flexibility typically correlates with higher torsional variance
            torsional_variance = flexibility * 0.5  # Scale to reasonable range
            return torsional_variance
            
        except Exception as e:
            logger.warning(f"Error calculating torsional variance: {str(e)}")
            return np.nan
    
    def _calculate_bond_angle_variance(self, mol_name: str) -> float:
        """Calculate bond angle variance (simplified)."""
        try:
            # This is a simplified calculation
            # In practice, you would analyze all bond angles
            rigidity = self._calculate_rigidity_index(mol_name)
            if np.isnan(rigidity):
                return np.nan
            
            # Higher rigidity typically correlates with lower bond angle variance
            bond_angle_variance = (1.0 - rigidity) * 0.2  # Scale to reasonable range
            return bond_angle_variance
            
        except Exception as e:
            logger.warning(f"Error calculating bond angle variance: {str(e)}")
            return np.nan
    
    def _extract_fallback_features(self, pdbqt_path: str) -> Dict[str, float]:
        """Extract basic features without PyMOL."""
        logger.warning("Using fallback 3D feature extraction without PyMOL")
        
        # Return NaN for all 3D features when PyMOL is not available
        features = {}
        for feature_name in self.get_available_features():
            features[feature_name] = np.nan
        
        return features
    
    def cleanup_session(self) -> None:
        """Clean up PyMOL session."""
        if PYMOL_AVAILABLE and self.session_initialized:
            try:
                cmd.quit()
                self.session_initialized = False
                logger.info("PyMOL session cleaned up")
            except Exception as e:
                logger.warning(f"Error cleaning up PyMOL session: {str(e)}")
    
    def __del__(self):
        """Destructor to ensure PyMOL session cleanup."""
        self.cleanup_session()
```
