## Supplementary material for "Machine Learning-Driven Prediction of TLR4 Binding Affinity: A Comprehensive Molecular Feature Analysis for Drug Discovery": Codebase: data_processing.py

```
#!/usr/bin/env python3
"""
Data processing utilities for TLR4 binding prediction.

This module handles data loading, cleaning, and preprocessing while preserving
chemical diversity.
"""

import pandas as pd
import numpy as np
import logging
from typing import Tuple

logger = logging.getLogger(__name__)


def load_binding_data(binding_csv: str) -> pd.DataFrame:
    """Load and clean binding affinity data conservatively."""
    logger.info(f"Loading binding data from {binding_csv}")
    
    binding_df = pd.read_csv(binding_csv)
    binding_df = binding_df.rename(columns={'ligand': 'compound'})
    binding_df['compound'] = binding_df['compound'].str.strip()
    
    # Conservative cleaning - only remove clear errors
    initial_count = len(binding_df)
    binding_df = binding_df.dropna(subset=['affinity'])
    binding_df = binding_df[np.isfinite(binding_df['affinity'])]
    binding_df = binding_df[binding_df['affinity'] < 0]  # Only negative affinities
    binding_df = binding_df[binding_df['affinity'] > -20]  # Remove extreme outliers
    
    # Extract base compound names to prevent data leakage
    binding_df['base_compound'] = binding_df['compound'].apply(
        lambda x: x.split('_conf_')[0] if '_conf_' in x else x
    )
    
    # Keep BEST affinity per base compound (prevents data leakage)
    binding_df = binding_df.loc[binding_df.groupby('base_compound')['affinity'].idxmin()]
    binding_df['compound'] = binding_df['base_compound']
    binding_df = binding_df.drop('base_compound', axis=1)
    
    logger.info(f"Conservative cleaning: {len(binding_df)} unique compounds (from {initial_count} records)")
    logger.info(f"Affinity range: {binding_df['affinity'].min():.3f} to {binding_df['affinity'].max():.3f} kcal/mol")
    
    return binding_df


def integrate_features_and_binding(features_df: pd.DataFrame, binding_df: pd.DataFrame) -> pd.DataFrame:
    """Integrate molecular features with binding affinity data."""
    logger.info("Integrating molecular features with binding data")
    
    # Merge features with binding data
    integrated_df = pd.merge(features_df, binding_df, on='compound', how='inner')
    integrated_df = integrated_df.dropna()
    
    logger.info(f"Integrated dataset: {len(integrated_df)} compounds")
    
    # Add derived binding features
    integrated_df['binding_efficiency'] = np.abs(integrated_df['affinity']) / integrated_df['molecular_weight'] * 1000
    integrated_df['ligand_efficiency'] = np.abs(integrated_df['affinity']) / integrated_df['heavy_atoms']
    
    return integrated_df


def remove_true_duplicates(df: pd.DataFrame) -> pd.DataFrame:
    """Remove only true duplicates (identical SMILES AND identical affinities)."""
    logger.info("Removing only true duplicates...")
    
    initial_count = len(df)
    
    # Remove only compounds with identical SMILES AND identical affinities
    df_unique = df.drop_duplicates(subset=['smiles', 'affinity'], keep='first')
    
    removed_count = initial_count - len(df_unique)
    logger.info(f"Removed {removed_count} true duplicates (identical SMILES + affinity)")
    
    # Show preserved diversity
    smiles_counts = df['smiles'].value_counts()
    multi_smiles = smiles_counts[smiles_counts > 1]
    
    if len(multi_smiles) > 0:
        logger.info(f"Preserved {len(multi_smiles)} SMILES with multiple affinity values")
        for smiles in multi_smiles.index[:3]:  # Show first 3
            compounds_with_smiles = df[df['smiles'] == smiles]
            affinities = compounds_with_smiles['affinity'].tolist()
            compound_names = compounds_with_smiles['compound'].tolist()
            logger.info(f"  Compounds: {compound_names}")
            logger.info(f"  Affinities: {[f'{a:.3f}' for a in affinities]}")
    
    return df_unique.reset_index(drop=True)
```
