## Supplementary material for "Machine Learning-Driven Prediction of TLR4 Binding Affinity: A Comprehensive Molecular Feature Analysis for Drug Discovery": Codebase: feature_extraction.py

```
#!/usr/bin/env python3
"""
Molecular feature extraction for TLR4 binding prediction.

This module extracts comprehensive molecular descriptors from SMILES strings
while preserving chemical diversity.
"""

import pandas as pd
import numpy as np
import logging
from pathlib import Path
from typing import Dict, List
from tqdm import tqdm

logger = logging.getLogger(__name__)

try:
    from rdkit import Chem
    from rdkit.Chem import Descriptors, Crippen, Fragments
    RDKIT_AVAILABLE = True
except ImportError:
    RDKIT_AVAILABLE = False
    logger.warning("RDKit not available - molecular descriptors will be limited")


class MolecularFeatureExtractor:
    """Extract comprehensive molecular features for TLR4 binding prediction."""
    
    def __init__(self, smiles_database):
        """Initialize with SMILES database."""
        self.smiles_db = smiles_database
        self.logger = logging.getLogger(__name__)
    
    def calculate_molecular_descriptors(self, smiles: str) -> Dict[str, float]:
        """Calculate comprehensive molecular descriptors from SMILES."""
        if not RDKIT_AVAILABLE:
            return {}
        
        try:
            mol = Chem.MolFromSmiles(smiles)
            if mol is None:
                return {}
            
            descriptors = {
                # Basic properties
                'molecular_weight': Descriptors.MolWt(mol),
                'logp': Crippen.MolLogP(mol),
                'tpsa': Descriptors.TPSA(mol),
                'rotatable_bonds': Descriptors.NumRotatableBonds(mol),
                'hbd': Descriptors.NumHDonors(mol),
                'hba': Descriptors.NumHAcceptors(mol),
                'heavy_atoms': Descriptors.HeavyAtomCount(mol),
                'molar_refractivity': Crippen.MolMR(mol),
                'fraction_csp3': Descriptors.FractionCSP3(mol),
                
                # Ring descriptors
                'aromatic_rings': Descriptors.NumAromaticRings(mol),
                'aliphatic_rings': Descriptors.NumAliphaticRings(mol),
                'ring_count': Descriptors.RingCount(mol),
                'num_aromatic_carbocycles': Descriptors.NumAromaticCarbocycles(mol),
                'num_aromatic_heterocycles': Descriptors.NumAromaticHeterocycles(mol),
                'num_aliphatic_carbocycles': Descriptors.NumAliphaticCarbocycles(mol),
                'num_saturated_carbocycles': Descriptors.NumSaturatedCarbocycles(mol),
                
                # Topological descriptors
                'balaban_j': Descriptors.BalabanJ(mol),
                'bertz_ct': Descriptors.BertzCT(mol),
                'chi0v': Descriptors.Chi0v(mol),
                'chi1v': Descriptors.Chi1v(mol),
                'chi2v': Descriptors.Chi2v(mol),
                'chi3v': Descriptors.Chi3v(mol),
                'kappa1': Descriptors.Kappa1(mol),
                'kappa2': Descriptors.Kappa2(mol),
                'kappa3': Descriptors.Kappa3(mol),
                'hall_kier_alpha': Descriptors.HallKierAlpha(mol),
                'ipc': Descriptors.Ipc(mol),
                
                # VSA descriptors
                'slogp_vsa1': Descriptors.SlogP_VSA1(mol),
                'slogp_vsa2': Descriptors.SlogP_VSA2(mol),
                'slogp_vsa3': Descriptors.SlogP_VSA3(mol),
                'smr_vsa1': Descriptors.SMR_VSA1(mol),
                'smr_vsa2': Descriptors.SMR_VSA2(mol),
                'smr_vsa3': Descriptors.SMR_VSA3(mol),
                'peoe_vsa1': Descriptors.PEOE_VSA1(mol),
                'peoe_vsa2': Descriptors.PEOE_VSA2(mol),
                
                # Fragment descriptors
                'fr_aliphatic_oh': Fragments.fr_Al_OH(mol),
                'fr_aromatic_oh': Fragments.fr_Ar_OH(mol),
                'fr_nh': Fragments.fr_NH0(mol) + Fragments.fr_NH1(mol) + Fragments.fr_NH2(mol),
                'fr_carbonyl': Fragments.fr_C_O(mol),
                'fr_ester': Fragments.fr_COO(mol),
                'fr_ether': Fragments.fr_ether(mol),
                'fr_halogen': Fragments.fr_halogen(mol),
                'fr_benzene': Fragments.fr_benzene(mol),
                'fr_amide': Fragments.fr_amide(mol),
                
                # Drug-likeness
                'qed': Descriptors.qed(mol),
                'lipinski_violations': (
                    (Descriptors.MolWt(mol) > 500) +
                    (Crippen.MolLogP(mol) > 5) +
                    (Descriptors.NumHDonors(mol) > 5) +
                    (Descriptors.NumHAcceptors(mol) > 10)
                ),
            }
            
            # Derived features
            descriptors.update({
                'aromatic_ratio': descriptors['aromatic_rings'] / max(descriptors['ring_count'], 1),
                'flexibility': descriptors['rotatable_bonds'] / max(descriptors['heavy_atoms'], 1),
                'complexity': descriptors['bertz_ct'] / max(descriptors['heavy_atoms'], 1),
                'polarity_ratio': descriptors['tpsa'] / max(descriptors['molecular_weight'], 1),
                'hb_ratio': (descriptors['hbd'] + descriptors['hba']) / max(descriptors['heavy_atoms'], 1),
                'ring_density': descriptors['ring_count'] / max(descriptors['heavy_atoms'], 1),
                'shape_index': descriptors['kappa1'] * descriptors['kappa2'] / max(descriptors['kappa3'], 0.1),
            })
            
            # Ensure finite values
            for key, value in descriptors.items():
                if not np.isfinite(value):
                    descriptors[key] = 0.0
            
            return descriptors
            
        except Exception as e:
            self.logger.error(f"Error calculating descriptors for SMILES {smiles}: {e}")
            return {}
    
    def extract_features_from_files(self, pdbqt_dir: str) -> pd.DataFrame:
        """Extract molecular features from PDBQT files."""
        pdbqt_files = list(Path(pdbqt_dir).glob("*.pdbqt"))
        features_list = []
        
        for pdbqt_file in tqdm(pdbqt_files, desc="Extracting molecular features"):
            compound_name = pdbqt_file.stem
            
            # Clean compound name (remove conformation suffixes)
            if '_conf_' in compound_name:
                base_name = compound_name.split('_conf_')[0]
            else:
                base_name = compound_name
            
            # Get SMILES from database
            smiles = self.smiles_db.get_smiles(base_name)
            
            if smiles:
                descriptors = self.calculate_molecular_descriptors(smiles)
                if descriptors:
                    descriptors['compound'] = compound_name
                    descriptors['smiles'] = smiles
                    features_list.append(descriptors)
        
        if not features_list:
            raise ValueError("No molecular features could be extracted")
        
        features_df = pd.DataFrame(features_list)
        logger.info(f"Extracted {len(features_df.columns)-2} descriptors for {len(features_df)} compounds")
        
        return features_df
```
