## Supplementary material for "Machine Learning-Driven Prediction of TLR4 Binding Affinity: A Comprehensive Molecular Feature Analysis for Drug Discovery": Codebase: feature_selection.py

```
#!/usr/bin/env python3
"""
Feature selection utilities for TLR4 binding prediction.

This module implements smart feature selection that balances performance
and interpretability while preventing overfitting.
"""

import pandas as pd
import numpy as np
import logging
from typing import Tuple, List
from sklearn.feature_selection import (
    VarianceThreshold, SelectKBest, f_regression, 
    SelectFromModel, RFE
)
from sklearn.ensemble import RandomForestRegressor

logger = logging.getLogger(__name__)


def smart_feature_selection(X: pd.DataFrame, y: pd.Series, target_features: int = 20) -> Tuple[pd.DataFrame, List[str]]:
    """Smart feature selection that balances performance and interpretability."""
    logger.info(f"Smart feature selection targeting {target_features} features")
    
    # Step 1: Remove low variance features
    variance_selector = VarianceThreshold(threshold=0.01)
    X_variance = pd.DataFrame(
        variance_selector.fit_transform(X),
        columns=X.columns[variance_selector.get_support()],
        index=X.index
    )
    logger.info(f"After variance threshold: {len(X_variance.columns)} features")
    
    # Step 2: Univariate selection
    n_univariate = min(40, len(X_variance.columns))
    univariate_selector = SelectKBest(score_func=f_regression, k=n_univariate)
    X_univariate = pd.DataFrame(
        univariate_selector.fit_transform(X_variance, y),
        columns=X_variance.columns[univariate_selector.get_support()],
        index=X_variance.index
    )
    logger.info(f"After univariate selection: {len(X_univariate.columns)} features")
    
    # Step 3: Model-based selection
    rf_selector = SelectFromModel(
        RandomForestRegressor(n_estimators=100, random_state=42),
        max_features=min(30, len(X_univariate.columns))
    )
    X_model = pd.DataFrame(
        rf_selector.fit_transform(X_univariate, y),
        columns=X_univariate.columns[rf_selector.get_support()],
        index=X_univariate.index
    )
    logger.info(f"After model-based selection: {len(X_model.columns)} features")
    
    # Step 4: Final RFE if needed
    if len(X_model.columns) > target_features:
        rfe_selector = RFE(
            RandomForestRegressor(n_estimators=100, random_state=42),
            n_features_to_select=target_features
        )
        X_selected = pd.DataFrame(
            rfe_selector.fit_transform(X_model, y),
            columns=X_model.columns[rfe_selector.get_support()],
            index=X_model.index
        )
        selected_features = X_model.columns[rfe_selector.get_support()].tolist()
    else:
        X_selected = X_model
        selected_features = X_model.columns.tolist()
    
    logger.info(f"Final selected features ({len(selected_features)}): {selected_features[:5]}...")
    return X_selected, selected_features
```
