## Supplementary material for "Machine Learning-Driven Prediction of TLR4 Binding Affinity: A Comprehensive Molecular Feature Analysis for Drug Discovery": Codebase: models.py

```
#!/usr/bin/env python3
"""
Machine learning models for TLR4 binding prediction.

This module implements ensemble learning approaches optimized for
small datasets with comprehensive validation.
"""

import pandas as pd
import numpy as np
import logging
from typing import Dict, List, Any, Tuple
from sklearn.model_selection import (
    train_test_split, cross_val_score, permutation_test_score
)
from sklearn.ensemble import RandomForestRegressor, VotingRegressor
from sklearn.linear_model import Ridge, ElasticNet, BayesianRidge
from sklearn.metrics import r2_score, mean_squared_error, mean_absolute_error
from sklearn.preprocessing import RobustScaler, PowerTransformer
from sklearn.pipeline import Pipeline

logger = logging.getLogger(__name__)


def create_ensemble_model() -> VotingRegressor:
    """Create a balanced ensemble for robust predictions."""
    
    models = [
        ('rf', RandomForestRegressor(
            n_estimators=300,
            max_depth=10,
            min_samples_split=3,
            min_samples_leaf=2,
            max_features='sqrt',
            random_state=42,
            n_jobs=-1
        )),
        
        ('elastic', ElasticNet(
            alpha=0.1,
            l1_ratio=0.5,
            random_state=42,
            max_iter=2000
        )),
        
        ('ridge', Ridge(
            alpha=1.0,
            random_state=42
        )),
        
        ('bayesian', BayesianRidge(
            alpha_1=1e-6,
            alpha_2=1e-6,
            lambda_1=1e-6,
            lambda_2=1e-6
        ))
    ]
    
    return VotingRegressor(models)


def train_binding_prediction_model(
    df: pd.DataFrame, 
    selected_features: List[str],
    test_size: float = 0.2, 
    random_state: int = 42
) -> Dict[str, Any]:
    """Train TLR4 binding prediction model with comprehensive validation."""
    logger.info("Training TLR4 binding prediction model")
    
    # Prepare data
    X = df[selected_features]
    y = df['affinity']
    
    logger.info(f"Dataset: {len(X)} samples, {len(selected_features)} features")
    logger.info(f"Samples per feature: {len(X) / len(selected_features):.1f}")
    
    # Split data
    X_train, X_test, y_train, y_test = train_test_split(
        X, y, test_size=test_size, random_state=random_state
    )
    
    logger.info(f"Train: {len(X_train)} samples, Test: {len(X_test)} samples")
    
    # Advanced scaling
    scaler = Pipeline([
        ('power', PowerTransformer(method='yeo-johnson')),
        ('robust', RobustScaler())
    ])
    
    X_train_scaled = pd.DataFrame(
        scaler.fit_transform(X_train),
        columns=X_train.columns,
        index=X_train.index
    )
    X_test_scaled = pd.DataFrame(
        scaler.transform(X_test),
        columns=X_test.columns,
        index=X_test.index
    )
    
    # Create and train ensemble
    ensemble = create_ensemble_model()
    
    logger.info("Training ensemble model...")
    ensemble.fit(X_train_scaled, y_train)
    
    # Predictions
    y_train_pred = ensemble.predict(X_train_scaled)
    y_test_pred = ensemble.predict(X_test_scaled)
    
    # Metrics
    train_r2 = r2_score(y_train, y_train_pred)
    test_r2 = r2_score(y_test, y_test_pred)
    train_rmse = np.sqrt(mean_squared_error(y_train, y_train_pred))
    test_rmse = np.sqrt(mean_squared_error(y_test, y_test_pred))
    train_mae = mean_absolute_error(y_train, y_train_pred)
    test_mae = mean_absolute_error(y_test, y_test_pred)
    
    # Cross-validation
    logger.info("Performing cross-validation...")
    cv_scores = cross_val_score(ensemble, X_train_scaled, y_train, cv=5, scoring='r2')
    
    # Permutation test for statistical significance
    perm_score, perm_scores, perm_pvalue = permutation_test_score(
        ensemble, X_train_scaled, y_train, scoring='r2', cv=5, n_permutations=100, random_state=42
    )
    
    # Feature importance
    rf_model = ensemble.named_estimators_['rf']
    feature_importance = pd.DataFrame({
        'feature': selected_features,
        'importance': rf_model.feature_importances_
    }).sort_values('importance', ascending=False)
    
    # Compile results
    metrics = {
        'train_r2': train_r2,
        'test_r2': test_r2,
        'train_rmse': train_rmse,
        'test_rmse': test_rmse,
        'train_mae': train_mae,
        'test_mae': test_mae,
        'cv_r2_mean': cv_scores.mean(),
        'cv_r2_std': cv_scores.std(),
        'permutation_pvalue': perm_pvalue,
        'overfitting_gap': train_r2 - test_r2
    }
    
    results = {
        'model': ensemble,
        'scaler': scaler,
        'selected_features': selected_features,
        'metrics': metrics,
        'feature_importance': feature_importance,
        'data_info': {
            'n_samples': len(df),
            'n_features': len(selected_features),
            'n_train': len(X_train),
            'n_test': len(X_test),
            'samples_per_feature': len(df) / len(selected_features)
        }
    }
    
    return results
```
