## Supplementary material for "Machine Learning-Driven Prediction of TLR4 Binding Affinity: A Comprehensive Molecular Feature Analysis for Drug Discovery": Codebase: pipeline.py

```
#!/usr/bin/env python3
"""
Main TLR4 binding prediction pipeline.

This module orchestrates the complete pipeline from feature extraction
to model training and evaluation.
"""

import pandas as pd
import numpy as np
import logging
import joblib
import json
from pathlib import Path
from datetime import datetime
from typing import Dict, Any

from ..molecular_analysis.smiles_database import get_smiles_database
from .feature_extraction import MolecularFeatureExtractor
from .data_processing import load_binding_data, integrate_features_and_binding, remove_true_duplicates
from .feature_selection import smart_feature_selection
from .models import train_binding_prediction_model

logger = logging.getLogger(__name__)


class TLR4BindingPredictor:
    """Complete TLR4 binding prediction pipeline."""
    
    def __init__(self):
        """Initialize the predictor."""
        self.smiles_db = get_smiles_database()
        self.feature_extractor = MolecularFeatureExtractor(self.smiles_db)
        self.model = None
        self.scaler = None
        self.selected_features = None
        self.results = None
    
    def train(
        self, 
        pdbqt_dir: str, 
        binding_csv: str, 
        target_features: int = 20,
        test_size: float = 0.2,
        random_state: int = 42
    ) -> Dict[str, Any]:
        """Train the complete TLR4 binding prediction model."""
        logger.info("=== TLR4 BINDING PREDICTION PIPELINE ===")
        
        try:
            # Step 1: Extract molecular features
            logger.info("Step 1: Extracting molecular features")
            features_df = self.feature_extractor.extract_features_from_files(pdbqt_dir)
            
            # Step 2: Load binding data
            logger.info("Step 2: Loading binding data")
            binding_df = load_binding_data(binding_csv)
            
            # Step 3: Integrate data
            logger.info("Step 3: Integrating features with binding data")
            integrated_df = integrate_features_and_binding(features_df, binding_df)
            
            # Step 4: Remove true duplicates
            logger.info("Step 4: Removing true duplicates")
            clean_df = remove_true_duplicates(integrated_df)
            
            # Step 5: Feature selection
            logger.info("Step 5: Performing feature selection")
            feature_cols = [col for col in clean_df.columns if col not in ['compound', 'affinity', 'smiles']]
            X = clean_df[feature_cols]
            y = clean_df['affinity']
            
            X_selected, selected_features = smart_feature_selection(X, y, target_features)
            
            # Add binding efficiency features
            if 'binding_efficiency' not in selected_features:
                X_selected['binding_efficiency'] = clean_df['binding_efficiency']
                selected_features.append('binding_efficiency')
            
            if 'ligand_efficiency' not in selected_features:
                X_selected['ligand_efficiency'] = clean_df['ligand_efficiency']
                selected_features.append('ligand_efficiency')
            
            # Prepare final dataset
            final_df = clean_df[['compound', 'affinity', 'smiles'] + selected_features].copy()
            
            # Step 6: Train model
            logger.info("Step 6: Training prediction model")
            results = train_binding_prediction_model(
                final_df, selected_features, test_size, random_state
            )
            
            # Store results
            self.model = results['model']
            self.scaler = results['scaler']
            self.selected_features = results['selected_features']
            self.results = results
            
            # Print results
            self._print_results(results)
            
            return results
            
        except Exception as e:
            logger.error(f"Pipeline failed: {e}")
            raise
    
    def predict(self, features: pd.DataFrame) -> np.ndarray:
        """Make predictions on new data."""
        if self.model is None:
            raise ValueError("Model not trained. Call train() first.")
        
        # Select features and scale
        X = features[self.selected_features]
        X_scaled = self.scaler.transform(X)
        
        # Predict
        predictions = self.model.predict(X_scaled)
        return predictions
    
    def save_model(self, output_dir: str = "results"):
        """Save the trained model and results."""
        if self.model is None:
            raise ValueError("No model to save. Train the model first.")
        
        output_path = Path(output_dir)
        output_path.mkdir(exist_ok=True)
        
        # Save model components
        joblib.dump(self.model, output_path / 'tlr4_model.pkl')
        joblib.dump(self.scaler, output_path / 'tlr4_scaler.pkl')
        
        # Save results
        save_data = {
            'timestamp': datetime.now().isoformat(),
            'selected_features': self.selected_features,
            'metrics': self.results['metrics'],
            'data_info': self.results['data_info'],
            'methodology': 'diversity_preserving_ensemble'
        }
        
        with open(output_path / 'tlr4_results.json', 'w') as f:
            json.dump(save_data, f, indent=2)
        
        # Save feature importance
        self.results['feature_importance'].to_csv(
            output_path / 'tlr4_feature_importance.csv', index=False
        )
        
        logger.info(f"Model saved to {output_path}")
    
    def load_model(self, model_dir: str = "results"):
        """Load a previously trained model."""
        model_path = Path(model_dir)
        
        # Load model components
        self.model = joblib.load(model_path / 'tlr4_model.pkl')
        self.scaler = joblib.load(model_path / 'tlr4_scaler.pkl')
        
        # Load results
        with open(model_path / 'tlr4_results.json', 'r') as f:
            results_data = json.load(f)
        
        self.selected_features = results_data['selected_features']
        
        logger.info(f"Model loaded from {model_path}")
    
    def _print_results(self, results: Dict[str, Any]):
        """Print training results."""
        logger.info("=" * 80)
        logger.info("TLR4 BINDING PREDICTION RESULTS")
        logger.info("=" * 80)
        logger.info(f"Training R²: {results['metrics']['train_r2']:.4f}")
        logger.info(f"Test R²: {results['metrics']['test_r2']:.4f}")
        logger.info(f"Cross-validation R²: {results['metrics']['cv_r2_mean']:.4f} ± {results['metrics']['cv_r2_std']:.4f}")
        logger.info(f"Overfitting gap: {results['metrics']['overfitting_gap']:.4f}")
        logger.info(f"Permutation p-value: {results['metrics']['permutation_pvalue']:.4f}")
        
        logger.info(f"\nDataset Info:")
        logger.info(f"  Samples: {results['data_info']['n_samples']}")
        logger.info(f"  Features: {results['data_info']['n_features']}")
        logger.info(f"  Samples per feature: {results['data_info']['samples_per_feature']:.1f}")
        
        logger.info(f"\nTop 10 Features:")
        for i, row in results['feature_importance'].head(10).iterrows():
            logger.info(f"  {i+1:2d}. {row['feature']}: {row['importance']:.4f}")
        
        # Assessment
        cv_r2 = results['metrics']['cv_r2_mean']
        if cv_r2 >= 0.6:
            logger.info(f"\n🎉 Excellent performance (CV R² = {cv_r2:.4f})")
        elif cv_r2 >= 0.5:
            logger.info(f"\n✅ Good performance (CV R² = {cv_r2:.4f})")
        elif cv_r2 >= 0.4:
            logger.info(f"\n✅ Acceptable performance (CV R² = {cv_r2:.4f})")
        else:
            logger.info(f"\n⚠️  Moderate performance (CV R² = {cv_r2:.4f})")


def run_tlr4_prediction_pipeline(
    pdbqt_dir: str = "data/raw/pdbqt",
    binding_csv: str = "data/processed/processed_logs.csv",
    output_dir: str = "results",
    target_features: int = 20
) -> Dict[str, Any]:
    """Run the complete TLR4 binding prediction pipeline."""
    
    # Setup logging
    logging.basicConfig(
        level=logging.INFO,
        format='%(asctime)s - %(name)s - %(levelname)s - %(message)s'
    )
    
    # Create and run predictor
    predictor = TLR4BindingPredictor()
    results = predictor.train(pdbqt_dir, binding_csv, target_features)
    
    # Save results
    predictor.save_model(output_dir)
    
    return results
```
