## Supplementary material for "Machine Learning-Driven Prediction of TLR4 Binding Affinity: A Comprehensive Molecular Feature Analysis for Drug Discovery": Codebase: __init__.py

```
"""
Utility functions and helper modules.

Contains utility functions for logging, file operations,
and other common tasks used throughout the TLR4 binding system.
"""

from .logging_config import setup_logging, get_logger
from .file_utils import ensure_directory, safe_file_operation
from .data_utils import validate_dataframe, clean_dataframe

__all__ = [
    "setup_logging",
    "get_logger", 
    "ensure_directory",
    "safe_file_operation",
    "validate_dataframe",
    "clean_dataframe"
]
```
