## Supplementary material for "Machine Learning-Driven Prediction of TLR4 Binding Affinity: A Comprehensive Molecular Feature Analysis for Drug Discovery": Codebase: data_quality.py

```
"""
Data quality validation and monitoring utilities.

This module provides comprehensive data quality assessment,
anomaly detection, and quality monitoring for the TLR4 binding pipeline.
"""

import logging
import numpy as np
import pandas as pd
from typing import Dict, List, Optional, Tuple, Any, Union
from pathlib import Path
from datetime import datetime
import json
from scipy import stats
from sklearn.ensemble import IsolationForest
from sklearn.preprocessing import StandardScaler
import warnings

from .error_handling import DataQualityError, PipelineError

logger = logging.getLogger(__name__)


class DataQualityValidator:
    """Comprehensive data quality validator for molecular data."""
    
    def __init__(self, config: Optional[Dict[str, Any]] = None):
        """
        Initialize data quality validator.
        
        Args:
            config: Configuration for validation rules
        """
        self.config = config or self._default_config()
        self.validation_history = []
        self.baseline_stats = None
        self.quality_thresholds = self._setup_quality_thresholds()
        
    def _default_config(self) -> Dict[str, Any]:
        """Get default validation configuration."""
        return {
            'enable_missing_value_checks': True,
            'enable_outlier_detection': True,
            'enable_data_type_validation': True,
            'enable_range_validation': True,
            'enable_distribution_validation': True,
            'enable_correlation_validation': True,
            'missing_value_threshold': 0.3,
            'outlier_threshold': 3.0,
            'correlation_threshold': 0.95,
            'strict_validation': False
        }
    
    def _setup_quality_thresholds(self) -> Dict[str, Any]:
        """Setup quality thresholds for molecular data."""
        return {
            'molecular_weight': {'min': 10, 'max': 2000},
            'logp': {'min': -10, 'max': 10},
            'tpsa': {'min': 0, 'max': 1000},
            'rotatable_bonds': {'min': 0, 'max': 50},
            'hbd': {'min': 0, 'max': 20},
            'hba': {'min': 0, 'max': 30},
            'molecular_volume': {'min': 0, 'max': 10000},
            'surface_area': {'min': 0, 'max': 5000},
            'radius_of_gyration': {'min': 0, 'max': 50},
            'asphericity': {'min': 0, 'max': 1}
        }
    
    def validate_dataset(self, data: pd.DataFrame, 
                        dataset_name: str = "dataset") -> Dict[str, Any]:
        """
        Perform comprehensive dataset validation.
        
        Args:
            data: Dataset to validate
            dataset_name: Name of dataset for logging
            
        Returns:
            Validation results dictionary
        """
        logger.info(f"Starting data quality validation for {dataset_name}")
        
        validation_results = {
            'dataset_name': dataset_name,
            'timestamp': datetime.now().isoformat(),
            'data_shape': data.shape,
            'validation_passed': True,
            'quality_score': 1.0,
            'issues': [],
            'warnings': [],
            'recommendations': []
        }
        
        try:
            # Basic data structure validation
            structure_results = self._validate_data_structure(data)
            validation_results.update(structure_results)
            
            # Missing value validation
            if self.config['enable_missing_value_checks']:
                missing_results = self._validate_missing_values(data)
                validation_results.update(missing_results)
            
            # Data type validation
            if self.config['enable_data_type_validation']:
                type_results = self._validate_data_types(data)
                validation_results.update(type_results)
            
            # Range validation
            if self.config['enable_range_validation']:
                range_results = self._validate_ranges(data)
                validation_results.update(range_results)
            
            # Outlier detection
            if self.config['enable_outlier_detection']:
                outlier_results = self._detect_outliers(data)
                validation_results.update(outlier_results)
            
            # Distribution validation
            if self.config['enable_distribution_validation']:
                distribution_results = self._validate_distributions(data)
                validation_results.update(distribution_results)
            
            # Correlation validation
            if self.config['enable_correlation_validation']:
                correlation_results = self._validate_correlations(data)
                validation_results.update(correlation_results)
            
            # Calculate overall quality score
            validation_results['quality_score'] = self._calculate_quality_score(validation_results)
            validation_results['validation_passed'] = validation_results['quality_score'] >= 0.7
            
            # Generate recommendations
            validation_results['recommendations'] = self._generate_recommendations(validation_results)
            
            # Store in history
            self.validation_history.append(validation_results)
            
            if not validation_results['validation_passed']:
                logger.warning(f"Data quality validation failed for {dataset_name}")
                logger.warning(f"Quality score: {validation_results['quality_score']:.2f}")
                logger.warning(f"Issues: {validation_results['issues']}")
            else:
                logger.info(f"Data quality validation passed for {dataset_name}")
                logger.info(f"Quality score: {validation_results['quality_score']:.2f}")
            
            return validation_results
            
        except Exception as e:
            error_msg = f"Data quality validation failed for {dataset_name}: {str(e)}"
            logger.error(error_msg)
            raise DataQualityError(error_msg, ['validation_error'])
    
    def _validate_data_structure(self, data: pd.DataFrame) -> Dict[str, Any]:
        """Validate basic data structure."""
        results = {
            'structure_issues': [],
            'structure_warnings': []
        }
        
        # Check for empty dataset
        if data.empty:
            results['structure_issues'].append("Dataset is empty")
            return results
        
        # Check for minimum number of samples
        if data.shape[0] < 10:
            results['structure_warnings'].append(f"Dataset has only {data.shape[0]} samples (minimum recommended: 10)")
        
        # Check for minimum number of features
        if data.shape[1] < 5:
            results['structure_warnings'].append(f"Dataset has only {data.shape[1]} features (minimum recommended: 5)")
        
        # Check for duplicate rows
        duplicate_count = data.duplicated().sum()
        if duplicate_count > 0:
            results['structure_issues'].append(f"Found {duplicate_count} duplicate rows")
        
        # Check for completely empty rows
        empty_rows = data.isnull().all(axis=1).sum()
        if empty_rows > 0:
            results['structure_issues'].append(f"Found {empty_rows} completely empty rows")
        
        return results
    
    def _validate_missing_values(self, data: pd.DataFrame) -> Dict[str, Any]:
        """Validate missing value patterns."""
        results = {
            'missing_value_issues': [],
            'missing_value_warnings': [],
            'missing_value_stats': {}
        }
        
        missing_counts = data.isnull().sum()
        missing_percentages = (missing_counts / len(data)) * 100
        
        # Check for high missing value rates
        high_missing_cols = missing_percentages[missing_percentages > self.config['missing_value_threshold'] * 100]
        if len(high_missing_cols) > 0:
            for col, pct in high_missing_cols.items():
                results['missing_value_issues'].append(f"Column '{col}' has {pct:.1f}% missing values")
        
        # Check for moderate missing value rates
        moderate_missing_cols = missing_percentages[
            (missing_percentages > 0.05 * 100) & 
            (missing_percentages <= self.config['missing_value_threshold'] * 100)
        ]
        if len(moderate_missing_cols) > 0:
            for col, pct in moderate_missing_cols.items():
                results['missing_value_warnings'].append(f"Column '{col}' has {pct:.1f}% missing values")
        
        # Store missing value statistics
        results['missing_value_stats'] = {
            'total_missing': missing_counts.sum(),
            'missing_percentage': (missing_counts.sum() / (len(data) * len(data.columns))) * 100,
            'columns_with_missing': missing_counts[missing_counts > 0].to_dict(),
            'missing_percentages': missing_percentages[missing_percentages > 0].to_dict()
        }
        
        return results
    
    def _validate_data_types(self, data: pd.DataFrame) -> Dict[str, Any]:
        """Validate data types and consistency."""
        results = {
            'data_type_issues': [],
            'data_type_warnings': [],
            'data_type_stats': {}
        }
        
        # Check for mixed data types in columns
        mixed_type_columns = []
        for column in data.columns:
            if data[column].dtype == 'object':
                non_null_values = data[column].dropna()
                if len(non_null_values) > 0:
                    type_counts = non_null_values.apply(type).value_counts()
                    if len(type_counts) > 1:
                        mixed_type_columns.append(column)
        
        if mixed_type_columns:
            results['data_type_issues'].append(f"Columns with mixed data types: {mixed_type_columns}")
        
        # Check for numeric columns that should be numeric
        expected_numeric_cols = ['molecular_weight', 'logp', 'tpsa', 'rotatable_bonds', 
                                'hbd', 'hba', 'molecular_volume', 'surface_area']
        for col in expected_numeric_cols:
            if col in data.columns and not pd.api.types.is_numeric_dtype(data[col]):
                results['data_type_issues'].append(f"Column '{col}' should be numeric but is {data[col].dtype}")
        
        # Store data type statistics
        results['data_type_stats'] = {
            'type_distribution': data.dtypes.value_counts().to_dict(),
            'numeric_columns': data.select_dtypes(include=[np.number]).columns.tolist(),
            'categorical_columns': data.select_dtypes(include=['object', 'category']).columns.tolist(),
            'mixed_type_columns': mixed_type_columns
        }
        
        return results
    
    def _validate_ranges(self, data: pd.DataFrame) -> Dict[str, Any]:
        """Validate value ranges for molecular properties."""
        results = {
            'range_issues': [],
            'range_warnings': [],
            'range_stats': {}
        }
        
        numeric_data = data.select_dtypes(include=[np.number])
        
        for column in numeric_data.columns:
            if column in self.quality_thresholds:
                thresholds = self.quality_thresholds[column]
                min_val = numeric_data[column].min()
                max_val = numeric_data[column].max()
                
                # Check for values outside expected ranges
                out_of_range_count = 0
                if min_val < thresholds['min']:
                    out_of_range_count += (numeric_data[column] < thresholds['min']).sum()
                if max_val > thresholds['max']:
                    out_of_range_count += (numeric_data[column] > thresholds['max']).sum()
                
                if out_of_range_count > 0:
                    if out_of_range_count > len(numeric_data) * 0.1:  # More than 10% out of range
                        results['range_issues'].append(f"Column '{column}' has {out_of_range_count} values outside expected range")
                    else:
                        results['range_warnings'].append(f"Column '{column}' has {out_of_range_count} values outside expected range")
                
                results['range_stats'][column] = {
                    'min_value': min_val,
                    'max_value': max_val,
                    'expected_min': thresholds['min'],
                    'expected_max': thresholds['max'],
                    'out_of_range_count': out_of_range_count
                }
        
        return results
    
    def _detect_outliers(self, data: pd.DataFrame) -> Dict[str, Any]:
        """Detect outliers using multiple methods."""
        results = {
            'outlier_issues': [],
            'outlier_warnings': [],
            'outlier_stats': {}
        }
        
        numeric_data = data.select_dtypes(include=[np.number])
        
        if numeric_data.empty:
            results['outlier_warnings'].append("No numeric columns for outlier detection")
            return results
        
        # IQR method
        iqr_outliers = self._detect_iqr_outliers(numeric_data)
        results['outlier_stats']['iqr_outliers'] = iqr_outliers
        
        # Z-score method
        zscore_outliers = self._detect_zscore_outliers(numeric_data)
        results['outlier_stats']['zscore_outliers'] = zscore_outliers
        
        # Isolation Forest method
        isolation_outliers = self._detect_isolation_forest_outliers(numeric_data)
        results['outlier_stats']['isolation_forest_outliers'] = isolation_outliers
        
        # Flag columns with high outlier rates
        for method, outliers in results['outlier_stats'].items():
            if isinstance(outliers, dict):
                for column, count in outliers.items():
                    outlier_rate = count / len(numeric_data)
                    if outlier_rate > 0.2:  # More than 20% outliers
                        results['outlier_issues'].append(f"Column '{column}' has {count} outliers ({outlier_rate:.1%}) using {method}")
                    elif outlier_rate > 0.1:  # More than 10% outliers
                        results['outlier_warnings'].append(f"Column '{column}' has {count} outliers ({outlier_rate:.1%}) using {method}")
        
        return results
    
    def _detect_iqr_outliers(self, data: pd.DataFrame) -> Dict[str, int]:
        """Detect outliers using IQR method."""
        outliers = {}
        
        for column in data.columns:
            Q1 = data[column].quantile(0.25)
            Q3 = data[column].quantile(0.75)
            IQR = Q3 - Q1
            lower_bound = Q1 - 1.5 * IQR
            upper_bound = Q3 + 1.5 * IQR
            outlier_count = ((data[column] < lower_bound) | (data[column] > upper_bound)).sum()
            outliers[column] = int(outlier_count)
        
        return outliers
    
    def _detect_zscore_outliers(self, data: pd.DataFrame) -> Dict[str, int]:
        """Detect outliers using Z-score method."""
        outliers = {}
        threshold = self.config['outlier_threshold']
        
        for column in data.columns:
            z_scores = np.abs(stats.zscore(data[column].dropna()))
            outlier_count = (z_scores > threshold).sum()
            outliers[column] = int(outlier_count)
        
        return outliers
    
    def _detect_isolation_forest_outliers(self, data: pd.DataFrame) -> Dict[str, int]:
        """Detect outliers using Isolation Forest."""
        try:
            # Fill missing values
            data_filled = data.fillna(data.median())
            
            # Fit Isolation Forest
            iso_forest = IsolationForest(contamination=0.1, random_state=42)
            outlier_labels = iso_forest.fit_predict(data_filled)
            
            # Count outliers per column
            outlier_counts = {}
            for i, column in enumerate(data.columns):
                column_outliers = (outlier_labels == -1).sum()
                outlier_counts[column] = int(column_outliers)
            
            return outlier_counts
            
        except Exception as e:
            logger.warning(f"Isolation Forest outlier detection failed: {str(e)}")
            return {column: 0 for column in data.columns}
    
    def _validate_distributions(self, data: pd.DataFrame) -> Dict[str, Any]:
        """Validate data distributions."""
        results = {
            'distribution_issues': [],
            'distribution_warnings': [],
            'distribution_stats': {}
        }
        
        numeric_data = data.select_dtypes(include=[np.number])
        
        if numeric_data.empty:
            results['distribution_warnings'].append("No numeric columns for distribution analysis")
            return results
        
        for column in numeric_data.columns:
            values = numeric_data[column].dropna()
            if len(values) > 0:
                # Calculate distribution statistics
                skewness = stats.skew(values)
                kurtosis = stats.kurtosis(values)
                
                # Check for normality if sample size allows
                if len(values) <= 5000:  # Shapiro-Wilk works best with smaller samples
                    shapiro_stat, shapiro_p = stats.shapiro(values)
                    is_normal = shapiro_p > 0.05
                else:
                    # For large samples, use Kolmogorov-Smirnov test
                    ks_stat, ks_p = stats.kstest(values, 'norm')
                    is_normal = ks_p > 0.05
                    shapiro_stat, shapiro_p = None, None
                
                results['distribution_stats'][column] = {
                    'skewness': skewness,
                    'kurtosis': kurtosis,
                    'is_normal': is_normal,
                    'shapiro_statistic': shapiro_stat,
                    'shapiro_p_value': shapiro_p
                }
                
                # Flag highly skewed distributions
                if abs(skewness) > 2:
                    results['distribution_warnings'].append(f"Column '{column}' is highly skewed (skewness: {skewness:.2f})")
                
                # Flag distributions with high kurtosis
                if abs(kurtosis) > 3:
                    results['distribution_warnings'].append(f"Column '{column}' has high kurtosis ({kurtosis:.2f})")
        
        return results
    
    def _validate_correlations(self, data: pd.DataFrame) -> Dict[str, Any]:
        """Validate correlation patterns."""
        results = {
            'correlation_issues': [],
            'correlation_warnings': [],
            'correlation_stats': {}
        }
        
        numeric_data = data.select_dtypes(include=[np.number])
        
        if numeric_data.empty or len(numeric_data.columns) < 2:
            results['correlation_warnings'].append("Not enough numeric columns for correlation analysis")
            return results
        
        corr_matrix = numeric_data.corr()
        
        # Find highly correlated pairs
        high_corr_pairs = []
        for i in range(len(corr_matrix.columns)):
            for j in range(i+1, len(corr_matrix.columns)):
                corr_val = corr_matrix.iloc[i, j]
                if abs(corr_val) > self.config['correlation_threshold']:
                    high_corr_pairs.append({
                        'feature_1': corr_matrix.columns[i],
                        'feature_2': corr_matrix.columns[j],
                        'correlation': corr_val
                    })
        
        if high_corr_pairs:
            results['correlation_issues'].append(f"Found {len(high_corr_pairs)} highly correlated feature pairs")
        
        results['correlation_stats'] = {
            'correlation_matrix': corr_matrix,
            'high_correlation_pairs': high_corr_pairs,
            'max_correlation': corr_matrix.abs().max().max(),
            'mean_correlation': corr_matrix.abs().mean().mean()
        }
        
        return results
    
    def _calculate_quality_score(self, validation_results: Dict[str, Any]) -> float:
        """Calculate overall data quality score."""
        score = 1.0
        
        # Penalize for issues
        issue_count = len(validation_results.get('issues', []))
        if issue_count > 0:
            score -= min(0.5, issue_count * 0.1)
        
        # Penalize for warnings
        warning_count = len(validation_results.get('warnings', []))
        if warning_count > 0:
            score -= min(0.2, warning_count * 0.02)
        
        # Penalize for high missing value rates
        missing_stats = validation_results.get('missing_value_stats', {})
        missing_pct = missing_stats.get('missing_percentage', 0)
        if missing_pct > 0:
            score -= min(0.3, missing_pct / 100)
        
        # Penalize for high outlier rates
        outlier_stats = validation_results.get('outlier_stats', {})
        if 'iqr_outliers' in outlier_stats:
            total_outliers = sum(outlier_stats['iqr_outliers'].values())
            data_shape = validation_results.get('data_shape', (0, 0))
            if isinstance(data_shape, tuple) and len(data_shape) >= 2:
                outlier_rate = total_outliers / (data_shape[0] * 10)
                score -= min(0.2, outlier_rate)
        
        return max(0.0, score)
    
    def _generate_recommendations(self, validation_results: Dict[str, Any]) -> List[str]:
        """Generate recommendations based on validation results."""
        recommendations = []
        
        # Missing value recommendations
        missing_stats = validation_results.get('missing_value_stats', {})
        if missing_stats.get('missing_percentage', 0) > 0.1:
            recommendations.append("Consider imputation strategies for missing values")
        
        # Outlier recommendations
        outlier_stats = validation_results.get('outlier_stats', {})
        if any(sum(outliers.values()) > 0 for outliers in outlier_stats.values() if isinstance(outliers, dict)):
            recommendations.append("Review and potentially remove or transform outliers")
        
        # Correlation recommendations
        correlation_stats = validation_results.get('correlation_stats', {})
        if correlation_stats.get('high_correlation_pairs'):
            recommendations.append("Consider removing highly correlated features to reduce redundancy")
        
        # Distribution recommendations
        distribution_stats = validation_results.get('distribution_stats', {})
        skewed_columns = [col for col, stats in distribution_stats.items() 
                         if abs(stats.get('skewness', 0)) > 2]
        if skewed_columns:
            recommendations.append(f"Consider log transformation for skewed columns: {skewed_columns}")
        
        return recommendations
    
    def establish_baseline(self, data: pd.DataFrame) -> Dict[str, Any]:
        """Establish quality baseline from training data."""
        logger.info("Establishing data quality baseline")
        
        baseline = {
            'timestamp': datetime.now().isoformat(),
            'data_shape': data.shape,
            'missing_rates': data.isnull().sum() / len(data),
            'numeric_stats': data.select_dtypes(include=[np.number]).describe(),
            'categorical_stats': data.select_dtypes(include=['object']).nunique(),
            'data_types': data.dtypes.to_dict(),
            'memory_usage': data.memory_usage(deep=True).sum(),
            'quality_thresholds': self.quality_thresholds
        }
        
        self.baseline_stats = baseline
        logger.info("Data quality baseline established")
        return baseline
    
    def compare_to_baseline(self, data: pd.DataFrame, 
                           data_name: str = "data") -> Dict[str, Any]:
        """Compare data quality to established baseline."""
        if not self.baseline_stats:
            raise ValueError("No baseline established. Call establish_baseline first.")
        
        logger.info(f"Comparing {data_name} to quality baseline")
        
        comparison_results = {
            'data_name': data_name,
            'timestamp': datetime.now().isoformat(),
            'baseline_comparison_passed': True,
            'comparison_issues': [],
            'comparison_warnings': []
        }
        
        # Compare shape
        baseline_shape = self.baseline_stats['data_shape']
        current_shape = data.shape
        
        if current_shape[1] != baseline_shape[1]:
            comparison_results['comparison_issues'].append(
                f"Feature count changed: baseline {baseline_shape[1]} -> current {current_shape[1]}"
            )
            comparison_results['baseline_comparison_passed'] = False
        
        # Compare missing value rates
        baseline_missing = self.baseline_stats['missing_rates']
        current_missing = data.isnull().sum() / len(data)
        
        for col in baseline_missing.index:
            if col in current_missing.index:
                missing_increase = current_missing[col] - baseline_missing[col]
                if missing_increase > 0.1:  # 10% increase in missing data
                    comparison_results['comparison_issues'].append(
                        f"Missing data increase in {col}: {missing_increase:.2%}"
                    )
                    comparison_results['baseline_comparison_passed'] = False
                elif missing_increase > 0.05:  # 5% increase
                    comparison_results['comparison_warnings'].append(
                        f"Missing data increase in {col}: {missing_increase:.2%}"
                    )
        
        return comparison_results


class DataAnomalyDetector:
    """Advanced anomaly detection for molecular data."""
    
    def __init__(self, config: Optional[Dict[str, Any]] = None):
        """
        Initialize anomaly detector.
        
        Args:
            config: Configuration for anomaly detection
        """
        self.config = config or self._default_config()
        self.detectors = {}
        self.baseline_data = None
        
    def _default_config(self) -> Dict[str, Any]:
        """Get default anomaly detection configuration."""
        return {
            'enable_statistical_detection': True,
            'enable_machine_learning_detection': True,
            'enable_temporal_detection': False,
            'contamination_rate': 0.1,
            'z_score_threshold': 3.0,
            'iqr_multiplier': 1.5
        }
    
    def fit_baseline(self, data: pd.DataFrame) -> None:
        """Fit anomaly detection models on baseline data."""
        logger.info("Fitting anomaly detection models on baseline data")
        
        self.baseline_data = data.copy()
        numeric_data = data.select_dtypes(include=[np.number])
        
        if numeric_data.empty:
            logger.warning("No numeric columns for anomaly detection")
            return
        
        # Statistical detectors
        if self.config['enable_statistical_detection']:
            self._fit_statistical_detectors(numeric_data)
        
        # Machine learning detectors
        if self.config['enable_machine_learning_detection']:
            self._fit_ml_detectors(numeric_data)
        
        logger.info("Anomaly detection models fitted successfully")
    
    def _fit_statistical_detectors(self, data: pd.DataFrame) -> None:
        """Fit statistical anomaly detectors."""
        # Z-score detector
        self.detectors['zscore'] = {
            'mean': data.mean(),
            'std': data.std(),
            'threshold': self.config['z_score_threshold']
        }
        
        # IQR detector
        Q1 = data.quantile(0.25)
        Q3 = data.quantile(0.75)
        IQR = Q3 - Q1
        multiplier = self.config['iqr_multiplier']
        
        self.detectors['iqr'] = {
            'Q1': Q1,
            'Q3': Q3,
            'IQR': IQR,
            'lower_bound': Q1 - multiplier * IQR,
            'upper_bound': Q3 + multiplier * IQR
        }
    
    def _fit_ml_detectors(self, data: pd.DataFrame) -> None:
        """Fit machine learning anomaly detectors."""
        try:
            # Isolation Forest
            iso_forest = IsolationForest(
                contamination=self.config['contamination_rate'],
                random_state=42
            )
            iso_forest.fit(data.fillna(data.median()))
            self.detectors['isolation_forest'] = iso_forest
            
        except Exception as e:
            logger.warning(f"Failed to fit Isolation Forest: {e}")
    
    def detect_anomalies(self, data: pd.DataFrame) -> Dict[str, Any]:
        """Detect anomalies in new data."""
        if self.baseline_data is None:
            raise ValueError("No baseline data fitted. Call fit_baseline first.")
        
        logger.info("Detecting anomalies in new data")
        
        numeric_data = data.select_dtypes(include=[np.number])
        
        if numeric_data.empty:
            return {'anomalies': {}, 'anomaly_count': 0}
        
        anomaly_results = {
            'anomaly_detection_methods': {},
            'total_anomalies': 0,
            'anomaly_indices': set(),
            'column_anomaly_counts': {}
        }
        
        # Statistical anomaly detection
        if self.config['enable_statistical_detection']:
            anomaly_results['anomaly_detection_methods'].update(
                self._detect_statistical_anomalies(numeric_data)
            )
        
        # Machine learning anomaly detection
        if self.config['enable_machine_learning_detection']:
            anomaly_results['anomaly_detection_methods'].update(
                self._detect_ml_anomalies(numeric_data)
            )
        
        # Aggregate results
        for method, anomalies in anomaly_results['anomaly_detection_methods'].items():
            if isinstance(anomalies, dict) and 'anomaly_indices' in anomalies:
                anomaly_results['anomaly_indices'].update(anomalies['anomaly_indices'])
        
        anomaly_results['total_anomalies'] = len(anomaly_results['anomaly_indices'])
        anomaly_results['anomaly_indices'] = list(anomaly_results['anomaly_indices'])
        
        # Count anomalies per column
        for column in numeric_data.columns:
            column_anomalies = 0
            for method, results in anomaly_results['anomaly_detection_methods'].items():
                if isinstance(results, dict) and column in results:
                    column_anomalies += results[column]
            anomaly_results['column_anomaly_counts'][column] = column_anomalies
        
        logger.info(f"Detected {anomaly_results['total_anomalies']} anomalies")
        return anomaly_results
    
    def _detect_statistical_anomalies(self, data: pd.DataFrame) -> Dict[str, Any]:
        """Detect anomalies using statistical methods."""
        results = {}
        
        # Z-score anomalies
        if 'zscore' in self.detectors:
            zscore_results = {'anomaly_indices': set()}
            detector = self.detectors['zscore']
            
            for column in data.columns:
                if column in detector['mean'] and detector['std'][column] > 0:
                    z_scores = np.abs((data[column] - detector['mean'][column]) / detector['std'][column])
                    anomalies = data.index[z_scores > detector['threshold']]
                    zscore_results[column] = len(anomalies)
                    zscore_results['anomaly_indices'].update(anomalies)
            
            results['zscore'] = zscore_results
        
        # IQR anomalies
        if 'iqr' in self.detectors:
            iqr_results = {'anomaly_indices': set()}
            detector = self.detectors['iqr']
            
            for column in data.columns:
                if column in detector['lower_bound']:
                    anomalies = data.index[
                        (data[column] < detector['lower_bound'][column]) |
                        (data[column] > detector['upper_bound'][column])
                    ]
                    iqr_results[column] = len(anomalies)
                    iqr_results['anomaly_indices'].update(anomalies)
            
            results['iqr'] = iqr_results
        
        return results
    
    def _detect_ml_anomalies(self, data: pd.DataFrame) -> Dict[str, Any]:
        """Detect anomalies using machine learning methods."""
        results = {}
        
        # Isolation Forest anomalies
        if 'isolation_forest' in self.detectors:
            try:
                iso_forest = self.detectors['isolation_forest']
                data_filled = data.fillna(data.median())
                anomaly_labels = iso_forest.predict(data_filled)
                anomaly_indices = data_filled.index[anomaly_labels == -1]
                
                results['isolation_forest'] = {
                    'anomaly_indices': set(anomaly_indices),
                    'total_anomalies': len(anomaly_indices)
                }
                
            except Exception as e:
                logger.warning(f"Isolation Forest anomaly detection failed: {e}")
                results['isolation_forest'] = {
                    'anomaly_indices': set(),
                    'total_anomalies': 0,
                    'error': str(e)
                }
        
        return results


def validate_molecular_data(data: pd.DataFrame, 
                          config: Optional[Dict[str, Any]] = None) -> Dict[str, Any]:
    """
    Convenience function for comprehensive molecular data validation.
    
    Args:
        data: Dataset to validate
        config: Validation configuration
        
    Returns:
        Validation results
    """
    validator = DataQualityValidator(config)
    return validator.validate_dataset(data)
```
