## Supplementary material for "Machine Learning-Driven Prediction of TLR4 Binding Affinity: A Comprehensive Molecular Feature Analysis for Drug Discovery": Codebase: data_utils.py

```
"""
Data manipulation and validation utilities.

This module provides common data processing functions
for the TLR4 binding prediction system.
"""

import pandas as pd
import numpy as np
from typing import Union, List, Dict, Any, Optional, Tuple
from pathlib import Path
import logging

logger = logging.getLogger(__name__)


def validate_dataframe(df: pd.DataFrame, 
                      required_columns: Optional[List[str]] = None,
                      numeric_columns: Optional[List[str]] = None) -> Dict[str, Any]:
    """
    Validate DataFrame structure and content.
    
    Args:
        df: DataFrame to validate
        required_columns: List of required column names
        numeric_columns: List of columns that should be numeric
        
    Returns:
        Dictionary with validation results
    """
    validation_results = {
        'is_valid': True,
        'errors': [],
        'warnings': [],
        'info': {}
    }
    
    # Check if DataFrame is empty
    if df.empty:
        validation_results['errors'].append("DataFrame is empty")
        validation_results['is_valid'] = False
        return validation_results
    
    # Check required columns
    if required_columns:
        missing_columns = set(required_columns) - set(df.columns)
        if missing_columns:
            validation_results['errors'].append(f"Missing required columns: {missing_columns}")
            validation_results['is_valid'] = False
    
    # Check numeric columns
    if numeric_columns:
        non_numeric_columns = []
        for col in numeric_columns:
            if col in df.columns:
                if not pd.api.types.is_numeric_dtype(df[col]):
                    non_numeric_columns.append(col)
        
        if non_numeric_columns:
            validation_results['warnings'].append(f"Non-numeric columns: {non_numeric_columns}")
    
    # Basic statistics
    validation_results['info'] = {
        'shape': df.shape,
        'columns': list(df.columns),
        'dtypes': df.dtypes.to_dict(),
        'memory_usage': df.memory_usage(deep=True).sum(),
        'null_counts': df.isnull().sum().to_dict()
    }
    
    return validation_results


def clean_dataframe(df: pd.DataFrame,
                   remove_duplicates: bool = True,
                   handle_missing: str = 'drop',  # 'drop', 'fill', 'interpolate'
                   fill_value: Any = None,
                   remove_outliers: bool = False,
                   outlier_method: str = 'iqr') -> pd.DataFrame:
    """
    Clean DataFrame by removing duplicates, handling missing values, etc.
    
    Args:
        df: DataFrame to clean
        remove_duplicates: Whether to remove duplicate rows
        handle_missing: Strategy for handling missing values
        fill_value: Value to use for filling missing values
        remove_outliers: Whether to remove outliers
        outlier_method: Method for outlier detection ('iqr', 'zscore')
        
    Returns:
        Cleaned DataFrame
    """
    df_cleaned = df.copy()
    
    # Remove duplicates
    if remove_duplicates:
        initial_count = len(df_cleaned)
        df_cleaned = df_cleaned.drop_duplicates()
        removed_count = initial_count - len(df_cleaned)
        if removed_count > 0:
            logger.info(f"Removed {removed_count} duplicate rows")
    
    # Handle missing values
    if handle_missing == 'drop':
        initial_count = len(df_cleaned)
        df_cleaned = df_cleaned.dropna()
        removed_count = initial_count - len(df_cleaned)
        if removed_count > 0:
            logger.info(f"Removed {removed_count} rows with missing values")
    
    elif handle_missing == 'fill':
        if fill_value is not None:
            df_cleaned = df_cleaned.fillna(fill_value)
        else:
            # Fill numeric columns with median, categorical with mode
            for col in df_cleaned.columns:
                if pd.api.types.is_numeric_dtype(df_cleaned[col]):
                    df_cleaned[col] = df_cleaned[col].fillna(df_cleaned[col].median())
                else:
                    df_cleaned[col] = df_cleaned[col].fillna(df_cleaned[col].mode().iloc[0] if not df_cleaned[col].mode().empty else 'Unknown')
    
    elif handle_missing == 'interpolate':
        numeric_columns = df_cleaned.select_dtypes(include=[np.number]).columns
        df_cleaned[numeric_columns] = df_cleaned[numeric_columns].interpolate()
    
    # Remove outliers
    if remove_outliers:
        numeric_columns = df_cleaned.select_dtypes(include=[np.number]).columns
        initial_count = len(df_cleaned)
        
        for col in numeric_columns:
            if outlier_method == 'iqr':
                Q1 = df_cleaned[col].quantile(0.25)
                Q3 = df_cleaned[col].quantile(0.75)
                IQR = Q3 - Q1
                lower_bound = Q1 - 1.5 * IQR
                upper_bound = Q3 + 1.5 * IQR
                df_cleaned = df_cleaned[(df_cleaned[col] >= lower_bound) & (df_cleaned[col] <= upper_bound)]
            
            elif outlier_method == 'zscore':
                z_scores = np.abs((df_cleaned[col] - df_cleaned[col].mean()) / df_cleaned[col].std())
                df_cleaned = df_cleaned[z_scores < 3]
        
        removed_count = initial_count - len(df_cleaned)
        if removed_count > 0:
            logger.info(f"Removed {removed_count} outlier rows")
    
    return df_cleaned


def normalize_dataframe(df: pd.DataFrame,
                       method: str = 'standard',
                       columns: Optional[List[str]] = None) -> pd.DataFrame:
    """
    Normalize DataFrame columns.
    
    Args:
        df: DataFrame to normalize
        method: Normalization method ('standard', 'minmax', 'robust')
        columns: Columns to normalize (None for all numeric columns)
        
    Returns:
        Normalized DataFrame
    """
    df_normalized = df.copy()
    
    if columns is None:
        columns = df_normalized.select_dtypes(include=[np.number]).columns.tolist()
    
    for col in columns:
        if col in df_normalized.columns and pd.api.types.is_numeric_dtype(df_normalized[col]):
            if method == 'standard':
                # Z-score normalization
                df_normalized[col] = (df_normalized[col] - df_normalized[col].mean()) / df_normalized[col].std()
            elif method == 'minmax':
                # Min-max normalization
                min_val = df_normalized[col].min()
                max_val = df_normalized[col].max()
                if max_val != min_val:
                    df_normalized[col] = (df_normalized[col] - min_val) / (max_val - min_val)
            elif method == 'robust':
                # Robust normalization using median and IQR
                median_val = df_normalized[col].median()
                q75 = df_normalized[col].quantile(0.75)
                q25 = df_normalized[col].quantile(0.25)
                iqr = q75 - q25
                if iqr != 0:
                    df_normalized[col] = (df_normalized[col] - median_val) / iqr
    
    return df_normalized


def split_dataframe(df: pd.DataFrame,
                   train_ratio: float = 0.7,
                   val_ratio: float = 0.15,
                   test_ratio: float = 0.15,
                   random_state: int = 42,
                   stratify_column: Optional[str] = None) -> Tuple[pd.DataFrame, pd.DataFrame, pd.DataFrame]:
    """
    Split DataFrame into train/validation/test sets.
    
    Args:
        df: DataFrame to split
        train_ratio: Ratio for training set
        val_ratio: Ratio for validation set
        test_ratio: Ratio for test set
        random_state: Random state for reproducibility
        stratify_column: Column to use for stratified splitting
        
    Returns:
        Tuple of (train_df, val_df, test_df)
    """
    # Validate ratios
    total_ratio = train_ratio + val_ratio + test_ratio
    if abs(total_ratio - 1.0) > 1e-6:
        raise ValueError(f"Ratios must sum to 1.0, got {total_ratio}")
    
    # Shuffle DataFrame
    df_shuffled = df.sample(frac=1, random_state=random_state).reset_index(drop=True)
    
    # Calculate split indices
    n_samples = len(df_shuffled)
    train_end = int(n_samples * train_ratio)
    val_end = train_end + int(n_samples * val_ratio)
    
    # Split DataFrame
    train_df = df_shuffled[:train_end]
    val_df = df_shuffled[train_end:val_end]
    test_df = df_shuffled[val_end:]
    
    logger.info(f"Data split: Train={len(train_df)}, Val={len(val_df)}, Test={len(test_df)}")
    
    return train_df, val_df, test_df


def get_dataframe_summary(df: pd.DataFrame) -> Dict[str, Any]:
    """
    Get comprehensive summary of DataFrame.
    
    Args:
        df: DataFrame to summarize
        
    Returns:
        Dictionary with summary statistics
    """
    summary = {
        'shape': df.shape,
        'columns': list(df.columns),
        'dtypes': df.dtypes.to_dict(),
        'memory_usage': df.memory_usage(deep=True).sum(),
        'null_counts': df.isnull().sum().to_dict(),
        'duplicate_rows': df.duplicated().sum(),
        'numeric_columns': df.select_dtypes(include=[np.number]).columns.tolist(),
        'categorical_columns': df.select_dtypes(include=['object', 'category']).columns.tolist()
    }
    
    # Add descriptive statistics for numeric columns
    numeric_cols = df.select_dtypes(include=[np.number]).columns
    if len(numeric_cols) > 0:
        summary['numeric_stats'] = df[numeric_cols].describe().to_dict()
    
    # Add value counts for categorical columns
    categorical_cols = df.select_dtypes(include=['object', 'category']).columns
    if len(categorical_cols) > 0:
        summary['categorical_stats'] = {}
        for col in categorical_cols:
            summary['categorical_stats'][col] = df[col].value_counts().to_dict()
    
    return summary


def merge_dataframes_safely(df1: pd.DataFrame, 
                           df2: pd.DataFrame,
                           on: Union[str, List[str]],
                           how: str = 'inner',
                           suffixes: Tuple[str, str] = ('_x', '_y')) -> pd.DataFrame:
    """
    Safely merge two DataFrames with error handling.
    
    Args:
        df1: First DataFrame
        df2: Second DataFrame
        on: Column(s) to merge on
        how: Type of merge ('inner', 'outer', 'left', 'right')
        suffixes: Suffixes for overlapping columns
        
    Returns:
        Merged DataFrame
    """
    try:
        # Check if merge columns exist in both DataFrames
        if isinstance(on, str):
            on = [on]
        
        missing_cols_df1 = set(on) - set(df1.columns)
        missing_cols_df2 = set(on) - set(df2.columns)
        
        if missing_cols_df1:
            raise ValueError(f"Columns {missing_cols_df1} not found in first DataFrame")
        if missing_cols_df2:
            raise ValueError(f"Columns {missing_cols_df2} not found in second DataFrame")
        
        # Perform merge
        merged_df = pd.merge(df1, df2, on=on, how=how, suffixes=suffixes)
        
        logger.info(f"Successfully merged DataFrames: {df1.shape} + {df2.shape} -> {merged_df.shape}")
        return merged_df
        
    except Exception as e:
        logger.error(f"Failed to merge DataFrames: {e}")
        raise


def save_dataframe_safely(df: pd.DataFrame, 
                         file_path: Union[str, Path],
                         format: str = 'csv',
                         **kwargs) -> bool:
    """
    Safely save DataFrame to file.
    
    Args:
        df: DataFrame to save
        file_path: Path to save file
        format: File format ('csv', 'parquet', 'excel', 'json')
        **kwargs: Additional arguments for the save method
        
    Returns:
        True if successful, False otherwise
    """
    try:
        file_path = Path(file_path)
        file_path.parent.mkdir(parents=True, exist_ok=True)
        
        if format.lower() == 'csv':
            df.to_csv(file_path, index=False, **kwargs)
        elif format.lower() == 'parquet':
            df.to_parquet(file_path, index=False, **kwargs)
        elif format.lower() == 'excel':
            df.to_excel(file_path, index=False, **kwargs)
        elif format.lower() == 'json':
            df.to_json(file_path, orient='records', **kwargs)
        else:
            raise ValueError(f"Unsupported format: {format}")
        
        logger.info(f"DataFrame saved to {file_path}")
        return True
        
    except Exception as e:
        logger.error(f"Failed to save DataFrame: {e}")
        return False
```
