## Supplementary material for "Machine Learning-Driven Prediction of TLR4 Binding Affinity: A Comprehensive Molecular Feature Analysis for Drug Discovery": Codebase: error_handling.py

```
"""
Comprehensive error handling and robustness utilities.

This module provides centralized error handling, recovery mechanisms,
and robustness features for the TLR4 binding prediction pipeline.
"""

import logging
import traceback
import functools
import time
import json
from typing import Any, Dict, List, Optional, Union, Callable, Type, Tuple
from pathlib import Path
from datetime import datetime
import pandas as pd
import numpy as np
from contextlib import contextmanager
import warnings

logger = logging.getLogger(__name__)


class PipelineError(Exception):
    """Base exception for pipeline-specific errors."""
    
    def __init__(self, message: str, error_code: str = "PIPELINE_ERROR", 
                 context: Optional[Dict[str, Any]] = None):
        super().__init__(message)
        self.message = message
        self.error_code = error_code
        self.context = context or {}
        self.timestamp = datetime.now()
    
    def to_dict(self) -> Dict[str, Any]:
        """Convert error to dictionary for logging."""
        return {
            'error_type': self.__class__.__name__,
            'error_code': self.error_code,
            'message': self.message,
            'context': self.context,
            'timestamp': self.timestamp.isoformat()
        }


class DataQualityError(PipelineError):
    """Raised when data quality issues are detected."""
    
    def __init__(self, message: str, quality_issues: List[str], 
                 context: Optional[Dict[str, Any]] = None):
        super().__init__(message, "DATA_QUALITY_ERROR", context)
        self.quality_issues = quality_issues


class ModelTrainingError(PipelineError):
    """Raised when model training fails."""
    
    def __init__(self, message: str, model_name: str, 
                 context: Optional[Dict[str, Any]] = None):
        super().__init__(message, "MODEL_TRAINING_ERROR", context)
        self.model_name = model_name


class FeatureExtractionError(PipelineError):
    """Raised when feature extraction fails."""
    
    def __init__(self, message: str, compound_name: Optional[str] = None,
                 context: Optional[Dict[str, Any]] = None):
        super().__init__(message, "FEATURE_EXTRACTION_ERROR", context)
        self.compound_name = compound_name


class RobustnessManager:
    """Manages robustness features and error recovery."""
    
    def __init__(self, config: Optional[Dict[str, Any]] = None):
        """
        Initialize robustness manager.
        
        Args:
            config: Configuration dictionary for robustness settings
        """
        self.config = config or self._default_config()
        self.error_log = []
        self.recovery_attempts = {}
        self.checkpoint_manager = CheckpointManager()
        
    def _default_config(self) -> Dict[str, Any]:
        """Get default robustness configuration."""
        return {
            'max_retry_attempts': 3,
            'retry_delay': 1.0,
            'enable_graceful_degradation': True,
            'enable_checkpointing': True,
            'checkpoint_interval': 100,
            'log_level': 'INFO',
            'enable_circuit_breaker': True,
            'circuit_breaker_threshold': 5,
            'circuit_breaker_timeout': 300,
            'enable_data_validation': True,
            'enable_outlier_detection': True,
            'outlier_threshold': 3.0
        }
    
    def log_error(self, error: Exception, context: Optional[Dict[str, Any]] = None):
        """Log error with context information."""
        error_entry = {
            'timestamp': datetime.now().isoformat(),
            'error_type': type(error).__name__,
            'error_message': str(error),
            'traceback': traceback.format_exc(),
            'context': context or {}
        }
        
        self.error_log.append(error_entry)
        logger.error(f"Error logged: {error_entry['error_type']} - {error_entry['error_message']}")
        
        # Store in file if configured
        if self.config.get('enable_error_logging', True):
            self._write_error_log(error_entry)
    
    def _write_error_log(self, error_entry: Dict[str, Any]):
        """Write error to log file."""
        try:
            log_dir = Path("logs/errors")
            log_dir.mkdir(parents=True, exist_ok=True)
            
            log_file = log_dir / f"error_log_{datetime.now().strftime('%Y%m%d')}.jsonl"
            
            with open(log_file, 'a') as f:
                f.write(json.dumps(error_entry) + '\n')
                
        except Exception as e:
            logger.warning(f"Failed to write error log: {e}")
    
    def should_retry(self, error: Exception, attempt_count: int) -> bool:
        """Determine if operation should be retried."""
        if attempt_count >= self.config['max_retry_attempts']:
            return False
        
        # Don't retry certain types of errors
        non_retryable_errors = (
            ValueError, TypeError, AttributeError, KeyError,
            FileNotFoundError, PermissionError
        )
        
        if isinstance(error, non_retryable_errors):
            return False
        
        return True
    
    def get_retry_delay(self, attempt_count: int) -> float:
        """Get delay before retry (exponential backoff)."""
        base_delay = self.config['retry_delay']
        return base_delay * (2 ** attempt_count)


class CheckpointManager:
    """Manages checkpointing and resume functionality."""
    
    def __init__(self, checkpoint_dir: str = "checkpoints"):
        """
        Initialize checkpoint manager.
        
        Args:
            checkpoint_dir: Directory for storing checkpoints
        """
        self.checkpoint_dir = Path(checkpoint_dir)
        self.checkpoint_dir.mkdir(parents=True, exist_ok=True)
        
    def save_checkpoint(self, checkpoint_id: str, data: Dict[str, Any]) -> bool:
        """
        Save checkpoint data.
        
        Args:
            checkpoint_id: Unique identifier for checkpoint
            data: Data to checkpoint
            
        Returns:
            True if successful, False otherwise
        """
        try:
            checkpoint_file = self.checkpoint_dir / f"{checkpoint_id}.json"
            
            checkpoint_data = {
                'timestamp': datetime.now().isoformat(),
                'data': data
            }
            
            with open(checkpoint_file, 'w') as f:
                json.dump(checkpoint_data, f, indent=2, default=str)
            
            logger.info(f"Checkpoint saved: {checkpoint_id}")
            return True
            
        except Exception as e:
            logger.error(f"Failed to save checkpoint {checkpoint_id}: {e}")
            return False
    
    def load_checkpoint(self, checkpoint_id: str) -> Optional[Dict[str, Any]]:
        """
        Load checkpoint data.
        
        Args:
            checkpoint_id: Unique identifier for checkpoint
            
        Returns:
            Checkpoint data or None if not found
        """
        try:
            checkpoint_file = self.checkpoint_dir / f"{checkpoint_id}.json"
            
            if not checkpoint_file.exists():
                return None
            
            with open(checkpoint_file, 'r') as f:
                checkpoint_data = json.load(f)
            
            logger.info(f"Checkpoint loaded: {checkpoint_id}")
            return checkpoint_data['data']
            
        except Exception as e:
            logger.error(f"Failed to load checkpoint {checkpoint_id}: {e}")
            return None
    
    def list_checkpoints(self) -> List[str]:
        """List available checkpoints."""
        try:
            checkpoint_files = list(self.checkpoint_dir.glob("*.json"))
            return [f.stem for f in checkpoint_files]
        except Exception as e:
            logger.error(f"Failed to list checkpoints: {e}")
            return []
    
    def delete_checkpoint(self, checkpoint_id: str) -> bool:
        """Delete checkpoint."""
        try:
            checkpoint_file = self.checkpoint_dir / f"{checkpoint_id}.json"
            if checkpoint_file.exists():
                checkpoint_file.unlink()
                logger.info(f"Checkpoint deleted: {checkpoint_id}")
                return True
            return False
        except Exception as e:
            logger.error(f"Failed to delete checkpoint {checkpoint_id}: {e}")
            return False


class CircuitBreaker:
    """Circuit breaker pattern for preventing cascading failures."""
    
    def __init__(self, failure_threshold: int = 5, timeout: int = 300):
        """
        Initialize circuit breaker.
        
        Args:
            failure_threshold: Number of failures before opening circuit
            timeout: Timeout in seconds before attempting to close circuit
        """
        self.failure_threshold = failure_threshold
        self.timeout = timeout
        self.failure_count = 0
        self.last_failure_time = None
        self.state = 'CLOSED'  # CLOSED, OPEN, HALF_OPEN
        
    def call(self, func: Callable, *args, **kwargs) -> Any:
        """
        Execute function with circuit breaker protection.
        
        Args:
            func: Function to execute
            *args: Function arguments
            **kwargs: Function keyword arguments
            
        Returns:
            Function result
            
        Raises:
            Exception: If circuit is open or function fails
        """
        if self.state == 'OPEN':
            if self._should_attempt_reset():
                self.state = 'HALF_OPEN'
                logger.info("Circuit breaker entering HALF_OPEN state")
            else:
                raise Exception("Circuit breaker is OPEN - operation blocked")
        
        try:
            result = func(*args, **kwargs)
            self._on_success()
            return result
            
        except Exception as e:
            self._on_failure()
            raise e
    
    def _should_attempt_reset(self) -> bool:
        """Check if circuit breaker should attempt reset."""
        if self.last_failure_time is None:
            return True
        
        time_since_failure = time.time() - self.last_failure_time
        return time_since_failure >= self.timeout
    
    def _on_success(self):
        """Handle successful operation."""
        self.failure_count = 0
        self.state = 'CLOSED'
    
    def _on_failure(self):
        """Handle failed operation."""
        self.failure_count += 1
        self.last_failure_time = time.time()
        
        if self.failure_count >= self.failure_threshold:
            self.state = 'OPEN'
            logger.warning(f"Circuit breaker opened after {self.failure_count} failures")


def robust_execution(max_retries: int = 3, 
                    delay: float = 1.0,
                    exceptions: Tuple[Type[Exception], ...] = (Exception,),
                    circuit_breaker: Optional[CircuitBreaker] = None):
    """
    Decorator for robust execution with retry logic.
    
    Args:
        max_retries: Maximum number of retry attempts
        delay: Initial delay between retries (exponential backoff)
        exceptions: Tuple of exception types to retry on
        circuit_breaker: Optional circuit breaker instance
    """
    def decorator(func: Callable) -> Callable:
        @functools.wraps(func)
        def wrapper(*args, **kwargs):
            last_exception = None
            
            for attempt in range(max_retries + 1):
                try:
                    if circuit_breaker:
                        return circuit_breaker.call(func, *args, **kwargs)
                    else:
                        return func(*args, **kwargs)
                        
                except exceptions as e:
                    last_exception = e
                    
                    if attempt < max_retries:
                        retry_delay = delay * (2 ** attempt)
                        logger.warning(
                            f"Attempt {attempt + 1} failed for {func.__name__}: {e}. "
                            f"Retrying in {retry_delay}s..."
                        )
                        time.sleep(retry_delay)
                    else:
                        logger.error(f"All {max_retries + 1} attempts failed for {func.__name__}")
            
            # If we get here, all retries failed
            raise last_exception
            
        return wrapper
    return decorator


@contextmanager
def safe_execution(context: str, 
                  error_handler: Optional[Callable] = None,
                  default_return: Any = None):
    """
    Context manager for safe execution with error handling.
    
    Args:
        context: Context description for logging
        error_handler: Optional error handler function
        default_return: Default return value if execution fails
    """
    try:
        logger.info(f"Starting safe execution: {context}")
        yield
        logger.info(f"Completed safe execution: {context}")
        
    except Exception as e:
        logger.error(f"Error in {context}: {e}")
        logger.error(traceback.format_exc())
        
        if error_handler:
            error_handler(e)
        
        if default_return is not None:
            return default_return
        
        raise


class DataQualityMonitor:
    """Monitors data quality and detects anomalies."""
    
    def __init__(self, config: Optional[Dict[str, Any]] = None):
        """
        Initialize data quality monitor.
        
        Args:
            config: Configuration for quality monitoring
        """
        self.config = config or {}
        self.baseline_stats = {}
        self.quality_history = []
        
    def establish_baseline(self, data: pd.DataFrame) -> Dict[str, Any]:
        """
        Establish quality baseline from training data.
        
        Args:
            data: Training data for baseline
            
        Returns:
            Baseline statistics
        """
        logger.info("Establishing data quality baseline")
        
        baseline = {
            'shape': data.shape,
            'missing_rates': data.isnull().sum() / len(data),
            'numeric_stats': data.select_dtypes(include=[np.number]).describe(),
            'categorical_stats': data.select_dtypes(include=['object']).nunique(),
            'data_types': data.dtypes.to_dict(),
            'memory_usage': data.memory_usage(deep=True).sum()
        }
        
        self.baseline_stats = baseline
        logger.info("Data quality baseline established")
        return baseline
    
    def validate_data_quality(self, data: pd.DataFrame, 
                            data_name: str = "data") -> Dict[str, Any]:
        """
        Validate data quality against baseline.
        
        Args:
            data: Data to validate
            data_name: Name of dataset for logging
            
        Returns:
            Quality validation results
        """
        if not self.baseline_stats:
            raise ValueError("No baseline established. Call establish_baseline first.")
        
        logger.info(f"Validating data quality for {data_name}")
        
        validation_results = {
            'data_name': data_name,
            'timestamp': datetime.now().isoformat(),
            'quality_issues': [],
            'quality_score': 1.0,
            'warnings': [],
            'passed': True
        }
        
        # Check shape consistency
        expected_shape = self.baseline_stats['shape'][1]  # Number of columns
        actual_shape = data.shape[1]
        
        if actual_shape != expected_shape:
            issue = f"Column count mismatch: expected {expected_shape}, got {actual_shape}"
            validation_results['quality_issues'].append(issue)
            validation_results['quality_score'] *= 0.8
        
        # Check for missing data
        baseline_missing = self.baseline_stats['missing_rates']
        current_missing = data.isnull().sum() / len(data)
        
        for col in baseline_missing.index:
            if col in current_missing.index:
                missing_increase = current_missing[col] - baseline_missing[col]
                if missing_increase > 0.1:  # 10% increase in missing data
                    issue = f"Missing data increase in {col}: {missing_increase:.2%}"
                    validation_results['quality_issues'].append(issue)
                    validation_results['quality_score'] *= 0.9
        
        # Check data types
        baseline_types = self.baseline_stats['data_types']
        current_types = data.dtypes.to_dict()
        
        for col, expected_type in baseline_types.items():
            if col in current_types:
                if current_types[col] != expected_type:
                    warning = f"Data type change in {col}: {expected_type} -> {current_types[col]}"
                    validation_results['warnings'].append(warning)
        
        # Overall quality assessment
        validation_results['passed'] = validation_results['quality_score'] >= 0.7
        
        self.quality_history.append(validation_results)
        
        if not validation_results['passed']:
            logger.warning(f"Data quality validation failed for {data_name}")
            logger.warning(f"Issues: {validation_results['quality_issues']}")
        
        return validation_results
    
    def detect_outliers(self, data: pd.DataFrame, 
                       method: str = 'iqr',
                       threshold: float = 3.0) -> Dict[str, Any]:
        """
        Detect outliers in data.
        
        Args:
            data: Data to analyze
            method: Outlier detection method ('iqr', 'zscore', 'isolation_forest')
            threshold: Threshold for outlier detection
            
        Returns:
            Outlier detection results
        """
        numeric_data = data.select_dtypes(include=[np.number])
        
        if numeric_data.empty:
            return {'outliers': {}, 'outlier_count': 0, 'method': method}
        
        outlier_results = {
            'method': method,
            'threshold': threshold,
            'outliers': {},
            'outlier_count': 0,
            'outlier_indices': []
        }
        
        if method == 'iqr':
            outlier_results = self._detect_iqr_outliers(numeric_data, threshold)
        elif method == 'zscore':
            outlier_results = self._detect_zscore_outliers(numeric_data, threshold)
        elif method == 'isolation_forest':
            outlier_results = self._detect_isolation_forest_outliers(numeric_data)
        
        logger.info(f"Outlier detection completed: {outlier_results['outlier_count']} outliers found")
        return outlier_results
    
    def _detect_iqr_outliers(self, data: pd.DataFrame, threshold: float) -> Dict[str, Any]:
        """Detect outliers using IQR method."""
        outliers = {}
        outlier_indices = set()
        
        for column in data.columns:
            Q1 = data[column].quantile(0.25)
            Q3 = data[column].quantile(0.75)
            IQR = Q3 - Q1
            lower_bound = Q1 - threshold * IQR
            upper_bound = Q3 + threshold * IQR
            
            column_outliers = data[(data[column] < lower_bound) | (data[column] > upper_bound)]
            outliers[column] = len(column_outliers)
            outlier_indices.update(column_outliers.index)
        
        return {
            'method': 'iqr',
            'threshold': threshold,
            'outliers': outliers,
            'outlier_count': len(outlier_indices),
            'outlier_indices': list(outlier_indices)
        }
    
    def _detect_zscore_outliers(self, data: pd.DataFrame, threshold: float) -> Dict[str, Any]:
        """Detect outliers using Z-score method."""
        outliers = {}
        outlier_indices = set()
        
        for column in data.columns:
            z_scores = np.abs(stats.zscore(data[column].dropna()))
            column_outliers = data[column][z_scores > threshold]
            outliers[column] = len(column_outliers)
            outlier_indices.update(column_outliers.index)
        
        return {
            'method': 'zscore',
            'threshold': threshold,
            'outliers': outliers,
            'outlier_count': len(outlier_indices),
            'outlier_indices': list(outlier_indices)
        }
    
    def _detect_isolation_forest_outliers(self, data: pd.DataFrame) -> Dict[str, Any]:
        """Detect outliers using Isolation Forest."""
        try:
            from sklearn.ensemble import IsolationForest
            
            # Fill missing values
            data_filled = data.fillna(data.median())
            
            # Fit Isolation Forest
            iso_forest = IsolationForest(contamination=0.1, random_state=42)
            outlier_labels = iso_forest.fit_predict(data_filled)
            
            outlier_indices = data_filled.index[outlier_labels == -1].tolist()
            
            return {
                'method': 'isolation_forest',
                'threshold': 0.1,
                'outliers': {'total': len(outlier_indices)},
                'outlier_count': len(outlier_indices),
                'outlier_indices': outlier_indices
            }
            
        except Exception as e:
            logger.warning(f"Isolation Forest outlier detection failed: {e}")
            return {
                'method': 'isolation_forest',
                'threshold': 0.1,
                'outliers': {},
                'outlier_count': 0,
                'outlier_indices': []
            }


def graceful_degradation(fallback_value: Any = None, 
                        fallback_func: Optional[Callable] = None):
    """
    Decorator for graceful degradation when operations fail.
    
    Args:
        fallback_value: Value to return if operation fails
        fallback_func: Function to call if operation fails
    """
    def decorator(func: Callable) -> Callable:
        @functools.wraps(func)
        def wrapper(*args, **kwargs):
            try:
                return func(*args, **kwargs)
            except Exception as e:
                logger.warning(f"Operation {func.__name__} failed, using graceful degradation: {e}")
                
                if fallback_func:
                    try:
                        return fallback_func(*args, **kwargs)
                    except Exception as fallback_error:
                        logger.error(f"Fallback function also failed: {fallback_error}")
                        return fallback_value
                
                return fallback_value
                
        return wrapper
    return decorator


# Global robustness manager instance
robustness_manager = RobustnessManager()
```
