## Supplementary material for "Machine Learning-Driven Prediction of TLR4 Binding Affinity: A Comprehensive Molecular Feature Analysis for Drug Discovery": Codebase: file_utils.py

```
"""
File operation utilities.

This module provides safe file operations and directory management
for the TLR4 binding prediction system.
"""

import os
import shutil
import tempfile
from pathlib import Path
from typing import Union, Optional, List
import logging

logger = logging.getLogger(__name__)


def ensure_directory(path: Union[str, Path], 
                    create_parents: bool = True) -> Path:
    """
    Ensure directory exists, create if necessary.
    
    Args:
        path: Directory path
        create_parents: Whether to create parent directories
        
    Returns:
        Path object of the directory
    """
    path = Path(path)
    
    if not path.exists():
        if create_parents:
            path.mkdir(parents=True, exist_ok=True)
            logger.info(f"Created directory: {path}")
        else:
            raise FileNotFoundError(f"Directory does not exist: {path}")
    
    return path


def safe_file_operation(operation_func, *args, **kwargs):
    """
    Safely execute file operation with error handling.
    
    Args:
        operation_func: Function to execute
        *args: Arguments for the function
        **kwargs: Keyword arguments for the function
        
    Returns:
        Result of the operation or None if failed
    """
    try:
        return operation_func(*args, **kwargs)
    except FileNotFoundError as e:
        logger.error(f"File not found: {e}")
        return None
    except PermissionError as e:
        logger.error(f"Permission denied: {e}")
        return None
    except Exception as e:
        logger.error(f"File operation failed: {e}")
        return None


def copy_file_safely(source: Union[str, Path], 
                    destination: Union[str, Path],
                    overwrite: bool = False) -> bool:
    """
    Safely copy file with error handling.
    
    Args:
        source: Source file path
        destination: Destination file path
        overwrite: Whether to overwrite existing file
        
    Returns:
        True if successful, False otherwise
    """
    source = Path(source)
    destination = Path(destination)
    
    if not source.exists():
        logger.error(f"Source file does not exist: {source}")
        return False
    
    if destination.exists() and not overwrite:
        logger.error(f"Destination file exists and overwrite=False: {destination}")
        return False
    
    try:
        # Ensure destination directory exists
        destination.parent.mkdir(parents=True, exist_ok=True)
        
        # Copy file
        shutil.copy2(source, destination)
        logger.info(f"File copied: {source} -> {destination}")
        return True
        
    except Exception as e:
        logger.error(f"Failed to copy file: {e}")
        return False


def move_file_safely(source: Union[str, Path], 
                    destination: Union[str, Path],
                    overwrite: bool = False) -> bool:
    """
    Safely move file with error handling.
    
    Args:
        source: Source file path
        destination: Destination file path
        overwrite: Whether to overwrite existing file
        
    Returns:
        True if successful, False otherwise
    """
    source = Path(source)
    destination = Path(destination)
    
    if not source.exists():
        logger.error(f"Source file does not exist: {source}")
        return False
    
    if destination.exists() and not overwrite:
        logger.error(f"Destination file exists and overwrite=False: {destination}")
        return False
    
    try:
        # Ensure destination directory exists
        destination.parent.mkdir(parents=True, exist_ok=True)
        
        # Move file
        shutil.move(str(source), str(destination))
        logger.info(f"File moved: {source} -> {destination}")
        return True
        
    except Exception as e:
        logger.error(f"Failed to move file: {e}")
        return False


def delete_file_safely(file_path: Union[str, Path]) -> bool:
    """
    Safely delete file with error handling.
    
    Args:
        file_path: Path to file to delete
        
    Returns:
        True if successful, False otherwise
    """
    file_path = Path(file_path)
    
    if not file_path.exists():
        logger.warning(f"File does not exist: {file_path}")
        return True
    
    try:
        file_path.unlink()
        logger.info(f"File deleted: {file_path}")
        return True
        
    except Exception as e:
        logger.error(f"Failed to delete file: {e}")
        return False


def get_file_size(file_path: Union[str, Path]) -> Optional[int]:
    """
    Get file size in bytes.
    
    Args:
        file_path: Path to file
        
    Returns:
        File size in bytes or None if error
    """
    file_path = Path(file_path)
    
    if not file_path.exists():
        logger.error(f"File does not exist: {file_path}")
        return None
    
    try:
        return file_path.stat().st_size
    except Exception as e:
        logger.error(f"Failed to get file size: {e}")
        return None


def list_files(directory: Union[str, Path], 
              pattern: str = "*",
              recursive: bool = False) -> List[Path]:
    """
    List files in directory matching pattern.
    
    Args:
        directory: Directory to search
        pattern: File pattern to match
        recursive: Whether to search recursively
        
    Returns:
        List of matching file paths
    """
    directory = Path(directory)
    
    if not directory.exists():
        logger.error(f"Directory does not exist: {directory}")
        return []
    
    try:
        if recursive:
            files = list(directory.rglob(pattern))
        else:
            files = list(directory.glob(pattern))
        
        # Filter to only files (not directories)
        files = [f for f in files if f.is_file()]
        
        logger.info(f"Found {len(files)} files matching pattern '{pattern}' in {directory}")
        return files
        
    except Exception as e:
        logger.error(f"Failed to list files: {e}")
        return []


def create_temp_file(suffix: str = ".tmp", 
                    prefix: str = "tlr4_",
                    directory: Optional[Union[str, Path]] = None) -> Path:
    """
    Create temporary file.
    
    Args:
        suffix: File suffix
        prefix: File prefix
        directory: Directory for temporary file
        
    Returns:
        Path to temporary file
    """
    try:
        if directory:
            directory = Path(directory)
            directory.mkdir(parents=True, exist_ok=True)
        
        fd, temp_path = tempfile.mkstemp(suffix=suffix, prefix=prefix, dir=directory)
        os.close(fd)  # Close file descriptor
        
        temp_path = Path(temp_path)
        logger.info(f"Created temporary file: {temp_path}")
        return temp_path
        
    except Exception as e:
        logger.error(f"Failed to create temporary file: {e}")
        raise


def cleanup_temp_files(temp_files: List[Union[str, Path]]) -> None:
    """
    Clean up temporary files.
    
    Args:
        temp_files: List of temporary file paths
    """
    for temp_file in temp_files:
        temp_file = Path(temp_file)
        if temp_file.exists():
            try:
                temp_file.unlink()
                logger.info(f"Cleaned up temporary file: {temp_file}")
            except Exception as e:
                logger.warning(f"Failed to clean up temporary file {temp_file}: {e}")


def get_file_extension(file_path: Union[str, Path]) -> str:
    """
    Get file extension.
    
    Args:
        file_path: Path to file
        
    Returns:
        File extension (including dot)
    """
    return Path(file_path).suffix


def change_file_extension(file_path: Union[str, Path], 
                         new_extension: str) -> Path:
    """
    Change file extension.
    
    Args:
        file_path: Original file path
        new_extension: New extension (with or without dot)
        
    Returns:
        New file path with changed extension
    """
    file_path = Path(file_path)
    
    # Ensure extension starts with dot
    if not new_extension.startswith('.'):
        new_extension = '.' + new_extension
    
    return file_path.with_suffix(new_extension)
```
