## Supplementary material for "Machine Learning-Driven Prediction of TLR4 Binding Affinity: A Comprehensive Molecular Feature Analysis for Drug Discovery": Codebase: fix_negative_r2.py

```
"""
Fixes for negative R² issues in model training.
"""

import pandas as pd
import numpy as np
from sklearn.feature_selection import SelectKBest, f_regression
from sklearn.preprocessing import StandardScaler
import logging

logger = logging.getLogger(__name__)


class FeatureQualityFixer:
    """Fixes feature quality issues that can cause negative R²."""
    
    def __init__(self):
        self.scaler = StandardScaler()
        self.feature_selector = None
    
    def fix_feature_quality(self, df: pd.DataFrame) -> pd.DataFrame:
        """Fix feature quality issues."""
        logger.info("Fixing feature quality issues...")
        
        # Identify target and feature columns
        target_col = 'affinity'
        if target_col not in df.columns:
            logger.warning(f"Target column '{target_col}' not found")
            return df
        
        feature_cols = [col for col in df.select_dtypes(include=[np.number]).columns 
                       if col not in [target_col, 'compound_name']]
        
        if len(feature_cols) == 0:
            logger.warning("No numeric feature columns found")
            return df
        
        # Remove features with zero variance
        original_count = len(feature_cols)
        for col in feature_cols.copy():
            if df[col].var() == 0 or df[col].isna().all():
                feature_cols.remove(col)
                df = df.drop(columns=[col])
        
        logger.info(f"Removed {original_count - len(feature_cols)} zero-variance features")
        
        # Remove highly correlated features
        if len(feature_cols) > 1:
            corr_matrix = df[feature_cols].corr().abs()
            upper_tri = corr_matrix.where(
                np.triu(np.ones(corr_matrix.shape), k=1).astype(bool)
            )
            
            to_drop = [column for column in upper_tri.columns if any(upper_tri[column] > 0.95)]
            df = df.drop(columns=to_drop)
            feature_cols = [col for col in feature_cols if col not in to_drop]
            
            logger.info(f"Removed {len(to_drop)} highly correlated features")
        
        # Scale features
        if len(feature_cols) > 0:
            df[feature_cols] = self.scaler.fit_transform(df[feature_cols])
            logger.info(f"Scaled {len(feature_cols)} features")
        
        return df
    
    def get_feature_importance_analysis(self, df: pd.DataFrame) -> pd.DataFrame:
        """Get feature importance analysis."""
        target_col = 'affinity'
        if target_col not in df.columns:
            return pd.DataFrame()
        
        feature_cols = [col for col in df.select_dtypes(include=[np.number]).columns 
                       if col not in [target_col, 'compound_name']]
        
        if len(feature_cols) == 0:
            return pd.DataFrame()
        
        # Remove rows with NaN in target
        clean_df = df.dropna(subset=[target_col])
        
        if len(clean_df) == 0:
            return pd.DataFrame()
        
        try:
            # Use SelectKBest to get feature scores
            selector = SelectKBest(score_func=f_regression, k='all')
            X = clean_df[feature_cols].fillna(0)
            y = clean_df[target_col]
            
            selector.fit(X, y)
            
            importance_df = pd.DataFrame({
                'feature': feature_cols,
                'score': selector.scores_,
                'p_value': selector.pvalues_
            }).sort_values('score', ascending=False)
            
            return importance_df
            
        except Exception as e:
            logger.error(f"Feature importance analysis failed: {e}")
            return pd.DataFrame()
```
