## Supplementary material for "Machine Learning-Driven Prediction of TLR4 Binding Affinity: A Comprehensive Molecular Feature Analysis for Drug Discovery": Codebase: logger.py

```
"""
Logging utilities for TLR4 binding prediction project.
"""

import logging
import sys
from typing import Optional


def get_logger(name: str, level: int = logging.INFO) -> logging.Logger:
    """
    Get a logger instance with consistent configuration.
    
    Args:
        name: Logger name (typically __name__)
        level: Logging level
        
    Returns:
        Configured logger instance
    """
    logger = logging.getLogger(name)
    
    # Avoid adding multiple handlers
    if logger.handlers:
        return logger
    
    logger.setLevel(level)
    
    # Create console handler
    console_handler = logging.StreamHandler(sys.stdout)
    console_handler.setLevel(level)
    
    # Create formatter
    formatter = logging.Formatter(
        '%(asctime)s - %(name)s - %(levelname)s - %(message)s',
        datefmt='%Y-%m-%d %H:%M:%S'
    )
    console_handler.setFormatter(formatter)
    
    # Add handler to logger
    logger.addHandler(console_handler)
    
    return logger


def setup_logging(level: int = logging.INFO, log_file: Optional[str] = None):
    """
    Set up logging configuration for the entire application.
    
    Args:
        level: Logging level
        log_file: Optional log file path
    """
    # Configure root logger
    logging.basicConfig(
        level=level,
        format='%(asctime)s - %(name)s - %(levelname)s - %(message)s',
        datefmt='%Y-%m-%d %H:%M:%S',
        handlers=[
            logging.StreamHandler(sys.stdout),
            *([logging.FileHandler(log_file)] if log_file else [])
        ]
    )
```
