## Supplementary material for "Machine Learning-Driven Prediction of TLR4 Binding Affinity: A Comprehensive Molecular Feature Analysis for Drug Discovery": Codebase: logging_config.py

```
"""
Logging configuration and utilities.

This module provides centralized logging configuration
for the TLR4 binding prediction system.
"""

import logging
import sys
from pathlib import Path
from typing import Optional, Union
from datetime import datetime


def setup_logging(log_level: str = "INFO", 
                 log_file: Optional[Union[str, Path]] = None,
                 log_format: Optional[str] = None) -> None:
    """
    Set up logging configuration for the application.
    
    Args:
        log_level: Logging level (DEBUG, INFO, WARNING, ERROR, CRITICAL)
        log_file: Path to log file (optional)
        log_format: Custom log format string (optional)
    """
    # Default log format
    if log_format is None:
        log_format = "%(asctime)s - %(name)s - %(levelname)s - %(message)s"
    
    # Configure root logger
    root_logger = logging.getLogger()
    root_logger.setLevel(getattr(logging, log_level.upper()))
    
    # Clear existing handlers
    root_logger.handlers.clear()
    
    # Create formatter
    formatter = logging.Formatter(log_format)
    
    # Console handler
    console_handler = logging.StreamHandler(sys.stdout)
    console_handler.setLevel(getattr(logging, log_level.upper()))
    console_handler.setFormatter(formatter)
    root_logger.addHandler(console_handler)
    
    # File handler (if specified)
    if log_file:
        log_file = Path(log_file)
        log_file.parent.mkdir(parents=True, exist_ok=True)
        
        file_handler = logging.FileHandler(log_file)
        file_handler.setLevel(getattr(logging, log_level.upper()))
        file_handler.setFormatter(formatter)
        root_logger.addHandler(file_handler)
    
    # Set specific logger levels
    logging.getLogger('matplotlib').setLevel(logging.WARNING)
    logging.getLogger('PIL').setLevel(logging.WARNING)
    logging.getLogger('urllib3').setLevel(logging.WARNING)
    
    logging.info(f"Logging configured with level: {log_level}")


def get_logger(name: str) -> logging.Logger:
    """
    Get logger instance for specific module.
    
    Args:
        name: Logger name (usually __name__)
        
    Returns:
        Logger instance
    """
    return logging.getLogger(name)


class LoggerMixin:
    """Mixin class to add logging functionality to any class."""
    
    @property
    def logger(self) -> logging.Logger:
        """Get logger for this class."""
        return logging.getLogger(self.__class__.__name__)
    
    def log_info(self, message: str) -> None:
        """Log info message."""
        self.logger.info(message)
    
    def log_warning(self, message: str) -> None:
        """Log warning message."""
        self.logger.warning(message)
    
    def log_error(self, message: str) -> None:
        """Log error message."""
        self.logger.error(message)
    
    def log_debug(self, message: str) -> None:
        """Log debug message."""
        self.logger.debug(message)
```
