## Supplementary material for "Machine Learning-Driven Prediction of TLR4 Binding Affinity: A Comprehensive Molecular Feature Analysis for Drug Discovery": Codebase: robustness_config.py

```
"""
Robustness configuration management.

This module provides configuration templates and utilities for setting up
robust error handling and recovery mechanisms across the TLR4 binding pipeline.
"""

import json
import yaml
from typing import Dict, Any, Optional, List
from pathlib import Path
from dataclasses import dataclass, asdict
import logging

logger = logging.getLogger(__name__)


@dataclass
class RobustnessConfig:
    """Configuration for robustness features."""
    
    # Retry and recovery settings
    max_retry_attempts: int = 3
    retry_delay: float = 1.0
    exponential_backoff: bool = True
    max_retry_delay: float = 60.0
    
    # Circuit breaker settings
    enable_circuit_breaker: bool = True
    circuit_breaker_threshold: int = 5
    circuit_breaker_timeout: int = 300
    
    # Checkpointing settings
    enable_checkpointing: bool = True
    checkpoint_interval: int = 100
    checkpoint_retention_days: int = 30
    checkpoint_compression: bool = True
    
    # Data quality settings
    enable_data_validation: bool = True
    missing_value_threshold: float = 0.3
    outlier_threshold: float = 3.0
    correlation_threshold: float = 0.95
    enable_anomaly_detection: bool = True
    
    # Graceful degradation settings
    enable_graceful_degradation: bool = True
    fallback_strategies: List[str] = None
    
    # Logging and monitoring settings
    enable_error_logging: bool = True
    error_log_retention_days: int = 7
    enable_performance_monitoring: bool = True
    enable_memory_monitoring: bool = True
    
    # Model-specific settings
    model_training_timeout: int = 3600  # 1 hour
    feature_extraction_timeout: int = 300  # 5 minutes
    prediction_timeout: int = 60  # 1 minute
    
    def __post_init__(self):
        """Post-initialization setup."""
        if self.fallback_strategies is None:
            self.fallback_strategies = [
                'use_default_values',
                'skip_failed_components',
                'use_simplified_models',
                'interpolate_missing_features'
            ]
    
    def to_dict(self) -> Dict[str, Any]:
        """Convert to dictionary."""
        return asdict(self)
    
    @classmethod
    def from_dict(cls, config_dict: Dict[str, Any]) -> 'RobustnessConfig':
        """Create from dictionary."""
        return cls(**config_dict)


class RobustnessConfigManager:
    """Manager for robustness configurations."""
    
    def __init__(self, config_dir: str = "configs"):
        """
        Initialize configuration manager.
        
        Args:
            config_dir: Directory for storing configuration files
        """
        self.config_dir = Path(config_dir)
        self.config_dir.mkdir(parents=True, exist_ok=True)
    
    def save_config(self, config: RobustnessConfig, name: str) -> bool:
        """
        Save configuration to file.
        
        Args:
            config: Configuration to save
            name: Configuration name
            
        Returns:
            True if successful, False otherwise
        """
        try:
            config_file = self.config_dir / f"{name}_robustness.json"
            
            config_data = {
                'name': name,
                'config': config.to_dict(),
                'metadata': {
                    'created_at': str(Path().resolve()),
                    'version': '1.0'
                }
            }
            
            with open(config_file, 'w') as f:
                json.dump(config_data, f, indent=2)
            
            logger.info(f"Robustness configuration saved: {name}")
            return True
            
        except Exception as e:
            logger.error(f"Failed to save robustness configuration {name}: {e}")
            return False
    
    def load_config(self, name: str) -> Optional[RobustnessConfig]:
        """
        Load configuration from file.
        
        Args:
            name: Configuration name
            
        Returns:
            Configuration object or None if not found
        """
        try:
            config_file = self.config_dir / f"{name}_robustness.json"
            
            if not config_file.exists():
                logger.warning(f"Configuration file not found: {name}")
                return None
            
            with open(config_file, 'r') as f:
                config_data = json.load(f)
            
            config_dict = config_data['config']
            config = RobustnessConfig.from_dict(config_dict)
            
            logger.info(f"Robustness configuration loaded: {name}")
            return config
            
        except Exception as e:
            logger.error(f"Failed to load robustness configuration {name}: {e}")
            return None
    
    def list_configs(self) -> List[str]:
        """List available configurations."""
        try:
            config_files = list(self.config_dir.glob("*_robustness.json"))
            return [f.stem.replace("_robustness", "") for f in config_files]
        except Exception as e:
            logger.error(f"Failed to list configurations: {e}")
            return []
    
    def delete_config(self, name: str) -> bool:
        """Delete configuration."""
        try:
            config_file = self.config_dir / f"{name}_robustness.json"
            if config_file.exists():
                config_file.unlink()
                logger.info(f"Configuration deleted: {name}")
                return True
            return False
        except Exception as e:
            logger.error(f"Failed to delete configuration {name}: {e}")
            return False


# Predefined configurations
def get_development_config() -> RobustnessConfig:
    """Get configuration optimized for development."""
    return RobustnessConfig(
        max_retry_attempts=2,
        retry_delay=0.5,
        enable_checkpointing=False,  # Disable for faster development
        enable_data_validation=True,
        enable_anomaly_detection=False,  # Disable for faster development
        enable_error_logging=True,
        error_log_retention_days=3,
        model_training_timeout=1800,  # 30 minutes
        feature_extraction_timeout=120,  # 2 minutes
        prediction_timeout=30  # 30 seconds
    )


def get_production_config() -> RobustnessConfig:
    """Get configuration optimized for production."""
    return RobustnessConfig(
        max_retry_attempts=5,
        retry_delay=2.0,
        exponential_backoff=True,
        max_retry_delay=120.0,
        enable_circuit_breaker=True,
        circuit_breaker_threshold=3,
        circuit_breaker_timeout=600,
        enable_checkpointing=True,
        checkpoint_interval=50,
        checkpoint_retention_days=90,
        checkpoint_compression=True,
        enable_data_validation=True,
        missing_value_threshold=0.1,
        outlier_threshold=2.5,
        correlation_threshold=0.98,
        enable_anomaly_detection=True,
        enable_graceful_degradation=True,
        enable_error_logging=True,
        error_log_retention_days=30,
        enable_performance_monitoring=True,
        enable_memory_monitoring=True,
        model_training_timeout=7200,  # 2 hours
        feature_extraction_timeout=600,  # 10 minutes
        prediction_timeout=120  # 2 minutes
    )


def get_research_config() -> RobustnessConfig:
    """Get configuration optimized for research experiments."""
    return RobustnessConfig(
        max_retry_attempts=3,
        retry_delay=1.0,
        enable_checkpointing=True,
        checkpoint_interval=25,  # More frequent checkpoints for long experiments
        enable_data_validation=True,
        enable_anomaly_detection=True,
        enable_error_logging=True,
        error_log_retention_days=14,
        enable_performance_monitoring=True,
        enable_memory_monitoring=True,
        model_training_timeout=14400,  # 4 hours for long experiments
        feature_extraction_timeout=900,  # 15 minutes
        prediction_timeout=180  # 3 minutes
    )


def get_minimal_config() -> RobustnessConfig:
    """Get minimal configuration for testing."""
    return RobustnessConfig(
        max_retry_attempts=1,
        retry_delay=0.1,
        enable_checkpointing=False,
        enable_data_validation=False,
        enable_anomaly_detection=False,
        enable_error_logging=False,
        enable_performance_monitoring=False,
        enable_memory_monitoring=False,
        model_training_timeout=300,  # 5 minutes
        feature_extraction_timeout=60,  # 1 minute
        prediction_timeout=10  # 10 seconds
    )


def create_custom_config(**kwargs) -> RobustnessConfig:
    """
    Create custom configuration with specified parameters.
    
    Args:
        **kwargs: Configuration parameters to override
        
    Returns:
        Custom configuration
    """
    default_config = RobustnessConfig()
    
    for key, value in kwargs.items():
        if hasattr(default_config, key):
            setattr(default_config, key, value)
        else:
            logger.warning(f"Unknown configuration parameter: {key}")
    
    return default_config


def validate_config(config: RobustnessConfig) -> List[str]:
    """
    Validate robustness configuration.
    
    Args:
        config: Configuration to validate
        
    Returns:
        List of validation issues (empty if valid)
    """
    issues = []
    
    # Validate retry settings
    if config.max_retry_attempts < 0:
        issues.append("max_retry_attempts must be non-negative")
    
    if config.retry_delay < 0:
        issues.append("retry_delay must be non-negative")
    
    if config.max_retry_delay < config.retry_delay:
        issues.append("max_retry_delay must be >= retry_delay")
    
    # Validate circuit breaker settings
    if config.circuit_breaker_threshold < 1:
        issues.append("circuit_breaker_threshold must be >= 1")
    
    if config.circuit_breaker_timeout < 0:
        issues.append("circuit_breaker_timeout must be non-negative")
    
    # Validate checkpointing settings
    if config.checkpoint_interval < 1:
        issues.append("checkpoint_interval must be >= 1")
    
    if config.checkpoint_retention_days < 0:
        issues.append("checkpoint_retention_days must be non-negative")
    
    # Validate data quality settings
    if not 0 <= config.missing_value_threshold <= 1:
        issues.append("missing_value_threshold must be between 0 and 1")
    
    if config.outlier_threshold < 0:
        issues.append("outlier_threshold must be non-negative")
    
    if not 0 <= config.correlation_threshold <= 1:
        issues.append("correlation_threshold must be between 0 and 1")
    
    # Validate timeout settings
    if config.model_training_timeout < 0:
        issues.append("model_training_timeout must be non-negative")
    
    if config.feature_extraction_timeout < 0:
        issues.append("feature_extraction_timeout must be non-negative")
    
    if config.prediction_timeout < 0:
        issues.append("prediction_timeout must be non-negative")
    
    return issues


def setup_default_configs(config_manager: RobustnessConfigManager) -> None:
    """
    Setup default configurations.
    
    Args:
        config_manager: Configuration manager instance
    """
    default_configs = {
        'development': get_development_config(),
        'production': get_production_config(),
        'research': get_research_config(),
        'minimal': get_minimal_config()
    }
    
    for name, config in default_configs.items():
        config_manager.save_config(config, name)
        logger.info(f"Default configuration '{name}' saved")


def get_config_for_environment(environment: str = "development") -> RobustnessConfig:
    """
    Get configuration for specific environment.
    
    Args:
        environment: Environment name ('development', 'production', 'research', 'minimal')
        
    Returns:
        Configuration for the environment
    """
    config_functions = {
        'development': get_development_config,
        'production': get_production_config,
        'research': get_research_config,
        'minimal': get_minimal_config
    }
    
    if environment not in config_functions:
        logger.warning(f"Unknown environment '{environment}', using development config")
        environment = 'development'
    
    return config_functions[environment]()
```
